# A Consensus Proteomic Analysis of Alzheimer’s Disease Brain and Cerebrospinal Fluid Reveals Early Changes in Energy Metabolism Associated with Microglia and Astrocyte Activation

**DOI:** 10.1101/802959

**Authors:** Erik C.B. Johnson, Eric B. Dammer, Duc M. Duong, Lingyan Ping, Maotian Zhou, Luming Yin, Lenora A. Higginbotham, Andrew Guajardo, Bartholomew White, Juan C. Troncoso, Madhav Thambisetty, Thomas J. Montine, Edward B. Lee, John Q. Trojanowski, Thomas G. Beach, Eric M. Reiman, Vahram Haroutunian, Minghui Wang, Eric Schadt, Bin Zhang, Dennis W. Dickson, Nilufer Ertekin-Taner, Todd E. Golde, Vladislav A. Petyuk, Philip L. De Jager, David A. Bennett, Thomas S. Wingo, Srikant Rangaraju, Ihab Hajjar, Joshua M. Shulman, James J. Lah, Allan I. Levey, Nicholas T. Seyfried

## Abstract

Our understanding of the biological changes in the brain associated with Alzheimer’s disease (AD) pathology and cognitive impairment remains incomplete. To increase our understanding of these changes, we analyzed dorsolateral prefrontal cortex of control, asymptomatic AD, and AD brains from four different centers by label-free quantitative mass spectrometry and weighted protein co-expression analysis to obtain a consensus protein co-expression network of AD brain. This network consisted of 13 protein co-expression modules. Six of these modules correlated with amyloid-β plaque burden, tau neurofibrillary tangle burden, cognitive function, and clinical functional status, and were altered in asymptomatic AD, AD, or in both disease states. These six modules reflected synaptic, mitochondrial, sugar metabolism, extracellular matrix, cytoskeletal, and RNA binding/splicing biological functions. The identified protein network modules were preserved in a community-based cohort analyzed by a different quantitative mass spectrometry approach. They were also preserved in temporal lobe and precuneus brain regions. Some of the modules were influenced by aging, and showed changes in other neurodegenerative diseases such as frontotemporal dementia and corticobasal degeneration. The module most strongly associated with AD pathology and cognitive impairment was the sugar metabolism module. This module was enriched in AD genetic risk factors, and was also highly enriched in microglia and astrocyte protein markers associated with an anti-inflammatory state, suggesting that the biological functions it represents serve a protective role in AD. Proteins from the sugar metabolism module were increased in cerebrospinal fluid from asymptomatic AD and AD cases, highlighting their potential as biomarkers of the altered brain network. In this study of >2000 brains and nearly 400 cerebrospinal fluid samples by quantitative proteomics, we identify proteins and biological processes in AD brain that may serve as therapeutic targets and fluid biomarkers for the disease.

## Introduction

Alzheimer’s disease (AD) is a leading cause of death worldwide, with increasing prevalence as global life expectancy increases^1^. Although AD is currently defined on the basis of amyloid-β plaque and tau neurofibrillary tangle deposition within the neocortex^2^, the biochemical and cellular changes in the brain that characterize the disease beyond amyloid-β and tau deposition remain incompletely understood. The genetic architecture of late-onset AD has been extensively studied, and the results of these studies implicate multiple biological pathways that contribute to development of the disease, including immune function, endocytic vesicle trafficking, and lipid homeostasis, among others^3-5^. In addition to genetic studies, transcriptomic studies on postmortem AD brain tissue have identified changes in mRNA co-expression that correlate with disease traits and cognitive decline^6,7^. However, given that mRNA levels correlate only modestly to protein levels^8,9^, and proteins are the effectors of most biological functions, it is important to understand the proteomic changes that occur in AD brain to help advance drug development and biomarker efforts for this disease^10^.

Protein co-expression analysis is a powerful tool to understand biological network, pathway, and cell type changes in human tissue^11,12^. Communities of co-expressed proteins can be linked to disease processes, and the most strongly correlated proteins, or “hubs,” within these co-expression modules are enriched in key drivers of disease pathogenesis^13-18^. Therefore, targeting hubs within protein co-expression modules most related to disease biology is a promising approach for drug and biomarker development^19-22^. We recently analyzed control, asymptomatic AD, and AD brain tissue, using both protein differential and co-expression approaches, in a cohort of 50 individuals from the Baltimore Longitudinal Study of Aging to better understand the proteomic changes that occur in AD brain^23,24^. We observed protein co-expression changes that correlated with AD phenotypes, and found that glial and inflammatory protein network alterations are important features of AD pathophysiology. Importantly, many of the observed AD network findings were unique to the proteome, and not observed in the transcriptome, consistent with recent findings in other diseases such as cancer^10^. Here, we describe a multi-center collaborative project to extend these initial findings by analyzing more than 2000 human brain tissues by quantitative mass spectrometry-based proteomics. We generate a consensus AD brain protein co-expression network, controlling for batch and other covariates, from 453 brains obtained from multiple research centers. We validate this protein network in a separate community-based cohort using a different mass spectrometry-based technology for protein quantitation, and show that the network is preserved in different brain regions affected in AD. By analyzing a separate cohort of normal aging brains, we are able to estimate the effect aging has on the observed AD brain protein co-expression network. We also analyze the disease specificity of the AD protein network changes by interrogating these changes in six other neurodegenerative diseases that encompass diverse brain pathologies, and validate the observed changes by targeted protein measurements. One of the most strongly altered AD protein co-expression modules, which we term the “astrocyte/microglial metabolism” module, is enriched in proteins linked to microglia, astrocytes, and sugar metabolism, and is enriched in protein products linked to AD genetic risk. Microglial protein markers within this module are biased toward an anti-inflammatory disease-associated state, suggesting that it reflects a protective or compensatory function in response to AD pathology. Proteins from this module are increased in cerebrospinal fluid in individuals with AD, including in the asymptomatic stage of the disease. Our results highlight the importance of inflammation, sugar metabolism, mitochondrial function, synaptic function, RNA-associated proteins, and glia in the pathogenesis of AD, and provide a robust framework for future proteomic and multi-omic studies on AD brain and biofluid biomarkers.

## Results

### Construction of a Consensus AD Protein Co-Expression Network

Our initial investigation into protein co-expression changes in AD was centered on analysis of 50 cases encompassing control, asymptomatic AD (AsymAD), and AD brains^23^. Proteomic analyses, like genomic and transcriptomic analyses, can be influenced by multiple technical and experimental variables that are difficult to completely control in any one experiment, as well as by the inherent variability in disease pathology among selected AD cases, and variability in pathological assessment among different investigators. Because these factors may influence the resulting AD protein co-expression network, we set out to build a network that was robust to such variability. To do so, we analyzed control, AsymAD, and AD brains from multiple research centers in multiple batches over different time periods. In addition to the original 47 cases from the Baltimore Longitudinal Study of Aging (BLSA), we analyzed dorsolateral prefrontal cortex (DLPFC) tissue in 178 cases from the Banner Sun Health Research Institute (Banner), 166 cases from the Mount Sinai School of Medicine Brain Bank (MSSB), and 65 cases from the Adult Changes in Thought Study (ACT), for a total of 453 control, AsymAD, and AD brains (Figure 1A). All cases had comprehensive clinical evaluations during life, and were classified according to a common diagnostic scheme as described in Methods. AsymAD was defined as postmortem pathology consistent with an AD diagnosis but without dementia, based on the NIA research framework for AD^2^. Tissues were analyzed by mass spectrometry-based proteomics using label-free quantitation (LFQ), and the resulting mass spectrometry data were processed using a common pipeline to arrive at 5688 total quantified proteins. Out of these 5688 proteins, 3334 were quantified with fewer than 50% missing values across the 453 cases. We included proteins with fewer than 50% missing values in the subsequent analyses, as it was determined that this threshold was robust to potential spurious correlations given the power of the study (Supplementary Figure 1). We also removed by regression the effects of age, sex, and post-mortem interval on the protein quantitative data, even though these covariates did not strongly influence the data (Supplementary Figure 2). The final adjusted 3334 proteins were used to generate a protein co-expression network using the weighted correlation network analysis (WGCNA) algorithm. The resulting network consisted of 13 protein co-expression “modules,” or communities of proteins with similar expression patterns across the cases analyzed (Figure 1B**, Extended Data Figures 1 and 2, Supplementary Table 3**). The modules ranged in size from 254 proteins (M1) to 20 proteins (M13). These modules could also be identified independently of the WGCNA algorithm using t-distributed stochastic neighbor embedding (t-SNE) analysis (Supplementary Figure 3), demonstrating that the protein communities identified by the WGCNA algorithm were robust. Gene ontology (GO) analysis of the protein module members revealed a clear ontology for eleven out of the thirteen modules, encompassing a diverse mix of biological functions, processes, and components (Figure 1B**, Extended Data Figure 3**). To assess whether a given co-expression module was related to AD, we correlated the module eigenprotein—or first principle component of the module protein expression level—to the neuropathological hallmarks of AD: amyloid-β plaques and neurofibrillary tangles. We also correlated the module eigenproteins to cognitive function as assessed by the Mini-Mental Status Examination (MMSE), and functional status as assessed by the Clinical Dementia Rating Scale (CDR), at the last research evaluations prior to death to capture module-disease relationships that may be independent of amyloid-β plaque or tau tangle pathology (Figure 1B**, Extended Data Figure 1**). We observed six modules that were significantly correlated with all pathological, cognitive, and functional measures, and whose ontologies could be best characterized by a structural component or a biologic process: modules M1 synapse, M3 mitochondrial, M4 glucose and carbohydrate metabolism (subsequently referred to as sugar metabolism), M5 extracellular matrix, M6 cytoskeleton, and M10 RNA binding/splicing. The M4 sugar metabolism module showed the strongest AD trait correlations (amyloid-β plaque r=0.46, p=1.3e^-23^; neurofibrillary tangle r=0.49, p=4.7e^-27^; cognition r=-0.67, p=8.5e^-23^; functional status r=0.52, p=2.6e^-12^). Two modules—M11 chaperone/protein folding and M12 of unknown function—correlated with AD pathology and cognitive function, but not with functional status as assessed by CDR. The M8 module, which contained many circulating blood components such as hemoglobins and fibrinogen, and the M2 module, which predominantly reflected myelin components, correlated only with cognitive status but not AD pathology or functional status, suggesting that these modules may influence dementia separately from amyloid-β plaques and tau tangles. Because AD neuropathology is not homogenous even within the same brain region, and because neuropathological measurements of AD pathology are semi-quantitative and subject to a certain degree of individual variability in assessment^25^, we also correlated module eigenproteins to mass spectrometry measurements of amyloid-β and the tau microtubule binding region, which comprises neurofibrillary tangles, within the DLPFC tissue used for proteomic analysis (Supplementary Figure 4). We observed strong concordance between neuropathological and molecular measurements of AD pathology. We were also able to quantify alpha-synuclein and TAR DNA-binding protein 43 (TDP-43) proteins and assess their module correlations (Supplementary Figure 4A). Alpha-synuclein correlated most strongly with the M6 cytoskeleton module, as well as the M1 synaptic module. TDP-43 correlated most strongly with the RNA-binding module. These observations lend additional validity to the co-expression results given their known functions at the synapse and in RNA-binding, respectively. Alpha-synuclein also positively correlated with the M1 synaptic module, suggesting that we measured mostly soluble and physiological forms of this protein, rather than alpha-synuclein present in Lewy body aggregates.

**Figure 1.**
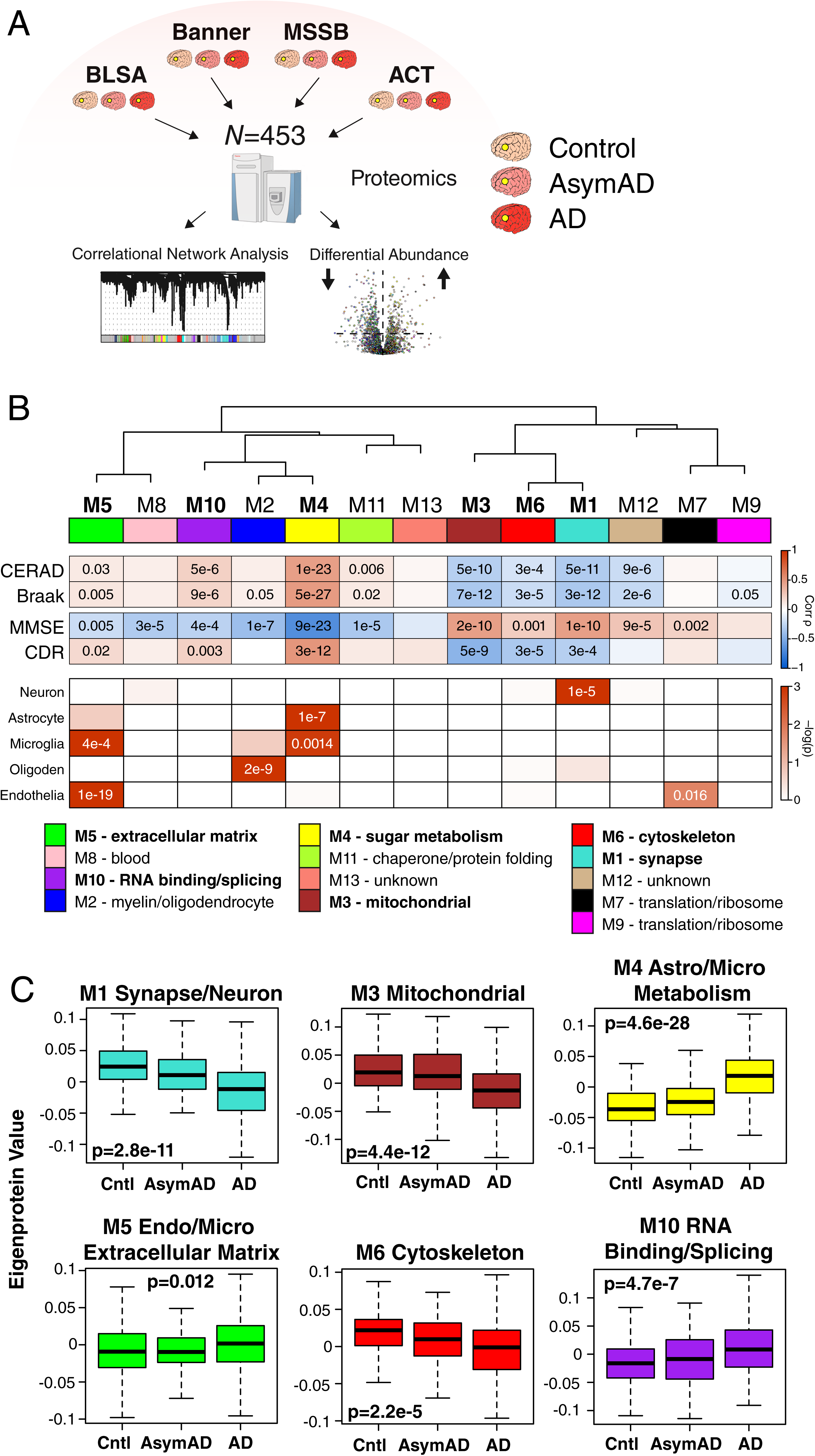
Protein Network Analysis of Asymptomatic and Symptomatic Alzheimer’s Disease Brain. (A-C) Protein levels in brain tissue from control, asymptomatic Alzheimer’s disease (AsymAD), and Alzheimer’s disease (AD) patients (*N*=453) were measured by label-free mass spectrometry and analyzed by weighted correlation network analysis (WGCNA) and differential abundance (A). Brain tissue was analyzed from postmortem dorsolateral prefrontal cortex (DLPFC, highlighted in yellow) in the Baltimore Longitudinal Study of Aging (BLSA, *n*=11 control, *n*=13 AsymAD, *n*=20 AD, *n*=44 total), Banner Sun Health Research Institute Brain Bank (Banner, *n*=26 control, *n*=58 AsymAD, *n*=94 AD, *n*=178 total), Mount Sinai School of Medicine Brain Bank (MSSB, *n*=46 control, *n*=17 AsymAD, *n*=103 AD, *n*=166 total), and the Adult Changes in Thought Study (ACT, *n*=11 control, *n*=14 AsymAD, *n*=40 AD, *n*=65). (B) A protein correlation network consisting of 13 protein modules was generated from 3334 proteins measured across four separate cohorts. (Top) Module eigenproteins, which represent the first principle component of the protein expression within each module, were correlated with neuropathological hallmarks of Alzheimer’s disease (CERAD, Consortium to Establish a Registry for Alzheimer’s disease amyloid-β plaque score, higher scores represent greater plaque burden; Braak, tau neurofibrillary tangle staging score, higher scores represent greater extent of tangle burden), cognitive function (MMSE, mini-mental status examination score, higher scores represent better cognitive function), and overall functional status (CDR, clinical dementia rating score, higher scores represent worse functional status). CERAD and Braak measures were from all cohorts, while MMSE was from Banner and CDR was from MSSB. Strength of positive (red) or negative (blue) correlation is shown by two-color heatmap, with *p* values provided for all correlations with *p* < 0.05. Modules that showed a significant correlation with all four traits are highlighted in bold. (Middle) The cell type nature of each protein module was assessed by module protein overlap with known neuron, astrocyte, microglia, oligodendrocyte (oligoden), and endothelia cell markers. Significance of overlap is shown by one-color heatmap, with *p* values provided for overlaps with *p* < 0.05. (Bottom) Gene ontology (GO) analysis of the proteins within each module clearly identified, for most modules, the biological processes associated with the module. (C) Module eigenprotein level by case status for each protein module that had significant correlation to all four traits in (B). Case status is from all cohorts (control, *n*=94; AsymAD, *n*=102; AD, *n*=257). *APOE* genotype effects and other trait correlations for all modules are provided in **Extended Data Figure 1**. Module eigenprotein correlations were performed using biweight midcorrelation and corrected by the Benjamini-Hochberg method. Protein module cell type overlap was performed using Fisher’s exact test with Benjamini-Hochberg correction. Differences in eigenprotein values were assessed by Kruskal-Wallis one-way ANOVA. Boxplots represent the median, 25^th^, and 75^th^ percentiles, and whiskers represent measurements to the 5^th^ and 95^th^ percentiles. Cntl, control; AsymAD, asymptomatic Alzheimer’s disease; AD, Alzheimer’s disease.

Because many protein co-expression changes in the brain can be driven by cell type changes^23,26^, we also assessed the cell type nature of each co-expression module by asking whether the module was enriched in particular cell type marker proteins (Figure 1B). We observed significant enrichment of neuronal proteins in the M1 synapse module and enrichment of oligodendrocyte markers in the M2 myelin module, as expected. We also observed enrichment of astrocyte and microglial proteins in the M4 sugar metabolism module, microglial and endothelial proteins in the M5 extracellular matrix module, and endothelial markers in the M7 translation/ribosome module. These findings suggest that the biological processes reflected by GO analysis for each module may be altered in AD within a particular cell type. To incorporate the cell type nature of each module into its description, we will subsequently refer to those modules with strong cell type enrichment as the “M1 synapse/neuron” module, the “M2 myelin/oligodendrocyte” module, the “M4 astrocyte/microglial metabolism” module, and the “M5 endo/micro extracellular matrix” module.

To assess the relationship of the network modules to diagnostic classification, we measured the module eigenprotein values by case status (Figure 1C **and Extended Data Figure 1**). In general, most modules that were increased or decreased in AD compared to control also showed a trend, or were significantly changed, in the same direction in the AsymAD group, indicating that these modules reflect pathophysiologic processes that begin early—in the preclinical phase—of AD. The M1 synapse/neuron, M3 mitochondrial, and M4 astrocyte/microglial metabolism modules showed the strongest differences by case status. We also assessed the influence of *APOE* genotype on module eigenproteins, but did not find strong effects except for the *APOE* ε2 allele on the M2 myelin/oligodendrocyte module, which appeared to attenuate the observed changes in AD (**Extended Data Figure 1**). In summary, we were able to construct a robust AD protein co-expression network from mass spectrometry-based proteomic analysis of greater than 450 human DLPFC brain tissues from multiple centers. We found that many of these modules correlated with AD neuropathology and cognitive function, reflected a number of different biological processes and cell types, and were altered in AsymAD.

### The AD Network Is Preserved in a Community-Based Observational Aging Cohort Analyzed by a Different Mass Spectrometry-Based Quantification Approach

The brains analyzed for the consensus AD network were obtained from studies or brain bank programs where brain donation is an optional aspect of the study, potentially introducing case selection bias. Furthermore, all of the cases analyzed for the consensus AD network were processed in a mass spectrometry-based pipeline with label-free quantitation (LFQ). Other forms of mass spectrometry-based protein quantification approaches exist, such as quantification using isobaric multiplex tandem mass tags (TMT)^27-29^. To assess whether a different mass spectrometry-based quantitation approach applied to a community-based aging cohort in which brain donation is a mandatory aspect of the study would yield a similar co-expression network, we analyzed 340 DLPFC brain tissues from the Religious Orders Study and Memory and Aging Project (ROS/MAP)^30-32^ using tandem mass tag mass spectrometry-based quantitation (TMT-MS) (Figure 2A). Cases in ROS/MAP were selected and classified for analysis using the same criteria as used for the LFQ-based cohorts. A protein co-expression network was constructed from the ROS/MAP cases, and network module preservation statistics were used to assess conservation of the consensus AD LFQ-based network in the ROS/MAP TMT-based network (Figure 2B). We found that all consensus LFQ modules were preserved in the ROS/MAP TMT-based network. To further investigate preservation of consensus AD network modules in ROS/MAP, we created “synthetic” eigenproteins for each consensus AD network module using the top 20% of module proteins by module eigenprotein correlation value (kME), and then tested if and how these synthetic eigenproteins were altered by case status in the ROS/MAP cohort. We found that the synthetic module eigenproteins showed similar changes by case status as observed in the consensus AD network, and showed consistent correlations with amyloid-β pathology, tau pathology, and cognitive function as assessed in ROS/MAP (Figure 2C **and Extended Data Figure 4**). Furthermore, targeted protein measurements in a cohort of 1016 ROS/MAP control, AsymAD, and AD brains by another mass spectrometry protein quantification approach—selected reaction monitoring (SRM)—showed that individual module proteins had the same direction of change as the AD LFQ-based network co-expression module of which they were a member (**Extended Data Figures 5 and 6**). These findings confirm that the consensus AD network is robust to different mass spectrometry-based protein quantification approaches, and reflects brain proteomic changes that occur in community-dwelling older individuals.

**Figure 2.**
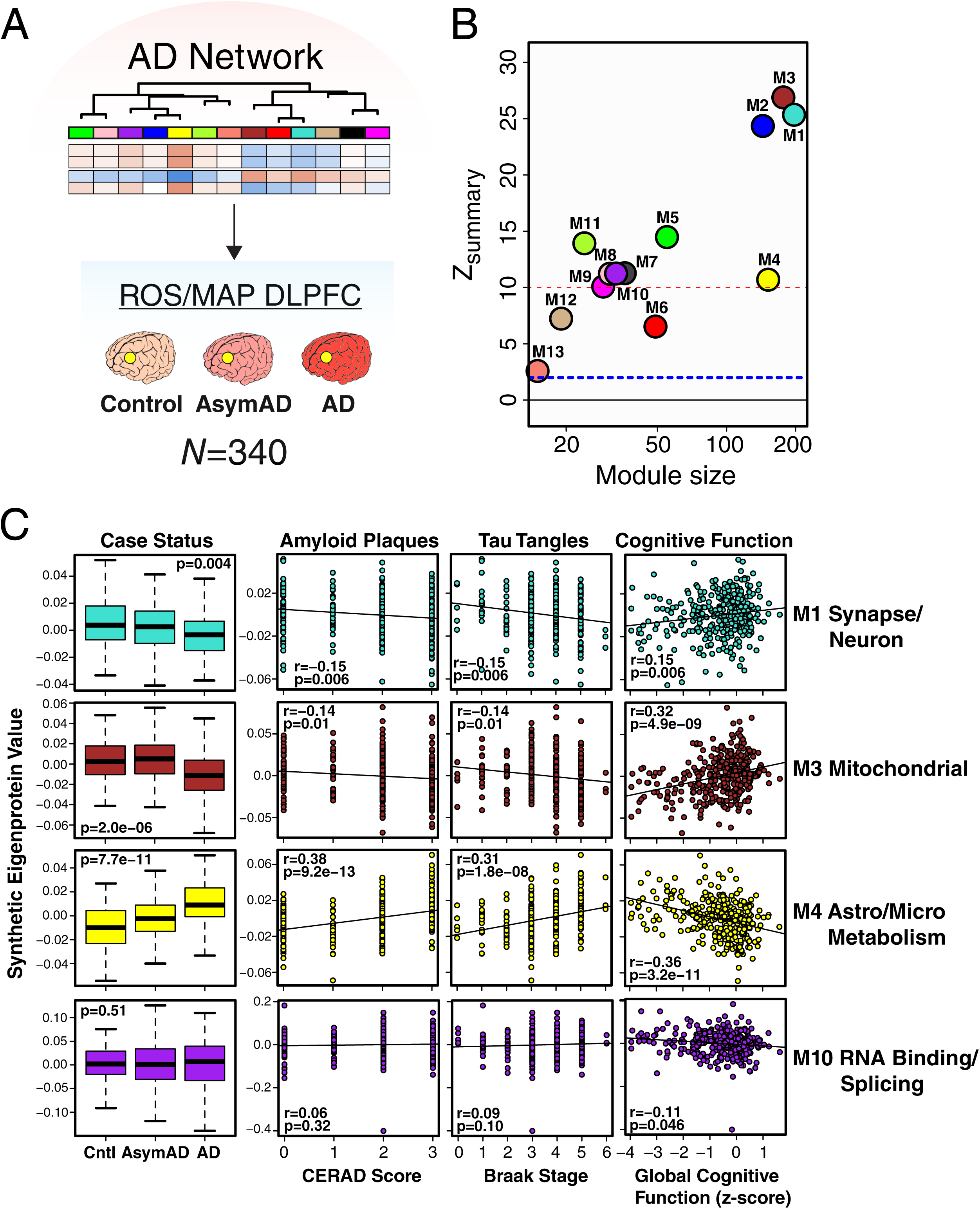
AD Protein Network Validation in a Longitudinal Cohort of Aging. (A-C) Preservation of AD protein network modules and trait correlations in the Religious Orders Study and Memory and Aging Project (ROS/MAP) cohorts. (A) Protein levels from dorsolateral prefrontal cortex (DLPFC) in a total of 340 control, AsymAD, and AD cases (control, *n*=84; AsymAD, *n*=148; AD, *n*=108) from the ROS/MAP cohorts were measured using a different mass spectrometry platform and quantification approach compared to the cases used to generate the AD network as shown in Figure 1. The resulting data were used to assess conservation of the AD brain protein network in the ROS/MAP cohorts. (B) AD brain protein network module preservation in the ROS/MAP cohorts. The dashed blue line indicates a *z*_summary_ score of 1.96, or FDR *q* value <0.05, above which module preservation was considered statistically significant. The dashed red line indicates a *z*_summary_ score of 10, or FDR *q* value ∼ 1e^-23^, above which module preservation was considered highly statistically significant. (C) Case status and trait preservation in the ROS/MAP cohorts. The top 20% of proteins by kME value in each AD brain protein network module was used to create a synthetic eigenprotein, which was then measured by case status in ROS/MAP and correlated with amyloid plaque load (CERAD score), tau neurofibrillary tangle burden (Braak stage), and cognitive function (global cognitive function composite z-score). Synthetic eigenprotein analyses for modules M1, M3, M4, and M10 are shown. Analyses for all modules, with additional trait correlations, are provided in **Extended Data Figure 4**. Differences in module synthetic eigenproteins by case status were assessed by Kruskal-Wallis one-way ANOVA. Module synthetic eigenprotein correlations were performed using biweight midcorrelation with Benjamini-Hochberg correction. Boxplots represent the median, 25^th^, and 75^th^ percentiles, and whiskers represent measurements to the 5^th^ and 95^th^ percentiles. Cntl, control; AsymAD, asymptomatic Alzheimer’s disease; AD, Alzheimer’s disease.

### The AD Network Is Preserved in Other Brain Regions

The consensus AD network was generated from analysis of DLPFC tissue. To assess whether the network was similar in other brain regions commonly affected in AD, we analyzed control and AD brain tissue from temporal cortex in a separate set of 111 brains from the Mayo Clinic, and control, AsymAD, and AD brain tissue from precuneus in the same set of brains from the BLSA (Figure 3A) using LFQ-MS. Co-expression networks were built for each brain region, and network preservation statistics were used to assess module preservation from DLPFC in temporal cortex (Figure 3B) and precuneus (Figure 3C). We found that all consensus AD network modules derived from DLPFC were preserved in temporal cortex, and twelve out of the thirteen modules were preserved in precuneus. While the larger modules by protein membership were all very highly preserved in both temporal cortex and precuneus, the M3 mitochondrial, M4 astrocyte/microglial metabolism, and M10 RNA binding/splicing modules were more highly preserved in temporal cortex, whereas the M2 myelin/oligodendrocyte and M5 endothelial/microglial extracellular matrix modules were more highly preserved in precuneus. Analysis of synthetic module eigenprotein values by case status showed similar differences between and among case groups in temporal cortex (Figure 3D**, Extended Data Figure 7**) and precuneus (Figure 3E**, Extended Data Figure 8**) brain regions, with changes in AsymAD more pronounced in precuneus than in DLPFC. These findings suggest that the consensus AD network is generalized across brain regions that are commonly affected in AD.

**Figure 3.**
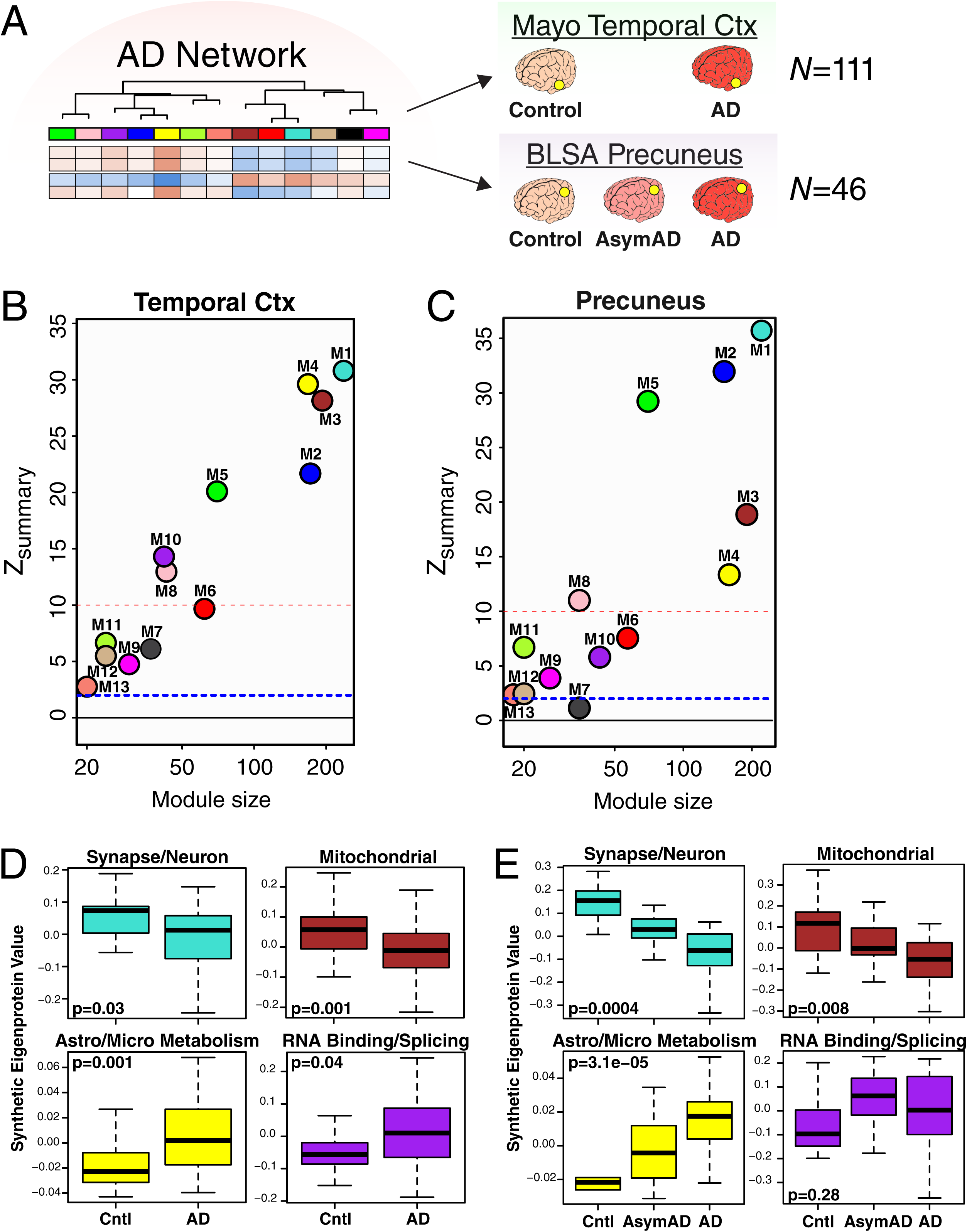
AD Protein Network Is Preserved in Different Brain Regions. (A-E) Preservation of AD protein network modules derived from analysis of DLPFC in other brain regions affected by AD. (A) Protein levels in temporal cortex from a total of 111 control and AD cases (control, *n*=28; AD, *n*=83) from the Mayo Brain Bank, and in precuneus from a total of 46 cases from the BLSA (control, *n*=12; AsymAD, *n*=14; AD, *n*=20) were measured by label-free mass spectrometry and used to assess conservation of the AD brain protein network derived from DLPFC. (B, C) AD brain protein network preservation in temporal cortex (B) and precuneus (C). The dashed blue line indicates a *z*_summary_ score of 1.96, or FDR *q* value <0.05, above which module preservation was considered statistically significant. The dashed red line indicates a *z*_summary_ score of 10, or FDR *q* value ∼ 1e^-23^, above which module preservation was considered highly statistically significant. (D, E) Case status preservation in temporal cortex and precuneus. A synthetic eigenprotein was created for each AD network module as described in Figure 2 and measured by case status in temporal cortex (D) and precuneus (E). Asymptomatic AD was not assessed in the Mayo cohort, and is therefore not included in the temporal cortex analyses. Synthetic eigenprotein analyses for modules M1, M3, M4, and M10 are shown. Analyses for all modules, with additional trait correlations, are provided in **Extended Data Figures 7 and 8**. Differences in module synthetic eigenproteins by case status were assessed by Welch’s *t* test (D) or Kruskal-Wallis one-way ANOVA (E). Boxplots represent the median, 25^th^, and 75^th^ percentiles, and whiskers represent measurements to the 5^th^ and 95^th^ percentiles. Cntl, control; AsymAD, asymptomatic Alzheimer’s disease; AD, Alzheimer’s disease.

### Effects of Aging on AD Network Modules

Aging is the strongest risk factor for AD, but the mechanistic relationship between aging and AD is unclear and is a topic of debate^33-38^. To better understand the influence aging may have on the consensus AD network, we analyzed DLPFC tissues from Johns Hopkins in 84 cases ages 30 to 69 (Figure 4A) by LFQ-MS. All cases had a final primary neuropathological diagnosis of control. We created synthetic eigenproteins in the aging cohort from the consensus AD network modules and asked whether the synthetic module eigenproteins changed with age (Figure 4B **and Extended Data Figure 9**). We found that the M1 synapse/neuron and M4 astrocyte/microglial metabolism modules decreased and increased with aging, respectively, while the M3 mitochondrial and M10 RNA binding/splicing modules were not affected by aging. Other modules that appeared to be affected by aging included the M6 cytoskeleton, M7 translation/ribosome, and M9 translation/ribosome modules (**Extended Data Figure 9**). Additional information on the correlation of individual proteins with age and overlap with markers of cellular senescence is provided in **Supplementary Table 4**. These findings indicate that the relationship between aging and AD at the proteomic level is complex, and that some, but not all, AD trait-associated modules are influenced by the aging process.

**Figure 4.**
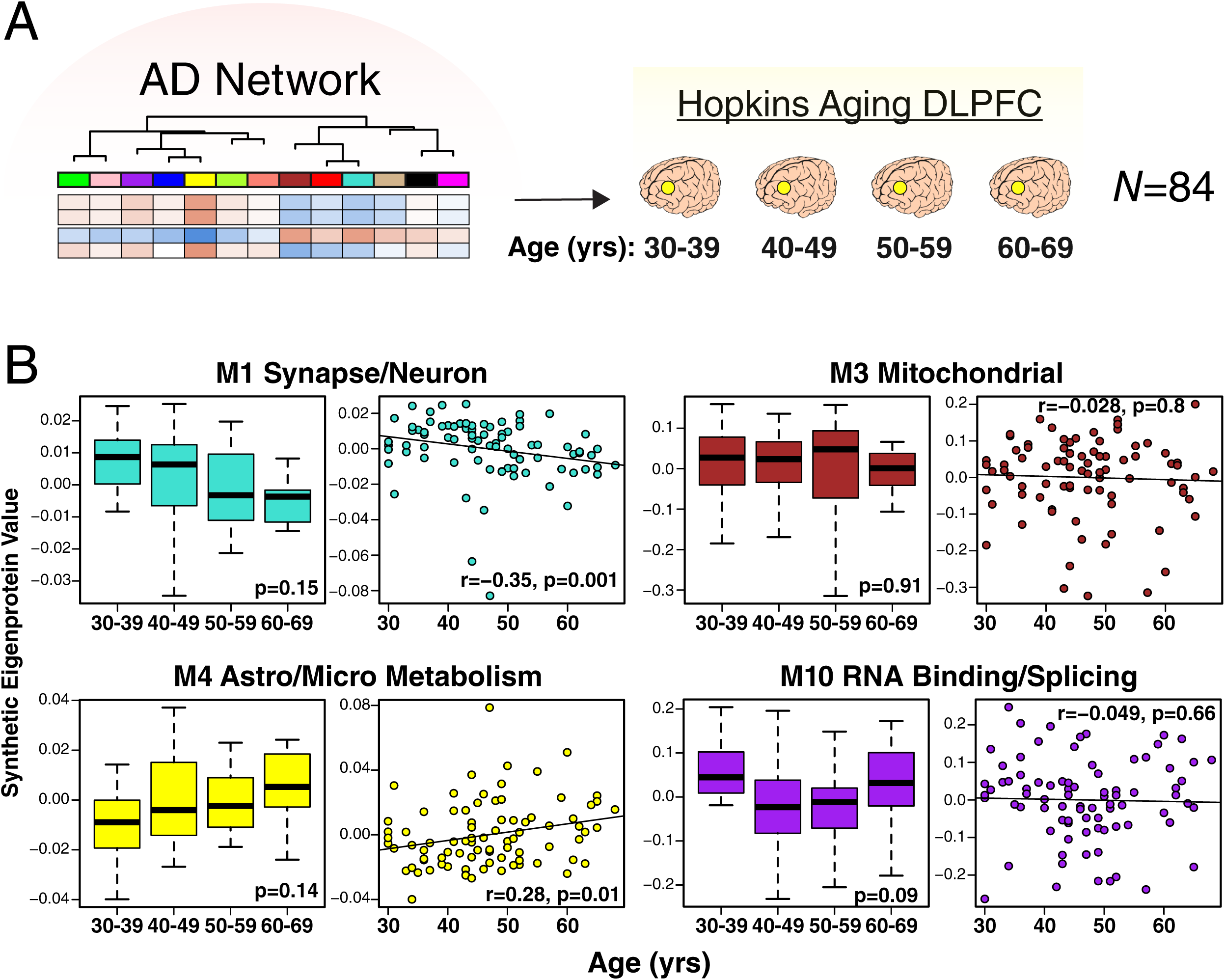
Effects of Aging on AD Protein Network Modules. (A, B) Protein levels were measured in DLPFC from cognitively normal people who died at different ages (age 30-39, *n*=20; age 40-49, *n*=34; age 50-59, *n*=17; age 60-69, *n*=13), and used to analyze AD protein network module changes with age. Brains were obtained from Johns Hopkins University. (B) A synthetic eigenprotein was created for each AD network module as described in Figure 2 and measured by age group (left boxplot) as well as correlated with age (right scatterplot) in the aging brain cohort. Synthetic eigenprotein analyses for modules M1, M3, M4, and M10 are shown. Analyses for all modules are provided in **Extended Data Figure 9**. Differences in module synthetic eigenproteins by age grouping were assessed by Kruskal-Wallis one-way ANOVA. Synthetic eigenprotein correlations were performed using biweight midcorrelation. Boxplots represent the median, 25^th^, and 75^th^ percentiles, and whiskers represent measurements to the 5^th^ and 95^th^ percentiles.

### AD Network Changes in Other Neurodegenerative Diseases

The extent to which AD network protein co-expression modules are altered in other neurodegenerative diseases is not fully understood. To explore the specificity of these network changes for AD, we analyzed 331 DLPFC tissues by LFQ-MS from control, AD, amyotrophic lateral sclerosis (ALS), frontotemporal lobar degeneration with TDP-43 pathology (FTLD-TDP), progressive supranuclear palsy (PSP), corticobasal degeneration (CBD), Parkinson’s disease and Parkinson’s disease dementia (PD/PDD), and multiple systems atrophy (MSA) cases (Figure 5A). We created synthetic eigenproteins for consensus AD network modules and assessed whether they changed in these different neurodegenerative diseases compared to AD (Figure 5B**, Extended Data Figure 10, Supplementary Table 5**). We found that the M1 synapse/neuron and M4 astrocyte/microglial metabolism modules showed significant changes in FTLD-TDP and CBD cases, similar to AD, whereas the M3 mitochondrial and M10 RNA binding/splicing modules showed more mixed changes across other diseases. To further validate these findings, we used a targeted mass spectrometry method called parallel reaction monitoring (PRM)^39^ to measure 323 individual proteins from approximately one-third of the cases analyzed in the untargeted experiments (**Extended Data Figure 11, Supplementary Table 5**). Protein levels across all cases were highly correlated (r=0.92, p=1.4e^-124^) between LFQ and PRM measurements (Supplementary Figure 5B). We created synthetic eigenproteins from these targeted PRM protein measurements by AD consensus module, and assessed eigenprotein changes by disease category (Supplementary Figure 5C**, Extended Data Figure 12, Supplementary Table 5**). We observed very similar AD network module changes across diseases compared to the untargeted measurements, validating the findings from the untargeted LFQ measurements. These results indicate that certain AD network modules are affected to a greater extent in AD compared to other neurodegenerative diseases, and that FTLD and CBD show many similar changes to AD, with the caveat that not all neurodegenerative diseases affect the DLPFC region equally at end-stages of disease.

**Figure 5.**
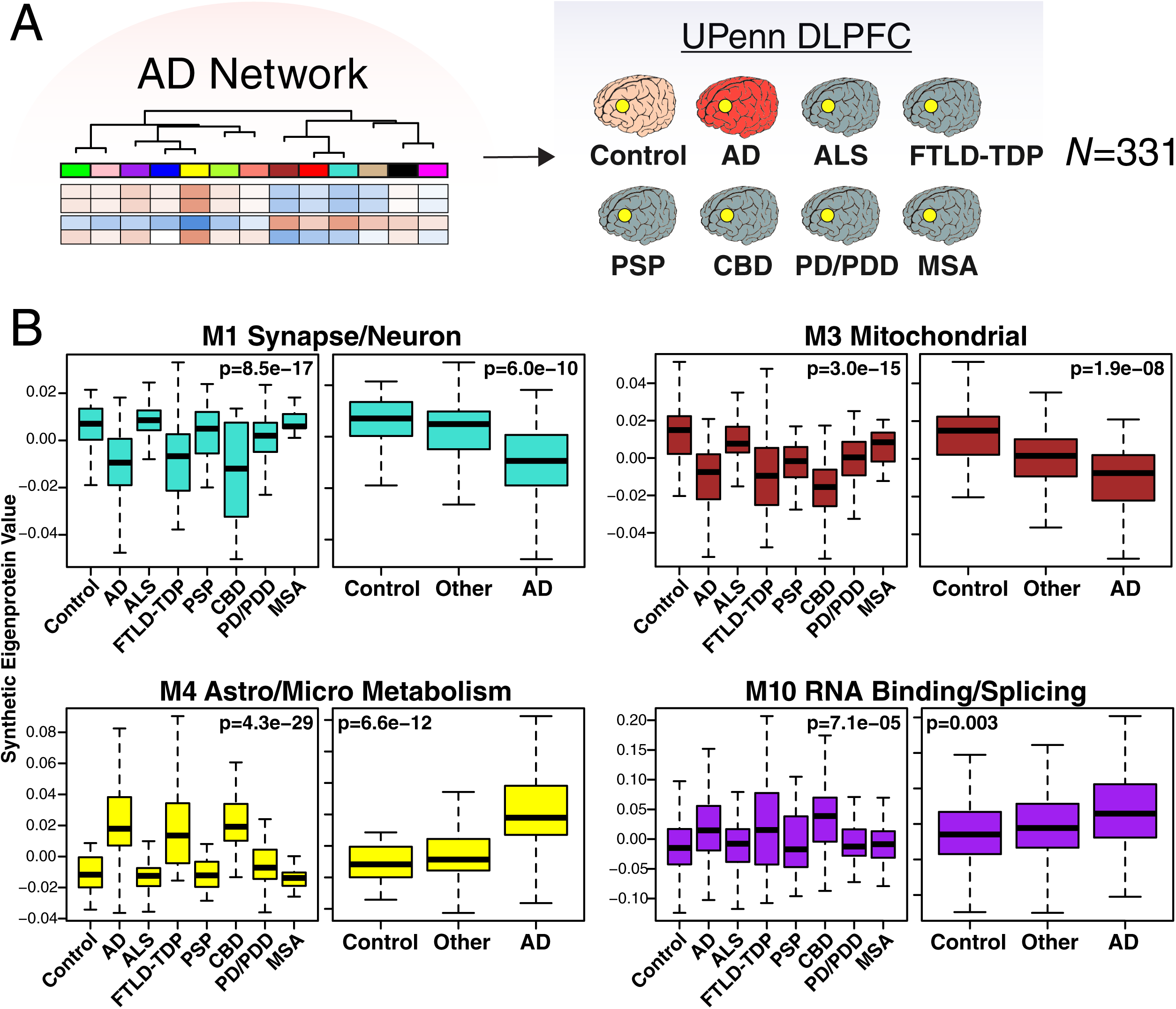
AD Protein Network Module Changes in Other Neurodegenerative Diseases. (A, B) Protein levels were measured in DLPFC from control (*n*=46), AD (*n*=49), amyotrophic lateral sclerosis (ALS, *n*=59), frontotemporal lobar degeneration with TAR DNA-binding protein 43 inclusions (FTLD-TDP, *n*=29), progressive supranuclear palsy (PSP, *n*=27), corticobasal degeneration (CBD, *n*=17), Parkinson’s disease and Parkinsons’s disease dementia (PD/PDD, *n*=81), and multiple system atrophy (MSA, *n*=23) cases from the University of Pennsylvania Brain Bank, and used to analyze AD protein network module changes in different neurodegenerative diseases. (B) A synthetic eigenprotein was created for each AD network module as described in Figure 2 and measured by disease group in the UPenn cohort. Synthetic eigenprotein analyses for modules M1, M3, M4, and M10 are shown. Analyses for all modules are provided in **Extended Data Figure 10**. Differences in module synthetic eigenproteins were assessed by Kruskal-Wallis one-way ANOVA. Differences between AD and other case groups were assessed by Dunnett’s test, the results of which are provided in **Supplementary Table 5**. Boxplots represent the median, 25^th^, and 75^th^ percentiles, and whiskers represent measurements to the 5^th^ and 95^th^ percentiles.

### The M4 Astrocyte/Microglial Metabolism Module is Enriched in AD Genetic Risk Factors and Markers of Anti-Inflammatory Disease-Associated Microglia

One difficulty in analyzing proteomic changes in AD brain, and indeed in any human disease post-mortem tissue, is ascribing the observed changes to causes or consequence of disease—in this case, neurodegeneration. A useful approach to this problem is to assess for enrichment of genetic risk factors that are associated with the disease in question across the disease network, on the assumption that network modules that are enriched in these risk factor gene products may contribute to or reflect upstream, rather than downstream, pathophysiology^40-42^. To this end, we applied an algorithm to calculate a weighted disease risk score for proteins according to their linkage disequilibrium with AD-associated single nucleotide polymorphisms (SNPs) discovered through AD genome wide association studies (GWAS)^43^. We then calculated whether a given AD network module was enriched in these risk factor proteins. We found that the M2 myelin/oligodendrocyte and M4 astrocyte/microglial metabolism modules were significantly enriched in gene products contained within AD risk factor loci (Figure 6A), suggesting that the biological functions or processes reflected by these protein co-expression modules may serve causative roles in AD.

**Figure 6.**
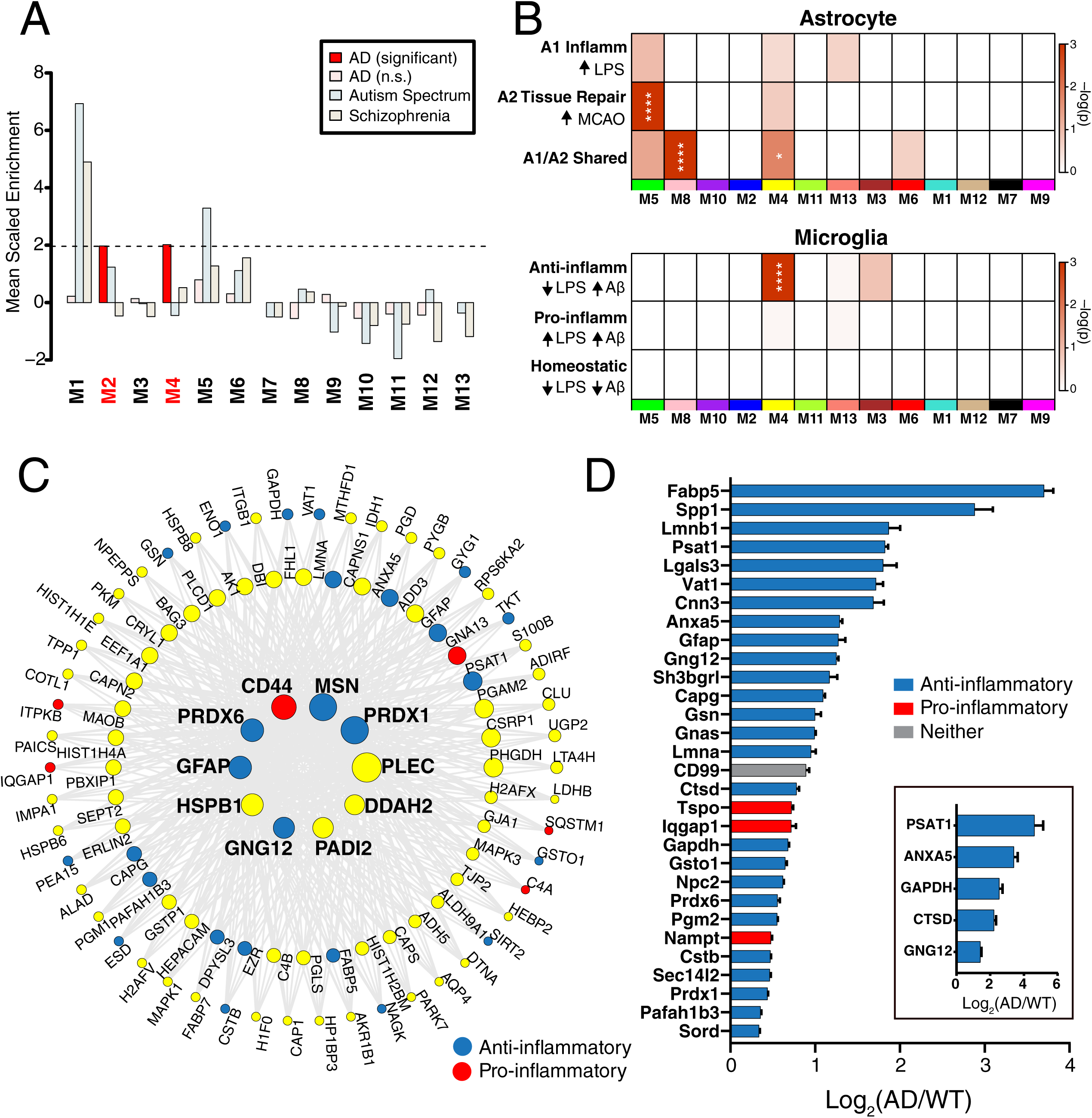
The M4 Astrocyte/Microglial Metabolism Module is Enriched in AD Genetic Risk Factors and Markers of Anti-Inflammatory Disease-Associated Microglia. (A-D) Enrichment of proteins contained within genomic regions identified by genome wide association studies (GWAS) as risk factors for AD, autism spectrum disorder, and schizophrenia was calculated for each module in the AD protein network (A). Modules highlighted in dark red were significantly enriched for AD risk factors, and not for risk factors associated with autism spectrum disorders or schizophrenia. The horizontal dotted line indicates a *z* score level of enrichment of 1.96, or false discovery rate (FDR) *q* value <0.05, above which enrichment was considered statistically significant. (B) Enrichment of astrocyte (top) and microglia (bottom) phenotypic markers in AD protein network modules. (Top) Astrocyte phenotype markers indicating upregulation in response to acute injury with lipopolysaccharide (LPS) (A1 Inflammatory), middle cerebral artery occlusion (MCAO) (A2 Tissue repair), or both types of acute injury (A1/A2 Shared) in a mouse model^46^ were assessed for enrichment in AD network modules. (Bottom) Microglia markers from an mRNA co-expression analysis that are altered after challenge with LPS and/or amyloid-β plaque deposition in mouse models^47^ were assessed for enrichment in AD network modules (Anti-inflammatory, decrease with LPS administration and increase with plaque deposition; Pro-inflammatory, increase with LPS administration and increase with plaque deposition; Homeostatic, decrease with LPS administration and decrease with plaque deposition). Module enrichment was determined by Fisher’s exact test with Benjamini-Hochberg correction. \**P* < 0.05, \*\**P* < 0.01, \*\*\**P* < 0.01, **** *P* < 0.0001. (C) The top 100 proteins by module eigenprotein correlation value (kME) in module M4. The size of each circle indicates the relative kME. Those proteins with the largest kME are considered “hub” proteins within the module. Proteins highlighted in blue are upregulated in A2 tissue repair astrocyte and anti-inflammatory microglia; proteins highlighted in red are upregulated in A1 inflammatory astrocyte and pro-inflammatory microglia. Additional such proteins are provided in **Supplementary Table 6**. (D) The top 30 most differentially abundant microglial transcripts in an AD mouse model^51^ that overlap with proteins in the M4 module, colored as shown in (C). M4 proteins that overlap with transcripts elevated in microglia undergoing active amyloid-β plaque phagocytosis^52^ are provided in Supplementary Figure 9. (Inset) Transcript elevations validated at the protein level in microglia undergoing active amyloid-β plaque phagocytosis^52^.

Given the strong AD trait associations of the M4 astrocyte/microglial metabolism module and its enrichment in AD genetic risk factors, we decided to more deeply investigate the cell type nature of this co-expression module. Astrocyte and microglia phenotypes are known to be heterogenous and dynamic, and dependent upon environmental context and stimuli^44,45^. One common categorization of astrocyte phenotypes is into deleterious pro-inflammatory “A1” astrocytes, such as the phenotype adopted in response to challenge with the pro-inflammatory molecule lipopolysaccharide (LPS), and protective “A2” astrocytes, such as the phenotype adopted after ischemic injury by middle cerebral artery occlusion^46^. Similarly, microglia are known to adopt a number of different phenotypes, both deleterious and protective^45,47-50^. Although expression of the M4 astrocyte/microglia metabolism module is increased with progression from a normal to an AD disease state, and a majority of the most significantly increased proteins in AD are members of this module (Supplementary Figure 6), it is unclear whether these glial responses are deleterious or protective. To better understand the role of these glial cell type responses in AD, we first examined differential expression of astrocyte and microglia protein markers in AD brain by the types of cellular phenotypes with which they are associated in AD animal models. We found that for both astrocytic markers (Supplementary Figure 7) and microglial markers (Supplementary Figure 8), there appeared to be a bias towards expression of markers that are generally considered to be protective. We formally tested this observation with marker over-representation analysis in the AD network (Figure 6B, **Supplementary Table 6**). Microglial protein markers that are increased in response to amyloid-β plaques but decreased in response to LPS—or markers of anti-inflammatory disease-associated microglia^47^—were significantly enriched in the M4 module. Astrocyte markers were more mixed in module M4, with a majority of markers being shared between A1 and A2 phenotypes. Interestingly, a related module to M4 that also had some enrichment in astrocyte cell-type markers and was increased in AD—the M5 endothelial/microglial extracellular matrix module—was significantly enriched in A2 markers. Astrocyte and microglia phenotype markers that overlap with the top 100 proteins by module eigenprotein correlation value in the M4 module are shown in Figure 6C. The majority of these markers were from microglia (**Supplementary Table 6**). To further validate these findings, we analyzed whether these markers were increased at both the transcript and protein levels in acutely isolated microglia from AD mouse models^51,52^. The top 30 most differentially abundant microglial transcripts corresponding to proteins in the M4 module were found to be heavily biased toward an anti-inflammatory phenotype (Figure 6D, **Supplementary Table 6**). Furthermore, many of the disease-associated M4 microglial protein markers were found to be increased in microglia undergoing active amyloid plaque phagocytosis (Supplementary Figure 9, **Supplementary Table 6**)^52^. In summary, we found that the M4 astrocyte/microglial metabolism module was enriched in AD genetic risk factors, and that microglia cell type markers within M4 appeared to be biased towards a protective anti-inflammatory, rather than a deleterious pro-inflammatory, microglial phenotype.

### M4 Astrocyte/Microglial Metabolism Module Proteins Can Be Measured in Cerebrospinal Fluid and May Serve as Potential AD Biomarkers

The ability to assess and monitor pathological brain changes in the preclinical and clinical stages of AD is limited, especially in the asymptomatic phase of the disease. A current focus of AD biomarker research is on measurement of brain amyloid-β and tau protein dynamics as assessed by radiolabeled tracers to aggregated forms of these proteins, in addition to established measurements of amyloid-β and tau protein levels in cerebrospinal fluid (CSF)^53,54^. Fluid biomarkers of neurodegeneration are also in development^55^. However, there are currently no fluid biomarkers that are clinically available to assess AD brain pathophysiology beyond amyloid-β and tau. To explore whether proteins from the M4 astrocyte/microglial metabolism module might also be able to serve as AD fluid biomarkers, we analyzed CSF from two separate cohorts: one cohort of 297 subjects consisting of controls and AD patients (Cohort 1), and a second cohort of 96 subjects classified into control, AsymAD, and AD (Cohort 2). Subjects in both cohorts were classified by the “A/T/N” AD biomarker classification framework (Figure 7A)^56^. We measured 532 CSF protein levels in discovery Cohort 1 using a TMT-MS approach without prior pre-fractionation and without depletion of highly abundant proteins in order to avoid potential depletion measurement artifacts^57,58^. Of these 532 proteins, we observed 22 that mapped to the M4 astrocyte/microglial metabolism module in brain (**Extended Data Figure 13**). All of them showed either an increase in AD or no change, with 10 reaching statistical significance at *p* < 0.05. Only one, cathepsin D (CTSD), showed a trend in the opposite direction. Of note, CTSD had the lowest module eigenprotein correlation value to M4 of all M4 proteins measured in CSF (kME 0.3). The most significantly increased M4 module proteins observed in Cohort 1 are shown in Figure 7B, and include the M4 hub proteins CD44, peroxiredoxin-1 (PRDX1), and dimethylarginine dimethylaminohydrolase-2 (DDAH2), in addition to the metabolic proteins lactate dehydrogenase B-chain (LDHB) and pyruvate kinase (PKM) involved in glycolysis. To validate these findings, and to assess whether the observed changes in CSF levels of M4 proteins occur prior to the development of cognitive impairment, we analyzed subjects in Cohort 2, approximately one-third of which had AsymAD. AsymAD was defined as CSF levels of amyloid-β, total tau, and phospho-tau consistent with an AD diagnosis, but without cognitive impairment. We were able to measure 792 proteins in Cohort 2; 27 mapped to the M4 astrocyte/microglial metabolism module in brain (**Extended Data Figure 14**). Of these 27 proteins, 17 overlapped with M4 proteins measured in discovery Cohort 1, and showed the same direction of change in AD CSF. In addition, many also showed significant or trend elevations in AsymAD, including CD44, LDHB, and PKM, and correlated with cognitive function (Figure 7C). In summary, we were able to measure multiple M4 astrocyte/microglial metabolism module protein members in human CSF by mass spectrometry without fractionation or prior depletion of highly abundant proteins. A number of these proteins were elevated in AsymAD and AD, including M4 hub proteins CD44, PRDX1, and DDAH2.

**Figure 7.**
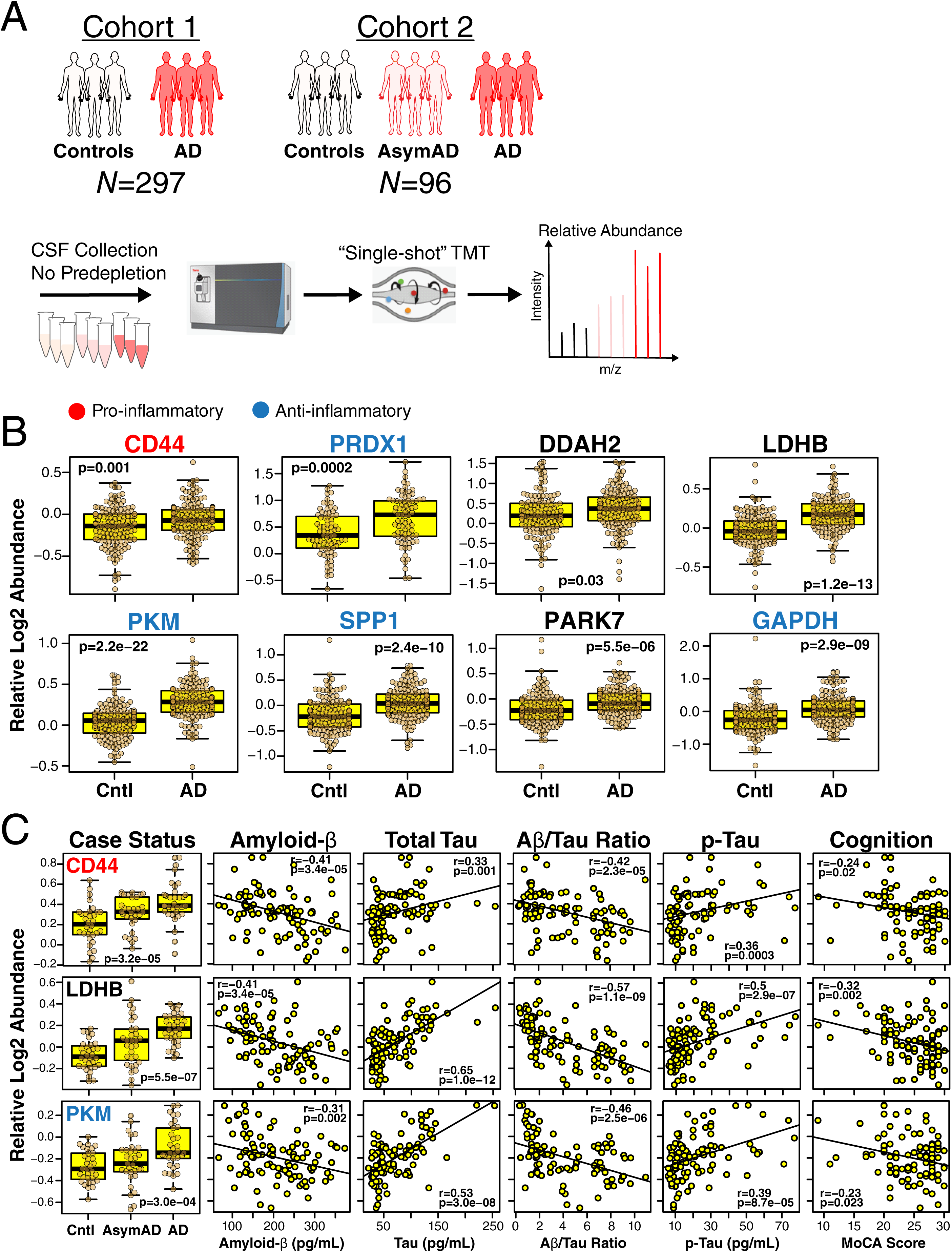
M4 Astrocyte/Microglial Metabolism Module Protein Levels Are Elevated in AsymAD and AD CSF. (A-C) Approach to analysis of M4 proteins in CSF from two different cohorts (A). CSF in Cohort 1 (*n*=297) was obtained from subjects with normal CSF amyloid-β and tau levels (controls) and patients with low amyloid-β, elevated tau levels, and cognitive impairment (AD). CSF in Cohort 2 (*n*=96) was obtained from control subjects and AD patients as defined in Cohort 1, as well as subjects with CSF amyloid-β and tau levels that met criteria for AD but who were cognitively normal at the time of collection. CSF was analyzed without prior pre-fractionation or depletion of highly abundant proteins; relative protein levels were measured by TMT-MS. (B) Relative CSF protein levels of selected M4 module members in Cohort 1. Protein names are colored according to pro-inflammatory (red) or anti-inflammatory (blue) classification. Additional M4 protein measurements, as well as trait correlations for the measured proteins, are provided in **Extended Data Figure 13**. (C) Relative CSF protein levels of selected M4 module members in Cohort 2. Protein names are colored as in (B). Additional measurements and trait correlations are provided in **Extended Data Figure 14**. Differences in protein levels were assessed by Welch’s *t* test (B) or Kruskal-Wallis one-way ANOVA (C). Correlations were performed using biweight midcorrelation. Boxplots represent the median, 25^th^, and 75^th^ percentiles, and whiskers represent measurements to the 5^th^ and 95^th^ percentiles. Cntl, control; AsymAD, asymptomatic Alzheimer’s disease; AD, Alzheimer’s disease; TMT, tandem mass tag; MoCA, Montreal Cognitive Assessment (higher scores represent better cognitive function).

## Discussion

In this study, we analyzed more than 2000 brains by mass spectrometry-based proteomics to arrive at a consensus view of the proteomic changes that occur in brain during progression from normal to asymptomatic and symptomatic AD states. We find that the protein co-expression families most strongly correlated to disease reflect synaptic, mitochondrial, RNA binding/splicing, and astrocyte/microglial metabolism biological functions, with astrocyte/microglial metabolism most significantly associated with AD compared to other biological processes and functions. Increases in expression level of the M4 astrocyte/microglial metabolism module are observed with aging, but are stronger in AD, reflecting shared biology between “normal” aging and AD. The M4 module is enriched in AD genetic risk factors, indicating a potential causative role for this protein co-expression module in disease pathogenesis, and appears to serve a protective anti-inflammatory function in model systems, suggesting that genetic risk factor polymorphisms that cluster in this module may induce a loss-of-function phenotype. M4 astrocyte/microglial module proteins are increased in AsymAD and AD CSF, suggesting that proteins within the M4 module may serve as useful biomarkers for staging AD progression and for development of novel therapeutic approaches to the disease.

The robustness of the AD brain protein co-expression network described in this study is derived from both the multi-center cohort study design and the proteomics analysis pipeline. Neuropathological examination and tissue dissection were performed at each center by different individuals prior to tissue transfer to one center for proteomic analysis. Tissue from each cohort was prepared and analyzed by mass spectrometry-based proteomics separately from other cohorts, oftentimes in multiple batches per cohort. All cohorts were searched together to enforce parsimony of protein identifications across cohorts. The AD protein co-expression network was generated from proteins that could be quantified in at least 50% of cases after batch and cohort normalization and regression of covariates, and therefore reflects the core co-expression patterns in AD brain. The protein co-expression modules are not sensitive to differences in mass spectrometry instrumentation or protein quantitation approaches, and are not significantly influenced by potential case selection bias given that all modules were significantly preserved in the ROS/MAP cohort, and were generated by including other well-established epidemiological cohorts from across the U.S. such as ACT and BLSA. Furthermore, the protein co-expression modules are not significantly influenced by regional tissue variation among temporal cortex, precuneus, and DLPFC brain regions. Indeed, we observed that all of the larger modules were highly preserved in both temporal cortex and precuneus, with preservation *p* values approaching zero in both regions. This suggests that the biological processes and cell types driving the co-expression patterns in AD brain are highly shared among these brain regions. Future proteomic analyses that include other brain regions less affected in late-onset AD (e.g, visual cortex) would be informative to further explore potential protective processes that may be important for regional vulnerability in AD. Finally, the co-expression modules are robust to the algorithm used to identify them, as an orthogonal data dimension reduction technique was able to separately identify the protein communities represented by the modules in the co-expression network.

Six co-expression modules showed significant and robust correlations with amyloid-β plaque load, tau tangle burden, and cognitive function: the M1 synapse/neuron, M3 mitochondrial, M4 astrocyte/microglial metabolism, M5 endothelial/microglial extracellular matrix, M6 cytoskeleton, and M10 RNA binding/splicing modules. We chose to highlight four of these six modules in the subsequent analyses, but the M5 endothelial/microglial extracellular matrix and M6 cytoskeleton modules also deserve attention as protein co-expression families relevant to AD. Similarly, the M11 chaperone/protein folding module and M12 module of unknown function also were significantly correlated with all measures except for CDR, and deserve further study. We did not observe strong independent effects of different *APOE* alleles on protein co-expression modules, although there appeared to be a trend for a protective effect of the E2 allele on many of the modules, as we have observed previously^59^, and a significant protective effect of E2 on the M2 myelin/oligodendrocyte module. Although ApoE4 can cause mitochondrial toxicity^60,61^ and altered brain energy metabolism^62-64^, and ApoE variants can have important effects on brain inflammation^59,65,66^, these changes may not be evident at end-stage AD. Further analysis of *APOE* effects on the proteome of the aging brain and in early AD disease states is warranted.

The M1 synapse/neuron and M4 astrocyte/microglial metabolism modules were significantly associated with aging, a finding consistent with mRNA co-expression studies^11,49^. In fact, the M1 module was the most strongly correlated module with aging among all network modules, consistent with the known generalized cortical volume loss observed in most aged individuals^67^. However, it is interesting to note that while the M1 module declined in aging, the M3 mitochondrial module did not, even though mitochondria are highly populated at the synapse^68^. This may suggest that “normal” aging is associated with maintenance of mitochondrial number and/or function, whereas AD is associated with loss of mitochondrial number/function. This finding stands in contrast to mRNA co-expression results in normal aging, where decreases in synapse/neuron modules are accompanied by decreases in mitochondrial metabolism^11^, and highlights a potential divergence between mitochondrial RNA and protein co-expression. Similar to M3, the M10 RNA binding/splicing module was also not changed in normal aging, but was altered in AD. RNA binding protein aggregation and dysfunctional protein splicing is a known feature of AD^24,69,70^, but has not been well-studied in the normal aging brain. A previous study found both common and unique changes in alternatively spliced transcripts between normal aging and AD^71^. We found that the M10 RNA binding/splicing module was only modestly elevated in AsymAD, and like the M3 mitochondrial module, M10 may more strongly reflect a late AD pathophysiologic process. However, we and others have also observed that certain RNA-binding proteins, such as the U1 spliceosome proteins, aggregate early in the disease^72,73^. Further study of brain RNA binding protein and splicing biology in aging and AD is clearly required.

We assessed the disease specificity of the AD protein co-expression network by analyzing how the protein network modules changed in six other neurodegenerative diseases encompassing diverse brain pathologies. One caveat to this analysis is that we analyzed only DLPFC, which is not equally affected in all the neurodegenerative diseases we assessed. With this caveat in mind, we observed that FTLD-TDP and CBD had the most similar network changes to AD, suggesting that these clinicopathologic entities are fundamentally related to AD at the brain proteomic level. It is interesting that TDP-43 pathology and a four-repeat tauopathy (CBD) led to similar network changes as AD, whereas synuclein pathology and a different four-repeat tauopathy (PSP) did not, at least at end-stage disease. A proteomic relationship between AD and FTD is supported by the fact that mutations in the triggering receptor expressed on myeloid cells 2 (TREM2) protein cause microglial dysfunction and lead to AD^74,75^, whereas mutations in the progranulin (PGRN) protein also cause microglial dysfunction and lead to FTD^76-78^. Further studies comparing frontal predominant AD, FTLD-TDP, and FTLD-tau cases would be informative to assess the degree to which the underlying neuropathology observed at autopsy is related to differences in proteomic network changes in the DLPFC region.

We found that two AD protein network modules—the M2 myelin/oligodendrocyte module and the M4 astrocyte/microglial metabolism module—were significantly enriched for AD genetic risk factors, suggesting that the biological processes reflected by these protein co-expression families are involved in disease etiology. This finding is consistent with other reports of genetic risk factor enrichment in AD networks^23,79^. Interestingly, while the M4 module showed the strongest trait relationships to AD, the M2 module was only slightly increased in AD and did not correlate as strongly with neuropathology or cognitive function as other AD network modules. This finding is consistent with our prior proteomic studies in DLPFC^23,24,59^. A previous study investigating the relationship of a similar myelin/oligodendrocyte mRNA brain co-expression module to AD found that expression of this module decreased in both AD and PSP in temporal cortex, with stronger changes in PSP^80^. Another mRNA network study observed downregulation of oligodendrocyte module hubs in AD^81^. We observed an increase in this module in AD, with no changes in PSP. The discrepancy in our findings may relate to possible differences between DLPFC and temporal cortex in this co-expression module, differences in module membership that may be important for disease relationship, or a divergence between mRNA and protein expression in M2 module members. In fact, the basis for disease causality regarding the M2 myelin/oligodendrocyte module may relate more to dysregulation of module member co-expression, rather than to the actual change in direction of expression^81^.

A key finding from our proteomic study is that glial biology—and microglial biology in particular— is a likely causal driver of AD pathogenesis. This finding is consistent with the results of other recent protein co-expression analyses of AD^23,79^. The AD protein network module most strongly associated with AD is enriched in astrocyte and microglial proteins, and is also enriched in proteins associated with genetic risk for AD. The M4 astrocyte/microglial metabolism module increases in AsymAD and correlates most strongly with cognitive impairment, suggesting that the biological changes reflected by this module occur early in the disease and have significant functional consequence on progression to dementia. A natural assumption would be that increases in M4 module expression levels are deleterious to brain health, and that potential therapies targeting reduction of M4 would likely be beneficial in AD. However, several lines of evidence support a possible protective role of this co-expression module. An important observation is that AD genetic risk alleles, which are more likely to cause loss-of-function changes rather than gain-of-function changes, are enriched in the M4 module. The M4 module is also enriched in microglial markers that are upregulated in response to amyloid-β deposition and downregulated in response to LPS, indicating that the microglial response as reflected in M4 module expression is likely biased towards an anti-inflammatory disease-associated phenotype^47^. Many M4 proteins are elevated in microglia that are undergoing plaque phagocytosis, which is consistent with the strong association of M4 expression with CERAD score. Notably, when we compare our findings to a prior proteomic study that quantified levels of plaque-associated proteins in normal *versus* rapidly-progressive AD^82^, 7 out of the top 10 plaque-associated proteins most significantly *decreased* in rapidly-progressive AD are found in the M4 module, including M4 hubs MSN and PLEC. This is consistent with the finding that early microglial activation in response to amyloid plaques, as assessed by *in vivo* microglial imaging studies, is correlated with increased grey matter volume and reduced rate of cognitive decline^83,84^. Interestingly, the degree of astrogliosis surrounding plaques seems to be positively correlated with improved cognitive function not only in AD, but also in normal aging individuals^85^. Taken together, these findings suggest that lack of an M4 astrocyte/microglial response to plaques in preclinical or clinical AD may lead to more rapid cognitive decline. In this context, it is worth noting that AD risk factor mutations in the microglial TREM2 receptor, which regulates the microglial response to amyloid-β plaques, lead to a reduced microglial response to plaque pathology^86-90^. Importantly, recent studies have suggested that TREM2 is also necessary for enabling microglia to increase their metabolic rate in response to various stimuli^91^, which include not only plaques, but also myelin debris and the abnormal lipid environment of apoptotic neurons^75,92,93^. The M4 module is highly enriched for metabolic proteins that are likely upregulated to meet the metabolic response demand to such stimuli. The relationship between different immune response states and AD is clearly complex, and is a subject that requires further investigation. The ability to fully recapitulate the M4 module in AD animal models would help to advance our mechanistic understanding of M4 module relationships with AD neuropathology and cognitive function, and whether it should be targeted for therapeutic reduction or enhancement. Collaborative efforts to generate new AD animal models that better reflect AD pathophysiology are currently underway^94^; however, recent work on microglial and astrocyte function in current mouse AD models generally supports the findings in this study, and a hypothesis that M4 is associated with a largely protective response^75,87,95,96^.

As many co-expression relationships in tissue are driven by cell type^23,26,97^, we infer that the biological processes reflected in the M4 module are related to sugar metabolism primarily in astrocytes and microglia, and reflect increased metabolic activity in these cell types in AsymAD and AD states. This is a hypothesis that requires further testing, likely through single cell studies, but is consistent with the astrogliosis response in AD and the fact that glycolysis is performed predominantly within glia in the brain^98^. In the context of M4 glycolytic metabolism, it is interesting to speculate on the origin of the reduced fluoro-2-deoxy-D-glucose positron emission tomography (FDG-PET) signal observed in AD brain. Recent animal model work has suggested that loss of a proper microglial response through mutations in either TREM2 or PGRN leads to reduced cerebral glucose metabolism^99^, and that normal astrocyte function is also important for brain glucose uptake^100,101^. Therefore, it is possible that the failure of a proper astroglial compensatory or protective response in AD may lead to general neuronal metabolic failure in susceptible brain regions, as assessed by the FDG-PET signal, and cognitive deterioration. The precise mechanism by which such metabolic failure occurs in AD is not known, but could involve breakdown of the astrocyte-neuron lactate shuttle, upon which neurons are highly dependent for normal synaptic activity^98^. Evidence also exists that primary metabolic failure could occur in neurons prior to involvement of astroglia, especially in ApoE4 carriers^61,102-104^. Notably, we observed larger relative changes in AsymAD in both the M4 astrocyte/microglial metabolism and M3 mitochondrial modules in precuneus compared to DLPFC, consistent with early metabolic dysfunction in this brain region in AD^105^. The primary cause of abnormal cerebral glucose metabolism in AD as measured by FDG-PET remains an area for continued study.

Given the pressing need for new biomarkers that reflect AD pathophysiology beyond amyloid and tau, we assessed whether M4 module proteins could be detected in CSF, and whether their levels changed in AD and AsymAD. We observed multiple M4 members in CSF that validated in two separate cohorts; nearly all of these proteins increased in AD, consistent with the direction of change of the M4 module in AD brain. Many of the proteins also were elevated in AsymAD, suggesting that measurement of M4 elevation in biofluids in early stages of the disease—when diagnostic information and therapeutic intervention are likely to be of highest utility—is feasible. Many of the most significantly elevated M4 protein in CSF are involved in glycolysis, including LDHB, PKM, and glyceraldehyde 3-phosphate dehydrogenase (GAPDH). Elevations in PRDX1, DDAH, and protein/nucleic acid deglycase DJ-1 (PARK7) were also observed, all of which are important anti-oxidant effector proteins^106-108^ and are likely elevated in concert with increased glycolytic flux. LDHB, PKM, and DDAH1 have recently been reported as promising AD CSF biomarkers^109,110^. Separate from the metabolic pathway, we observed increased levels of M4 proteins osteopontin (SPP1), dickkopf-related protein 3 (DKK3), and CD44. SPP1 has been nominated as an AD CSF biomarker in previous studies^111-114^. SPP1 is known to be involved in tissue repair^115^, and promotes phagocytosis of amyloid-β and an anti-inflammatory microglial phenotype^116,117^. It is closely associated with the CD44 receptor^118^, which is highly expressed on the surface of activated astrocytes^119^. While M4 markers may not be entirely specific for AD given elevation of the M4 module in FTD and CBD, they may allow for assessment of an injury response in AD in conjunction with amyloid and tau biomarkers, and serve as useful biomarkers for other neurodegenerative dementias in addition to AD. The degree of overlap between and among groups for many of the potential M4 markers was large, which may be due in part to reduction in the absolute differences in relative measured protein levels caused by batch correction. Despite this possibility, it is likely that panels consisting of multiple markers in CSF that map to M4 and other modules will provide better discrimination among groups than individual proteins^120^. Measurement of additional M4 markers in biofluids is undoubtedly possible, as our mass spectrometry measurements were performed on unfractionated CSF not depleted of highly abundant proteins. Monitoring multiple M4 protein levels in biofluid may provide a robust measure of target engagement for AD therapies.

In summary, our comprehensive study on more than 2000 brains and nearly 400 CSF samples provides a consensus view of the proteomic network landscape of AD and the biological changes associated with asymptomatic and symptomatic stages of the disease, and highlights the central role of glial biology in the pathogenesis of the disease. Programs that target this biology hold promise for AD drug therapy and biomarker development, especially those that target pro- and anti-inflammatory astrocytes and microglia.

## Supporting information

Johnson et al 2020 Extended Data Figures

## Acknowledgments

We are grateful to those who agreed to donate their brains and CSF for research, and who participated in the described observational studies. This study was supported by the following NIH funding mechanisms: R01AG053960, R01AG057911, R01AG061800, RF1AG057471, RF1AG057470, R01AG061800, R01AG057911, R01AG057339, U01AG061357, P50AG025688, RF1AG057470, RF1AG051633, P30AG10161, R01AG15819, R01AG17917, U01AG61356, R01AG056533, K08NS099474, U01AG046170, RF1AG054014, RF1AG057440, R01AG057907, U01AG052411, P30AG10124, U01AG046161, R01AG050631, R01AG053960, R01AG057339, U01AG061357, P50AG005146, U24NS072026, and P30AG19610.

## Data Availability

All raw data, case traits and analyses (differential and co-expression) related to this manuscript are available at https://www.synapse.org/consensus. The results published here are in whole or in part based on data obtained from the AMP-AD Knowledge Portal (https://adknowledgeportal.synapse.org). The AMP-AD Knowledge Portal is a platform for accessing data, analyses, and tools generated by the Accelerating Medicines Partnership (AMP-AD) Target Discovery Program and other National Institute on Aging (NIA)-supported programs to enable open-science practices and accelerate translational learning. The data, analyses and tools are shared early in the research cycle without a publication embargo on secondary use. Data is available for general research use according to the following requirements for data access and data attribution (https://adknowledgeportal.synapse.org/#/DataAccess/Instructions). ROSMAP resources can be requested at www.radc.rush.edu.

## Code Availability

The algorithm used for batch correction is fully documented and available as an R function, which can be downloaded from https://github.com/edammer/TAMPOR.

## Author Contributions

ECBJ, EBD, DMD, LP, MZ, AG, VAP, and NTS and designed experiments; DMD, LP, MZ, LY, AG, and VAP carried out experiments; ECBJ, EBD, MZ, VAP, and NTS analyzed data; DMD, LP, LAH, BW, JCT, MT, TJM, EBL, JQT, TGB, EMR, VH, MW, ES, BZ, DWD, NE, TG, VAP, PLD, DAB, TSW, SR, IH, and JMS provided advice on the interpretation of data; ECBJ wrote the manuscript with input from coauthors; JCT, TJM, JQT, TGB, VH, MW, DWD, DAB, and IH provided tissue samples; AIL, JJL, and NTS supervised the study. All authors approved the final manuscript.

## Methods

### Brain Tissue Samples and Case Classification

Brain tissue used in this study was obtained from the autopsy collections of the Baltimore Longitudinal Study of Aging^121^, Banner Sun Health Research Institute^122^, Mount Sinai School of Medicine Brain Bank, Adult Changes in Thought Study, Mayo Clinic Brain Bank, Religious Orders Study and Rush Memory and Aging Project^123^, University of Pennsylvania School of Medicine Brain Bank, and the Baltimore Coroner’s Office. Tissue was from the dorsolateral prefrontal cortex (Brodmann Area 9 where available), or temporal cortex and precuneus regions where indicated. Human postmortem tissues were acquired under proper Institutional Review Board (IRB) protocols at each respective institution. Postmortem neuropathological evaluation of neuritic plaque distribution was performed according to the Consortium to Establish a Registry for Alzheimer’s Disease (CERAD) criteria^25^, while extent of spread of neurofibrillary tangle pathology was assessed with the Braak staging system^124^. Other neuropathologic diagnoses were made in accordance with established criteria and guidelines^125,126^. All case metadata, including age, sex, post-mortem interval, cognitive function, *APOE* genotype, neuropathological criteria, and disease status, are provided in **Supplementary Table 1**. Case classification harmonization across cohorts was performed using the following rubric: cases with CERAD 0-1 and Braak 0-3 without dementia at last evaluation were defined as control (if Braak equals 3, then CERAD must equal 0); cases with CERAD 1-3 and Braak 3-6 without dementia at last evaluation were defined as AsymAD; cases with CERAD 2-3 and Braak 3-6 with dementia at last evaluation were defined as AD. Dementia was defined as MMSE <24, CASI score <81, or CDR ≥1, based on prior comparative study^127^. Mayo and UPenn cases were not included in the case harmonization scheme, and therefore preservation of consensus network modules in these cohorts provides an additional degree of robustness.

### Brain Tissue Homogenization and Protein Digestion

Procedures for tissue homogenization for all tissues were performed essentially as described^23,27^. Approximately 100 mg (wet tissue weight) of brain tissue was homogenize in 8 M urea lysis buffer (8 M urea, 10 mM Tris, 100 mM NaHPO4, pH 8.5) with HALT protease and phosphatase inhibitor cocktail (ThermoFisher) using a Bullet Blender (NextAdvance). Each Rino sample tube (NextAdvance) was supplemented with ∼100 μL of stainless steel beads (0.9 to 2.0 mm blend, NextAdvance) and 500 μL of lysis buffer. Tissues were added immediately after excision and samples were then placed into the bullet blender at 4 °C. The samples were homogenized for 2 full 5 min cycles, and the lysates transferred to new Eppendorf Lobind tubes. Each sample was then sonicated for 3 cycles consisting of 5 s of active sonication at 30% amplitude, followed by 15 s on ice. Samples were then centrifuged for 5 min at 15,000 x *g* and the supernatant transferred to a new tube. Protein concentration was determined by bicinchoninic acid (BCA) assay (Pierce). For protein digestion, 100 μg of each sample was aliquoted and volumes normalized with additional lysis buffer. For the ROS/MAP cohort, an equal amount of protein from each sample was aliquoted and digested in parallel to serve as the global pooled internal standard (GIS) in each TMT batch, as described below. Similarly, GIS pooled standards were generated from the Banner, MSSB, Mayo, Aging, and UPenn cohorts. Samples were reduced with 1 mM dithiothreitol (DTT) at room temperature for 30 min, followed by 5 mM iodoacetamide (IAA) alkylation in the dark for another 30 min. Lysyl endopeptidase (Wako) at 1:100 (w/w) was added and digestion allowed to proceed overnight. Samples were then 7-fold diluted with 50 mM ammonium bicarbonate. Trypsin (Promega) was then added at 1:50 (w/w) and digestion was carried out for another 16 h. The peptide solutions were acidified to a final concentration of 1% (vol/vol) formic acid (FA) and 0.1% (vol/vol) trifluoroacetic acid (TFA), and desalted with a 30 mg HLB column (Oasis). Each HLB column was first rinsed with 1 mL of methanol, washed with 1 mL 50% (vol/vol) acetonitrile (ACN), and equilibrated with 2×1 mL 0.1% (vol/vol) TFA. The samples were then loaded onto the column and washed with 2×1 mL 0.1% (vol/vol) TFA. Elution was performed with 2 volumes of 0.5 mL 50% (vol/vol) ACN.

### Mass Spectrometry Analysis for Label-free Proteomics

Mass spectrometry analyses of MSSB, ACT, BLSA, Banner, Mayo, and UPenn cohorts were performed on a Q-Exactive Plus mass spectrometer essentially as described^23^. Brain-derived tryptic peptides (2 μg) were resuspended in peptide loading buffer (0.1% FA, 0.03% TFA, 1% ACN) containing 0.2 pmol of isotopically labeled peptide calibrants (ThermoFisher 88321). Peptide mixtures were separated on a self-packed C18 (1.9 μm, Dr. Maisch, Germany) fused silica column (25 cm x 75 μM internal diameter; New Objective, Woburn, MA) by a NanoAcquity UHPLC (Waters, Milford, MA) and monitored on a Q-Exactive Plus mass spectrometer (ThermoFisher Scientific, San Jose, CA). Elution was performed over a 120 minute gradient at a rate of 400 nL/min with buffer B ranging from 3% to 80% (buffer A: 0.1% FA and 5% DMSO in water, buffer B: 0.1 % FA and 5% DMSO in ACN). The mass spectrometer cycle was programmed to collect one full MS scan followed by 10 data dependent MS/MS scans. The MS scans (300-1800 m/z range, 1,000,000 automatic gain control (AGC), 150 ms maximum ion time) were collected at a resolution of 70,000 at m/z 200 in profile mode, and the MS/MS spectra (2 m/z isolation width, 25% collision energy, 100,000 AGC target, 50 ms maximum ion time) were acquired at a resolution of 17,500 at m/z 200. Dynamic exclusion was set to exclude previous sequenced precursor ions for 30 seconds within a 10 ppm window. Precursor ions with +1 and +6 or higher charge states were excluded from sequencing.

### Label-free Quantification

For the consensus LFQ search, 645 RAW files, including individual cases and pooled GIS samples from the MSSB, ACT, Banner and BLSA cohorts, were uploaded onto the Amazon Web Services (AWS) Cloud and analyzed using MaxQuant v1.6.3.4 with Thermo Foundation 2.0 for RAW file reading capability. The Mayo, BLSA precuneus, Aging, and UPenn cohorts were each searched separately using MaxQuant. The search engine Andromeda was used to build and search a concatenated target-decoy UniProt Knowledgebase (UniProtKB) containing both Swiss-Prot and TrEMBL human reference protein sequences (90,411 target sequences downloaded April 21, 2015), plus 245 contaminant proteins included as a parameter for the Andromeda search within MaxQuant^128^. Methionine oxidation (+15.9949 Da), asparagine and glutamine deamidation (+0.9840 Da), and protein N-terminal acetylation (+42.0106 Da) were variable modifications (up to 5 allowed per peptide); cysteine was assigned a fixed carbamidomethyl modification (+57.0215 Da). Only fully tryptic peptides with up to 2 miscleavages were considered in the database search. A precursor mass tolerance of ±20 ppm was applied prior to mass accuracy calibration, and ±4.5 ppm after internal MaxQuant calibration. Other search settings included a maximum peptide mass of 6,000 Da, a minimum peptide length of 6 residues, and 0.05 Da tolerance for high resolution MS/MS scans. The false discovery rate (FDR) for peptide spectral matches, proteins, and site decoy fraction were each set to 1 percent. Quantification settings were as follows: re-quantify with a second peak-finding attempt after protein identification is complete; match full MS1 peaks between runs; use a 0.7 min retention time match window after an alignment function was found with a 20 minute retention time search space. The label free quantitation (LFQ) algorithm in MaxQuant^129,130^ was used for protein quantitation. The quantitation method considered only razor and unique peptides for protein level quantitation. The total summed protein intensity was also used to assess overall signal drift across samples prior to LFQ normalization.

### Isobaric Tandem Mass Tag (TMT) Peptide Labeling of ROS/MAP Brain Tissues

Prior to TMT labeling, cases were randomized by covariates (age, sex, PMI, diagnosis, etc.), into 50 total batches (8 cases per batch). Peptides from each individual case (n=400) and the GIS pooled standard (n=100) were labeled using the TMT 10-plex kit (ThermoFisher 90406). In each batch, TMT channels 126 and 131 were used to label GIS standards, while the 8 middle TMT channels were reserved for individual samples following randomization. Labeling was performed as previously described^24,27^. Briefly, each sample (containing 100 μg of peptides) was re-suspended in 100 mM TEAB buffer (100 μL). The TMT labeling reagents were equilibrated to room temperature, and anhydrous ACN (256 μL) was added to each reagent channel. Each channel was gently vortexed for 5 min, and then 41 μL from each TMT channel was transferred to the peptide solutions and allowed to incubate for 1 h at room temperature. The reaction was quenched with 5% (vol/vol) hydroxylamine (8 μl) (Pierce). All 10 channels were then combined and dried by SpeedVac (LabConco) to approximately 150 μL and diluted with 1 mL of 0.1% (vol/vol) TFA, then acidified to a final concentration of 1% (vol/vol) FA and 0.1% (vol/vol) TFA. Peptides were desalted with a 200 mg C18 Sep-Pak column (Waters). Each Sep-Pak column was activated with 3 mL of methanol, washed with 3 mL of 50% (vol/vol) ACN, and equilibrated with 2×3 mL of 0.1% TFA. The samples were then loaded and each column was washed with 2×3 mL 0.1% (vol/vol) TFA, followed by 2 mL of 1% (vol/vol) FA. Elution was performed with 2 volumes of 1.5 mL 50% (vol/vol) ACN. The eluates were then dried to completeness using a SpeedVac.

### High-pH Off-line Fractionation of ROS/MAP Brain Tissues

High pH fractionation was performed essentially as described^131^ with slight modification. Dried samples were re-suspended in high pH loading buffer (0.07% vol/vol NH_4_OH, 0.045% vol/vol FA, 2% vol/vol ACN) and loaded onto an Agilent ZORBAX 300 Extend-C18 column (2.1mm x 150 mm with 3.5 µm beads). An Agilent 1100 HPLC system was used to carry out the fractionation. Solvent A consisted of 0.0175% (vol/vol) NH_4_OH, 0.01125% (vol/vol) FA, and 2% (vol/vol) ACN; solvent B consisted of 0.0175% (vol/vol) NH_4_OH, 0.01125% (vol/vol) FA, and 90% (vol/vol) ACN. The sample elution was performed over a 58.6 min gradient with a flow rate of 0.4 mL/min. The gradient consisted of 100% solvent A for 2 min, then 0% to 12% solvent B over 6 min, then 12% to 40 % over 28 min, then 40% to 44% over 4 min, then 44% to 60% over 5 min, and then held constant at 60% solvent B for 13.6 min. A total of 96 individual equal volume fractions were collected across the gradient and subsequently pooled by concatenation^131^ into 24 fractions and dried to completeness using a SpeedVac.

### TMT Mass Spectrometry of ROS/MAP Brain Tissues

All fractions were resuspended in an equal volume of loading buffer (0.1% FA, 0.03% TFA, 1% ACN) and analyzed by liquid chromatography coupled to tandem mass spectrometry essentially as described^132^, with slight modifications. Peptide eluents were separated on a self-packed C18 (1.9 μm, Dr. Maisch, Germany) fused silica column (25 cm × 75 μM internal diameter (ID); New Objective, Woburn, MA) by an Dionex UltiMate 3000 RSLCnano liquid chromatography system (ThermoFisher Scientific) and monitored on an Orbitrap Fusion mass spectrometer (ThermoFisher Scientific). Sample elution was performed over a 180 min gradient with flow rate at 225 nL/min. The gradient was from 3% to 7% buffer B over 5 min, then 7% to 30% over 140 min, then 30% to 60% over 5 min, then 60% to 99% over 2 min, then held constant at 99% solvent B for 8 min, and then back to 1% B for an additional 20 min to equilibrate the column. Buffer A was water with 0.1% (vol/vol) formic acid, and buffer B was 80% (vol/vol) acetonitrile in water with 0.1% (vol/vol) formic acid. The mass spectrometer was set to acquire in data dependent mode using the top speed workflow with a cycle time of 3 seconds. Each cycle consisted of 1 full scan followed by as many MS/MS (MS2) scans that could fit within the time window. The full scan (MS1) was performed with an m/z range of 350-1500 at 120,000 resolution (at 200 m/z) with AGC set at 4×10^5^ and maximum injection time 50 ms. The most intense ions were selected for higher energy collision-induced dissociation (HCD) at 38% collision energy with an isolation of 0.7 m/z, a resolution of 30,000, an AGC setting of 5×10^4^, and a maximum injection time of 100 ms. Five of the 50 TMT batches were run on the Orbitrap Fusion mass spectrometer using the SPS-MS3 method as previously described^27^.

### TMT ROS/MAP Database Searches and Protein Quantification

All RAW files (1,200 RAW files generated from 50 TMT 10-plexes) were analyzed using the Proteome Discoverer suite (version 2.3, ThermoFisher Scientific). MS2 spectra were searched against the UniProtKB human proteome database containing both Swiss-Prot and TrEMBL human reference protein sequences (90,411 target sequences downloaded April 21, 2015), plus 245 contaminant proteins. The Sequest HT search engine was used and parameters were specified as follows: fully tryptic specificity, maximum of two missed cleavages, minimum peptide length of 6, fixed modifications for TMT tags on lysine residues and peptide N-termini (+229.162932 Da) and carbamidomethylation of cysteine residues (+57.02146 Da), variable modifications for oxidation of methionine residues (+15.99492 Da) and deamidation of asparagine and glutamine (+0.984 Da), precursor mass tolerance of 20 ppm, and a fragment mass tolerance of 0.05 Da for MS2 spectra collected in the Orbitrap (0.5 Da for the MS2 from the SPS-MS3 batches). Percolator was used to filter peptide spectral matches (PSMs) and peptides to a false discovery rate (FDR) of less than 1%. Following spectral assignment, peptides were assembled into proteins and were further filtered based on the combined probabilities of their constituent peptides to a final FDR of 1%. In cases of redundancy, shared peptides were assigned to the protein sequence in adherence with the principles of parsimony. Reporter ions were quantified from MS2 or MS3 scans using an integration tolerance of 20 ppm with the most confident centroid setting.

### Parallel Reaction Monitoring (PRM) Analysis

Peptides from brain digests used for the first 3 batches of the untargeted UPenn cohort analysis (equal to 1 µg protein digestion) were used for targeted analysis on an Orbitrap Lumos™ Tribrid™ Mass Spectrometer (ThermoFisher Scientific) fitted with a Nanospray Flex ion source and coupled to a NanoAcuity liquid chromatography system (Waters). The tryptic peptides were resuspended in loading buffer (0.1% TFA, 500 ng/µl), and an external reference peptide mix (Promega) was spiked into the sample at the concentration of 0.5 pmol/µl. The solution (2 µl) was loaded onto a self-packed 1.9 µm ReproSil-Pur C18 (Dr. Maisch) analytical column (New Objective, 50 cm × 75 µm inner diameter; 360 µm outer diameter) heated to 60 °C. The capillary temperature and spray voltage was set at 300 °C and 2.0 kV, respectively. Elution was performed over a 100 min gradient at a rate of 350 nL/min with buffer B ranging from 1% to 32% (buffer A: 0.1% FA in water, buffer B: 0.1% FA in ACN). The column was then washed with 99% buffer B for 10 minutes and equilibrated with 1% B for 15 minutes. The mass spectrometer was set to collect in PRM mode using an inclusion peptide list (**Supplementary Table 5**). An additional full survey scan was collected to assess for possible interference. Full scans were collected at a resolution of 120,000 at 200 m/z with an AGC setting of 2×10^5^ ion and a maximum ion transfer (IT) time of 50 ms. For PRM scans, the settings were: resolution at 30,000 at 200 m/z, AGC target of 1×10^5^ ions, maximum IT time of 50 ms, microscans count of 1, isolation width of 1.6 m/z, and isolation offset of 0 m/z. A pre-optimized normalized collision energy of 32% was used to obtain the maximal recovery of target product ions. The top 5-10 product ions from this collision energy optimization were used for downstream peptide quantification.

### Peptide Quantification

A spectral library was built using Skyline^133^ (version 4.2) based on tandem mass spectra gathered from previous data dependent acquisition methods. A Skyline template was then created to quantify the endogenous peptides. The template parameters were: centroided precursor mass analyzer, MS1 mass accuracy of 20 ppm; centroided product mass analyzer, MS/MS mass accuracy of 20 ppm; include all matching scans. All rawfiles were then imported and processed accordingly. The resulting extracted ion chromatograms (XICs) of selected fragments were manually inspected and peak picking adjustments were made accordingly. The sum of all product ion peak areas was calculated in Skyline and extracted for further statistical analyses. The peak areas were normalized using the peak areas of external reference peptides. Raw peptide intensities are provided in **Supplementary Table 5**.

### Selected Reaction Monitoring (SRM) of ROS/MAP Brain Tissues

Samples were prepared for LC-SRM analysis using a standard protocol described elsewhere^134,135^. Briefly, on average ∼20 mg of DLPFC brain tissue from each subject was homogenized in denaturation buffer. After denaturation with DTT, 400 µg protein aliquots were taken for further alkylation with iodoacetamide followed by digestion with trypsin as described. The digests were cleaned using C_18_ solid phase extraction, and 30 µL aliquots at 1 µg/µL concentrations were mixed with 30 µL synthetic peptide mix. LC-SRM experiments were performed on a nanoACQUITY UPLC (Waters) coupled to a TSQ Vantage mass spectrometer (ThermoScientific), with 2 µL of peptide injection for each brain sample. Buffer A was 0.1% FA in water and buffer B was 0.1% FA in 90% ACN. Peptide separations were performed on an Acquity UPLC BEH 1.7 µm C_18_ column (75 µm i.d. × 25 cm) at a flow rate 350 nL/min using a gradient of 0.5% buffer B over 0 to 14.5 min, then 0.5% to 15% over 14.5 to 15.0 min, then 15% to 40% over 15 to 30 min, and then 45% to 90% B over 30 to 32 min. The heated capillary temperature and spray voltage was set at 350 °C and 2.4 kV, respectively. Both the Q1 and Q3 were set as 0.7 FWHM. A scan width of 0.002 m/z and a dwell time of 10 ms were used. All SRM data were analyzed using the Skyline software package^133^. All data were manually inspected to ensure correct peak assignment and peak boundaries. The peak area ratios of endogenous light peptides and their heavy isotope-labeled internal standards (i.e., L/H peak area ratios) were then automatically calculated by the Skyline software, and the best transition without matrix interference was used for accurate quantification. Following homogenization of all tissues, small aliquots of protein from each of the samples was pooled, which were then digested and served as a global external pooled reference standard. Peptides generated from this pooled standard were scattered throughout the study (8 samples per 96-well plate) and were used to capture the technical variance that is due to sample preparation steps (except homogenization) and instrument measurements. The signal-to-noise ratio in quantification of each peptide was calculated as the ratio of variances across the human subject samples *versus* the technical controls. Peptides with a signal-to-noise ratio less than 2 were excluded from further analysis. The peptide relative abundances were log_2_ transformed and centered at the median. The abundance of endogenous peptides was quantified as a ratio to spiked-in synthetic peptides containing stable heavy isotopes. The “light/heavy” ratios were log_2_ transformed and shifted such that median log_2_-ratio was zero. Normalization adjusted for differences in protein amounts among the samples. During normalization, the log_2_-ratios were shifted for each sample to make sure the median was set at zero.

### CSF Samples

All participants from whom CSF samples were collected provided informed consent under protocols approved by the Institutional Review Board (IRB) at Emory University. All patients received standardized cognitive assessments (including MoCA) in the Emory Cognitive Neurology clinic, the Emory Goizueta Alzheimer’s Disease Research Center (ADRC), and affiliated research studies (Emory Healthy Brain Study [EHBS] and Emory M^2^OVE-AD study). All diagnostic data were supplied by the ADRC and the Emory Cognitive Neurology Program. CSF was collected by lumbar puncture and banked according to 2014 ADC/NIA best practices guidelines. For patients recruited from the Emory Cognitive Neurology Clinic, CSF samples were sent to Athena Diagnostics and assayed for Aβ42, total-Tau, and phospho-Tau (CSF ADmark®) using the INNOTEST® assay platform. CSF samples collected from research participants in the ADRC, EHBS, and M^2^OVE-AD were assayed using the INNO-BIA AlzBio3 Luminex assay. In total, there were two cohorts of CSF samples that were used in the proteomics studies. Cohort 1 contained CSF samples from 150 healthy controls and 150 MCI/AD patients. Cohort 2 included CSF obtained from three groups: 32 cognitively normal, 31 AsymAD, and 33 MCI/AD. Cases and normal individuals with AsymAD were defined using established biomarker cutoff criteria for AD for each assay platform^136,137^. Cohort information is provided in **Supplementary Table 2**.

### CSF Protein Digestion

To generate peptides, all crude CSF samples were digested with LysC and trypsin. Briefly, 20 µL CSF from each sample was reduced and alkylated with 0.4 µL 0.5 M tris-2(-carboxyethyl)-phosphine (TCEP) and 2 µL 0.4 M chloroacetamide (CAA) with heating at 90°C for 10 min, followed by a 15 min water bath sonication. The samples were then further denatured by the addition of 67.2 µL of 8 M urea buffer (8 M urea, 100 mM NaHPO_4_, pH 8.5) and digested overnight with 1.9 µg LysC (Wako) (1:10 enzyme to protein ratio according to the highest amount of sample). Following LysC digestion, the samples were diluted to 1 M urea using 50 mM ammonium bicarbonate. The same amount of trypsin (Promega) was then added (1:10 enzyme to protein ratio) and digestion was carried out for another 12 h. After trypsin digestion, the peptide solutions were acidified with a 1% TFA and 10% FA solution to a final concentration of 0.1% TFA and 1% FA. Peptides were desalted with a 30 mg C18 HLB column (Waters) and eluted in 1 mL of 50% ACN. Aliquots (120 µL) from cohort 1 (*n*=297) or cohort 2 (*n*=96) samples were pooled together and split into equal volume aliquots (880 µL) for use as the global internal standard (GIS) for TMT labeling. All samples and GIS were dried using a SpeedVac.

### TMT Boost Channel

Signals of low abundant proteins in the TMT 11-plex were amplified using a boost channel, as previously described^138,139^. A pooled CSF sample was created separately for each cohort by combining 50 µL from each sample in cohort 1 or cohort 2 into a pool for each cohort. Abundant proteins were removed using the High Select Top14 Abundant Proteins Depletion Resin (Thermo Scientific A36372BR) according to the manufacturer’s protocol, using a CSF-to-resin volume ratio of 1:1 and an incubation time of 15 min. After immunodepletion, protein concentrations were determined by BCA. Proteins were then reduced and alkylated (10 mM TCEP, 40 mM CAA) for 10 minutes at 90 °C. The samples were then subjected to bath sonication for 15 min and dried under vacuum in a SpeedVac. The immunodepleted pooled samples were re-suspended in 6 M urea buffer (6 M urea, 75 mM NaHPO_4_, pH 8.5) at half the volume of the pooled sample prior to evaporation. Samples were digested overnight with LysC at an enzyme to protein ratio of 1:10. The following day, samples were diluted with 50 mM ammonium bicarbonate to reduce the urea concentration to 1M, and trypsin (Promega) was added (1:10 enzyme to protein ratio). Digestion was allowed to proceed for 12 hr. Peptides were then desalted using a 200 mg C18 Sep-Pak column, and the eluate was dried using a SpeedVac. Aliquots (600 µg) of the immunodepleted pooled CSF samples were separately dissolved in 100 mM TEAB buffer (625 μL) and labeled with 5 mg of TMT 126 channel reagent (cohort 1 lot# TF266326, cohort 2 lot# SG253447, ThermoFisher Scientific) in anhydrous ACN (256 μL). The reactions were allowed to proceed for 1 hr, and were subsequently quenched by adding 5% hydroxylamine (50 μL) and incubating for 15 min. The 126 channel was then added to the other channels, as described below.

### TMT Labeling of Individual and GIS CSF Samples

All samples, including the GIS, were labeled with the 10-plex TMT kit plus an additional channel, for a total of 11 TMT channels (cohort 1 lot# TG273545 for 10-plex, TG273555 for channel 131C; cohort 2 lot# SI258088 for 10-plex, SJ258847 for channel 131C, ThermoFisher Scientific). Samples were grouped into batches as shown in **Supplementary Table 2**. The TMT labeling kit was equilibrated to room temperature and dissolved in anhydrous ACN (256 μL). The samples were reconstituted in 100 mM TEAB buffer (50 μL) and mixed with 0.4 mg (20.5 μL) of the corresponding labeling reagent. The labeling reactions were allowed to proceed for 1 hr, and were subsequently quenched with 5% hydroxylamine (4 μL). Per each TMT batch, labeled peptides from 9 channels (127N, 128N, 128C, 129N, 129C, 130N, 130C, 131, 131C) were mixed, desalted using a 100 mg C18 Sep-Pak column, and dried using a SpeedVac. The immunodepleted pooled sample labeled with the 126 channel (boost channel) was then added to each 9-channel TMT mixture at a ratio of 50:1 pooled to individual CSF sample by original volume:volume prior to evaporation. The sample mixtures were desalted using a 200 mg C18 Sep-Pak column, and dried using a SpeedVac.

### Mass Spectrometry Analysis of CSF

All samples were resuspended in equal volume of loading buffer (0.1% FA, 0.03% TFA, 1% ACN). Peptide eluents were separated on a self-packed C18 (1.9 µm, Dr. Maisch, Germany) fused silica column (25 cm × 75 μM internal diameter (ID): New Objective, Woburn, MA) by an Easy-nLC system (ThermoFisher Scientific) and monitored on an Orbitrap Fusion Lumos mass spectrometer (ThermoFisher Scientific) interfaced with a high-field asymmetric waveform ion mobility spectrometry (FAIMS) Pro. Sample elution was performed over a 180 min gradient (buffer A: 0.1% FA in water, buffer B: 0.1% FA in 80% ACN) with flow rate at 225 nL/min. The gradient was from 1% to 8% buffer B over 3 min, then from 8% to 40% over 160 min, then from 40% to 99% over 10 min, and then held at 99% B for 10 min. The mass spectrometer was set to acquire data in positive ion mode using data dependent acquisition and three (−50, −65 and −85 V) different compensation voltages (CV)^140^. Data were acquired at each CV for 1 s during each cycle. Each cycle consisted of 1 full scan followed by as many MS2 and MS3 scans as possible within a 1 s timeframe. The full scan was performed with an m/z range of 450-1500 at 120,000 resolution (at 200 m/z) with an AGC setting of 4×10^5^ and maximum injection time 50 ms. The collision induced dissociation (CID) MS/MS scans were collected in the ion trap with an isolation window of 0.7 m/z, a collision energy of 35%, AGC setting of 1×10^4^, and a maximum injection time of 50 ms. The top 10 product ions were subjected to HCD synchronous precursor selection-based MS3 (SPS-MS3) as previously described^27^. For SPS-MS3 scans the isolation window was set to 2 m/z, the resolution to 50,000, the AGC to 1×10^5^, and the maximum injection time to 105 ms. For both cohorts, a single preliminary run of TMT batch 1 using the above parameters was used to create a target inclusion list of peptides that specifically excluded those from the top 15 most abundant proteins. This inclusion list was used for all TMT batches in cohort 1 (*n*=38) and in cohort 2 (*n*=12).

### Database Searches and Protein Quantification of CSF

All RAW files were analyzed using the Proteome Discoverer Suite (version 2.3, ThermoFisher Scientific). MS/MS spectra were searched against the UniProtKB human proteome database (downloaded April 2015 with 90,411 total sequences). The Sequest HT search engine was used to search the RAW files, with search parameters specified as follows: fully tryptic specificity, maximum of two missed cleavages, minimum peptide length of 6, fixed modifications for TMT tags on lysine residues and peptide N-termini (+229.162932 Da) and carbamidomethylation of cysteine residues (+57.02146 Da), variable modifications for oxidation of methionine residues (+15.99492 Da), serine, threonine and tyrosine phosphorylation (+79.966 Da) and deamidation of asparagine and glutamine (+0.984 Da), precursor mass tolerance of 20 ppm, and a fragment mass tolerance of 0.6 Da. Percolator was used to filter PSMs and peptides to an FDR of less than 1%. Following spectral assignment, peptides were assembled into proteins and were further filtered based on the combined probabilities of their constituent peptides to a final FDR of 1%. In cases of redundancy, shared peptides were assigned to the protein sequence in adherence with the principles of parsimony. Reporter ions were quantified from MS3 scans using an integration tolerance of 20 ppm with the most confident centroid setting, as previously described^27^.

### Controlling for Batch-specific Variance

We implemented a median polish algorithm for removing technical variance (e.g., due to tissue collection, cohort, or batch effects) from a two-way abundance-sample data table as originally described by Tukey^141^. The algorithm is fully documented and available as an R function, which can be downloaded from https://github.com/edammer/TAMPOR. The algorithm implements iterations of the below equation, where batch and cohort are interchangeable.

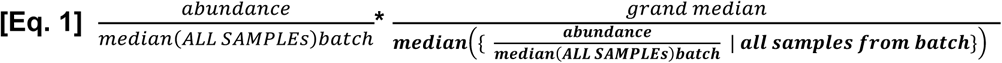

Briefly, Equation 1 is applied to each protein measurement (LFQ or TMT reporter abundance) across all samples individually where the first term represents batch-wise median-centered abundance, and the second term is a batch-specific normalization factor comprised of the grand median of all batch-specific medians, divided by the appropriate batch-specific median of median-centered abundances. The data matrix is then log_2_-transformed, and each log_2_(ratio) is adjusted by subtraction of sample (column)-wise median log_2_(ratio) for all proteins. Then, ratios are anti-logged and multiplied by the protein (row)-wise median of all samples used for the Eq. 1, term 1, denominator, extracted before Eq. 1 was executed. This process is iterated until convergence. The use of median polish ensures that the reduction of variance is robust to outliers while the overall algorithm preserves biological variance, given that batches have been randomized to avoid confounding batch with diagnosis or other biological traits. Prior to matrix assembly for the consensus analysis, intra-cohort batch effects were first removed in the MSSB (batch correction with 166 case samples across 7 batches) and Banner (batch correction with 178 case samples across 4 batches) cohorts. All remaining batch corrections restricted the first term denominator to global pooled (within cohort) standard sample abundances, and the second term used all individual case samples. Following removal of intra-cohort batch effects in MSSB and Banner, all samples were processed jointly with the algorithm in the same sample-protein matrix to capture biological variance across all samples in all four cohorts (ACT, Banner, BLSA, and MSSB) for the consensus analysis. The above algorithm was applied to a matrix in which proteins that had ≥ 50% missing values were removed. For the consensus LFQ network, 450 case samples (3 ACT outliers were removed prior to inclusion, as described below) classified as control, AsymAD, or AD by our unified criteria (see case classification methods above) were considered as “all samples” for denominators in Eq. 1. All remaining batch corrections listed as follows restricted the first term denominator to global pooled (within cohort) standard sample abundances, and the second term used all individual case samples.

For ROSMAP 50-batch TMT protein abundances, there were two pooled global internal standard channels in each TMT batch (*n*=64), and 400 individual case samples (non-internal standard samples). For the Hopkins aging cohort (84 case samples), global pool mixture samples (3 each per 3 batches) were used for the first term denominator, with the second term using all non-global pool mixture samples. For the UPenn PRM analysis (3 batches, 114 case samples, and 9 pooled controls), data were likewise batch corrected using 3 global pool mixture samples per batch for the first term denominator and all within-batch non-pooled samples for the second term. UPenn LFQ data (10 batches, 330 case samples, and 29 control pools) were similarly batch corrected as described. CSF 96-case and 300-case TMT normalized abundances were also batch corrected using the above algorithm, with equation 1 first term denominator restricted to global pooled (within cohort) standard sample abundances, while the second term used all individual non-internal standard case samples.

### Regression for Covariates and Outlier Removal

No imputation of missing values was performed in any cohort. Nonparametric bootstrap regression was performed separately in each cohort by subtracting the trait of interest (age at death, sex, or postmortem interval (PMI)) times the median estimated coefficient from 1000 iterations of fitting for each protein in the cohort-specific log_2_(abundance) matrix. Ages at death used for regression were uncensored. Case status/diagnosis was also explicitly modeled (i.e., protected) in each regression. Following regression of each individual cohort, we assessed whether any cohort-specific tissue dissection bias was present by performing a Spearman rank correlation of traits including age, sex, PMI, and white matter markers to the top five principle components (PC) of log_2_(abundance). Network outlier case samples were not considered in the PCs, and were identified prior to PC analysis using Oldham’s ‘SampleNetworks’ v1.06 R script^12^ as previously published^142^ using a 3 fold-SD cutoff of Z-transformed sample connectivity. The Spearman rank correlation was performed prior to correction of cohort-specific batch effects as described above, and after intra-cohort batch correction of the MSSB and Banner cohorts. All four of the cohorts were confirmed to have no significant PC correlation to age, sex, or PMI; however, ACT was observed to have a first PC significantly correlated (average rho=0.94) to protein abundance of white matter markers identified previously as oligodendrocyte coexpression network hubs^23^. These markers were BCAS1, SIRT2, MBP, and MAG. This white matter PC represented 27 percent of variance in the ACT cohort, whereas the white matter marker-correlated PC represented 7 to 12 percent variance in the other three cohorts. To adjust for this white matter variance in ACT, we applied a second round of bootstrap regression to the 62 non-outlier ACT case sample log_2_(abundances), using the white matter PC as a regression covariate, and subtracted 28 percent of the white matter marker correlated variance to achieve a final variance of 12 percent after recalculation of the top 5 PCs. Abundance data for the 450 case samples were then assembled into a matrix of 3334 proteins, and cross-cohort batch correction by median polish was performed as described above. Finally, network outlier detection was performed as described above, which removed 31/450 cases from consideration in the four-cohort consensus network and differential abundance analyses. All outliers are listed in **Supplementary Table 3**. In all other cohorts that were not combined for the consensus network analysis, batch correction was performed first, followed by outlier removal, followed by removal of proteins with ≥ 50% missing values, and then regression of age, gender, and PMI prior to coexpression network and differential abundance analyses. In the Hopkins aging cohort, age was not considered as a trait for regression. In the CSF cohorts, only age at time of collection and sex were considered for regression.

### Differential Expression Analysis

Differentially expressed proteins were found using one-way ANOVA followed by Tukey’s comparison post-hoc test across control, AsymAD and AD cases. Significantly altered proteins with corresponding *p* value are provided in **Supplementary Tables 3 and 6** for consensus AD network proteins and astrocyte/microglial phenotype proteins, respectively. Differential expression is presented as volcano plots, which were generated with the ggplot2 package in R v3.5.2.

### Protein Correlations to Aging

Protein expression levels for all 2,756 proteins measured in the aging cohort with fewer than 50% missing values were correlated to age at death using the bicor function, after regression for sex and PMI. In addition to bicor rho, the Student’s *p* value for significance of the correlation, FDR (*q* value), and signed Z-score for the correlation were calculated. The CSGene database of cellular senescence genes^143^ was cross-referenced with the protein data to annote proteins involved in cellular senescence. This information is provided in **Supplementary Table 4**, along with differences between AD, control, or AsymAD for the same proteins in the consensus network, and their corresponding consensus network module relationships. Individual correlations for proteins in the consensus network to all traits provided in **Extended Data Figure 1** are provided online through the Synapse web portal.

### Weighted Correlation Network Analysis (WGCNA)

We used the WeiGhted Correlation Network Analysis (WGCNA) algorithm for our network analysis pipeline, as previously described^81^. A weighted protein co-expression network for the four-cohort consensus data was generated using the 3,334 log_2_ protein abundance x 419 case-sample matrix that had undergone covariate and batch correction as described above. The WGCNA::blockwiseModules() function was used with the following settings: soft threshold power beta=5.5, deepSplit=4, minimum module size of 14, merge cut height of 0.07, mean TOM denominator, a signed network with partitioning about medioids (PAM) respecting the dendrogram, a minimum kME to remain in a module of 0.30, and a reassignment threshold of p<0.05. Specifically, we calculated pair-wise biweight mid-correlations (bicor, a robust correlation metric) between each protein pair, and transformed this correlation matrix into a signed adjacency matrix^144^. The connection strength of components within this matrix was used to calculate a topological overlap matrix, which represents measurements of protein expression pattern similarity across cohort samples constructed on the pairwise correlations for all proteins within the network^145^. Hierarchical protein correlation clustering analysis by this approach was conducted using 1-TOM, and initial module identifications were established using dynamic tree cutting as implemented in the WGCNA::blockwiseModules() function^146^. Module eigenproteins were defined, which represent the most representative abundance value for a module and which explain covariance of all proteins within a module^26^. Pearson correlations between each protein and each module eigenprotein were performed; this module membership measure is defined as k_ME_. After the initial network construction, 18 modules consisting of 14 or more proteins were detected. Given high k_ME_ similarity between some modules, we used the WGCNA::moduleMergeUsingKME() function to reduce the number of modules to 13, with the following parameters: percentage of module members checked for k_ME_ overlap of 50 percent (threshPercent=50), threshold for merging modules with high common k_ME.intramodule_ of 25 percent (mergePercent=25), and all other parameters default for the function. After module merging, MEs and signed kMEs were recalculated with the WGCNA::moduleEigengenes() and WGCNA::signedKME() functions, respectively. Finally, we ‘cleaned’ the network of assignments with aberrant k_ME_ (since WGCNA clustering into modules and merging of those modules use hybrid approaches not solely dependent on kME) by applying the following algorithm: remove module members with k_ME.intramodule_ <0.28, then check all grey (unclustered) proteins for any k_ME_ >0.35 and assign to the module with the highest k_ME.intramodule_. For BLSA precuneus, Mayo temporal cortex, and ROS/MAP cohorts, we confirmed that they built networks using the pipeline described above, including batch correction, outlier removal, and age/sex/PMI regression. Network building was employed without module merging. The blockwiseModules function for ROS/MAP used power=10, minimum module size=30 (11,225 proteins in 323 non-excluded and non-outlier case samples); for BLSA precuneus used power=8, minimum module size=14 (3,348 proteins in 46 case samples); and for Mayo used power=5, minimum module size=14 (3,951 proteins in 107 case samples). For analysis of sex effects on the consensus network, the identical cases and analysis pipeline as described above were used to build the consensus network, except that covariance due to sex was not removed by regression prior to WGNCA analysis.

### Network Preservation

We used the WGCNA::modulePreservation() function to assess network module preservation across cohorts. We also used this function to assess the effect of missing values on the consensus network. Z-summary composite preservation scores were obtained using the consensus network as the template versus each other cohort or missing value threshold tested, with 500 permutations. Random seed was set to 1 for reproducibility, and the quickCor option was set to 0. We also assessed network module preservation using synthetic eigenproteins. Briefly, protein module members in the consensus network template with a k_ME.intramodule_ among the top 20th percentile were assembled into a synthetic module in each target cohort, and synthetic modules with at least 4 members were used to calculate synthetic weighted eigengenes representing the variance of all members in the target network across case samples via the WGCNA::moduleEigengenes() function. Statistics and correlation scatterplots involving target cohort traits were then calculated and visualized.

### Gene Ontology and Cell Type Marker Enrichment Analyses

To characterize differentially expressed proteins and co-expressed proteins based on gene ontology annotation, we used GO Elite v1.2.5 as previously published^81^, with pruned output visualized using an in-house R script. Cell type enrichment was also investigated as previously published^81^. Astrocyte subtype markers were obtained from Zamanian *et al*.^46^. Microglia subtype markers were obtained from Rangaraju *et al*.^47^.

### GWAS Module Association

To determine if any protein products of GWAS targets were enriched in a particular module, we used the single nucleotide polymorphism (SNP) summary statistics from the International Genomics of Alzheimer’s Project (http://www.pasteur-lille.fr/en/recherche/u744/igap/igap_download.php) ^5^ to calculate the gene level association value using MAGMA^43^, as previously described^23^. *APOE* was added to the gene list and assigned a - log *p* value of 50, given its known strong association with AD^147^. ApoE did not fall within a network module, and therefore did not influence the reported module enrichment results. Similar analyses were performed with GWAS candidates for Schizophrenia (SCZ) and Autism Spectrum Disorders (ASD)^148^. These GWAS datasets were provided and downloaded from the Psychiatric Genomics Consortium (http://www.med.unc.edu/pgc/downloads).

### Curation of AD Mouse Model Purified Glial Transcriptomic and Proteomic Data

Orre *et al.*^51^ microarray data files (n=11; files last updated on July 20, 2015) were downloaded from the NCBI gene expression omnibus (GEO) datasets website (GEO identifier: GSE74615). The data, consisting of 22 samples, was normalized and centered based on limma R package user guidelines with backgroundCorrect method=”normexp” and normalizeBetweenArrays method=”quantile”. Transcripts with signal at least 110% of the 95^th^ percentile of all normalized red negative control spots on the arrays in at least 4/22 samples were considered as expressed above noise in at least one cell type (n=28,157 gene transcripts across N=14 purified microglial measurements and N=8 purified astrocyte measurements). WGCNA collapseRows function was used to handle multiple probe mappings to any gene, with probe selection method=”maxRowVariance”. Grubman *et al*.^52^ peptide-level total peak area data for XO4^+^ (amyloid-β plaque phagocytosing) and XO4^−^ (amyloid-β plaque non-phagocytosing) acutely purified microglia from AD and WT mice, respectively, were summed and log_2_-transformed to achieve protein-level data for 94 gene product proteins. Grubman *et al*. mRNA-level measurements were obtained directly and used without further processing.

### Other Statistics

All statistical analyses were performed in R (v. 3.5.2). Boxplots represent the median, 25^th^, and 75^th^ percentiles, and whiskers with staples represent measurements to the 5^th^ and 95^th^ percentiles of non-outlier samples; outlier samples are plotted beyond these whiskers as open circles. Correlations were performed using the biweight midcorrelation function as implemented in the WGCNA R package. Comparisons between two groups were performed by *t* test. Comparisons among three or more groups were performed with Kruskal-Wallis nonparametric ANOVA or standard ANOVA with Tukey or Dunnett *post hoc* pairwise comparison of significance. *P* values were adjusted for multiple comparisons by false discovery rate (FDR) correction where indicated. T-distributed Stochastic Neighbor Embedding (t-SNE) analysis was implemented with R package Rtsne, as previously published^47^. Module membership graphs were generated using the igraph R package as previously described^23,149^.

---

**Supplementary Figure 1.**
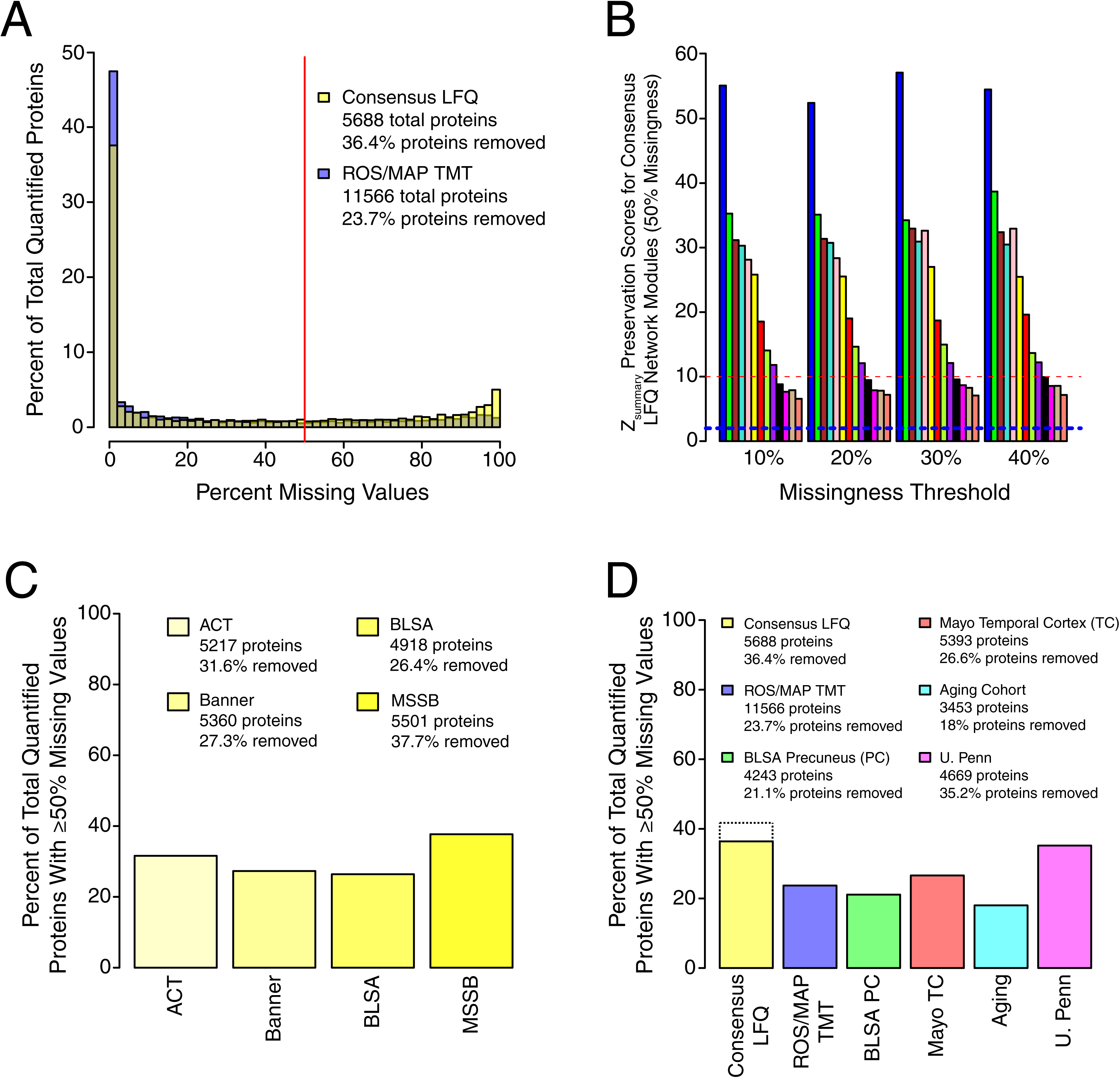
Analysis of Missing Protein Quantitative Measurements and their Effect on the AD Network. (A-D) The percentage of quantified proteins with a given level of missing quantitative measurements was analyzed for both the consensus LFQ and ROS/MAP TMT networks (A). Each bar represents a bin of 2%. The red line indicates the 50% missing measurement threshold used in this study. The total number of quantified proteins for each dataset, and the percentage of quantified proteins removed due to ≥50% missing measurements prior to construction of the respective protein networks, is provided in the legend. (B) The effect of missing value threshold on AD network modules. The AD network was constructed using different allowed levels of missing protein measurements. Preservation of AD network modules (50% missingness threshold) in each network generated using a more stringent threshold (10-40% missingness) was assessed by Z_summary_ score. The dashed blue line indicates a *z*_summary_ score of 1.96, or FDR *q* value <0.05, above which module preservation was considered statistically significant. The dashed red line indicates a *z*_summary_ score of 10, or FDR *q* value ∼ 1e^-23^, above which module preservation was considered highly statistically significant. Each module is color coded as shown in Figure 1. (C) Percentage of total quantified proteins with ≥50% missing measurements in each cohort used for the AD consensus network. The total number of quantified proteins, and the percentage removed by applying the ≥50% missingness threshold, is provided in the legend for each cohort. (D) Percentage of total quantified proteins with ≥50% missing measurements in each cohort used in this study. The total number of quantified proteins, and the percentage removed by applying the ≥50% missingness threshold, is provided in the legend for each cohort. For the consensus LFQ cohort, the dotted line indicates the percent removed (41.3%) when missingness is controlled separately in each cohort prior to combination for construction of the AD network, as was done in this study. The solid bar is provided for direct method comparison to other cohorts used in the study. LFQ, label-free quantitation; TMT, tandem-mass tag; BLSA, Baltimore Longitudinal Study of Aging, Banner, Banner Sun Health Research Institute; MSSB, Mount Sinai School of Medicine Brain Bank; ACT, Adult Changes in Thought Study; ROS/MAP, Religious Orders Study and Memory and Aging Project; PC, precuneus; TC, temporal cortex.

**Supplementary Figure 2.**
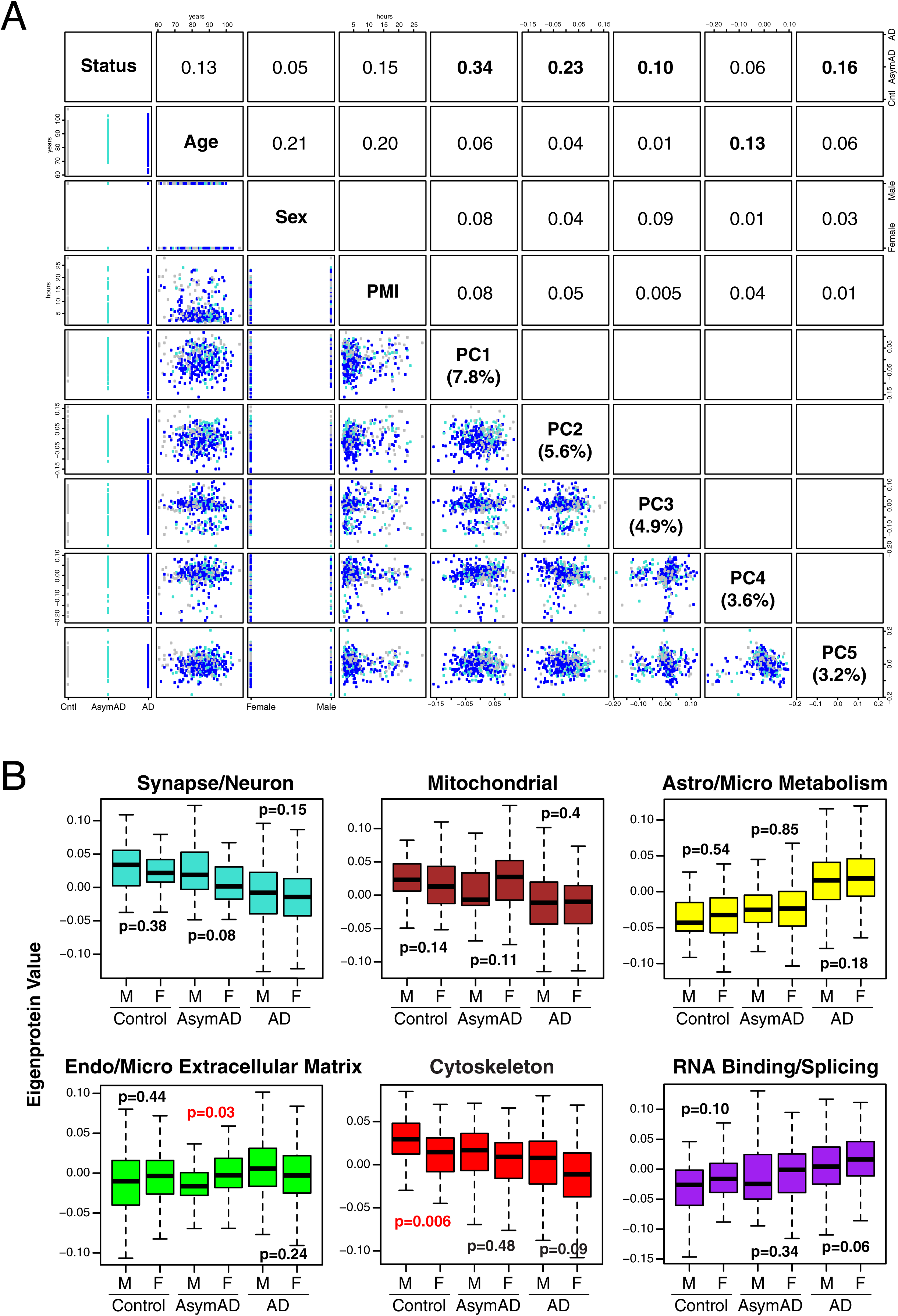
Covariate Effects on AD Network Protein Quantitative Values and Modules. (A, B) Principal component analysis was performed on AD network protein quantitative values after batch correction but prior to regression for age, sex, and post-mortem interval (PMI) covariates (A). Correlation values between case status (control, AsymAD, or AD), age, sex, PMI, and the first five principal components of the data are shown. The covariate most strongly correlated to each principal component is highlighted in bold. The percentage of variance in the data explained by each principal component is given in parentheses. (B) Effects of sex on AD network modules shown in Figure 1C. The AD network was built without regression for sex, and module eigenprotein levels were compared between male and female sex for each case group. Statistically significant differences are highlighted in red. Correlations were performed using Spearman’s rank correlation. Differences in protein levels were assessed by Kruskal-Wallis one-way ANOVA. Boxplots represent the median, 25^th^, and 75^th^ percentiles, and whiskers represent measurements to the 5^th^ and 95^th^ percentiles. PC, principal component; PMI, post-mortem interval; Cntl, control; AsymAD, asymptomatic Alzheimer’s disease; AD, Alzheimer’s disease.

**Supplementary Figure 3.**
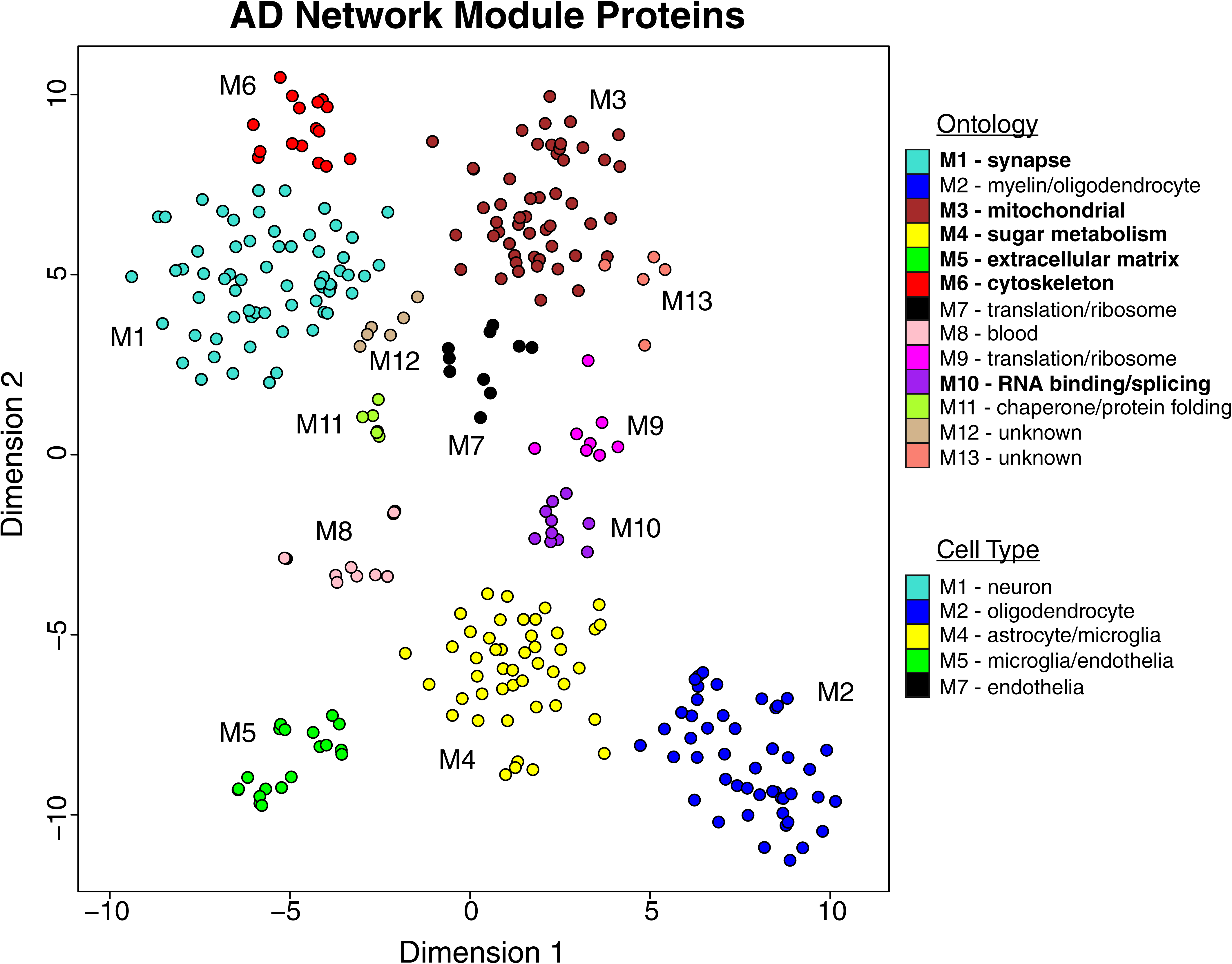
Relationship of AD Network Proteins by t-SNE Analysis. Dimensionality reduction and visualization by t-distributed stochastic neighbor embedding (t-SNE) was applied to proteins that were in the top 25% by kME value within each AD network module. Proteins are color coded as shown in Figure 1B according to the network module in which they reside. Network module ontologies and cell type enrichments are provided as shown in Figure 1B. Ontologies are highlighted based on the most robust AD trait correlations as shown in Figure 1B.

**Supplementary Figure 4.**
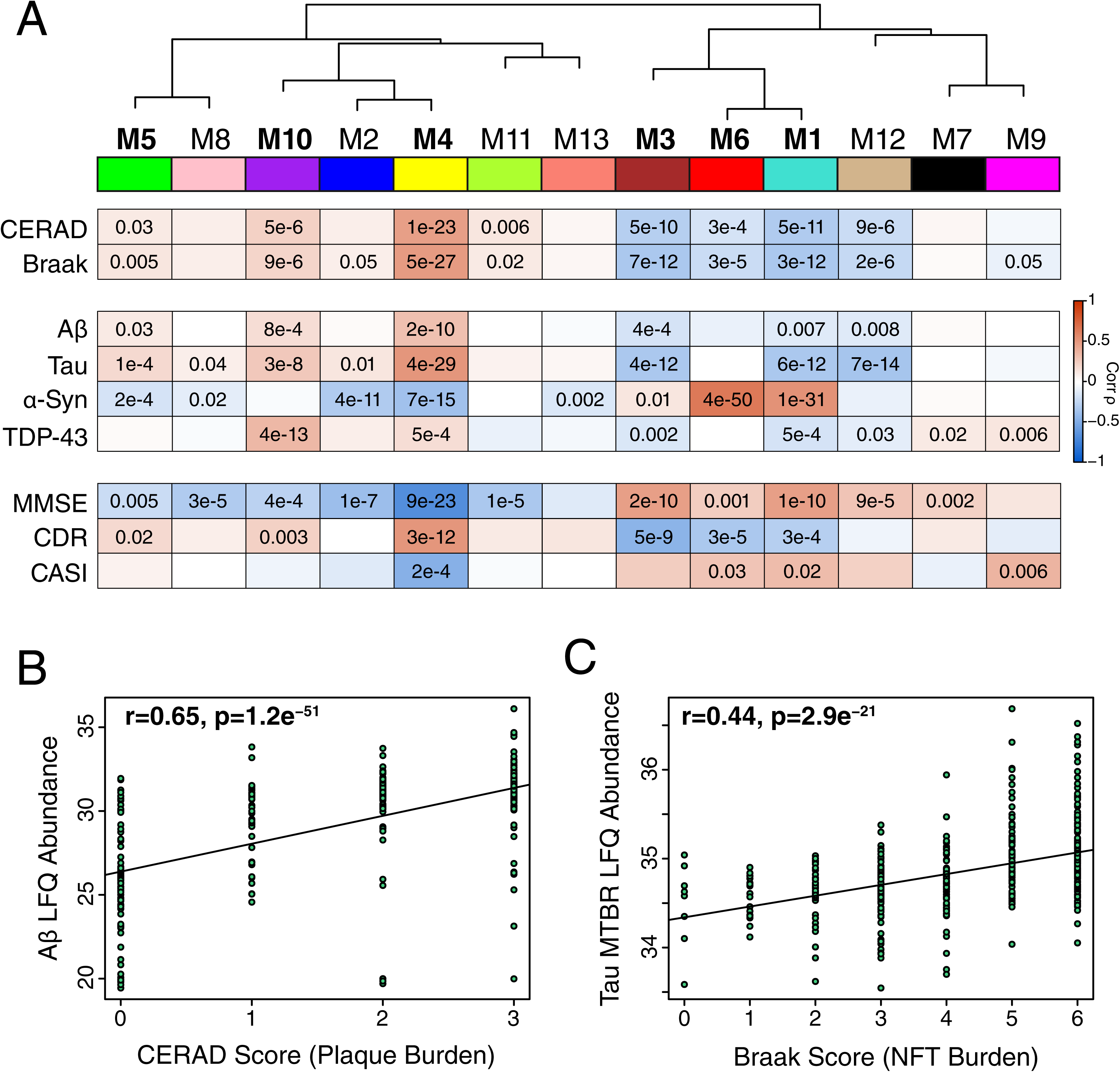
AD Protein Network Module Trait and Pathology Correlations. (A-C) The eigenprotein of each protein network module was correlated with neuropathological, molecular, and cognitive/functional traits (A). Protein modules are bolded as in Figure 1B using CERAD, Braak, MMSE, and CDR correlations. Strength of positive (red) or negative (blue) correlation is shown by two-color heatmap, with *p* values provided for all correlations with *p* < 0.05. Neuropathological, molecular, and cognitive/functional correlations were corrected using the Benjamini-Hochberg method. (B) Correlation between CERAD plaque score and Aβ levels measured by label free quantification (LFQ) mass spectrometry. (C) Correlation between Braak score (NFT, neurofibrillary tangle) and tau levels measured by LFQ of the microtubule binding region (MTBR). CERAD, Consortium to Establish a Registry for Alzheimer’s disease Aβ plaque score (higher scores represent greater plaque burden); Braak, tau neurofibrillary tangle staging score (higher scores represent greater extent of tangle burden); Aβ, amyloid-β; α-Syn, alpha synuclein; TDP-43, TAR DNA-binding protein 43; MMSE, mini-mental status examination score (higher scores represent better cognitive function); CDR, clinical dementia rating score (higher scores representing worse functional status); CASI, Cognitive Abilities Screening Instrument (higher scores represent better cognitive function). MMSE is from Banner, CDR is from MSSB, and CASI is from ACT.

**Supplementary Figure 5.**
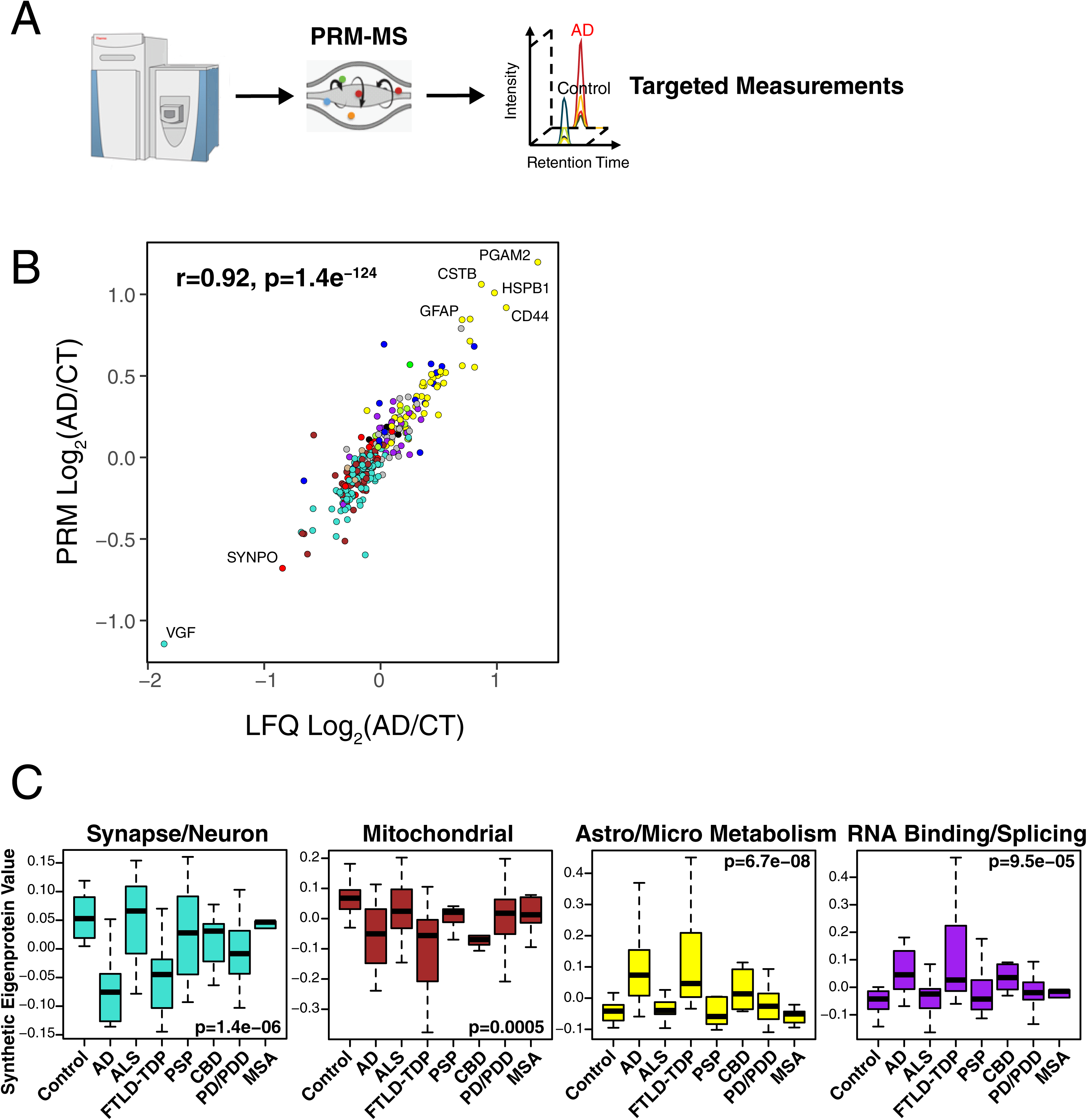
AD Protein Network Module Changes in Other Neurodegenerative Diseases by PRM Analysis. (A-C) Protein levels for 323 proteins across 108 brains from the UPenn cohort were measured by parallel reaction monitoring targeted mass spectrometry (PRM-MS) (A). Targeted peptides and individual protein measurements by disease group are provided in **Supplementary Table 5** and **Extended Data Figure 11**, respectively. (B) Protein levels across all cases were highly correlated between LFQ and PRM measurements. (C) A synthetic eigenprotein was created from proteins that mapped to an AD network module and measured across the different disease groups. Analyses for all modules are provided in **Extended Data Figure 12**. Differences in module synthetic eigenproteins were assessed by Kruskal-Wallis one-way ANOVA. Differences between AD and other case groups were assessed by Dunnett’s test, the results of which are provided in **Supplementary Table 5**. Boxplots represent the median, 25^th^, and 75^th^ percentiles, and whiskers represent measurements to the 5^th^ and 95^th^ percentiles.

**Supplementary Figure 6.**
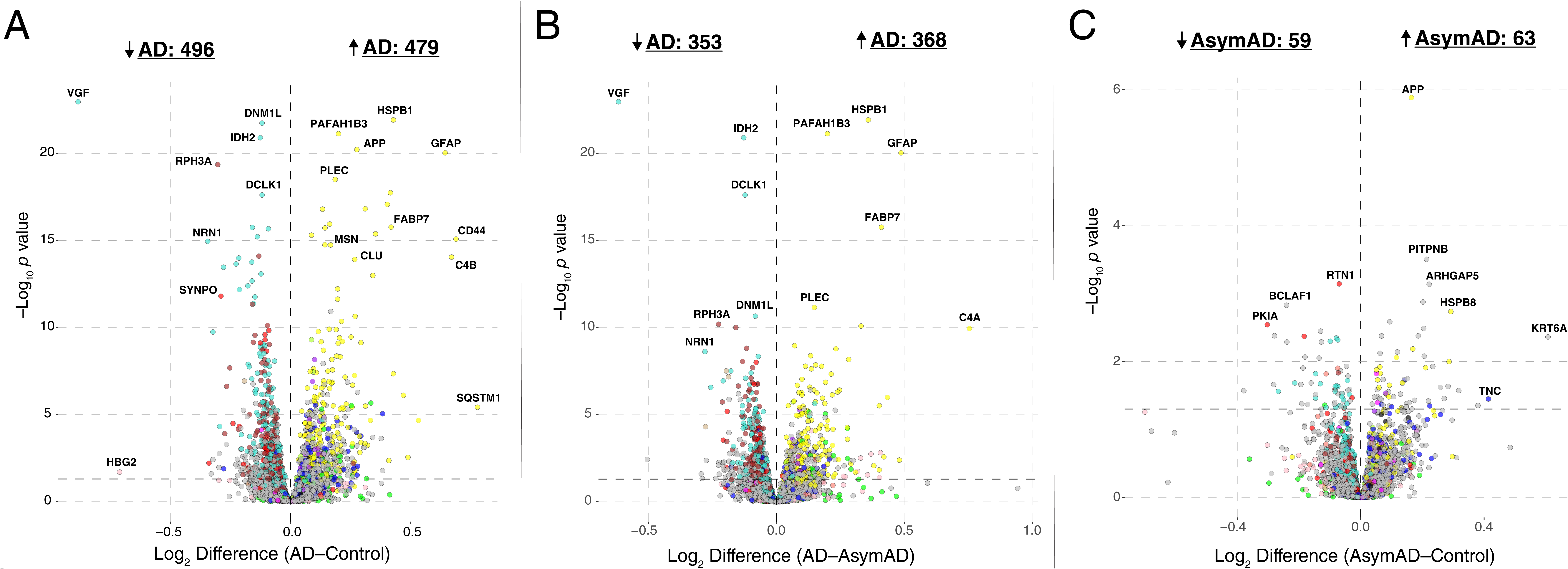
Protein Differential Abundance in AD Brain. (A-C) Differential protein abundance for AD *versus* control (A), AD *versus* AsymAD (B), and AsymAD *versus* control (C) brain, represented by fold-change versus *t* statistic for the given comparison. Proteins are colored by the module in which they reside according to the scheme shown in Figure 1B. The bold horizontal dashed line represents *p* < 0.05. *P* values are corrected by the Benjamini-Hochberg method. AsymAD, asymptomatic Alzheimer’s disease; AD, Alzheimer’s disease.

**Supplementary Figure 7.**
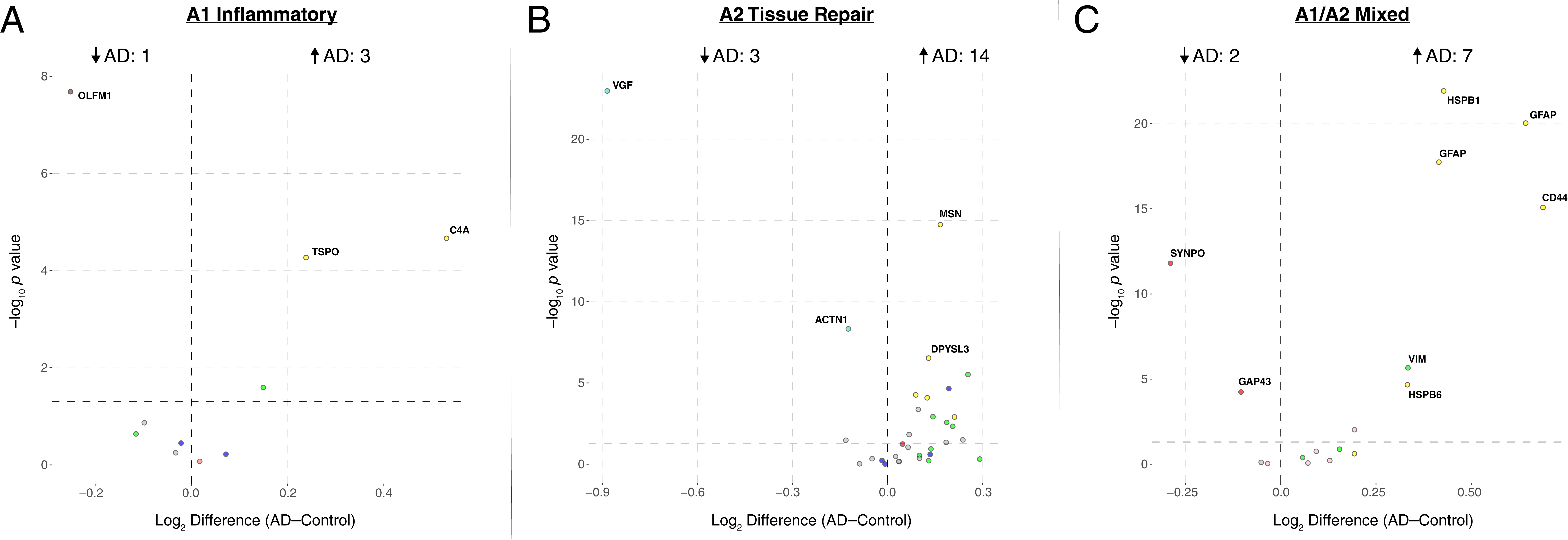
Differential Abundance of Reactive Astrocyte Protein Markers in AD Brain. (A-C) Proteins expressed in different astrocytic response states to acute injury^46^ were analyzed for changes in AD. Astrocyte mRNAs that were upregulated greater than four-fold after acute injury by LPS administration (“A1” Inflammatory) (A), middle cerebral artery occlusion (“A2” Tissue Repair) (B), or both (“A1/A2 Mixed”) (C) were analyzed for changes in abundance between AD and control. Results are shown as protein fold-change versus *t* statistic for the given comparison. The bold horizontal dashed line represents *p* < 0.05. *P* values are corrected by the Benjamini-Hochberg method. Proteins are colored by the module in which they reside according to the scheme shown in Figure 1B. AD, Alzheimer’s disease.

**Supplementary Figure 8.**
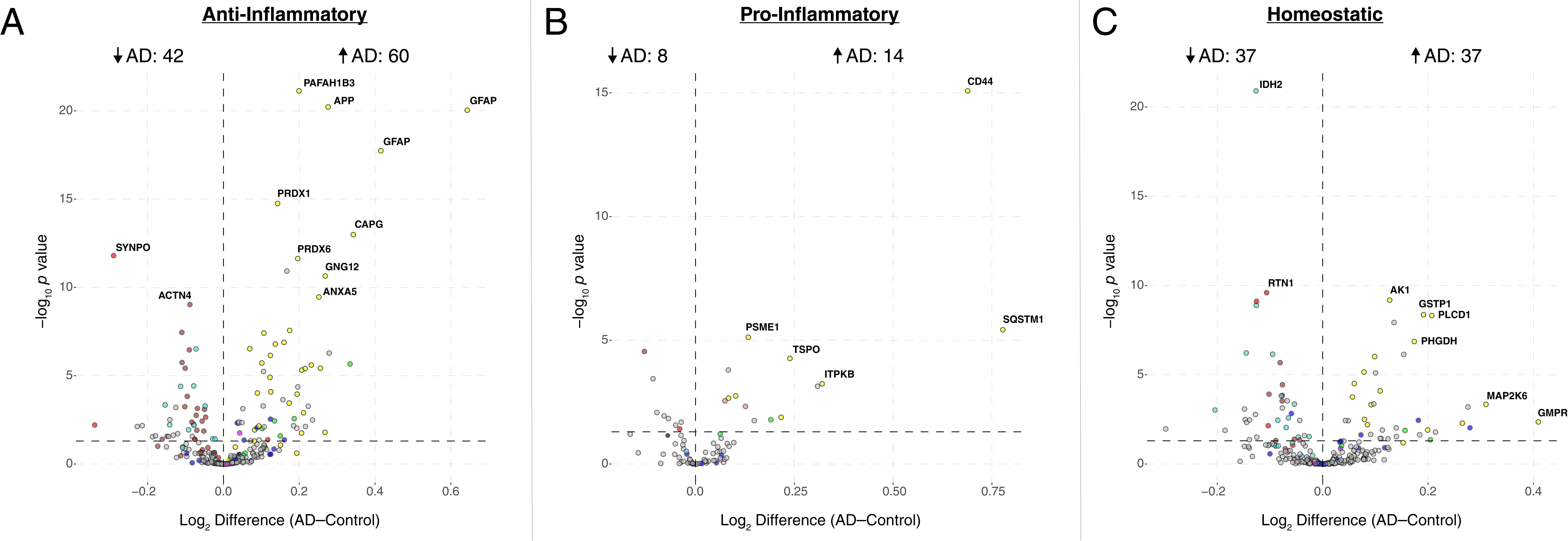
Differential Abundance of Microglial Phenotypic Protein Markers in AD Brain. (A-C) Proteins corresponding to microglial mRNAs that were found to be associated with different microglial phenotypic states^47^ were analyzed for changes in AD. Proteins from microglial co-expression modules corresponding to a disease-associated anti-inflammatory (A), disease-associated pro-inflammatory (B), and homeostatic (C) response phenotype were analyzed for changes in abundance between AD and control. Results are shown as protein fold-change versus *t* statistic for the given comparison. The bold horizontal dashed line represents *p* < 0.05. *P* values are corrected by the Benjamini-Hochberg method. Proteins are colored by the module in which they reside according to the scheme shown in Figure 1B. AD, Alzheimer’s disease.

**Supplementary Figure 9.**
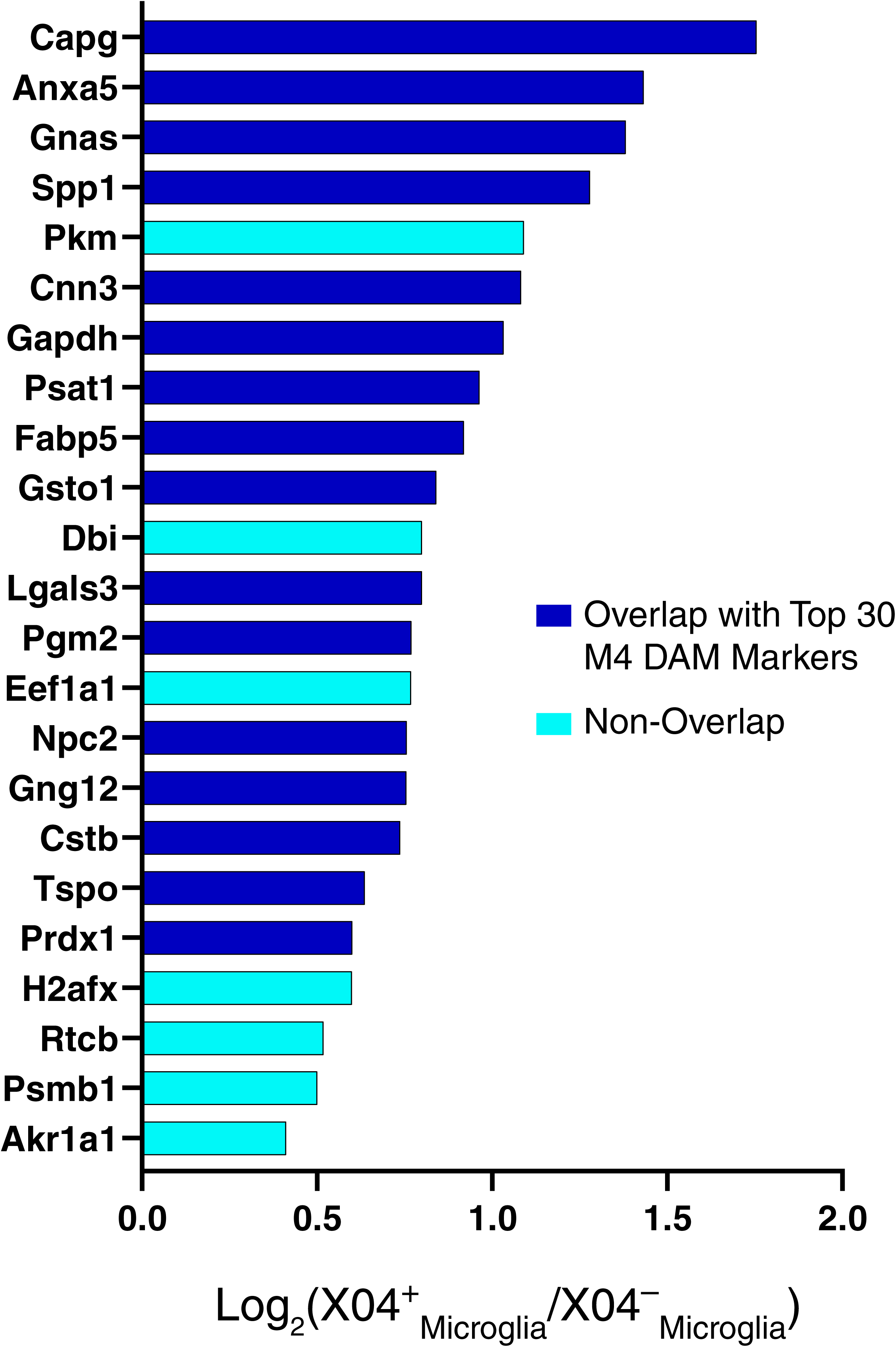
M4 Astrocyte/Microglial Metabolism Module Members Increased at the Transcript Level in Microglia Undergoing Active Amyloid-β Plaque Phagocytosis. mRNA transcripts increased in microglia undergoing active amyloid-β plaque phagocytosis (X04^+^)^52^ were overlapped with cognate proteins in the M4 module. There were 23 transcripts that overlapped with M4 module members. Proteins that also overlapped with the top 30 disease-associated microglia (DAM) markers in the M4 module (Figure 6D) are shown in blue. Proteins that did not overlap with the top 30 DAM markers are shown in cyan. Proteins in cyan are therefore M4 members that may be more specifically elevated in microglia undergoing active amyloid-β plaque phagocytosis.

**Extended Data Figure 1. AD Network Module Phenotype Correlations.** Each module eigenprotein was assessed for group differences by case status and *APOE* genotype, and correlated to genetic, neuropathological, cognitive/functional, and molecular traits. *APOE* genotype information is from the Banner and BLSA cohorts. Control and AsymAD cases were grouped into non-demented (ND) in the *APOE* analysis for sufficient statistical power. Eigenprotein differences by case status or *APOE* genotype were assessed by Kruskal-Wallis (K-W) one-way ANOVA. Statistical significance at *p* <0.05 for case status is highlighted in red. Correlations were performed using both Pearson correlation (cor) and biweight midcorrelation (bicor), which is more robust to outliers. Boxplots represent the median, 25^th^, and 75^th^ percentiles, and whiskers represent measurements to the 5^th^ and 95^th^ percentiles.

**Extended Data Figure 2. AD Network Module Protein Memberships.** The top 100 proteins by module eigenprotein correlation value (kME) in each AD protein network module. The size of each circle indicates the relative kME. Those proteins with the largest kME are considered “hub” proteins within the module, and explain the largest variance in module expression.

**Extended Data Figure 3. GO Analysis on AD Network Modules.** Gene ontology (GO) analysis was performed to gain insight into the biological meaning of each AD protein network module. Enrichment for a given ontology is shown by *z* score.

**Extended Data Figure 4. AD Network Validation in ROS/MAP.** The top 20% of proteins by kME value in each AD brain protein network module was used to create a synthetic eigenprotein, which was then measured by case status in ROS/MAP and correlated with different traits as assessed in the ROS/MAP cohorts. The first boxplot for each module is the AD network eigenprotein by case status, given as reference for the second boxplot, which is the synthetic eigenprotein in the ROS/MAP cohorts. Control and AsymAD cases were grouped into non-demented (ND) in the *APOE* analysis for sufficient statistical power. Synthetic eigenprotein differences by case status or *APOE* genotype in ROS/MAP were assessed by Kruskal-Wallis (K-W) one-way ANOVA. Correlations were performed using both Pearson correlation (cor) and biweight midcorrelation (bicor), which is more robust to outliers. Statistical significance at *p* <0.05 is highlighted in red. Boxplots represent the median, 25^th^, and 75^th^ percentiles, and whiskers represent measurements to the 5^th^ and 95^th^ percentiles.

**Extended Data Figure 5. Round 1 Targeted Measurements of AD Network Proteins in ROS/MAP**. Levels of 67 proteins were measured by selected reaction monitoring (SRM) mass spectrometry across 1016 ROS/MAP cases. One peptide per protein was selected for measurement. Peptide sequences can be found at https://www.synapse.org/#!Synapse:syn10468856. Protein levels were correlated to Aβ levels as measured by SRM, CERAD score, Braak stage, and MMSE. Protein level differences by case status were assessed by Kruskal-Wallis (K-W) one-way ANOVA with Tukey test. Correlations were performed using both Pearson correlation (cor) and biweight midcorrelation (bicor), which is more robust to outliers. Data for each protein is colored by the AD network module in which it resides. Statistical significance at *p* <0.05 is highlighted in red. Boxplots represent the median, 25^th^, and 75^th^ percentiles, and whiskers represent measurements to the 5^th^ and 95^th^ percentiles.

**Extended Data Figure 6. Round 2 Targeted Measurements in ROS/MAP**. Levels of 37 proteins were measured by selected reaction monitoring (SRM) mass spectrometry across 1016 ROS/MAP cases. Most proteins were measured with at least two peptides per protein, and therefore protein level measurements for these proteins are calculated as the average of the individual peptide measurements. Individual peptide measurements are shown after the protein level measurements. Peptide sequences can be found at https://www.synapse.org/#!Synapse:syn10468856. Protein and peptide levels were correlated to tau levels as measured by SRM, CERAD score, Braak stage, and MMSE. Differences in levels by case status were assessed by Kruskal-Wallis (K-W) one-way ANOVA with Tukey test. Correlations were performed using both Pearson correlation (cor) and biweight midcorrelation (bicor), which is more robust to outliers. Data for each protein and peptide is colored by the AD network module in which it resides. Statistical significance at *p* <0.05 is highlighted in red. Boxplots represent the median, 25^th^, and 75^th^ percentiles, and whiskers represent measurements to the 5^th^ and 95^th^ percentiles.

**Extended Data Figure 7. AD Network Preservation in Temporal Cortex.** The top 20% of proteins by kME value in each AD brain protein network module was used to create a synthetic eigenprotein, which was then measured by case status in the Mayo cohort (temporal cortex) and correlated with Braak score. The first boxplot for each module is the AD network eigenprotein by case status, given as reference for the second boxplot, which is the synthetic eigenprotein in the Mayo cohort. Synthetic eigenprotein differences by case status in the Mayo cohort were assessed by Kruskal-Wallis (K-W) one-way ANOVA. Correlations were performed using both Pearson correlation (cor) and biweight midcorrelation (bicor), which is more robust to outliers. Statistical significance at *p* <0.05 is highlighted in red. Boxplots represent the median, 25^th^, and 75^th^ percentiles, and whiskers represent measurements to the 5^th^ and 95^th^ percentiles.

**Extended Data Figure 8. AD Network Preservation in Precuneus.** The top 20% of proteins by kME value in each AD brain protein network module was used to create a synthetic eigenprotein, which was then measured by case status in the BLSA cohort (precuneus) and correlated with CERAD and Braak scores. The first boxplot for each module is the AD network eigenprotein by case status, given as reference for the second boxplot, which is the synthetic eigenprotein in the BLSA precuneus cohort. Synthetic eigenprotein differences by case status in the BLSA precuneus cohort were assessed by Kruskal-Wallis (K-W) one-way ANOVA. Correlations were performed using both Pearson correlation (cor) and biweight midcorrelation (bicor), which is more robust to outliers. Statistical significance at *p* <0.05 is highlighted in red. Boxplots represent the median, 25^th^, and 75^th^ percentiles, and whiskers represent measurements to the 5^th^ and 95^th^ percentiles.

**Extended Data Figure 9. Effect of Aging on AD Network Modules.** The top 20% of proteins by kME value in each AD brain protein network module was used to create a synthetic eigenprotein, which was then measured by age grouping and correlated with age in the aging cohort from the Baltimore coroner’s office. The first boxplot for each module is the AD network eigenprotein by case status, given as reference for the second and third boxplots, which is the synthetic eigenprotein in the BLSA precuneus cohort measured in two different age groupings. Synthetic eigenproteins were also correlated directly with age, using both Pearson correlation (cor) and biweight midcorrelation (bicor), which is more robust to outliers. Synthetic eigenprotein differences by case status were assessed by Kruskal-Wallis (K-W) one-way ANOVA. Statistical significance at *p* <0.05 is highlighted in red. Boxplots represent the median, 25^th^, and 75^th^ percentiles, and whiskers represent measurements to the 5^th^ and 95^th^ percentiles.

**Extended Data Figure 10. AD Protein Network Module Changes in Other Neurodegenerative Diseases**. The top 20% of proteins by kME value in each AD brain protein network module was used to create a synthetic eigenprotein, which was then measured in different neurodegenerative diseases in the UPenn cohort. The first boxplot for each module is the AD network eigenprotein by case status, given as reference for the second and third boxplots. Other neurodegenerative diseases (otherND) included amyotrophic lateral sclerosis (ALS), frontotemporal lobar degeneration with TAR DNA-binding protein 43 inclusions (FTLD-TDP), progressive supranuclear palsy (PSP), corticobasal degeneration (CBD), Parkinson’s disease and Parkinsons’s disease dementia (PD/PDD), and multiple system atrophy (MSA). Synthetic eigenproteins were also correlated with CERAD and Braak scores, using both Pearson correlation (cor) and biweight midcorrelation (bicor), which is more robust to outliers. Synthetic eigenprotein differences by case status were assessed by Kruskal-Wallis (K-W) one-way ANOVA. Differences between AD and other case groups were assessed by Dunnett’s test and are provided in **Supplementary Table 5**. Statistical significance at *p* <0.05 is highlighted in red. Boxplots represent the median, 25^th^, and 75^th^ percentiles, and whiskers represent measurements to the 5^th^ and 95^th^ percentiles.

**Extended Data Figure 11. Targeted Individual Protein Measurements in Other Neurodegenerative Diseases.** Peptides from proteins across all modules in the AD network were targeted for measurement by parallel reaction monitoring mass spectrometry (PRM) in a subset of cases in the UPenn cohort encompassing all disease groups. Other neurodegenerative diseases (otherND) included amyotrophic lateral sclerosis (ALS), frontotemporal lobar degeneration with TAR DNA-binding protein 43 inclusions (FTLD-TDP), progressive supranuclear palsy (PSP), corticobasal degeneration (CBD), Parkinson’s disease and Parkinsons’s disease dementia (PD/PDD), and multiple system atrophy (MSA). Protein levels were correlated with CERAD and Braak scores, using both Pearson correlation (cor) and biweight midcorrelation (bicor), which is more robust to outliers. Protein level differences by case status were assessed by Kruskal-Wallis (K-W) one-way ANOVA. Differences between AD and other case groups were assessed by Dunnett’s test and are provided in **Supplementary Table 5**. Data for each protein is colored by the AD network module in which it resides. Statistical significance at *p* <0.05 is highlighted in red. Boxplots represent the median, 25^th^, and 75^th^ percentiles, and whiskers represent measurements to the 5^th^ and 95^th^ percentiles.

**Extended Data Figure 12. AD Network Module Changes in Other Neurodegenerative Diseases by Targeted Mass Spectrometry Measurements.** A synthetic eigenprotein for each AD network module was created from proteins measured by parallel reaction monitoring mass spectrometry (PRM) that mapped to an AD network module. Individual protein measurements by disease group are provided in **Extended Data Figure 11**. Other neurodegenerative diseases (otherND) included amyotrophic lateral sclerosis (ALS), frontotemporal lobar degeneration with TAR DNA-binding protein 43 inclusions (FTLD-TDP), progressive supranuclear palsy (PSP), corticobasal degeneration (CBD), Parkinson’s disease and Parkinsons’s disease dementia (PD/PDD), and multiple system atrophy (MSA). Synthetic eigenproteins were correlated with CERAD and Braak scores, using both Pearson correlation (cor) and biweight midcorrelation (bicor), which is more robust to outliers. Synthetic eigenprotein differences by case status were assessed by Kruskal-Wallis (K-W) one-way ANOVA. Differences between AD and other case groups were assessed by Dunnett’s test and are provided in **Supplementary Table 5**. Statistical significance at *p* <0.05 is highlighted in red. Boxplots represent the median, 25^th^, and 75^th^ percentiles, and whiskers represent measurements to the 5^th^ and 95^th^ percentiles.

**Extended Data Figure 13. Relative Levels and Trait Correlations of Module M4 Proteins in AD CSF Cohort 1.** Relative levels of each protein were measured in control and AD CSF by tandem mass tag mass spectrometry (TMT-MS), and correlated with ELISA measurements of Aβ42, Tau, and phosphorylated tau, as well as Aβ42/Tau ratio and MoCA score. ELISA measurements are given in pg/mL. Differences in protein levels were assessed by Welch’s *t* test. Correlations were performed using both Pearson correlation (cor) and biweight midcorrelation (bicor), which is more robust to outliers. Statistical significance at *p* <0.05 is highlighted in red. Boxplots represent the median, 25^th^, and 75^th^ percentiles, and whiskers represent measurements to the 5^th^ and 95^th^ percentiles. MoCA, Montreal Cognitive Assessment (higher scores represent better cognitive function).

**Extended Data Figure 14. Relative Levels and Trait Correlations of Module M4 Proteins in AsymAD and AD CSF Cohort 2.** Relative levels of each protein were measured in control, AsymAD, and AD CSF by tandem mass tag mass spectrometry (TMT-MS), and correlated with ELISA measurements of Aβ42, Tau, and phosphorylated tau, as well as Aβ42/Tau ratio and MoCA score. ELISA measurements are given in pg/mL. Differences in protein levels were assessed by Kruskal-Wallis (K-W) one-way ANOVA with Tukey test. Correlations were performed using both Pearson correlation (cor) and biweight midcorrelation (bicor), which is more robust to outliers. Statistical significance at *p* <0.05 is highlighted in red. Boxplots represent the median, 25^th^, and 75^th^ percentiles, and whiskers represent measurements to the 5^th^ and 95^th^ percentiles. MoCA, Montreal Cognitive Assessment (higher scores represent better cognitive function).

