## Supplementary material for "A Consensus Proteomic Analysis of Alzheimer’s Disease Brain and Cerebrospinal Fluid Reveals Early Changes in Energy Metabolism Associated with Microglia and Astrocyte Activation": Johnson et al 2020 Extended Data Figures

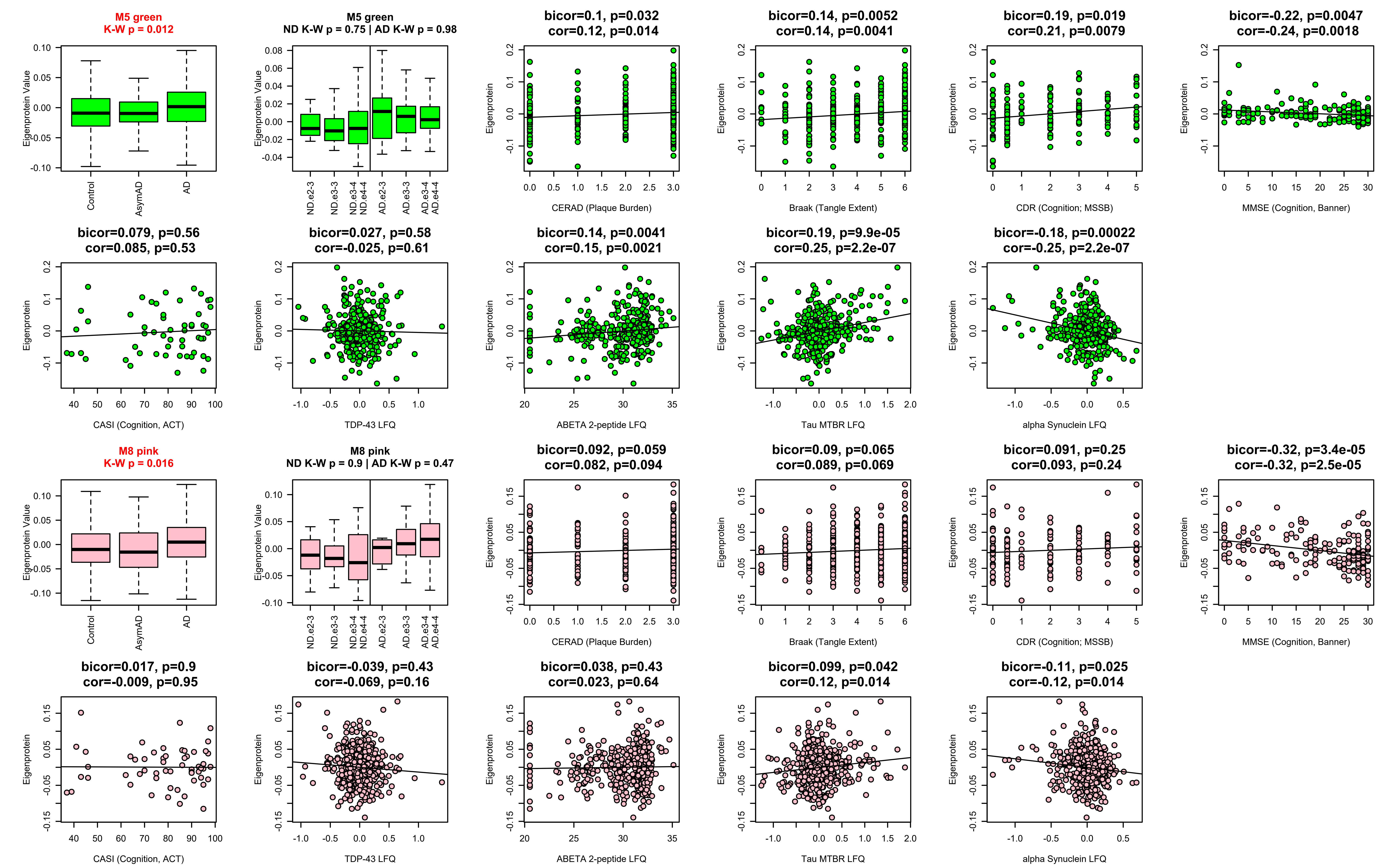
M5 green
K-W p = 0.012
Eigenprotein Value
Control
AsymAD
AD
M5 green
ND K-W p = 0.75 | AD K-W p = 0.98
Eigenprotein Value
ND e2-3
ND e3-3
ND e3-4
ND e4-4
AD e2-3
AD e3-3
AD e3-4
AD e4-4
bicor=0.1, p=0.032
cor=0.12, p=0.014
Eigenprotein
CERAD (Plaque Burden)
bicor=0.14, p=0.0052
cor=0.14, p=0.0041
Eigenprotein
Braak (Tangle Extent)
bicor=0.19, p=0.019
cor=0.21, p=0.0079
Eigenprotein
CDR (Cognition; MSSB)
bicor=-0.22, p=0.0047
cor=-0.24, p=0.0018
Eigenprotein
MMSE (Cognition, Banner)
bicor=0.079, p=0.56
cor=0.085, p=0.53
Eigenprotein
CASI (Cognition, ACT)
bicor=0.027, p=0.58
cor=-0.025, p=0.61
Eigenprotein
TDP-43 LFQ
bicor=0.14, p=0.0041
cor=0.15, p=0.0021
Eigenprotein
ABETA 2-peptide LFQ
bicor=0.19, p=9.9e-05
cor=0.25, p=2.2e-07
Eigenprotein
Tau MTBR LFQ
bicor=-0.18, p=0.00022
cor=-0.25, p=2.2e-07
Eigenprotein
alpha Synuclein LFQ
M8 pink
K-W p = 0.016
Eigenprotein Value
Control
AsymAD
AD
M8 pink
ND K-W p = 0.9 | AD K-W p = 0.47
Eigenprotein Value
ND e2-3
ND e3-3
ND e3-4
ND e4-4
AD e2-3
AD e3-3
AD e3-4
AD e4-4
bicor=0.092, p=0.059
cor=0.082, p=0.094
Eigenprotein
CERAD (Plaque Burden)
bicor=0.09, p=0.065
cor=0.089, p=0.069
Eigenprotein
Braak (Tangle Extent)
bicor=0.091, p=0.25
cor=0.093, p=0.24
Eigenprotein
CDR (Cognition; MSSB)
bicor=-0.32, p=3.4e-05
cor=-0.32, p=2.5e-05
Eigenprotein
MMSE (Cognition, Banner)
bicor=0.017, p=0.9
cor=-0.009, p=0.95
Eigenprotein
CASI (Cognition, ACT)
bicor=-0.039, p=0.43
cor=-0.069, p=0.16
Eigenprotein
TDP-43 LFQ
bicor=0.038, p=0.43
cor=0.023, p=0.64
Eigenprotein
ABETA 2-peptide LFQ
bicor=0.099, p=0.042
cor=0.12, p=0.014
Eigenprotein
Tau MTBR LFQ
bicor=-0.11, p=0.025
cor=-0.12, p=0.014
Eigenprotein
alpha Synuclein LFQ

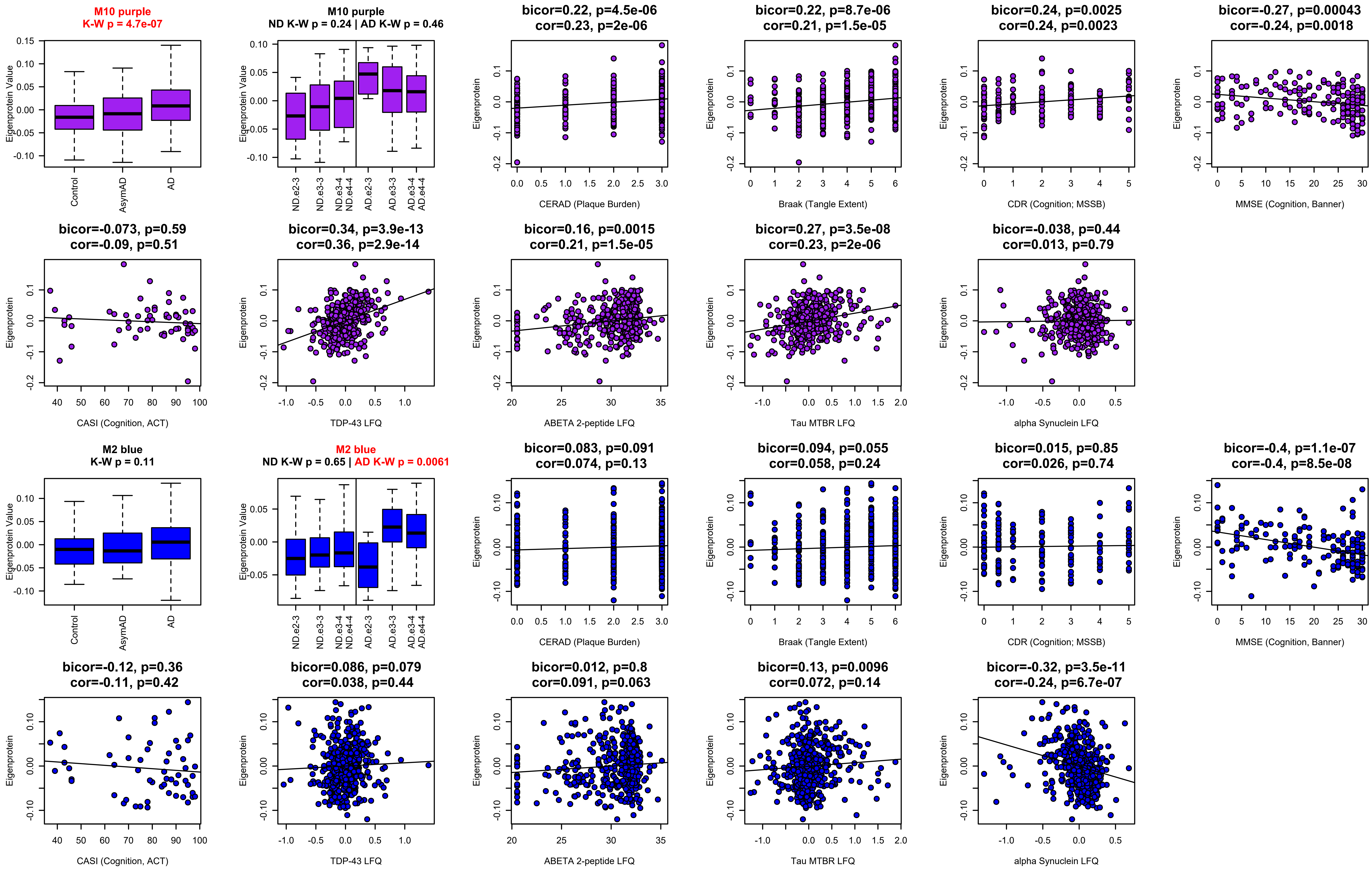

M10 purple
K-W p = 4.7e-07
Eigenprotein Value
Control
AsymAD
AD
M10 purple
ND K-W p = 0.24 | AD K-W p = 0.46
ND e2-3
ND e3-3
ND e3-4 ND e4-4
AD e2-3
AD e3-3
AD e3-4 AD e4-4
bicor=0.22, p=4.5e-06
cor=0.23, p=2e-06
CERAD (Plaque Burden)
bicor=0.22, p=8.7e-06
cor=0.21, p=1.5e-05
Braak (Tangle Extent)
bicor=0.24, p=0.0025
cor=0.24, p=0.0023
CDR (Cognition; MSSB)
bicor=-0.27, p=0.00043
cor=-0.24, p=0.0018
MMSE (Cognition, Banner)
bicor=-0.073, p=0.59
cor=-0.09, p=0.51
CASI (Cognition, ACT)
bicor=0.34, p=3.9e-13
cor=0.36, p=2.9e-14
TDP-43 LFQ
bicor=0.16, p=0.0015
cor=0.21, p=1.5e-05
ABETA 2-peptide LFQ
bicor=0.27, p=3.5e-08
cor=0.23, p=2e-06
Tau MTBR LFQ
bicor=-0.038, p=0.44
cor=0.013, p=0.79
alpha Synuclein LFQ
Eigenprotein
M2 blue
K-W p = 0.11
Eigenprotein Value
Control
AsymAD
AD
M2 blue
ND K-W p = 0.65 | AD K-W p = 0.0061
ND e2-3
ND e3-3
ND e3-4 ND e4-4
AD e2-3
AD e3-3
AD e3-4 AD e4-4
bicor=0.083, p=0.091
cor=0.074, p=0.13
CERAD (Plaque Burden)
bicor=0.094, p=0.055
cor=0.058, p=0.24
Braak (Tangle Extent)
bicor=0.015, p=0.85
cor=0.026, p=0.74
CDR (Cognition; MSSB)
bicor=-0.4, p=1.1e-07
cor=-0.4, p=8.5e-08
MMSE (Cognition, Banner)
bicor=-0.12, p=0.36
cor=-0.11, p=0.42
CASI (Cognition, ACT)
bicor=0.086, p=0.079
cor=0.038, p=0.44
TDP-43 LFQ
bicor=0.012, p=0.8
cor=0.091, p=0.063
ABETA 2-peptide LFQ
bicor=0.13, p=0.0096
cor=0.072, p=0.14
Tau MTBR LFQ
bicor=-0.32, p=3.5e-11
cor=-0.24, p=6.7e-07
alpha Synuclein LFQ
Eigenprotein

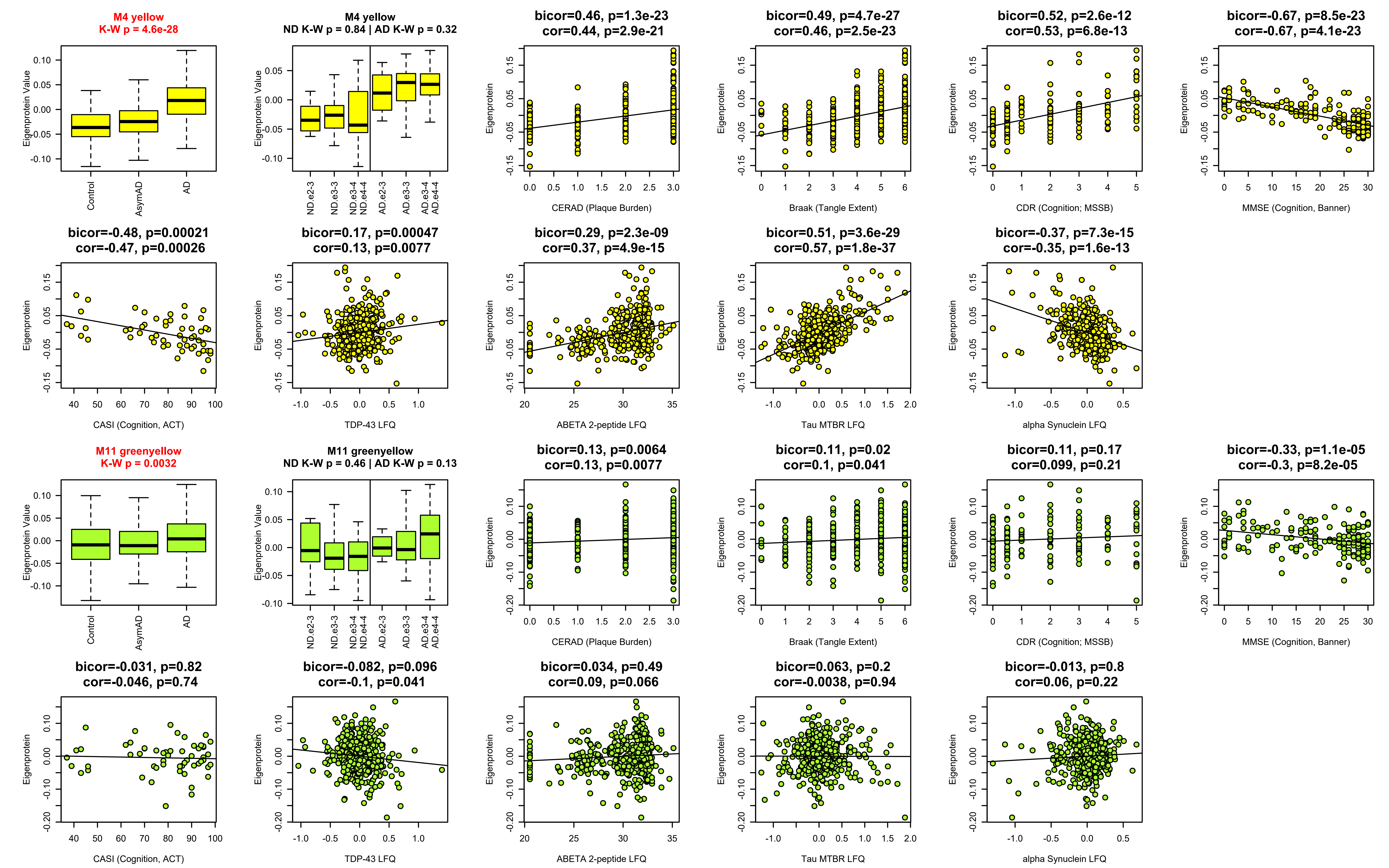

M4 yellow
K-W p = 4.6e-28
Eigenprotein Value
Control
AsymAD
AD
M4 yellow
ND K-W p = 0.84 | AD K-W p = 0.32
ND e2-3
ND e3-3
ND e3-4
ND e4-4
AD e2-3
AD e3-3
AD e3-4
AD e4-4
bicor=0.46, p=1.3e-23
cor=0.44, p=2.9e-21
CERAD (Plaque Burden)
bicor=0.49, p=4.7e-27
cor=0.46, p=2.5e-23
Braak (Tangle Extent)
bicor=0.52, p=2.6e-12
cor=0.53, p=6.8e-13
CDR (Cognition; MSSB)
bicor=-0.67, p=8.5e-23
cor=-0.67, p=4.1e-23
MMSE (Cognition, Banner)
Eigenprotein
bicor=-0.48, p=0.00021
cor=-0.47, p=0.00026
CASI (Cognition, ACT)
bicor=0.17, p=0.00047
cor=0.13, p=0.0077
TDP-43 LFQ
bicor=0.29, p=2.3e-09
cor=0.37, p=4.9e-15
ABETA 2-peptide LFQ
bicor=0.51, p=3.6e-29
cor=0.57, p=1.8e-37
Tau MTBR LFQ
bicor=-0.37, p=7.3e-15
cor=-0.35, p=1.6e-13
alpha Synuclein LFQ
M11 greenyellow
K-W p = 0.0032
M11 greenyellow
ND K-W p = 0.46 | AD K-W p = 0.13
bicor=0.13, p=0.0064
cor=0.13, p=0.0077
bicor=0.11, p=0.02
cor=0.1, p=0.041
bicor=0.11, p=0.17
cor=0.099, p=0.21
bicor=-0.33, p=1.1e-05
cor=-0.3, p=8.2e-05
bicor=-0.031, p=0.82
cor=-0.046, p=0.74
bicor=-0.082, p=0.096
cor=-0.1, p=0.041
bicor=0.034, p=0.49
cor=0.09, p=0.066
bicor=0.063, p=0.2
cor=-0.0038, p=0.94
bicor=-0.013, p=0.8
cor=0.06, p=0.22

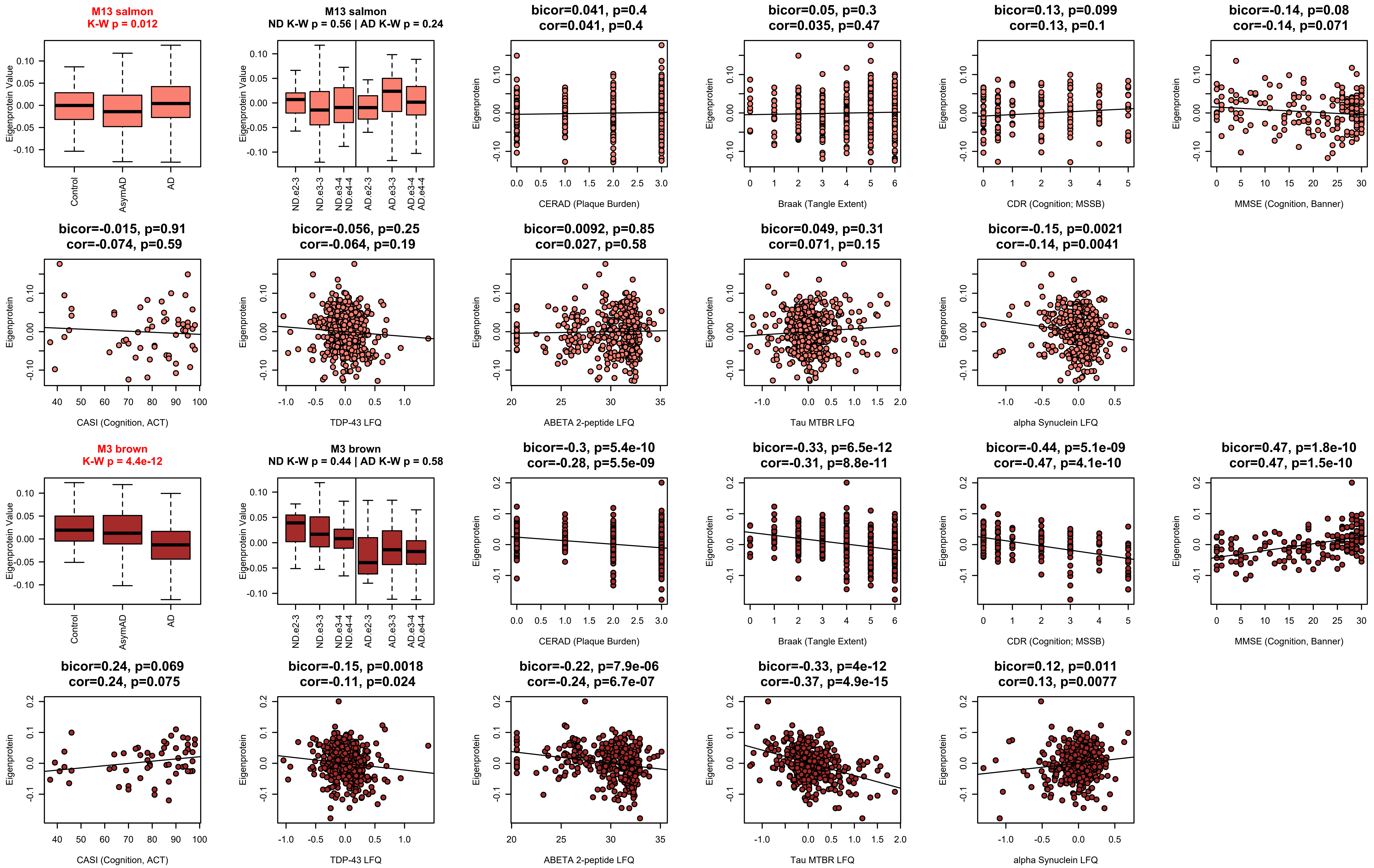
M13 salmon
K-W p = 0.012
Eigenprotein Value
Control
AsymAD
AD
M13 salmon
ND K-W p = 0.56 | AD K-W p = 0.24
ND e2-3
ND e3-3
ND e3-4 ND e4-4
AD e2-3
AD e3-3
AD e3-4 AD e4-4
bicor=0.041, p=0.4
cor=0.041, p=0.4
Eigenprotein
CERAD (Plaque Burden)
bicor=0.05, p=0.3
cor=0.035, p=0.47
Braak (Tangle Extent)
bicor=0.13, p=0.099
cor=0.13, p=0.1
CDR (Cognition; MSSB)
bicor=-0.14, p=0.08
cor=-0.14, p=0.071
MMSE (Cognition, Banner)
bicor=-0.015, p=0.91
cor=-0.074, p=0.59
CASI (Cognition, ACT)
bicor=-0.056, p=0.25
cor=-0.064, p=0.19
TDP-43 LFQ
bicor=0.0092, p=0.85
cor=0.027, p=0.58
ABETA 2-peptide LFQ
bicor=0.049, p=0.31
cor=0.071, p=0.15
Tau MTBR LFQ
bicor=-0.15, p=0.0021
cor=-0.14, p=0.0041
alpha Synuclein LFQ
M3 brown
K-W p = 4.4e-12
M3 brown
ND K-W p = 0.44 | AD K-W p = 0.58
bicor=-0.3, p=5.4e-10
cor=-0.28, p=5.5e-09
bicor=-0.33, p=6.5e-12
cor=-0.31, p=8.8e-11
bicor=-0.44, p=5.1e-09
cor=-0.47, p=4.1e-10
bicor=0.47, p=1.8e-10
cor=0.47, p=1.5e-10
bicor=0.24, p=0.069
cor=0.24, p=0.075
bicor=-0.15, p=0.0018
cor=-0.11, p=0.024
bicor=-0.22, p=7.9e-06
cor=-0.24, p=6.7e-07
bicor=-0.33, p=4e-12
cor=-0.37, p=4.9e-15
bicor=0.12, p=0.011
cor=0.13, p=0.0077

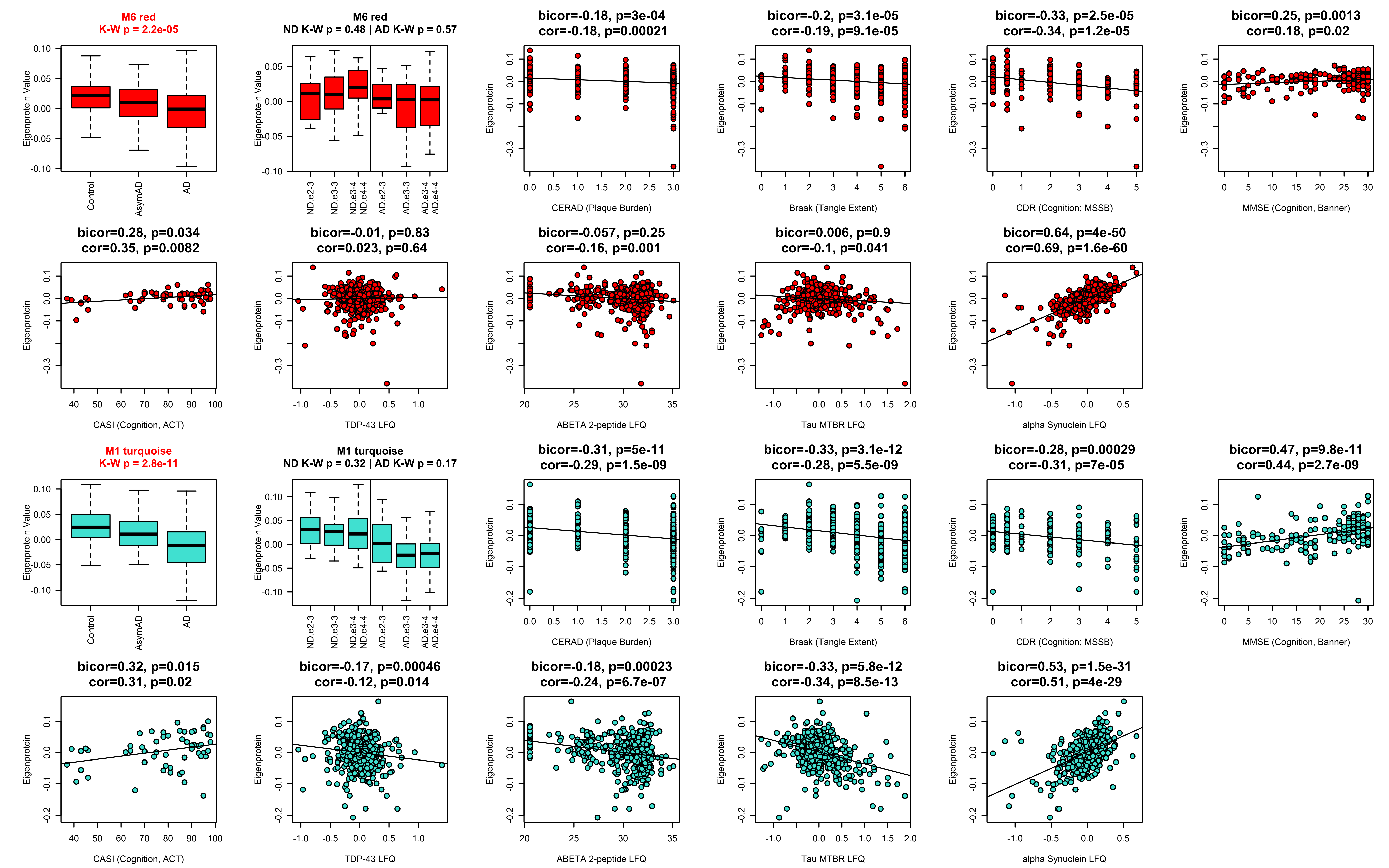
M6 red
K-W p = 2.2e-05
Eigenprotein Value
Control
AsymAD
AD
M6 red
ND K-W p = 0.48 | AD K-W p = 0.57
ND e2-3
ND e3-3
ND e3-4
ND e4-4
AD e2-3
AD e3-3
AD e3-4
AD e4-4
bicor=-0.18, p=3e-04
cor=-0.18, p=0.00021
CERAD (Plaque Burden)
bicor=-0.2, p=3.1e-05
cor=-0.19, p=9.1e-05
Braak (Tangle Extent)
bicor=-0.33, p=2.5e-05
cor=-0.34, p=1.2e-05
CDR (Cognition; MSSB)
bicor=0.25, p=0.0013
cor=0.18, p=0.02
MMSE (Cognition, Banner)
bicor=0.28, p=0.034
cor=0.35, p=0.0082
CASI (Cognition, ACT)
bicor=-0.01, p=0.83
cor=0.023, p=0.64
TDP-43 LFQ
bicor=-0.057, p=0.25
cor=-0.16, p=0.001
ABETA 2-peptide LFQ
bicor=0.006, p=0.9
cor=-0.1, p=0.041
Tau MTBR LFQ
bicor=0.64, p=4e-50
cor=0.69, p=1.6e-60
alpha Synuclein LFQ
Eigenprotein
M1 turquoise
K-W p = 2.8e-11
M1 turquoise
ND K-W p = 0.32 | AD K-W p = 0.17
bicor=-0.31, p=5e-11
cor=-0.29, p=1.5e-09
bicor=-0.33, p=3.1e-12
cor=-0.28, p=5.5e-09
bicor=-0.28, p=0.00029
cor=-0.31, p=7e-05
bicor=0.47, p=9.8e-11
cor=0.44, p=2.7e-09
bicor=0.32, p=0.015
cor=0.31, p=0.02
bicor=-0.17, p=0.00046
cor=-0.12, p=0.014
bicor=-0.18, p=0.00023
cor=-0.24, p=6.7e-07
bicor=-0.33, p=5.8e-12
cor=-0.34, p=8.5e-13
bicor=0.53, p=1.5e-31
cor=0.51, p=4e-29

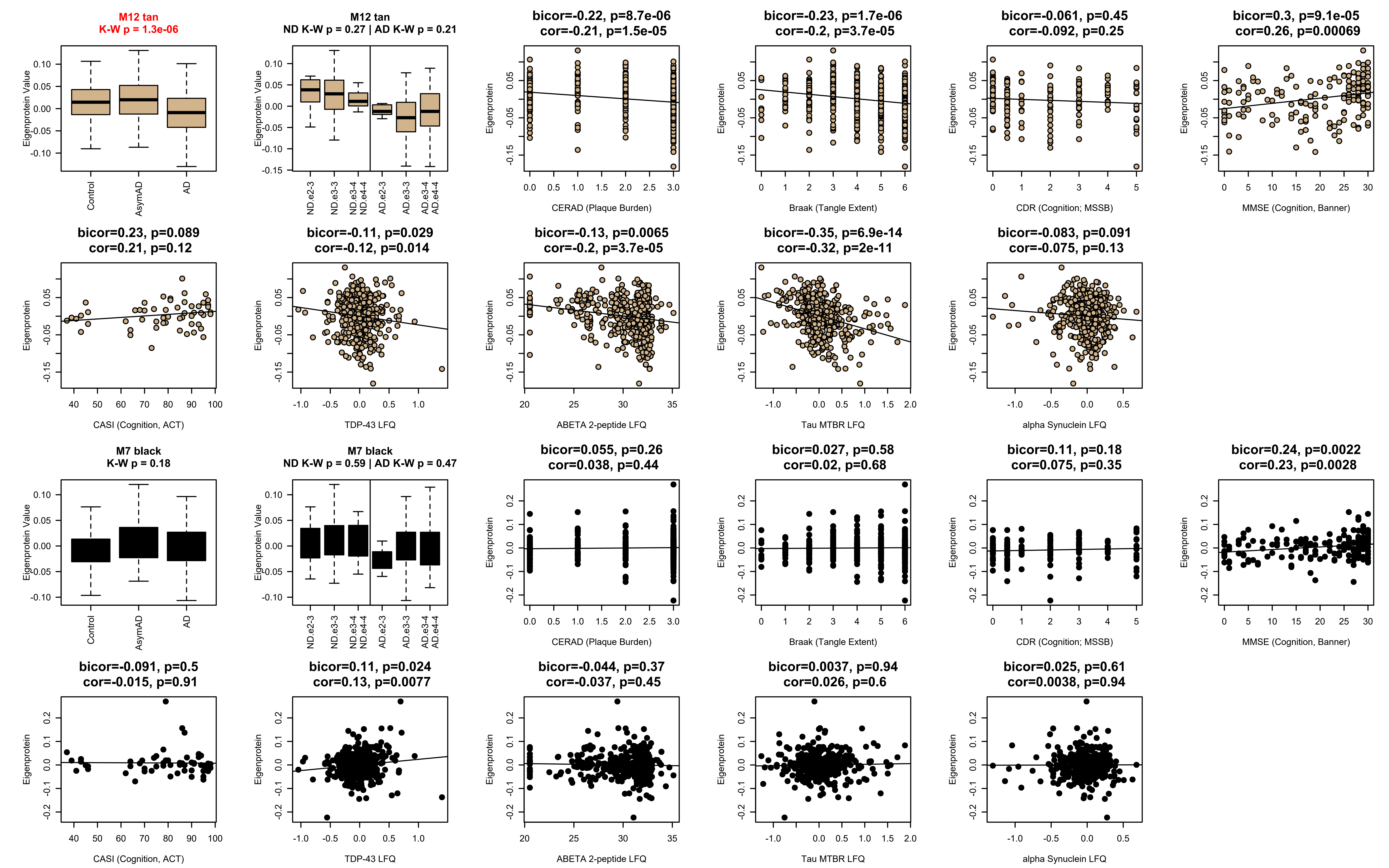

M12 tan
K-W p = 1.3e-06
Eigenprotein Value
Control
AsymAD
AD
M12 tan
ND K-W p = 0.27 | AD K-W p = 0.21
ND e2-3
ND e3-3
ND e3-4 ND e4-4
AD e2-3
AD e3-3
AD e3-4 AD e4-4
bicor=-0.22, p=8.7e-06
cor=-0.21, p=1.5e-05
CERAD (Plaque Burden)
bicor=-0.23, p=1.7e-06
cor=-0.2, p=3.7e-05
Braak (Tangle Extent)
bicor=-0.061, p=0.45
cor=-0.092, p=0.25
CDR (Cognition; MSSB)
bicor=0.3, p=9.1e-05
cor=0.26, p=0.00069
MMSE (Cognition, Banner)
bicor=0.23, p=0.089
cor=0.21, p=0.12
CASI (Cognition, ACT)
bicor=-0.11, p=0.029
cor=-0.12, p=0.014
TDP-43 LFQ
bicor=-0.13, p=0.0065
cor=-0.2, p=3.7e-05
ABETA 2-peptide LFQ
bicor=-0.35, p=6.9e-14
cor=-0.32, p=2e-11
Tau MTBR LFQ
bicor=-0.083, p=0.091
cor=-0.075, p=0.13
alpha Synuclein LFQ
Eigenprotein
M7 black
K-W p = 0.18
Eigenprotein Value
Control
AsymAD
AD
M7 black
ND K-W p = 0.59 | AD K-W p = 0.47
ND e2-3
ND e3-3
ND e3-4 ND e4-4
AD e2-3
AD e3-3
AD e3-4 AD e4-4
bicor=0.055, p=0.26
cor=0.038, p=0.44
CERAD (Plaque Burden)
bicor=0.027, p=0.58
cor=0.02, p=0.68
Braak (Tangle Extent)
bicor=0.11, p=0.18
cor=0.075, p=0.35
CDR (Cognition; MSSB)
bicor=0.24, p=0.0022
cor=0.23, p=0.0028
MMSE (Cognition, Banner)
bicor=-0.091, p=0.5
cor=-0.015, p=0.91
CASI (Cognition, ACT)
bicor=0.11, p=0.024
cor=0.13, p=0.0077
TDP-43 LFQ
bicor=-0.044, p=0.37
cor=-0.037, p=0.45
ABETA 2-peptide LFQ
bicor=0.0037, p=0.94
cor=0.026, p=0.6
Tau MTBR LFQ
bicor=0.025, p=0.61
cor=0.0038, p=0.94
alpha Synuclein LFQ
Eigenprotein

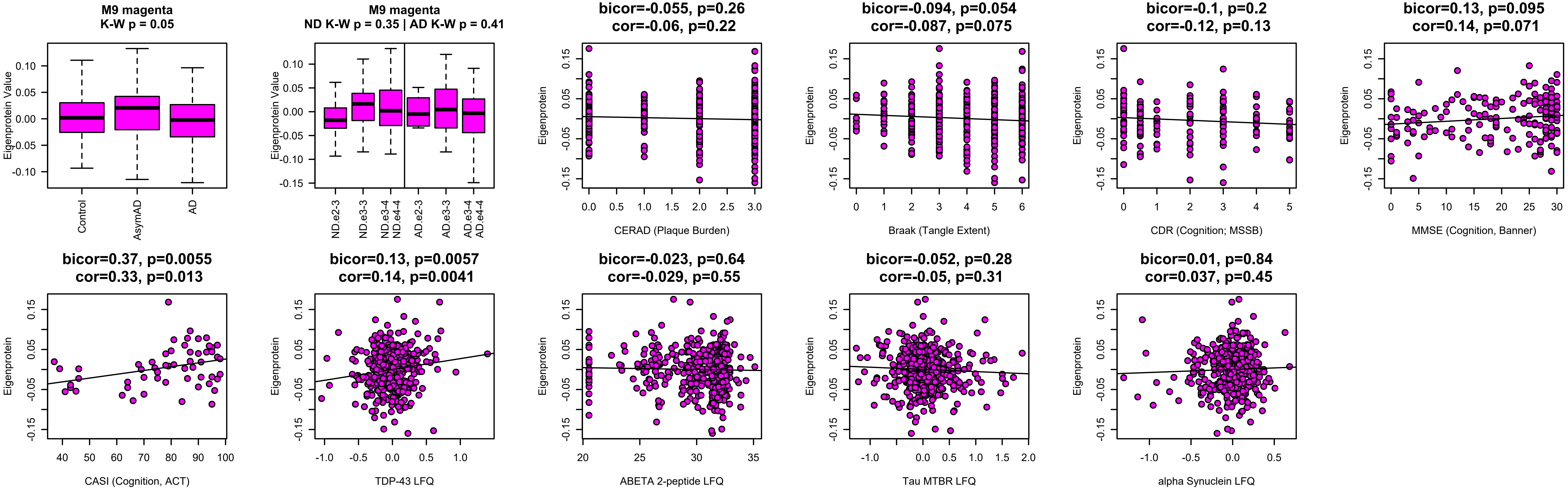
M9 magenta
K-W p = 0.05
Eigenprotein Value
Control
AsymAD
AD
M9 magenta
ND K-W p = 0.35 | AD K-W p = 0.41
ND e2-3
ND e3-3
ND e3-4
ND e4-4
AD e2-3
AD e3-3
AD e3-4
AD e4-4
bicor=-0.055, p=0.26
cor=-0.06, p=0.22
CERAD (Plaque Burden)
bicor=-0.094, p=0.054
cor=-0.087, p=0.075
Braak (Tangle Extent)
bicor=-0.1, p=0.2
cor=-0.12, p=0.13
CDR (Cognition; MSSB)
bicor=0.13, p=0.095
cor=0.14, p=0.071
MMSE (Cognition, Banner)
bicor=0.37, p=0.0055
cor=0.33, p=0.013
CASI (Cognition, ACT)
bicor=0.13, p=0.0057
cor=0.14, p=0.0041
TDP-43 LFQ
bicor=-0.023, p=0.64
cor=-0.029, p=0.55
ABETA 2-peptide LFQ
bicor=-0.052, p=0.28
cor=-0.05, p=0.31
Tau MTBR LFQ
bicor=0.01, p=0.84
cor=0.037, p=0.45
alpha Synuclein LFQ
Eigenprotein

**M5 Extracellular Matrix**

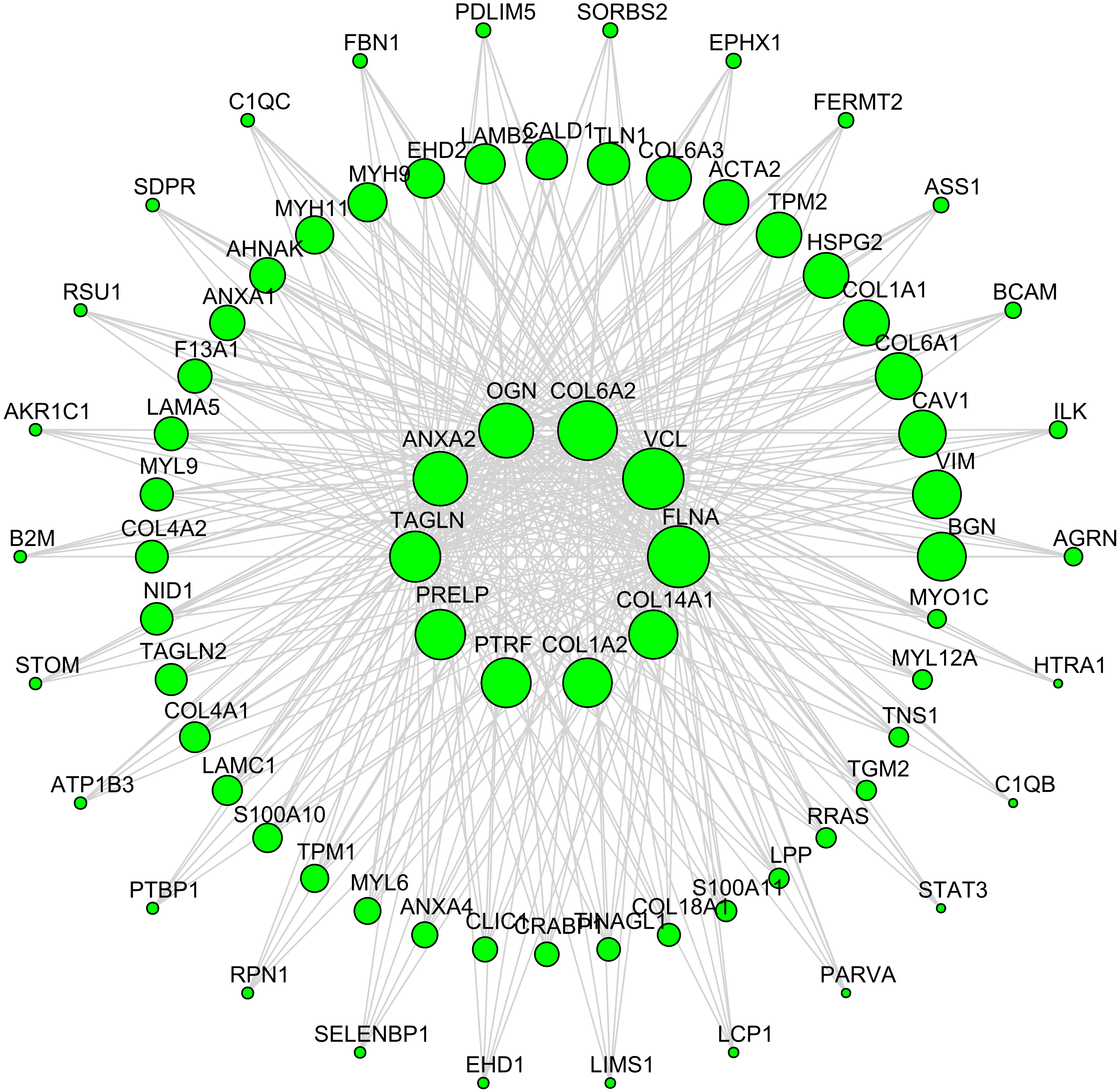

### M8 Blood

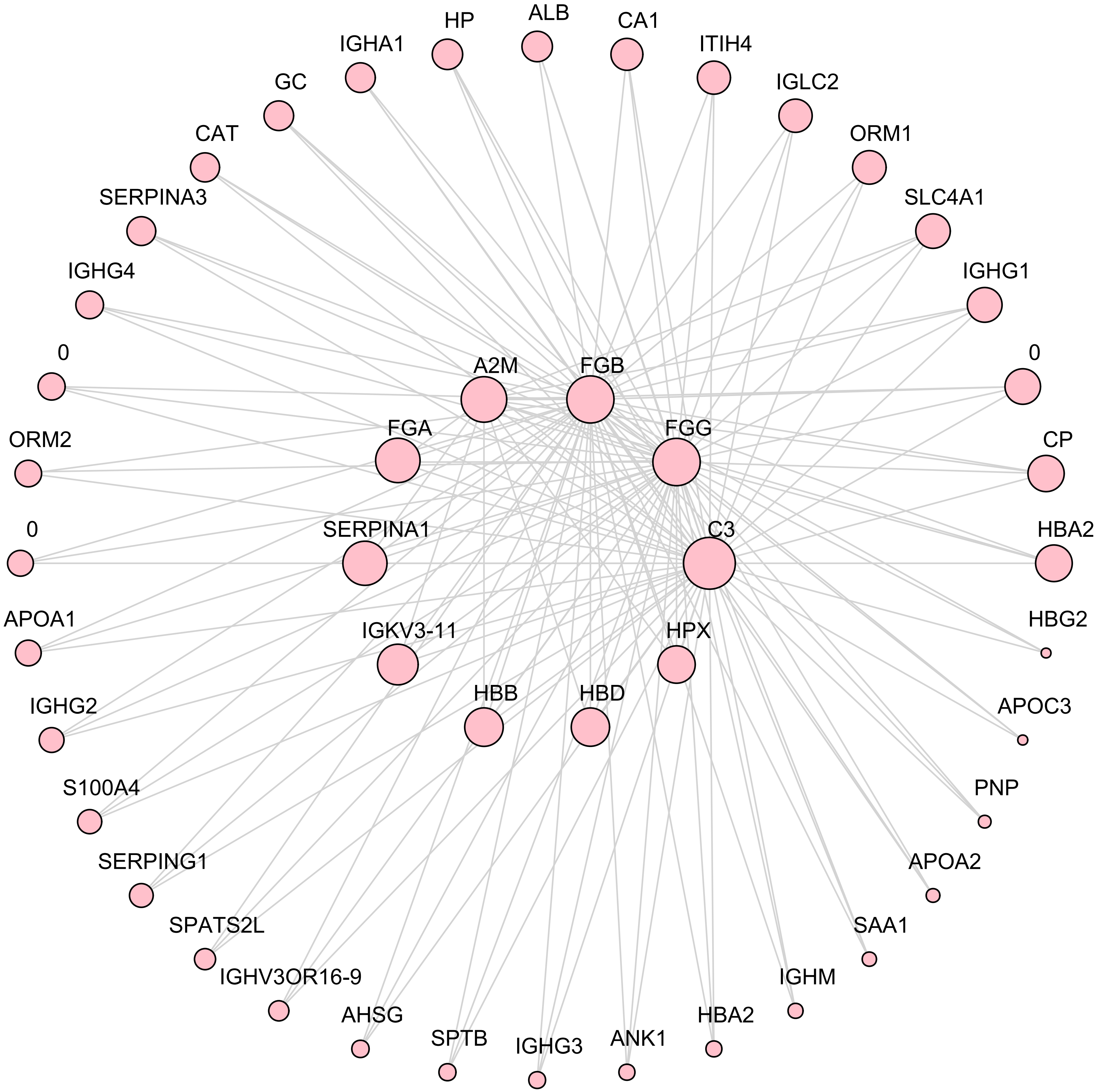
HP
ALB
CA1
IGHA1
ITIH4
GC
IGLC2
CAT
ORM1
SERPINA3
SLC4A1
IGHG4
IGHG1
0
A2M
FGB
0
ORM2
FGA
FGG
CP
0
SERPINA1
C3
HBA2
APOA1
IGKV3-11
HPX
HBG2
HBB
HBD
IGHG2
APOC3
S100A4
PNP
SERPING1
APOA2
SPATS2L
SAA1
IGHV3OR16-9
IGHM
AHSG
HBA2
SPTB
ANK1
IGHG3

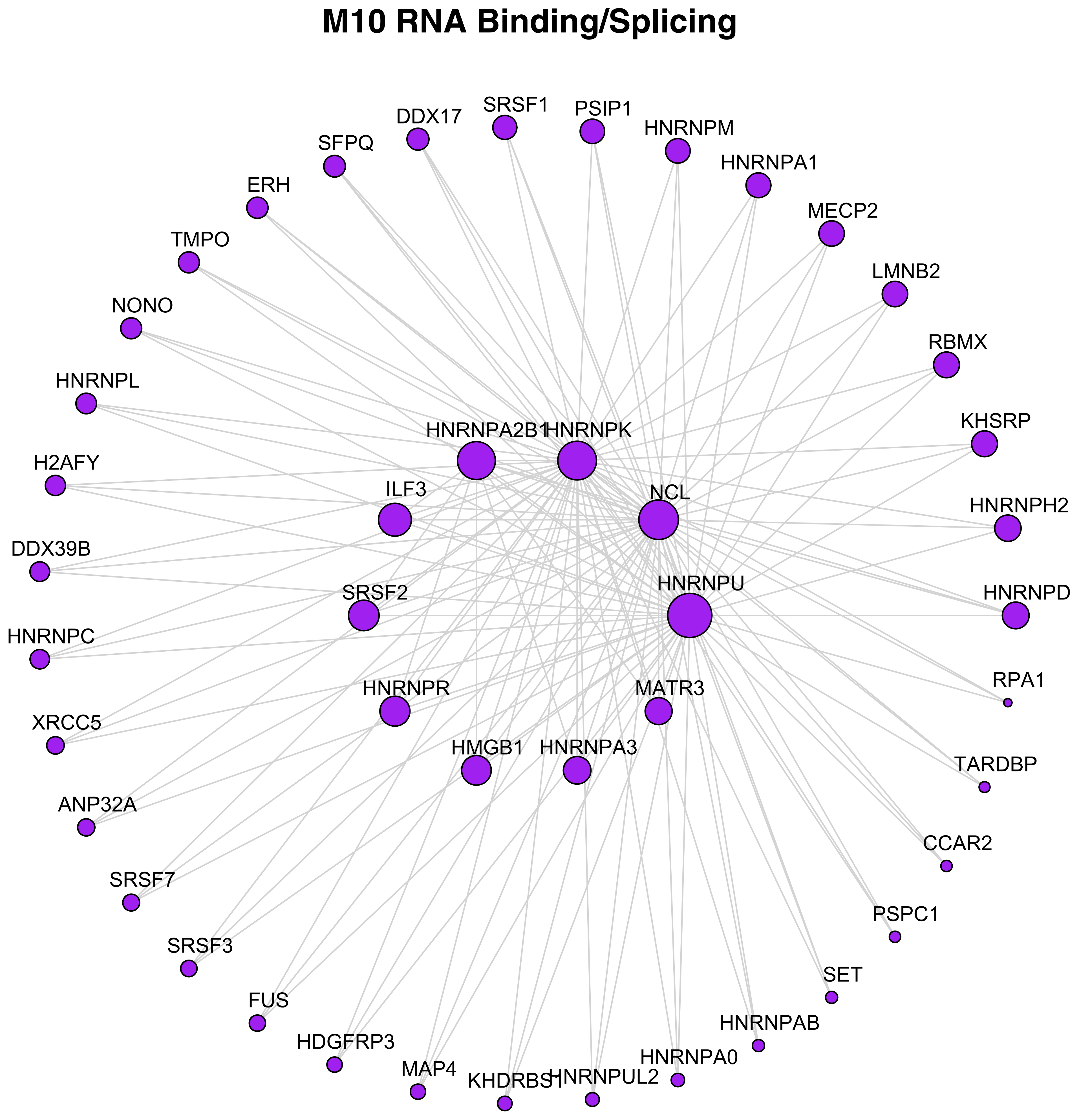
M10 RNA Binding/Splicing
SRSF1
PSIP1
DDX17
HNRNPM
SFPQ
HNRNPA1
ERH
MECP2
TMPO
LMNB2
NONO
RBMX
HNRNPL
KHSRP
HNRNPA2B1
HNRNPK
H2AFY
ILF3
NCL
HNRNPH2
DDX39B
HNRNPU
SRSF2
HNRNPD
HNRNPC
RPA1
HNRNPR
MATR3
XRCC5
HMGB1
HNRNPA3
TARDBP
ANP32A
CCAR2
SRSF7
PSPC1
SRSF3
SET
FUS
HNRNPAB
HDGFRP3
HNRNPA0
MAP4
KHDRBS1
HNRNPUL2

### M2 Myelin/Oligodendrocyte

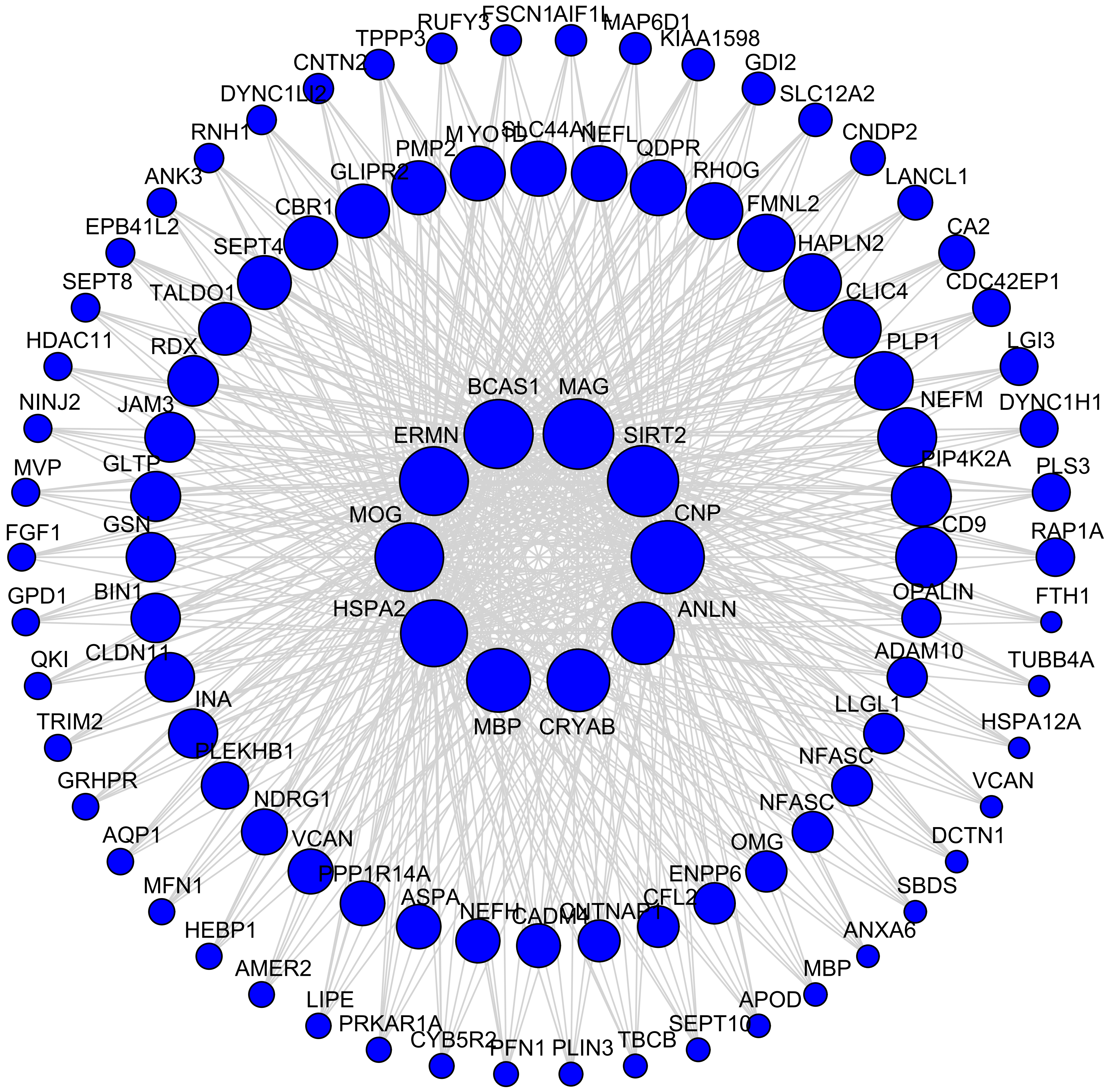

#### M4 Sugar Metabolism

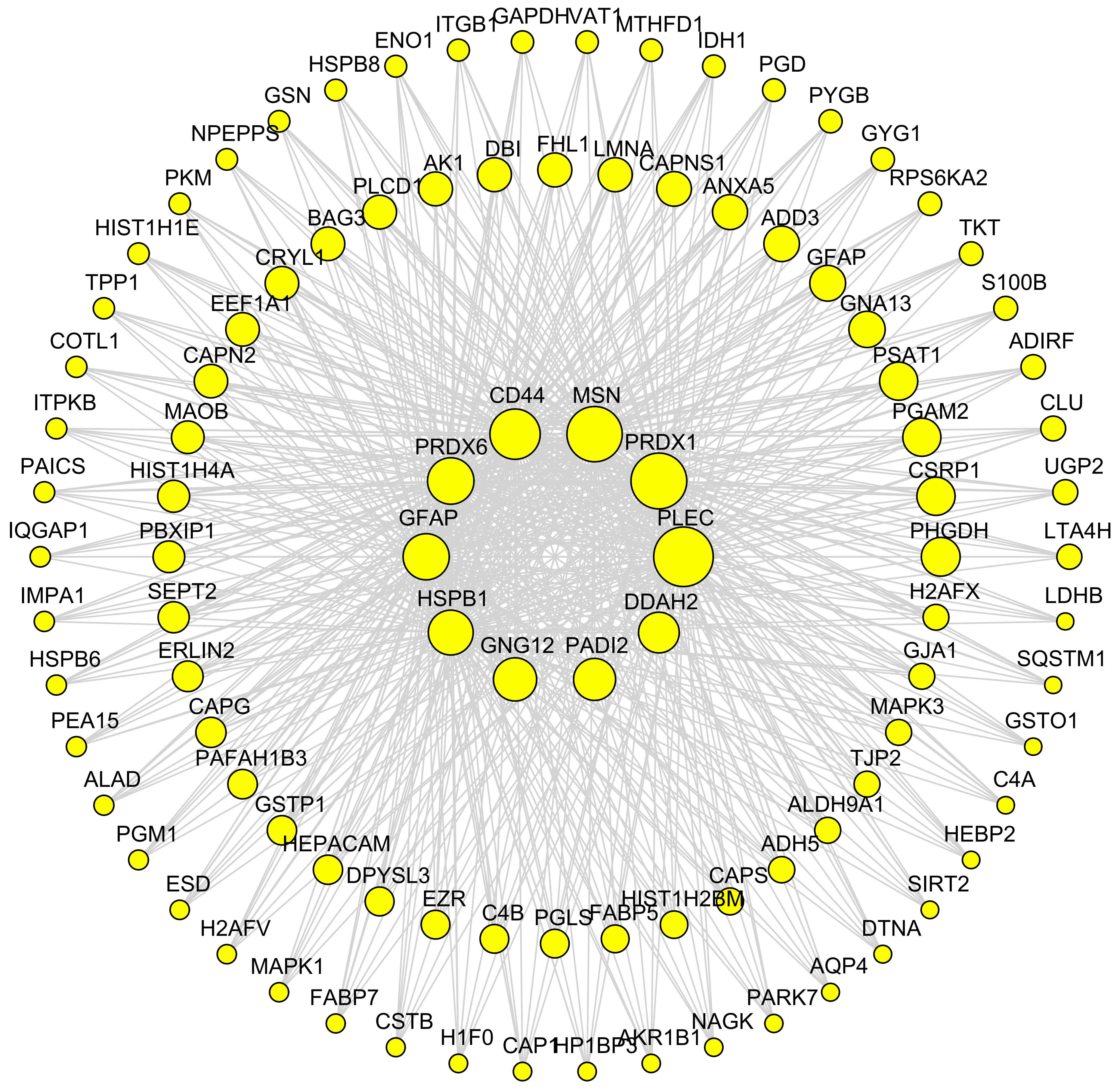

### M11 Chaperone/Protein Folding

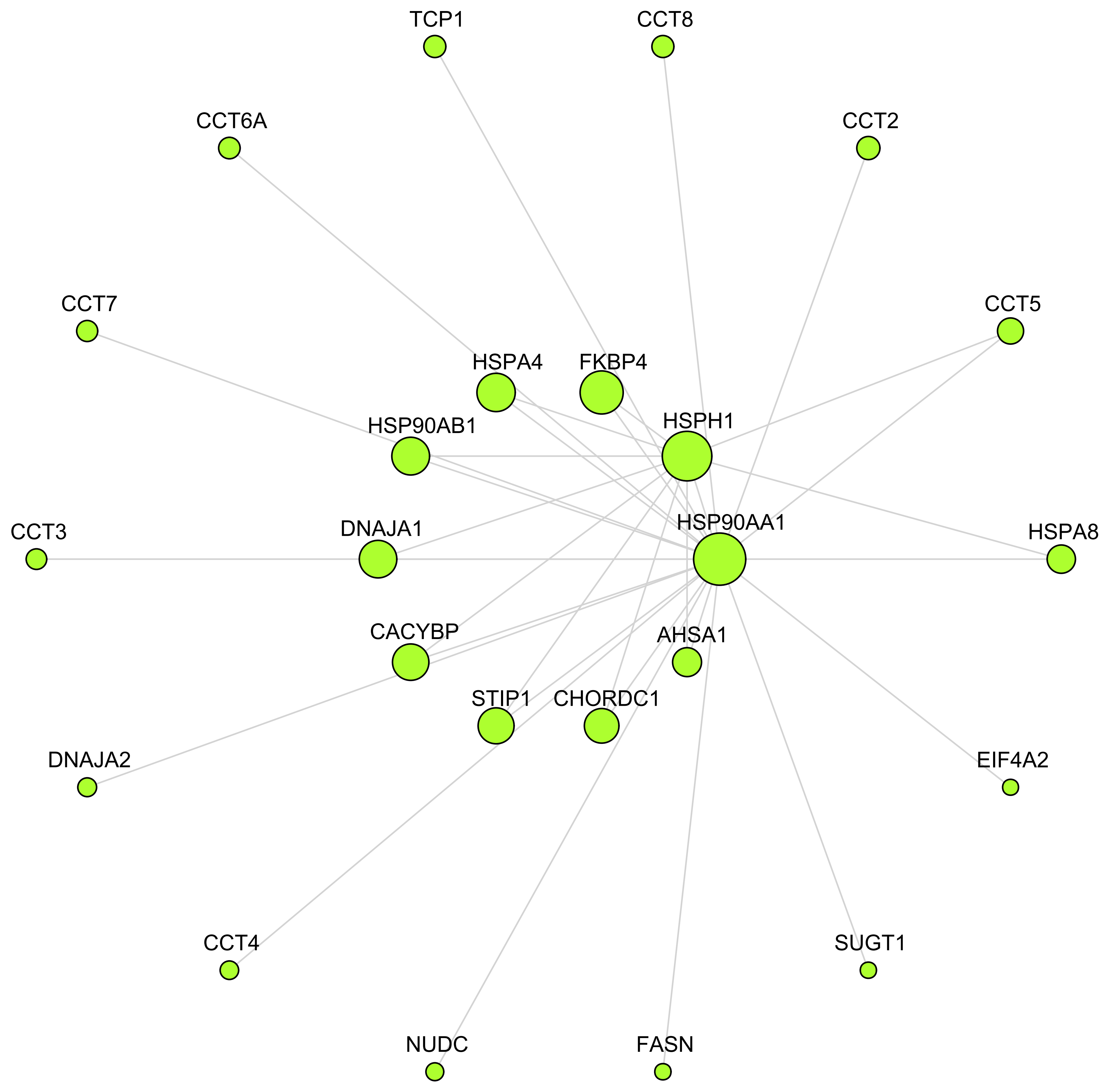

### M13 Unknown

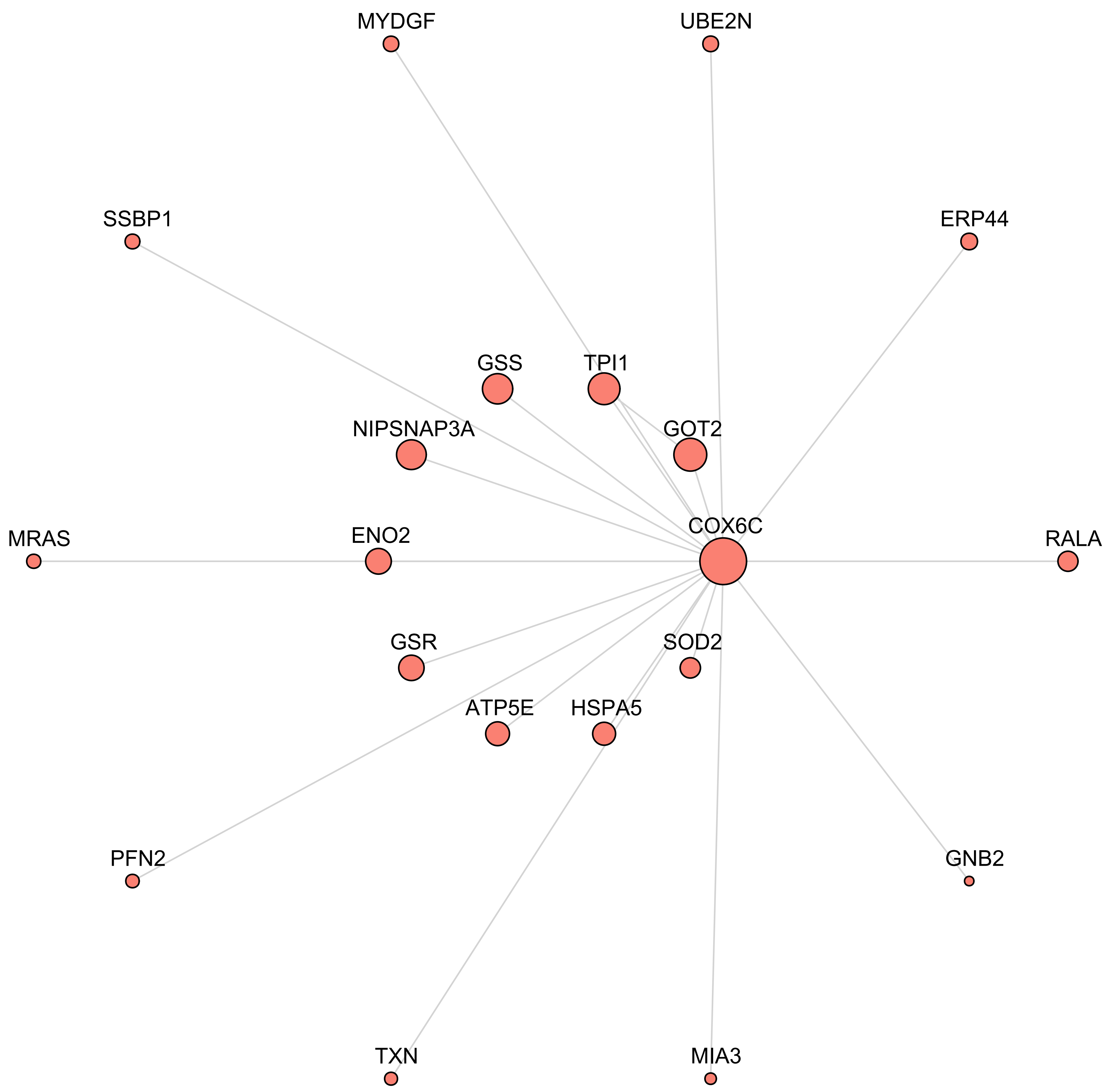
MYDGF
UBE2N
SSBP1
ERP44
GSS
TPI1
NIPSNAP3A
GOT2
MRAS
ENO2
COX6C
RALA
GSR
SOD2
ATP5E
HSPA5
PFN2
GNB2
TXN
MIA3

**M3 Mitochondrial**

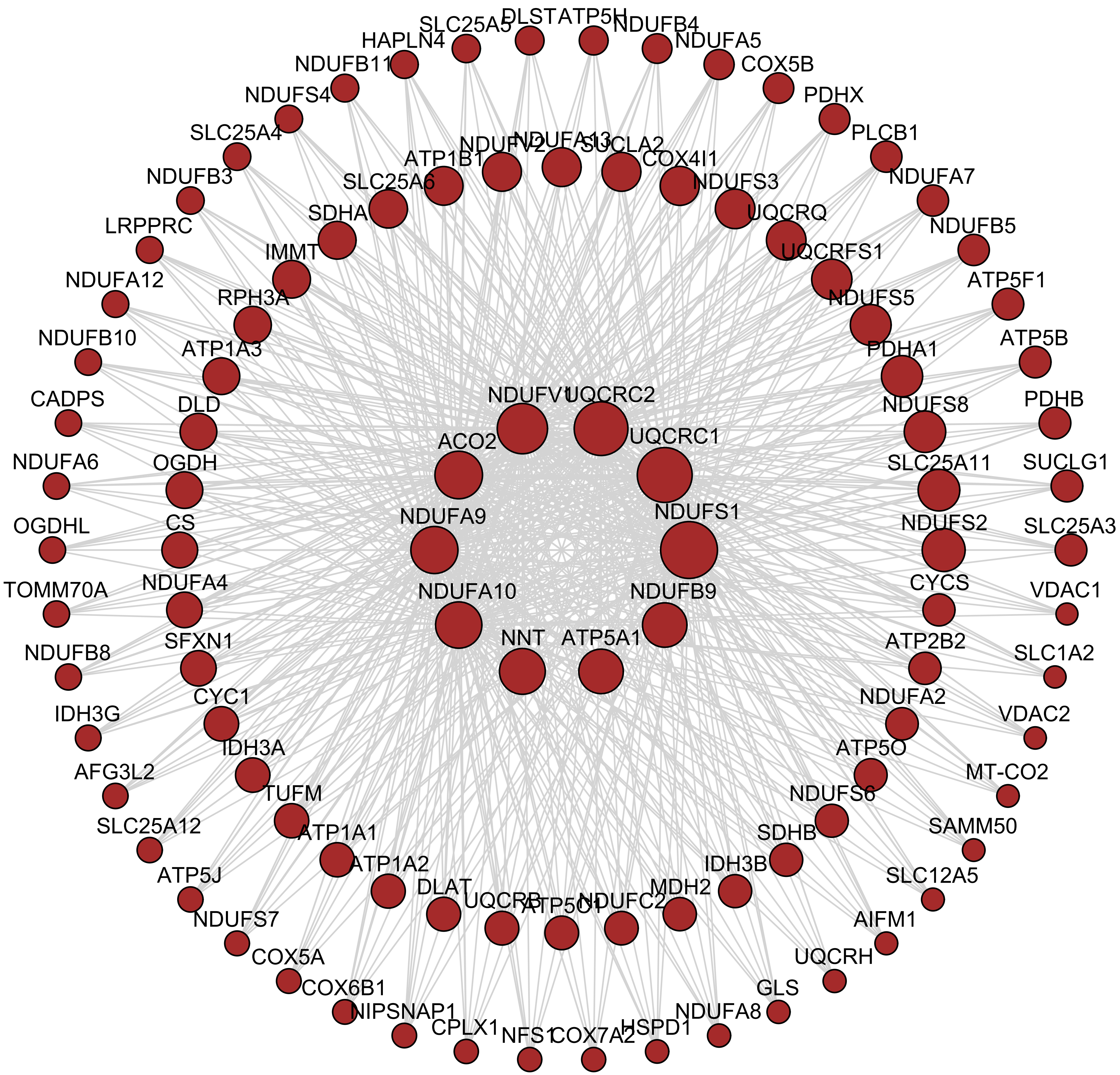

#### M6 Cytoskeleton

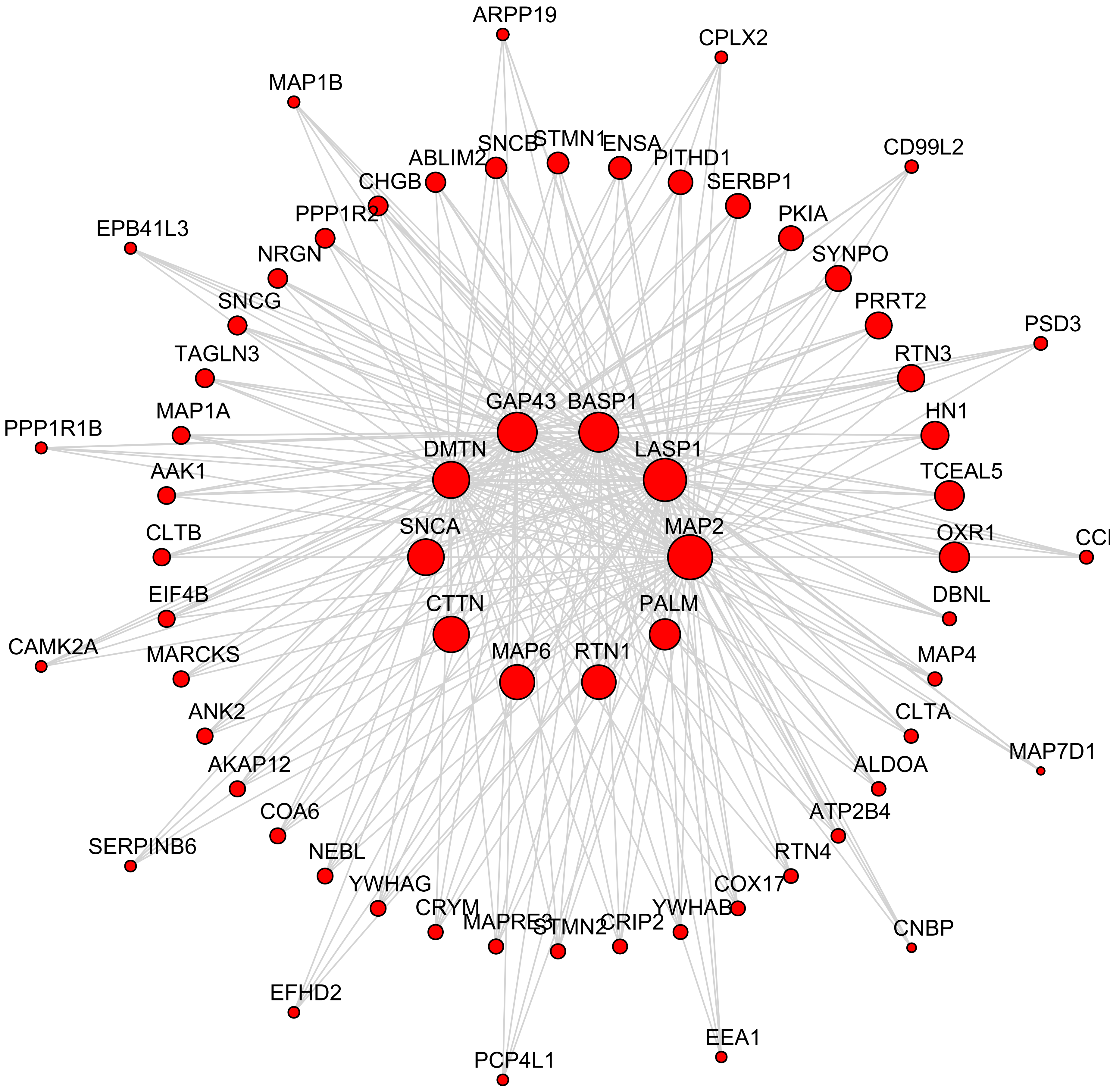

#### M1 Synapse

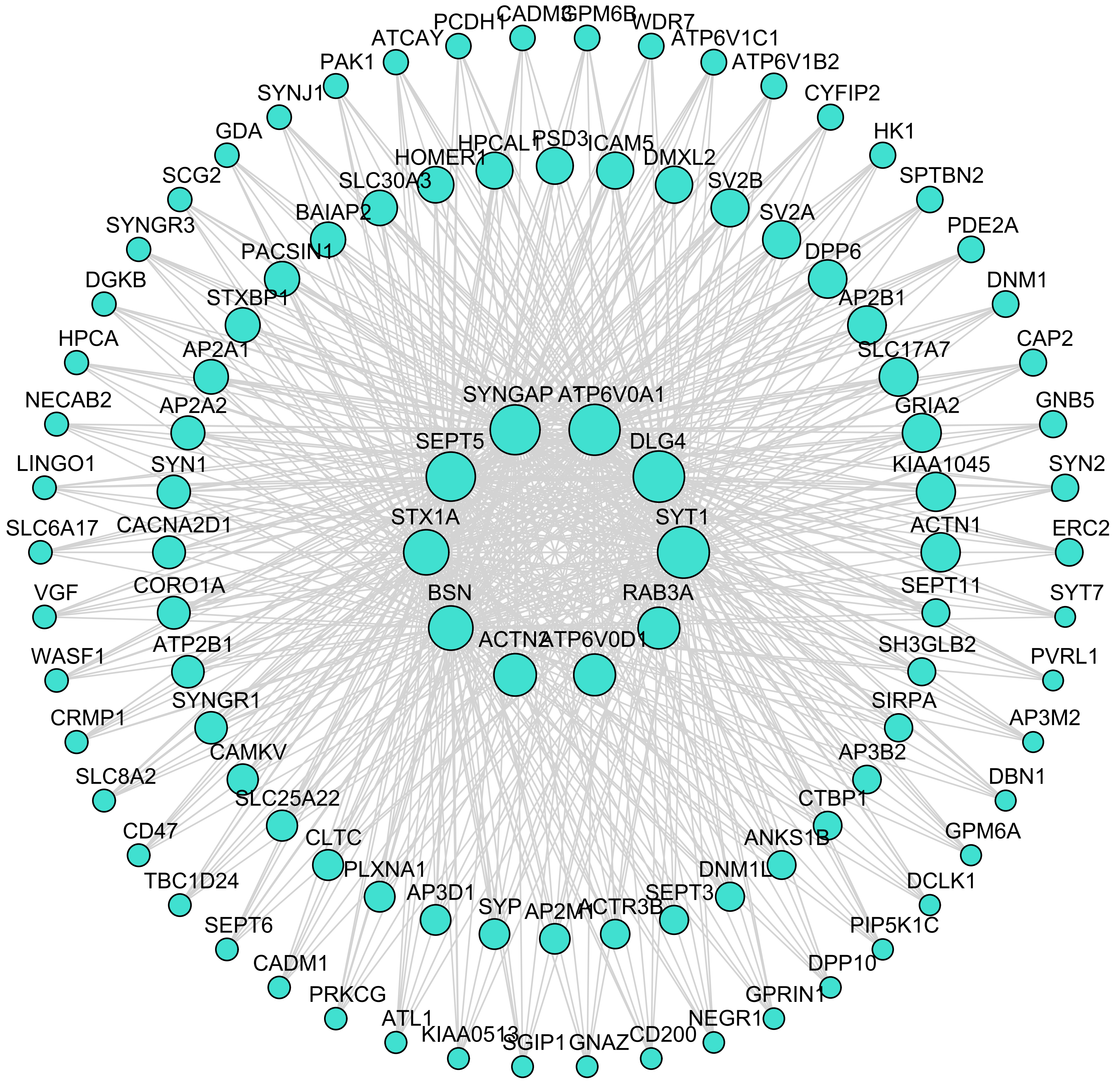
SYNGAP
ATP6V0A1
SEPT5
DLG4
STX1A
SYT1
BSN
RAB3A
ACTN2
ATP6V0D1
HOMER1
HPCAL1
PSD3
ICAM5
DMXL2
SV2B
SV2A
DPP6
AP2B1
SLC17A7
GRIA2
KIAA1045
ACTN1
SEPT11
SH3GLB2
SIRPA
AP3B2
CTBP1
ANKS1B
DNM1L
SEPT3
ACTR3B
AP2M1
SYP
AP3D1
PLXNA1
CLTC
SLC25A22
CAMKV
SYNGR1
ATP2B1
CORO1A
CACNA2D1
SYN1
AP2A2
AP2A1
STXBP1
PACSIN1
BAIAP2
SLC30A3
ATCAY
PCDH1
CADM3
GPM6B
WDR7
ATP6V1C1
ATP6V1B2
CYFIP2
HK1
SPTBN2
PDE2A
DNM1
CAP2
GNB5
SYN2
ERC2
SYT7
PVRL1
AP3M2
DBN1
GPM6A
DCLK1
PIP5K1C
DPP10
GPRIN1
NEGR1
CD200
GNAZ
SGIP1
KIAA0513
ATL1
PRKCG
CADM1
SEPT6
TBC1D24
CD47
SLC8A2
CRMP1
WASF1
VGF
SLC6A17
LINGO1
NECAB2
HPCA
DGKB
SYNGR3
SCG2
GDA
SYNJ1
PAK1

### M12 Unknown

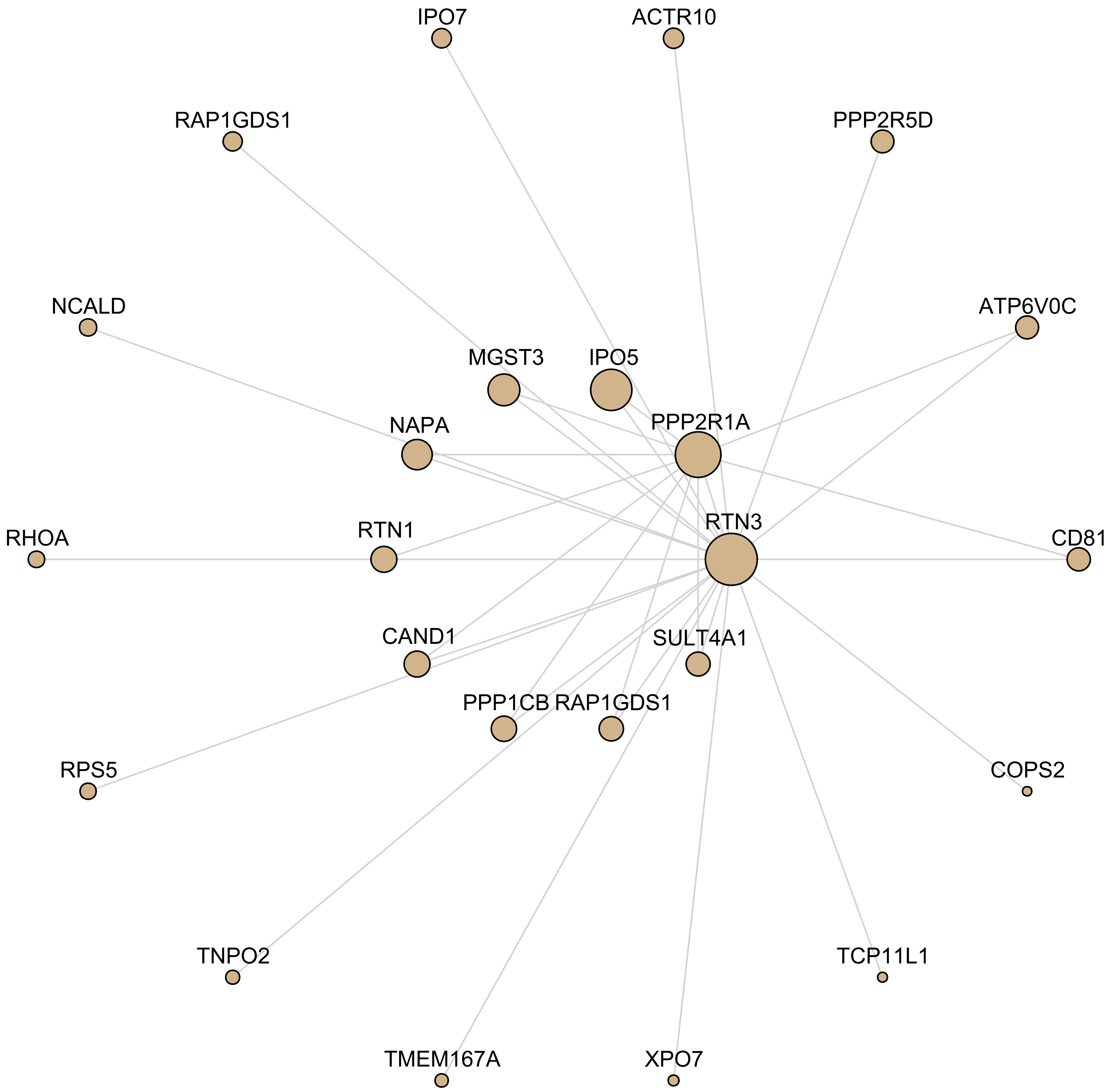

IPO7
ACTR10
RAP1GDS1
PPP2R5D
NCALD
ATP6V0C
MGST3
IPO5
NAPA
PPP2R1A
RTN1
RTN3
RHOA
CD81
CAND1
SULT4A1
PPP1CB
RAP1GDS1
RPS5
COPS2
TNPO2
TCP11L1
TMEM167A
XPO7

#### M7 Translation/Ribosome

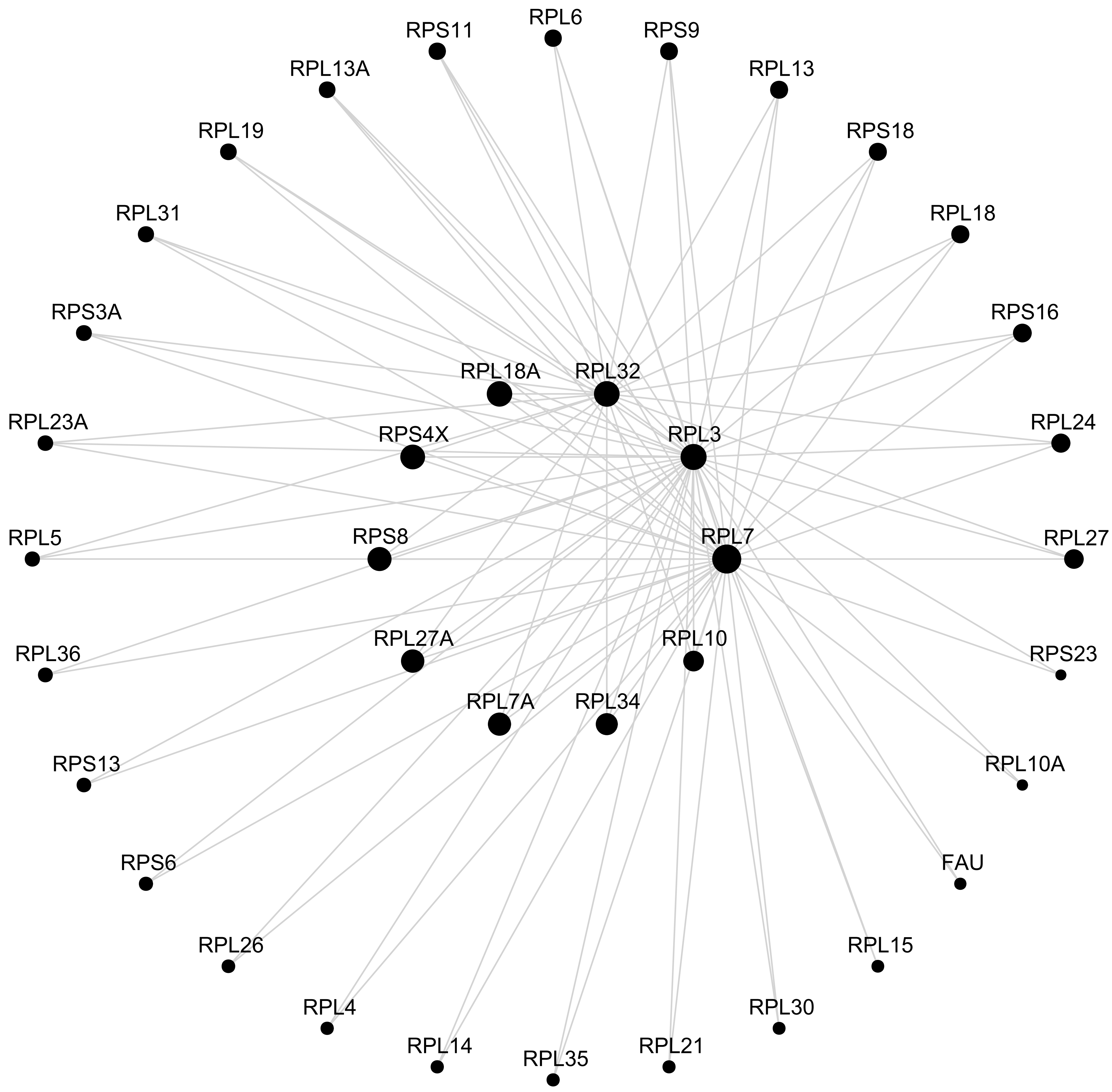

### M9 Translation/Ribosome

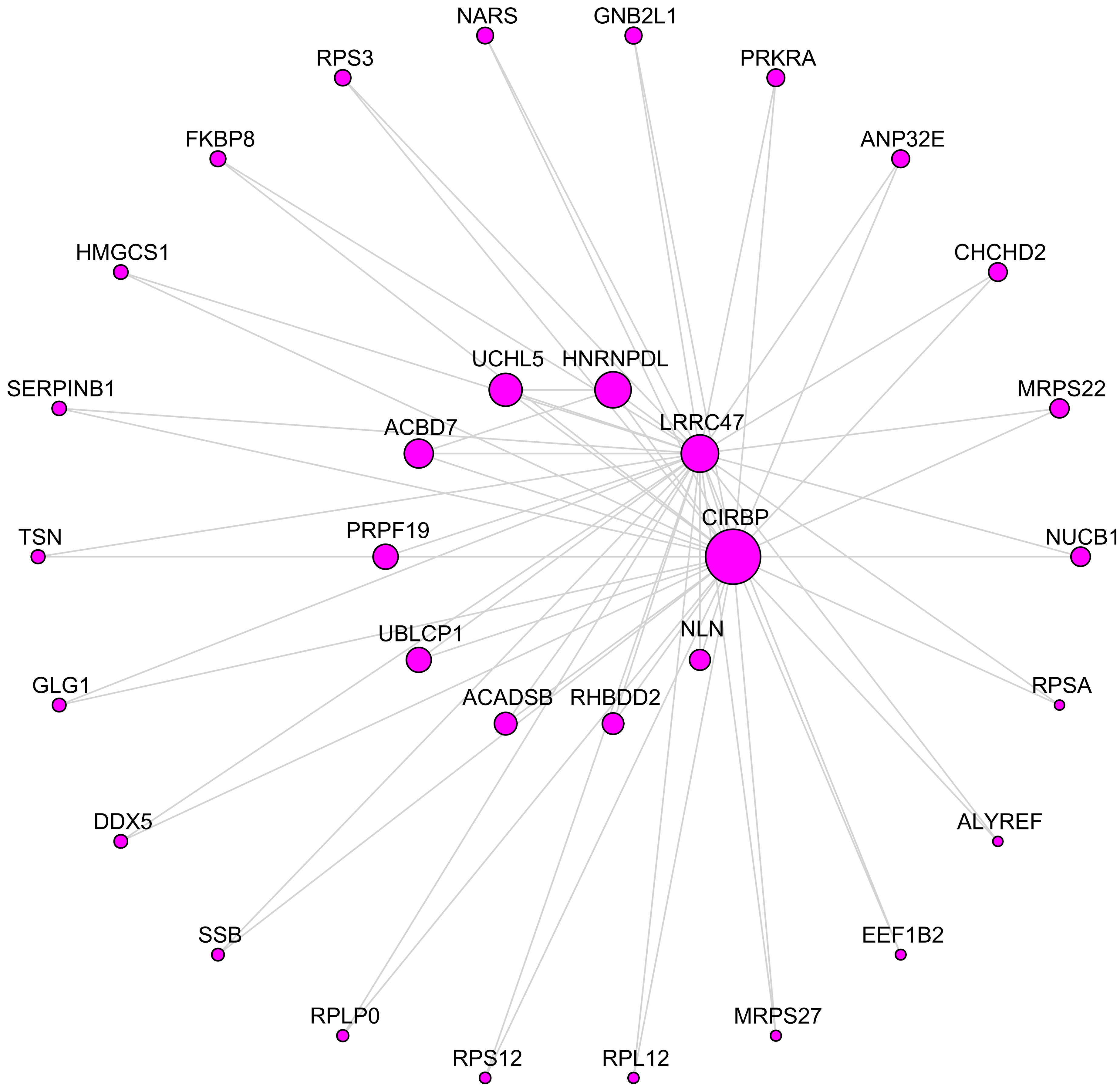

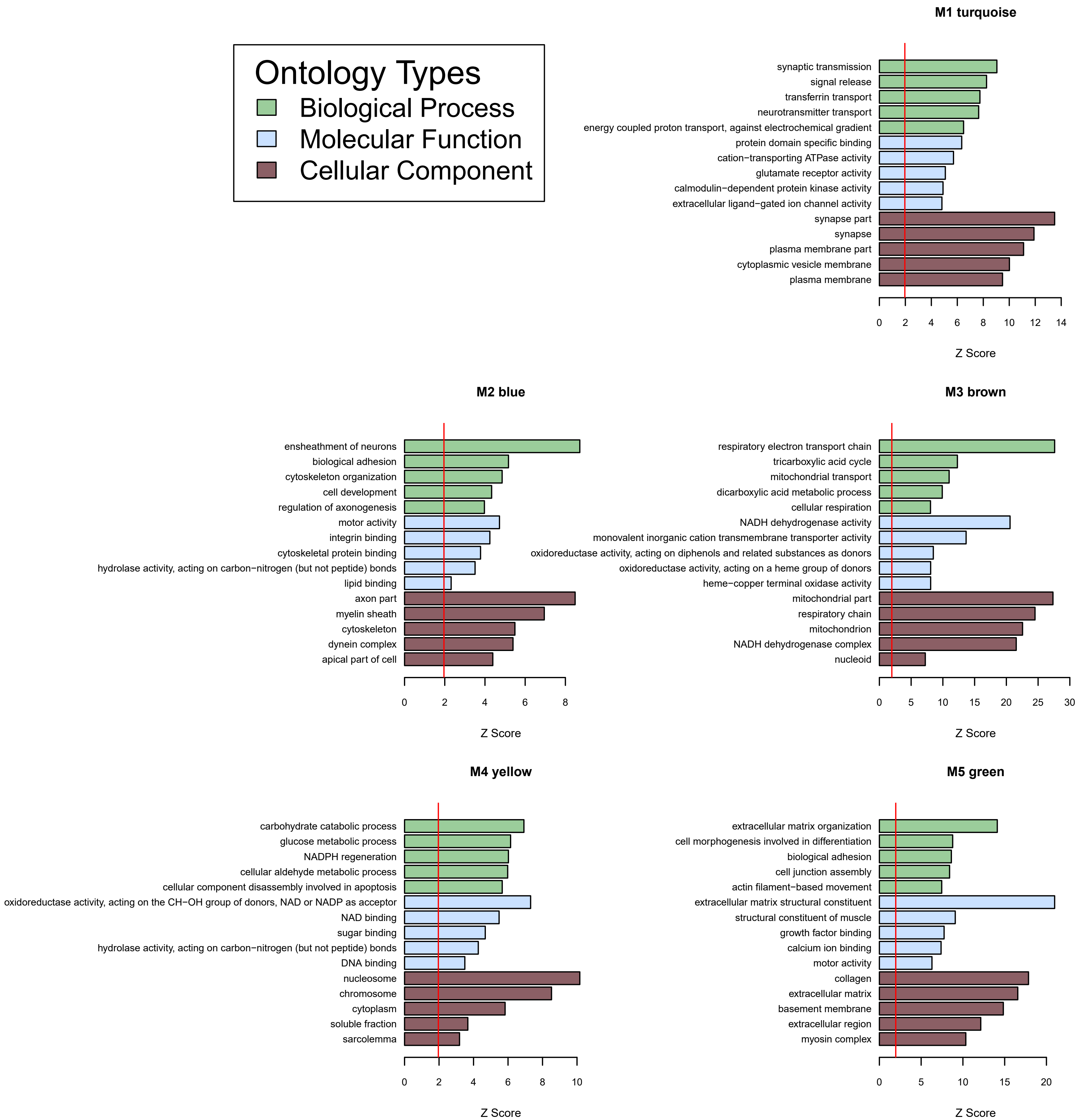
Ontology Types
Biological Process
Molecular Function
Cellular Component
M1 turquoise
synaptic transmission
signal release
transferrin transport
neurotransmitter transport
energy coupled proton transport, against electrochemical gradient
protein domain specific binding
cation−transporting ATPase activity
glutamate receptor activity
calmodulin−dependent protein kinase activity
extracellular ligand−gated ion channel activity
synapse part
synapse
plasma membrane part
cytoplasmic vesicle membrane
plasma membrane
0 2 4 6 8 10 12 14
Z Score
M2 blue
ensheathment of neurons
biological adhesion
cytoskeleton organization
cell development
regulation of axonogenesis
motor activity
integrin binding
cytoskeletal protein binding
hydrolase activity, acting on carbon−nitrogen (but not peptide) bonds
lipid binding
axon part
myelin sheath
cytoskeleton
dynein complex
apical part of cell
0 2 4 6 8
Z Score
M3 brown
respiratory electron transport chain
tricarboxylic acid cycle
mitochondrial transport
dicarboxylic acid metabolic process
cellular respiration
NADH dehydrogenase activity
monovalent inorganic cation transmembrane transporter activity
oxidoreductase activity, acting on diphenols and related substances as donors
oxidoreductase activity, acting on a heme group of donors
heme−copper terminal oxidase activity
mitochondrial part
respiratory chain
mitochondrion
NADH dehydrogenase complex
nucleoid
0 5 10 15 20 25 30
Z Score
M4 yellow
carbohydrate catabolic process
glucose metabolic process
NADPH regeneration
cellular aldehyde metabolic process
cellular component disassembly involved in apoptosis
oxidoreductase activity, acting on the CH−OH group of donors, NAD or NADP as acceptor
NAD binding
sugar binding
hydrolase activity, acting on carbon−nitrogen (but not peptide) bonds
DNA binding
nucleosome
chromosome
cytoplasm
soluble fraction
sarcolemma
0 2 4 6 8 10
Z Score
M5 green
extracellular matrix organization
cell morphogenesis involved in differentiation
biological adhesion
cell junction assembly
actin filament−based movement
extracellular matrix structural constituent
structural constituent of muscle
growth factor binding
calcium ion binding
motor activity
collagen
extracellular matrix
basement membrane
extracellular region
myosin complex
0 5 10 15 20
Z Score

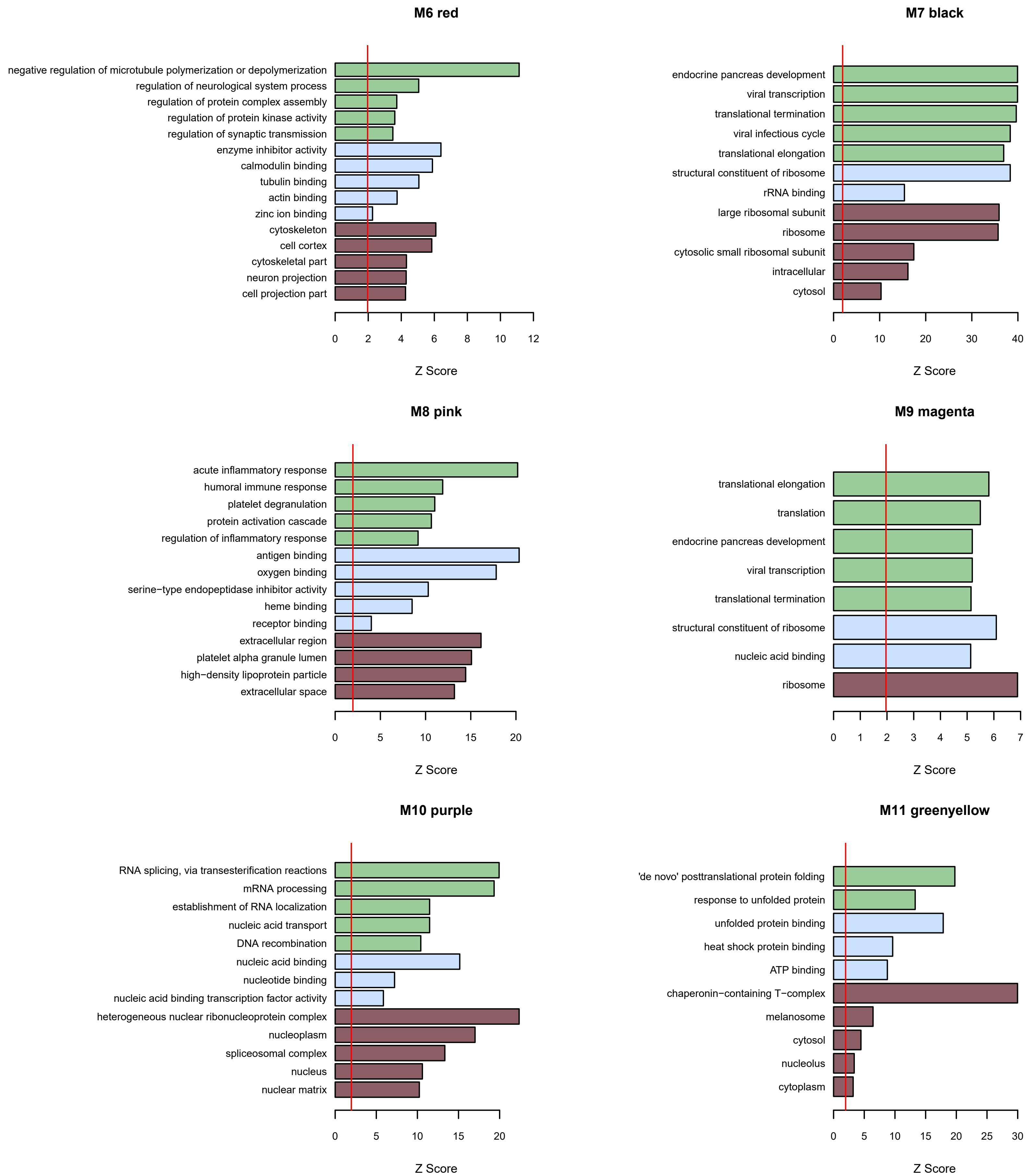
M6 red
negative regulation of microtubule polymerization or depolymerization
regulation of neurological system process
regulation of protein complex assembly
regulation of protein kinase activity
regulation of synaptic transmission
enzyme inhibitor activity
calmodulin binding
tubulin binding
actin binding
zinc ion binding
cytoskeleton
cell cortex
cytoskeletal part
neuron projection
cell projection part
0 2 4 6 8 10 12
Z Score
M7 black
endocrine pancreas development
viral transcription
translational termination
viral infectious cycle
translational elongation
structural constituent of ribosome
rRNA binding
large ribosomal subunit
ribosome
cytosolic small ribosomal subunit
intracellular
cytosol
0 10 20 30 40
Z Score
M8 pink
acute inflammatory response
humoral immune response
platelet degranulation
protein activation cascade
regulation of inflammatory response
antigen binding
oxygen binding
serine−type endopeptidase inhibitor activity
heme binding
receptor binding
extracellular region
platelet alpha granule lumen
high−density lipoprotein particle
extracellular space
0 5 10 15 20
Z Score
M9 magenta
translational elongation
translation
endocrine pancreas development
viral transcription
translational termination
structural constituent of ribosome
nucleic acid binding
ribosome
0 1 2 3 4 5 6 7
Z Score
M10 purple
RNA splicing, via transesterification reactions
mRNA processing
establishment of RNA localization
nucleic acid transport
DNA recombination
nucleic acid binding
nucleotide binding
nucleic acid binding transcription factor activity
heterogeneous nuclear ribonucleoprotein complex
nucleoplasm
spliceosomal complex
nucleus
nuclear matrix
0 5 10 15 20
Z Score
M11 greenyellow
'de novo' posttranslational protein folding
response to unfolded protein
unfolded protein binding
heat shock protein binding
ATP binding
chaperonin−containing T−complex
melanosome
cytosol
nucleolus
cytoplasm
0 5 10 15 20 25 30
Z Score

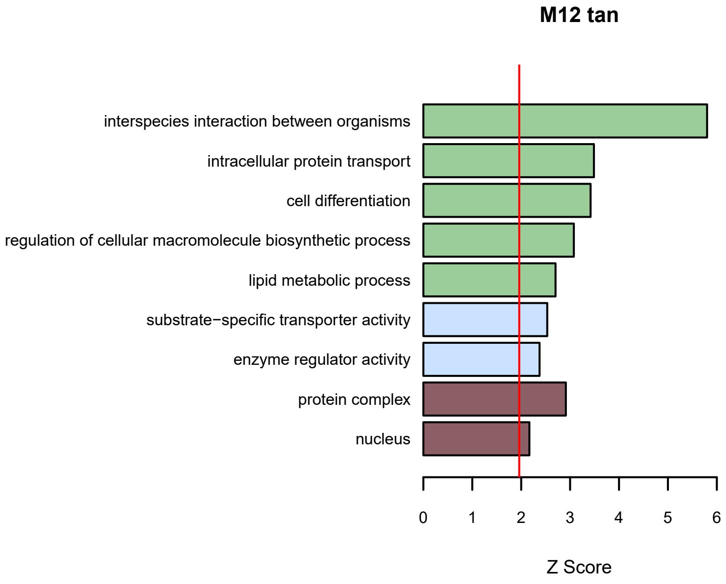

M12 tan
interspecies interaction between organisms
intracellular protein transport
cell differentiation
regulation of cellular macromolecule biosynthetic process
lipid metabolic process
substrate−specific transporter activity
enzyme regulator activity
protein complex
nucleus
0
1
2
3
4
5
6
Z Score


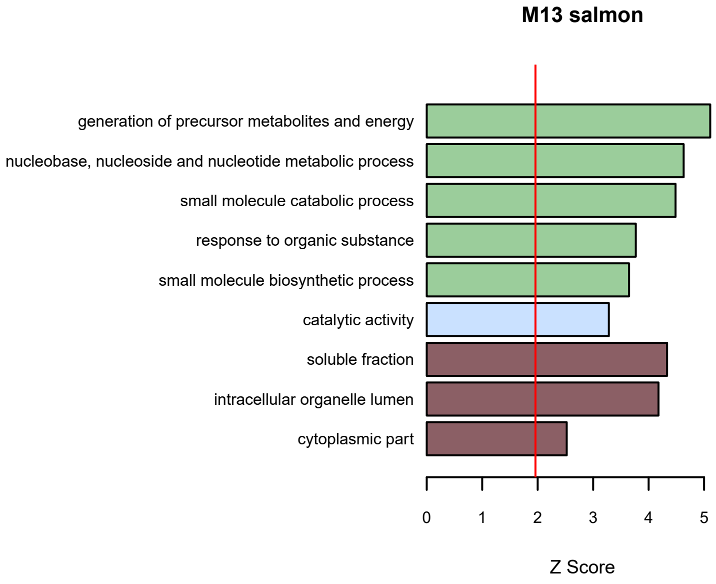

M13 salmon
generation of precursor metabolites and energy
nucleobase, nucleoside and nucleotide metabolic process
small molecule catabolic process
response to organic substance
small molecule biosynthetic process
catalytic activity
soluble fraction
intracellular organelle lumen
cytoplasmic part
0
1
2
3
4
5
Z Score

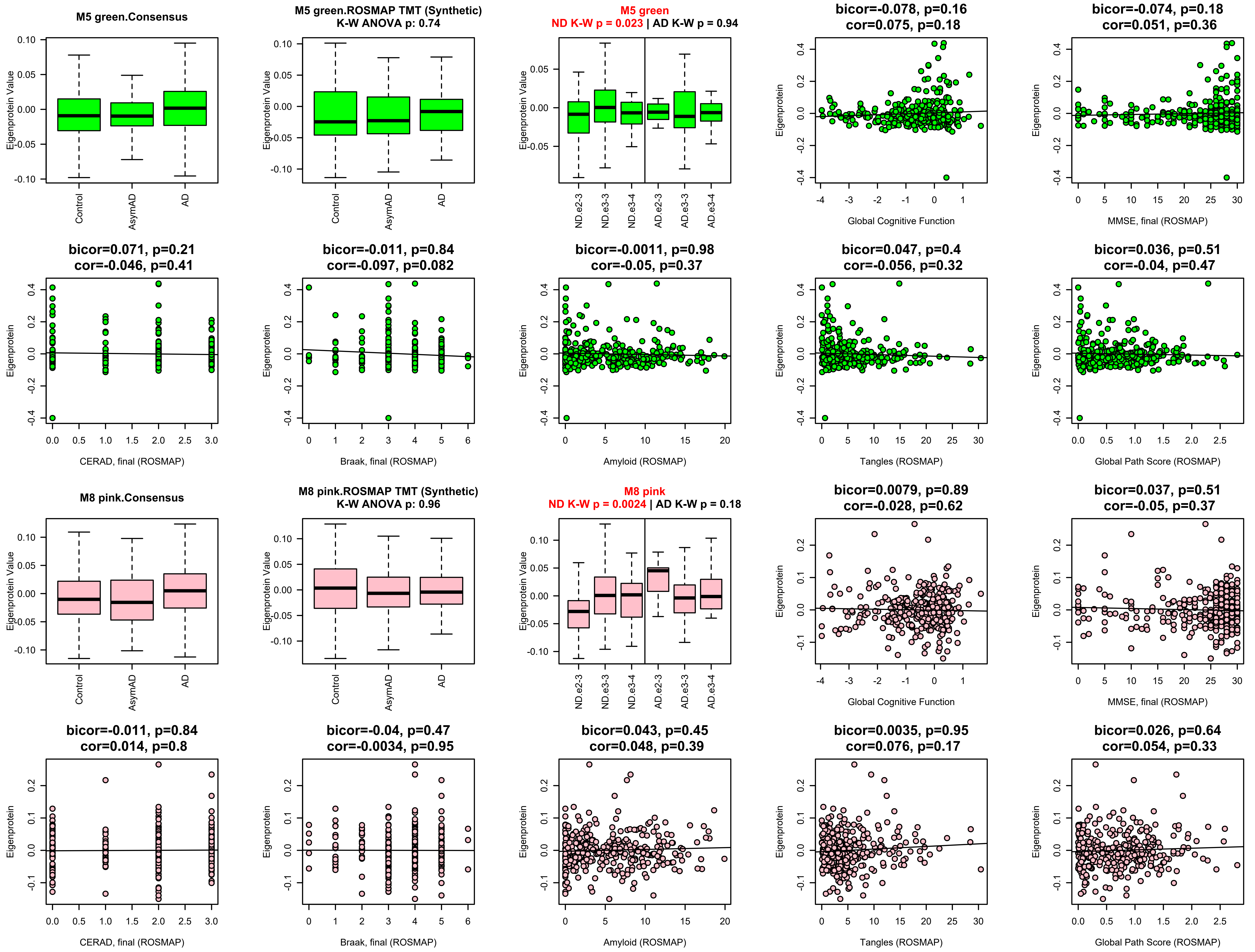
M5 green.Consensus
Eigenprotein Value
Control
AsymAD
AD
M5 green.ROSMAP TMT (Synthetic)
K-W ANOVA p: 0.74
M5 green
ND K-W p = 0.023 | AD K-W p = 0.94
ND.e2-3
ND.e3-3
ND.e3-4
AD.e2-3
AD.e3-3
AD.e3-4
bicor=-0.078, p=0.16
cor=0.075, p=0.18
Eigenprotein
Global Cognitive Function
bicor=-0.074, p=0.18
cor=0.051, p=0.36
MMSE, final (ROSMAP)
bicor=0.071, p=0.21
cor=-0.046, p=0.41
CERAD, final (ROSMAP)
bicor=-0.011, p=0.84
cor=-0.097, p=0.082
Braak, final (ROSMAP)
bicor=-0.0011, p=0.98
cor=-0.05, p=0.37
Amyloid (ROSMAP)
bicor=0.047, p=0.4
cor=-0.056, p=0.32
Tangles (ROSMAP)
bicor=0.036, p=0.51
cor=-0.04, p=0.47
Global Path Score (ROSMAP)
M8 pink.Consensus
M8 pink.ROSMAP TMT (Synthetic)
K-W ANOVA p: 0.96
M8 pink
ND K-W p = 0.0024 | AD K-W p = 0.18
bicor=0.0079, p=0.89
cor=-0.028, p=0.62
bicor=0.037, p=0.51
cor=-0.05, p=0.37
bicor=-0.011, p=0.84
cor=0.014, p=0.8
bicor=-0.04, p=0.47
cor=-0.0034, p=0.95
bicor=0.043, p=0.45
cor=0.048, p=0.39
bicor=0.0035, p=0.95
cor=0.076, p=0.17
bicor=0.026, p=0.64
cor=0.054, p=0.33

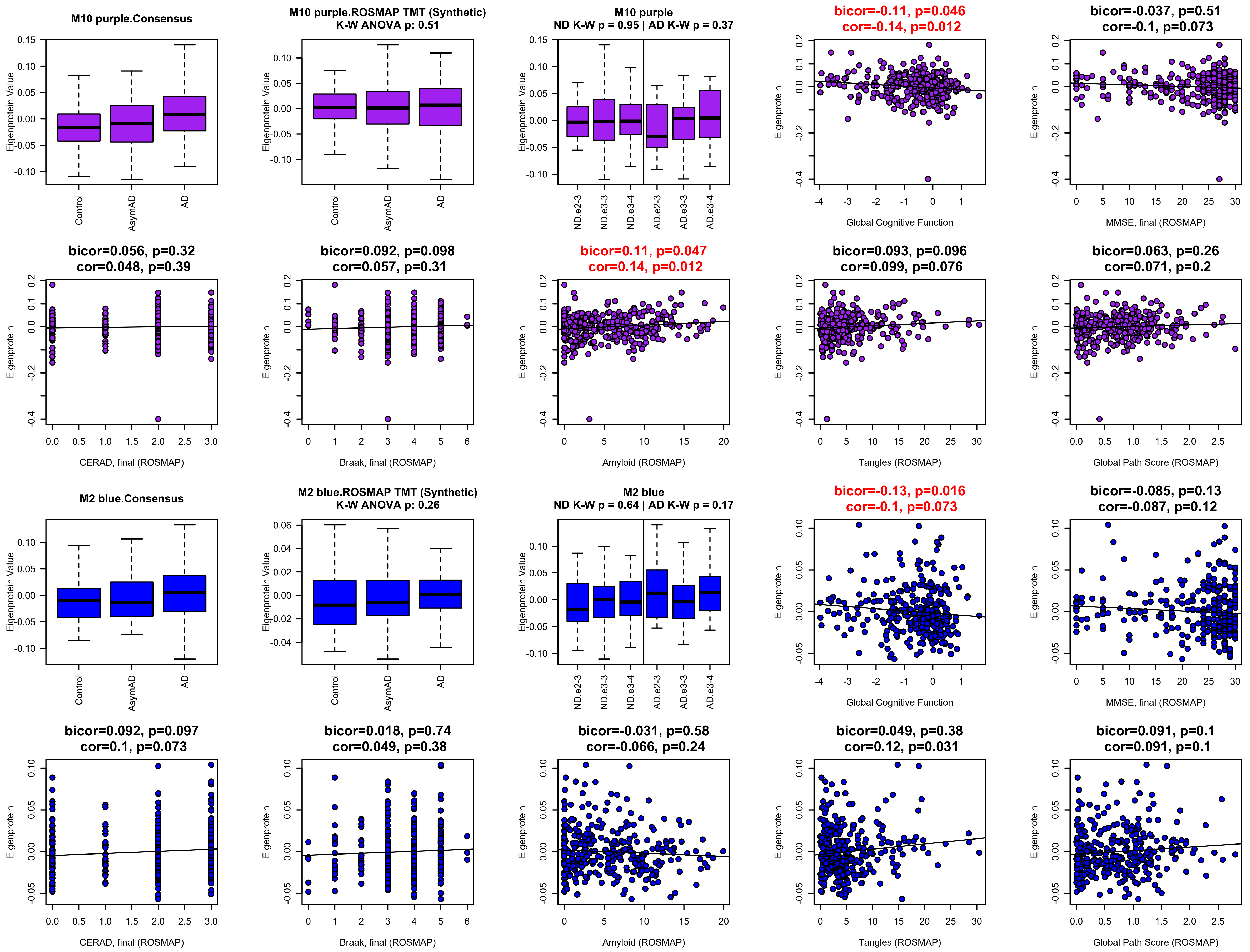
M10 purple.Consensus
Eigenprotein Value
Control
AsymAD
AD
M10 purple.ROSMAP TMT (Synthetic)
K-W ANOVA p: 0.51
M10 purple
ND K-W p = 0.95 | AD K-W p = 0.37
ND.e2-3
ND.e3-3
ND.e3-4
AD.e2-3
AD.e3-3
AD.e3-4
bicor=-0.11, p=0.046
cor=-0.14, p=0.012
Eigenprotein
Global Cognitive Function
bicor=-0.037, p=0.51
cor=-0.1, p=0.073
MMSE, final (ROSMAP)
bicor=0.056, p=0.32
cor=0.048, p=0.39
CERAD, final (ROSMAP)
bicor=0.092, p=0.098
cor=0.057, p=0.31
Braak, final (ROSMAP)
bicor=0.11, p=0.047
cor=0.14, p=0.012
Amyloid (ROSMAP)
bicor=0.093, p=0.096
cor=0.099, p=0.076
Tangles (ROSMAP)
bicor=0.063, p=0.26
cor=0.071, p=0.2
Global Path Score (ROSMAP)
M2 blue.Consensus
M2 blue.ROSMAP TMT (Synthetic)
K-W ANOVA p: 0.26
M2 blue
ND K-W p = 0.64 | AD K-W p = 0.17
bicor=-0.13, p=0.016
cor=-0.1, p=0.073
bicor=-0.085, p=0.13
cor=-0.087, p=0.12
bicor=0.092, p=0.097
cor=0.1, p=0.073
bicor=0.018, p=0.74
cor=0.049, p=0.38
bicor=-0.031, p=0.58
cor=-0.066, p=0.24
bicor=0.049, p=0.38
cor=0.12, p=0.031
bicor=0.091, p=0.1
cor=0.091, p=0.1

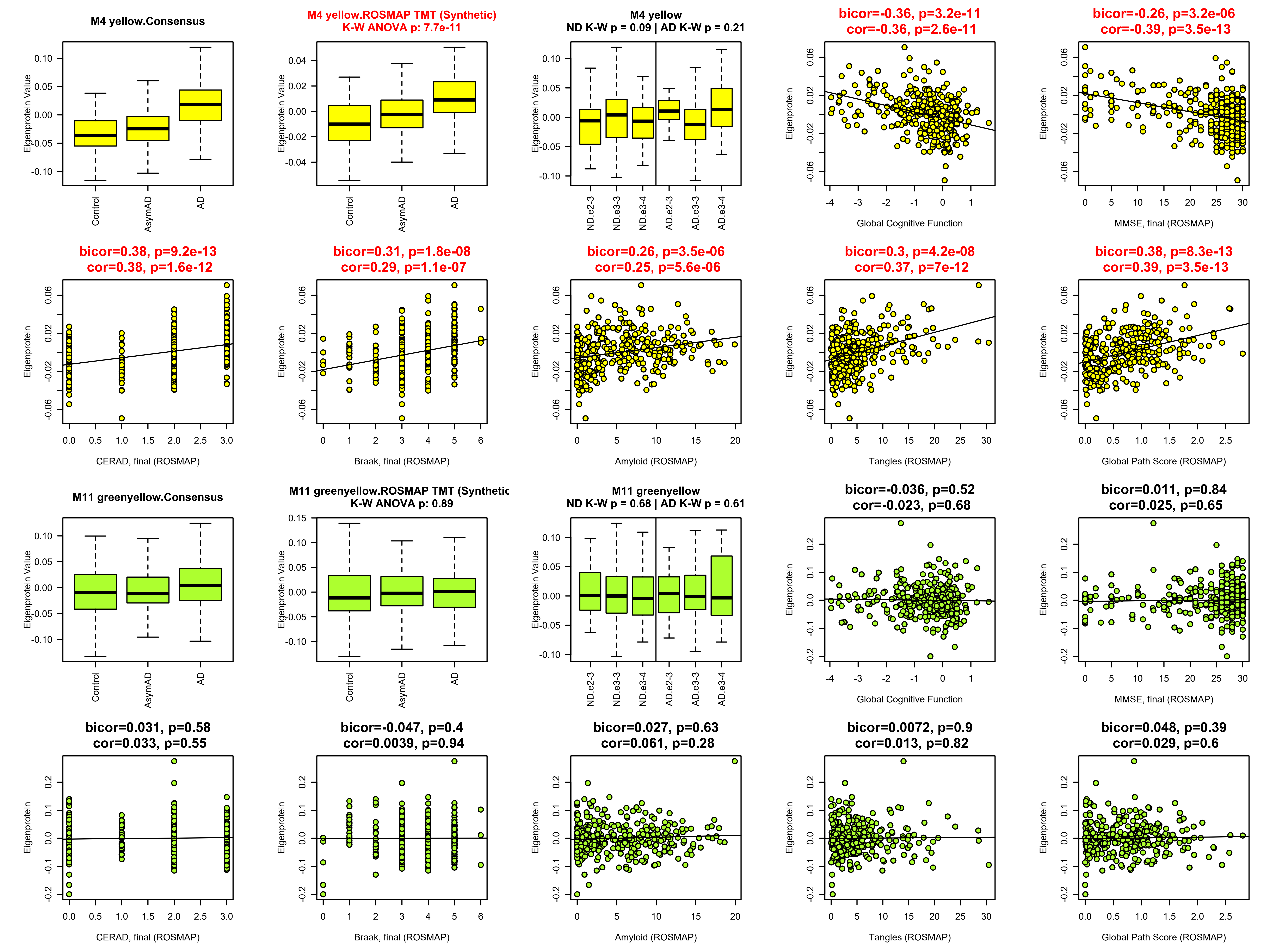
M4 yellow.Consensus
Eigenprotein Value
Control
AsymAD
AD
M4 yellow.ROSMAP TMT (Synthetic)
K-W ANOVA p: 7.7e-11
M4 yellow
ND K-W p = 0.09 | AD K-W p = 0.21
ND.e2-3
ND.e3-3
ND.e3-4
AD.e2-3
AD.e3-3
AD.e3-4
bicor=-0.36, p=3.2e-11
cor=-0.36, p=2.6e-11
Eigenprotein
Global Cognitive Function
bicor=-0.26, p=3.2e-06
cor=-0.39, p=3.5e-13
MMSE, final (ROSMAP)
bicor=0.38, p=9.2e-13
cor=0.38, p=1.6e-12
CERAD, final (ROSMAP)
bicor=0.31, p=1.8e-08
cor=0.29, p=1.1e-07
Braak, final (ROSMAP)
bicor=0.26, p=3.5e-06
cor=0.25, p=5.6e-06
Amyloid (ROSMAP)
bicor=0.3, p=4.2e-08
cor=0.37, p=7e-12
Tangles (ROSMAP)
bicor=0.38, p=8.3e-13
cor=0.39, p=3.5e-13
Global Path Score (ROSMAP)
M11 greenyellow.Consensus
M11 greenyellow.ROSMAP TMT (Synthetic
K-W ANOVA p: 0.89
M11 greenyellow
ND K-W p = 0.68 | AD K-W p = 0.61
bicor=-0.036, p=0.52
cor=-0.023, p=0.68
bicor=0.011, p=0.84
cor=0.025, p=0.65
bicor=0.031, p=0.58
cor=0.033, p=0.55
bicor=-0.047, p=0.4
cor=0.0039, p=0.94
bicor=0.027, p=0.63
cor=0.061, p=0.28
bicor=0.0072, p=0.9
cor=0.013, p=0.82
bicor=0.048, p=0.39
cor=0.029, p=0.6

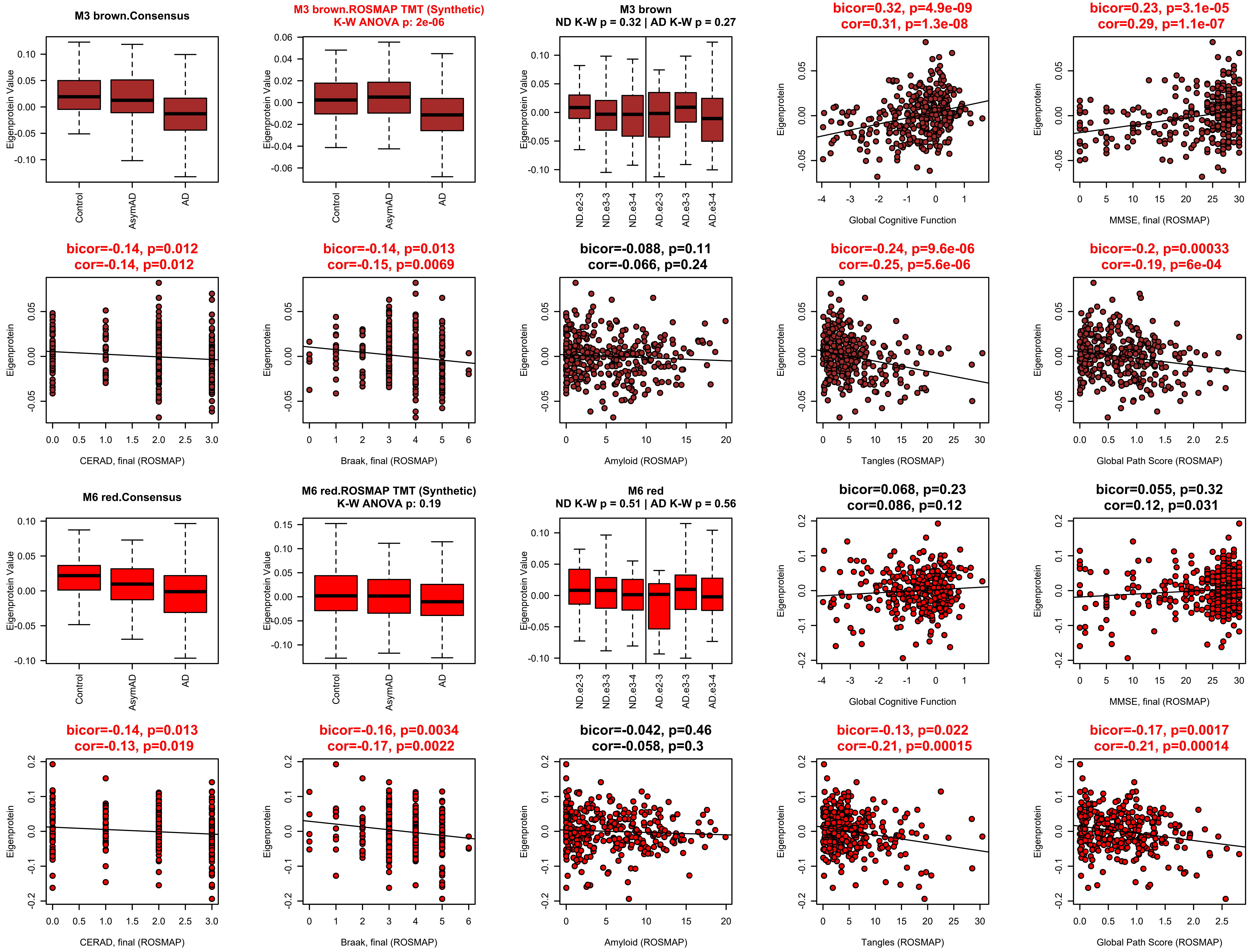

M3 brown.Consensus
M3 brown.ROSMAP TMT (Synthetic)
K-W ANOVA p: 2e-06
M3 brown
ND K-W p = 0.32 | AD K-W p = 0.27
bicor=0.32, p=4.9e-09
cor=0.31, p=1.3e-08
bicor=0.23, p=3.1e-05
cor=0.29, p=1.1e-07
Eigenprotein Value
Eigenprotein
Control
AsymAD
AD
ND.e2-3
ND.e3-3
ND.e3-4
AD.e2-3
AD.e3-3
AD.e3-4
Global Cognitive Function
MMSE, final (ROSMAP)
bicor=-0.14, p=0.012
cor=-0.14, p=0.012
bicor=-0.14, p=0.013
cor=-0.15, p=0.0069
bicor=-0.088, p=0.11
cor=-0.066, p=0.24
bicor=-0.24, p=9.6e-06
cor=-0.25, p=5.6e-06
bicor=-0.2, p=0.00033
cor=-0.19, p=6e-04
CERAD, final (ROSMAP)
Braak, final (ROSMAP)
Amyloid (ROSMAP)
Tangles (ROSMAP)
Global Path Score (ROSMAP)
M6 red.Consensus
M6 red.ROSMAP TMT (Synthetic)
K-W ANOVA p: 0.19
M6 red
ND K-W p = 0.51 | AD K-W p = 0.56
bicor=0.068, p=0.23
cor=0.086, p=0.12
bicor=0.055, p=0.32
cor=0.12, p=0.031
bicor=-0.14, p=0.013
cor=-0.13, p=0.019
bicor=-0.16, p=0.0034
cor=-0.17, p=0.0022
bicor=-0.042, p=0.46
cor=-0.058, p=0.3
bicor=-0.13, p=0.022
cor=-0.21, p=0.00015
bicor=-0.17, p=0.0017
cor=-0.21, p=0.00014

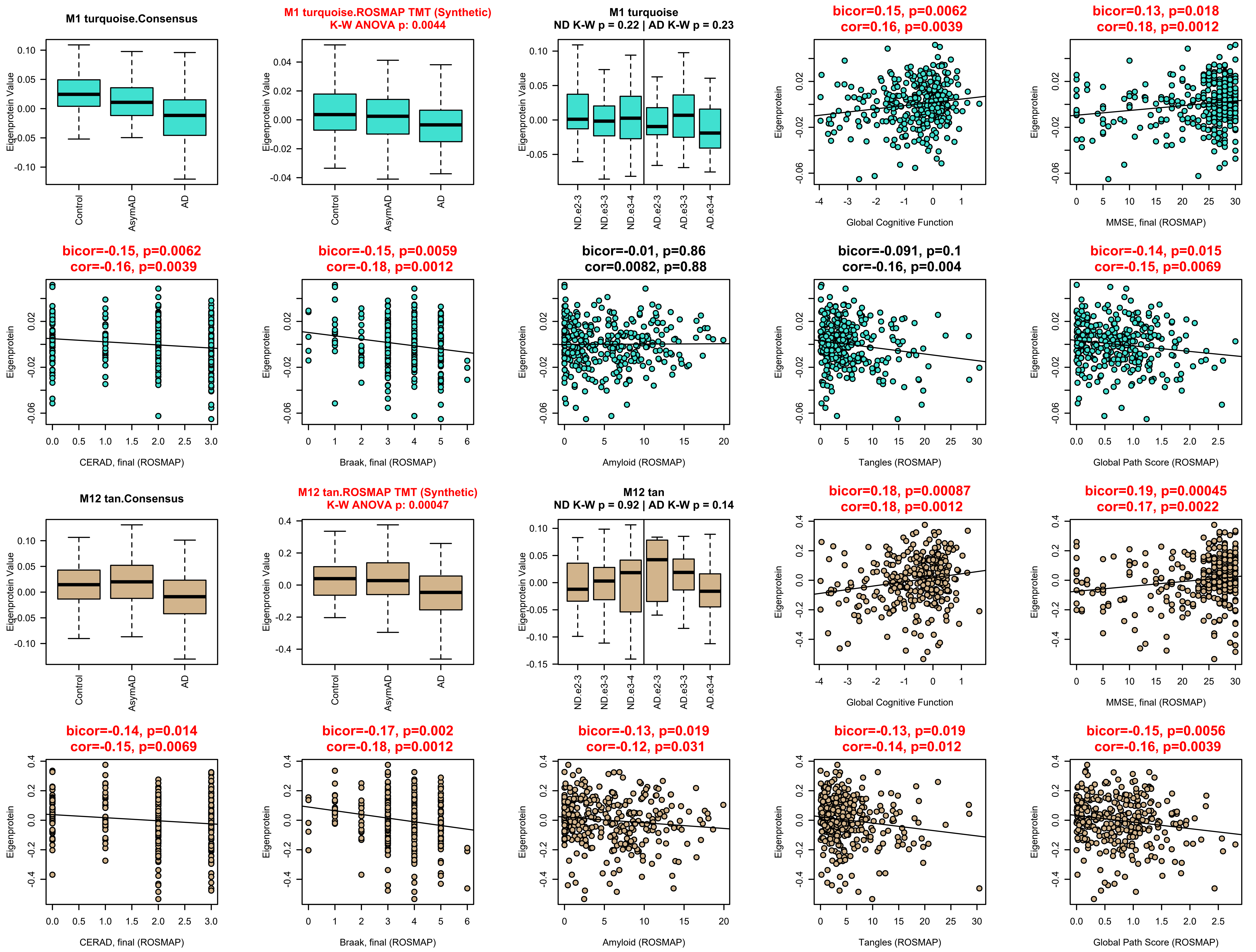
M1 turquoise.Consensus
Eigenprotein Value
Control
AsymAD
AD
M1 turquoise.ROSMAP TMT (Synthetic)
K-W ANOVA p: 0.0044
Eigenprotein Value
Control
AsymAD
AD
M1 turquoise
ND K-W p = 0.22 | AD K-W p = 0.23
Eigenprotein Value
ND.e2-3
ND.e3-3
ND.e3-4
AD.e2-3
AD.e3-3
AD.e3-4
bicor=0.15, p=0.0062
cor=0.16, p=0.0039
Eigenprotein
Global Cognitive Function
bicor=0.13, p=0.018
cor=0.18, p=0.0012
Eigenprotein
MMSE, final (ROSMAP)
bicor=-0.15, p=0.0062
cor=-0.16, p=0.0039
Eigenprotein
CERAD, final (ROSMAP)
bicor=-0.15, p=0.0059
cor=-0.18, p=0.0012
Eigenprotein
Braak, final (ROSMAP)
bicor=-0.01, p=0.86
cor=0.0082, p=0.88
Eigenprotein
Amyloid (ROSMAP)
bicor=-0.091, p=0.1
cor=-0.16, p=0.004
Eigenprotein
Tangles (ROSMAP)
bicor=-0.14, p=0.015
cor=-0.15, p=0.0069
Eigenprotein
Global Path Score (ROSMAP)
M12 tan.Consensus
Eigenprotein Value
Control
AsymAD
AD
M12 tan.ROSMAP TMT (Synthetic)
K-W ANOVA p: 0.00047
Eigenprotein Value
Control
AsymAD
AD
M12 tan
ND K-W p = 0.92 | AD K-W p = 0.14
Eigenprotein Value
ND.e2-3
ND.e3-3
ND.e3-4
AD.e2-3
AD.e3-3
AD.e3-4
bicor=0.18, p=0.00087
cor=0.18, p=0.0012
Eigenprotein
Global Cognitive Function
bicor=0.19, p=0.00045
cor=0.17, p=0.0022
Eigenprotein
MMSE, final (ROSMAP)
bicor=-0.14, p=0.014
cor=-0.15, p=0.0069
Eigenprotein
CERAD, final (ROSMAP)
bicor=-0.17, p=0.002
cor=-0.18, p=0.0012
Eigenprotein
Braak, final (ROSMAP)
bicor=-0.13, p=0.019
cor=-0.12, p=0.031
Eigenprotein
Amyloid (ROSMAP)
bicor=-0.13, p=0.019
cor=-0.14, p=0.012
Eigenprotein
Tangles (ROSMAP)
bicor=-0.15, p=0.0056
cor=-0.16, p=0.0039
Eigenprotein
Global Path Score (ROSMAP)

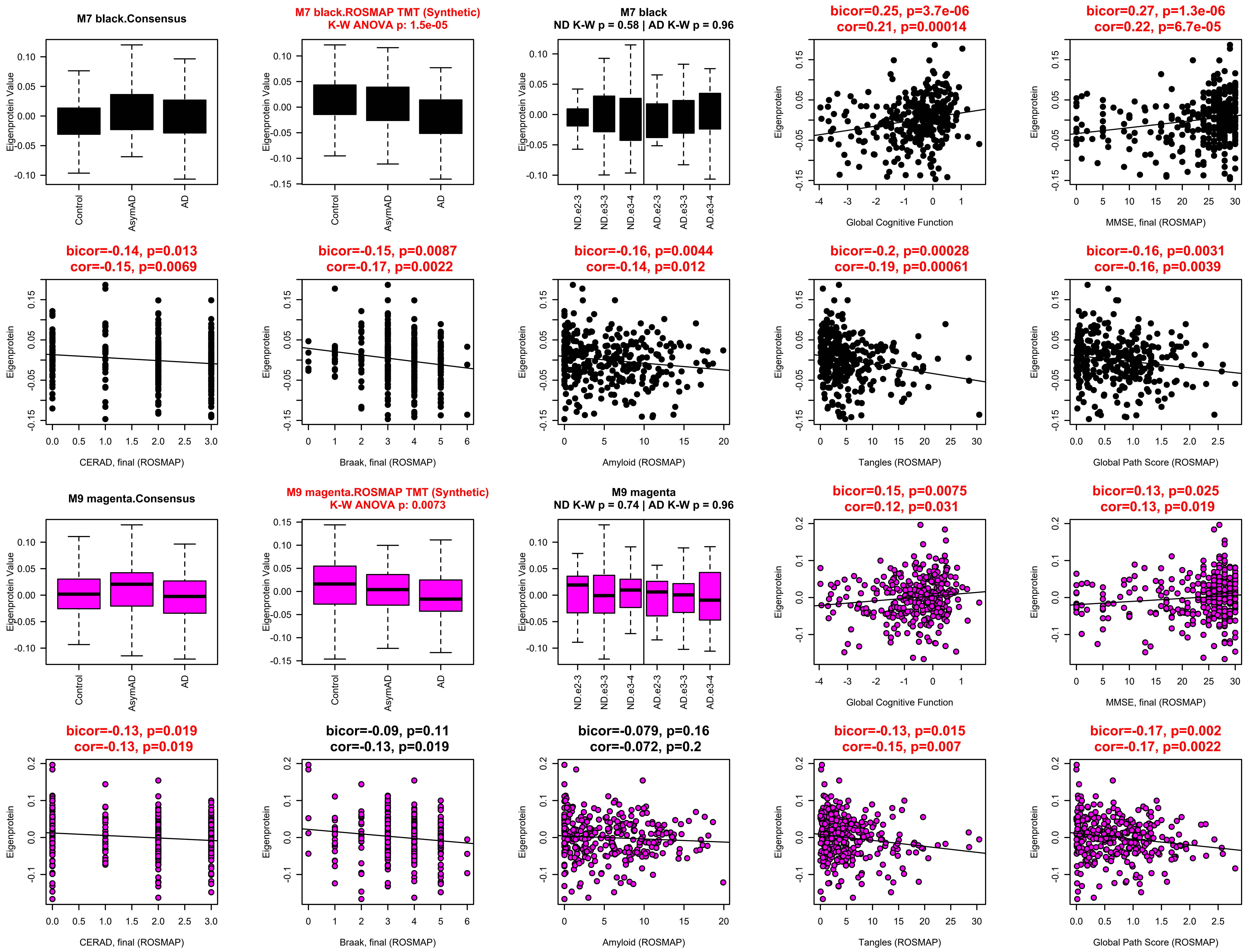

M7 black.Consensus
Eigenprotein Value
Control
AsymAD
AD
M7 black.ROSMAP TMT (Synthetic)
K-W ANOVA p: 1.5e-05
M7 black
ND K-W p = 0.58 | AD K-W p = 0.96
ND.e2-3
ND.e3-3
ND.e3-4
AD.e2-3
AD.e3-3
AD.e3-4
bicor=0.25, p=3.7e-06
cor=0.21, p=0.00014
Eigenprotein
Global Cognitive Function
bicor=0.27, p=1.3e-06
cor=0.22, p=6.7e-05
MMSE, final (ROSMAP)
bicor=-0.14, p=0.013
cor=-0.15, p=0.0069
CERAD, final (ROSMAP)
bicor=-0.15, p=0.0087
cor=-0.17, p=0.0022
Braak, final (ROSMAP)
bicor=-0.16, p=0.0044
cor=-0.14, p=0.012
Amyloid (ROSMAP)
bicor=-0.2, p=0.00028
cor=-0.19, p=0.00061
Tangles (ROSMAP)
bicor=-0.16, p=0.0031
cor=-0.16, p=0.0039
Global Path Score (ROSMAP)
M9 magenta.Consensus
M9 magenta.ROSMAP TMT (Synthetic)
K-W ANOVA p: 0.0073
M9 magenta
ND K-W p = 0.74 | AD K-W p = 0.96
bicor=0.15, p=0.0075
cor=0.12, p=0.031
bicor=0.13, p=0.025
cor=0.13, p=0.019
bicor=-0.13, p=0.019
cor=-0.13, p=0.019
bicor=-0.09, p=0.11
cor=-0.13, p=0.019
bicor=-0.079, p=0.16
cor=-0.072, p=0.2
bicor=-0.13, p=0.015
cor=-0.15, p=0.007
bicor=-0.17, p=0.002
cor=-0.17, p=0.0022

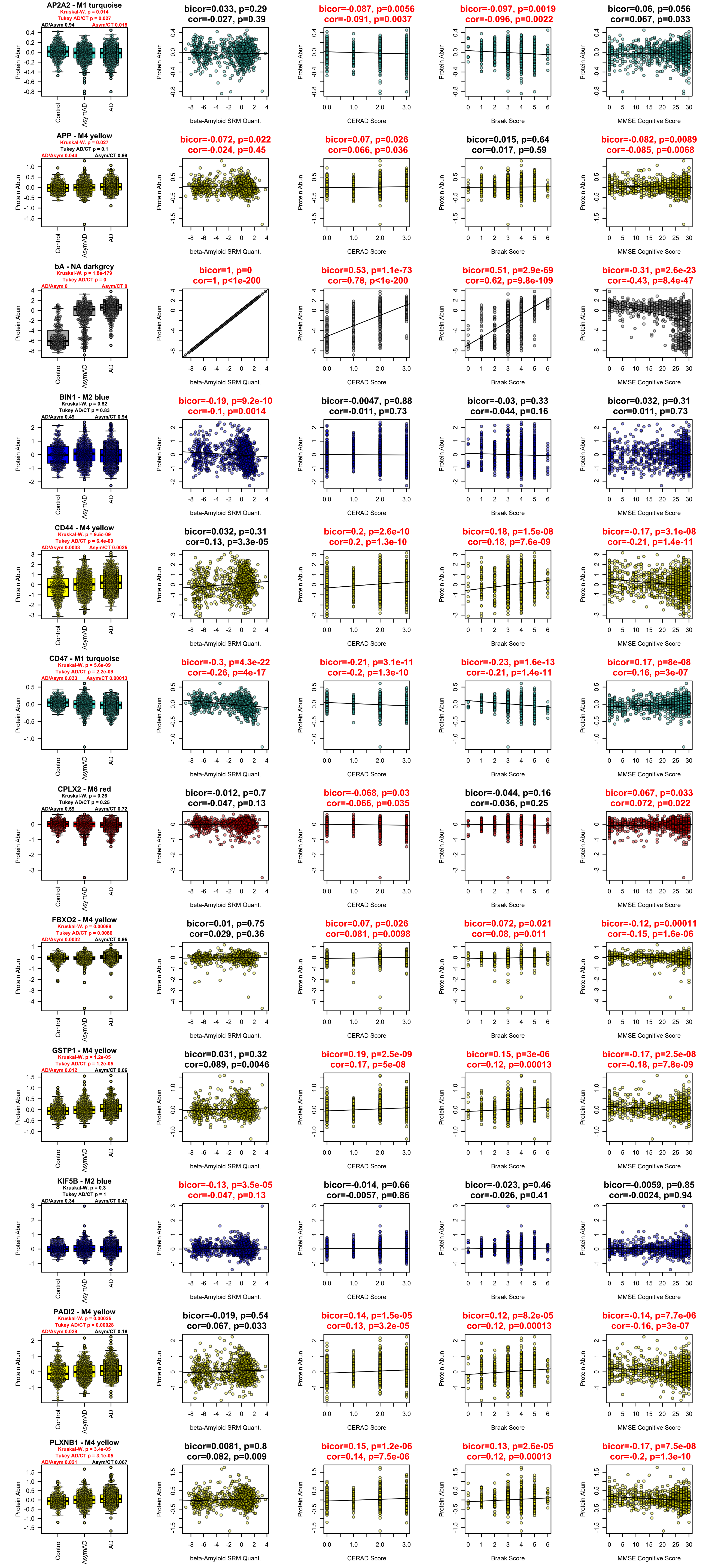

AP2A2 - M1 turquoise
Kruskal-W. p = 0.014
Tukey AD/CT p = 0.027
AD/Asym 0.94
Asym/CT 0.015
bicor=0.033, p=0.29
cor=-0.027, p=0.39
bicor=-0.087, p=0.0056
cor=-0.091, p=0.0037
bicor=-0.097, p=0.0019
cor=-0.096, p=0.0022
bicor=0.06, p=0.056
cor=0.067, p=0.033
APP - M4 yellow
Kruskal-W. p = 0.027
Tukey AD/CT p = 0.1
AD/Asym 0.044
Asym/CT 0.99
bicor=-0.072, p=0.022
cor=-0.024, p=0.45
bicor=0.07, p=0.026
cor=0.066, p=0.036
bicor=0.015, p=0.64
cor=0.017, p=0.59
bicor=-0.082, p=0.0089
cor=-0.085, p=0.0068
bA - NA darkgrey
Kruskal-W. p = 1.8e-179
Tukey AD/CT p = 0
AD/Asym 0
Asym/CT 0
bicor=1, p=0
cor=1, p<1e-200
bicor=0.53, p=1.1e-73
cor=0.78, p<1e-200
bicor=0.51, p=2.9e-69
cor=0.62, p=9.8e-109
bicor=-0.31, p=2.6e-23
cor=-0.43, p=8.4e-47
BIN1 - M2 blue
Kruskal-W. p = 0.52
Tukey AD/CT p = 0.83
AD/Asym 0.49
Asym/CT 0.94
bicor=-0.19, p=9.2e-10
cor=-0.1, p=0.0014
bicor=-0.0047, p=0.88
cor=-0.011, p=0.73
bicor=-0.03, p=0.33
cor=-0.044, p=0.16
bicor=0.032, p=0.31
cor=0.011, p=0.73
CD44 - M4 yellow
Kruskal-W. p = 9.5e-09
Tukey AD/CT p = 6.4e-09
AD/Asym 0.0033
Asym/CT 0.0025
bicor=0.032, p=0.31
cor=0.13, p=3.3e-05
bicor=0.2, p=2.6e-10
cor=0.2, p=1.3e-10
bicor=0.18, p=1.5e-08
cor=0.18, p=7.6e-09
bicor=-0.17, p=3.1e-08
cor=-0.21, p=1.4e-11
CD47 - M1 turquoise
Kruskal-W. p = 5.6e-09
Tukey AD/CT p = 2.2e-09
AD/Asym 0.033
Asym/CT 0.00013
bicor=-0.3, p=4.3e-22
cor=-0.26, p=4e-17
bicor=-0.21, p=3.1e-11
cor=-0.2, p=1.3e-10
bicor=-0.23, p=1.6e-13
cor=-0.21, p=1.4e-11
bicor=0.17, p=8e-08
cor=0.16, p=3e-07
CPLX2 - M6 red
Kruskal-W. p = 0.26
Tukey AD/CT p = 0.25
AD/Asym 0.59
Asym/CT 0.72
bicor=-0.012, p=0.7
cor=-0.047, p=0.13
bicor=-0.068, p=0.03
cor=-0.066, p=0.035
bicor=-0.044, p=0.16
cor=-0.036, p=0.25
bicor=0.067, p=0.033
cor=0.072, p=0.022
FBXO2 - M4 yellow
Kruskal-W. p = 0.00088
Tukey AD/CT p = 0.0086
AD/Asym 0.0032
Asym/CT 0.95
bicor=0.01, p=0.75
cor=0.029, p=0.36
bicor=0.07, p=0.026
cor=0.081, p=0.0098
bicor=0.072, p=0.021
cor=0.08, p=0.011
bicor=-0.12, p=0.00011
cor=-0.15, p=1.6e-06
GSTP1 - M4 yellow
Kruskal-W. p = 1.2e-05
Tukey AD/CT p = 1.2e-05
AD/Asym 0.012
Asym/CT 0.06
bicor=0.031, p=0.32
cor=0.089, p=0.0046
bicor=0.19, p=2.5e-09
cor=0.17, p=5e-08
bicor=0.15, p=3e-06
cor=0.12, p=0.00013
bicor=-0.17, p=2.5e-08
cor=-0.18, p=7.8e-09
KIF5B - M2 blue
Kruskal-W. p = 0.3
Tukey AD/CT p = 1
AD/Asym 0.34
Asym/CT 0.47
bicor=-0.13, p=3.5e-05
cor=-0.047, p=0.13
bicor=-0.014, p=0.66
cor=-0.0057, p=0.86
bicor=-0.023, p=0.46
cor=-0.026, p=0.41
bicor=-0.0059, p=0.85
cor=-0.0024, p=0.94
PADI2 - M4 yellow
Kruskal-W. p = 0.00025
Tukey AD/CT p = 0.00028
AD/Asym 0.029
Asym/CT 0.16
bicor=-0.019, p=0.54
cor=0.067, p=0.033
bicor=0.14, p=1.5e-05
cor=0.13, p=3.2e-05
bicor=0.12, p=8.2e-05
cor=0.12, p=0.00013
bicor=-0.14, p=7.7e-06
cor=-0.16, p=3e-07
PLXNB1 - M4 yellow
Kruskal-W. p = 3.4e-05
Tukey AD/CT p = 3.1e-05
AD/Asym 0.021
Asym/CT 0.067
bicor=0.0081, p=0.8
cor=0.082, p=0.009
bicor=0.15, p=1.2e-06
cor=0.14, p=7.5e-06
bicor=0.13, p=2.6e-05
cor=0.12, p=0.00013
bicor=-0.17, p=7.5e-08
cor=-0.2, p=1.3e-10
Protein Abun
Control
AsymAD
AD
beta-Amyloid SRM Quant.
CERAD Score
Braak Score
MMSE Cognitive Score

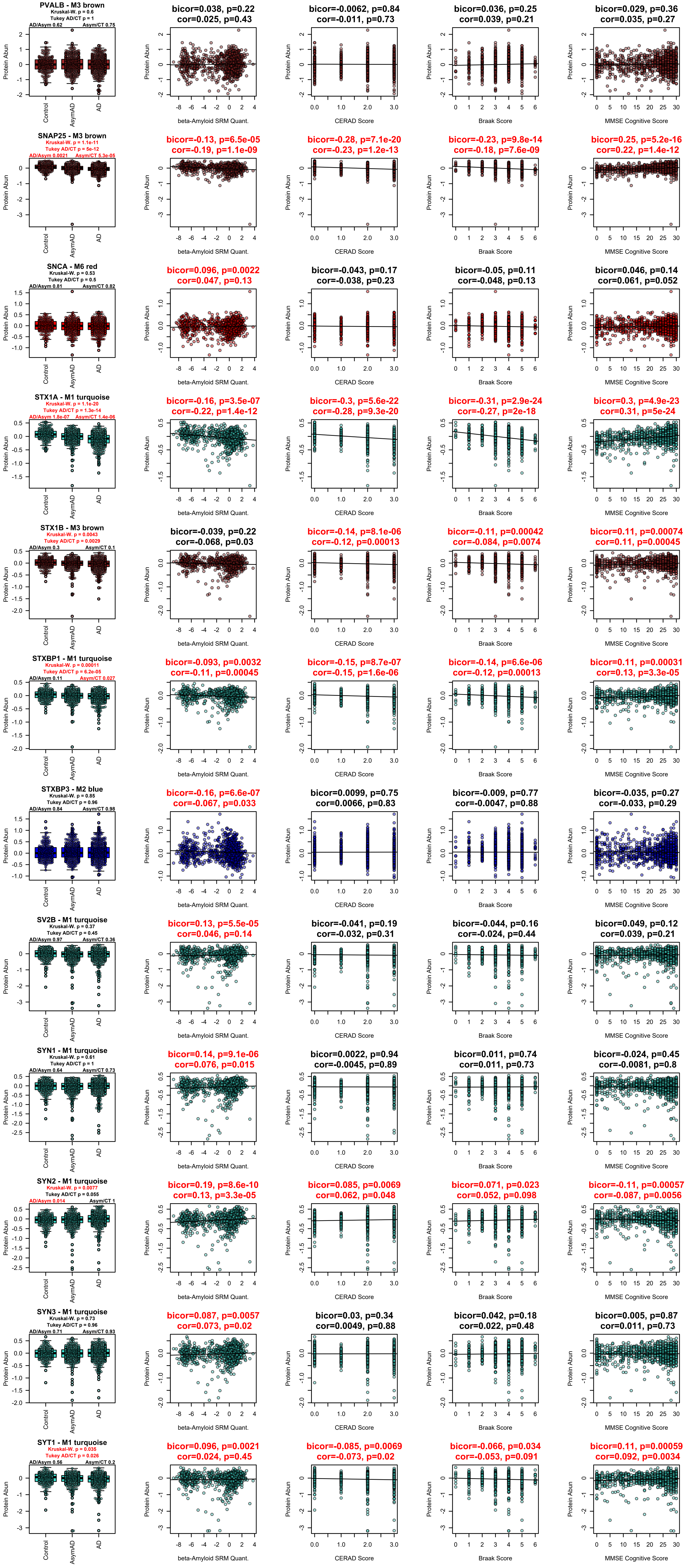

PVALB - M3 brown
Kruskal-W. p = 0.6
Tukey AD/CT p = 1
AD/Asym 0.62
Asym/CT 0.75
bicor=0.038, p=0.22
cor=0.025, p=0.43
bicor=-0.0062, p=0.84
cor=-0.011, p=0.73
bicor=0.036, p=0.25
cor=0.039, p=0.21
bicor=0.029, p=0.36
cor=0.035, p=0.27
SNAP25 - M3 brown
Kruskal-W. p = 1.1e-11
Tukey AD/CT p = 5e-12
AD/Asym 0.0021
Asym/CT 5.3e-05
bicor=-0.13, p=6.5e-05
cor=-0.19, p=1.1e-09
bicor=-0.28, p=7.1e-20
cor=-0.23, p=1.2e-13
bicor=-0.23, p=9.8e-14
cor=-0.18, p=7.6e-09
bicor=0.25, p=5.2e-16
cor=0.22, p=1.4e-12
SNCA - M6 red
Kruskal-W. p = 0.53
Tukey AD/CT p = 0.5
AD/Asym 0.81
Asym/CT 0.82
bicor=0.096, p=0.0022
cor=0.047, p=0.13
bicor=-0.043, p=0.17
cor=-0.038, p=0.23
bicor=-0.05, p=0.11
cor=-0.048, p=0.13
bicor=0.046, p=0.14
cor=0.061, p=0.052
STX1A - M1 turquoise
Kruskal-W. p = 1.1e-20
Tukey AD/CT p = 1.3e-14
AD/Asym 1.8e-07
Asym/CT 1.4e-06
bicor=-0.16, p=3.5e-07
cor=-0.22, p=1.4e-12
bicor=-0.3, p=5.6e-22
cor=-0.28, p=9.3e-20
bicor=-0.31, p=2.9e-24
cor=-0.27, p=2e-18
bicor=0.3, p=4.9e-23
cor=0.31, p=5e-24
STX1B - M3 brown
Kruskal-W. p = 0.0043
Tukey AD/CT p = 0.0029
AD/Asym 0.3
Asym/CT 0.1
bicor=-0.039, p=0.22
cor=-0.068, p=0.03
bicor=-0.14, p=8.1e-06
cor=-0.12, p=0.00013
bicor=-0.11, p=0.00042
cor=-0.084, p=0.0074
bicor=0.11, p=0.00074
cor=0.11, p=0.00045
STXBP1 - M1 turquoise
Kruskal-W. p = 0.00011
Tukey AD/CT p = 6.2e-05
AD/Asym 0.11
Asym/CT 0.027
bicor=-0.093, p=0.0032
cor=-0.11, p=0.00045
bicor=-0.15, p=8.7e-07
cor=-0.15, p=1.6e-06
bicor=-0.14, p=6.6e-06
cor=-0.12, p=0.00013
bicor=0.11, p=0.00031
cor=0.13, p=3.3e-05
STXBP3 - M2 blue
Kruskal-W. p = 0.85
Tukey AD/CT p = 0.96
AD/Asym 0.84
Asym/CT 0.98
bicor=-0.16, p=6.6e-07
cor=-0.067, p=0.033
bicor=0.0099, p=0.75
cor=0.0066, p=0.83
bicor=-0.009, p=0.77
cor=-0.0047, p=0.88
bicor=-0.035, p=0.27
cor=-0.033, p=0.29
SV2B - M1 turquoise
Kruskal-W. p = 0.37
Tukey AD/CT p = 0.45
AD/Asym 0.97
Asym/CT 0.36
bicor=0.13, p=5.5e-05
cor=0.046, p=0.14
bicor=-0.041, p=0.19
cor=-0.032, p=0.31
bicor=-0.044, p=0.16
cor=-0.024, p=0.44
bicor=0.049, p=0.12
cor=0.039, p=0.21
SYN1 - M1 turquoise
Kruskal-W. p = 0.61
Tukey AD/CT p = 1
AD/Asym 0.64
Asym/CT 0.73
bicor=0.14, p=9.1e-06
cor=0.076, p=0.015
bicor=0.0022, p=0.94
cor=-0.0045, p=0.89
bicor=0.011, p=0.74
cor=0.011, p=0.73
bicor=-0.024, p=0.45
cor=-0.0081, p=0.8
SYN2 - M1 turquoise
Kruskal-W. p = 0.0077
Tukey AD/CT p = 0.055
AD/Asym 0.014
Asym/CT 1
bicor=0.19, p=8.6e-10
cor=0.13, p=3.3e-05
bicor=0.085, p=0.0069
cor=0.062, p=0.048
bicor=0.071, p=0.023
cor=0.052, p=0.098
bicor=-0.11, p=0.00057
cor=-0.087, p=0.0056
SYN3 - M1 turquoise
Kruskal-W. p = 0.73
Tukey AD/CT p = 0.96
AD/Asym 0.71
Asym/CT 0.93
bicor=0.087, p=0.0057
cor=0.073, p=0.02
bicor=0.03, p=0.34
cor=0.0049, p=0.88
bicor=0.042, p=0.18
cor=0.022, p=0.48
bicor=0.005, p=0.87
cor=0.011, p=0.73
SYT1 - M1 turquoise
Kruskal-W. p = 0.035
Tukey AD/CT p = 0.026
AD/Asym 0.56
Asym/CT 0.2
bicor=0.096, p=0.0021
cor=0.024, p=0.45
bicor=-0.085, p=0.0069
cor=-0.073, p=0.02
bicor=-0.066, p=0.034
cor=-0.053, p=0.091
bicor=0.11, p=0.00059
cor=0.092, p=0.0034
Protein Abun
Control
AsymAD
AD
beta-Amyloid SRM Quant.
CERAD Score
Braak Score
MMSE Cognitive Score

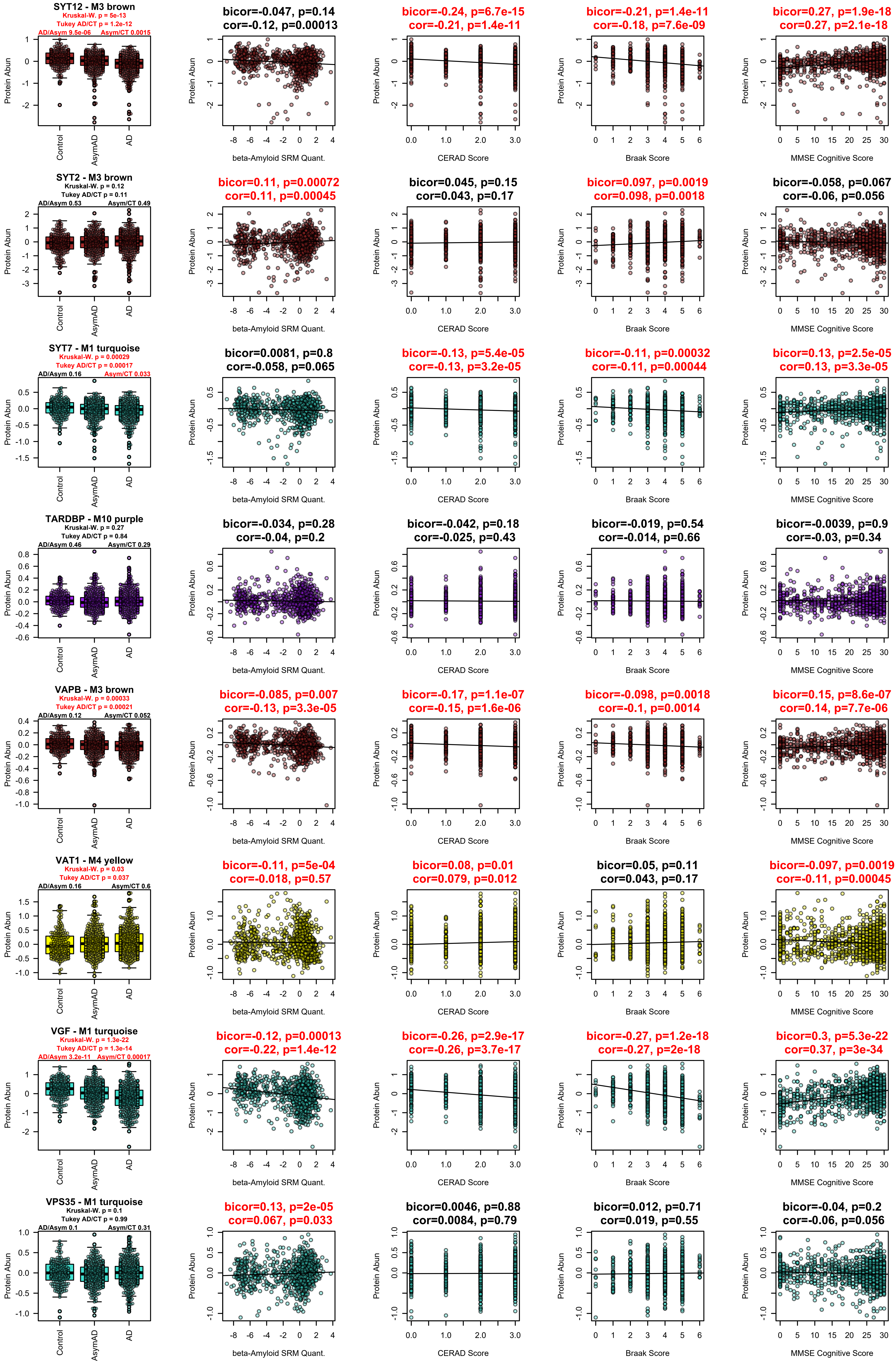

SYT12 - M3 brown
Kruskal-W. p = 5e-13
Tukey AD/CT p = 1.2e-12
AD/Asym 9.5e-06
Asym/CT 0.0015
bicor=-0.047, p=0.14
cor=-0.12, p=0.00013
bicor=-0.24, p=6.7e-15
cor=-0.21, p=1.4e-11
bicor=-0.21, p=1.4e-11
cor=-0.18, p=7.6e-09
bicor=0.27, p=1.9e-18
cor=0.27, p=2.1e-18
SYT2 - M3 brown
Kruskal-W. p = 0.12
Tukey AD/CT p = 0.11
AD/Asym 0.53
Asym/CT 0.49
bicor=0.11, p=0.00072
cor=0.11, p=0.00045
bicor=0.045, p=0.15
cor=0.043, p=0.17
bicor=0.097, p=0.0019
cor=0.098, p=0.0018
bicor=-0.058, p=0.067
cor=-0.06, p=0.056
SYT7 - M1 turquoise
Kruskal-W. p = 0.00029
Tukey AD/CT p = 0.00017
AD/Asym 0.16
Asym/CT 0.033
bicor=0.0081, p=0.8
cor=-0.058, p=0.065
bicor=-0.13, p=5.4e-05
cor=-0.13, p=3.2e-05
bicor=-0.11, p=0.00032
cor=-0.11, p=0.00044
bicor=0.13, p=2.5e-05
cor=0.13, p=3.3e-05
TARDBP - M10 purple
Kruskal-W. p = 0.27
Tukey AD/CT p = 0.84
AD/Asym 0.46
Asym/CT 0.29
bicor=-0.034, p=0.28
cor=-0.04, p=0.2
bicor=-0.042, p=0.18
cor=-0.025, p=0.43
bicor=-0.019, p=0.54
cor=-0.014, p=0.66
bicor=-0.0039, p=0.9
cor=-0.03, p=0.34
VAPB - M3 brown
Kruskal-W. p = 0.00033
Tukey AD/CT p = 0.00021
AD/Asym 0.12
Asym/CT 0.052
bicor=-0.085, p=0.007
cor=-0.13, p=3.3e-05
bicor=-0.17, p=1.1e-07
cor=-0.15, p=1.6e-06
bicor=-0.098, p=0.0018
cor=-0.1, p=0.0014
bicor=0.15, p=8.6e-07
cor=0.14, p=7.7e-06
VAT1 - M4 yellow
Kruskal-W. p = 0.03
Tukey AD/CT p = 0.037
AD/Asym 0.16
Asym/CT 0.6
bicor=-0.11, p=5e-04
cor=-0.018, p=0.57
bicor=0.08, p=0.01
cor=0.079, p=0.012
bicor=0.05, p=0.11
cor=0.043, p=0.17
bicor=-0.097, p=0.0019
cor=-0.11, p=0.00045
VGF - M1 turquoise
Kruskal-W. p = 1.3e-22
Tukey AD/CT p = 1.3e-14
AD/Asym 3.2e-11
Asym/CT 0.00017
bicor=-0.12, p=0.00013
cor=-0.22, p=1.4e-12
bicor=-0.26, p=2.9e-17
cor=-0.26, p=3.7e-17
bicor=-0.27, p=1.2e-18
cor=-0.27, p=2e-18
bicor=0.3, p=5.3e-22
cor=0.37, p=3e-34
VPS35 - M1 turquoise
Kruskal-W. p = 0.1
Tukey AD/CT p = 0.99
AD/Asym 0.1
Asym/CT 0.31
bicor=0.13, p=2e-05
cor=0.067, p=0.033
bicor=0.0046, p=0.88
cor=0.0084, p=0.79
bicor=0.012, p=0.71
cor=0.019, p=0.55
bicor=-0.04, p=0.2
cor=-0.06, p=0.056
Protein Abun
Control
AsymAD
AD
beta-Amyloid SRM Quant.
CERAD Score
Braak Score
MMSE Cognitive Score

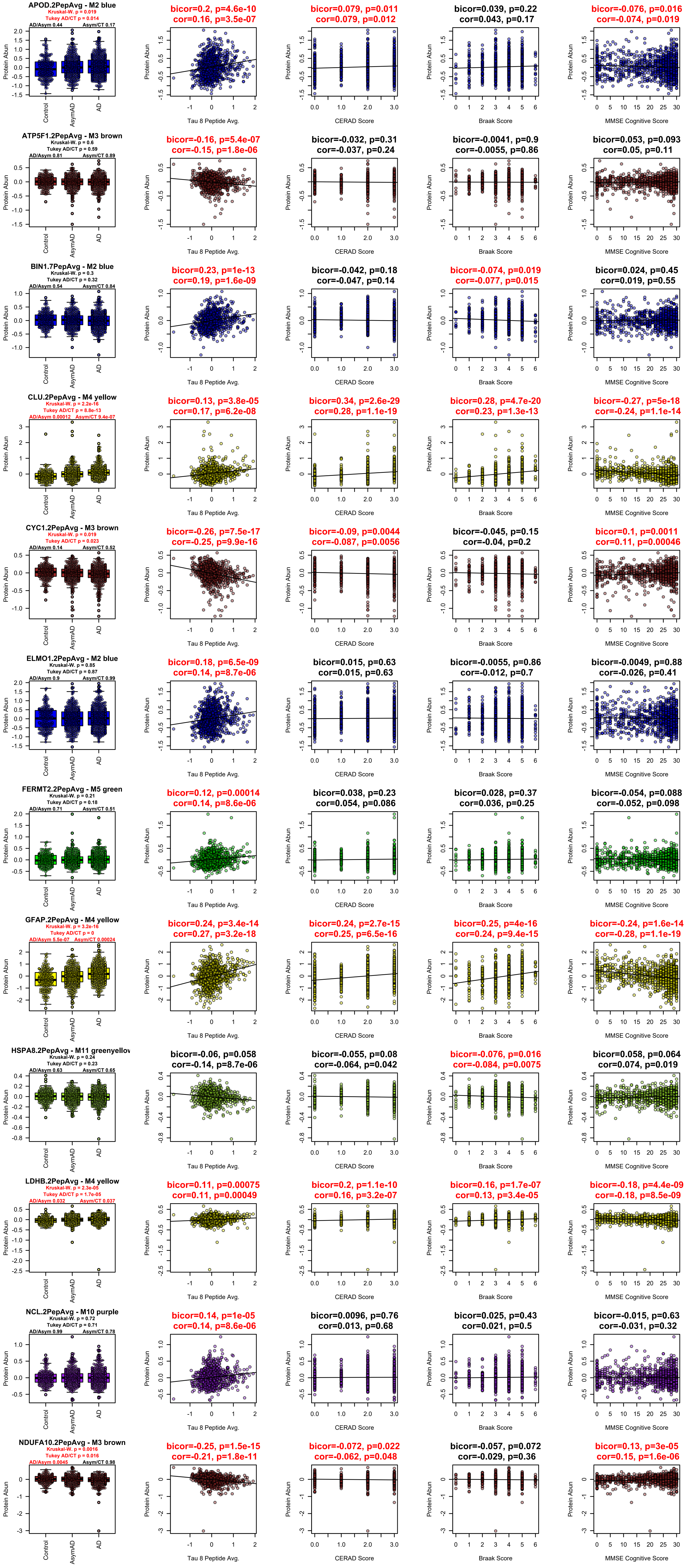

APOD.2PepAvg - M2 blue
Kruskal-W. p = 0.019
Tukey AD/CT p = 0.014
AD/Asym 0.44
Asym/CT 0.17
bicor=0.2, p=4.6e-10
cor=0.16, p=3.5e-07
bicor=0.079, p=0.011
cor=0.079, p=0.012
bicor=0.039, p=0.22
cor=0.043, p=0.17
bicor=-0.076, p=0.016
cor=-0.074, p=0.019
ATP5F1.2PepAvg - M3 brown
Kruskal-W. p = 0.6
Tukey AD/CT p = 0.59
AD/Asym 0.81
Asym/CT 0.89
bicor=-0.16, p=5.4e-07
cor=-0.15, p=1.8e-06
bicor=-0.032, p=0.31
cor=-0.037, p=0.24
bicor=-0.0041, p=0.9
cor=-0.0055, p=0.86
bicor=0.053, p=0.093
cor=0.05, p=0.11
BIN1.7PepAvg - M2 blue
Kruskal-W. p = 0.3
Tukey AD/CT p = 0.32
AD/Asym 0.54
Asym/CT 0.84
bicor=0.23, p=1e-13
cor=0.19, p=1.6e-09
bicor=-0.042, p=0.18
cor=-0.047, p=0.14
bicor=-0.074, p=0.019
cor=-0.077, p=0.015
bicor=0.024, p=0.45
cor=0.019, p=0.55
CLU.2PepAvg - M4 yellow
Kruskal-W. p = 2.2e-16
Tukey AD/CT p = 8.8e-13
AD/Asym 0.00012
Asym/CT 9.4e-07
bicor=0.13, p=3.8e-05
cor=0.17, p=6.2e-08
bicor=0.34, p=2.6e-29
cor=0.28, p=1.1e-19
bicor=0.28, p=4.7e-20
cor=0.23, p=1.3e-13
bicor=-0.27, p=5e-18
cor=-0.24, p=1.1e-14
CYC1.2PepAvg - M3 brown
Kruskal-W. p = 0.019
Tukey AD/CT p = 0.023
AD/Asym 0.14
Asym/CT 0.52
bicor=-0.26, p=7.5e-17
cor=-0.25, p=9.9e-16
bicor=-0.09, p=0.0044
cor=-0.087, p=0.0056
bicor=-0.045, p=0.15
cor=-0.04, p=0.2
bicor=0.1, p=0.0011
cor=0.11, p=0.00046
ELMO1.2PepAvg - M2 blue
Kruskal-W. p = 0.85
Tukey AD/CT p = 0.87
AD/Asym 0.9
Asym/CT 0.99
bicor=0.18, p=6.5e-09
cor=0.14, p=8.7e-06
bicor=0.015, p=0.63
cor=0.015, p=0.63
bicor=-0.0055, p=0.86
cor=-0.012, p=0.7
bicor=-0.0049, p=0.88
cor=-0.026, p=0.41
FERMT2.2PepAvg - M5 green
Kruskal-W. p = 0.21
Tukey AD/CT p = 0.18
AD/Asym 0.71
Asym/CT 0.51
bicor=0.12, p=0.00014
cor=0.14, p=8.6e-06
bicor=0.038, p=0.23
cor=0.054, p=0.086
bicor=0.028, p=0.37
cor=0.036, p=0.25
bicor=-0.054, p=0.088
cor=-0.052, p=0.098
GFAP.2PepAvg - M4 yellow
Kruskal-W. p = 3.2e-16
Tukey AD/CT p = 0
AD/Asym 5.5e-07
Asym/CT 0.00024
bicor=0.24, p=3.4e-14
cor=0.27, p=3.2e-18
bicor=0.24, p=2.7e-15
cor=0.25, p=6.5e-16
bicor=0.25, p=4e-16
cor=0.24, p=9.4e-15
bicor=-0.24, p=1.6e-14
cor=-0.28, p=1.1e-19
HSPA8.2PepAvg - M11 greenyellow
Kruskal-W. p = 0.24
Tukey AD/CT p = 0.23
AD/Asym 0.63
Asym/CT 0.65
bicor=-0.06, p=0.058
cor=-0.14, p=8.7e-06
bicor=-0.055, p=0.08
cor=-0.064, p=0.042
bicor=-0.076, p=0.016
cor=-0.084, p=0.0075
bicor=0.058, p=0.064
cor=0.074, p=0.019
LDHB.2PepAvg - M4 yellow
Kruskal-W. p = 2.3e-05
Tukey AD/CT p = 1.7e-05
AD/Asym 0.032
Asym/CT 0.037
bicor=0.11, p=0.00075
cor=0.11, p=0.00049
bicor=0.2, p=1.1e-10
cor=0.16, p=3.2e-07
bicor=0.16, p=1.7e-07
cor=0.13, p=3.4e-05
bicor=-0.18, p=4.4e-09
cor=-0.18, p=8.5e-09
NCL.2PepAvg - M10 purple
Kruskal-W. p = 0.72
Tukey AD/CT p = 0.71
AD/Asym 0.99
Asym/CT 0.78
bicor=0.14, p=1e-05
cor=0.14, p=8.6e-06
bicor=0.0096, p=0.76
cor=0.013, p=0.68
bicor=0.025, p=0.43
cor=0.021, p=0.5
bicor=-0.015, p=0.63
cor=-0.031, p=0.32
NDUFA10.2PepAvg - M3 brown
Kruskal-W. p = 0.0016
Tukey AD/CT p = 0.016
AD/Asym 0.0045
Asym/CT 0.98
bicor=-0.25, p=1.5e-15
cor=-0.21, p=1.8e-11
bicor=-0.072, p=0.022
cor=-0.062, p=0.048
bicor=-0.057, p=0.072
cor=-0.029, p=0.36
bicor=0.13, p=3e-05
cor=0.15, p=1.6e-06
Protein Abun
Control
AsymAD
AD
Tau 8 Peptide Avg.
CERAD Score
Braak Score
MMSE Cognitive Score

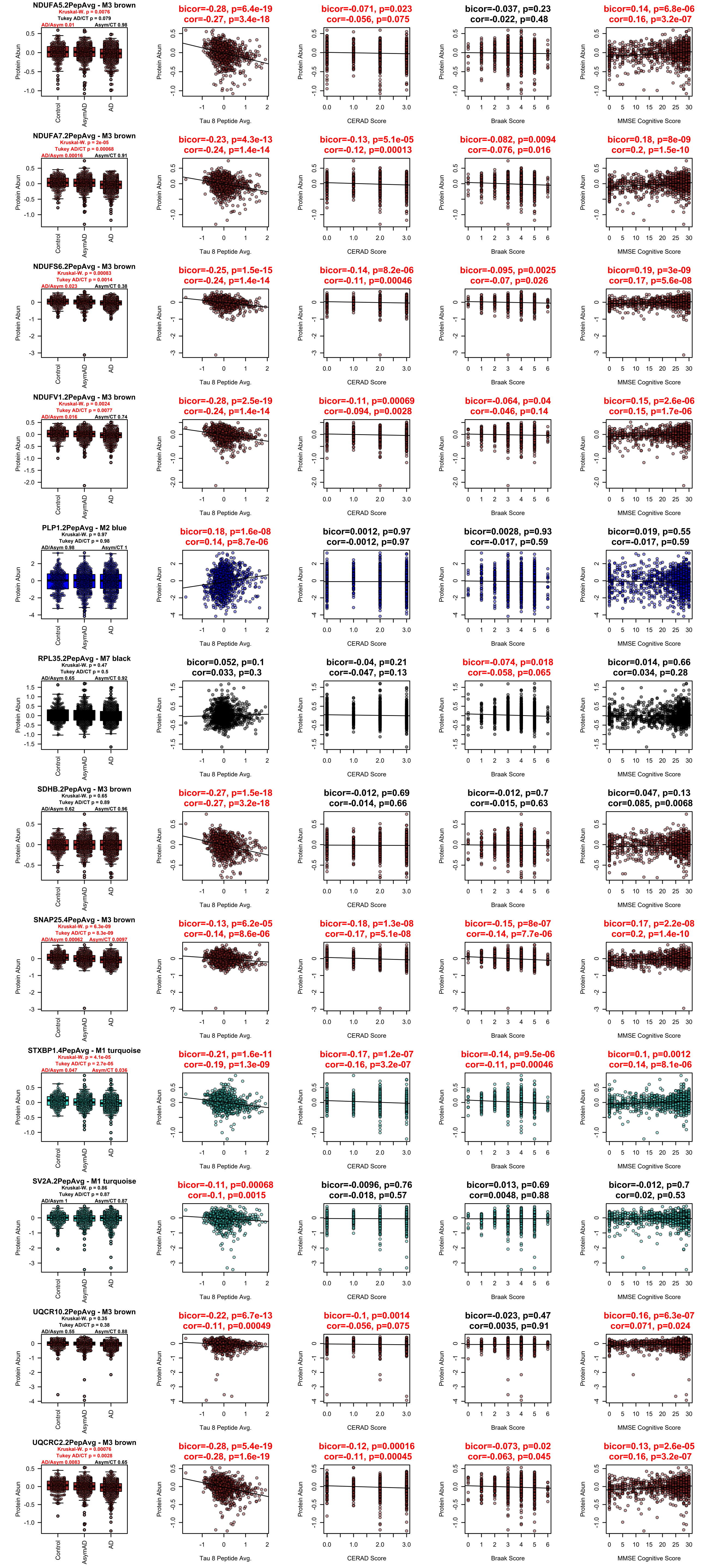

NDUFA5.2PepAvg - M3 brown
Kruskal-W. p = 0.0076
Tukey AD/CT p = 0.079
AD/Asym 0.01
Asym/CT 0.98
bicor=-0.28, p=6.4e-19
cor=-0.27, p=3.4e-18
bicor=-0.071, p=0.023
cor=-0.056, p=0.075
bicor=-0.037, p=0.23
cor=-0.022, p=0.48
bicor=0.14, p=6.8e-06
cor=0.16, p=3.2e-07
NDUFA7.2PepAvg - M3 brown
Kruskal-W. p = 2e-05
Tukey AD/CT p = 0.00068
AD/Asym 0.00016
Asym/CT 0.91
bicor=-0.23, p=4.3e-13
cor=-0.24, p=1.4e-14
bicor=-0.13, p=5.1e-05
cor=-0.12, p=0.00013
bicor=-0.082, p=0.0094
cor=-0.076, p=0.016
bicor=0.18, p=8e-09
cor=0.2, p=1.5e-10
NDUFS6.2PepAvg - M3 brown
Kruskal-W. p = 0.00083
Tukey AD/CT p = 0.0014
AD/Asym 0.023
Asym/CT 0.38
bicor=-0.25, p=1.5e-15
cor=-0.24, p=1.4e-14
bicor=-0.14, p=8.2e-06
cor=-0.11, p=0.00046
bicor=-0.095, p=0.0025
cor=-0.07, p=0.026
bicor=0.19, p=3e-09
cor=0.17, p=5.6e-08
NDUFV1.2PepAvg - M3 brown
Kruskal-W. p = 0.0024
Tukey AD/CT p = 0.0077
AD/Asym 0.016
Asym/CT 0.74
bicor=-0.28, p=2.5e-19
cor=-0.24, p=1.4e-14
bicor=-0.11, p=0.00069
cor=-0.094, p=0.0028
bicor=-0.064, p=0.04
cor=-0.046, p=0.14
bicor=0.15, p=2.6e-06
cor=0.15, p=1.7e-06
PLP1.2PepAvg - M2 blue
Kruskal-W. p = 0.97
Tukey AD/CT p = 0.98
AD/Asym 0.98
Asym/CT 1
bicor=0.18, p=1.6e-08
cor=0.14, p=8.7e-06
bicor=0.0012, p=0.97
cor=-0.0012, p=0.97
bicor=0.0028, p=0.93
cor=-0.017, p=0.59
bicor=0.019, p=0.55
cor=-0.017, p=0.59
RPL35.2PepAvg - M7 black
Kruskal-W. p = 0.47
Tukey AD/CT p = 0.5
AD/Asym 0.65
Asym/CT 0.92
bicor=0.052, p=0.1
cor=0.033, p=0.3
bicor=-0.04, p=0.21
cor=-0.047, p=0.13
bicor=-0.074, p=0.018
cor=-0.058, p=0.065
bicor=0.014, p=0.66
cor=0.034, p=0.28
SDHB.2PepAvg - M3 brown
Kruskal-W. p = 0.65
Tukey AD/CT p = 0.89
AD/Asym 0.62
Asym/CT 0.96
bicor=-0.27, p=1.5e-18
cor=-0.27, p=3.2e-18
bicor=-0.012, p=0.69
cor=-0.014, p=0.66
bicor=-0.012, p=0.7
cor=-0.015, p=0.63
bicor=0.047, p=0.13
cor=0.085, p=0.0068
SNAP25.4PepAvg - M3 brown
Kruskal-W. p = 6.3e-09
Tukey AD/CT p = 8.3e-09
AD/Asym 0.00062
Asym/CT 0.0097
bicor=-0.13, p=6.2e-05
cor=-0.14, p=8.6e-06
bicor=-0.18, p=1.3e-08
cor=-0.17, p=5.1e-08
bicor=-0.15, p=8e-07
cor=-0.14, p=7.7e-06
bicor=0.17, p=2.2e-08
cor=0.2, p=1.4e-10
STXBP1.4PepAvg - M1 turquoise
Kruskal-W. p = 4.1e-05
Tukey AD/CT p = 2.7e-05
AD/Asym 0.047
Asym/CT 0.036
bicor=-0.21, p=1.6e-11
cor=-0.19, p=1.3e-09
bicor=-0.17, p=1.2e-07
cor=-0.16, p=3.2e-07
bicor=-0.14, p=9.5e-06
cor=-0.11, p=0.00046
bicor=0.1, p=0.0012
cor=0.14, p=8.1e-06
SV2A.2PepAvg - M1 turquoise
Kruskal-W. p = 0.86
Tukey AD/CT p = 0.87
AD/Asym 1
Asym/CT 0.87
bicor=-0.11, p=0.00068
cor=-0.1, p=0.0015
bicor=-0.0096, p=0.76
cor=-0.018, p=0.57
bicor=0.013, p=0.69
cor=0.0048, p=0.88
bicor=-0.012, p=0.7
cor=0.02, p=0.53
UQCR10.2PepAvg - M3 brown
Kruskal-W. p = 0.35
Tukey AD/CT p = 0.38
AD/Asym 0.55
Asym/CT 0.88
bicor=-0.22, p=6.7e-13
cor=-0.11, p=0.00049
bicor=-0.1, p=0.0014
cor=-0.056, p=0.075
bicor=-0.023, p=0.47
cor=0.0035, p=0.91
bicor=0.16, p=6.3e-07
cor=0.071, p=0.024
UQCRC2.2PepAvg - M3 brown
Kruskal-W. p = 0.00076
Tukey AD/CT p = 0.0028
AD/Asym 0.0083
Asym/CT 0.65
bicor=-0.28, p=5.4e-19
cor=-0.28, p=1.6e-19
bicor=-0.12, p=0.00016
cor=-0.11, p=0.00045
bicor=-0.073, p=0.02
cor=-0.063, p=0.045
bicor=0.13, p=2.6e-05
cor=0.16, p=3.2e-07
Protein Abun
Control
AsymAD
AD
Tau 8 Peptide Avg.
CERAD Score
Braak Score
MMSE Cognitive Score

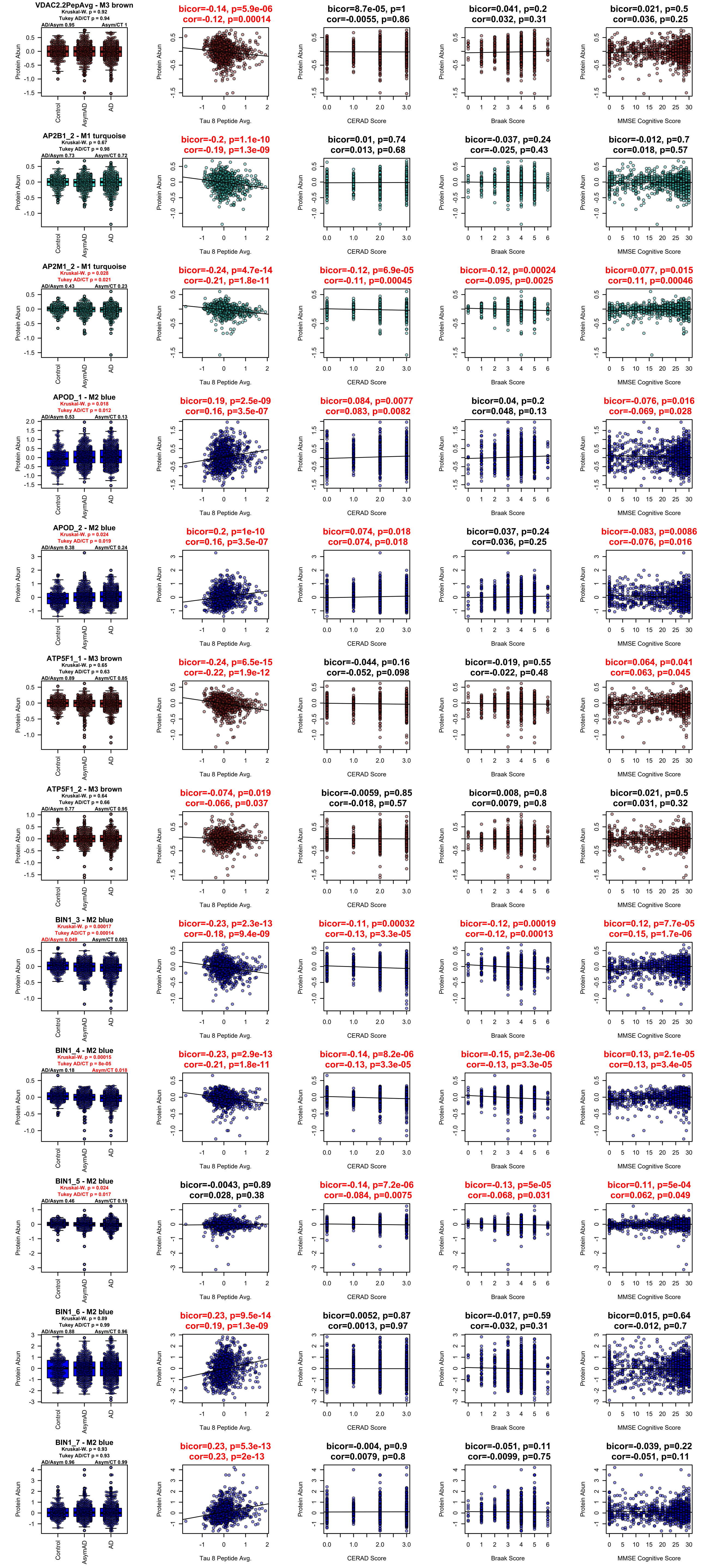
VDAC2.2PepAvg - M3 brown
Kruskal-W. p = 0.92
Tukey AD/CT p = 0.94
AD/Asym 0.95
Asym/CT 1
bicor=-0.14, p=5.9e-06
cor=-0.12, p=0.00014
bicor=8.7e-05, p=1
cor=-0.0055, p=0.86
bicor=0.041, p=0.2
cor=0.032, p=0.31
bicor=0.021, p=0.5
cor=0.036, p=0.25
AP2B1_2 - M1 turquoise
Kruskal-W. p = 0.67
Tukey AD/CT p = 0.98
AD/Asym 0.73
Asym/CT 0.72
bicor=-0.2, p=1.1e-10
cor=-0.19, p=1.3e-09
bicor=0.01, p=0.74
cor=0.013, p=0.68
bicor=-0.037, p=0.24
cor=-0.025, p=0.43
bicor=-0.012, p=0.7
cor=0.018, p=0.57
AP2M1_2 - M1 turquoise
Kruskal-W. p = 0.028
Tukey AD/CT p = 0.021
AD/Asym 0.43
Asym/CT 0.23
bicor=-0.24, p=4.7e-14
cor=-0.21, p=1.8e-11
bicor=-0.12, p=6.9e-05
cor=-0.11, p=0.00045
bicor=-0.12, p=0.00024
cor=-0.095, p=0.0025
bicor=0.077, p=0.015
cor=0.11, p=0.00046
APOD_1 - M2 blue
Kruskal-W. p = 0.018
Tukey AD/CT p = 0.012
AD/Asym 0.53
Asym/CT 0.13
bicor=0.19, p=2.5e-09
cor=0.16, p=3.5e-07
bicor=0.084, p=0.0077
cor=0.083, p=0.0082
bicor=0.04, p=0.2
cor=0.048, p=0.13
bicor=-0.076, p=0.016
cor=-0.069, p=0.028
APOD_2 - M2 blue
Kruskal-W. p = 0.024
Tukey AD/CT p = 0.019
AD/Asym 0.38
Asym/CT 0.24
bicor=0.2, p=1e-10
cor=0.16, p=3.5e-07
bicor=0.074, p=0.018
cor=0.074, p=0.018
bicor=0.037, p=0.24
cor=0.036, p=0.25
bicor=-0.083, p=0.0086
cor=-0.076, p=0.016
ATP5F1_1 - M3 brown
Kruskal-W. p = 0.65
Tukey AD/CT p = 0.63
AD/Asym 0.89
Asym/CT 0.85
bicor=-0.24, p=6.5e-15
cor=-0.22, p=1.9e-12
bicor=-0.044, p=0.16
cor=-0.052, p=0.098
bicor=-0.019, p=0.55
cor=-0.022, p=0.48
bicor=0.064, p=0.041
cor=0.063, p=0.045
ATP5F1_2 - M3 brown
Kruskal-W. p = 0.64
Tukey AD/CT p = 0.66
AD/Asym 0.77
Asym/CT 0.95
bicor=-0.074, p=0.019
cor=-0.066, p=0.037
bicor=-0.0059, p=0.85
cor=-0.018, p=0.57
bicor=0.008, p=0.8
cor=0.0079, p=0.8
bicor=0.021, p=0.5
cor=0.031, p=0.32
BIN1_3 - M2 blue
Kruskal-W. p = 0.00017
Tukey AD/CT p = 0.00014
AD/Asym 0.049
Asym/CT 0.083
bicor=-0.23, p=2.3e-13
cor=-0.18, p=9.4e-09
bicor=-0.11, p=0.00032
cor=-0.13, p=3.3e-05
bicor=-0.12, p=0.00019
cor=-0.12, p=0.00013
bicor=0.12, p=7.7e-05
cor=0.15, p=1.7e-06
BIN1_4 - M2 blue
Kruskal-W. p = 0.00015
Tukey AD/CT p = 8e-05
AD/Asym 0.18
Asym/CT 0.018
bicor=-0.23, p=2.9e-13
cor=-0.21, p=1.8e-11
bicor=-0.14, p=8.2e-06
cor=-0.13, p=3.3e-05
bicor=-0.15, p=2.3e-06
cor=-0.13, p=3.3e-05
bicor=0.13, p=2.1e-05
cor=0.13, p=3.4e-05
BIN1_5 - M2 blue
Kruskal-W. p = 0.024
Tukey AD/CT p = 0.017
AD/Asym 0.46
Asym/CT 0.19
bicor=-0.0043, p=0.89
cor=0.028, p=0.38
bicor=-0.14, p=7.2e-06
cor=-0.084, p=0.0075
bicor=-0.13, p=5e-05
cor=-0.068, p=0.031
bicor=0.11, p=5e-04
cor=0.062, p=0.049
BIN1_6 - M2 blue
Kruskal-W. p = 0.89
Tukey AD/CT p = 0.99
AD/Asym 0.88
Asym/CT 0.96
bicor=0.23, p=9.5e-14
cor=0.19, p=1.3e-09
bicor=0.0052, p=0.87
cor=0.0013, p=0.97
bicor=-0.017, p=0.59
cor=-0.032, p=0.31
bicor=0.015, p=0.64
cor=-0.012, p=0.7
BIN1_7 - M2 blue
Kruskal-W. p = 0.93
Tukey AD/CT p = 0.93
AD/Asym 0.96
Asym/CT 0.99
bicor=0.23, p=5.3e-13
cor=0.23, p=2e-13
bicor=-0.004, p=0.9
cor=0.0079, p=0.8
bicor=-0.051, p=0.11
cor=-0.0099, p=0.75
bicor=-0.039, p=0.22
cor=-0.051, p=0.11
Protein Abun
Control
AsymAD
AD
Tau 8 Peptide Avg.
CERAD Score
Braak Score
MMSE Cognitive Score

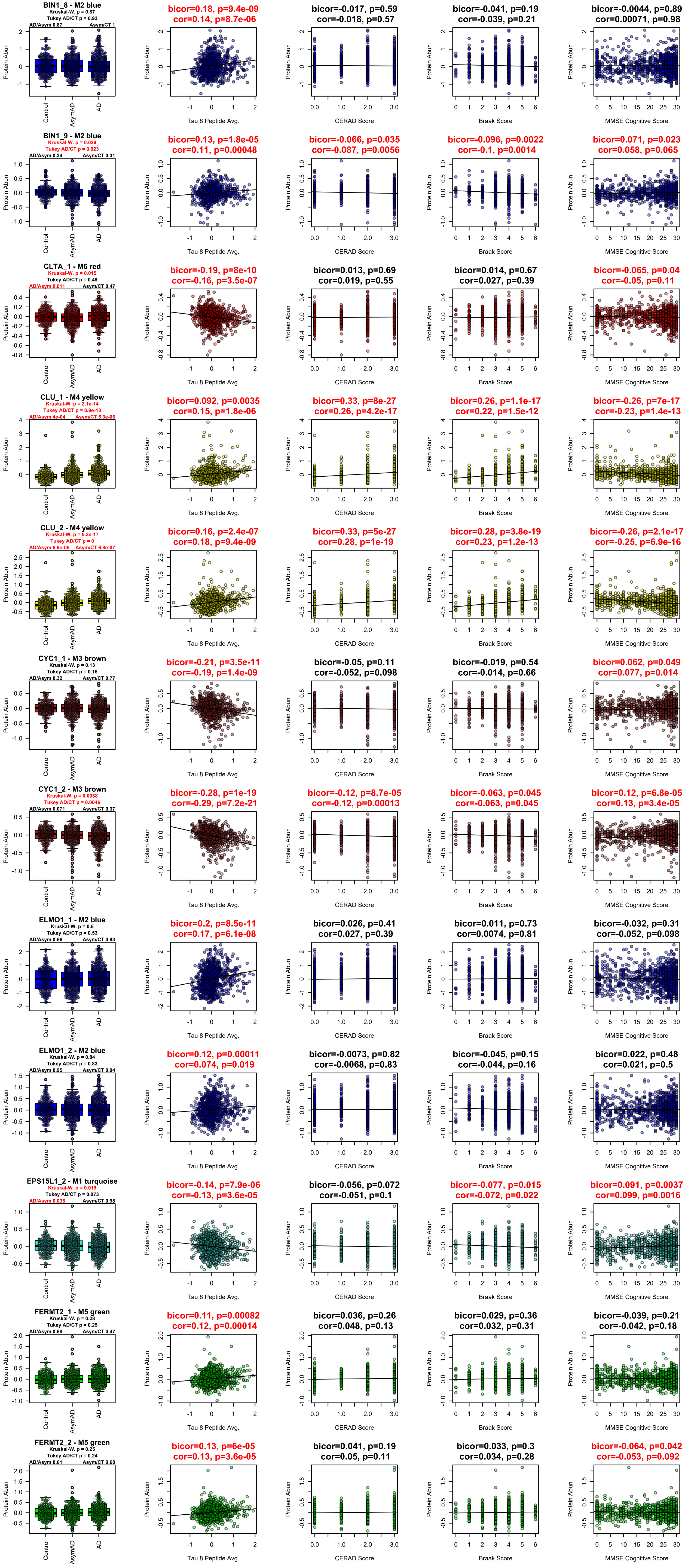

BIN1_8 - M2 blue
Kruskal-W. p = 0.87
Tukey AD/CT p = 0.93
AD/Asym 0.87
Asym/CT 1
bicor=0.18, p=9.4e-09
cor=0.14, p=8.7e-06
bicor=-0.017, p=0.59
cor=-0.018, p=0.57
bicor=-0.041, p=0.19
cor=-0.039, p=0.21
bicor=-0.0044, p=0.89
cor=0.00071, p=0.98
BIN1_9 - M2 blue
Kruskal-W. p = 0.029
Tukey AD/CT p = 0.023
AD/Asym 0.34
Asym/CT 0.31
bicor=0.13, p=1.8e-05
cor=0.11, p=0.00048
bicor=-0.066, p=0.035
cor=-0.087, p=0.0056
bicor=-0.096, p=0.0022
cor=-0.1, p=0.0014
bicor=0.071, p=0.023
cor=0.058, p=0.065
CLTA_1 - M6 red
Kruskal-W. p = 0.015
Tukey AD/CT p = 0.49
AD/Asym 0.011
Asym/CT 0.47
bicor=-0.19, p=8e-10
cor=-0.16, p=3.5e-07
bicor=0.013, p=0.69
cor=0.019, p=0.55
bicor=0.014, p=0.67
cor=0.027, p=0.39
bicor=-0.065, p=0.04
cor=-0.05, p=0.11
CLU_1 - M4 yellow
Kruskal-W. p = 2.1e-14
Tukey AD/CT p = 8.9e-13
AD/Asym 4e-04
Asym/CT 5.3e-06
bicor=0.092, p=0.0035
cor=0.15, p=1.8e-06
bicor=0.33, p=8e-27
cor=0.26, p=4.2e-17
bicor=0.26, p=1.1e-17
cor=0.22, p=1.5e-12
bicor=-0.26, p=7e-17
cor=-0.23, p=1.4e-13
CLU_2 - M4 yellow
Kruskal-W. p = 5.3e-17
Tukey AD/CT p = 0
AD/Asym 6.9e-05
Asym/CT 6.8e-07
bicor=0.16, p=2.4e-07
cor=0.18, p=9.4e-09
bicor=0.33, p=5e-27
cor=0.28, p=1e-19
bicor=0.28, p=3.8e-19
cor=0.23, p=1.2e-13
bicor=-0.26, p=2.1e-17
cor=-0.25, p=6.9e-16
CYC1_1 - M3 brown
Kruskal-W. p = 0.13
Tukey AD/CT p = 0.15
AD/Asym 0.32
Asym/CT 0.77
bicor=-0.21, p=3.5e-11
cor=-0.19, p=1.4e-09
bicor=-0.05, p=0.11
cor=-0.052, p=0.098
bicor=-0.019, p=0.54
cor=-0.014, p=0.66
bicor=0.062, p=0.049
cor=0.077, p=0.014
CYC1_2 - M3 brown
Kruskal-W. p = 0.0038
Tukey AD/CT p = 0.0046
AD/Asym 0.071
Asym/CT 0.37
bicor=-0.28, p=1e-19
cor=-0.29, p=7.2e-21
bicor=-0.12, p=8.7e-05
cor=-0.12, p=0.00013
bicor=-0.063, p=0.045
cor=-0.063, p=0.045
bicor=0.12, p=6.8e-05
cor=0.13, p=3.4e-05
ELMO1_1 - M2 blue
Kruskal-W. p = 0.5
Tukey AD/CT p = 0.53
AD/Asym 0.66
Asym/CT 0.93
bicor=0.2, p=8.5e-11
cor=0.17, p=6.1e-08
bicor=0.026, p=0.41
cor=0.027, p=0.39
bicor=0.011, p=0.73
cor=0.0074, p=0.81
bicor=-0.032, p=0.31
cor=-0.052, p=0.098
ELMO1_2 - M2 blue
Kruskal-W. p = 0.84
Tukey AD/CT p = 0.83
AD/Asym 0.95
Asym/CT 0.94
bicor=0.12, p=0.00011
cor=0.074, p=0.019
bicor=-0.0073, p=0.82
cor=-0.0068, p=0.83
bicor=-0.045, p=0.15
cor=-0.044, p=0.16
bicor=0.022, p=0.48
cor=0.021, p=0.5
EPS15L1_2 - M1 turquoise
Kruskal-W. p = 0.019
Tukey AD/CT p = 0.073
AD/Asym 0.035
Asym/CT 0.98
bicor=-0.14, p=7.9e-06
cor=-0.13, p=3.6e-05
bicor=-0.056, p=0.072
cor=-0.051, p=0.1
bicor=-0.077, p=0.015
cor=-0.072, p=0.022
bicor=0.091, p=0.0037
cor=0.099, p=0.0016
FERMT2_1 - M5 green
Kruskal-W. p = 0.28
Tukey AD/CT p = 0.25
AD/Asym 0.88
Asym/CT 0.47
bicor=0.11, p=0.00082
cor=0.12, p=0.00014
bicor=0.036, p=0.26
cor=0.048, p=0.13
bicor=0.029, p=0.36
cor=0.032, p=0.31
bicor=-0.039, p=0.21
cor=-0.042, p=0.18
FERMT2_2 - M5 green
Kruskal-W. p = 0.25
Tukey AD/CT p = 0.24
AD/Asym 0.61
Asym/CT 0.68
bicor=0.13, p=6e-05
cor=0.13, p=3.6e-05
bicor=0.041, p=0.19
cor=0.05, p=0.11
bicor=0.033, p=0.3
cor=0.034, p=0.28
bicor=-0.064, p=0.042
cor=-0.053, p=0.092
Protein Abun
Control
AsymAD
AD
Tau 8 Peptide Avg.
CERAD Score
Braak Score
MMSE Cognitive Score

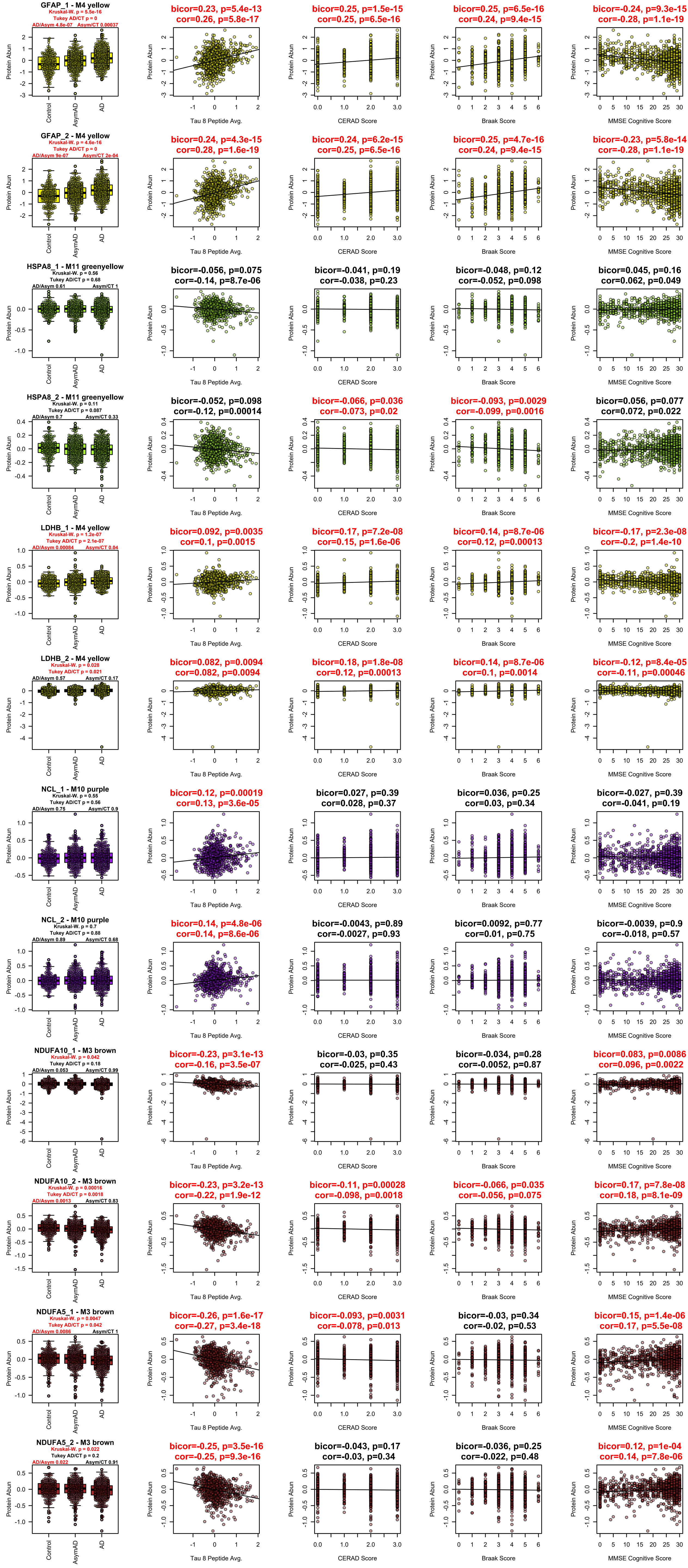

GFAP_1 - M4 yellow
Kruskal-W. p = 5.5e-16
Tukey AD/CT p = 0
AD/Asym 4.8e-07
Asym/CT 0.00037
bicor=0.23, p=5.4e-13
cor=0.26, p=5.8e-17
bicor=0.25, p=1.5e-15
cor=0.25, p=6.5e-16
bicor=0.25, p=6.5e-16
cor=0.24, p=9.4e-15
bicor=-0.24, p=9.3e-15
cor=-0.28, p=1.1e-19
GFAP_2 - M4 yellow
Kruskal-W. p = 4.6e-16
Tukey AD/CT p = 0
AD/Asym 9e-07
Asym/CT 2e-04
bicor=0.24, p=4.3e-15
cor=0.28, p=1.6e-19
bicor=0.24, p=6.2e-15
cor=0.25, p=6.5e-16
bicor=0.25, p=4.7e-16
cor=0.24, p=9.4e-15
bicor=-0.23, p=5.8e-14
cor=-0.28, p=1.1e-19
HSPA8_1 - M11 greenyellow
Kruskal-W. p = 0.56
Tukey AD/CT p = 0.68
AD/Asym 0.61
Asym/CT 1
bicor=-0.056, p=0.075
cor=-0.14, p=8.7e-06
bicor=-0.041, p=0.19
cor=-0.038, p=0.23
bicor=-0.048, p=0.12
cor=-0.052, p=0.098
bicor=0.045, p=0.16
cor=0.062, p=0.049
HSPA8_2 - M11 greenyellow
Kruskal-W. p = 0.11
Tukey AD/CT p = 0.087
AD/Asym 0.7
Asym/CT 0.33
bicor=-0.052, p=0.098
cor=-0.12, p=0.00014
bicor=-0.066, p=0.036
cor=-0.073, p=0.02
bicor=-0.093, p=0.0029
cor=-0.099, p=0.0016
bicor=0.056, p=0.077
cor=0.072, p=0.022
LDHB_1 - M4 yellow
Kruskal-W. p = 1.2e-07
Tukey AD/CT p = 2.1e-07
AD/Asym 0.00084
Asym/CT 0.04
bicor=0.092, p=0.0035
cor=0.1, p=0.0015
bicor=0.17, p=7.2e-08
cor=0.15, p=1.6e-06
bicor=0.14, p=8.7e-06
cor=0.12, p=0.00013
bicor=-0.17, p=2.3e-08
cor=-0.2, p=1.4e-10
LDHB_2 - M4 yellow
Kruskal-W. p = 0.028
Tukey AD/CT p = 0.021
AD/Asym 0.57
Asym/CT 0.17
bicor=0.082, p=0.0094
cor=0.082, p=0.0094
bicor=0.18, p=1.8e-08
cor=0.12, p=0.00013
bicor=0.14, p=8.7e-06
cor=0.1, p=0.0014
bicor=-0.12, p=8.4e-05
cor=-0.11, p=0.00046
NCL_1 - M10 purple
Kruskal-W. p = 0.55
Tukey AD/CT p = 0.56
AD/Asym 0.75
Asym/CT 0.9
bicor=0.12, p=0.00019
cor=0.13, p=3.6e-05
bicor=0.027, p=0.39
cor=0.028, p=0.37
bicor=0.036, p=0.25
cor=0.03, p=0.34
bicor=-0.027, p=0.39
cor=-0.041, p=0.19
NCL_2 - M10 purple
Kruskal-W. p = 0.7
Tukey AD/CT p = 0.88
AD/Asym 0.89
Asym/CT 0.68
bicor=0.14, p=4.8e-06
cor=0.14, p=8.6e-06
bicor=-0.0043, p=0.89
cor=-0.0027, p=0.93
bicor=0.0092, p=0.77
cor=0.01, p=0.75
bicor=-0.0039, p=0.9
cor=-0.018, p=0.57
NDUFA10_1 - M3 brown
Kruskal-W. p = 0.042
Tukey AD/CT p = 0.18
AD/Asym 0.053
Asym/CT 0.99
bicor=-0.23, p=3.1e-13
cor=-0.16, p=3.5e-07
bicor=-0.03, p=0.35
cor=-0.025, p=0.43
bicor=-0.034, p=0.28
cor=-0.0052, p=0.87
bicor=0.083, p=0.0086
cor=0.096, p=0.0022
NDUFA10_2 - M3 brown
Kruskal-W. p = 0.00016
Tukey AD/CT p = 0.0018
AD/Asym 0.0013
Asym/CT 0.83
bicor=-0.23, p=3.2e-13
cor=-0.22, p=1.9e-12
bicor=-0.11, p=0.00028
cor=-0.098, p=0.0018
bicor=-0.066, p=0.035
cor=-0.056, p=0.075
bicor=0.17, p=7.8e-08
cor=0.18, p=8.1e-09
NDUFA5_1 - M3 brown
Kruskal-W. p = 0.0047
Tukey AD/CT p = 0.042
AD/Asym 0.0086
Asym/CT 1
bicor=-0.26, p=1.6e-17
cor=-0.27, p=3.4e-18
bicor=-0.093, p=0.0031
cor=-0.078, p=0.013
bicor=-0.03, p=0.34
cor=-0.02, p=0.53
bicor=0.15, p=1.4e-06
cor=0.17, p=5.5e-08
NDUFA5_2 - M3 brown
Kruskal-W. p = 0.022
Tukey AD/CT p = 0.2
AD/Asym 0.022
Asym/CT 0.91
bicor=-0.25, p=3.5e-16
cor=-0.25, p=9.3e-16
bicor=-0.043, p=0.17
cor=-0.03, p=0.34
bicor=-0.036, p=0.25
cor=-0.022, p=0.48
bicor=0.12, p=1e-04
cor=0.14, p=7.8e-06
Protein Abun
Control
AsymAD
AD
Tau 8 Peptide Avg.
CERAD Score
Braak Score
MMSE Cognitive Score

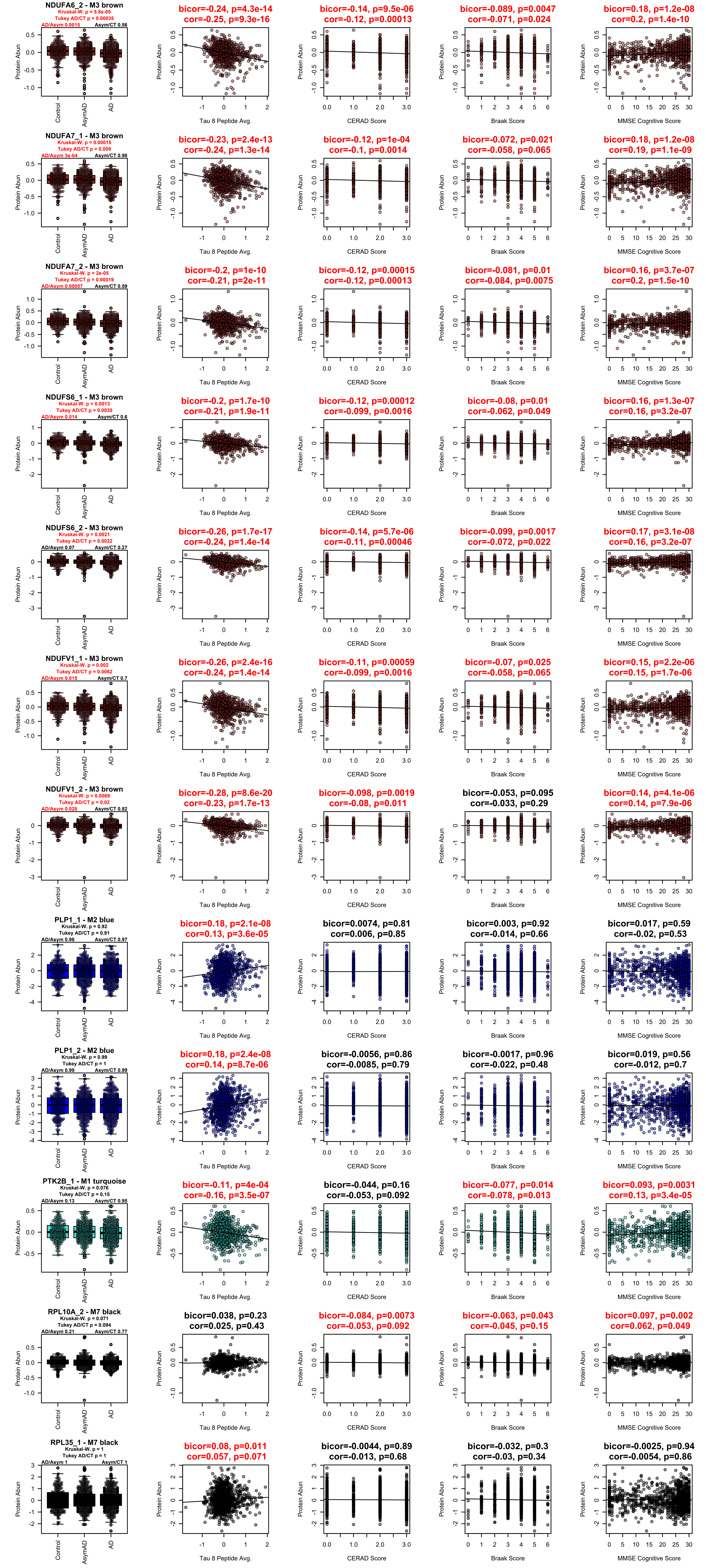
NDUFA6_2 - M3 brown
Kruskal-W. p = 5.8e-05
Tukey AD/CT p = 0.00035
AD/Asym 0.0015
Asym/CT 0.56
bicor=-0.24, p=4.3e-14
cor=-0.25, p=9.3e-16
bicor=-0.14, p=9.5e-06
cor=-0.12, p=0.00013
bicor=-0.089, p=0.0047
cor=-0.071, p=0.024
bicor=0.18, p=1.2e-08
cor=0.2, p=1.4e-10
NDUFA7_1 - M3 brown
Kruskal-W. p = 0.00015
Tukey AD/CT p = 0.009
AD/Asym 3e-04
Asym/CT 0.98
bicor=-0.23, p=2.4e-13
cor=-0.24, p=1.3e-14
bicor=-0.12, p=1e-04
cor=-0.1, p=0.0014
bicor=-0.072, p=0.021
cor=-0.058, p=0.065
bicor=0.18, p=1.2e-08
cor=0.19, p=1.1e-09
NDUFA7_2 - M3 brown
Kruskal-W. p = 2e-05
Tukey AD/CT p = 0.00019
AD/Asym 0.00057
Asym/CT 0.59
bicor=-0.2, p=1e-10
cor=-0.21, p=2e-11
bicor=-0.12, p=0.00015
cor=-0.12, p=0.00013
bicor=-0.081, p=0.01
cor=-0.084, p=0.0075
bicor=0.16, p=3.7e-07
cor=0.2, p=1.5e-10
NDUFS6_1 - M3 brown
Kruskal-W. p = 0.0013
Tukey AD/CT p = 0.0035
AD/Asym 0.014
Asym/CT 0.6
bicor=-0.2, p=1.7e-10
cor=-0.21, p=1.9e-11
bicor=-0.12, p=0.00012
cor=-0.099, p=0.0016
bicor=-0.08, p=0.01
cor=-0.062, p=0.049
bicor=0.16, p=1.3e-07
cor=0.16, p=3.2e-07
NDUFS6_2 - M3 brown
Kruskal-W. p = 0.0021
Tukey AD/CT p = 0.0022
AD/Asym 0.07
Asym/CT 0.27
bicor=-0.26, p=1.7e-17
cor=-0.24, p=1.4e-14
bicor=-0.14, p=5.7e-06
cor=-0.11, p=0.00046
bicor=-0.099, p=0.0017
cor=-0.072, p=0.022
bicor=0.17, p=3.1e-08
cor=0.16, p=3.2e-07
NDUFV1_1 - M3 brown
Kruskal-W. p = 0.002
Tukey AD/CT p = 0.0062
AD/Asym 0.015
Asym/CT 0.7
bicor=-0.26, p=2.4e-16
cor=-0.24, p=1.4e-14
bicor=-0.11, p=0.00059
cor=-0.099, p=0.0016
bicor=-0.07, p=0.025
cor=-0.058, p=0.065
bicor=0.15, p=2.2e-06
cor=0.15, p=1.7e-06
NDUFV1_2 - M3 brown
Kruskal-W. p = 0.0069
Tukey AD/CT p = 0.02
AD/Asym 0.028
Asym/CT 0.82
bicor=-0.28, p=8.6e-20
cor=-0.23, p=1.7e-13
bicor=-0.098, p=0.0019
cor=-0.08, p=0.011
bicor=-0.053, p=0.095
cor=-0.033, p=0.29
bicor=0.14, p=4.1e-06
cor=0.14, p=7.9e-06
PLP1_1 - M2 blue
Kruskal-W. p = 0.92
Tukey AD/CT p = 0.91
AD/Asym 0.98
Asym/CT 0.97
bicor=0.18, p=2.1e-08
cor=0.13, p=3.6e-05
bicor=0.0074, p=0.81
cor=0.006, p=0.85
bicor=0.003, p=0.92
cor=-0.014, p=0.66
bicor=0.017, p=0.59
cor=-0.02, p=0.53
PLP1_2 - M2 blue
Kruskal-W. p = 0.99
Tukey AD/CT p = 1
AD/Asym 0.99
Asym/CT 0.99
bicor=0.18, p=2.4e-08
cor=0.14, p=8.7e-06
bicor=-0.0056, p=0.86
cor=-0.0085, p=0.79
bicor=-0.0017, p=0.96
cor=-0.022, p=0.48
bicor=0.019, p=0.56
cor=-0.012, p=0.7
PTK2B_1 - M1 turquoise
Kruskal-W. p = 0.076
Tukey AD/CT p = 0.15
AD/Asym 0.13
Asym/CT 0.95
bicor=-0.11, p=4e-04
cor=-0.16, p=3.5e-07
bicor=-0.044, p=0.16
cor=-0.053, p=0.092
bicor=-0.077, p=0.014
cor=-0.078, p=0.013
bicor=0.093, p=0.0031
cor=0.13, p=3.4e-05
RPL10A_2 - M7 black
Kruskal-W. p = 0.071
Tukey AD/CT p = 0.094
AD/Asym 0.21
Asym/CT 0.77
bicor=0.038, p=0.23
cor=0.025, p=0.43
bicor=-0.084, p=0.0073
cor=-0.053, p=0.092
bicor=-0.063, p=0.043
cor=-0.045, p=0.15
bicor=0.097, p=0.002
cor=0.062, p=0.049
RPL35_1 - M7 black
Kruskal-W. p = 1
Tukey AD/CT p = 1
AD/Asym 1
Asym/CT 1
bicor=0.08, p=0.011
cor=0.057, p=0.071
bicor=-0.0044, p=0.89
cor=-0.013, p=0.68
bicor=-0.032, p=0.3
cor=-0.03, p=0.34
bicor=-0.0025, p=0.94
cor=-0.0054, p=0.86
Protein Abun
Control
AsymAD
AD
Tau 8 Peptide Avg.
CERAD Score
Braak Score
MMSE Cognitive Score

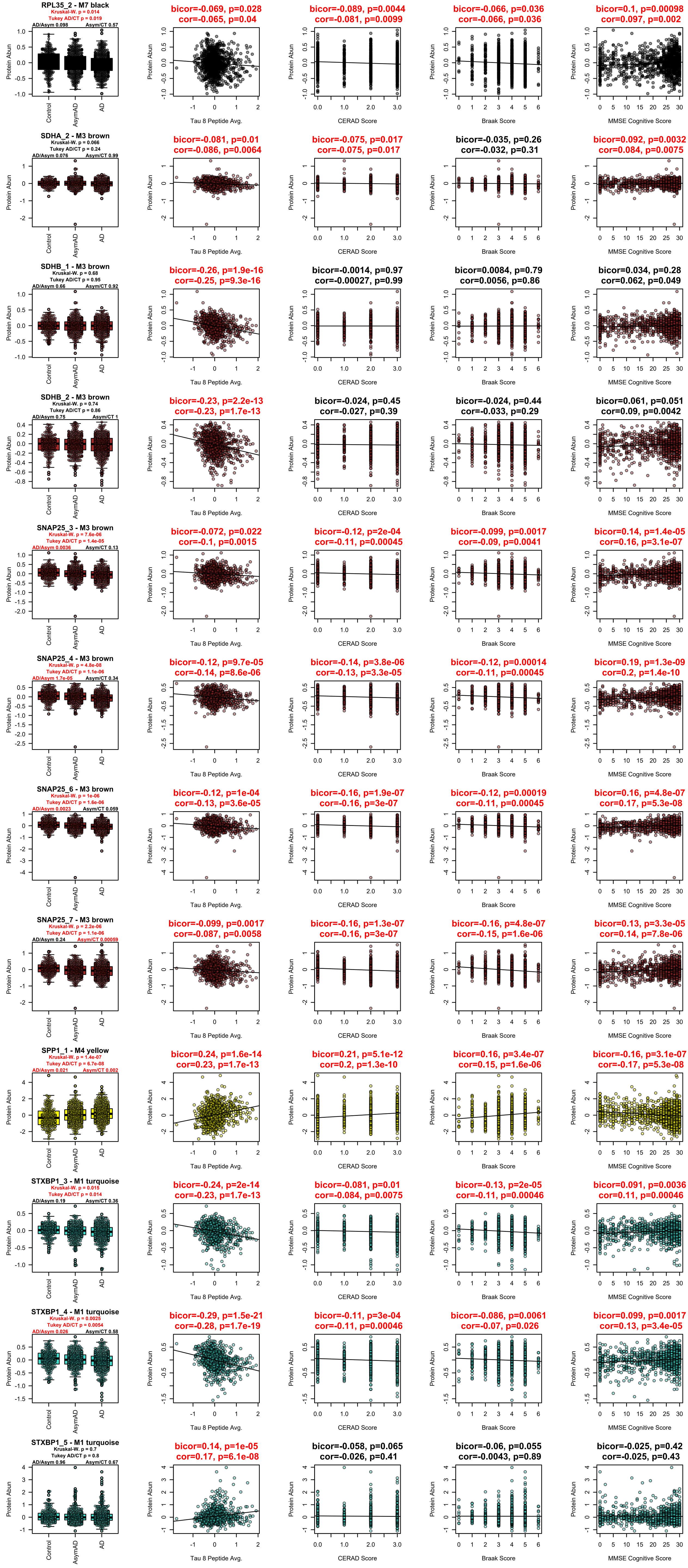
RPL35_2 - M7 black
Kruskal-W. p = 0.014
Tukey AD/CT p = 0.019
AD/Asym 0.098
Asym/CT 0.57
bicor=-0.069, p=0.028
cor=-0.065, p=0.04
bicor=-0.089, p=0.0044
cor=-0.081, p=0.0099
bicor=-0.066, p=0.036
cor=-0.066, p=0.036
bicor=0.1, p=0.00098
cor=0.097, p=0.002
SDHA_2 - M3 brown
Kruskal-W. p = 0.066
Tukey AD/CT p = 0.24
AD/Asym 0.076
Asym/CT 0.99
bicor=-0.081, p=0.01
cor=-0.086, p=0.0064
bicor=-0.075, p=0.017
cor=-0.075, p=0.017
bicor=-0.035, p=0.26
cor=-0.032, p=0.31
bicor=0.092, p=0.0032
cor=0.084, p=0.0075
SDHB_1 - M3 brown
Kruskal-W. p = 0.68
Tukey AD/CT p = 0.95
AD/Asym 0.66
Asym/CT 0.92
bicor=-0.26, p=1.9e-16
cor=-0.25, p=9.3e-16
bicor=-0.0014, p=0.97
cor=-0.00027, p=0.99
bicor=0.0084, p=0.79
cor=0.0056, p=0.86
bicor=0.034, p=0.28
cor=0.062, p=0.049
SDHB_2 - M3 brown
Kruskal-W. p = 0.74
Tukey AD/CT p = 0.86
AD/Asym 0.75
Asym/CT 1
bicor=-0.23, p=2.2e-13
cor=-0.23, p=1.7e-13
bicor=-0.024, p=0.45
cor=-0.027, p=0.39
bicor=-0.024, p=0.44
cor=-0.033, p=0.29
bicor=0.061, p=0.051
cor=0.09, p=0.0042
SNAP25_3 - M3 brown
Kruskal-W. p = 7.6e-06
Tukey AD/CT p = 1.4e-05
AD/Asym 0.0036
Asym/CT 0.13
bicor=-0.072, p=0.022
cor=-0.1, p=0.0015
bicor=-0.12, p=2e-04
cor=-0.11, p=0.00045
bicor=-0.099, p=0.0017
cor=-0.09, p=0.0041
bicor=0.14, p=1.4e-05
cor=0.16, p=3.1e-07
SNAP25_4 - M3 brown
Kruskal-W. p = 4.8e-08
Tukey AD/CT p = 1.1e-06
AD/Asym 1.7e-05
Asym/CT 0.34
bicor=-0.12, p=9.7e-05
cor=-0.14, p=8.6e-06
bicor=-0.14, p=3.8e-06
cor=-0.13, p=3.3e-05
bicor=-0.12, p=0.00014
cor=-0.11, p=0.00045
bicor=0.19, p=1.3e-09
cor=0.2, p=1.4e-10
SNAP25_6 - M3 brown
Kruskal-W. p = 1e-06
Tukey AD/CT p = 1.6e-06
AD/Asym 0.0023
Asym/CT 0.059
bicor=-0.12, p=1e-04
cor=-0.13, p=3.6e-05
bicor=-0.16, p=1.9e-07
cor=-0.16, p=3e-07
bicor=-0.12, p=0.00019
cor=-0.11, p=0.00045
bicor=0.16, p=4.8e-07
cor=0.17, p=5.3e-08
SNAP25_7 - M3 brown
Kruskal-W. p = 2.2e-06
Tukey AD/CT p = 1.1e-06
AD/Asym 0.24
Asym/CT 0.00059
bicor=-0.099, p=0.0017
cor=-0.087, p=0.0058
bicor=-0.16, p=1.3e-07
cor=-0.16, p=3e-07
bicor=-0.16, p=4.8e-07
cor=-0.15, p=1.6e-06
bicor=0.13, p=3.3e-05
cor=0.14, p=7.8e-06
SPP1_1 - M4 yellow
Kruskal-W. p = 1.4e-07
Tukey AD/CT p = 6.7e-08
AD/Asym 0.021
Asym/CT 0.002
bicor=0.24, p=1.6e-14
cor=0.23, p=1.7e-13
bicor=0.21, p=5.1e-12
cor=0.2, p=1.3e-10
bicor=0.16, p=3.4e-07
cor=0.15, p=1.6e-06
bicor=-0.16, p=3.1e-07
cor=-0.17, p=5.3e-08
STXBP1_3 - M1 turquoise
Kruskal-W. p = 0.015
Tukey AD/CT p = 0.014
AD/Asym 0.19
Asym/CT 0.36
bicor=-0.24, p=2e-14
cor=-0.23, p=1.7e-13
bicor=-0.081, p=0.01
cor=-0.084, p=0.0075
bicor=-0.13, p=2e-05
cor=-0.11, p=0.00046
bicor=0.091, p=0.0036
cor=0.11, p=0.00046
STXBP1_4 - M1 turquoise
Kruskal-W. p = 0.0025
Tukey AD/CT p = 0.0054
AD/Asym 0.026
Asym/CT 0.58
bicor=-0.29, p=1.5e-21
cor=-0.28, p=1.7e-19
bicor=-0.11, p=3e-04
cor=-0.11, p=0.00046
bicor=-0.086, p=0.0061
cor=-0.07, p=0.026
bicor=0.099, p=0.0017
cor=0.13, p=3.4e-05
STXBP1_5 - M1 turquoise
Kruskal-W. p = 0.7
Tukey AD/CT p = 0.8
AD/Asym 0.96
Asym/CT 0.67
bicor=0.14, p=1e-05
cor=0.17, p=6.1e-08
bicor=-0.058, p=0.065
cor=-0.026, p=0.41
bicor=-0.06, p=0.055
cor=-0.0043, p=0.89
bicor=-0.025, p=0.42
cor=-0.025, p=0.43
Protein Abun
Control
AsymAD
AD
Tau 8 Peptide Avg.
CERAD Score
Braak Score
MMSE Cognitive Score

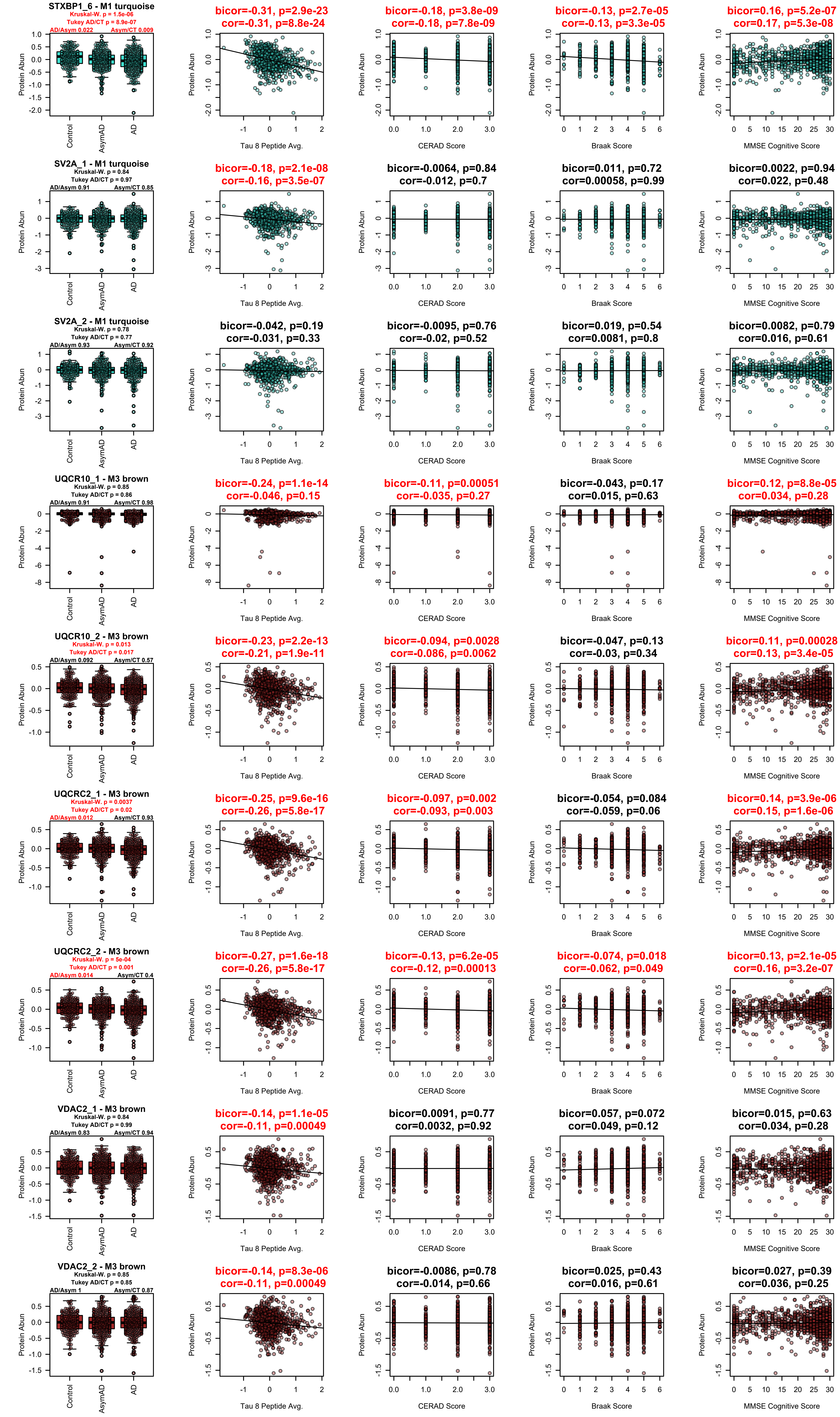
STXBP1_6 - M1 turquoise
Kruskal-W. p = 1.5e-06
Tukey AD/CT p = 8.9e-07
AD/Asym 0.022
Asym/CT 0.009
bicor=-0.31, p=2.9e-23
cor=-0.31, p=8.8e-24
bicor=-0.18, p=3.8e-09
cor=-0.18, p=7.8e-09
bicor=-0.13, p=2.7e-05
cor=-0.13, p=3.3e-05
bicor=0.16, p=5.2e-07
cor=0.17, p=5.3e-08
SV2A_1 - M1 turquoise
Kruskal-W. p = 0.84
Tukey AD/CT p = 0.97
AD/Asym 0.91
Asym/CT 0.85
bicor=-0.18, p=2.1e-08
cor=-0.16, p=3.5e-07
bicor=-0.0064, p=0.84
cor=-0.012, p=0.7
bicor=0.011, p=0.72
cor=0.00058, p=0.99
bicor=0.0022, p=0.94
cor=0.022, p=0.48
SV2A_2 - M1 turquoise
Kruskal-W. p = 0.78
Tukey AD/CT p = 0.77
AD/Asym 0.93
Asym/CT 0.92
bicor=-0.042, p=0.19
cor=-0.031, p=0.33
bicor=-0.0095, p=0.76
cor=-0.02, p=0.52
bicor=0.019, p=0.54
cor=0.0081, p=0.8
bicor=0.0082, p=0.79
cor=0.016, p=0.61
UQCR10_1 - M3 brown
Kruskal-W. p = 0.85
Tukey AD/CT p = 0.86
AD/Asym 0.91
Asym/CT 0.98
bicor=-0.24, p=1.1e-14
cor=-0.046, p=0.15
bicor=-0.11, p=0.00051
cor=-0.035, p=0.27
bicor=-0.043, p=0.17
cor=0.015, p=0.63
bicor=0.12, p=8.8e-05
cor=0.034, p=0.28
UQCR10_2 - M3 brown
Kruskal-W. p = 0.013
Tukey AD/CT p = 0.017
AD/Asym 0.092
Asym/CT 0.57
bicor=-0.23, p=2.2e-13
cor=-0.21, p=1.9e-11
bicor=-0.094, p=0.0028
cor=-0.086, p=0.0062
bicor=-0.047, p=0.13
cor=-0.03, p=0.34
bicor=0.11, p=0.00028
cor=0.13, p=3.4e-05
UQCRC2_1 - M3 brown
Kruskal-W. p = 0.0037
Tukey AD/CT p = 0.02
AD/Asym 0.012
Asym/CT 0.93
bicor=-0.25, p=9.6e-16
cor=-0.26, p=5.8e-17
bicor=-0.097, p=0.002
cor=-0.093, p=0.003
bicor=-0.054, p=0.084
cor=-0.059, p=0.06
bicor=0.14, p=3.9e-06
cor=0.15, p=1.6e-06
UQCRC2_2 - M3 brown
Kruskal-W. p = 5e-04
Tukey AD/CT p = 0.001
AD/Asym 0.014
Asym/CT 0.4
bicor=-0.27, p=1.6e-18
cor=-0.26, p=5.8e-17
bicor=-0.13, p=6.2e-05
cor=-0.12, p=0.00013
bicor=-0.074, p=0.018
cor=-0.062, p=0.049
bicor=0.13, p=2.1e-05
cor=0.16, p=3.2e-07
VDAC2_1 - M3 brown
Kruskal-W. p = 0.84
Tukey AD/CT p = 0.99
AD/Asym 0.83
Asym/CT 0.94
bicor=-0.14, p=1.1e-05
cor=-0.11, p=0.00049
bicor=0.0091, p=0.77
cor=0.0032, p=0.92
bicor=0.057, p=0.072
cor=0.049, p=0.12
bicor=0.015, p=0.63
cor=0.034, p=0.28
VDAC2_2 - M3 brown
Kruskal-W. p = 0.85
Tukey AD/CT p = 0.85
AD/Asym 1
Asym/CT 0.87
bicor=-0.14, p=8.3e-06
cor=-0.11, p=0.00049
bicor=-0.0086, p=0.78
cor=-0.014, p=0.66
bicor=0.025, p=0.43
cor=0.016, p=0.61
bicor=0.027, p=0.39
cor=0.036, p=0.25
Protein Abun
Control
AsymAD
AD
Tau 8 Peptide Avg.
CERAD Score
Braak Score
MMSE Cognitive Score

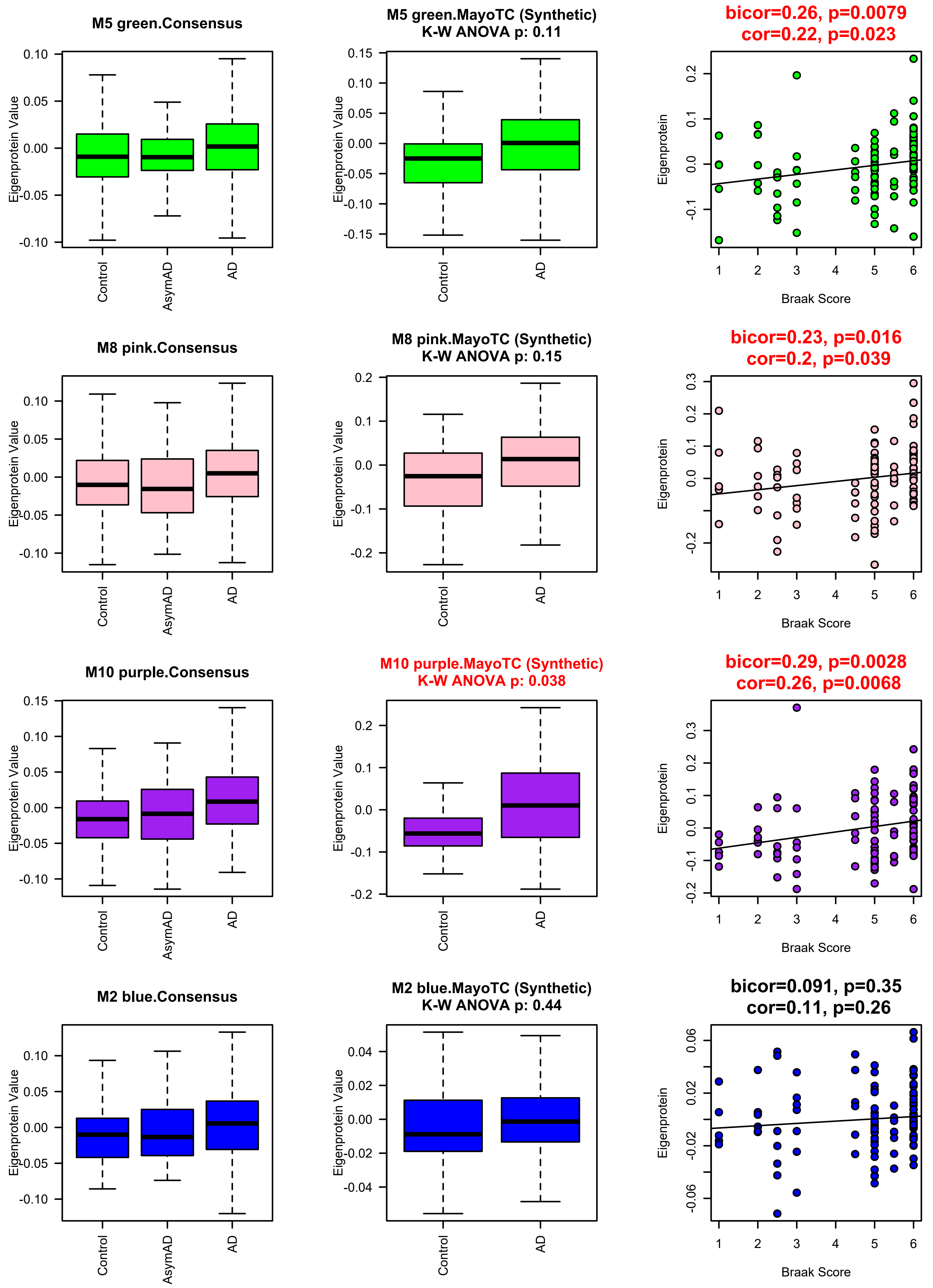

M5 green.Consensus
Eigenprotein Value
Control
AsymAD
AD
M5 green.MayoTC (Synthetic)
K-W ANOVA p: 0.11
bicor=0.26, p=0.0079
cor=0.22, p=0.023
Eigenprotein
Braak Score
M8 pink.Consensus
M8 pink.MayoTC (Synthetic)
K-W ANOVA p: 0.15
bicor=0.23, p=0.016
cor=0.2, p=0.039
M10 purple.Consensus
M10 purple.MayoTC (Synthetic)
K-W ANOVA p: 0.038
bicor=0.29, p=0.0028
cor=0.26, p=0.0068
M2 blue.Consensus
M2 blue.MayoTC (Synthetic)
K-W ANOVA p: 0.44
bicor=0.091, p=0.35
cor=0.11, p=0.26

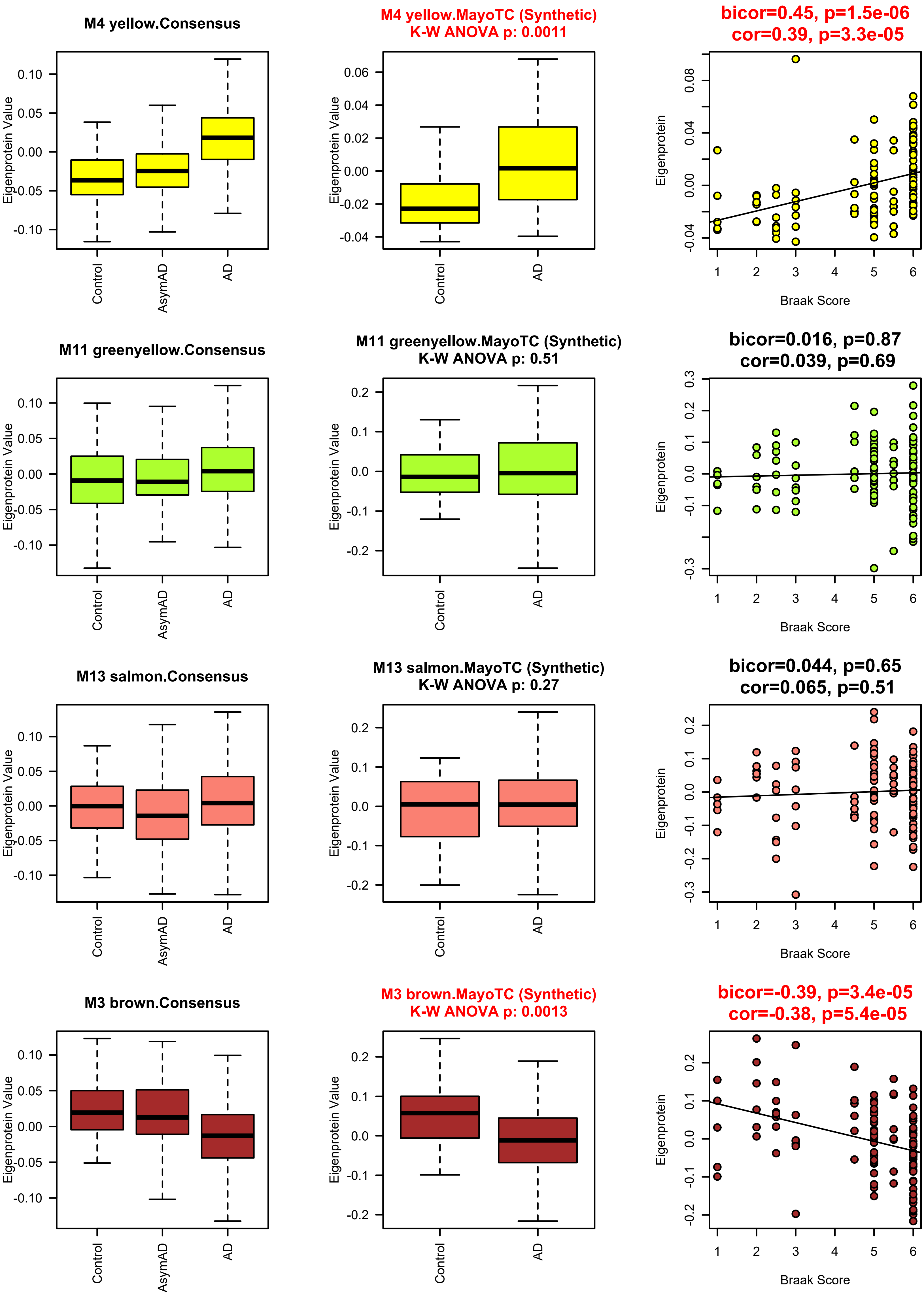
M4 yellow.Consensus
Eigenprotein Value
Control
AsymAD
AD
M4 yellow.MayoTC (Synthetic)
K-W ANOVA p: 0.0011
bicor=0.45, p=1.5e-06
cor=0.39, p=3.3e-05
Eigenprotein
Braak Score
M11 greenyellow.Consensus
M11 greenyellow.MayoTC (Synthetic)
K-W ANOVA p: 0.51
bicor=0.016, p=0.87
cor=0.039, p=0.69
M13 salmon.Consensus
M13 salmon.MayoTC (Synthetic)
K-W ANOVA p: 0.27
bicor=0.044, p=0.65
cor=0.065, p=0.51
M3 brown.Consensus
M3 brown.MayoTC (Synthetic)
K-W ANOVA p: 0.0013
bicor=-0.39, p=3.4e-05
cor=-0.38, p=5.4e-05

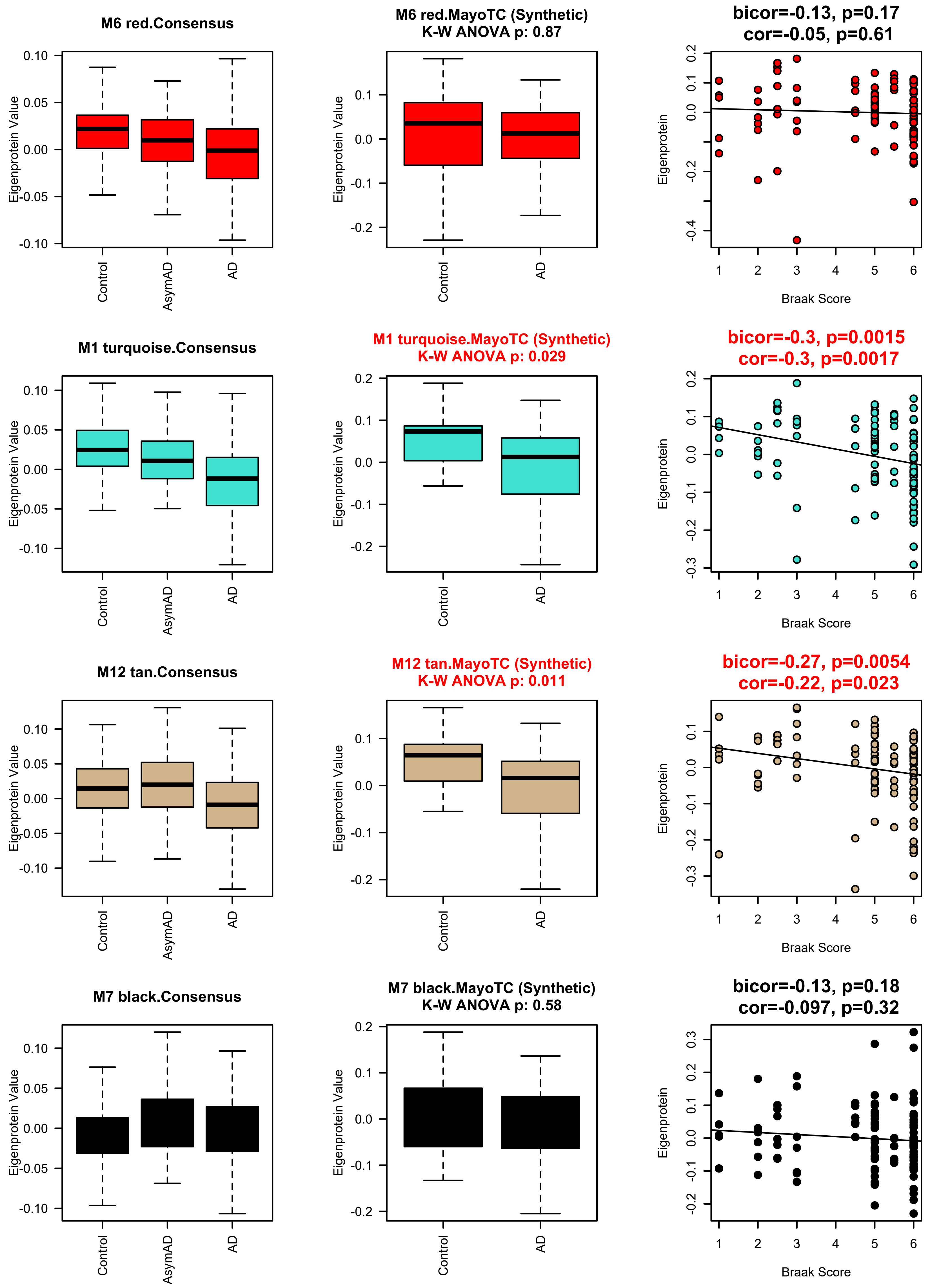
M6 red.Consensus
Eigenprotein Value
Control
AsymAD
AD
M6 red.MayoTC (Synthetic)
K-W ANOVA p: 0.87
Eigenprotein Value
Control
AD
bicor=-0.13, p=0.17
cor=-0.05, p=0.61
Eigenprotein
Braak Score
M1 turquoise.Consensus
Eigenprotein Value
Control
AsymAD
AD
M1 turquoise.MayoTC (Synthetic)
K-W ANOVA p: 0.029
Eigenprotein Value
Control
AD
bicor=-0.3, p=0.0015
cor=-0.3, p=0.0017
Eigenprotein
Braak Score
M12 tan.Consensus
Eigenprotein Value
Control
AsymAD
AD
M12 tan.MayoTC (Synthetic)
K-W ANOVA p: 0.011
Eigenprotein Value
Control
AD
bicor=-0.27, p=0.0054
cor=-0.22, p=0.023
Eigenprotein
Braak Score
M7 black.Consensus
Eigenprotein Value
Control
AsymAD
AD
M7 black.MayoTC (Synthetic)
K-W ANOVA p: 0.58
Eigenprotein Value
Control
AD
bicor=-0.13, p=0.18
cor=-0.097, p=0.32
Eigenprotein
Braak Score

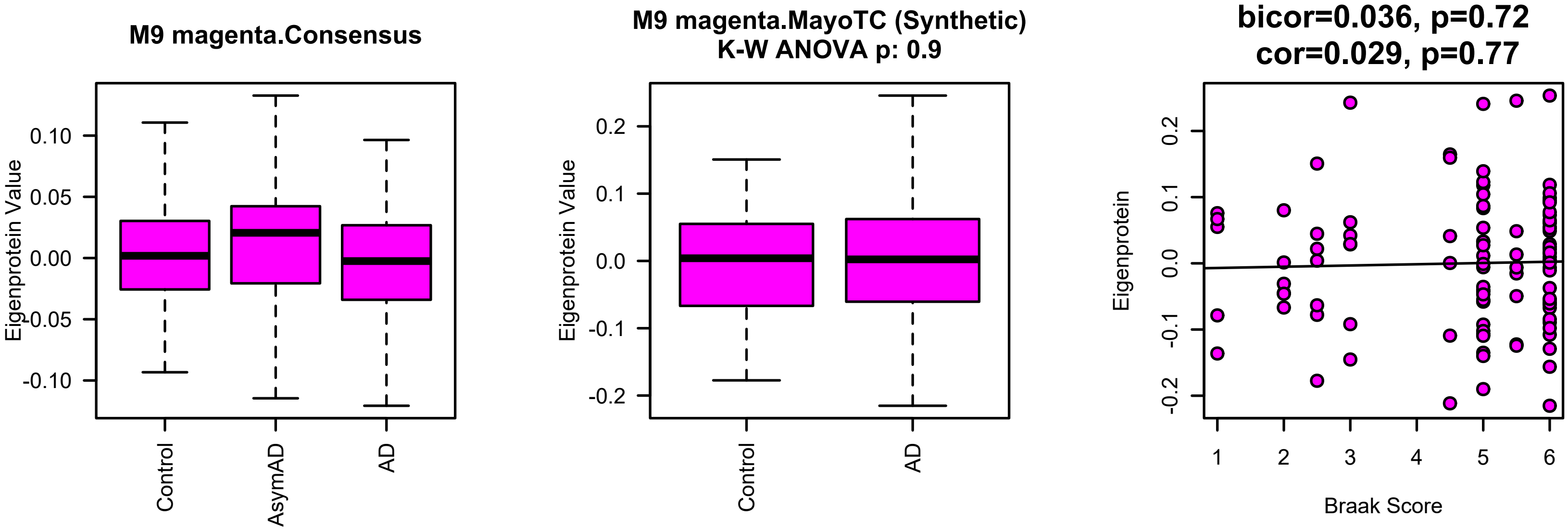

M9 magenta.Consensus
Eigenprotein Value
0.10
0.05
0.00
-0.05
-0.10
Control
AsymAD
AD
M9 magenta.MayoTC (Synthetic)
K-W ANOVA p: 0.9
Eigenprotein Value
0.2
0.1
0.0
-0.1
-0.2
Control
AD
bicor=0.036, p=0.72
cor=0.029, p=0.77
Eigenprotein
0.2
0.1
0.0
-0.1
-0.2
1
2
3
4
5
6
Braak Score

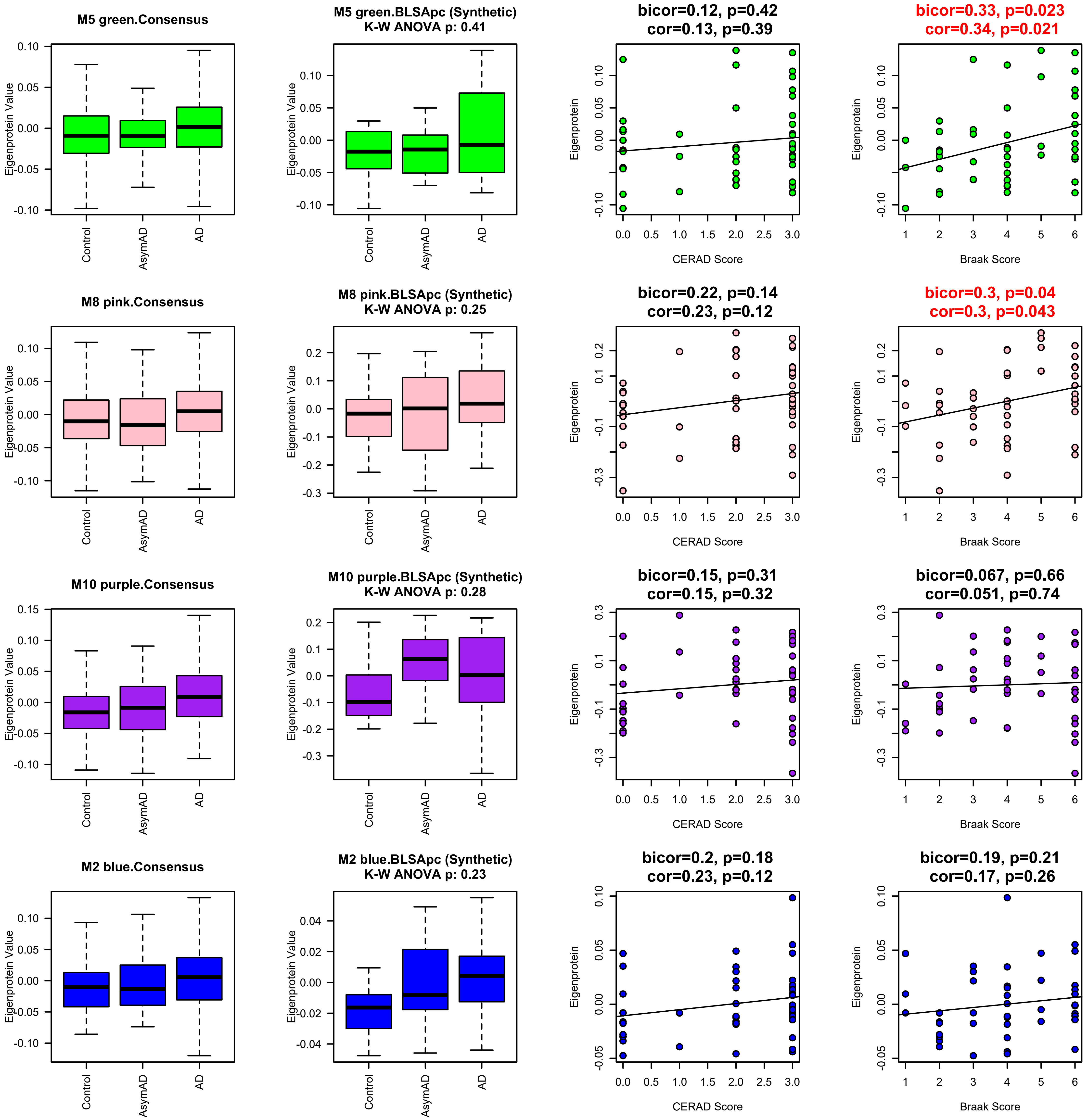
M5 green.Consensus
M5 green.BLSApc (Synthetic)
K-W ANOVA p: 0.41
bicor=0.12, p=0.42
cor=0.13, p=0.39
bicor=0.33, p=0.023
cor=0.34, p=0.021
M8 pink.Consensus
M8 pink.BLSApc (Synthetic)
K-W ANOVA p: 0.25
bicor=0.22, p=0.14
cor=0.23, p=0.12
bicor=0.3, p=0.04
cor=0.3, p=0.043
M10 purple.Consensus
M10 purple.BLSApc (Synthetic)
K-W ANOVA p: 0.28
bicor=0.15, p=0.31
cor=0.15, p=0.32
bicor=0.067, p=0.66
cor=0.051, p=0.74
M2 blue.Consensus
M2 blue.BLSApc (Synthetic)
K-W ANOVA p: 0.23
bicor=0.2, p=0.18
cor=0.23, p=0.12
bicor=0.19, p=0.21
cor=0.17, p=0.26
Eigenprotein Value
Eigenprotein
Control
AsymAD
AD
CERAD Score
Braak Score

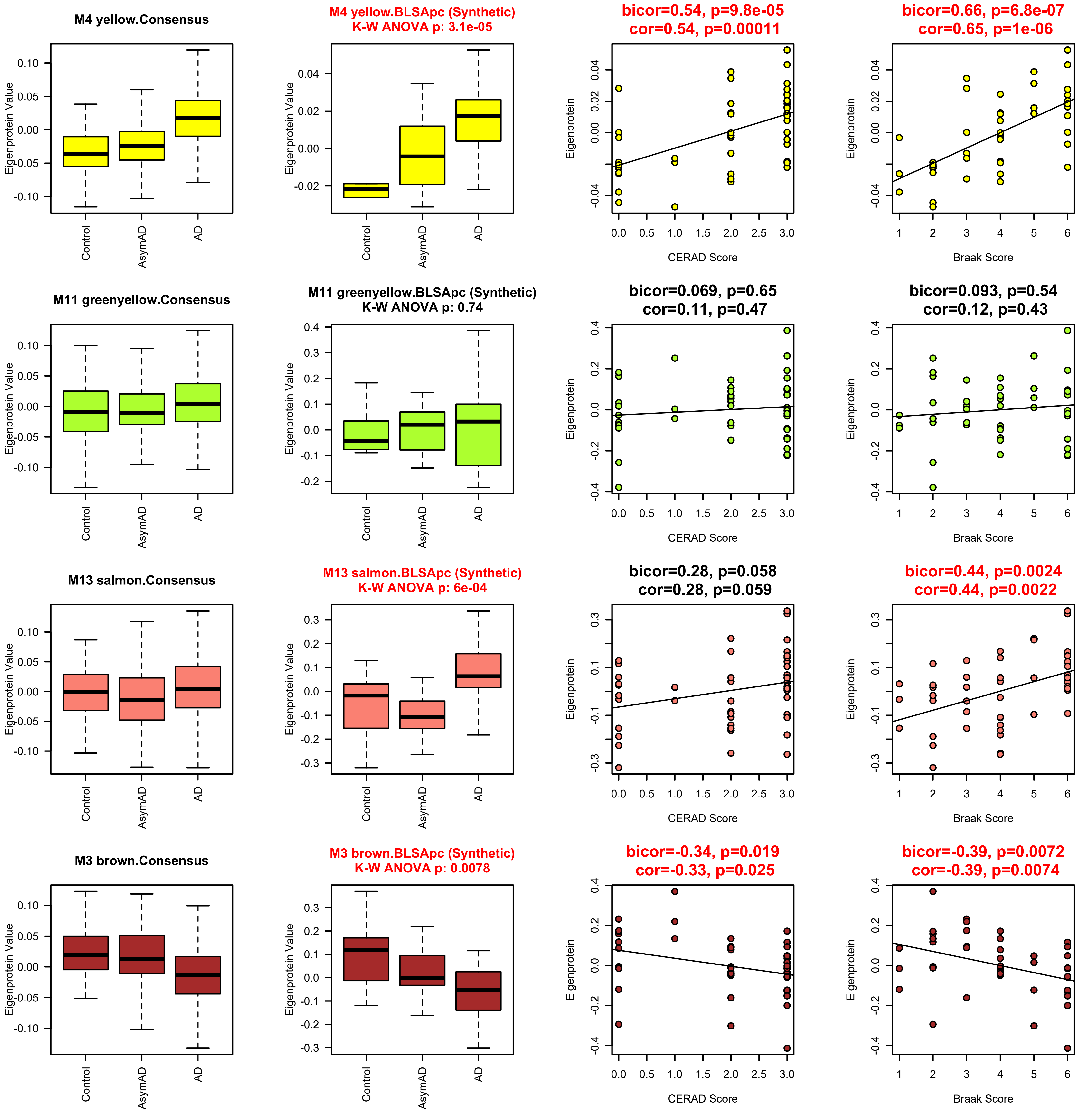

M4 yellow.Consensus
M4 yellow.BLSApc (Synthetic)
K-W ANOVA p: 3.1e-05
bicor=0.54, p=9.8e-05
cor=0.54, p=0.00011
bicor=0.66, p=6.8e-07
cor=0.65, p=1e-06
M11 greenyellow.Consensus
M11 greenyellow.BLSApc (Synthetic)
K-W ANOVA p: 0.74
bicor=0.069, p=0.65
cor=0.11, p=0.47
bicor=0.093, p=0.54
cor=0.12, p=0.43
M13 salmon.Consensus
M13 salmon.BLSApc (Synthetic)
K-W ANOVA p: 6e-04
bicor=0.28, p=0.058
cor=0.28, p=0.059
bicor=0.44, p=0.0024
cor=0.44, p=0.0022
M3 brown.Consensus
M3 brown.BLSApc (Synthetic)
K-W ANOVA p: 0.0078
bicor=-0.34, p=0.019
cor=-0.33, p=0.025
bicor=-0.39, p=0.0072
cor=-0.39, p=0.0074
Eigenprotein Value
Eigenprotein
Control
AsymAD
AD
CERAD Score
Braak Score

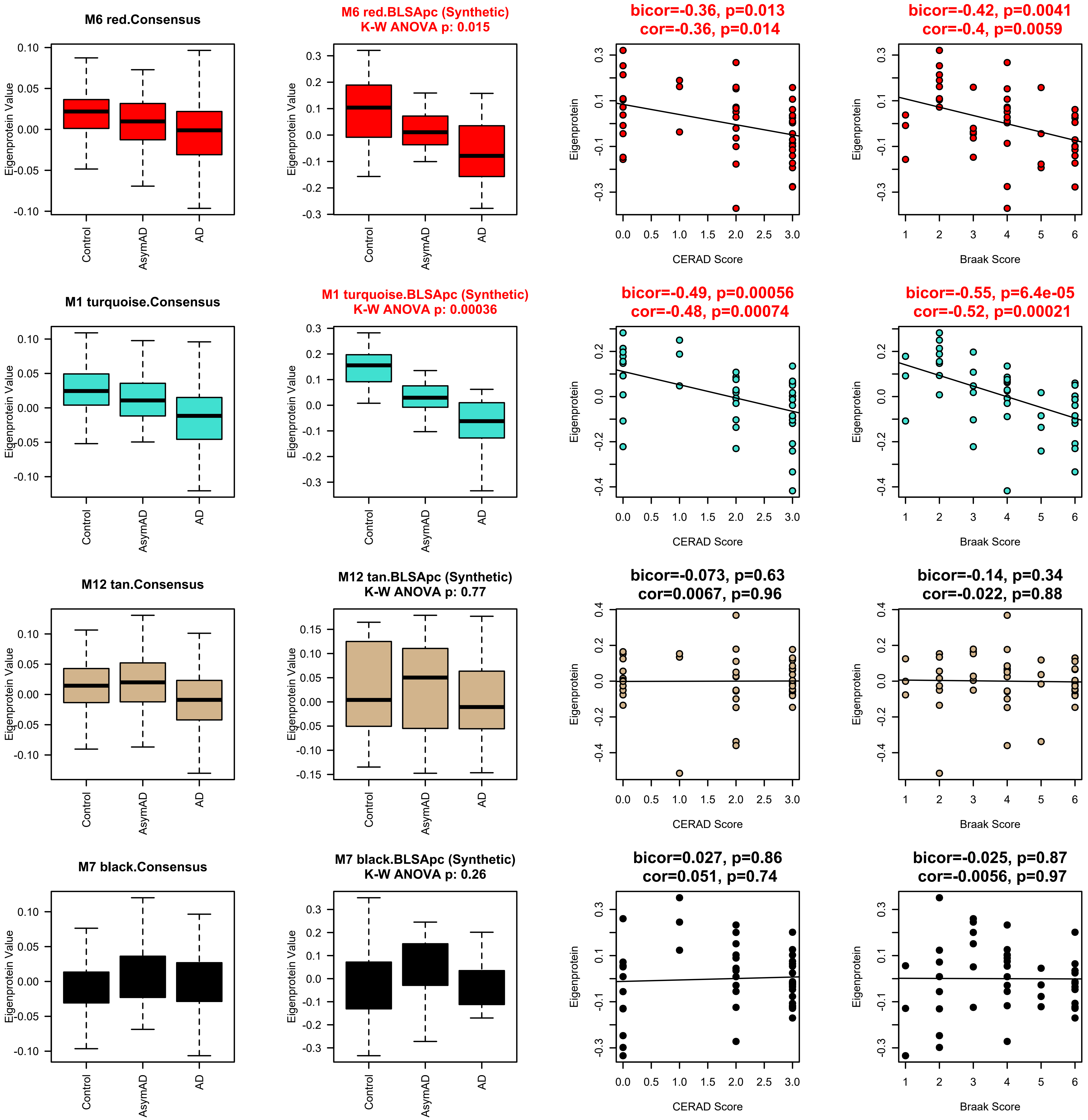

M6 red.Consensus
M6 red.BLSApc (Synthetic)
K-W ANOVA p: 0.015
bicor=-0.36, p=0.013
cor=-0.36, p=0.014
bicor=-0.42, p=0.0041
cor=-0.4, p=0.0059
M1 turquoise.Consensus
M1 turquoise.BLSApc (Synthetic)
K-W ANOVA p: 0.00036
bicor=-0.49, p=0.00056
cor=-0.48, p=0.00074
bicor=-0.55, p=6.4e-05
cor=-0.52, p=0.00021
M12 tan.Consensus
M12 tan.BLSApc (Synthetic)
K-W ANOVA p: 0.77
bicor=-0.073, p=0.63
cor=0.0067, p=0.96
bicor=-0.14, p=0.34
cor=-0.022, p=0.88
M7 black.Consensus
M7 black.BLSApc (Synthetic)
K-W ANOVA p: 0.26
bicor=0.027, p=0.86
cor=0.051, p=0.74
bicor=-0.025, p=0.87
cor=-0.0056, p=0.97
Eigenprotein Value
Eigenprotein
Control
AsymAD
AD
CERAD Score
Braak Score

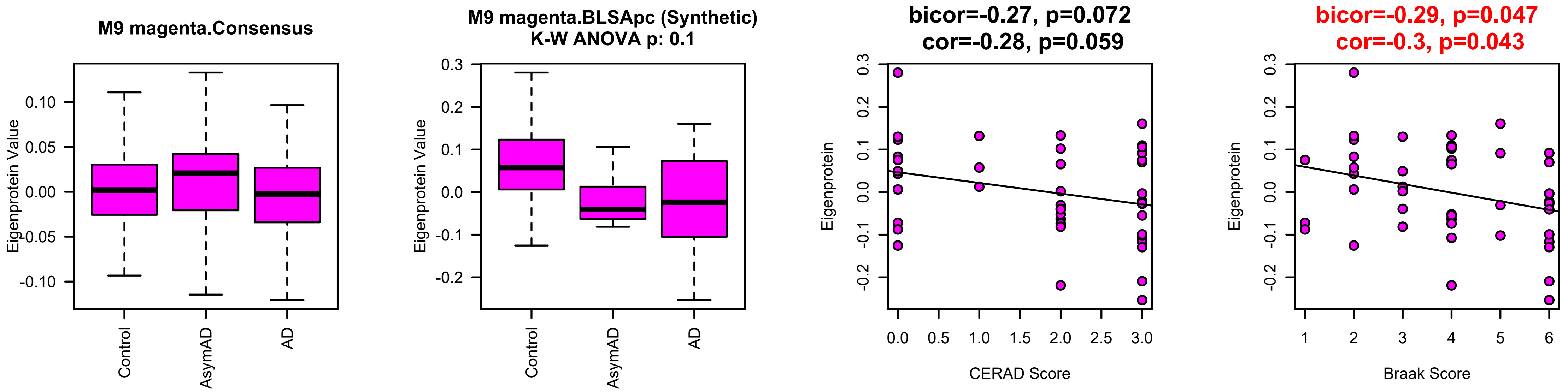
M9 magenta.Consensus
Eigenprotein Value
Control
AsymAD
AD
M9 magenta.BLSApc (Synthetic)
K-W ANOVA p: 0.1
bicor=-0.27, p=0.072
cor=-0.28, p=0.059
Eigenprotein
CERAD Score
bicor=-0.29, p=0.047
cor=-0.3, p=0.043
Braak Score

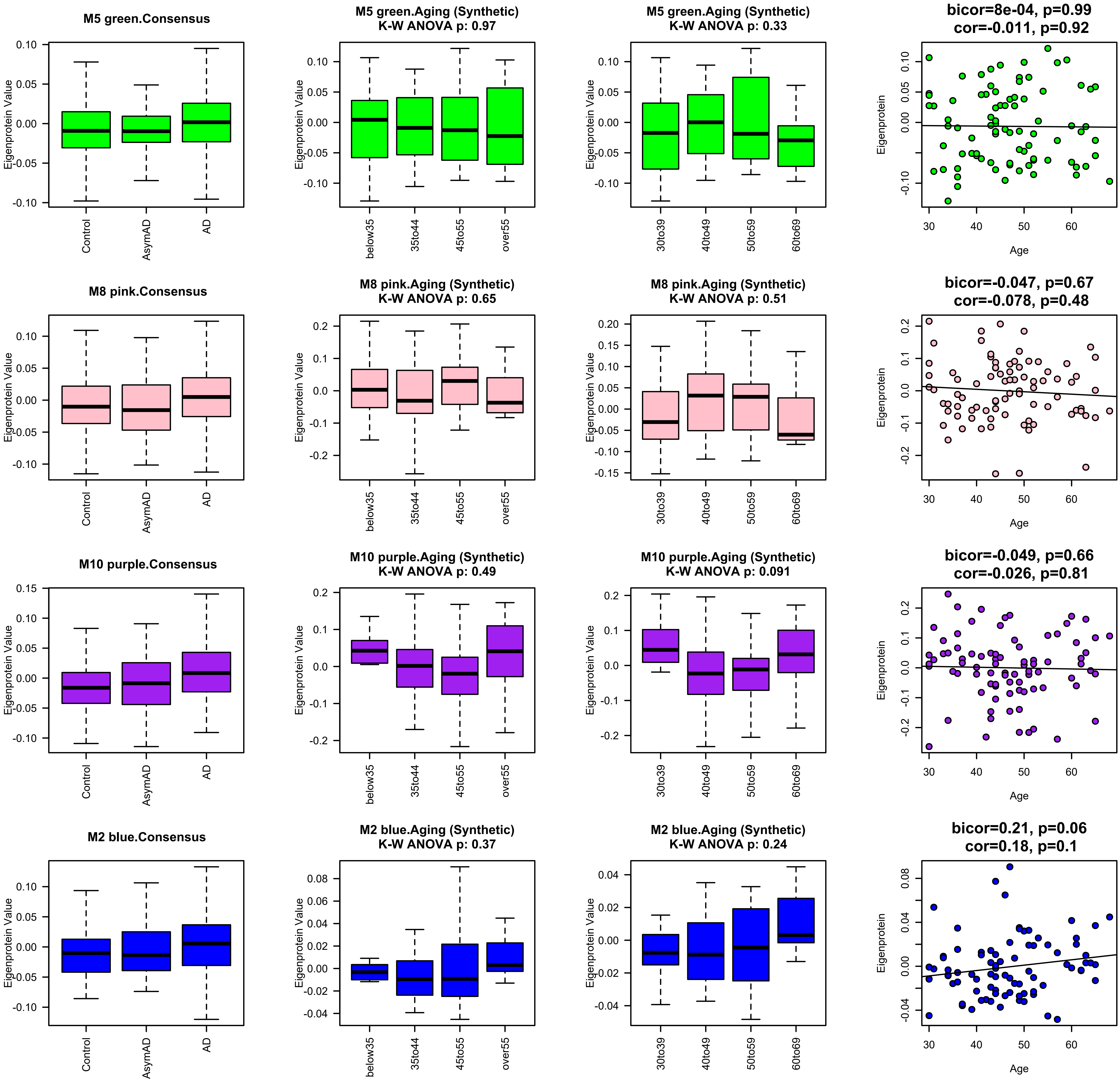

M5 green.Consensus
M5 green.Aging (Synthetic)
K-W ANOVA p: 0.97
M5 green.Aging (Synthetic)
K-W ANOVA p: 0.33
bicor=8e-04, p=0.99
cor=-0.011, p=0.92
M8 pink.Consensus
M8 pink.Aging (Synthetic)
K-W ANOVA p: 0.65
M8 pink.Aging (Synthetic)
K-W ANOVA p: 0.51
bicor=-0.047, p=0.67
cor=-0.078, p=0.48
M10 purple.Consensus
M10 purple.Aging (Synthetic)
K-W ANOVA p: 0.49
M10 purple.Aging (Synthetic)
K-W ANOVA p: 0.091
bicor=-0.049, p=0.66
cor=-0.026, p=0.81
M2 blue.Consensus
M2 blue.Aging (Synthetic)
K-W ANOVA p: 0.37
M2 blue.Aging (Synthetic)
K-W ANOVA p: 0.24
bicor=0.21, p=0.06
cor=0.18, p=0.1
Eigenprotein Value
Eigenprotein
Control
AsymAD
AD
below35
35to44
45to55
over55
30to39
40to49
50to59
60to69
Age

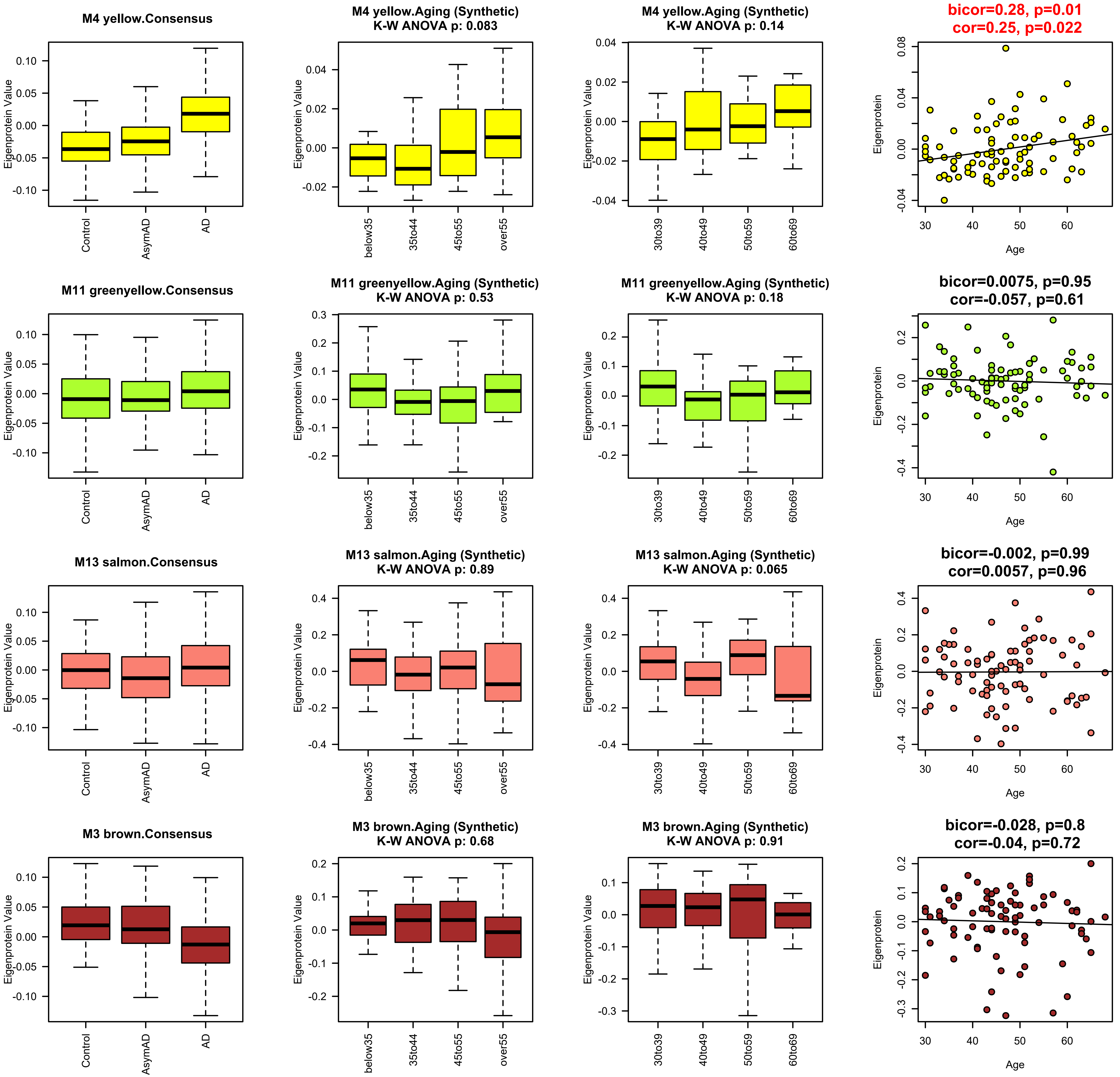
M4 yellow.Consensus
M4 yellow.Aging (Synthetic)
K-W ANOVA p: 0.083
M4 yellow.Aging (Synthetic)
K-W ANOVA p: 0.14
bicor=0.28, p=0.01
cor=0.25, p=0.022
M11 greenyellow.Consensus
M11 greenyellow.Aging (Synthetic)
K-W ANOVA p: 0.53
M11 greenyellow.Aging (Synthetic)
K-W ANOVA p: 0.18
bicor=0.0075, p=0.95
cor=-0.057, p=0.61
M13 salmon.Consensus
M13 salmon.Aging (Synthetic)
K-W ANOVA p: 0.89
M13 salmon.Aging (Synthetic)
K-W ANOVA p: 0.065
bicor=-0.002, p=0.99
cor=0.0057, p=0.96
M3 brown.Consensus
M3 brown.Aging (Synthetic)
K-W ANOVA p: 0.68
M3 brown.Aging (Synthetic)
K-W ANOVA p: 0.91
bicor=-0.028, p=0.8
cor=-0.04, p=0.72
Eigenprotein Value
Eigenprotein
Control
AsymAD
AD
below35
35to44
45to55
over55
30to39
40to49
50to59
60to69
Age

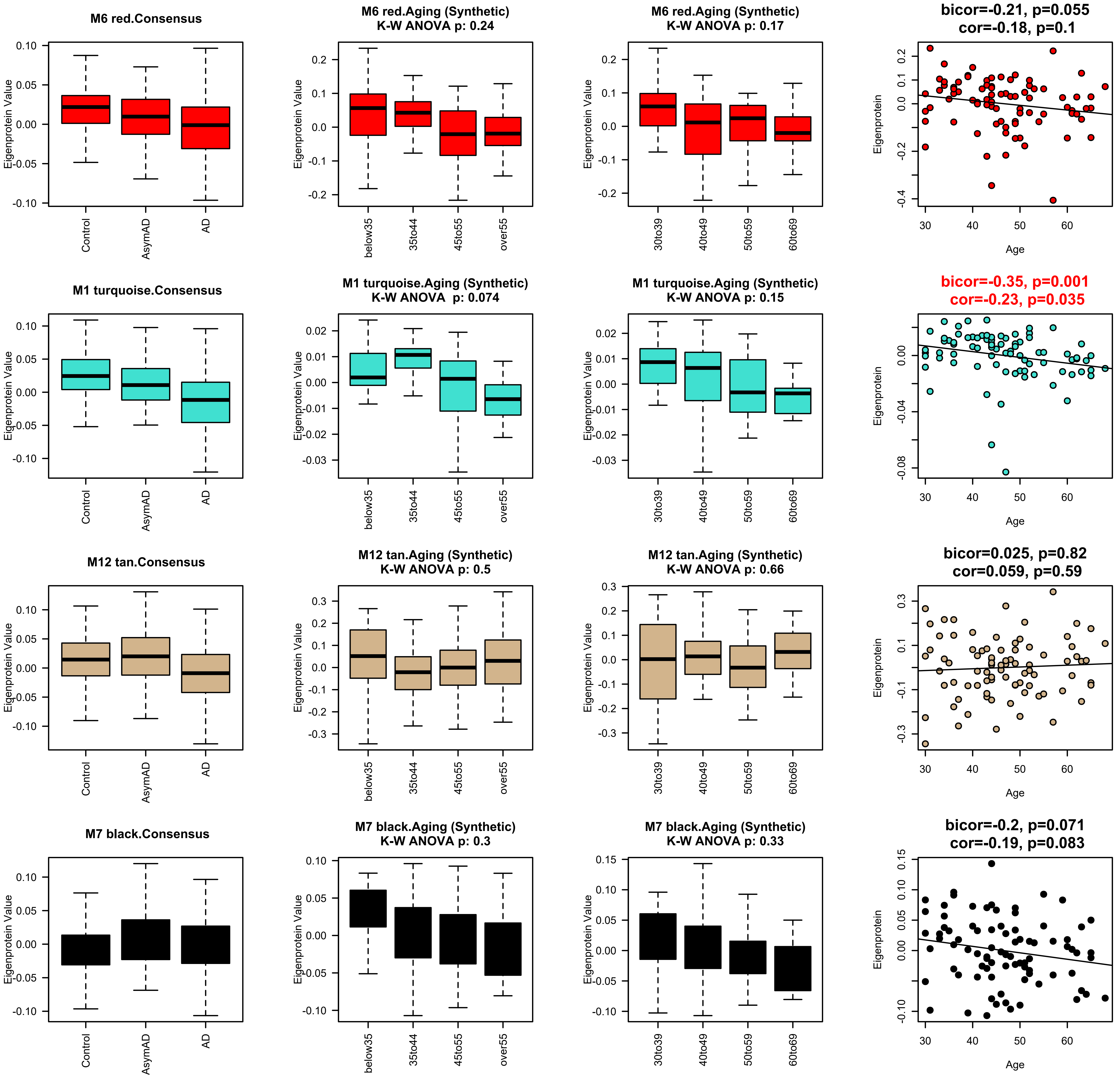
M6 red.Consensus
M6 red.Aging (Synthetic)
K-W ANOVA p: 0.24
M6 red.Aging (Synthetic)
K-W ANOVA p: 0.17
bicor=-0.21, p=0.055
cor=-0.18, p=0.1
M1 turquoise.Consensus
M1 turquoise.Aging (Synthetic)
K-W ANOVA p: 0.074
M1 turquoise.Aging (Synthetic)
K-W ANOVA p: 0.15
bicor=-0.35, p=0.001
cor=-0.23, p=0.035
M12 tan.Consensus
M12 tan.Aging (Synthetic)
K-W ANOVA p: 0.5
M12 tan.Aging (Synthetic)
K-W ANOVA p: 0.66
bicor=0.025, p=0.82
cor=0.059, p=0.59
M7 black.Consensus
M7 black.Aging (Synthetic)
K-W ANOVA p: 0.3
M7 black.Aging (Synthetic)
K-W ANOVA p: 0.33
bicor=-0.2, p=0.071
cor=-0.19, p=0.083
Eigenprotein Value
Eigenprotein
Control
AsymAD
AD
below35
35to44
45to55
over55
30to39
40to49
50to59
60to69
Age
30
40
50
60

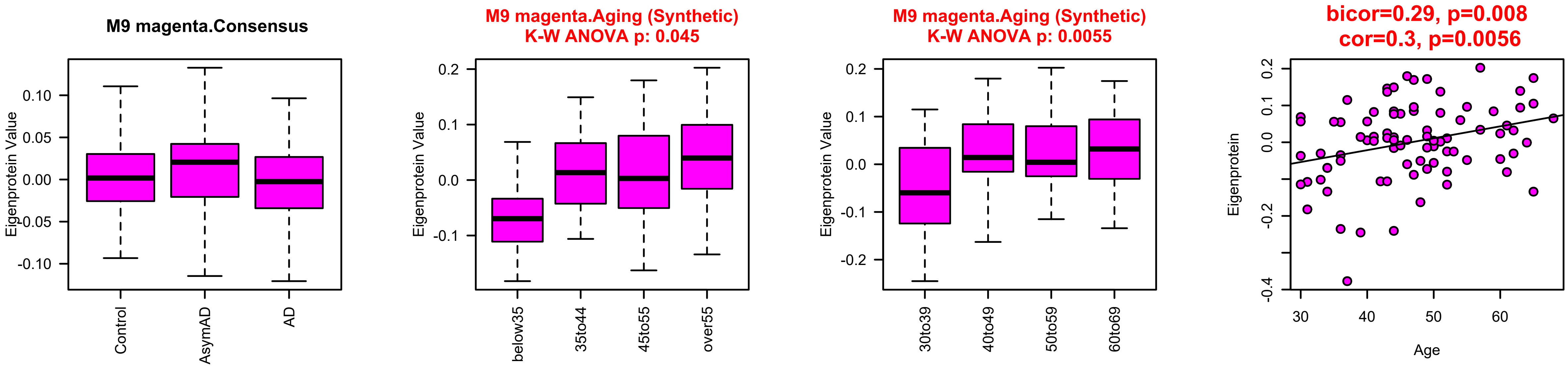
M9 magenta.Consensus
Eigenprotein Value
Control
AsymAD
AD
M9 magenta.Aging (Synthetic)
K-W ANOVA p: 0.045
below35
35to44
45to55
over55
M9 magenta.Aging (Synthetic)
K-W ANOVA p: 0.0055
30to39
40to49
50to59
60to69
bicor=0.29, p=0.008
cor=0.3, p=0.0056
Eigenprotein
Age

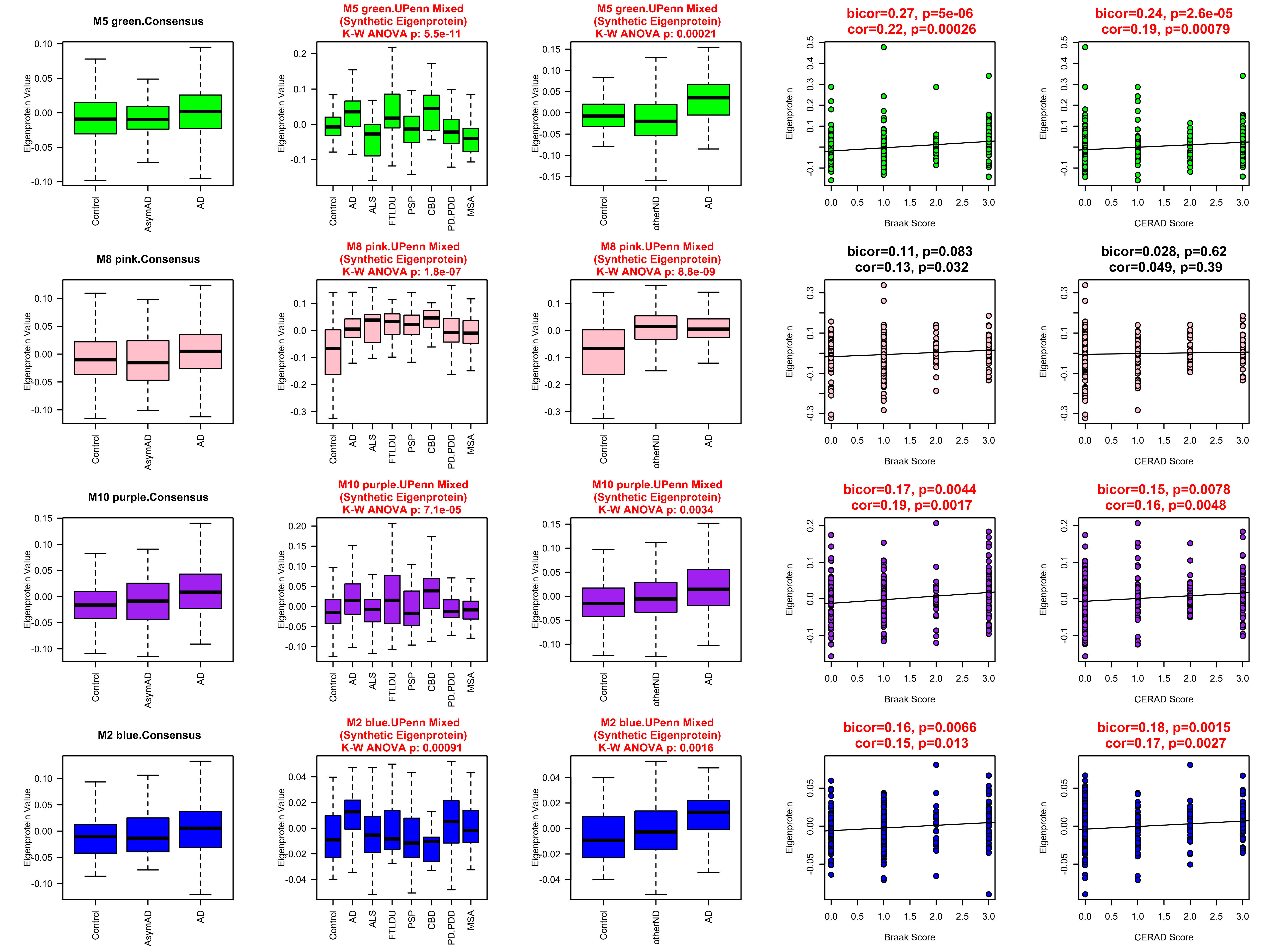
M5 green.Consensus
M5 green.UPenn Mixed (Synthetic Eigenprotein) K-W ANOVA p: 5.5e-11
M5 green.UPenn Mixed (Synthetic Eigenprotein) K-W ANOVA p: 0.00021
bicor=0.27, p=5e-06 cor=0.22, p=0.00026
bicor=0.24, p=2.6e-05 cor=0.19, p=0.00079
M8 pink.Consensus
M8 pink.UPenn Mixed (Synthetic Eigenprotein) K-W ANOVA p: 1.8e-07
M8 pink.UPenn Mixed (Synthetic Eigenprotein) K-W ANOVA p: 8.8e-09
bicor=0.11, p=0.083 cor=0.13, p=0.032
bicor=0.028, p=0.62 cor=0.049, p=0.39
M10 purple.Consensus
M10 purple.UPenn Mixed (Synthetic Eigenprotein) K-W ANOVA p: 7.1e-05
M10 purple.UPenn Mixed (Synthetic Eigenprotein) K-W ANOVA p: 0.0034
bicor=0.17, p=0.0044 cor=0.19, p=0.0017
bicor=0.15, p=0.0078 cor=0.16, p=0.0048
M2 blue.Consensus
M2 blue.UPenn Mixed (Synthetic Eigenprotein) K-W ANOVA p: 0.00091
M2 blue.UPenn Mixed (Synthetic Eigenprotein) K-W ANOVA p: 0.0016
bicor=0.16, p=0.0066 cor=0.15, p=0.013
bicor=0.18, p=0.0015 cor=0.17, p=0.0027
Eigenprotein Value
Eigenprotein
Control
AsymAD
AD
ALS
FTLDU
PSP
CBD
PD.PDD
MSA
otherND
Braak Score
CERAD Score

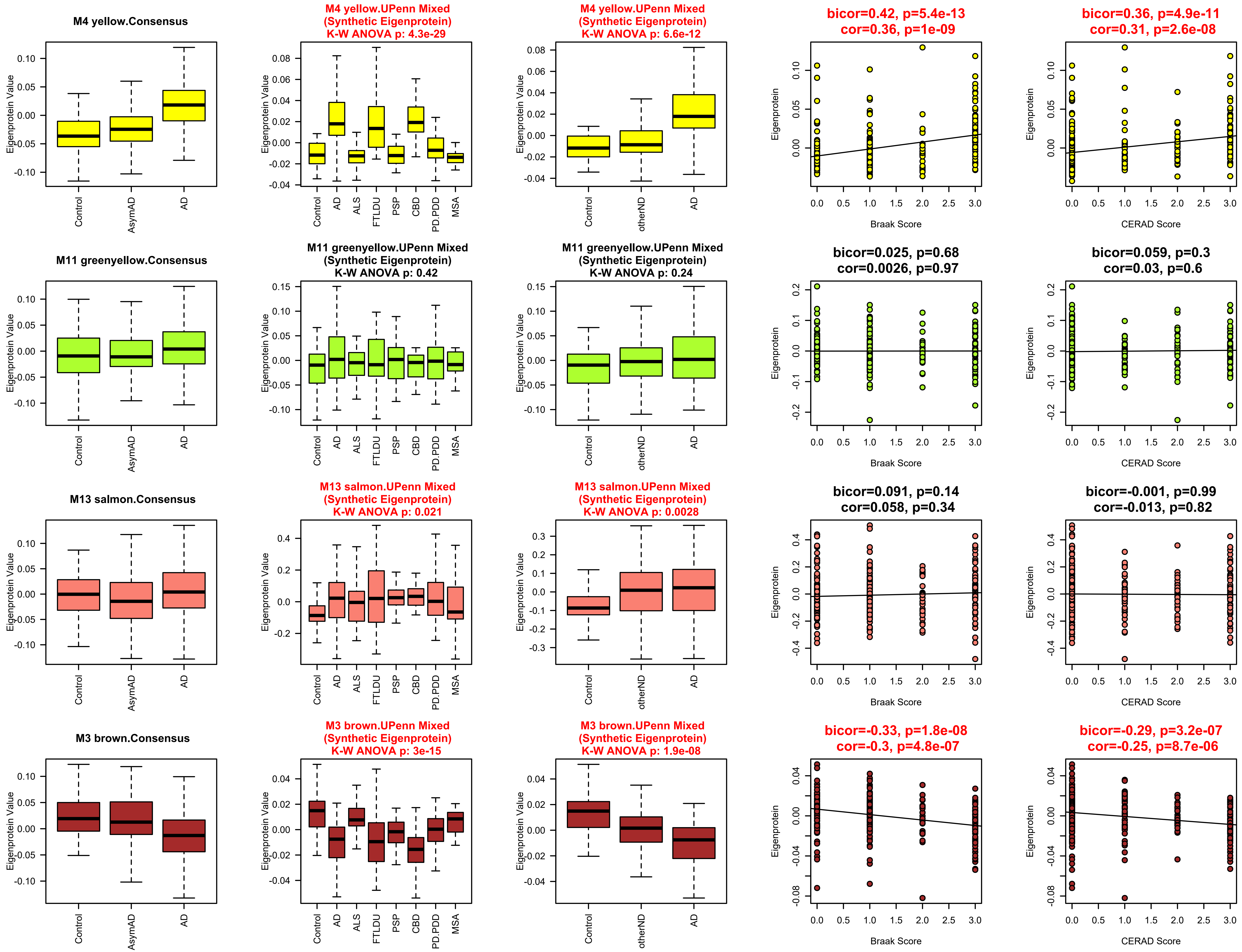

M4 yellow.Consensus
M4 yellow.UPenn Mixed (Synthetic Eigenprotein) K-W ANOVA p: 4.3e-29
M4 yellow.UPenn Mixed (Synthetic Eigenprotein) K-W ANOVA p: 6.6e-12
bicor=0.42, p=5.4e-13 cor=0.36, p=1e-09
bicor=0.36, p=4.9e-11 cor=0.31, p=2.6e-08
M11 greenyellow.Consensus
M11 greenyellow.UPenn Mixed (Synthetic Eigenprotein) K-W ANOVA p: 0.42
M11 greenyellow.UPenn Mixed (Synthetic Eigenprotein) K-W ANOVA p: 0.24
bicor=0.025, p=0.68 cor=0.0026, p=0.97
bicor=0.059, p=0.3 cor=0.03, p=0.6
M13 salmon.Consensus
M13 salmon.UPenn Mixed (Synthetic Eigenprotein) K-W ANOVA p: 0.021
M13 salmon.UPenn Mixed (Synthetic Eigenprotein) K-W ANOVA p: 0.0028
bicor=0.091, p=0.14 cor=0.058, p=0.34
bicor=-0.001, p=0.99 cor=-0.013, p=0.82
M3 brown.Consensus
M3 brown.UPenn Mixed (Synthetic Eigenprotein) K-W ANOVA p: 3e-15
M3 brown.UPenn Mixed (Synthetic Eigenprotein) K-W ANOVA p: 1.9e-08
bicor=-0.33, p=1.8e-08 cor=-0.3, p=4.8e-07
bicor=-0.29, p=3.2e-07 cor=-0.25, p=8.7e-06
Eigenprotein Value
Eigenprotein
Control
AsymAD
AD
ALS
FTLDU
PSP
CBD
PD.PDD
MSA
otherND
Braak Score
CERAD Score

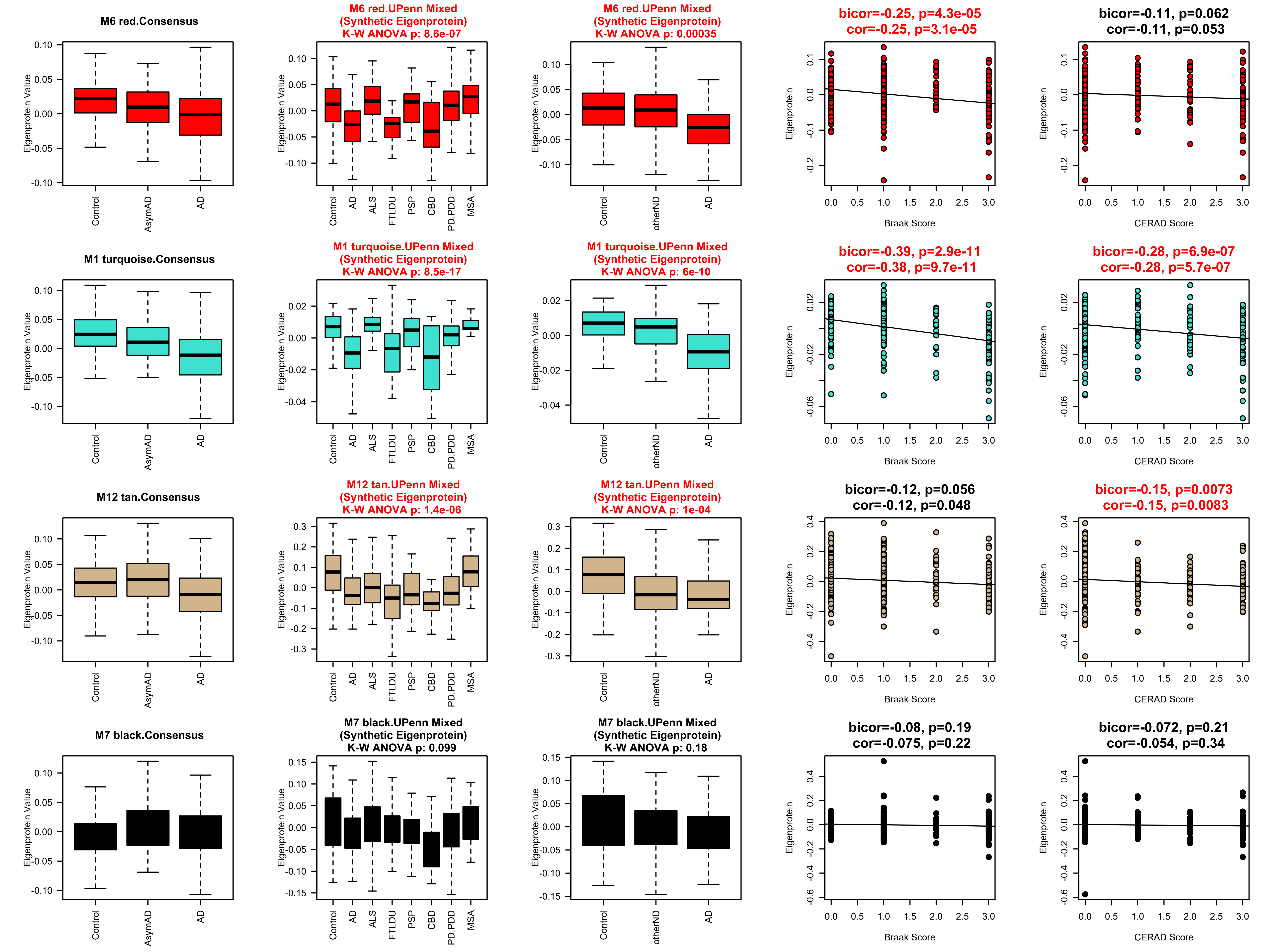
M6 red.Consensus
M6 red.UPenn Mixed
(Synthetic Eigenprotein)
K-W ANOVA p: 8.6e-07
M6 red.UPenn Mixed
(Synthetic Eigenprotein)
K-W ANOVA p: 0.00035
bicor=-0.25, p=4.3e-05
cor=-0.25, p=3.1e-05
bicor=-0.11, p=0.062
cor=-0.11, p=0.053
M1 turquoise.Consensus
M1 turquoise.UPenn Mixed
(Synthetic Eigenprotein)
K-W ANOVA p: 8.5e-17
M1 turquoise.UPenn Mixed
(Synthetic Eigenprotein)
K-W ANOVA p: 6e-10
bicor=-0.39, p=2.9e-11
cor=-0.38, p=9.7e-11
bicor=-0.28, p=6.9e-07
cor=-0.28, p=5.7e-07
M12 tan.Consensus
M12 tan.UPenn Mixed
(Synthetic Eigenprotein)
K-W ANOVA p: 1.4e-06
M12 tan.UPenn Mixed
(Synthetic Eigenprotein)
K-W ANOVA p: 1e-04
bicor=-0.12, p=0.056
cor=-0.12, p=0.048
bicor=-0.15, p=0.0073
cor=-0.15, p=0.0083
M7 black.Consensus
M7 black.UPenn Mixed
(Synthetic Eigenprotein)
K-W ANOVA p: 0.099
M7 black.UPenn Mixed
(Synthetic Eigenprotein)
K-W ANOVA p: 0.18
bicor=-0.08, p=0.19
cor=-0.075, p=0.22
bicor=-0.072, p=0.21
cor=-0.054, p=0.34
Eigenprotein Value
Eigenprotein
Control
AsymAD
AD
ALS
FTLDU
PSP
CBD
PD.PDD
MSA
otherND
Braak Score
CERAD Score

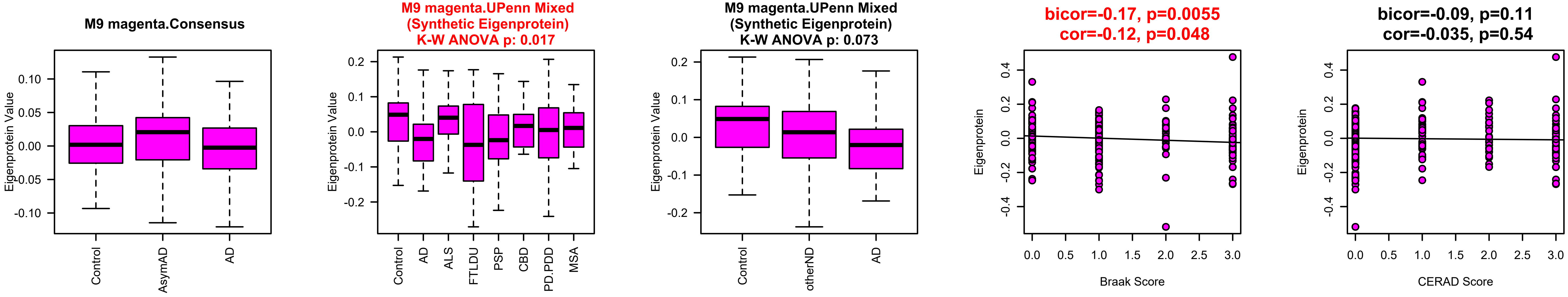
M9 magenta.Consensus
Eigenprotein Value
Control
AsymAD
AD
M9 magenta.UPenn Mixed
(Synthetic Eigenprotein)
K-W ANOVA p: 0.017
Control
AD
ALS
FTLDU
PSP
CBD
PD.PDD
MSA
M9 magenta.UPenn Mixed
(Synthetic Eigenprotein)
K-W ANOVA p: 0.073
Control
otherND
AD
bicor=-0.17, p=0.0055
cor=-0.12, p=0.048
Eigenprotein
Braak Score
bicor=-0.09, p=0.11
cor=-0.035, p=0.54
CERAD Score

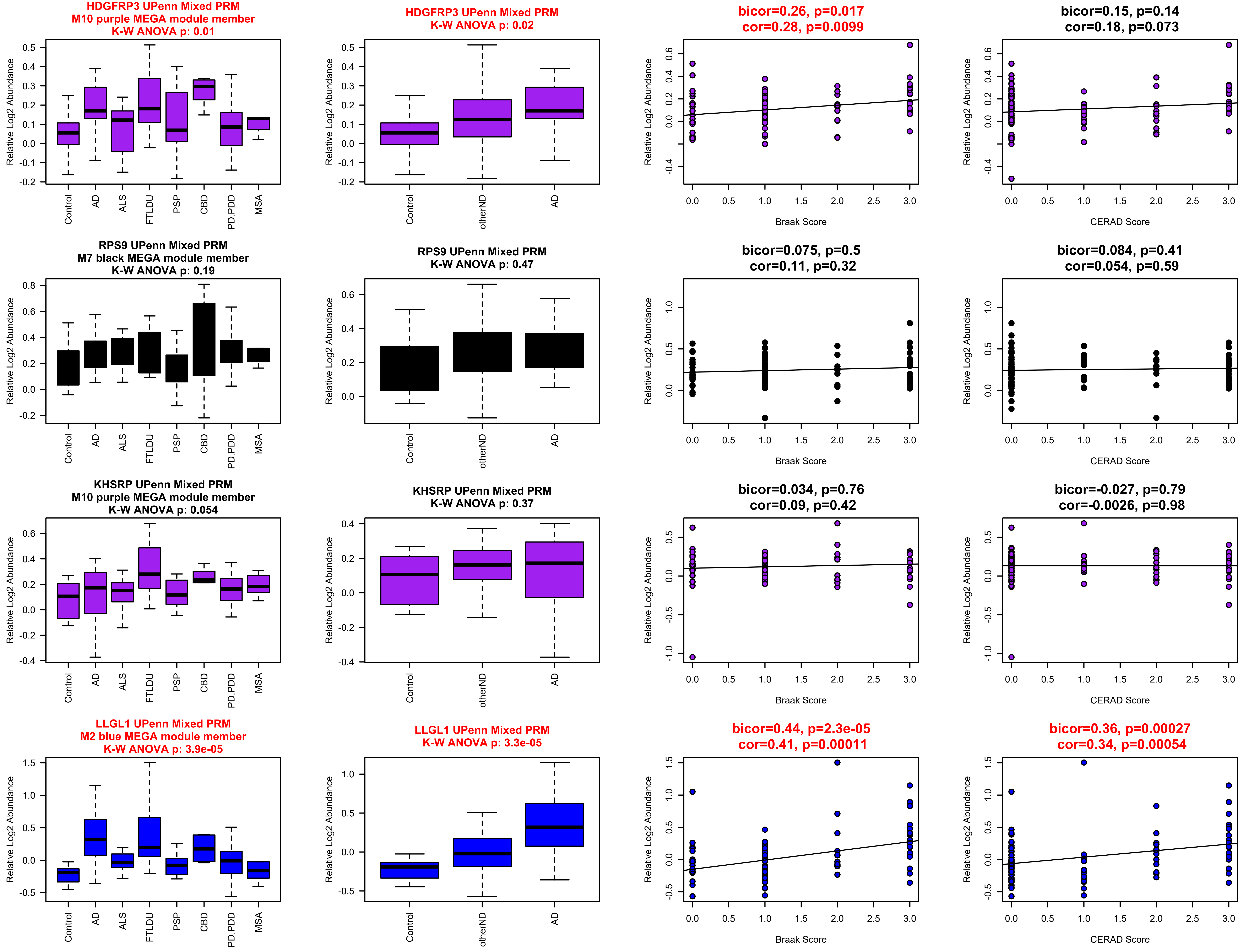
HDGFRP3 UPenn Mixed PRM
M10 purple MEGA module member
K-W ANOVA p: 0.01
HDGFRP3 UPenn Mixed PRM
K-W ANOVA p: 0.02
bicor=0.26, p=0.017
cor=0.28, p=0.0099
bicor=0.15, p=0.14
cor=0.18, p=0.073
RPS9 UPenn Mixed PRM
M7 black MEGA module member
K-W ANOVA p: 0.19
RPS9 UPenn Mixed PRM
K-W ANOVA p: 0.47
bicor=0.075, p=0.5
cor=0.11, p=0.32
bicor=0.084, p=0.41
cor=0.054, p=0.59
KHSRP UPenn Mixed PRM
M10 purple MEGA module member
K-W ANOVA p: 0.054
KHSRP UPenn Mixed PRM
K-W ANOVA p: 0.37
bicor=0.034, p=0.76
cor=0.09, p=0.42
bicor=-0.027, p=0.79
cor=-0.0026, p=0.98
LLGL1 UPenn Mixed PRM
M2 blue MEGA module member
K-W ANOVA p: 3.9e-05
LLGL1 UPenn Mixed PRM
K-W ANOVA p: 3.3e-05
bicor=0.44, p=2.3e-05
cor=0.41, p=0.00011
bicor=0.36, p=0.00027
cor=0.34, p=0.00054
Relative Log2 Abundance
Control
AD
ALS
FTLDU
PSP
CBD
PD.PDD
MSA
otherND
Braak Score
CERAD Score

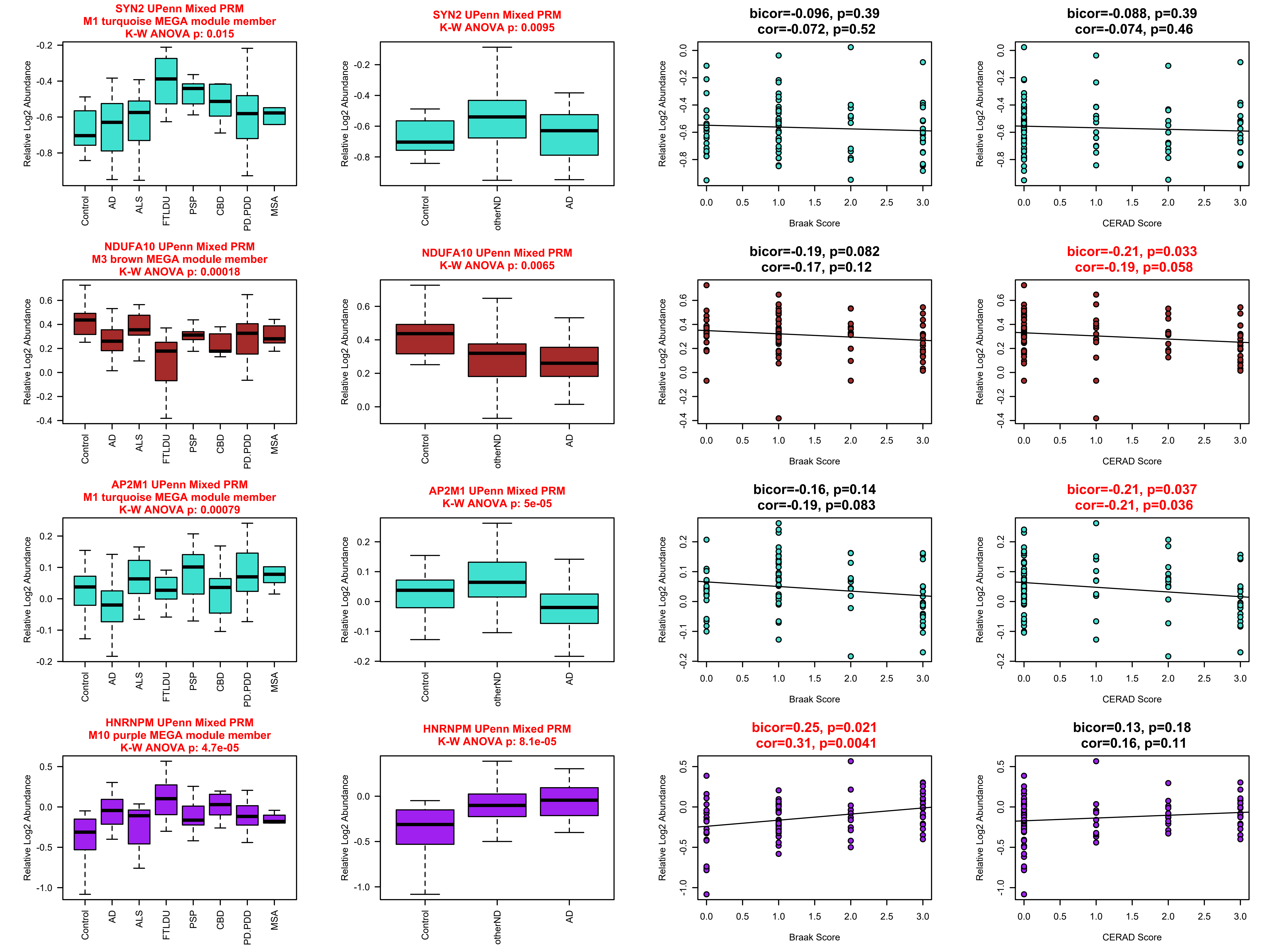

SYN2 UPenn Mixed PRM
M1 turquoise MEGA module member
K-W ANOVA p: 0.015
SYN2 UPenn Mixed PRM
K-W ANOVA p: 0.0095
bicor=-0.096, p=0.39
cor=-0.072, p=0.52
bicor=-0.088, p=0.39
cor=-0.074, p=0.46
NDUFA10 UPenn Mixed PRM
M3 brown MEGA module member
K-W ANOVA p: 0.00018
NDUFA10 UPenn Mixed PRM
K-W ANOVA p: 0.0065
bicor=-0.19, p=0.082
cor=-0.17, p=0.12
bicor=-0.21, p=0.033
cor=-0.19, p=0.058
AP2M1 UPenn Mixed PRM
M1 turquoise MEGA module member
K-W ANOVA p: 0.00079
AP2M1 UPenn Mixed PRM
K-W ANOVA p: 5e-05
bicor=-0.16, p=0.14
cor=-0.19, p=0.083
bicor=-0.21, p=0.037
cor=-0.21, p=0.036
HNRNPM UPenn Mixed PRM
M10 purple MEGA module member
K-W ANOVA p: 4.7e-05
HNRNPM UPenn Mixed PRM
K-W ANOVA p: 8.1e-05
bicor=0.25, p=0.021
cor=0.31, p=0.0041
bicor=0.13, p=0.18
cor=0.16, p=0.11
Relative Log2 Abundance
Control
AD
ALS
FTLDU
PSP
CBD
PD.PDD
MSA
otherND
Braak Score
CERAD Score

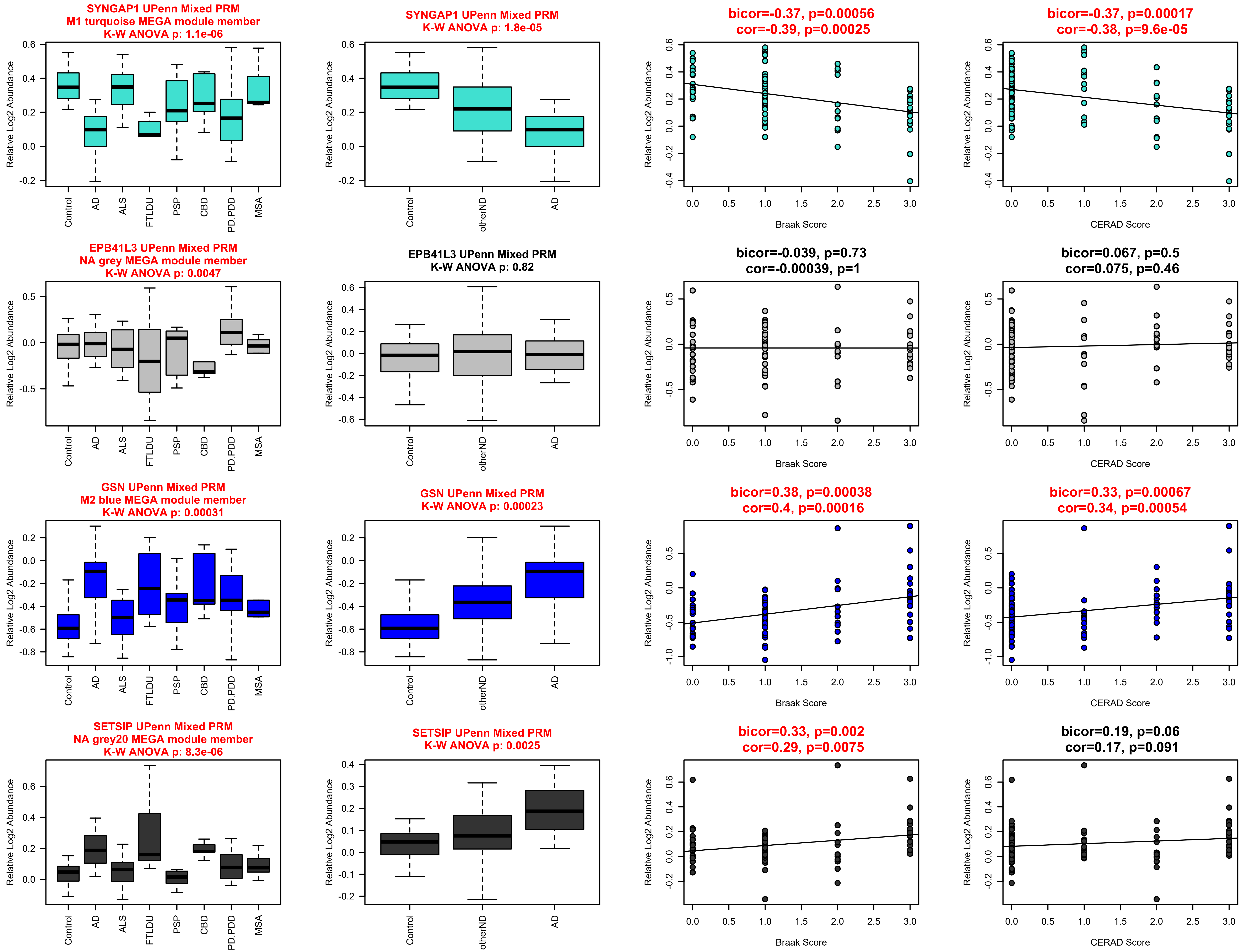
SYNGAP1 UPenn Mixed PRM
M1 turquoise MEGA module member
K-W ANOVA p: 1.1e-06
SYNGAP1 UPenn Mixed PRM
K-W ANOVA p: 1.8e-05
bicor=-0.37, p=0.00056
cor=-0.39, p=0.00025
bicor=-0.37, p=0.00017
cor=-0.38, p=9.6e-05
EPB41L3 UPenn Mixed PRM
NA grey MEGA module member
K-W ANOVA p: 0.0047
EPB41L3 UPenn Mixed PRM
K-W ANOVA p: 0.82
bicor=-0.039, p=0.73
cor=-0.00039, p=1
bicor=0.067, p=0.5
cor=0.075, p=0.46
GSN UPenn Mixed PRM
M2 blue MEGA module member
K-W ANOVA p: 0.00031
GSN UPenn Mixed PRM
K-W ANOVA p: 0.00023
bicor=0.38, p=0.00038
cor=0.4, p=0.00016
bicor=0.33, p=0.00067
cor=0.34, p=0.00054
SETSIP UPenn Mixed PRM
NA grey20 MEGA module member
K-W ANOVA p: 8.3e-06
SETSIP UPenn Mixed PRM
K-W ANOVA p: 0.0025
bicor=0.33, p=0.002
cor=0.29, p=0.0075
bicor=0.19, p=0.06
cor=0.17, p=0.091
Relative Log2 Abundance
Control
AD
ALS
FTLDU
PSP
CBD
PD.PDD
MSA
otherND
Braak Score
CERAD Score

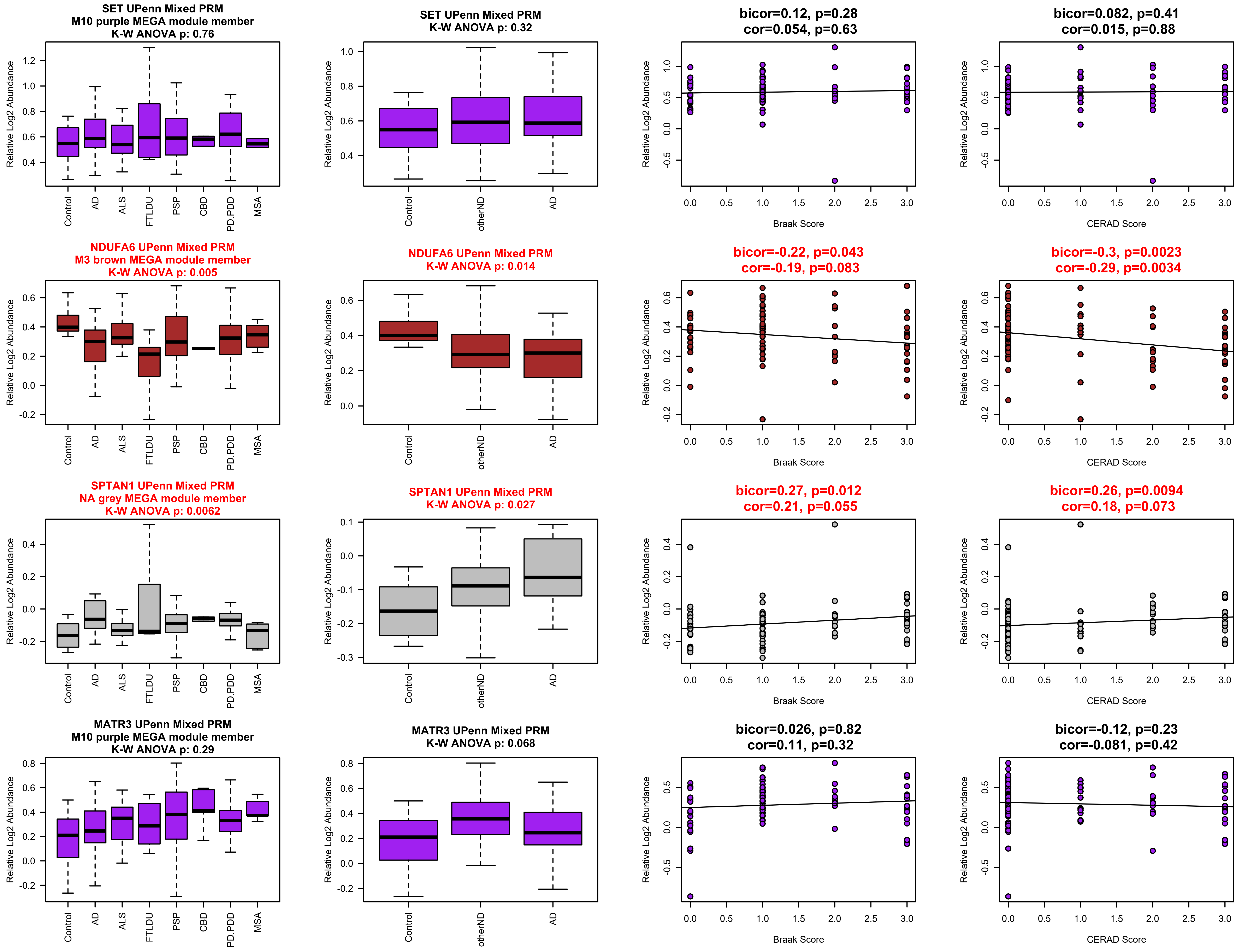
SET UPenn Mixed PRM
M10 purple MEGA module member
K-W ANOVA p: 0.76
SET UPenn Mixed PRM
K-W ANOVA p: 0.32
bicor=0.12, p=0.28
cor=0.054, p=0.63
bicor=0.082, p=0.41
cor=0.015, p=0.88
NDUFA6 UPenn Mixed PRM
M3 brown MEGA module member
K-W ANOVA p: 0.005
NDUFA6 UPenn Mixed PRM
K-W ANOVA p: 0.014
bicor=-0.22, p=0.043
cor=-0.19, p=0.083
bicor=-0.3, p=0.0023
cor=-0.29, p=0.0034
SPTAN1 UPenn Mixed PRM
NA grey MEGA module member
K-W ANOVA p: 0.0062
SPTAN1 UPenn Mixed PRM
K-W ANOVA p: 0.027
bicor=0.27, p=0.012
cor=0.21, p=0.055
bicor=0.26, p=0.0094
cor=0.18, p=0.073
MATR3 UPenn Mixed PRM
M10 purple MEGA module member
K-W ANOVA p: 0.29
MATR3 UPenn Mixed PRM
K-W ANOVA p: 0.068
bicor=0.026, p=0.82
cor=0.11, p=0.32
bicor=-0.12, p=0.23
cor=-0.081, p=0.42
Relative Log2 Abundance
Control
AD
ALS
FTLDU
PSP
CBD
PD.PDD
MSA
otherND
Braak Score
CERAD Score

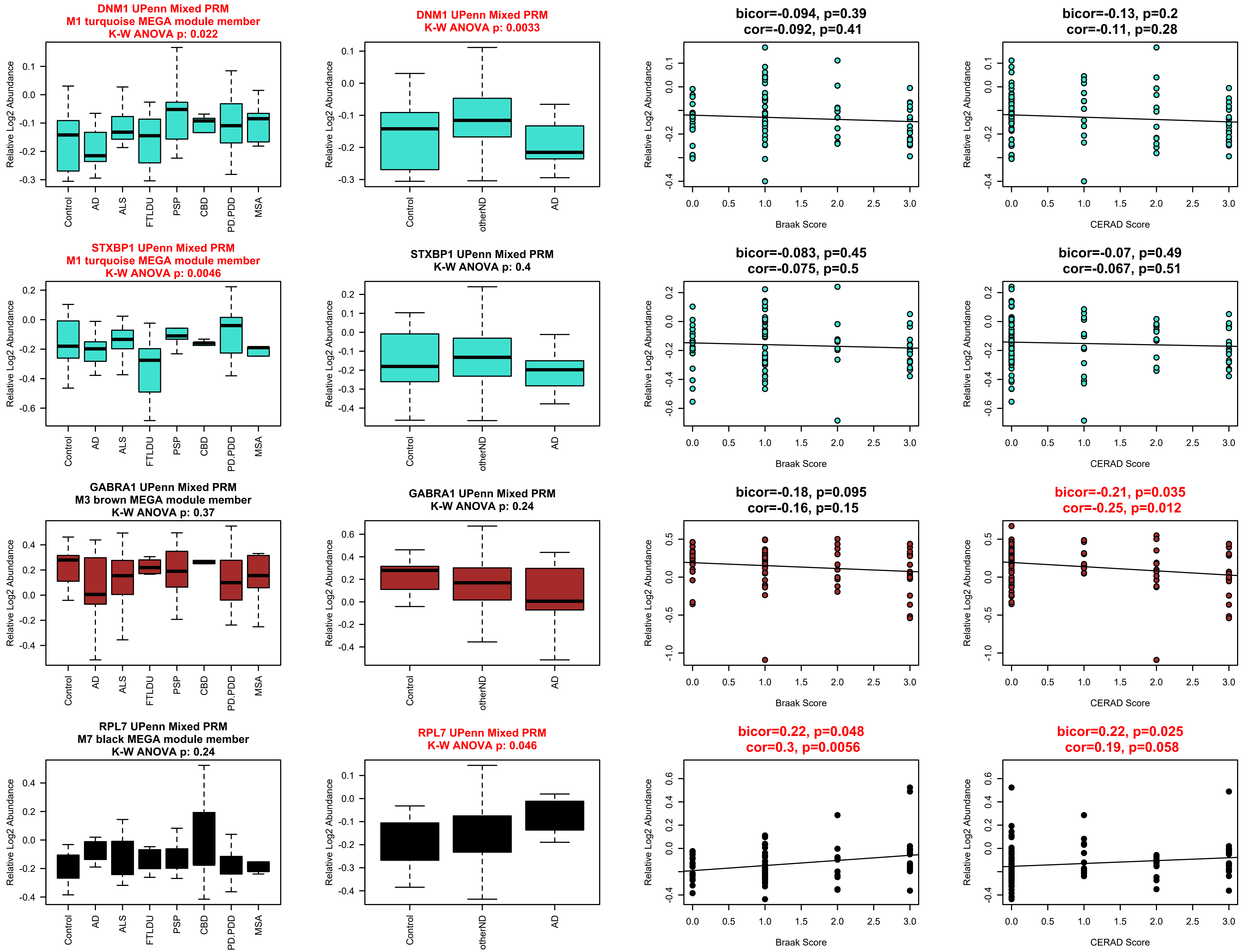
DNM1 UPenn Mixed PRM
M1 turquoise MEGA module member
K-W ANOVA p: 0.022
DNM1 UPenn Mixed PRM
K-W ANOVA p: 0.0033
bicor=-0.094, p=0.39
cor=-0.092, p=0.41
bicor=-0.13, p=0.2
cor=-0.11, p=0.28
STXBP1 UPenn Mixed PRM
M1 turquoise MEGA module member
K-W ANOVA p: 0.0046
STXBP1 UPenn Mixed PRM
K-W ANOVA p: 0.4
bicor=-0.083, p=0.45
cor=-0.075, p=0.5
bicor=-0.07, p=0.49
cor=-0.067, p=0.51
GABRA1 UPenn Mixed PRM
M3 brown MEGA module member
K-W ANOVA p: 0.37
GABRA1 UPenn Mixed PRM
K-W ANOVA p: 0.24
bicor=-0.18, p=0.095
cor=-0.16, p=0.15
bicor=-0.21, p=0.035
cor=-0.25, p=0.012
RPL7 UPenn Mixed PRM
M7 black MEGA module member
K-W ANOVA p: 0.24
RPL7 UPenn Mixed PRM
K-W ANOVA p: 0.046
bicor=0.22, p=0.048
cor=0.3, p=0.0056
bicor=0.22, p=0.025
cor=0.19, p=0.058
Relative Log2 Abundance
Control
AD
ALS
FTLDU
PSP
CBD
PD.PDD
MSA
otherND
Braak Score
CERAD Score

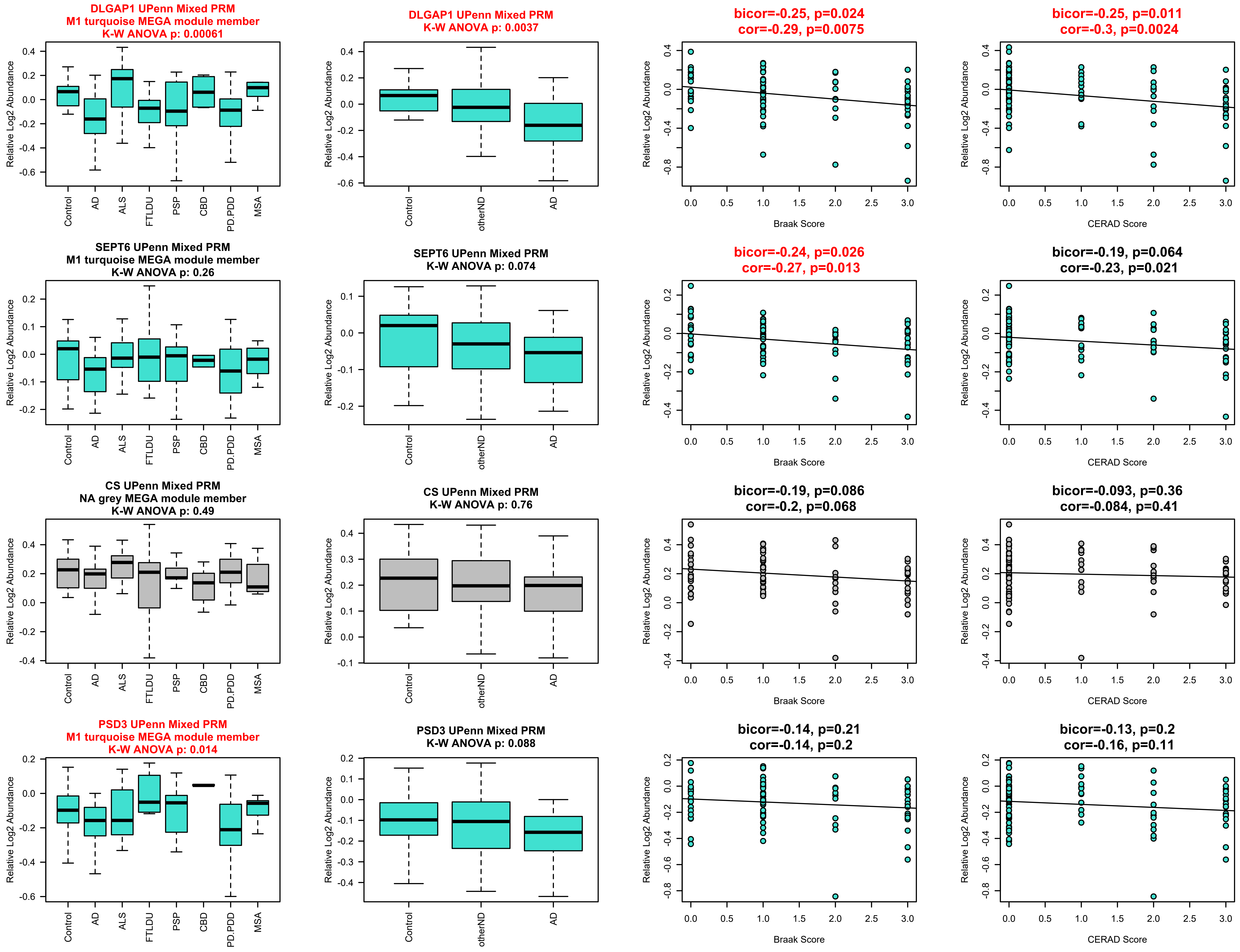
DLGAP1 UPenn Mixed PRM
M1 turquoise MEGA module member
K-W ANOVA p: 0.00061
DLGAP1 UPenn Mixed PRM
K-W ANOVA p: 0.0037
bicor=-0.25, p=0.024
cor=-0.29, p=0.0075
bicor=-0.25, p=0.011
cor=-0.3, p=0.0024
SEPT6 UPenn Mixed PRM
M1 turquoise MEGA module member
K-W ANOVA p: 0.26
SEPT6 UPenn Mixed PRM
K-W ANOVA p: 0.074
bicor=-0.24, p=0.026
cor=-0.27, p=0.013
bicor=-0.19, p=0.064
cor=-0.23, p=0.021
CS UPenn Mixed PRM
NA grey MEGA module member
K-W ANOVA p: 0.49
CS UPenn Mixed PRM
K-W ANOVA p: 0.76
bicor=-0.19, p=0.086
cor=-0.2, p=0.068
bicor=-0.093, p=0.36
cor=-0.084, p=0.41
PSD3 UPenn Mixed PRM
M1 turquoise MEGA module member
K-W ANOVA p: 0.014
PSD3 UPenn Mixed PRM
K-W ANOVA p: 0.088
bicor=-0.14, p=0.21
cor=-0.14, p=0.2
bicor=-0.13, p=0.2
cor=-0.16, p=0.11
Relative Log2 Abundance
Control
AD
ALS
FTLDU
PSP
CBD
PD.PDD
MSA
otherND
Braak Score
CERAD Score

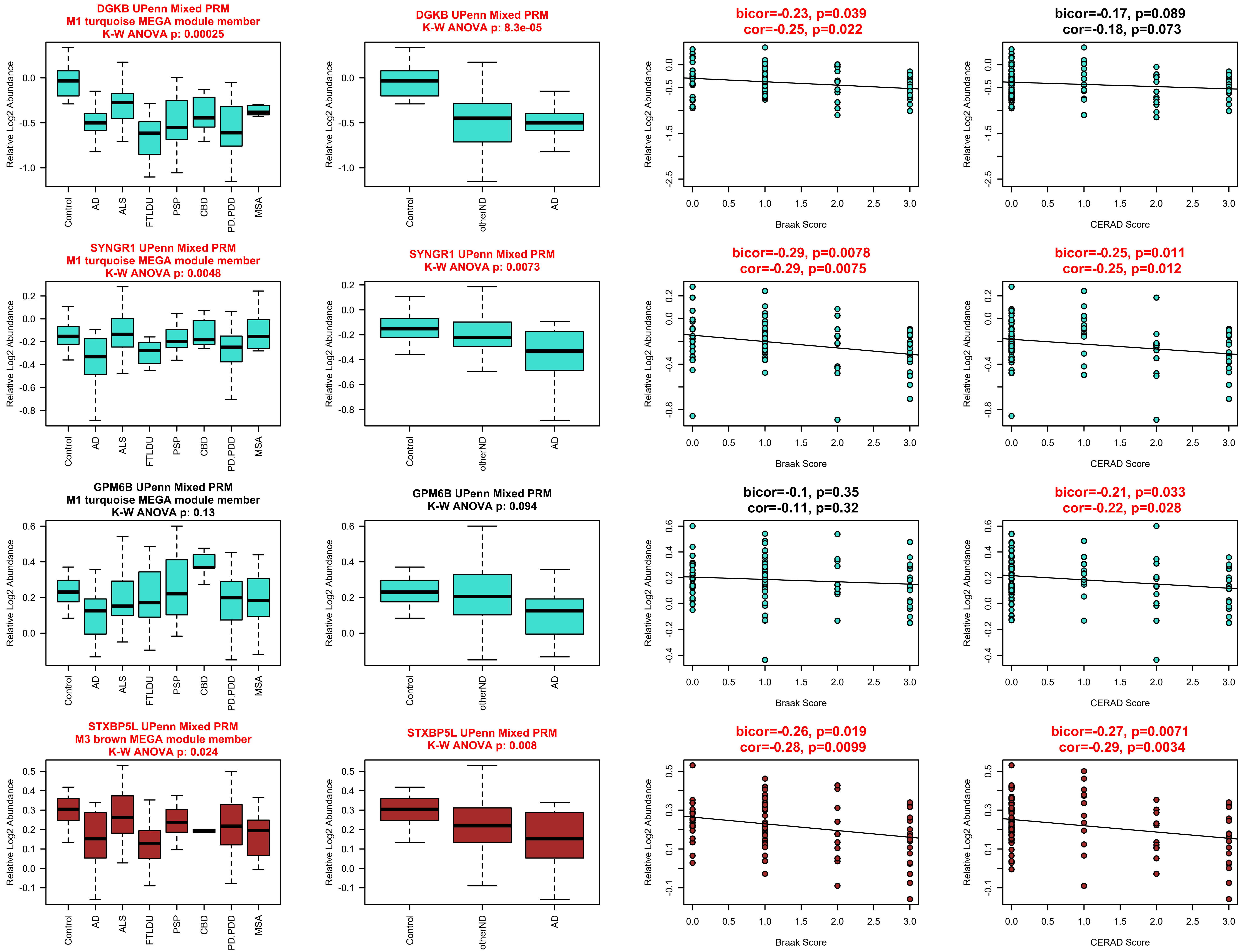
DGKB UPenn Mixed PRM
M1 turquoise MEGA module member
K-W ANOVA p: 0.00025
DGKB UPenn Mixed PRM
K-W ANOVA p: 8.3e-05
bicor=-0.23, p=0.039
cor=-0.25, p=0.022
bicor=-0.17, p=0.089
cor=-0.18, p=0.073
SYNGR1 UPenn Mixed PRM
M1 turquoise MEGA module member
K-W ANOVA p: 0.0048
SYNGR1 UPenn Mixed PRM
K-W ANOVA p: 0.0073
bicor=-0.29, p=0.0078
cor=-0.29, p=0.0075
bicor=-0.25, p=0.011
cor=-0.25, p=0.012
GPM6B UPenn Mixed PRM
M1 turquoise MEGA module member
K-W ANOVA p: 0.13
GPM6B UPenn Mixed PRM
K-W ANOVA p: 0.094
bicor=-0.1, p=0.35
cor=-0.11, p=0.32
bicor=-0.21, p=0.033
cor=-0.22, p=0.028
STXBP5L UPenn Mixed PRM
M3 brown MEGA module member
K-W ANOVA p: 0.024
STXBP5L UPenn Mixed PRM
K-W ANOVA p: 0.008
bicor=-0.26, p=0.019
cor=-0.28, p=0.0099
bicor=-0.27, p=0.0071
cor=-0.29, p=0.0034
Relative Log2 Abundance
Control
AD
ALS
FTLDU
PSP
CBD
PD.PDD
MSA
otherND
Braak Score
CERAD Score

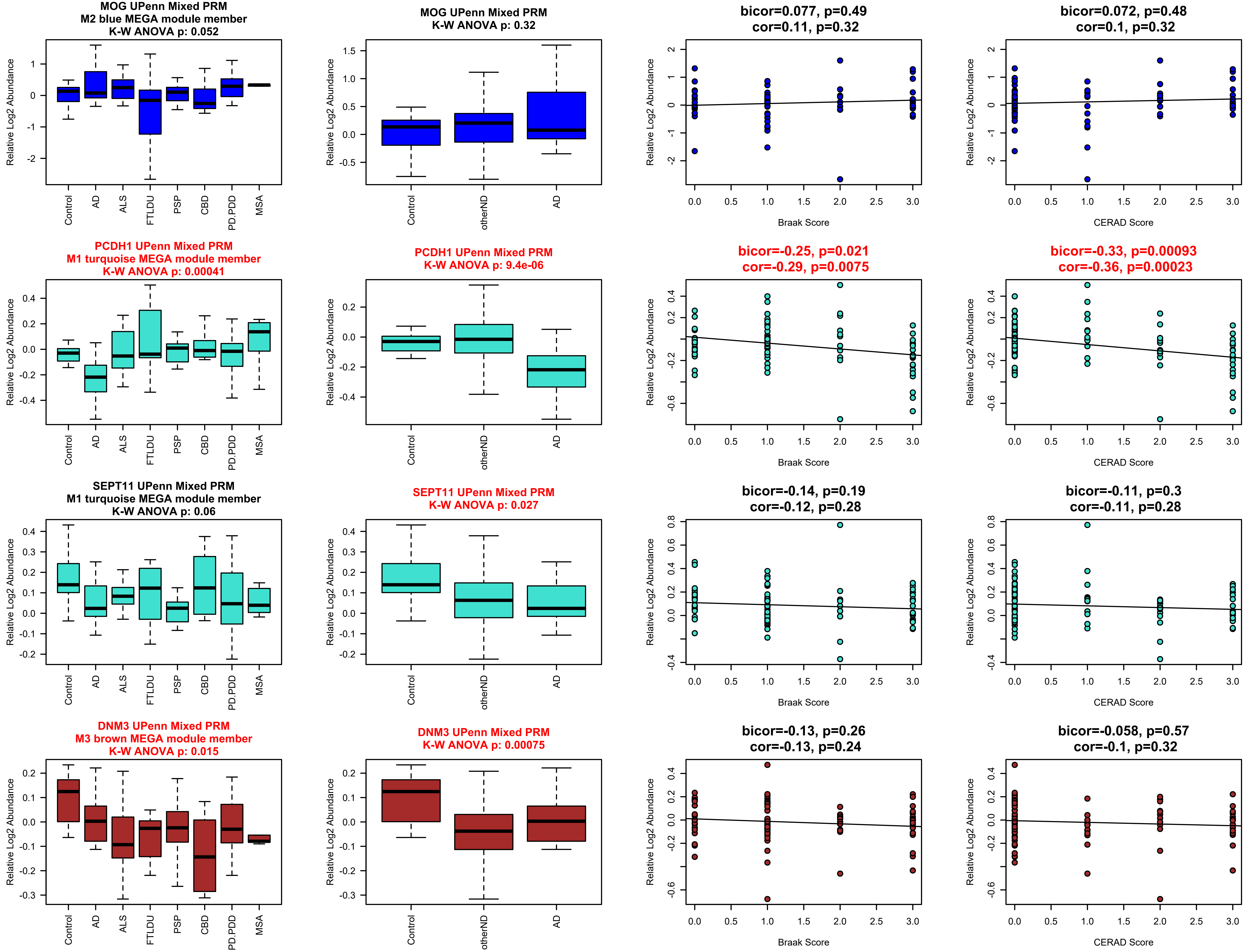

MOG UPenn Mixed PRM
M2 blue MEGA module member
K-W ANOVA p: 0.052
MOG UPenn Mixed PRM
K-W ANOVA p: 0.32
bicor=0.077, p=0.49
cor=0.11, p=0.32
bicor=0.072, p=0.48
cor=0.1, p=0.32
PCDH1 UPenn Mixed PRM
M1 turquoise MEGA module member
K-W ANOVA p: 0.00041
PCDH1 UPenn Mixed PRM
K-W ANOVA p: 9.4e-06
bicor=-0.25, p=0.021
cor=-0.29, p=0.0075
bicor=-0.33, p=0.00093
cor=-0.36, p=0.00023
SEPT11 UPenn Mixed PRM
M1 turquoise MEGA module member
K-W ANOVA p: 0.06
SEPT11 UPenn Mixed PRM
K-W ANOVA p: 0.027
bicor=-0.14, p=0.19
cor=-0.12, p=0.28
bicor=-0.11, p=0.3
cor=-0.11, p=0.28
DNM3 UPenn Mixed PRM
M3 brown MEGA module member
K-W ANOVA p: 0.015
DNM3 UPenn Mixed PRM
K-W ANOVA p: 0.00075
bicor=-0.13, p=0.26
cor=-0.13, p=0.24
bicor=-0.058, p=0.57
cor=-0.1, p=0.32
Relative Log2 Abundance
Control
AD
ALS
FTLDU
PSP
CBD
PD.PDD
MSA
otherND
Braak Score
CERAD Score

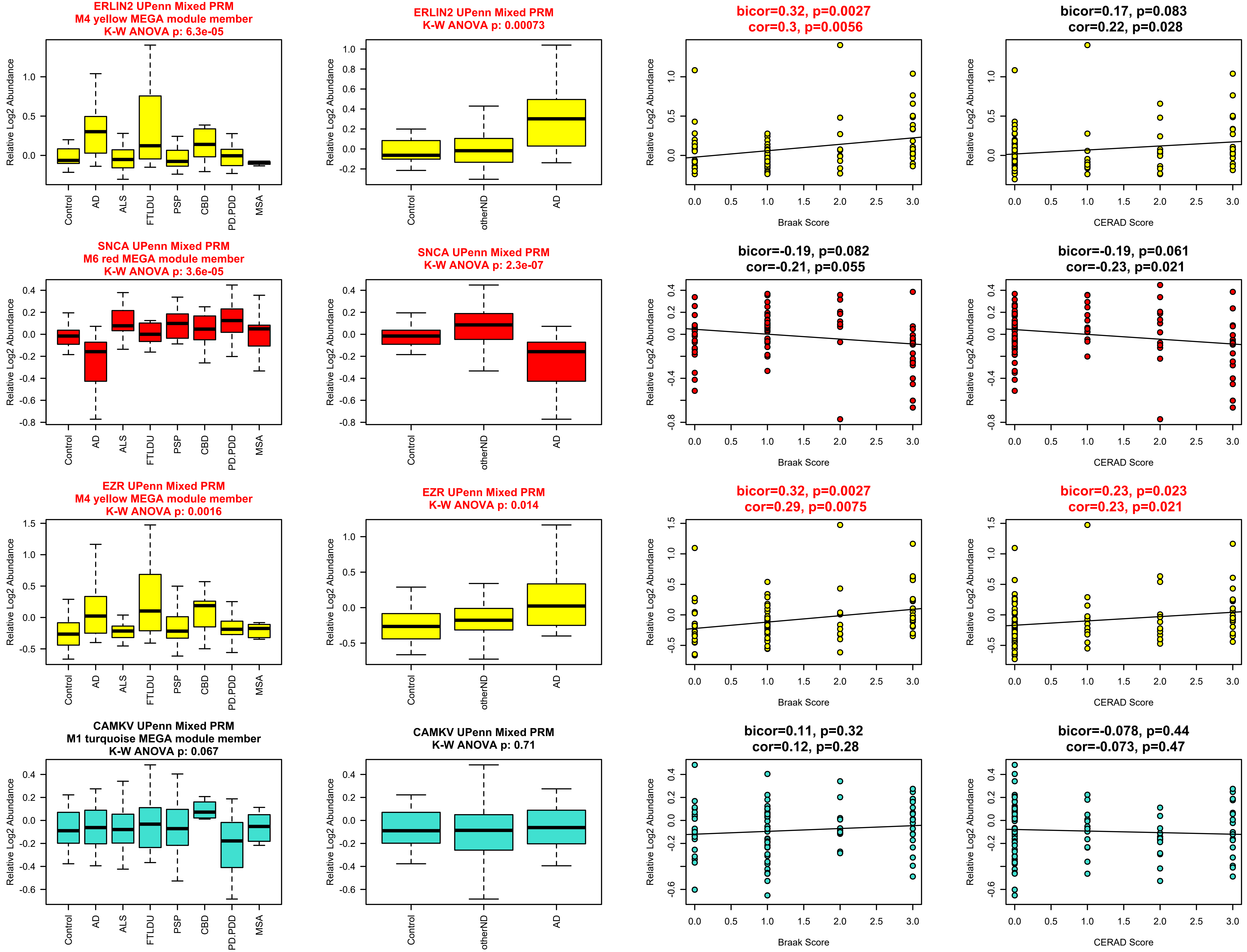
ERLIN2 UPenn Mixed PRM
M4 yellow MEGA module member
K-W ANOVA p: 6.3e-05
ERLIN2 UPenn Mixed PRM
K-W ANOVA p: 0.00073
bicor=0.32, p=0.0027
cor=0.3, p=0.0056
bicor=0.17, p=0.083
cor=0.22, p=0.028
SNCA UPenn Mixed PRM
M6 red MEGA module member
K-W ANOVA p: 3.6e-05
SNCA UPenn Mixed PRM
K-W ANOVA p: 2.3e-07
bicor=-0.19, p=0.082
cor=-0.21, p=0.055
bicor=-0.19, p=0.061
cor=-0.23, p=0.021
EZR UPenn Mixed PRM
M4 yellow MEGA module member
K-W ANOVA p: 0.0016
EZR UPenn Mixed PRM
K-W ANOVA p: 0.014
bicor=0.32, p=0.0027
cor=0.29, p=0.0075
bicor=0.23, p=0.023
cor=0.23, p=0.021
CAMKV UPenn Mixed PRM
M1 turquoise MEGA module member
K-W ANOVA p: 0.067
CAMKV UPenn Mixed PRM
K-W ANOVA p: 0.71
bicor=0.11, p=0.32
cor=0.12, p=0.28
bicor=-0.078, p=0.44
cor=-0.073, p=0.47
Relative Log2 Abundance
Control
AD
ALS
FTLDU
PSP
CBD
PD.PDD
MSA
otherND
Braak Score
CERAD Score

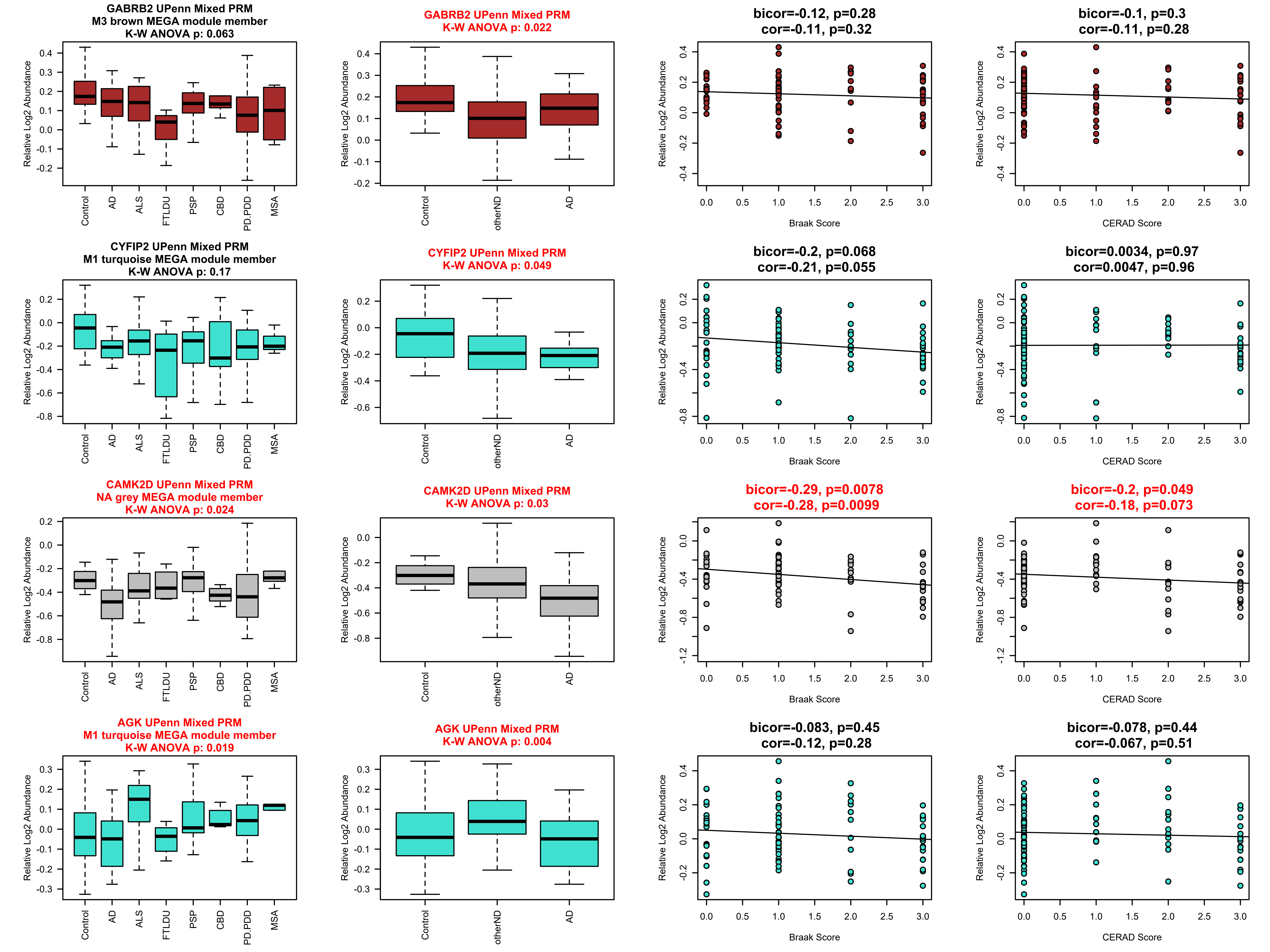
GABRB2 UPenn Mixed PRM
M3 brown MEGA module member
K-W ANOVA p: 0.063
GABRB2 UPenn Mixed PRM
K-W ANOVA p: 0.022
bicor=-0.12, p=0.28
cor=-0.11, p=0.32
bicor=-0.1, p=0.3
cor=-0.11, p=0.28
CYFIP2 UPenn Mixed PRM
M1 turquoise MEGA module member
K-W ANOVA p: 0.17
CYFIP2 UPenn Mixed PRM
K-W ANOVA p: 0.049
bicor=-0.2, p=0.068
cor=-0.21, p=0.055
bicor=0.0034, p=0.97
cor=0.0047, p=0.96
CAMK2D UPenn Mixed PRM
NA grey MEGA module member
K-W ANOVA p: 0.024
CAMK2D UPenn Mixed PRM
K-W ANOVA p: 0.03
bicor=-0.29, p=0.0078
cor=-0.28, p=0.0099
bicor=-0.2, p=0.049
cor=-0.18, p=0.073
AGK UPenn Mixed PRM
M1 turquoise MEGA module member
K-W ANOVA p: 0.019
AGK UPenn Mixed PRM
K-W ANOVA p: 0.004
bicor=-0.083, p=0.45
cor=-0.12, p=0.28
bicor=-0.078, p=0.44
cor=-0.067, p=0.51
Relative Log2 Abundance
Control
AD
ALS
FTLDU
PSP
CBD
PD.PDD
MSA
otherND
Braak Score
CERAD Score

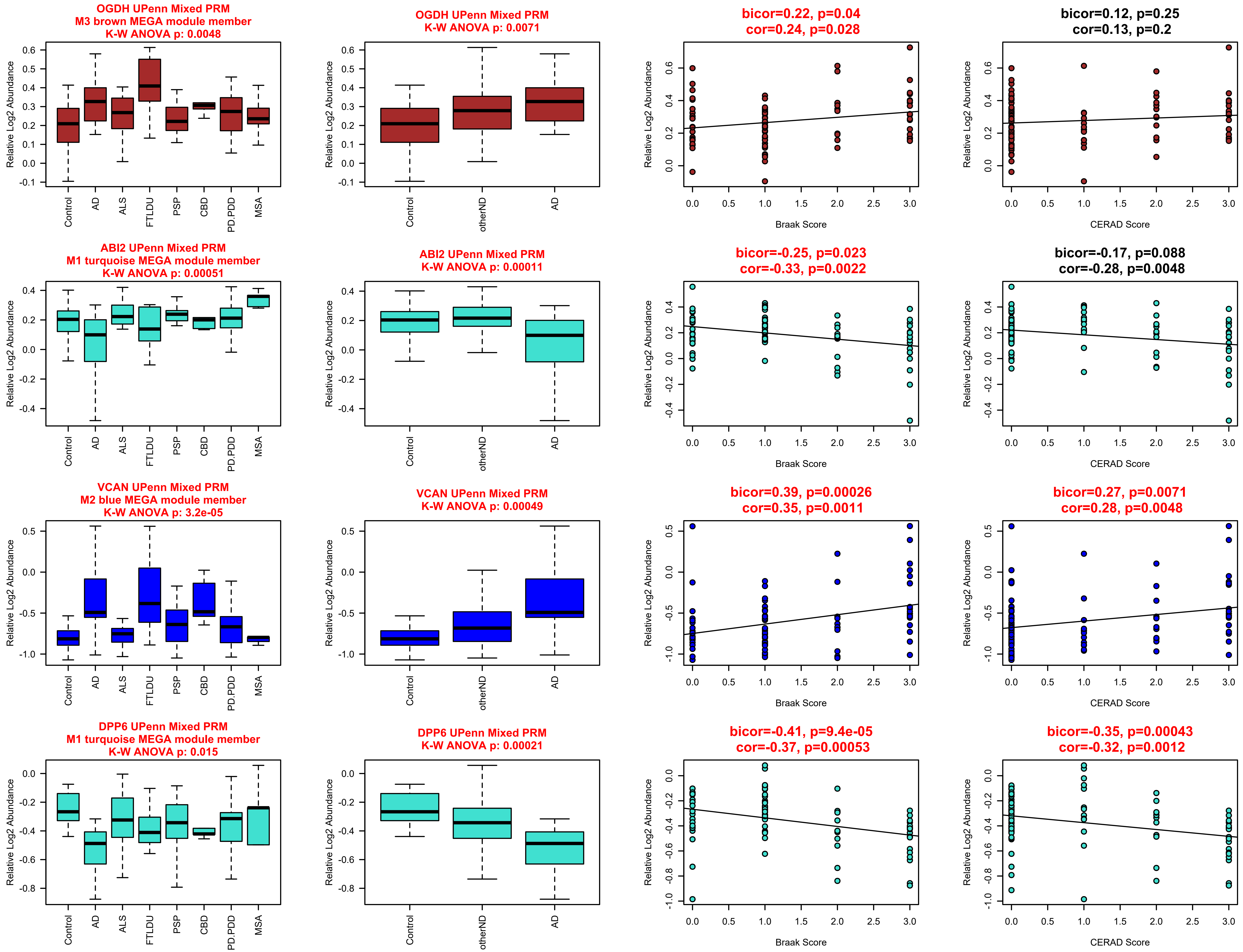
OGDH UPenn Mixed PRM
M3 brown MEGA module member
K-W ANOVA p: 0.0048
OGDH UPenn Mixed PRM
K-W ANOVA p: 0.0071
bicor=0.22, p=0.04
cor=0.24, p=0.028
bicor=0.12, p=0.25
cor=0.13, p=0.2
ABI2 UPenn Mixed PRM
M1 turquoise MEGA module member
K-W ANOVA p: 0.00051
ABI2 UPenn Mixed PRM
K-W ANOVA p: 0.00011
bicor=-0.25, p=0.023
cor=-0.33, p=0.0022
bicor=-0.17, p=0.088
cor=-0.28, p=0.0048
VCAN UPenn Mixed PRM
M2 blue MEGA module member
K-W ANOVA p: 3.2e-05
VCAN UPenn Mixed PRM
K-W ANOVA p: 0.00049
bicor=0.39, p=0.00026
cor=0.35, p=0.0011
bicor=0.27, p=0.0071
cor=0.28, p=0.0048
DPP6 UPenn Mixed PRM
M1 turquoise MEGA module member
K-W ANOVA p: 0.015
DPP6 UPenn Mixed PRM
K-W ANOVA p: 0.00021
bicor=-0.41, p=9.4e-05
cor=-0.37, p=0.00053
bicor=-0.35, p=0.00043
cor=-0.32, p=0.0012
Relative Log2 Abundance
Control
AD
ALS
FTLDU
PSP
CBD
PD.PDD
MSA
otherND
Braak Score
CERAD Score

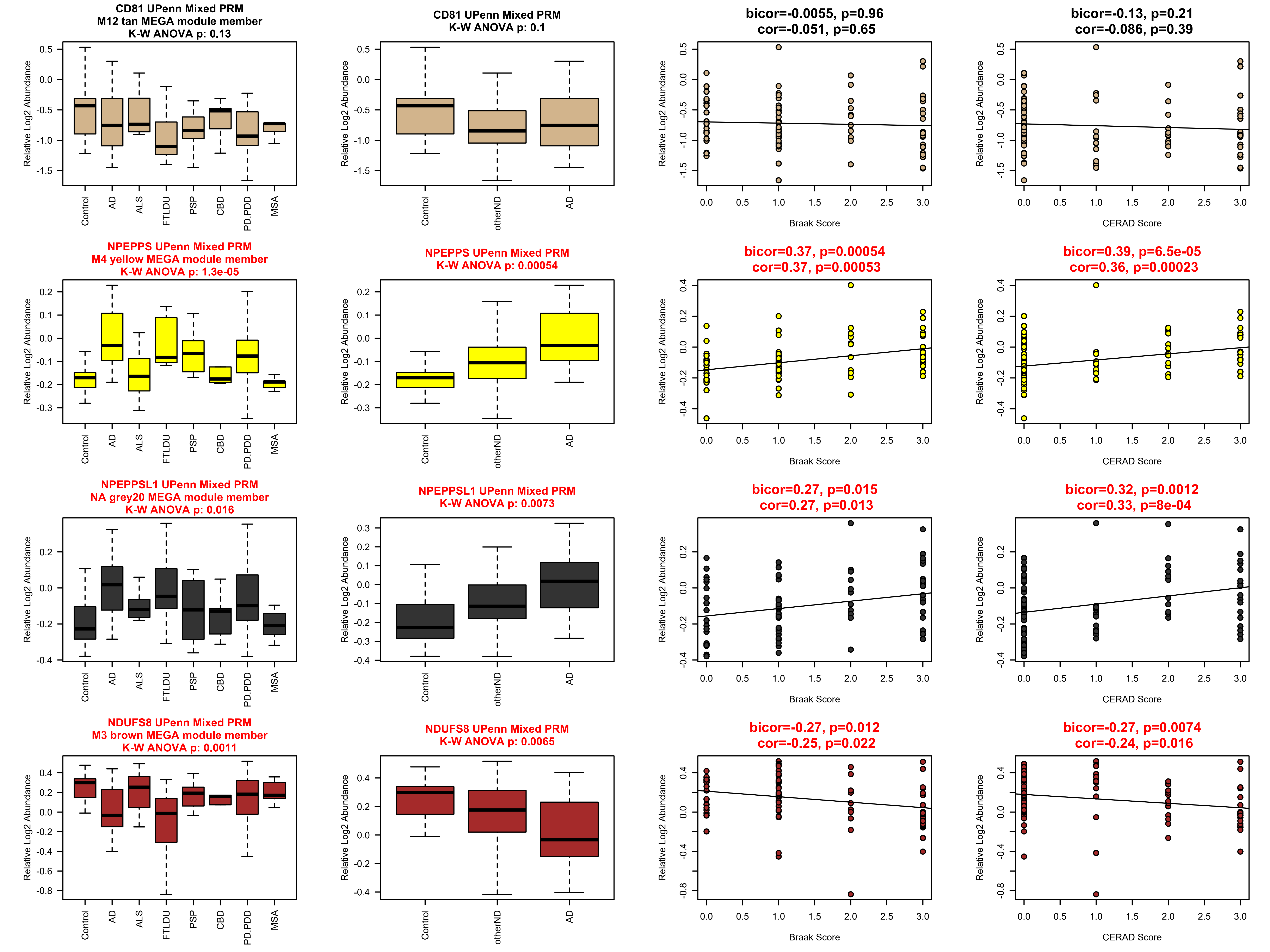

CD81 UPenn Mixed PRM
M12 tan MEGA module member
K-W ANOVA p: 0.13
CD81 UPenn Mixed PRM
K-W ANOVA p: 0.1
bicor=-0.0055, p=0.96
cor=-0.051, p=0.65
bicor=-0.13, p=0.21
cor=-0.086, p=0.39
NPEPPS UPenn Mixed PRM
M4 yellow MEGA module member
K-W ANOVA p: 1.3e-05
NPEPPS UPenn Mixed PRM
K-W ANOVA p: 0.00054
bicor=0.37, p=0.00054
cor=0.37, p=0.00053
bicor=0.39, p=6.5e-05
cor=0.36, p=0.00023
NPEPPSL1 UPenn Mixed PRM
NA grey20 MEGA module member
K-W ANOVA p: 0.016
NPEPPSL1 UPenn Mixed PRM
K-W ANOVA p: 0.0073
bicor=0.27, p=0.015
cor=0.27, p=0.013
bicor=0.32, p=0.0012
cor=0.33, p=8e-04
NDUFS8 UPenn Mixed PRM
M3 brown MEGA module member
K-W ANOVA p: 0.0011
NDUFS8 UPenn Mixed PRM
K-W ANOVA p: 0.0065
bicor=-0.27, p=0.012
cor=-0.25, p=0.022
bicor=-0.27, p=0.0074
cor=-0.24, p=0.016
Relative Log2 Abundance
Control
AD
ALS
FTLDU
PSP
CBD
PD.PDD
MSA
otherND
Braak Score
CERAD Score

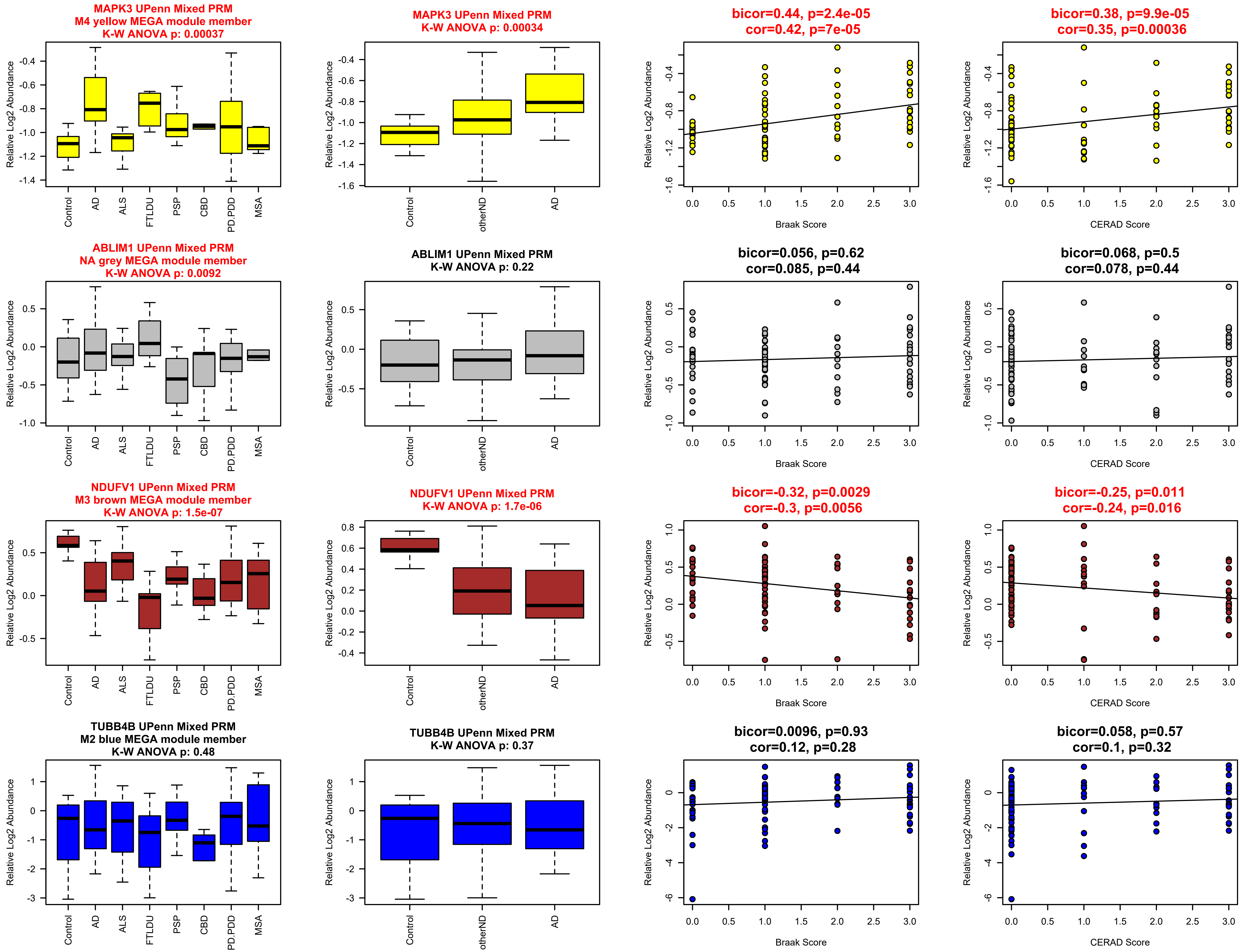

MAPK3 UPenn Mixed PRM
M4 yellow MEGA module member
K-W ANOVA p: 0.00037
MAPK3 UPenn Mixed PRM
K-W ANOVA p: 0.00034
bicor=0.44, p=2.4e-05
cor=0.42, p=7e-05
bicor=0.38, p=9.9e-05
cor=0.35, p=0.00036
ABLIM1 UPenn Mixed PRM
NA grey MEGA module member
K-W ANOVA p: 0.0092
ABLIM1 UPenn Mixed PRM
K-W ANOVA p: 0.22
bicor=0.056, p=0.62
cor=0.085, p=0.44
bicor=0.068, p=0.5
cor=0.078, p=0.44
NDUFV1 UPenn Mixed PRM
M3 brown MEGA module member
K-W ANOVA p: 1.5e-07
NDUFV1 UPenn Mixed PRM
K-W ANOVA p: 1.7e-06
bicor=-0.32, p=0.0029
cor=-0.3, p=0.0056
bicor=-0.25, p=0.011
cor=-0.24, p=0.016
TUBB4B UPenn Mixed PRM
M2 blue MEGA module member
K-W ANOVA p: 0.48
TUBB4B UPenn Mixed PRM
K-W ANOVA p: 0.37
bicor=0.0096, p=0.93
cor=0.12, p=0.28
bicor=0.058, p=0.57
cor=0.1, p=0.32
Relative Log2 Abundance
Control
AD
ALS
FTLDU
PSP
CBD
PD.PDD
MSA
otherND
Braak Score
CERAD Score

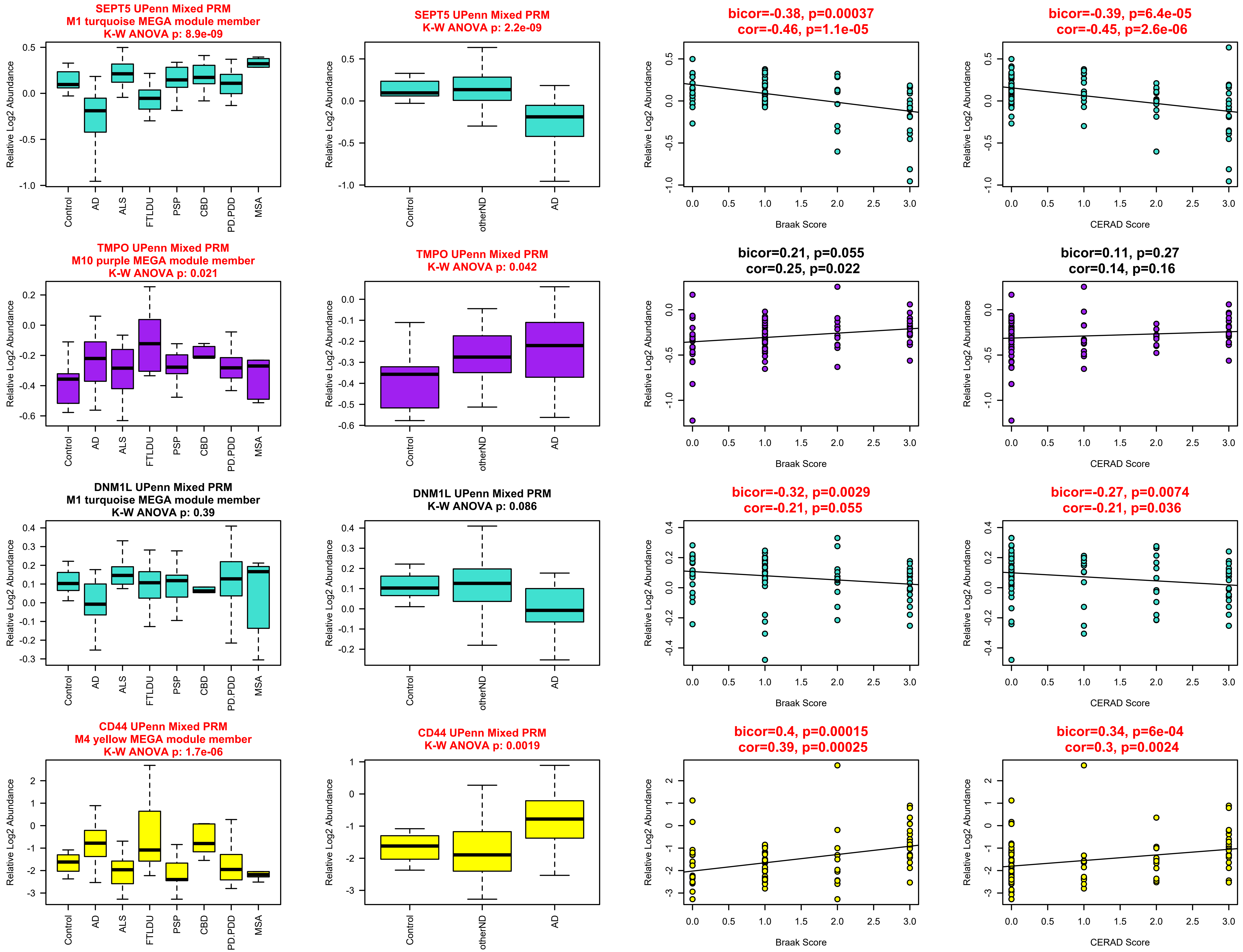
SEPT5 UPenn Mixed PRM
M1 turquoise MEGA module member
K-W ANOVA p: 8.9e-09
SEPT5 UPenn Mixed PRM
K-W ANOVA p: 2.2e-09
bicor=-0.38, p=0.00037
cor=-0.46, p=1.1e-05
bicor=-0.39, p=6.4e-05
cor=-0.45, p=2.6e-06
TMPO UPenn Mixed PRM
M10 purple MEGA module member
K-W ANOVA p: 0.021
TMPO UPenn Mixed PRM
K-W ANOVA p: 0.042
bicor=0.21, p=0.055
cor=0.25, p=0.022
bicor=0.11, p=0.27
cor=0.14, p=0.16
DNM1L UPenn Mixed PRM
M1 turquoise MEGA module member
K-W ANOVA p: 0.39
DNM1L UPenn Mixed PRM
K-W ANOVA p: 0.086
bicor=-0.32, p=0.0029
cor=-0.21, p=0.055
bicor=-0.27, p=0.0074
cor=-0.21, p=0.036
CD44 UPenn Mixed PRM
M4 yellow MEGA module member
K-W ANOVA p: 1.7e-06
CD44 UPenn Mixed PRM
K-W ANOVA p: 0.0019
bicor=0.4, p=0.00015
cor=0.39, p=0.00025
bicor=0.34, p=6e-04
cor=0.3, p=0.0024
Relative Log2 Abundance
Control
AD
ALS
FTLDU
PSP
CBD
PD.PDD
MSA
otherND
Braak Score
CERAD Score

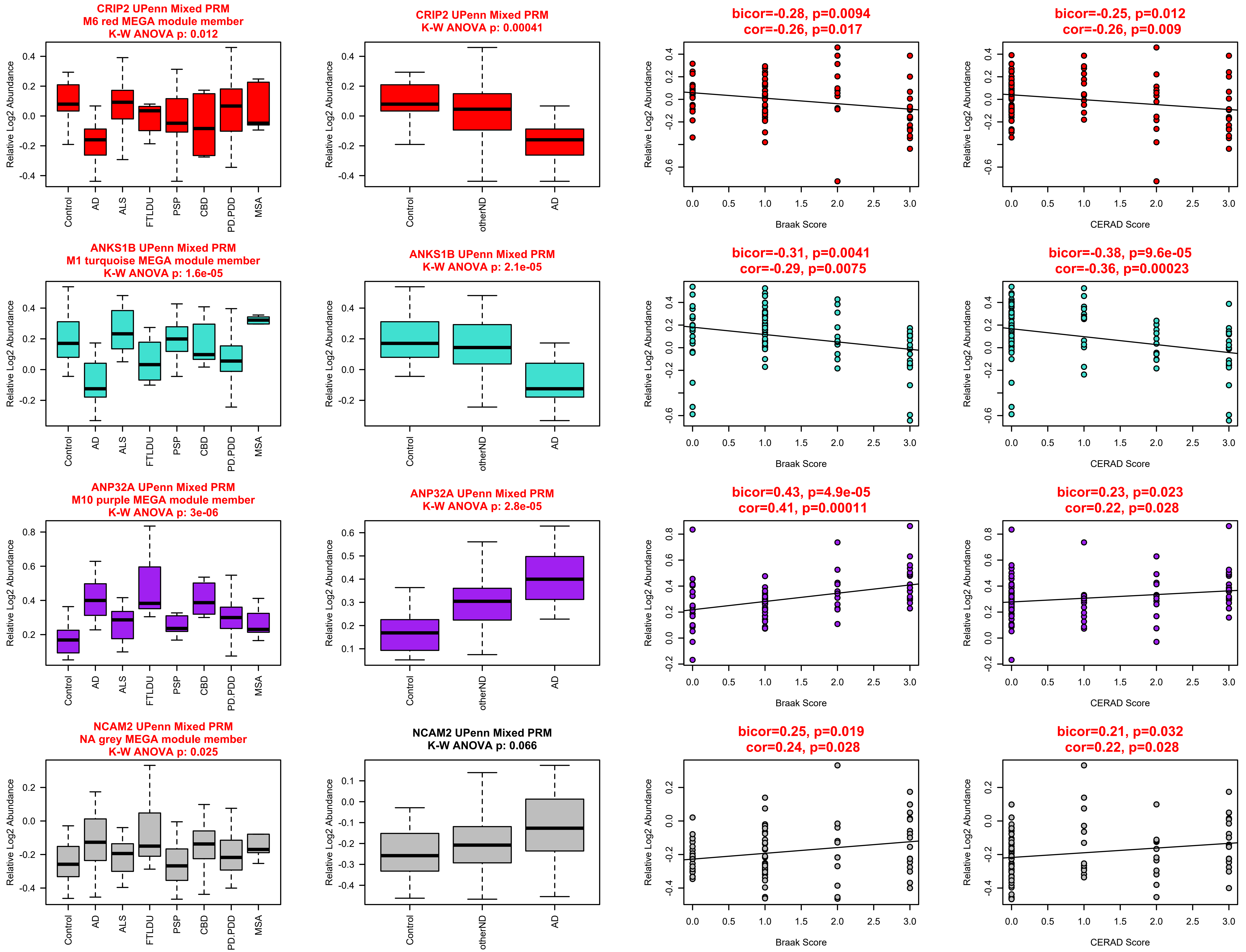
CRIP2 UPenn Mixed PRM
M6 red MEGA module member
K-W ANOVA p: 0.012
CRIP2 UPenn Mixed PRM
K-W ANOVA p: 0.00041
bicor=-0.28, p=0.0094
cor=-0.26, p=0.017
bicor=-0.25, p=0.012
cor=-0.26, p=0.009
ANKS1B UPenn Mixed PRM
M1 turquoise MEGA module member
K-W ANOVA p: 1.6e-05
ANKS1B UPenn Mixed PRM
K-W ANOVA p: 2.1e-05
bicor=-0.31, p=0.0041
cor=-0.29, p=0.0075
bicor=-0.38, p=9.6e-05
cor=-0.36, p=0.00023
ANP32A UPenn Mixed PRM
M10 purple MEGA module member
K-W ANOVA p: 3e-06
ANP32A UPenn Mixed PRM
K-W ANOVA p: 2.8e-05
bicor=0.43, p=4.9e-05
cor=0.41, p=0.00011
bicor=0.23, p=0.023
cor=0.22, p=0.028
NCAM2 UPenn Mixed PRM
NA grey MEGA module member
K-W ANOVA p: 0.025
NCAM2 UPenn Mixed PRM
K-W ANOVA p: 0.066
bicor=0.25, p=0.019
cor=0.24, p=0.028
bicor=0.21, p=0.032
cor=0.22, p=0.028
Relative Log2 Abundance
Control
AD
ALS
FTLDU
PSP
CBD
PD.PDD
MSA
otherND
Braak Score
CERAD Score

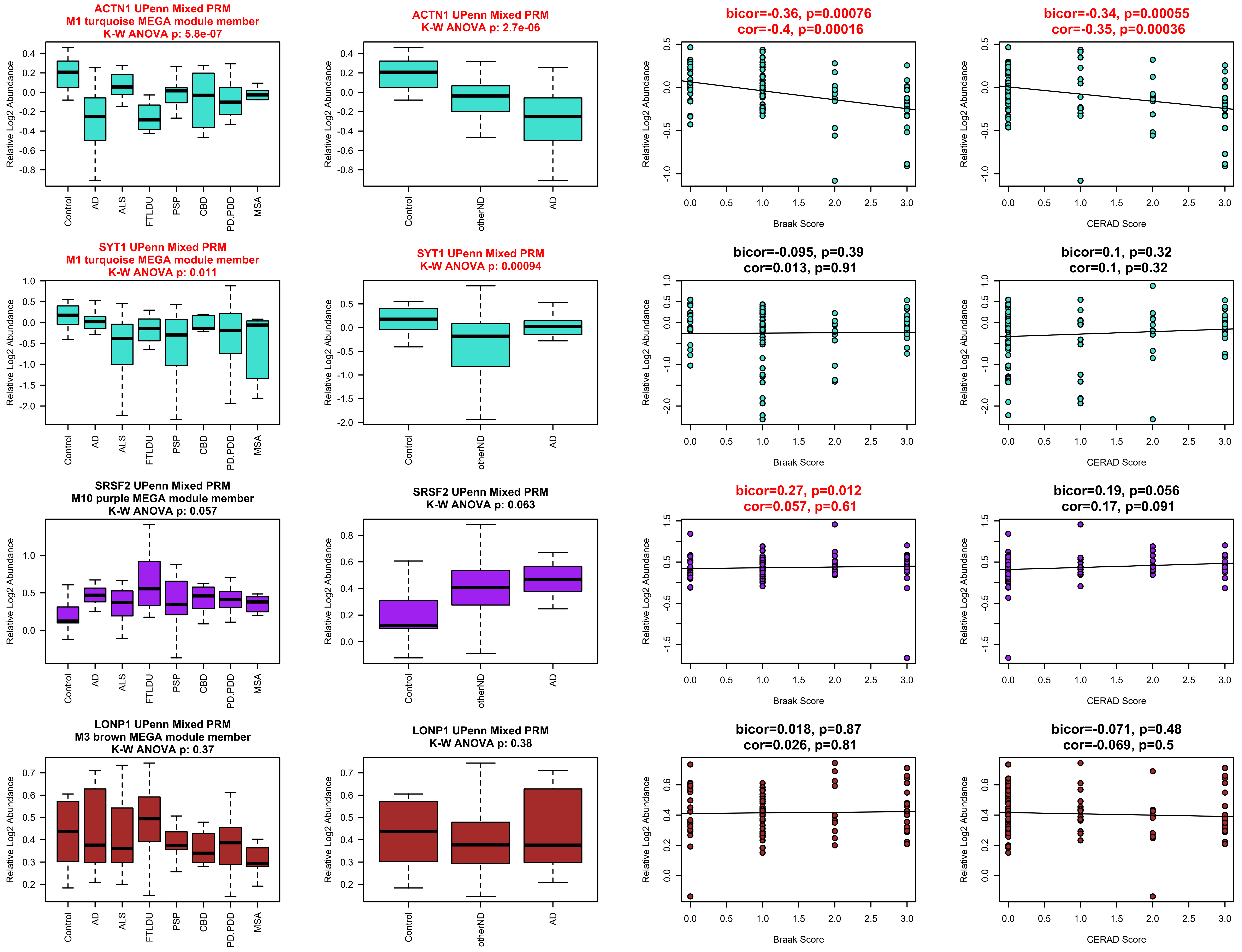
ACTN1 UPenn Mixed PRM
M1 turquoise MEGA module member
K-W ANOVA p: 5.8e-07
ACTN1 UPenn Mixed PRM
K-W ANOVA p: 2.7e-06
bicor=-0.36, p=0.00076
cor=-0.4, p=0.00016
bicor=-0.34, p=0.00055
cor=-0.35, p=0.00036
SYT1 UPenn Mixed PRM
M1 turquoise MEGA module member
K-W ANOVA p: 0.011
SYT1 UPenn Mixed PRM
K-W ANOVA p: 0.00094
bicor=-0.095, p=0.39
cor=0.013, p=0.91
bicor=0.1, p=0.32
cor=0.1, p=0.32
SRSF2 UPenn Mixed PRM
M10 purple MEGA module member
K-W ANOVA p: 0.057
SRSF2 UPenn Mixed PRM
K-W ANOVA p: 0.063
bicor=0.27, p=0.012
cor=0.057, p=0.61
bicor=0.19, p=0.056
cor=0.17, p=0.091
LONP1 UPenn Mixed PRM
M3 brown MEGA module member
K-W ANOVA p: 0.37
LONP1 UPenn Mixed PRM
K-W ANOVA p: 0.38
bicor=0.018, p=0.87
cor=0.026, p=0.81
bicor=-0.071, p=0.48
cor=-0.069, p=0.5
Relative Log2 Abundance
Control
AD
ALS
FTLDU
PSP
CBD
PD.PDD
MSA
otherND
Braak Score
CERAD Score

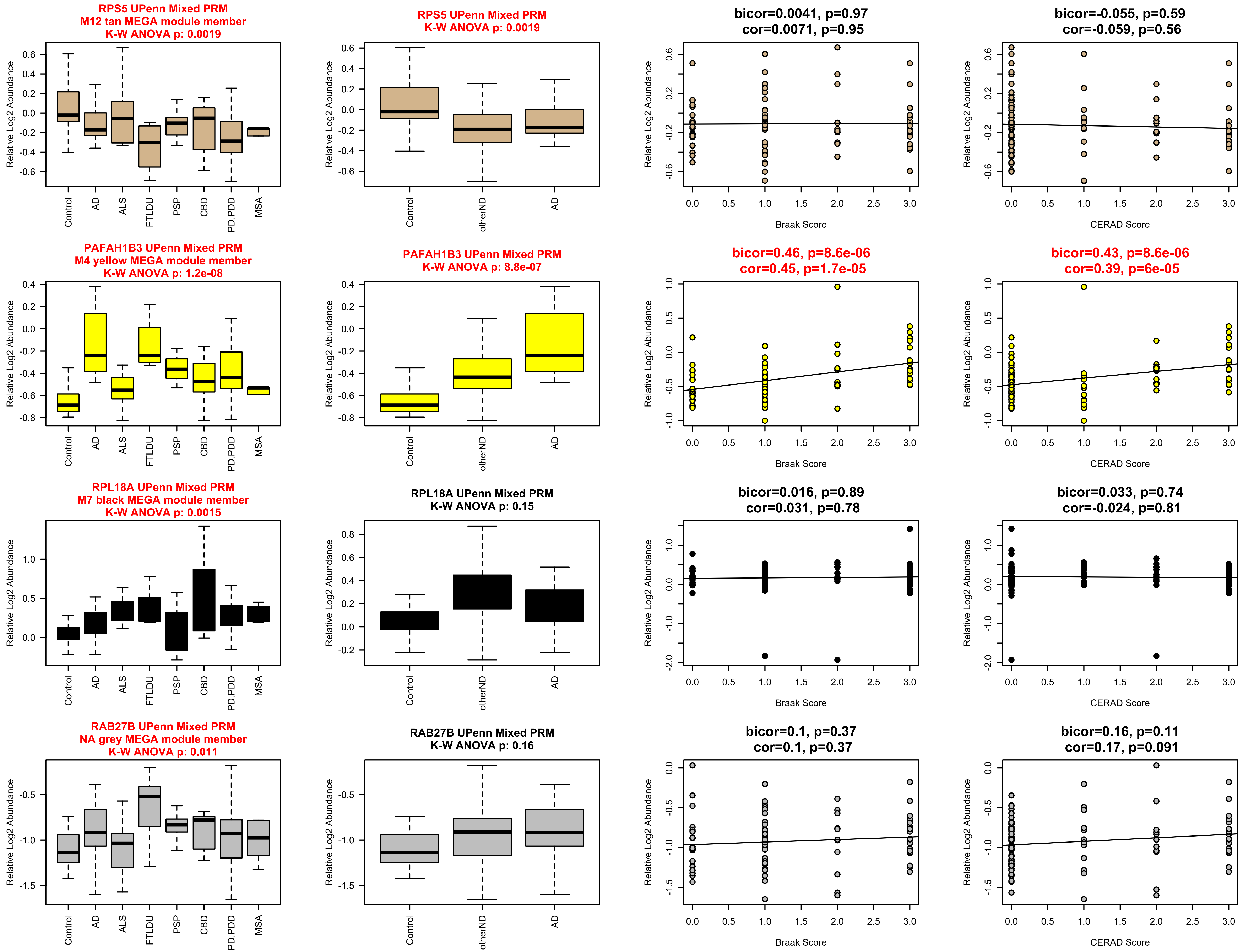
RPS5 UPenn Mixed PRM
M12 tan MEGA module member
K-W ANOVA p: 0.0019
RPS5 UPenn Mixed PRM
K-W ANOVA p: 0.0019
bicor=0.0041, p=0.97
cor=0.0071, p=0.95
bicor=-0.055, p=0.59
cor=-0.059, p=0.56
PAFAH1B3 UPenn Mixed PRM
M4 yellow MEGA module member
K-W ANOVA p: 1.2e-08
PAFAH1B3 UPenn Mixed PRM
K-W ANOVA p: 8.8e-07
bicor=0.46, p=8.6e-06
cor=0.45, p=1.7e-05
bicor=0.43, p=8.6e-06
cor=0.39, p=6e-05
RPL18A UPenn Mixed PRM
M7 black MEGA module member
K-W ANOVA p: 0.0015
RPL18A UPenn Mixed PRM
K-W ANOVA p: 0.15
bicor=0.016, p=0.89
cor=0.031, p=0.78
bicor=0.033, p=0.74
cor=-0.024, p=0.81
RAB27B UPenn Mixed PRM
NA grey MEGA module member
K-W ANOVA p: 0.011
RAB27B UPenn Mixed PRM
K-W ANOVA p: 0.16
bicor=0.1, p=0.37
cor=0.1, p=0.37
bicor=0.16, p=0.11
cor=0.17, p=0.091
Relative Log2 Abundance
Control
AD
ALS
FTLDU
PSP
CBD
PD.PDD
MSA
otherND
Braak Score
CERAD Score

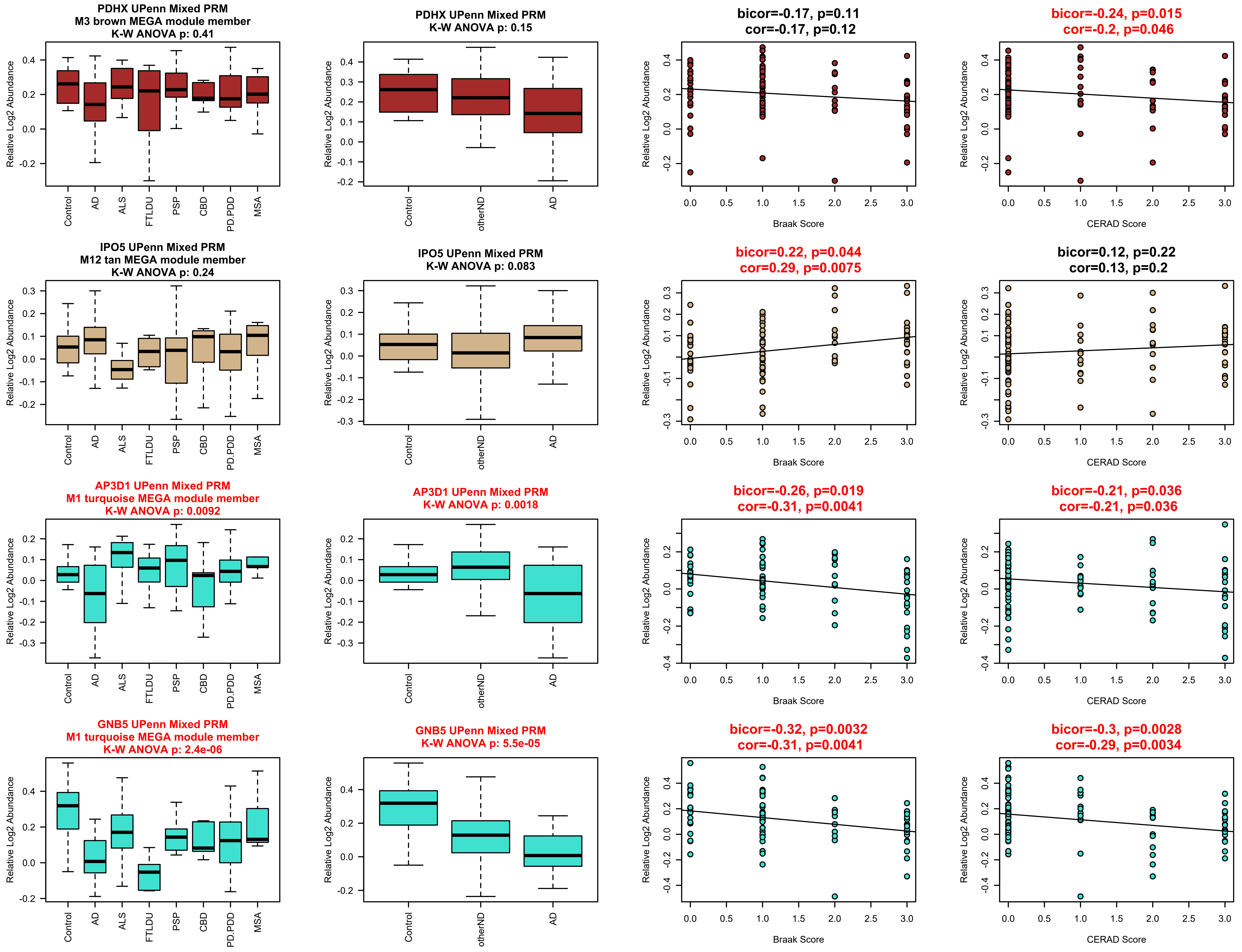
PDHX UPenn Mixed PRM
M3 brown MEGA module member
K-W ANOVA p: 0.41
PDHX UPenn Mixed PRM
K-W ANOVA p: 0.15
bicor=-0.17, p=0.11
cor=-0.17, p=0.12
bicor=-0.24, p=0.015
cor=-0.2, p=0.046
IPO5 UPenn Mixed PRM
M12 tan MEGA module member
K-W ANOVA p: 0.24
IPO5 UPenn Mixed PRM
K-W ANOVA p: 0.083
bicor=0.22, p=0.044
cor=0.29, p=0.0075
bicor=0.12, p=0.22
cor=0.13, p=0.2
AP3D1 UPenn Mixed PRM
M1 turquoise MEGA module member
K-W ANOVA p: 0.0092
AP3D1 UPenn Mixed PRM
K-W ANOVA p: 0.0018
bicor=-0.26, p=0.019
cor=-0.31, p=0.0041
bicor=-0.21, p=0.036
cor=-0.21, p=0.036
GNB5 UPenn Mixed PRM
M1 turquoise MEGA module member
K-W ANOVA p: 2.4e-06
GNB5 UPenn Mixed PRM
K-W ANOVA p: 5.5e-05
bicor=-0.32, p=0.0032
cor=-0.31, p=0.0041
bicor=-0.3, p=0.0028
cor=-0.29, p=0.0034
Relative Log2 Abundance
Control
AD
ALS
FTLDU
PSP
CBD
PD.PDD
MSA
otherND
Braak Score
CERAD Score

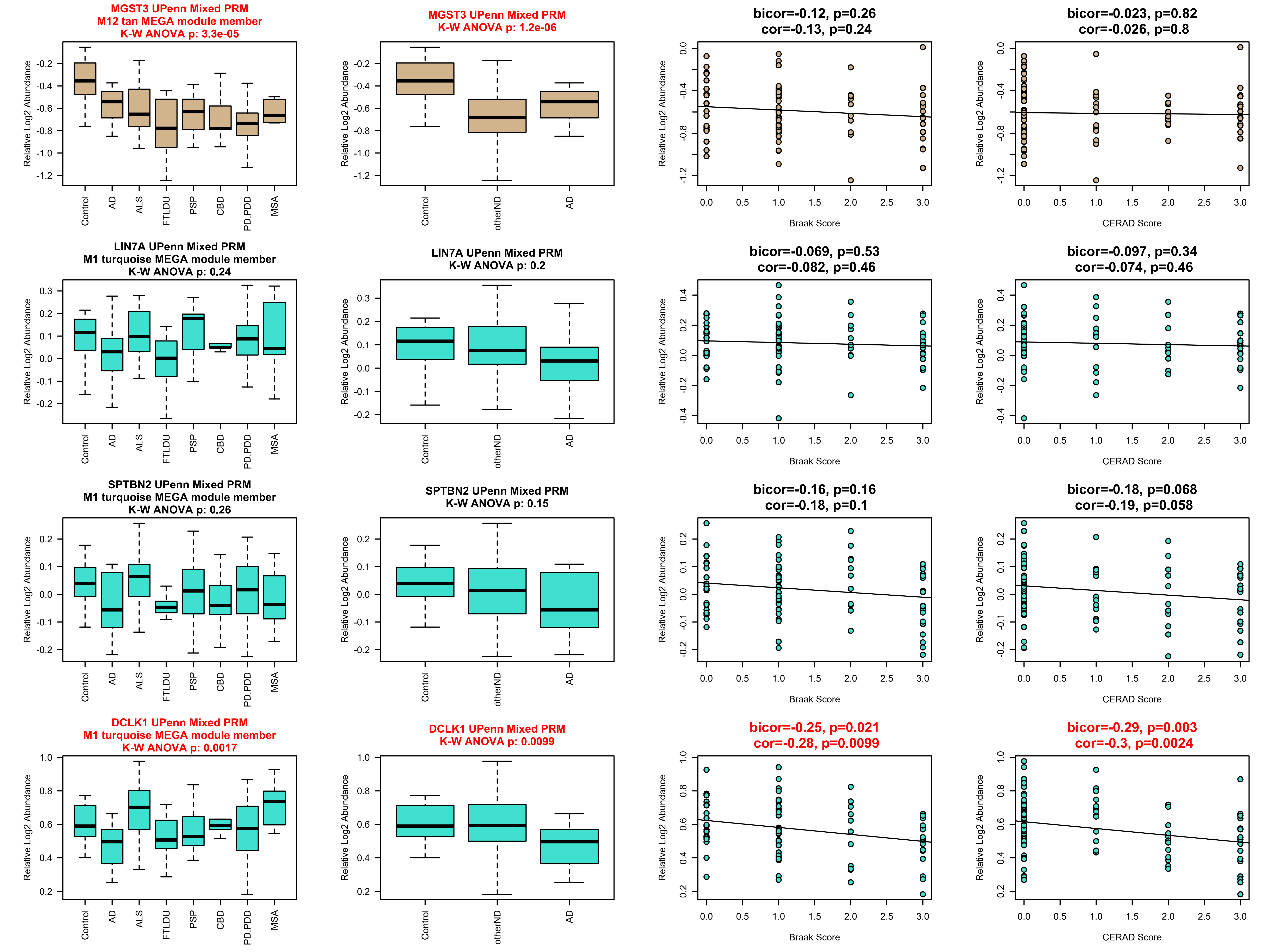
MGST3 UPenn Mixed PRM
M12 tan MEGA module member
K-W ANOVA p: 3.3e-05
MGST3 UPenn Mixed PRM
K-W ANOVA p: 1.2e-06
bicor=-0.12, p=0.26
cor=-0.13, p=0.24
bicor=-0.023, p=0.82
cor=-0.026, p=0.8
LIN7A UPenn Mixed PRM
M1 turquoise MEGA module member
K-W ANOVA p: 0.24
LIN7A UPenn Mixed PRM
K-W ANOVA p: 0.2
bicor=-0.069, p=0.53
cor=-0.082, p=0.46
bicor=-0.097, p=0.34
cor=-0.074, p=0.46
SPTBN2 UPenn Mixed PRM
M1 turquoise MEGA module member
K-W ANOVA p: 0.26
SPTBN2 UPenn Mixed PRM
K-W ANOVA p: 0.15
bicor=-0.16, p=0.16
cor=-0.18, p=0.1
bicor=-0.18, p=0.068
cor=-0.19, p=0.058
DCLK1 UPenn Mixed PRM
M1 turquoise MEGA module member
K-W ANOVA p: 0.0017
DCLK1 UPenn Mixed PRM
K-W ANOVA p: 0.0099
bicor=-0.25, p=0.021
cor=-0.28, p=0.0099
bicor=-0.29, p=0.003
cor=-0.3, p=0.0024
Relative Log2 Abundance
Control
AD
ALS
FTLDU
PSP
CBD
PD.PDD
MSA
otherND
Braak Score
CERAD Score

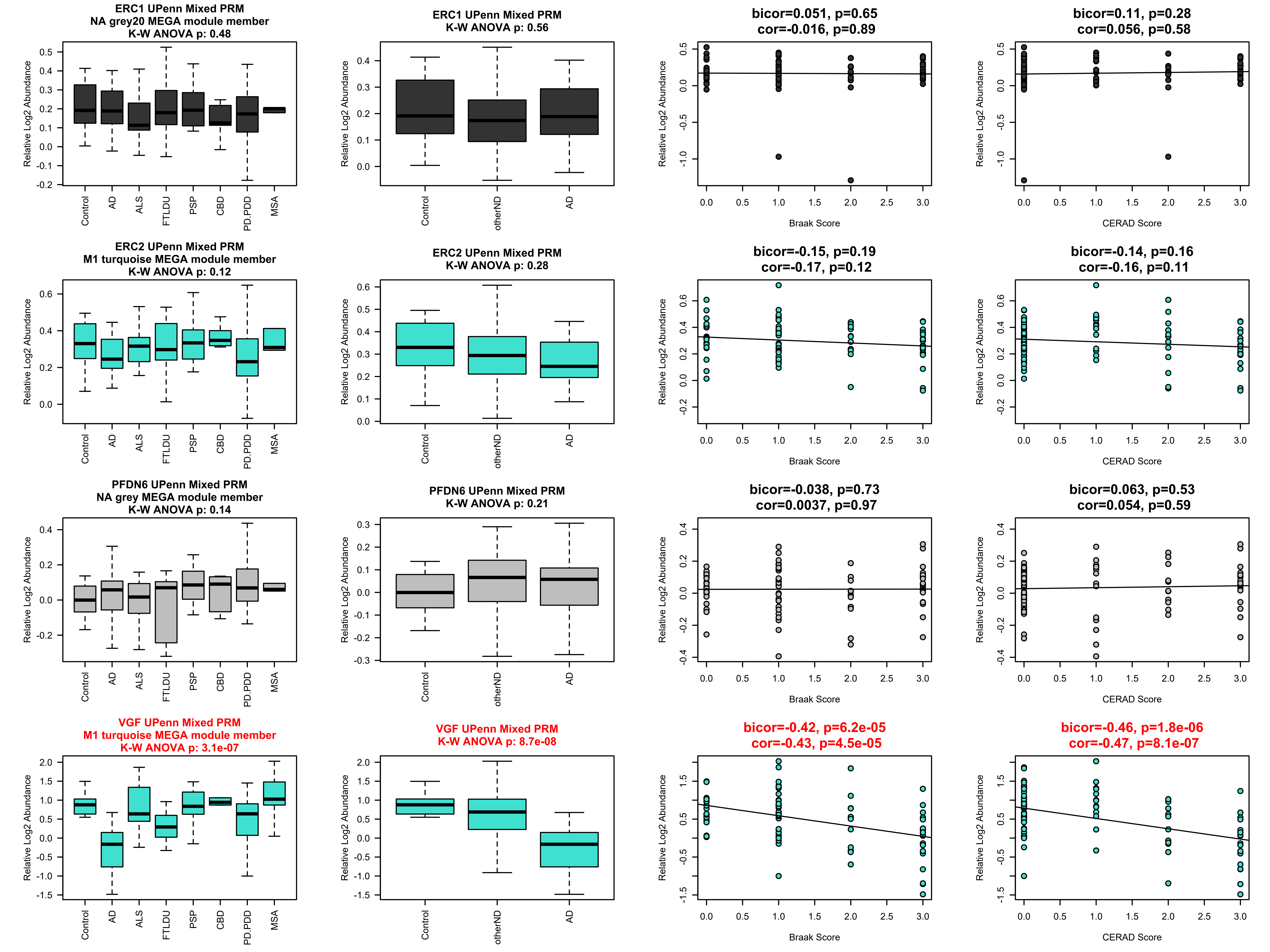
ERC1 UPenn Mixed PRM
NA grey20 MEGA module member
K-W ANOVA p: 0.48
ERC1 UPenn Mixed PRM
K-W ANOVA p: 0.56
bicor=0.051, p=0.65
cor=-0.016, p=0.89
bicor=0.11, p=0.28
cor=0.056, p=0.58
ERC2 UPenn Mixed PRM
M1 turquoise MEGA module member
K-W ANOVA p: 0.12
ERC2 UPenn Mixed PRM
K-W ANOVA p: 0.28
bicor=-0.15, p=0.19
cor=-0.17, p=0.12
bicor=-0.14, p=0.16
cor=-0.16, p=0.11
PFDN6 UPenn Mixed PRM
NA grey MEGA module member
K-W ANOVA p: 0.14
PFDN6 UPenn Mixed PRM
K-W ANOVA p: 0.21
bicor=-0.038, p=0.73
cor=0.0037, p=0.97
bicor=0.063, p=0.53
cor=0.054, p=0.59
VGF UPenn Mixed PRM
M1 turquoise MEGA module member
K-W ANOVA p: 3.1e-07
VGF UPenn Mixed PRM
K-W ANOVA p: 8.7e-08
bicor=-0.42, p=6.2e-05
cor=-0.43, p=4.5e-05
bicor=-0.46, p=1.8e-06
cor=-0.47, p=8.1e-07
Relative Log2 Abundance
Control
AD
ALS
FTLDU
PSP
CBD
PD.PDD
MSA
otherND
Braak Score
CERAD Score

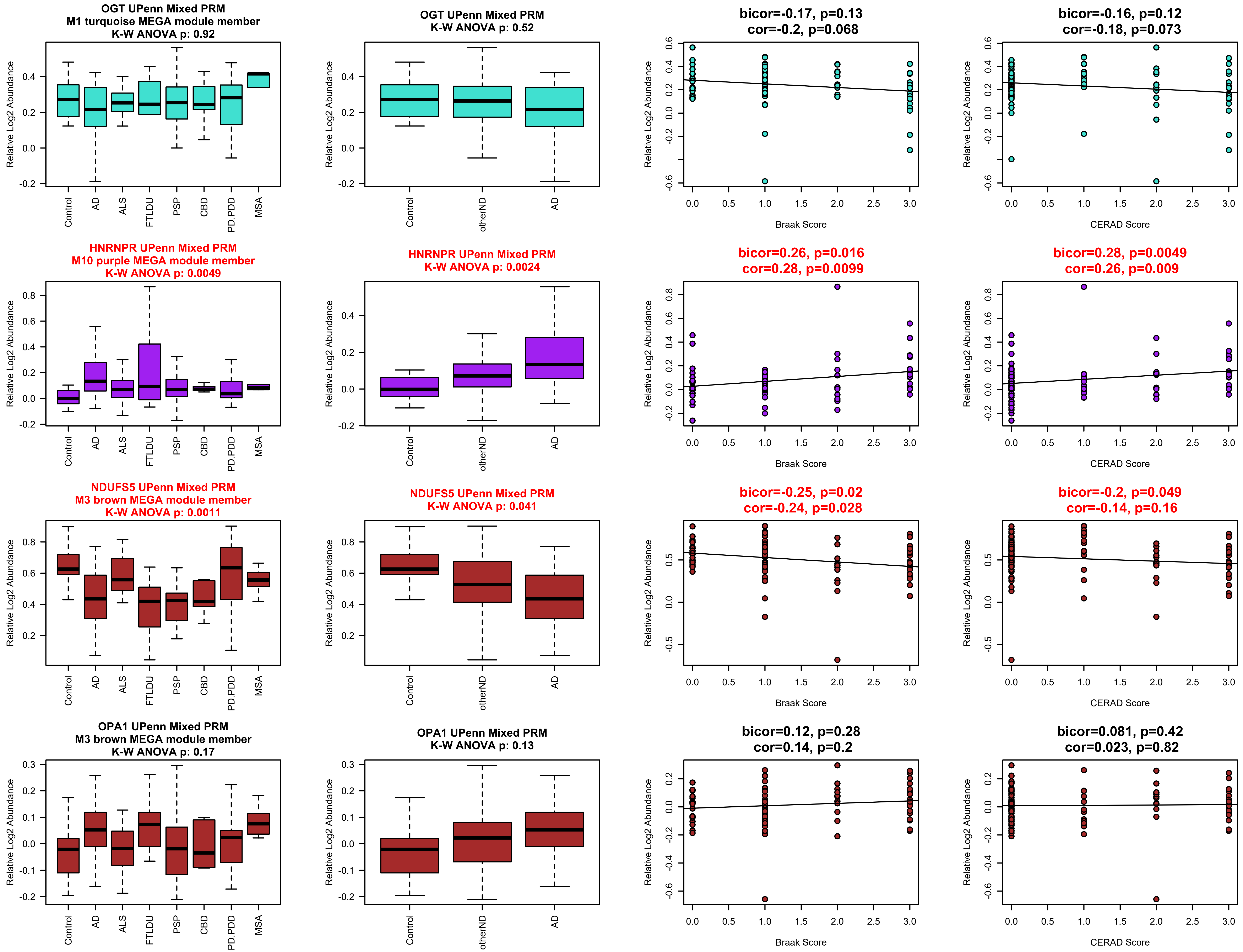
OGT UPenn Mixed PRM
M1 turquoise MEGA module member
K-W ANOVA p: 0.92
OGT UPenn Mixed PRM
K-W ANOVA p: 0.52
bicor=-0.17, p=0.13
cor=-0.2, p=0.068
bicor=-0.16, p=0.12
cor=-0.18, p=0.073
HNRNPR UPenn Mixed PRM
M10 purple MEGA module member
K-W ANOVA p: 0.0049
HNRNPR UPenn Mixed PRM
K-W ANOVA p: 0.0024
bicor=0.26, p=0.016
cor=0.28, p=0.0099
bicor=0.28, p=0.0049
cor=0.26, p=0.009
NDUFS5 UPenn Mixed PRM
M3 brown MEGA module member
K-W ANOVA p: 0.0011
NDUFS5 UPenn Mixed PRM
K-W ANOVA p: 0.041
bicor=-0.25, p=0.02
cor=-0.24, p=0.028
bicor=-0.2, p=0.049
cor=-0.14, p=0.16
OPA1 UPenn Mixed PRM
M3 brown MEGA module member
K-W ANOVA p: 0.17
OPA1 UPenn Mixed PRM
K-W ANOVA p: 0.13
bicor=0.12, p=0.28
cor=0.14, p=0.2
bicor=0.081, p=0.42
cor=0.023, p=0.82
Relative Log2 Abundance
Control
AD
ALS
FTLDU
PSP
CBD
PD.PDD
MSA
otherND
Braak Score
CERAD Score

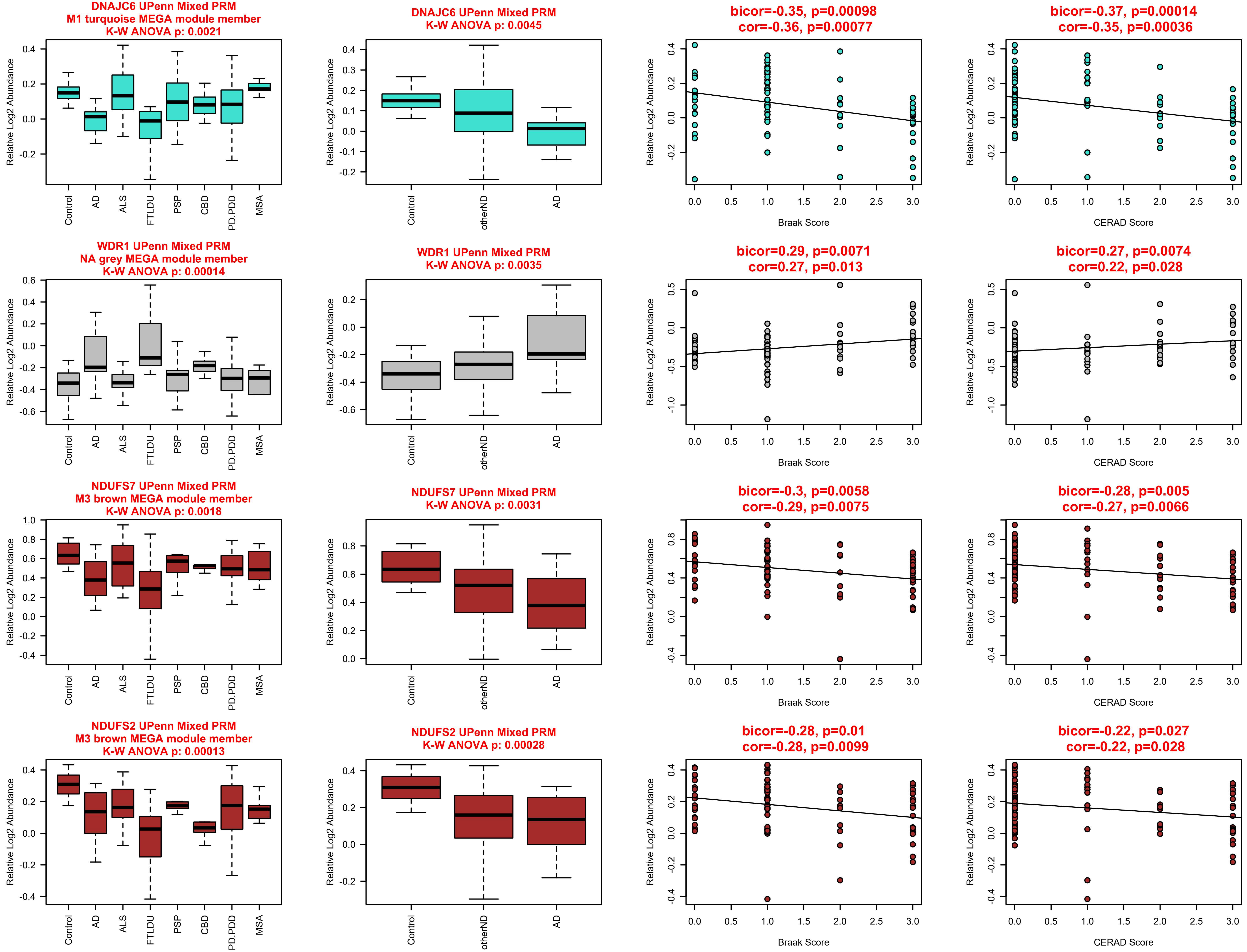
DNAJC6 UPenn Mixed PRM
M1 turquoise MEGA module member
K-W ANOVA p: 0.0021
DNAJC6 UPenn Mixed PRM
K-W ANOVA p: 0.0045
bicor=-0.35, p=0.00098
cor=-0.36, p=0.00077
bicor=-0.37, p=0.00014
cor=-0.35, p=0.00036
WDR1 UPenn Mixed PRM
NA grey MEGA module member
K-W ANOVA p: 0.00014
WDR1 UPenn Mixed PRM
K-W ANOVA p: 0.0035
bicor=0.29, p=0.0071
cor=0.27, p=0.013
bicor=0.27, p=0.0074
cor=0.22, p=0.028
NDUFS7 UPenn Mixed PRM
M3 brown MEGA module member
K-W ANOVA p: 0.0018
NDUFS7 UPenn Mixed PRM
K-W ANOVA p: 0.0031
bicor=-0.3, p=0.0058
cor=-0.29, p=0.0075
bicor=-0.28, p=0.005
cor=-0.27, p=0.0066
NDUFS2 UPenn Mixed PRM
M3 brown MEGA module member
K-W ANOVA p: 0.00013
NDUFS2 UPenn Mixed PRM
K-W ANOVA p: 0.00028
bicor=-0.28, p=0.01
cor=-0.28, p=0.0099
bicor=-0.22, p=0.027
cor=-0.22, p=0.028
Relative Log2 Abundance
Control
AD
ALS
FTLDU
PSP
CBD
PD.PDD
MSA
otherND
Braak Score
CERAD Score
0.0
0.5
1.0
1.5
2.0
2.5
3.0

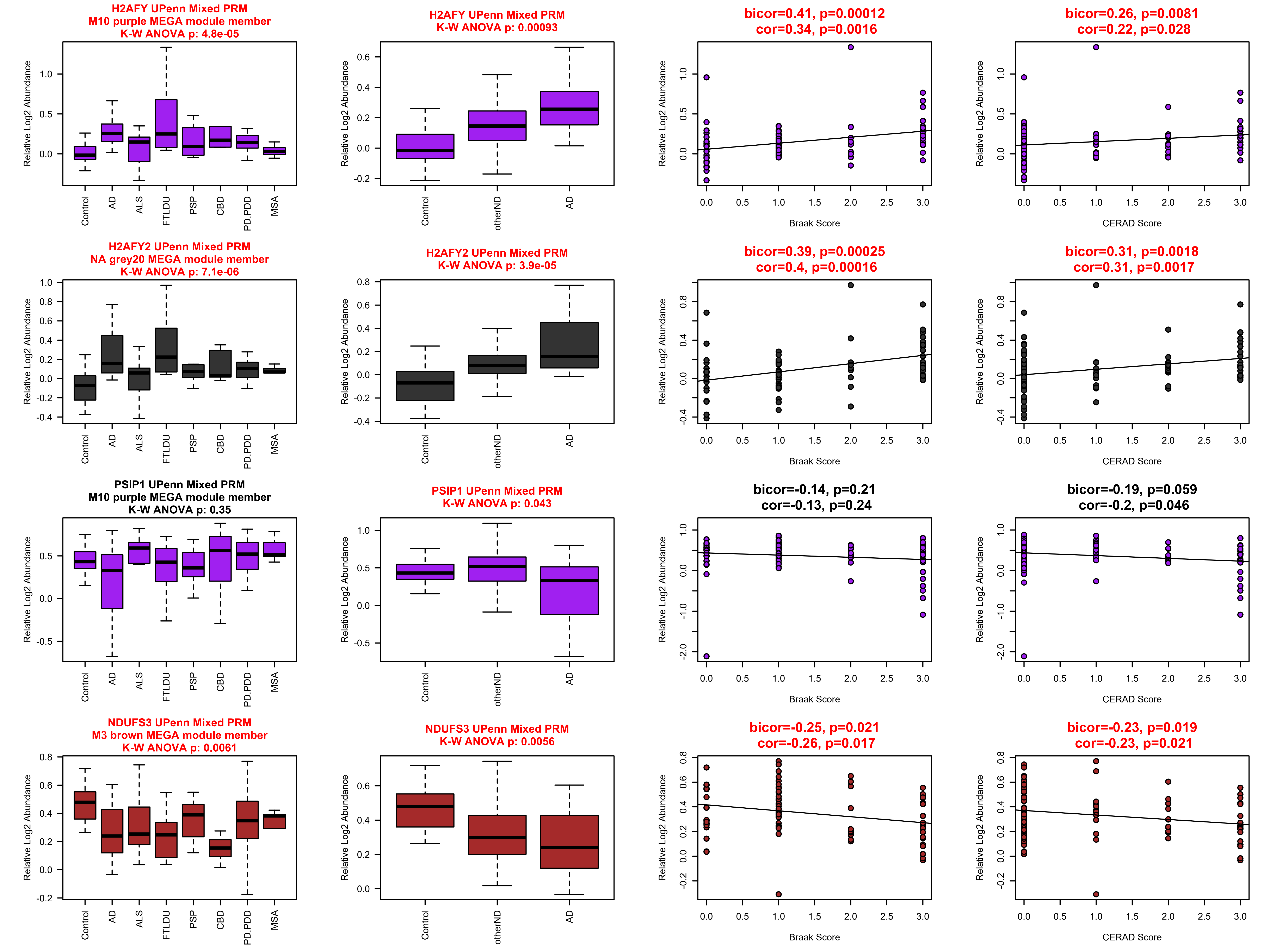
H2AFY UPenn Mixed PRM
M10 purple MEGA module member
K-W ANOVA p: 4.8e-05
H2AFY UPenn Mixed PRM
K-W ANOVA p: 0.00093
bicor=0.41, p=0.00012
cor=0.34, p=0.0016
bicor=0.26, p=0.0081
cor=0.22, p=0.028
H2AFY2 UPenn Mixed PRM
NA grey20 MEGA module member
K-W ANOVA p: 7.1e-06
H2AFY2 UPenn Mixed PRM
K-W ANOVA p: 3.9e-05
bicor=0.39, p=0.00025
cor=0.4, p=0.00016
bicor=0.31, p=0.0018
cor=0.31, p=0.0017
PSIP1 UPenn Mixed PRM
M10 purple MEGA module member
K-W ANOVA p: 0.35
PSIP1 UPenn Mixed PRM
K-W ANOVA p: 0.043
bicor=-0.14, p=0.21
cor=-0.13, p=0.24
bicor=-0.19, p=0.059
cor=-0.2, p=0.046
NDUFS3 UPenn Mixed PRM
M3 brown MEGA module member
K-W ANOVA p: 0.0061
NDUFS3 UPenn Mixed PRM
K-W ANOVA p: 0.0056
bicor=-0.25, p=0.021
cor=-0.26, p=0.017
bicor=-0.23, p=0.019
cor=-0.23, p=0.021
Relative Log2 Abundance
Control
AD
ALS
FTLDU
PSP
CBD
PD.PDD
MSA
otherND
Braak Score
CERAD Score

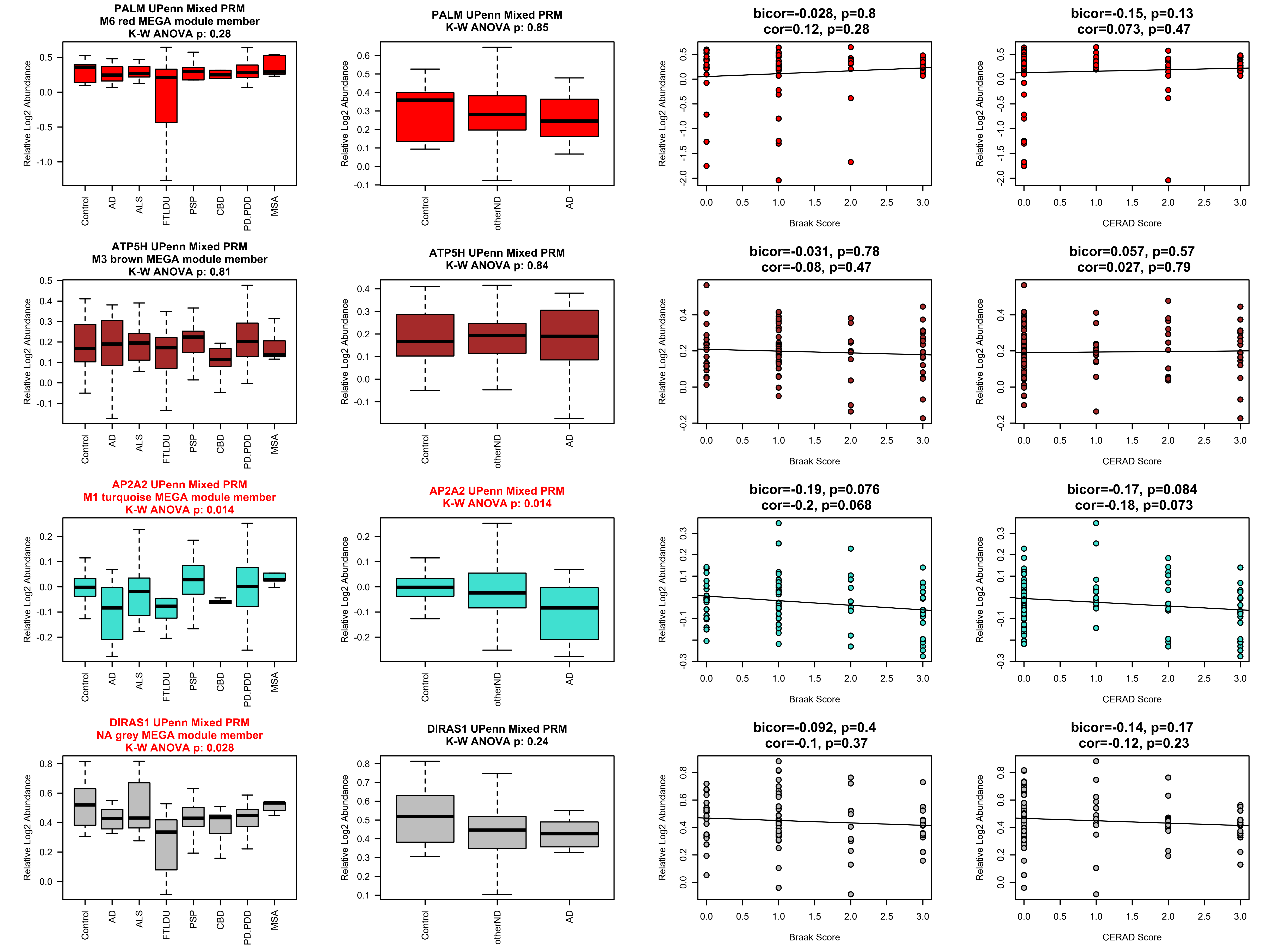

PALM UPenn Mixed PRM
M6 red MEGA module member
K-W ANOVA p: 0.28
PALM UPenn Mixed PRM
K-W ANOVA p: 0.85
bicor=-0.028, p=0.8
cor=0.12, p=0.28
bicor=-0.15, p=0.13
cor=0.073, p=0.47
ATP5H UPenn Mixed PRM
M3 brown MEGA module member
K-W ANOVA p: 0.81
ATP5H UPenn Mixed PRM
K-W ANOVA p: 0.84
bicor=-0.031, p=0.78
cor=-0.08, p=0.47
bicor=0.057, p=0.57
cor=0.027, p=0.79
AP2A2 UPenn Mixed PRM
M1 turquoise MEGA module member
K-W ANOVA p: 0.014
AP2A2 UPenn Mixed PRM
K-W ANOVA p: 0.014
bicor=-0.19, p=0.076
cor=-0.2, p=0.068
bicor=-0.17, p=0.084
cor=-0.18, p=0.073
DIRAS1 UPenn Mixed PRM
NA grey MEGA module member
K-W ANOVA p: 0.028
DIRAS1 UPenn Mixed PRM
K-W ANOVA p: 0.24
bicor=-0.092, p=0.4
cor=-0.1, p=0.37
bicor=-0.14, p=0.17
cor=-0.12, p=0.23
Relative Log2 Abundance
Control
AD
ALS
FTLDU
PSP
CBD
PD.PDD
MSA
otherND
Braak Score
CERAD Score

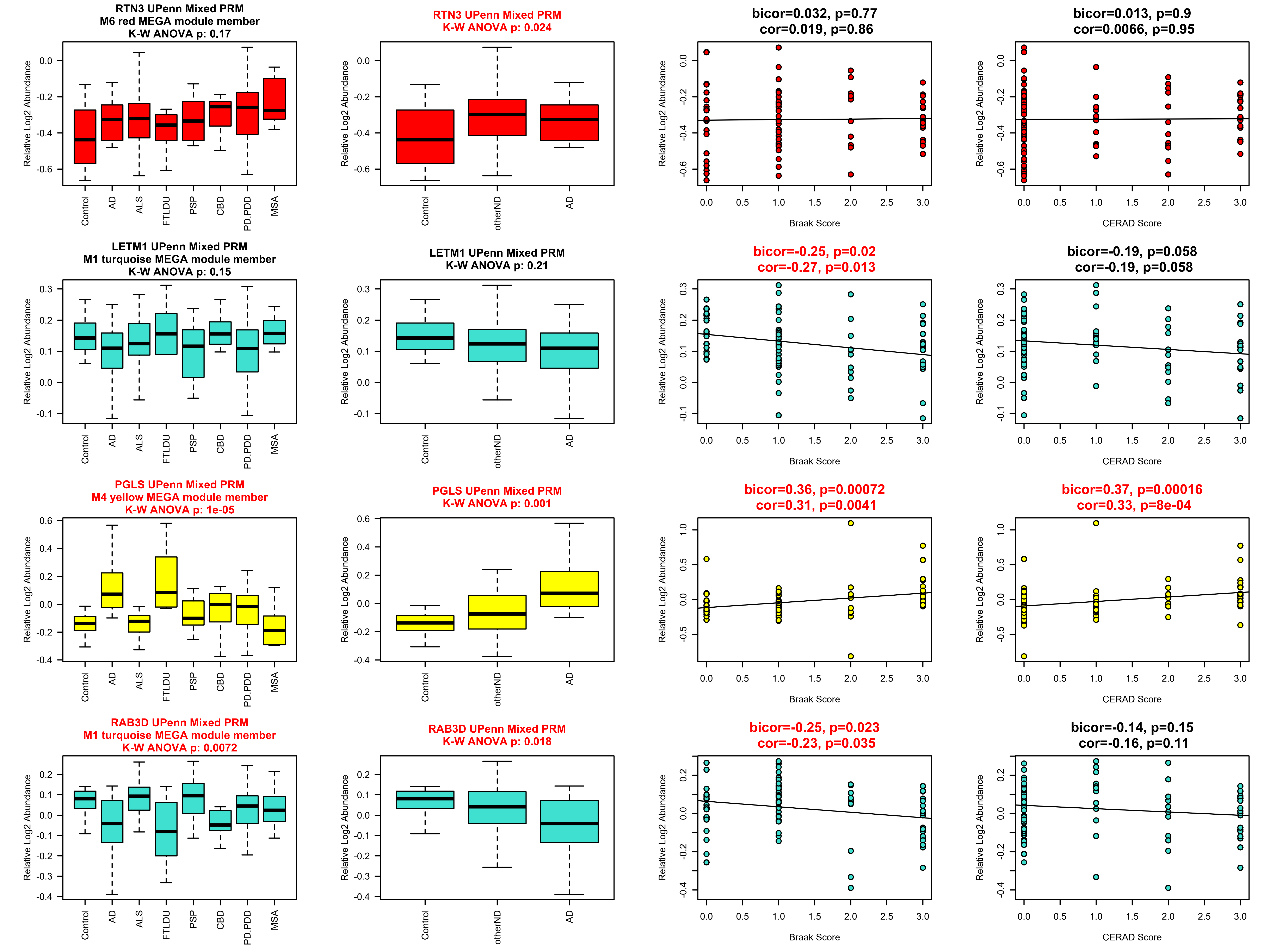
RTN3 UPenn Mixed PRM
M6 red MEGA module member
K-W ANOVA p: 0.17
RTN3 UPenn Mixed PRM
K-W ANOVA p: 0.024
bicor=0.032, p=0.77
cor=0.019, p=0.86
bicor=0.013, p=0.9
cor=0.0066, p=0.95
LETM1 UPenn Mixed PRM
M1 turquoise MEGA module member
K-W ANOVA p: 0.15
LETM1 UPenn Mixed PRM
K-W ANOVA p: 0.21
bicor=-0.25, p=0.02
cor=-0.27, p=0.013
bicor=-0.19, p=0.058
cor=-0.19, p=0.058
PGLS UPenn Mixed PRM
M4 yellow MEGA module member
K-W ANOVA p: 1e-05
PGLS UPenn Mixed PRM
K-W ANOVA p: 0.001
bicor=0.36, p=0.00072
cor=0.31, p=0.0041
bicor=0.37, p=0.00016
cor=0.33, p=8e-04
RAB3D UPenn Mixed PRM
M1 turquoise MEGA module member
K-W ANOVA p: 0.0072
RAB3D UPenn Mixed PRM
K-W ANOVA p: 0.018
bicor=-0.25, p=0.023
cor=-0.23, p=0.035
bicor=-0.14, p=0.15
cor=-0.16, p=0.11
Relative Log2 Abundance
Control
AD
ALS
FTLDU
PSP
CBD
PD.PDD
MSA
otherND
Braak Score
CERAD Score

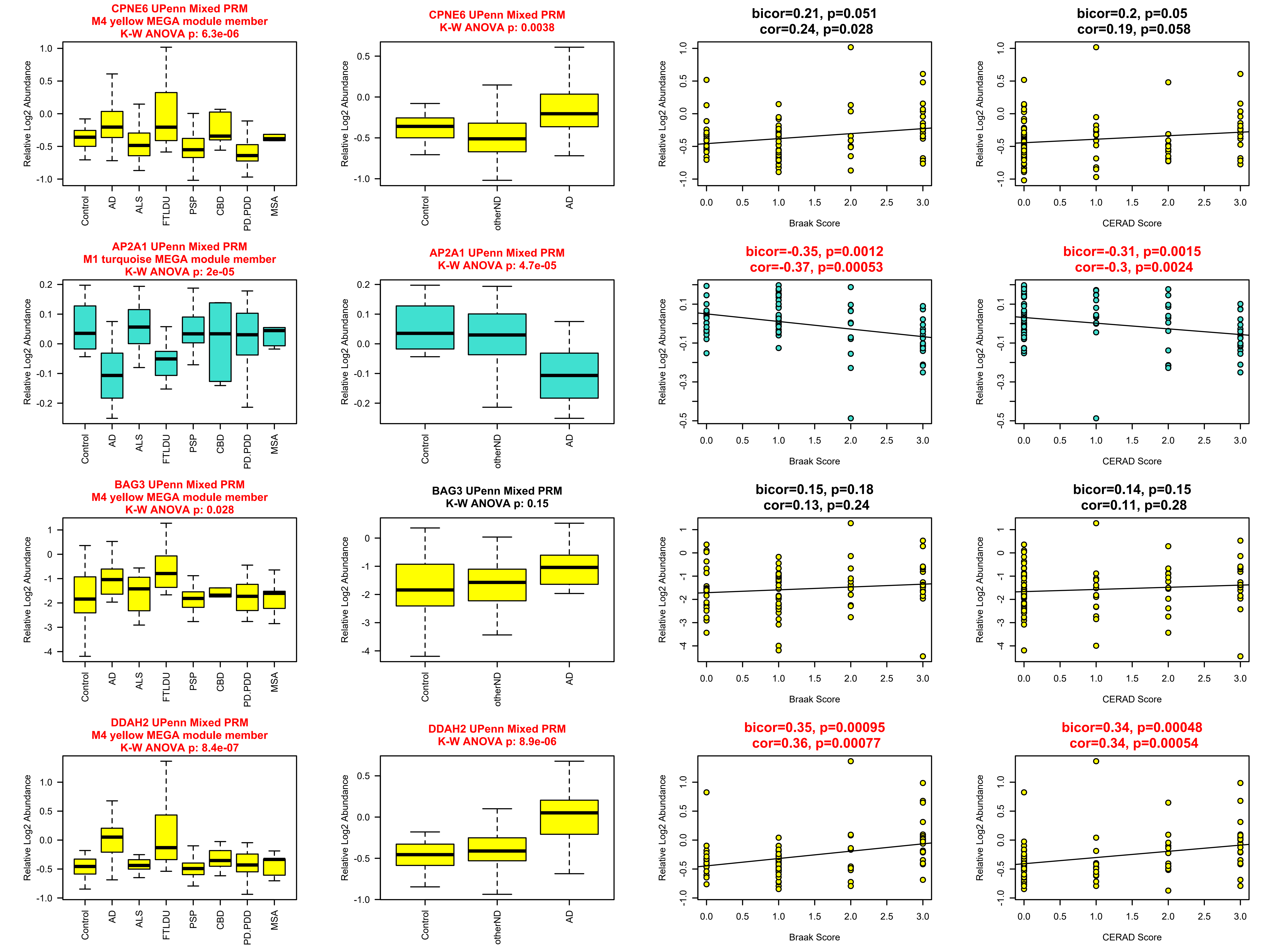
CPNE6 UPenn Mixed PRM
M4 yellow MEGA module member
K-W ANOVA p: 6.3e-06
CPNE6 UPenn Mixed PRM
K-W ANOVA p: 0.0038
bicor=0.21, p=0.051
cor=0.24, p=0.028
bicor=0.2, p=0.05
cor=0.19, p=0.058
AP2A1 UPenn Mixed PRM
M1 turquoise MEGA module member
K-W ANOVA p: 2e-05
AP2A1 UPenn Mixed PRM
K-W ANOVA p: 4.7e-05
bicor=-0.35, p=0.0012
cor=-0.37, p=0.00053
bicor=-0.31, p=0.0015
cor=-0.3, p=0.0024
BAG3 UPenn Mixed PRM
M4 yellow MEGA module member
K-W ANOVA p: 0.028
BAG3 UPenn Mixed PRM
K-W ANOVA p: 0.15
bicor=0.15, p=0.18
cor=0.13, p=0.24
bicor=0.14, p=0.15
cor=0.11, p=0.28
DDAH2 UPenn Mixed PRM
M4 yellow MEGA module member
K-W ANOVA p: 8.4e-07
DDAH2 UPenn Mixed PRM
K-W ANOVA p: 8.9e-06
bicor=0.35, p=0.00095
cor=0.36, p=0.00077
bicor=0.34, p=0.00048
cor=0.34, p=0.00054
Relative Log2 Abundance
Control
AD
ALS
FTLDU
PSP
CBD
PD.PDD
MSA
otherND
Braak Score
CERAD Score

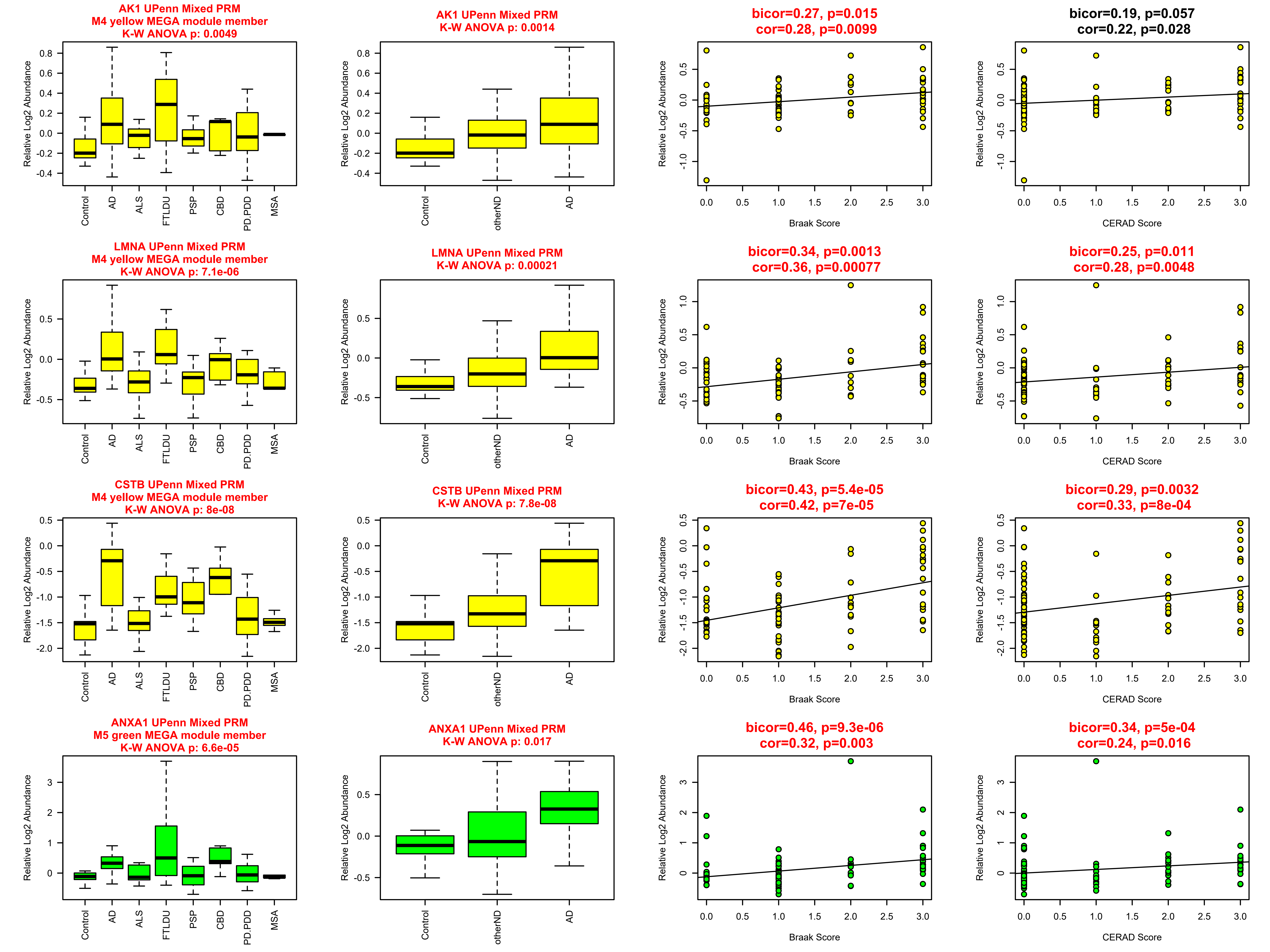
AK1 UPenn Mixed PRM
M4 yellow MEGA module member
K-W ANOVA p: 0.0049
AK1 UPenn Mixed PRM
K-W ANOVA p: 0.0014
bicor=0.27, p=0.015
cor=0.28, p=0.0099
bicor=0.19, p=0.057
cor=0.22, p=0.028
LMNA UPenn Mixed PRM
M4 yellow MEGA module member
K-W ANOVA p: 7.1e-06
LMNA UPenn Mixed PRM
K-W ANOVA p: 0.00021
bicor=0.34, p=0.0013
cor=0.36, p=0.00077
bicor=0.25, p=0.011
cor=0.28, p=0.0048
CSTB UPenn Mixed PRM
M4 yellow MEGA module member
K-W ANOVA p: 8e-08
CSTB UPenn Mixed PRM
K-W ANOVA p: 7.8e-08
bicor=0.43, p=5.4e-05
cor=0.42, p=7e-05
bicor=0.29, p=0.0032
cor=0.33, p=8e-04
ANXA1 UPenn Mixed PRM
M5 green MEGA module member
K-W ANOVA p: 6.6e-05
ANXA1 UPenn Mixed PRM
K-W ANOVA p: 0.017
bicor=0.46, p=9.3e-06
cor=0.32, p=0.003
bicor=0.34, p=5e-04
cor=0.24, p=0.016
Relative Log2 Abundance
Control
AD
ALS
FTLDU
PSP
CBD
PD.PDD
MSA
otherND
Braak Score
CERAD Score

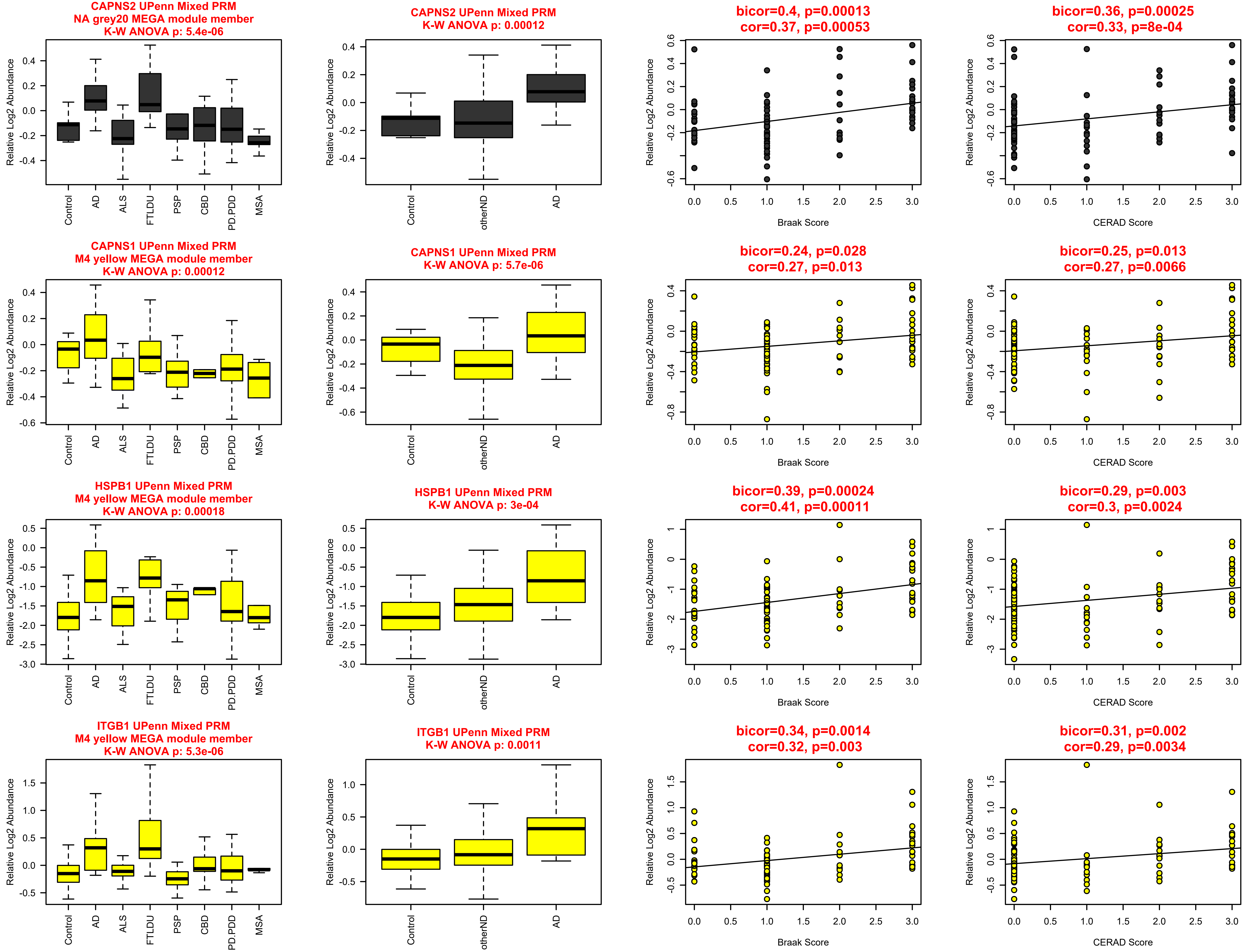
CAPNS2 UPenn Mixed PRM
NA grey20 MEGA module member
K-W ANOVA p: 5.4e-06
CAPNS2 UPenn Mixed PRM
K-W ANOVA p: 0.00012
bicor=0.4, p=0.00013
cor=0.37, p=0.00053
bicor=0.36, p=0.00025
cor=0.33, p=8e-04
CAPNS1 UPenn Mixed PRM
M4 yellow MEGA module member
K-W ANOVA p: 0.00012
CAPNS1 UPenn Mixed PRM
K-W ANOVA p: 5.7e-06
bicor=0.24, p=0.028
cor=0.27, p=0.013
bicor=0.25, p=0.013
cor=0.27, p=0.0066
HSPB1 UPenn Mixed PRM
M4 yellow MEGA module member
K-W ANOVA p: 0.00018
HSPB1 UPenn Mixed PRM
K-W ANOVA p: 3e-04
bicor=0.39, p=0.00024
cor=0.41, p=0.00011
bicor=0.29, p=0.003
cor=0.3, p=0.0024
ITGB1 UPenn Mixed PRM
M4 yellow MEGA module member
K-W ANOVA p: 5.3e-06
ITGB1 UPenn Mixed PRM
K-W ANOVA p: 0.0011
bicor=0.34, p=0.0014
cor=0.32, p=0.003
bicor=0.31, p=0.002
cor=0.29, p=0.0034
Relative Log2 Abundance
Control
AD
ALS
FTLDU
PSP
CBD
PD.PDD
MSA
otherND
Braak Score
CERAD Score

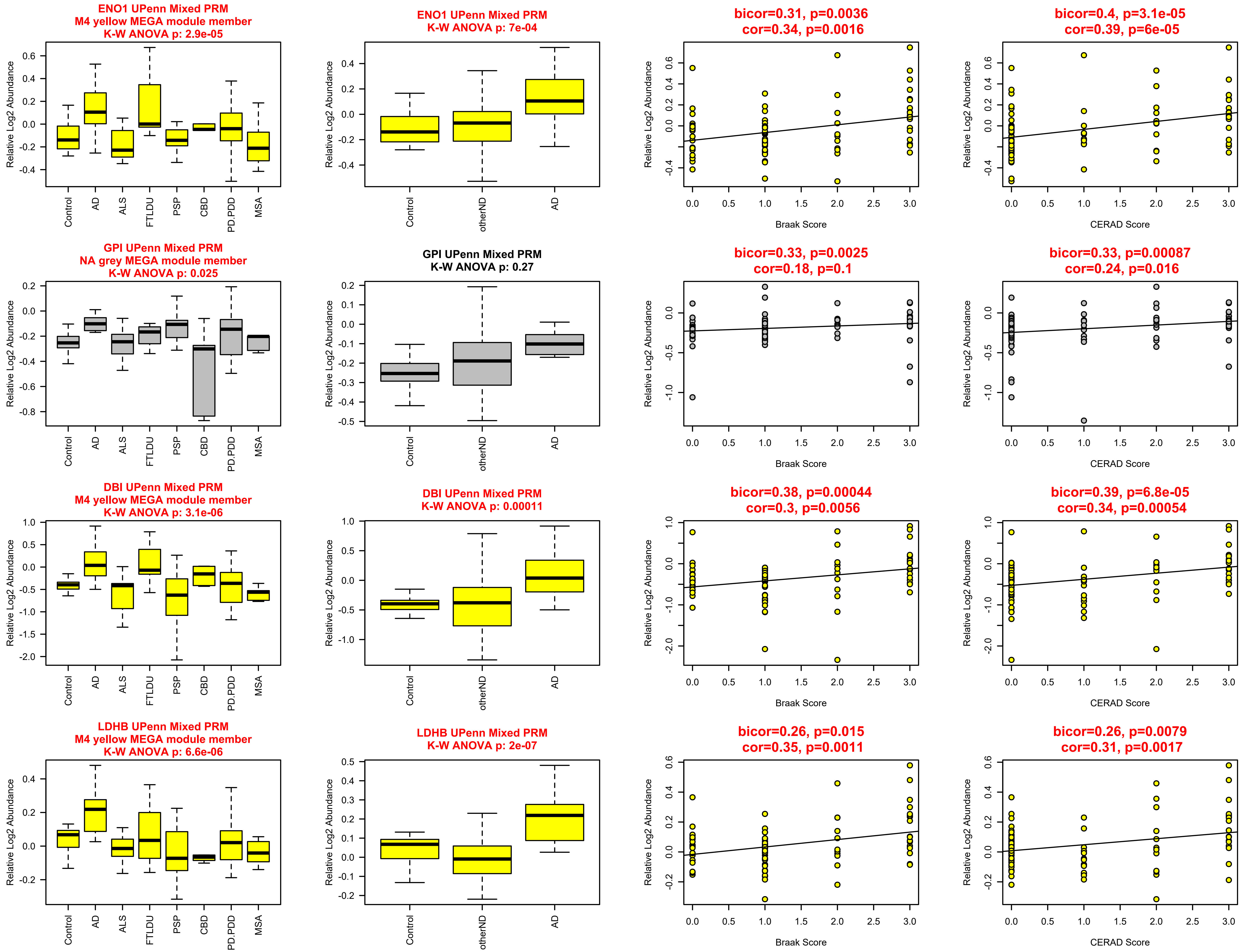

ENO1 UPenn Mixed PRM
M4 yellow MEGA module member
K-W ANOVA p: 2.9e-05
Relative Log2 Abundance
Control
AD
ALS
FTLDU
PSP
CBD
PD.PDD
MSA
ENO1 UPenn Mixed PRM
K-W ANOVA p: 7e-04
otherND
bicor=0.31, p=0.0036
cor=0.34, p=0.0016
Braak Score
bicor=0.4, p=3.1e-05
cor=0.39, p=6e-05
CERAD Score
GPI UPenn Mixed PRM
NA grey MEGA module member
K-W ANOVA p: 0.025
GPI UPenn Mixed PRM
K-W ANOVA p: 0.27
bicor=0.33, p=0.0025
cor=0.18, p=0.1
bicor=0.33, p=0.00087
cor=0.24, p=0.016
DBI UPenn Mixed PRM
M4 yellow MEGA module member
K-W ANOVA p: 3.1e-06
DBI UPenn Mixed PRM
K-W ANOVA p: 0.00011
bicor=0.38, p=0.00044
cor=0.3, p=0.0056
bicor=0.39, p=6.8e-05
cor=0.34, p=0.00054
LDHB UPenn Mixed PRM
M4 yellow MEGA module member
K-W ANOVA p: 6.6e-06
LDHB UPenn Mixed PRM
K-W ANOVA p: 2e-07
bicor=0.26, p=0.015
cor=0.35, p=0.0011
bicor=0.26, p=0.0079
cor=0.31, p=0.0017

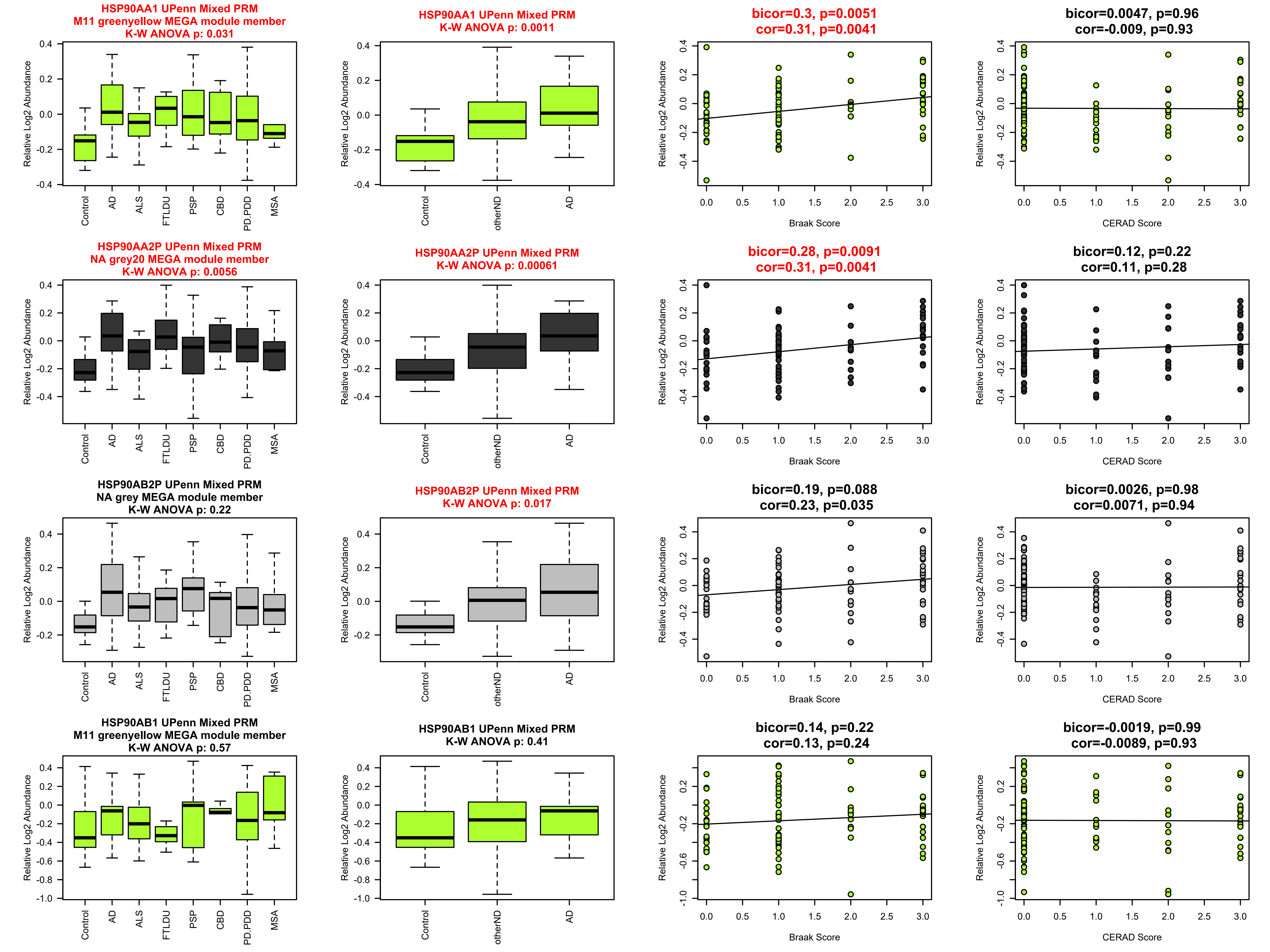
HSP90AA1 UPenn Mixed PRM
M11 greenyellow MEGA module member
K-W ANOVA p: 0.031
HSP90AA1 UPenn Mixed PRM
K-W ANOVA p: 0.0011
bicor=0.3, p=0.0051
cor=0.31, p=0.0041
bicor=0.0047, p=0.96
cor=-0.009, p=0.93
HSP90AA2P UPenn Mixed PRM
NA grey20 MEGA module member
K-W ANOVA p: 0.0056
HSP90AA2P UPenn Mixed PRM
K-W ANOVA p: 0.00061
bicor=0.28, p=0.0091
cor=0.31, p=0.0041
bicor=0.12, p=0.22
cor=0.11, p=0.28
HSP90AB2P UPenn Mixed PRM
NA grey MEGA module member
K-W ANOVA p: 0.22
HSP90AB2P UPenn Mixed PRM
K-W ANOVA p: 0.017
bicor=0.19, p=0.088
cor=0.23, p=0.035
bicor=0.0026, p=0.98
cor=0.0071, p=0.94
HSP90AB1 UPenn Mixed PRM
M11 greenyellow MEGA module member
K-W ANOVA p: 0.57
HSP90AB1 UPenn Mixed PRM
K-W ANOVA p: 0.41
bicor=0.14, p=0.22
cor=0.13, p=0.24
bicor=-0.0019, p=0.99
cor=-0.0089, p=0.93
Relative Log2 Abundance
Control
AD
ALS
FTLDU
PSP
CBD
PD.PDD
MSA
otherND
Braak Score
CERAD Score

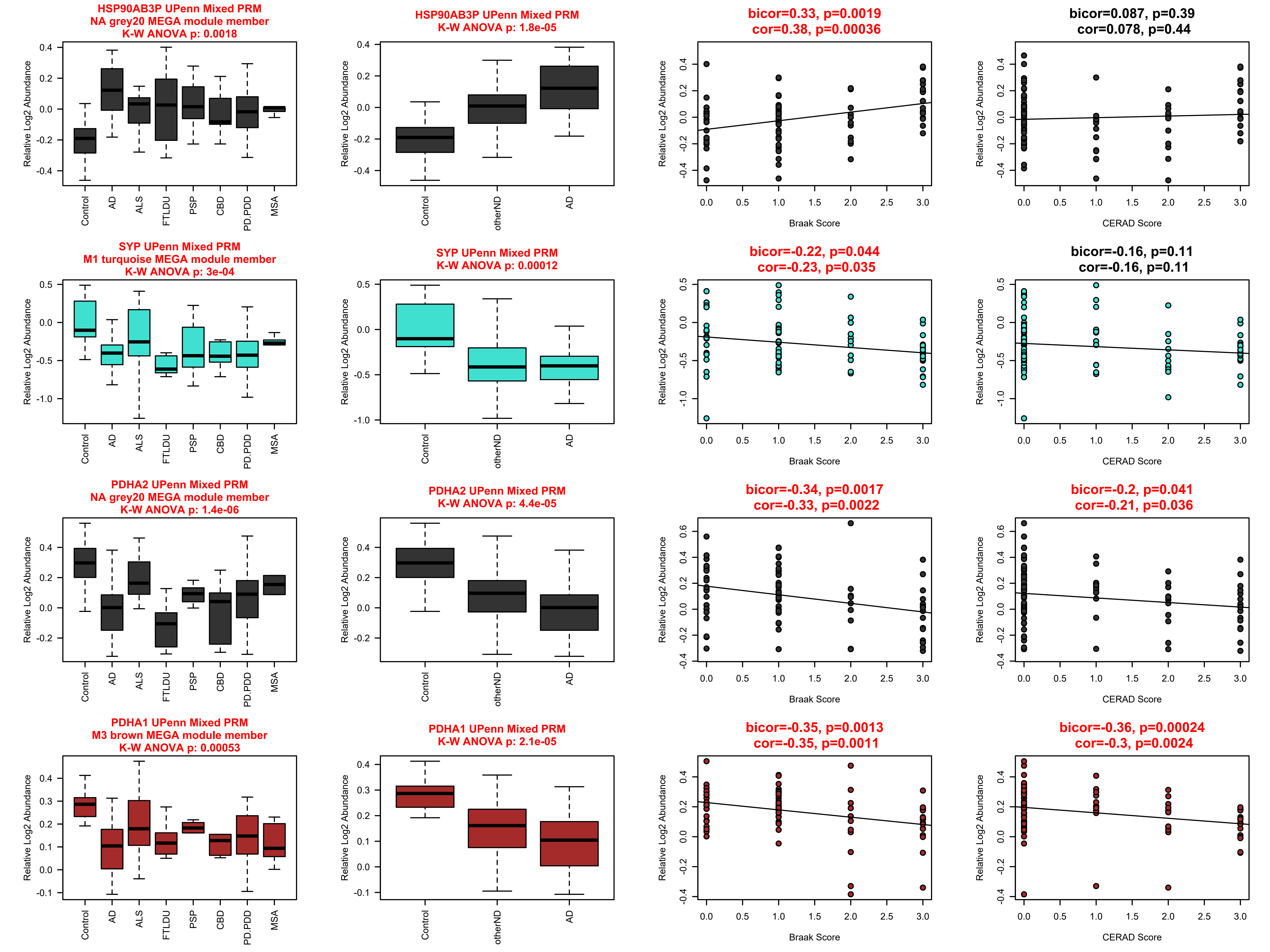
HSP90AB3P UPenn Mixed PRM
NA grey20 MEGA module member
K-W ANOVA p: 0.0018
HSP90AB3P UPenn Mixed PRM
K-W ANOVA p: 1.8e-05
bicor=0.33, p=0.0019
cor=0.38, p=0.00036
bicor=0.087, p=0.39
cor=0.078, p=0.44
SYP UPenn Mixed PRM
M1 turquoise MEGA module member
K-W ANOVA p: 3e-04
SYP UPenn Mixed PRM
K-W ANOVA p: 0.00012
bicor=-0.22, p=0.044
cor=-0.23, p=0.035
bicor=-0.16, p=0.11
cor=-0.16, p=0.11
PDHA2 UPenn Mixed PRM
NA grey20 MEGA module member
K-W ANOVA p: 1.4e-06
PDHA2 UPenn Mixed PRM
K-W ANOVA p: 4.4e-05
bicor=-0.34, p=0.0017
cor=-0.33, p=0.0022
bicor=-0.2, p=0.041
cor=-0.21, p=0.036
PDHA1 UPenn Mixed PRM
M3 brown MEGA module member
K-W ANOVA p: 0.00053
PDHA1 UPenn Mixed PRM
K-W ANOVA p: 2.1e-05
bicor=-0.35, p=0.0013
cor=-0.35, p=0.0011
bicor=-0.36, p=0.00024
cor=-0.3, p=0.0024
Relative Log2 Abundance
Control
AD
ALS
FTLDU
PSP
CBD
PD.PDD
MSA
otherND
Braak Score
CERAD Score

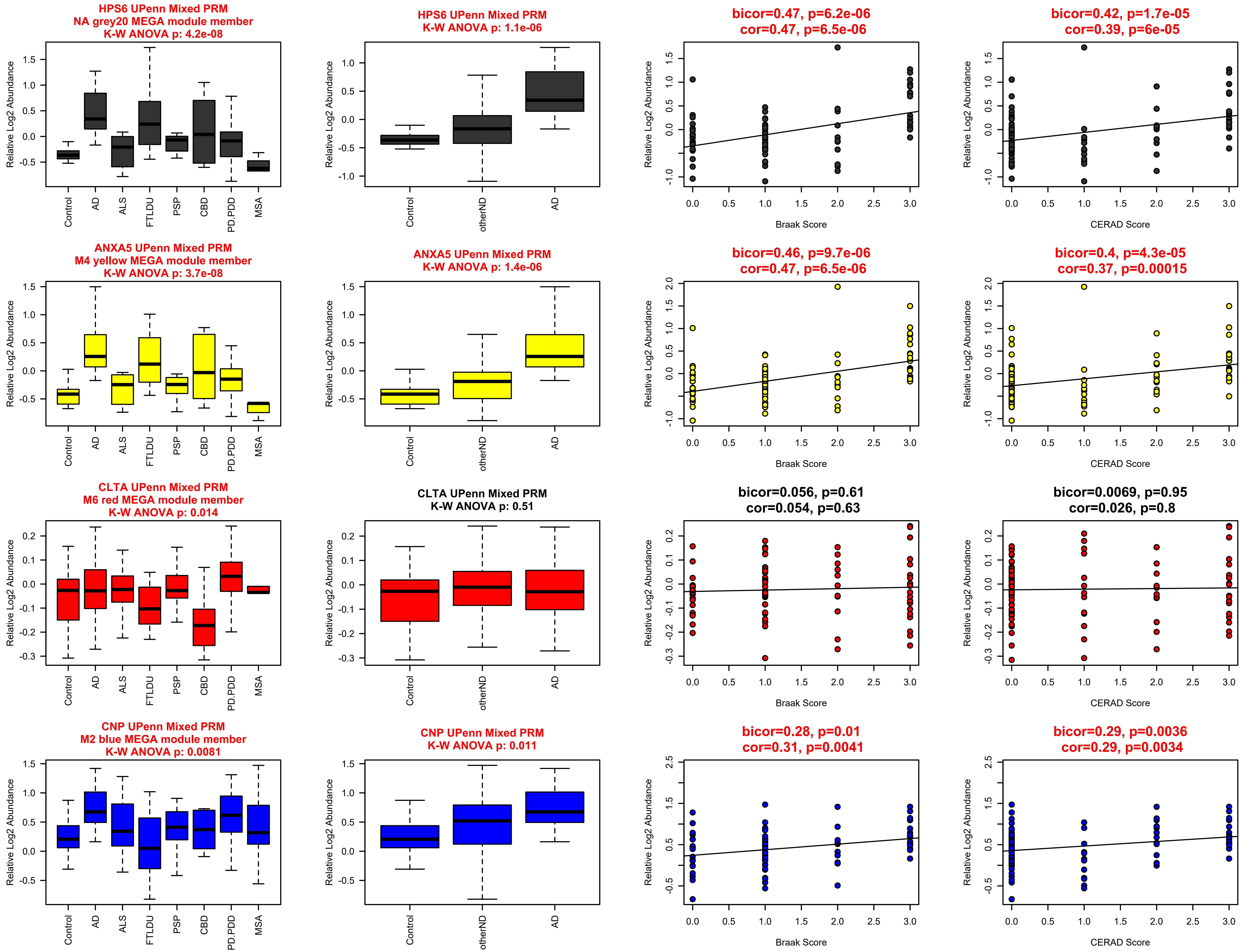
HPS6 UPenn Mixed PRM
NA grey20 MEGA module member
K-W ANOVA p: 4.2e-08
HPS6 UPenn Mixed PRM
K-W ANOVA p: 1.1e-06
bicor=0.47, p=6.2e-06
cor=0.47, p=6.5e-06
bicor=0.42, p=1.7e-05
cor=0.39, p=6e-05
ANXA5 UPenn Mixed PRM
M4 yellow MEGA module member
K-W ANOVA p: 3.7e-08
ANXA5 UPenn Mixed PRM
K-W ANOVA p: 1.4e-06
bicor=0.46, p=9.7e-06
cor=0.47, p=6.5e-06
bicor=0.4, p=4.3e-05
cor=0.37, p=0.00015
CLTA UPenn Mixed PRM
M6 red MEGA module member
K-W ANOVA p: 0.014
CLTA UPenn Mixed PRM
K-W ANOVA p: 0.51
bicor=0.056, p=0.61
cor=0.054, p=0.63
bicor=0.0069, p=0.95
cor=0.026, p=0.8
CNP UPenn Mixed PRM
M2 blue MEGA module member
K-W ANOVA p: 0.0081
CNP UPenn Mixed PRM
K-W ANOVA p: 0.011
bicor=0.28, p=0.01
cor=0.31, p=0.0041
bicor=0.29, p=0.0036
cor=0.29, p=0.0034
Relative Log2 Abundance
Control
AD
ALS
FTLDU
PSP
CBD
PD.PDD
MSA
otherND
Braak Score
CERAD Score

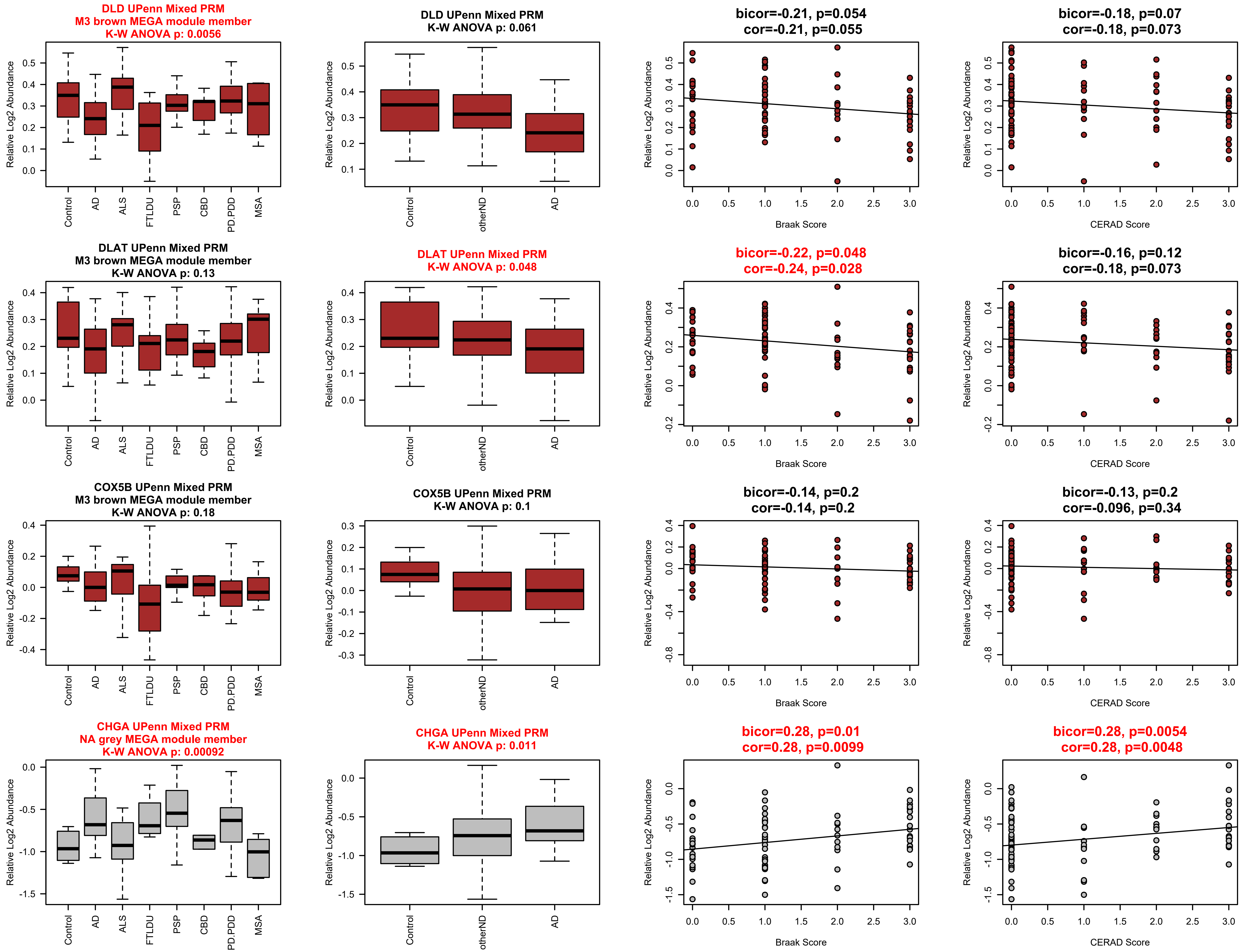
DLD UPenn Mixed PRM
M3 brown MEGA module member
K-W ANOVA p: 0.0056
Relative Log2 Abundance
Control
AD
ALS
FTLDU
PSP
CBD
PD.PDD
MSA
DLD UPenn Mixed PRM
K-W ANOVA p: 0.061
Control
otherND
AD
bicor=-0.21, p=0.054
cor=-0.21, p=0.055
Braak Score
bicor=-0.18, p=0.07
cor=-0.18, p=0.073
CERAD Score
DLAT UPenn Mixed PRM
M3 brown MEGA module member
K-W ANOVA p: 0.13
DLAT UPenn Mixed PRM
K-W ANOVA p: 0.048
bicor=-0.22, p=0.048
cor=-0.24, p=0.028
bicor=-0.16, p=0.12
cor=-0.18, p=0.073
COX5B UPenn Mixed PRM
M3 brown MEGA module member
K-W ANOVA p: 0.18
COX5B UPenn Mixed PRM
K-W ANOVA p: 0.1
bicor=-0.14, p=0.2
cor=-0.14, p=0.2
bicor=-0.13, p=0.2
cor=-0.096, p=0.34
CHGA UPenn Mixed PRM
NA grey MEGA module member
K-W ANOVA p: 0.00092
CHGA UPenn Mixed PRM
K-W ANOVA p: 0.011
bicor=0.28, p=0.01
cor=0.28, p=0.0099
bicor=0.28, p=0.0054
cor=0.28, p=0.0048

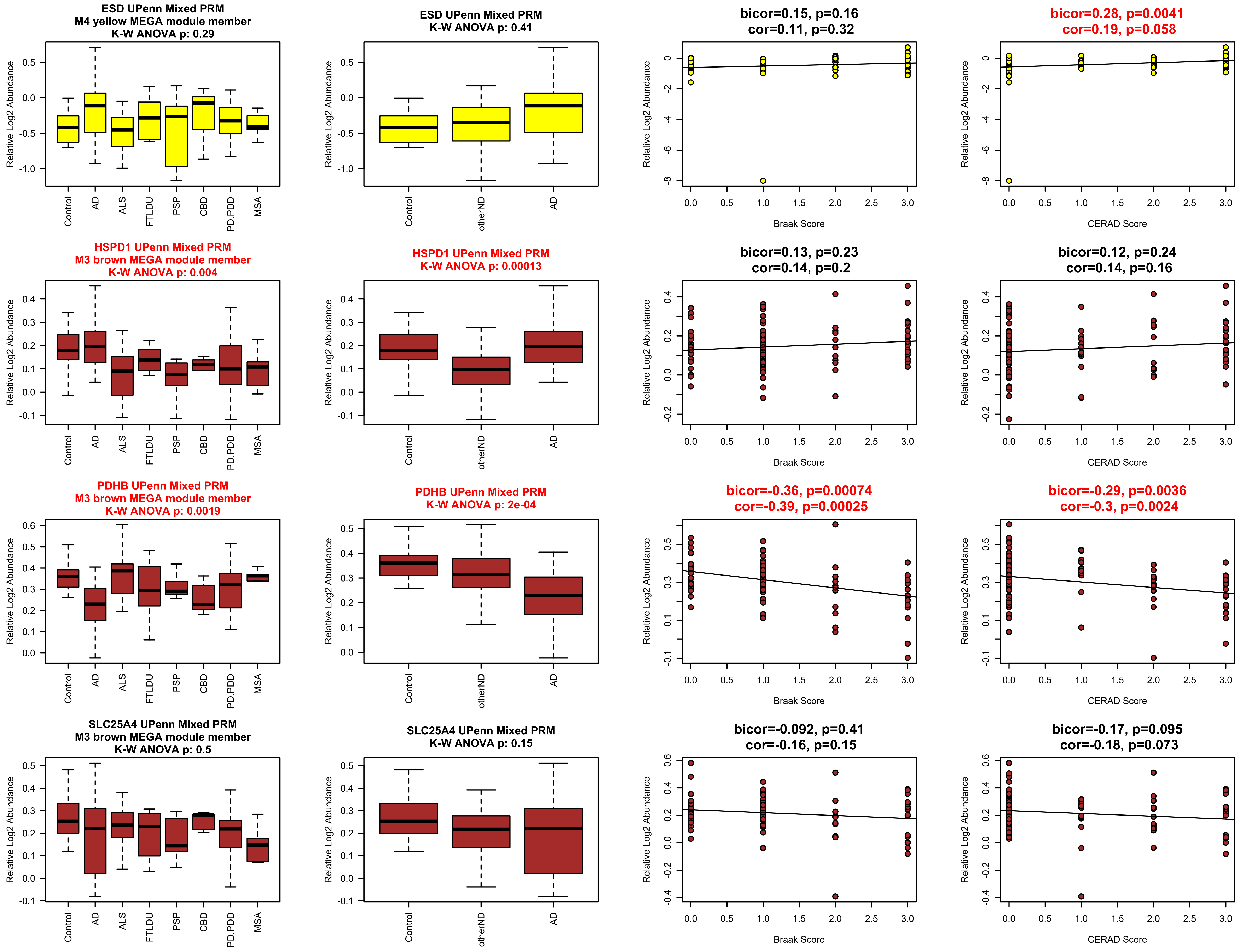
ESD UPenn Mixed PRM
M4 yellow MEGA module member
K-W ANOVA p: 0.29
Relative Log2 Abundance
Control
AD
ALS
FTLDU
PSP
CBD
PD.PDD
MSA
ESD UPenn Mixed PRM
K-W ANOVA p: 0.41
otherND
bicor=0.15, p=0.16
cor=0.11, p=0.32
Braak Score
bicor=0.28, p=0.0041
cor=0.19, p=0.058
CERAD Score
HSPD1 UPenn Mixed PRM
M3 brown MEGA module member
K-W ANOVA p: 0.004
HSPD1 UPenn Mixed PRM
K-W ANOVA p: 0.00013
bicor=0.13, p=0.23
cor=0.14, p=0.2
bicor=0.12, p=0.24
cor=0.14, p=0.16
PDHB UPenn Mixed PRM
M3 brown MEGA module member
K-W ANOVA p: 0.0019
PDHB UPenn Mixed PRM
K-W ANOVA p: 2e-04
bicor=-0.36, p=0.00074
cor=-0.39, p=0.00025
bicor=-0.29, p=0.0036
cor=-0.3, p=0.0024
SLC25A4 UPenn Mixed PRM
M3 brown MEGA module member
K-W ANOVA p: 0.5
SLC25A4 UPenn Mixed PRM
K-W ANOVA p: 0.15
bicor=-0.092, p=0.41
cor=-0.16, p=0.15
bicor=-0.17, p=0.095
cor=-0.18, p=0.073

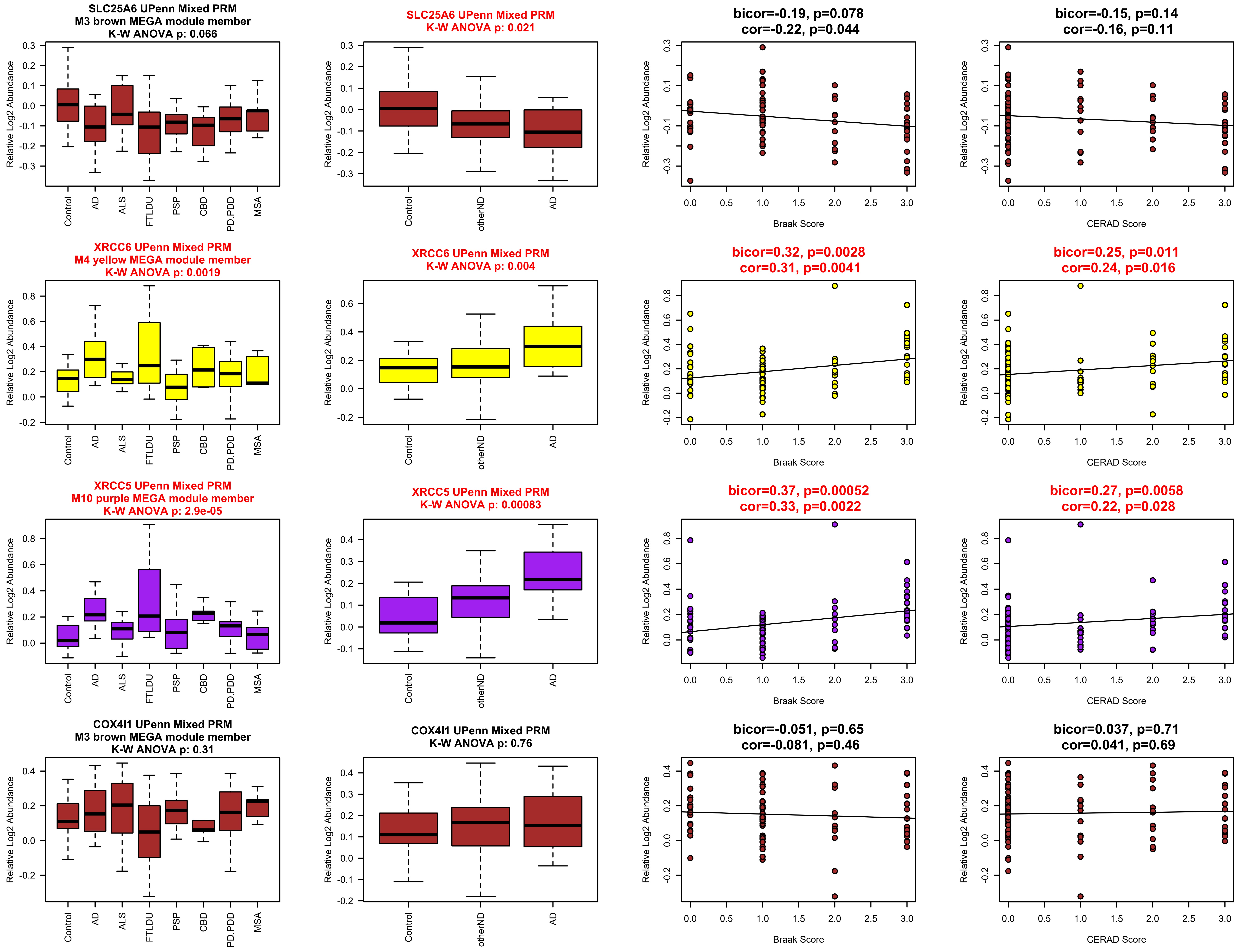
SLC25A6 UPenn Mixed PRM
M3 brown MEGA module member
K-W ANOVA p: 0.066
SLC25A6 UPenn Mixed PRM
K-W ANOVA p: 0.021
bicor=-0.19, p=0.078
cor=-0.22, p=0.044
bicor=-0.15, p=0.14
cor=-0.16, p=0.11
XRCC6 UPenn Mixed PRM
M4 yellow MEGA module member
K-W ANOVA p: 0.0019
XRCC6 UPenn Mixed PRM
K-W ANOVA p: 0.004
bicor=0.32, p=0.0028
cor=0.31, p=0.0041
bicor=0.25, p=0.011
cor=0.24, p=0.016
XRCC5 UPenn Mixed PRM
M10 purple MEGA module member
K-W ANOVA p: 2.9e-05
XRCC5 UPenn Mixed PRM
K-W ANOVA p: 0.00083
bicor=0.37, p=0.00052
cor=0.33, p=0.0022
bicor=0.27, p=0.0058
cor=0.22, p=0.028
COX4I1 UPenn Mixed PRM
M3 brown MEGA module member
K-W ANOVA p: 0.31
COX4I1 UPenn Mixed PRM
K-W ANOVA p: 0.76
bicor=-0.051, p=0.65
cor=-0.081, p=0.46
bicor=0.037, p=0.71
cor=0.041, p=0.69
Relative Log2 Abundance
Control
AD
ALS
FTLDU
PSP
CBD
PD.PDD
MSA
otherND
Braak Score
CERAD Score

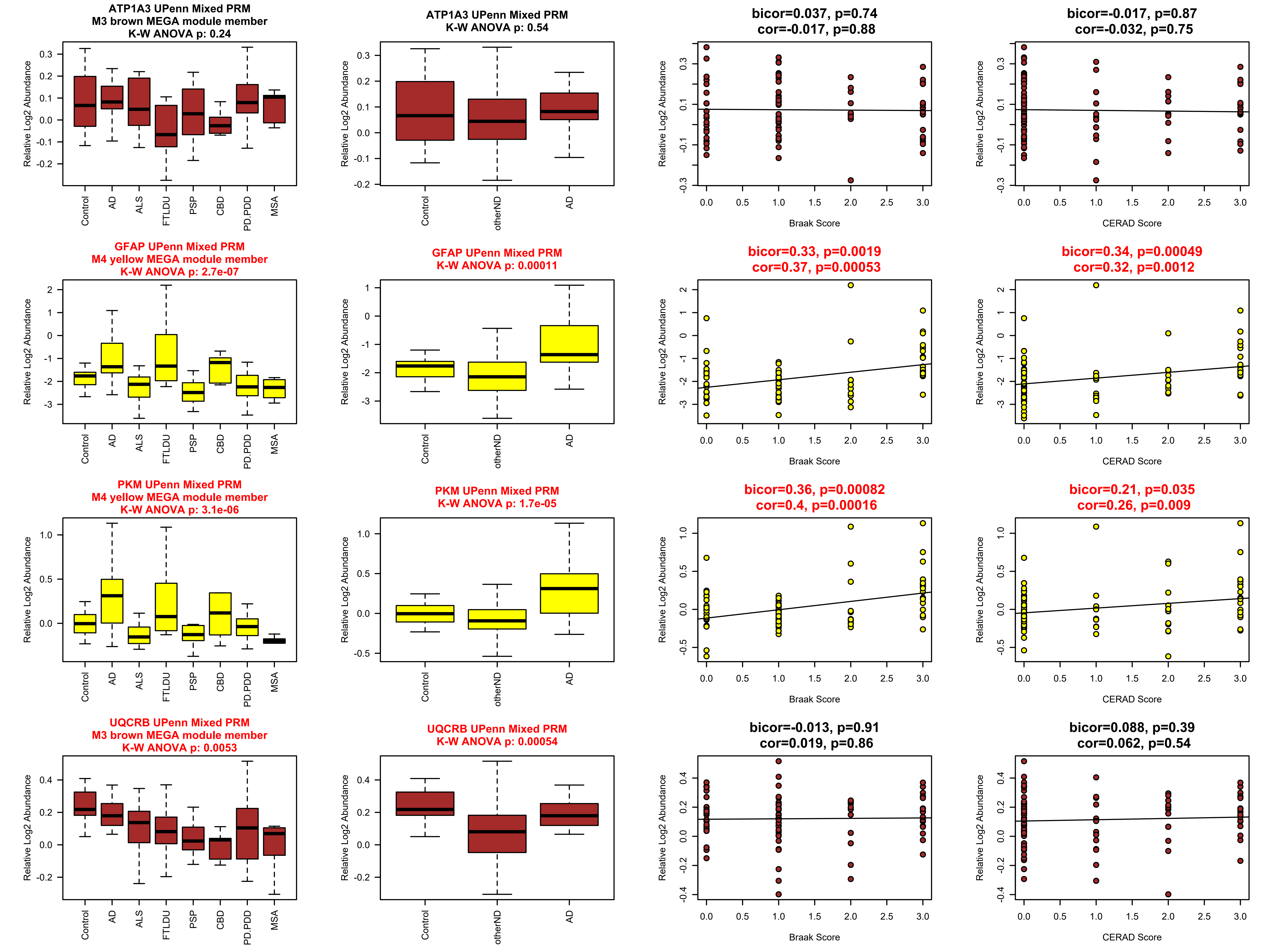
ATP1A3 UPenn Mixed PRM
M3 brown MEGA module member
K-W ANOVA p: 0.24
ATP1A3 UPenn Mixed PRM
K-W ANOVA p: 0.54
bicor=0.037, p=0.74
cor=-0.017, p=0.88
bicor=-0.017, p=0.87
cor=-0.032, p=0.75
GFAP UPenn Mixed PRM
M4 yellow MEGA module member
K-W ANOVA p: 2.7e-07
GFAP UPenn Mixed PRM
K-W ANOVA p: 0.00011
bicor=0.33, p=0.0019
cor=0.37, p=0.00053
bicor=0.34, p=0.00049
cor=0.32, p=0.0012
PKM UPenn Mixed PRM
M4 yellow MEGA module member
K-W ANOVA p: 3.1e-06
PKM UPenn Mixed PRM
K-W ANOVA p: 1.7e-05
bicor=0.36, p=0.00082
cor=0.4, p=0.00016
bicor=0.21, p=0.035
cor=0.26, p=0.009
UQCRB UPenn Mixed PRM
M3 brown MEGA module member
K-W ANOVA p: 0.0053
UQCRB UPenn Mixed PRM
K-W ANOVA p: 0.00054
bicor=-0.013, p=0.91
cor=0.019, p=0.86
bicor=0.088, p=0.39
cor=0.062, p=0.54
Relative Log2 Abundance
Control
AD
ALS
FTLDU
PSP
CBD
PD.PDD
MSA
otherND
Braak Score
CERAD Score

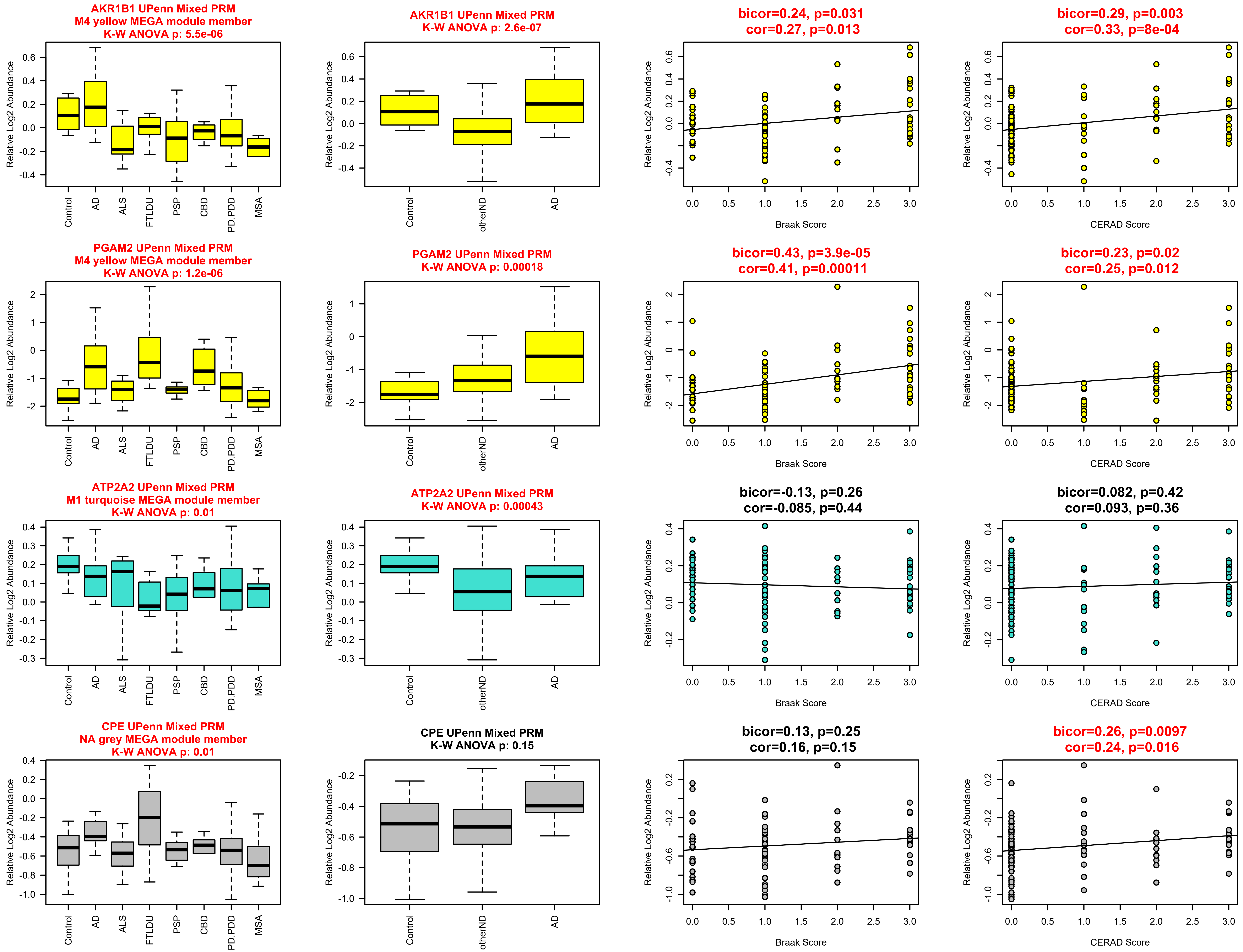
AKR1B1 UPenn Mixed PRM
M4 yellow MEGA module member
K-W ANOVA p: 5.5e-06
AKR1B1 UPenn Mixed PRM
K-W ANOVA p: 2.6e-07
bicor=0.24, p=0.031
cor=0.27, p=0.013
bicor=0.29, p=0.003
cor=0.33, p=8e-04
PGAM2 UPenn Mixed PRM
M4 yellow MEGA module member
K-W ANOVA p: 1.2e-06
PGAM2 UPenn Mixed PRM
K-W ANOVA p: 0.00018
bicor=0.43, p=3.9e-05
cor=0.41, p=0.00011
bicor=0.23, p=0.02
cor=0.25, p=0.012
ATP2A2 UPenn Mixed PRM
M1 turquoise MEGA module member
K-W ANOVA p: 0.01
ATP2A2 UPenn Mixed PRM
K-W ANOVA p: 0.00043
bicor=-0.13, p=0.26
cor=-0.085, p=0.44
bicor=0.082, p=0.42
cor=0.093, p=0.36
CPE UPenn Mixed PRM
NA grey MEGA module member
K-W ANOVA p: 0.01
CPE UPenn Mixed PRM
K-W ANOVA p: 0.15
bicor=0.13, p=0.25
cor=0.16, p=0.15
bicor=0.26, p=0.0097
cor=0.24, p=0.016
Relative Log2 Abundance
Control
AD
ALS
FTLDU
PSP
CBD
PD.PDD
MSA
otherND
Braak Score
CERAD Score

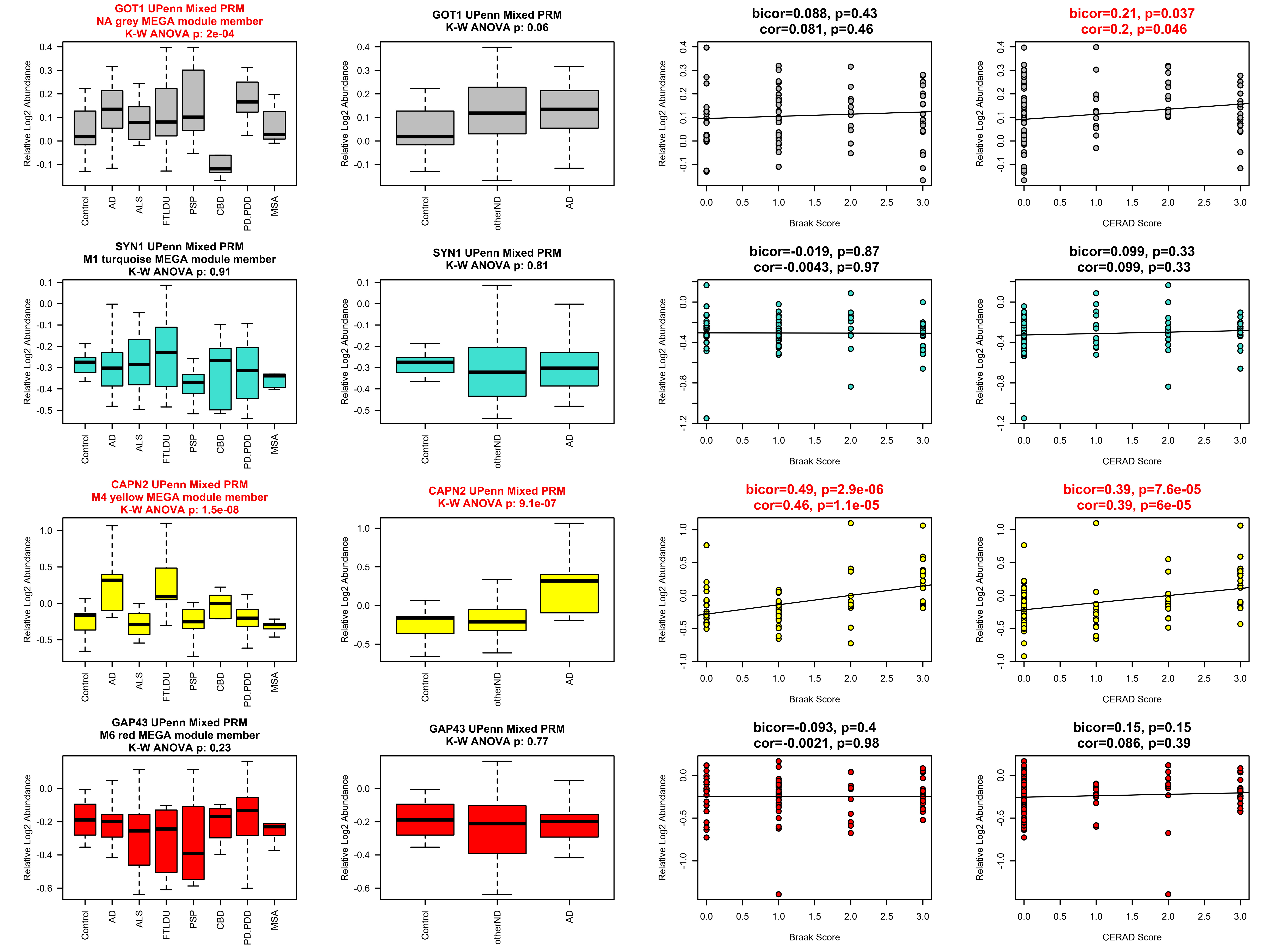
GOT1 UPenn Mixed PRM
NA grey MEGA module member
K-W ANOVA p: 2e-04
GOT1 UPenn Mixed PRM
K-W ANOVA p: 0.06
bicor=0.088, p=0.43
cor=0.081, p=0.46
bicor=0.21, p=0.037
cor=0.2, p=0.046
SYN1 UPenn Mixed PRM
M1 turquoise MEGA module member
K-W ANOVA p: 0.91
SYN1 UPenn Mixed PRM
K-W ANOVA p: 0.81
bicor=-0.019, p=0.87
cor=-0.0043, p=0.97
bicor=0.099, p=0.33
cor=0.099, p=0.33
CAPN2 UPenn Mixed PRM
M4 yellow MEGA module member
K-W ANOVA p: 1.5e-08
CAPN2 UPenn Mixed PRM
K-W ANOVA p: 9.1e-07
bicor=0.49, p=2.9e-06
cor=0.46, p=1.1e-05
bicor=0.39, p=7.6e-05
cor=0.39, p=6e-05
GAP43 UPenn Mixed PRM
M6 red MEGA module member
K-W ANOVA p: 0.23
GAP43 UPenn Mixed PRM
K-W ANOVA p: 0.77
bicor=-0.093, p=0.4
cor=-0.0021, p=0.98
bicor=0.15, p=0.15
cor=0.086, p=0.39
Relative Log2 Abundance
Control
AD
ALS
FTLDU
PSP
CBD
PD.PDD
MSA
otherND
Braak Score
CERAD Score

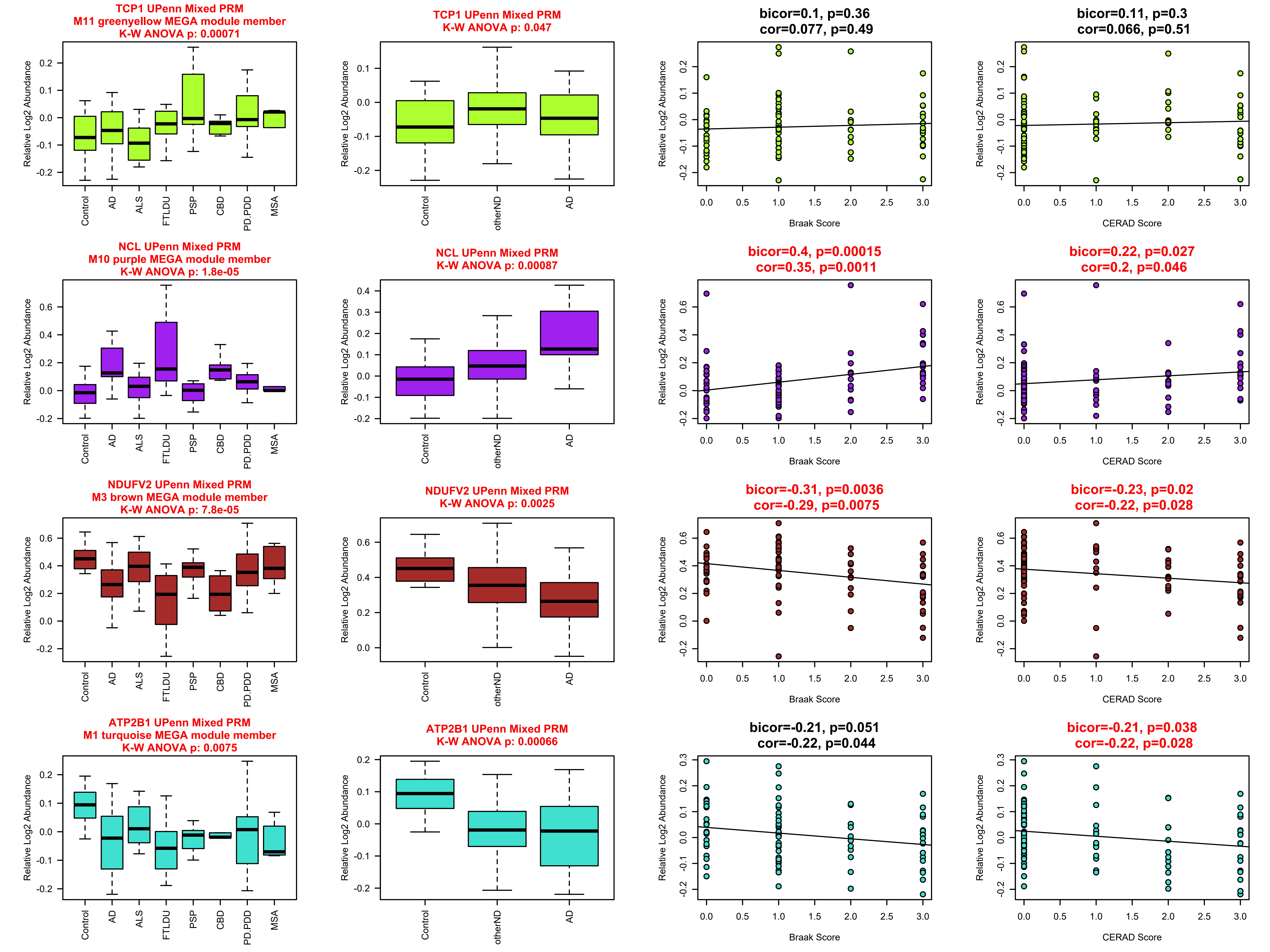

TCP1 UPenn Mixed PRM
M11 greenyellow MEGA module member
K-W ANOVA p: 0.00071
TCP1 UPenn Mixed PRM
K-W ANOVA p: 0.047
bicor=0.1, p=0.36
cor=0.077, p=0.49
bicor=0.11, p=0.3
cor=0.066, p=0.51
NCL UPenn Mixed PRM
M10 purple MEGA module member
K-W ANOVA p: 1.8e-05
NCL UPenn Mixed PRM
K-W ANOVA p: 0.00087
bicor=0.4, p=0.00015
cor=0.35, p=0.0011
bicor=0.22, p=0.027
cor=0.2, p=0.046
NDUFV2 UPenn Mixed PRM
M3 brown MEGA module member
K-W ANOVA p: 7.8e-05
NDUFV2 UPenn Mixed PRM
K-W ANOVA p: 0.0025
bicor=-0.31, p=0.0036
cor=-0.29, p=0.0075
bicor=-0.23, p=0.02
cor=-0.22, p=0.028
ATP2B1 UPenn Mixed PRM
M1 turquoise MEGA module member
K-W ANOVA p: 0.0075
ATP2B1 UPenn Mixed PRM
K-W ANOVA p: 0.00066
bicor=-0.21, p=0.051
cor=-0.22, p=0.044
bicor=-0.21, p=0.038
cor=-0.22, p=0.028
Relative Log2 Abundance
Control
AD
ALS
FTLDU
PSP
CBD
PD.PDD
MSA
otherND
Braak Score
CERAD Score

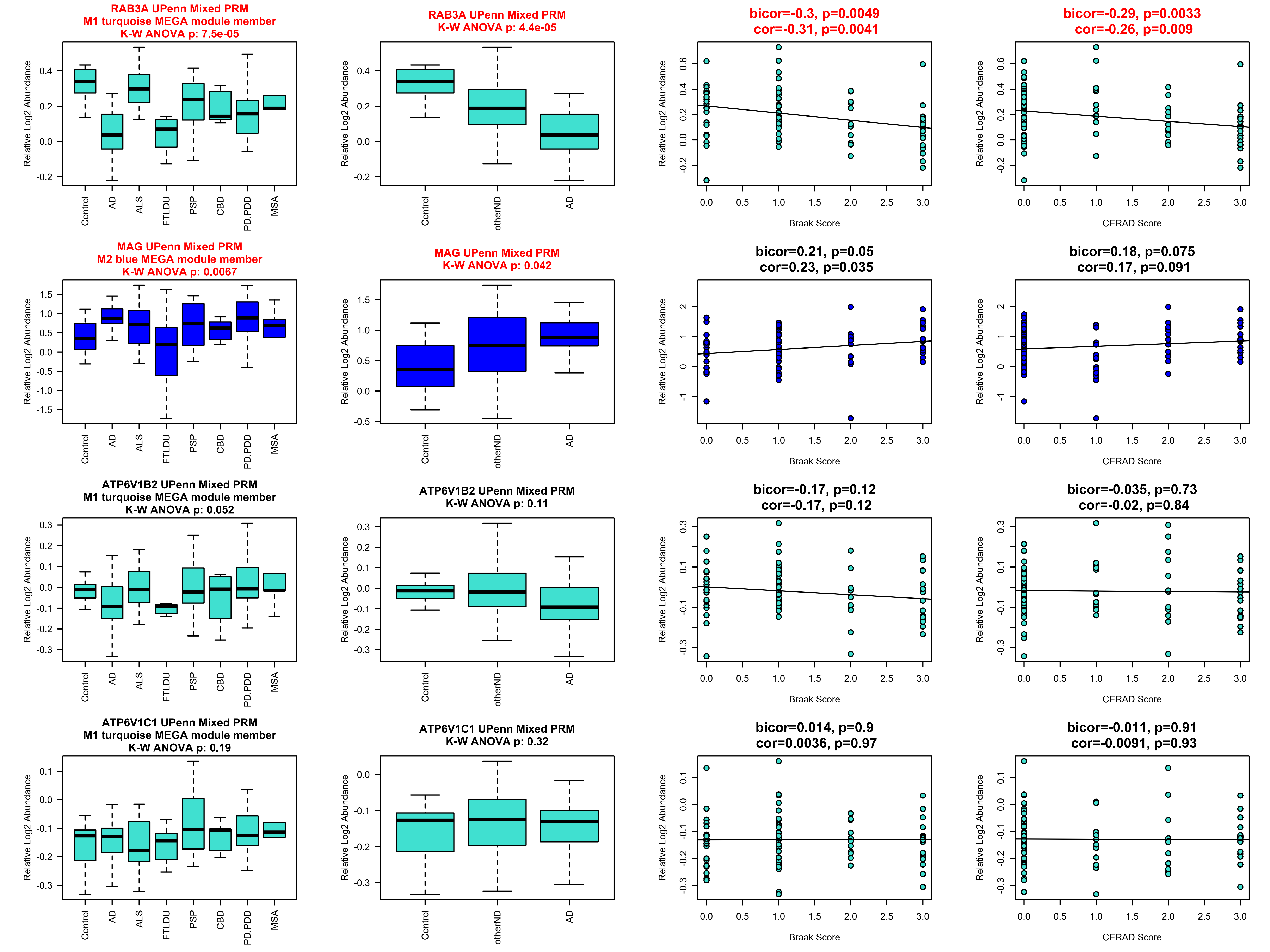
RAB3A UPenn Mixed PRM
M1 turquoise MEGA module member
K-W ANOVA p: 7.5e-05
RAB3A UPenn Mixed PRM
K-W ANOVA p: 4.4e-05
bicor=-0.3, p=0.0049
cor=-0.31, p=0.0041
bicor=-0.29, p=0.0033
cor=-0.26, p=0.009
MAG UPenn Mixed PRM
M2 blue MEGA module member
K-W ANOVA p: 0.0067
MAG UPenn Mixed PRM
K-W ANOVA p: 0.042
bicor=0.21, p=0.05
cor=0.23, p=0.035
bicor=0.18, p=0.075
cor=0.17, p=0.091
ATP6V1B2 UPenn Mixed PRM
M1 turquoise MEGA module member
K-W ANOVA p: 0.052
ATP6V1B2 UPenn Mixed PRM
K-W ANOVA p: 0.11
bicor=-0.17, p=0.12
cor=-0.17, p=0.12
bicor=-0.035, p=0.73
cor=-0.02, p=0.84
ATP6V1C1 UPenn Mixed PRM
M1 turquoise MEGA module member
K-W ANOVA p: 0.19
ATP6V1C1 UPenn Mixed PRM
K-W ANOVA p: 0.32
bicor=0.014, p=0.9
cor=0.0036, p=0.97
bicor=-0.011, p=0.91
cor=-0.0091, p=0.93
Relative Log2 Abundance
Control
AD
ALS
FTLDU
PSP
CBD
PD.PDD
MSA
otherND
Braak Score
CERAD Score

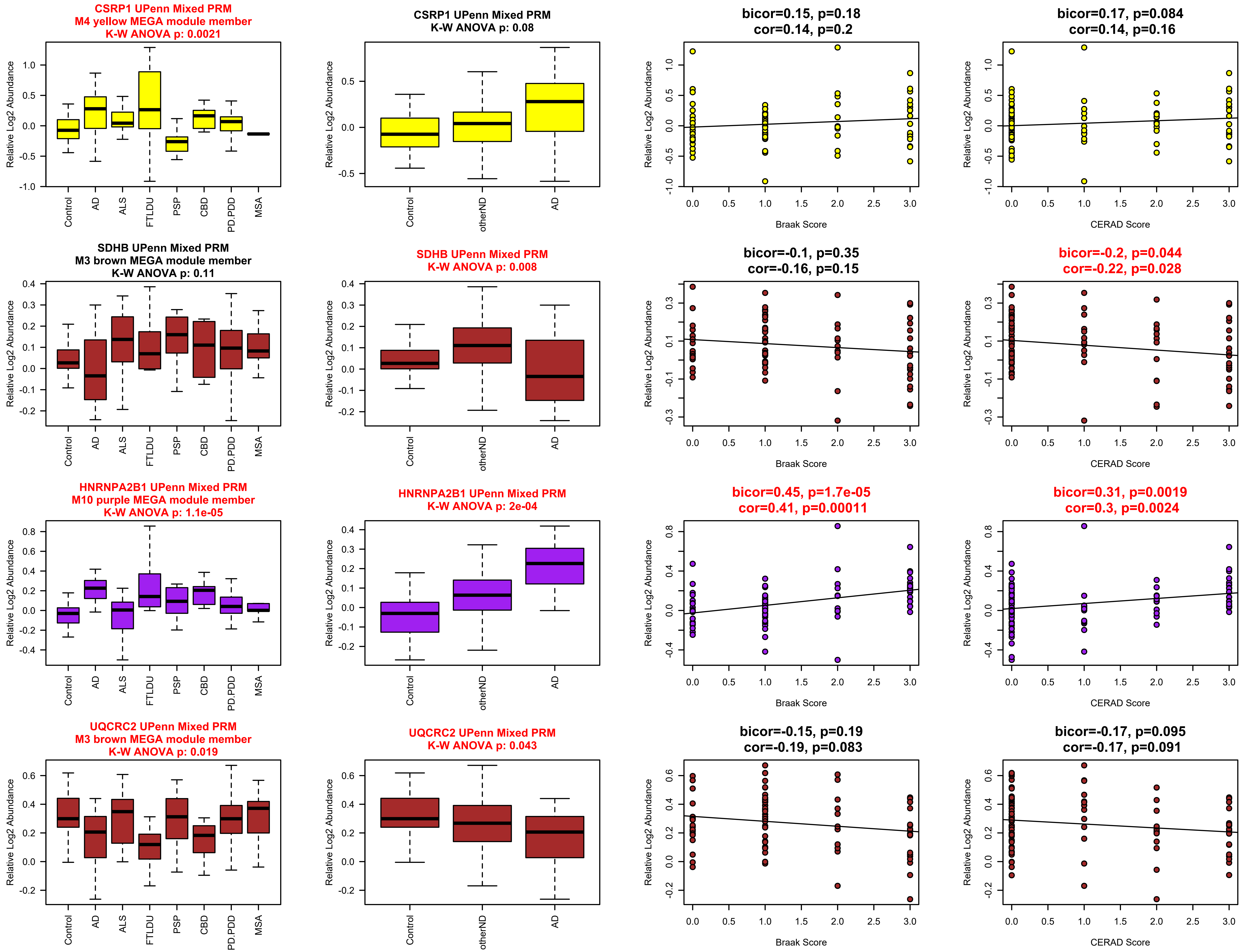
CSRP1 UPenn Mixed PRM
M4 yellow MEGA module member
K-W ANOVA p: 0.0021
CSRP1 UPenn Mixed PRM
K-W ANOVA p: 0.08
bicor=0.15, p=0.18
cor=0.14, p=0.2
bicor=0.17, p=0.084
cor=0.14, p=0.16
SDHB UPenn Mixed PRM
M3 brown MEGA module member
K-W ANOVA p: 0.11
SDHB UPenn Mixed PRM
K-W ANOVA p: 0.008
bicor=-0.1, p=0.35
cor=-0.16, p=0.15
bicor=-0.2, p=0.044
cor=-0.22, p=0.028
HNRNPA2B1 UPenn Mixed PRM
M10 purple MEGA module member
K-W ANOVA p: 1.1e-05
HNRNPA2B1 UPenn Mixed PRM
K-W ANOVA p: 2e-04
bicor=0.45, p=1.7e-05
cor=0.41, p=0.00011
bicor=0.31, p=0.0019
cor=0.3, p=0.0024
UQCRC2 UPenn Mixed PRM
M3 brown MEGA module member
K-W ANOVA p: 0.019
UQCRC2 UPenn Mixed PRM
K-W ANOVA p: 0.043
bicor=-0.15, p=0.19
cor=-0.19, p=0.083
bicor=-0.17, p=0.095
cor=-0.17, p=0.091
Relative Log2 Abundance
Control
AD
ALS
FTLDU
PSP
CBD
PD.PDD
MSA
otherND
Braak Score
CERAD Score

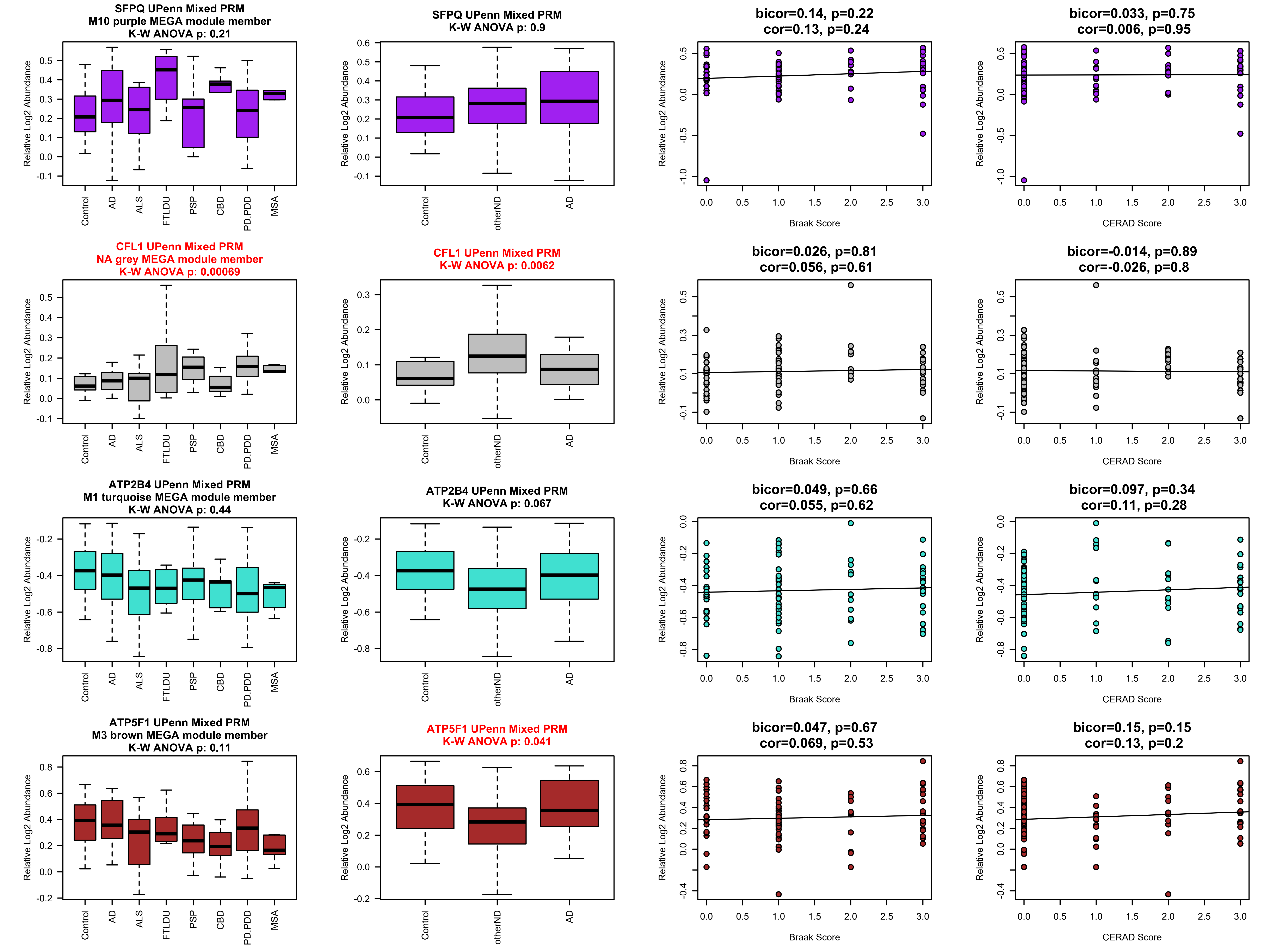
SFPQ UPenn Mixed PRM
M10 purple MEGA module member
K-W ANOVA p: 0.21
SFPQ UPenn Mixed PRM
K-W ANOVA p: 0.9
bicor=0.14, p=0.22
cor=0.13, p=0.24
bicor=0.033, p=0.75
cor=0.006, p=0.95
CFL1 UPenn Mixed PRM
NA grey MEGA module member
K-W ANOVA p: 0.00069
CFL1 UPenn Mixed PRM
K-W ANOVA p: 0.0062
bicor=0.026, p=0.81
cor=0.056, p=0.61
bicor=-0.014, p=0.89
cor=-0.026, p=0.8
ATP2B4 UPenn Mixed PRM
M1 turquoise MEGA module member
K-W ANOVA p: 0.44
ATP2B4 UPenn Mixed PRM
K-W ANOVA p: 0.067
bicor=0.049, p=0.66
cor=0.055, p=0.62
bicor=0.097, p=0.34
cor=0.11, p=0.28
ATP5F1 UPenn Mixed PRM
M3 brown MEGA module member
K-W ANOVA p: 0.11
ATP5F1 UPenn Mixed PRM
K-W ANOVA p: 0.041
bicor=0.047, p=0.67
cor=0.069, p=0.53
bicor=0.15, p=0.15
cor=0.13, p=0.2
Relative Log2 Abundance
Control
AD
ALS
FTLDU
PSP
CBD
PD.PDD
MSA
otherND
Braak Score
CERAD Score

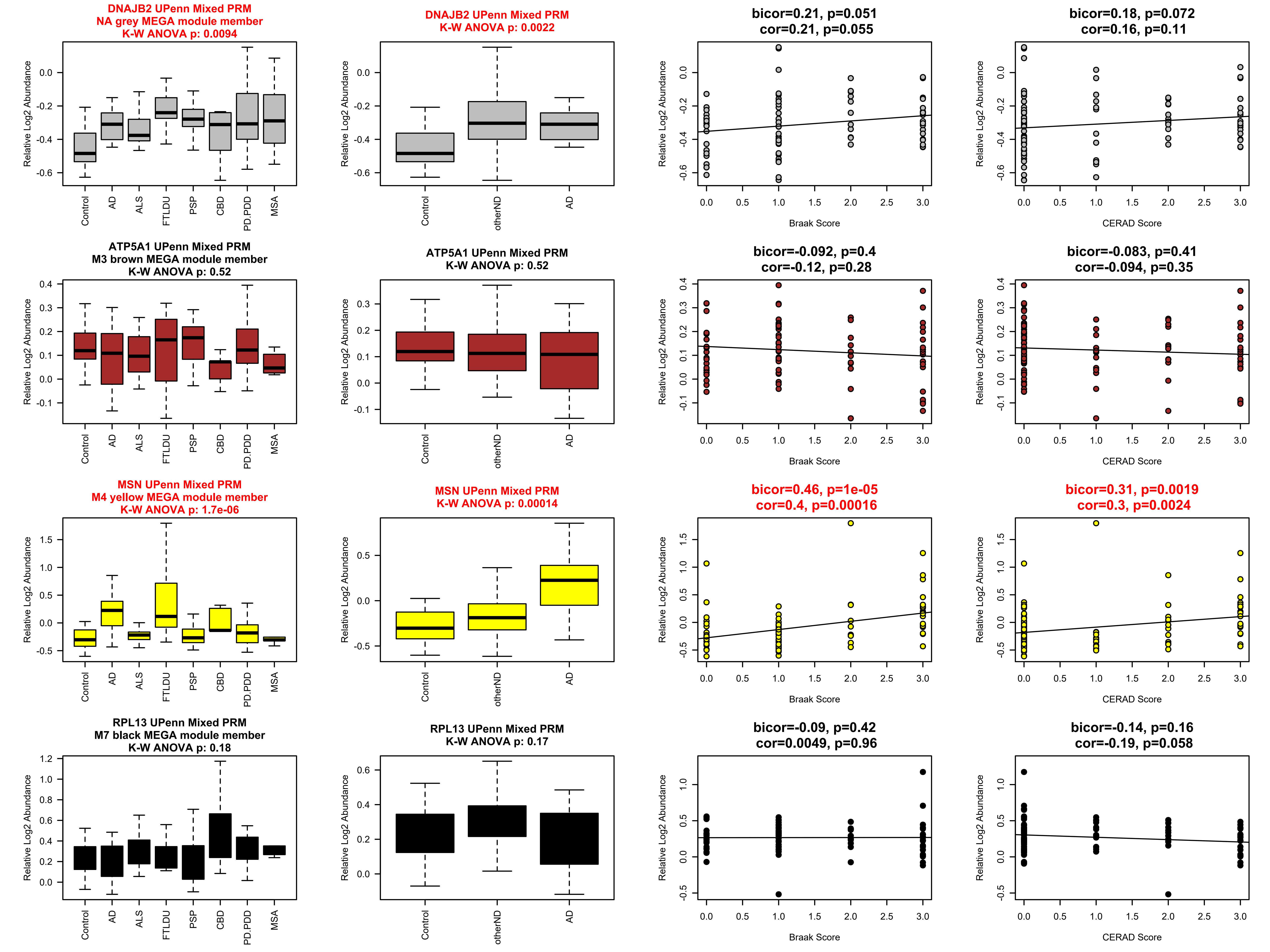
DNAJB2 UPenn Mixed PRM
NA grey MEGA module member
K-W ANOVA p: 0.0094
DNAJB2 UPenn Mixed PRM
K-W ANOVA p: 0.0022
bicor=0.21, p=0.051
cor=0.21, p=0.055
bicor=0.18, p=0.072
cor=0.16, p=0.11
ATP5A1 UPenn Mixed PRM
M3 brown MEGA module member
K-W ANOVA p: 0.52
ATP5A1 UPenn Mixed PRM
K-W ANOVA p: 0.52
bicor=-0.092, p=0.4
cor=-0.12, p=0.28
bicor=-0.083, p=0.41
cor=-0.094, p=0.35
MSN UPenn Mixed PRM
M4 yellow MEGA module member
K-W ANOVA p: 1.7e-06
MSN UPenn Mixed PRM
K-W ANOVA p: 0.00014
bicor=0.46, p=1e-05
cor=0.4, p=0.00016
bicor=0.31, p=0.0019
cor=0.3, p=0.0024
RPL13 UPenn Mixed PRM
M7 black MEGA module member
K-W ANOVA p: 0.18
RPL13 UPenn Mixed PRM
K-W ANOVA p: 0.17
bicor=-0.09, p=0.42
cor=0.0049, p=0.96
bicor=-0.14, p=0.16
cor=-0.19, p=0.058
Relative Log2 Abundance
Control
AD
ALS
FTLDU
PSP
CBD
PD.PDD
MSA
otherND
Braak Score
CERAD Score

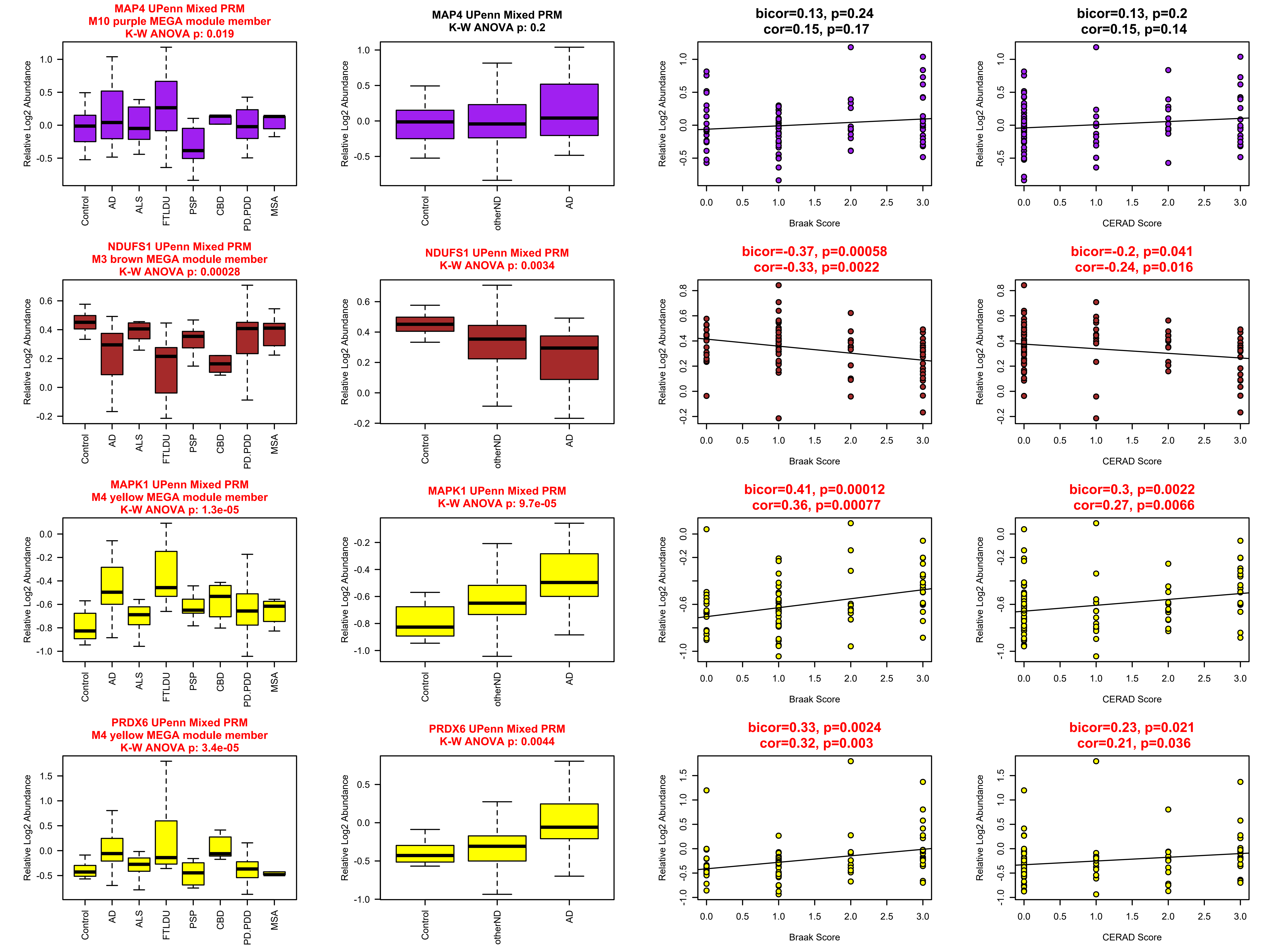
MAP4 UPenn Mixed PRM
M10 purple MEGA module member
K-W ANOVA p: 0.019
MAP4 UPenn Mixed PRM
K-W ANOVA p: 0.2
bicor=0.13, p=0.24
cor=0.15, p=0.17
bicor=0.13, p=0.2
cor=0.15, p=0.14
NDUFS1 UPenn Mixed PRM
M3 brown MEGA module member
K-W ANOVA p: 0.00028
NDUFS1 UPenn Mixed PRM
K-W ANOVA p: 0.0034
bicor=-0.37, p=0.00058
cor=-0.33, p=0.0022
bicor=-0.2, p=0.041
cor=-0.24, p=0.016
MAPK1 UPenn Mixed PRM
M4 yellow MEGA module member
K-W ANOVA p: 1.3e-05
MAPK1 UPenn Mixed PRM
K-W ANOVA p: 9.7e-05
bicor=0.41, p=0.00012
cor=0.36, p=0.00077
bicor=0.3, p=0.0022
cor=0.27, p=0.0066
PRDX6 UPenn Mixed PRM
M4 yellow MEGA module member
K-W ANOVA p: 3.4e-05
PRDX6 UPenn Mixed PRM
K-W ANOVA p: 0.0044
bicor=0.33, p=0.0024
cor=0.32, p=0.003
bicor=0.23, p=0.021
cor=0.21, p=0.036
Relative Log2 Abundance
Control
AD
ALS
FTLDU
PSP
CBD
PD.PDD
MSA
otherND
Braak Score
CERAD Score

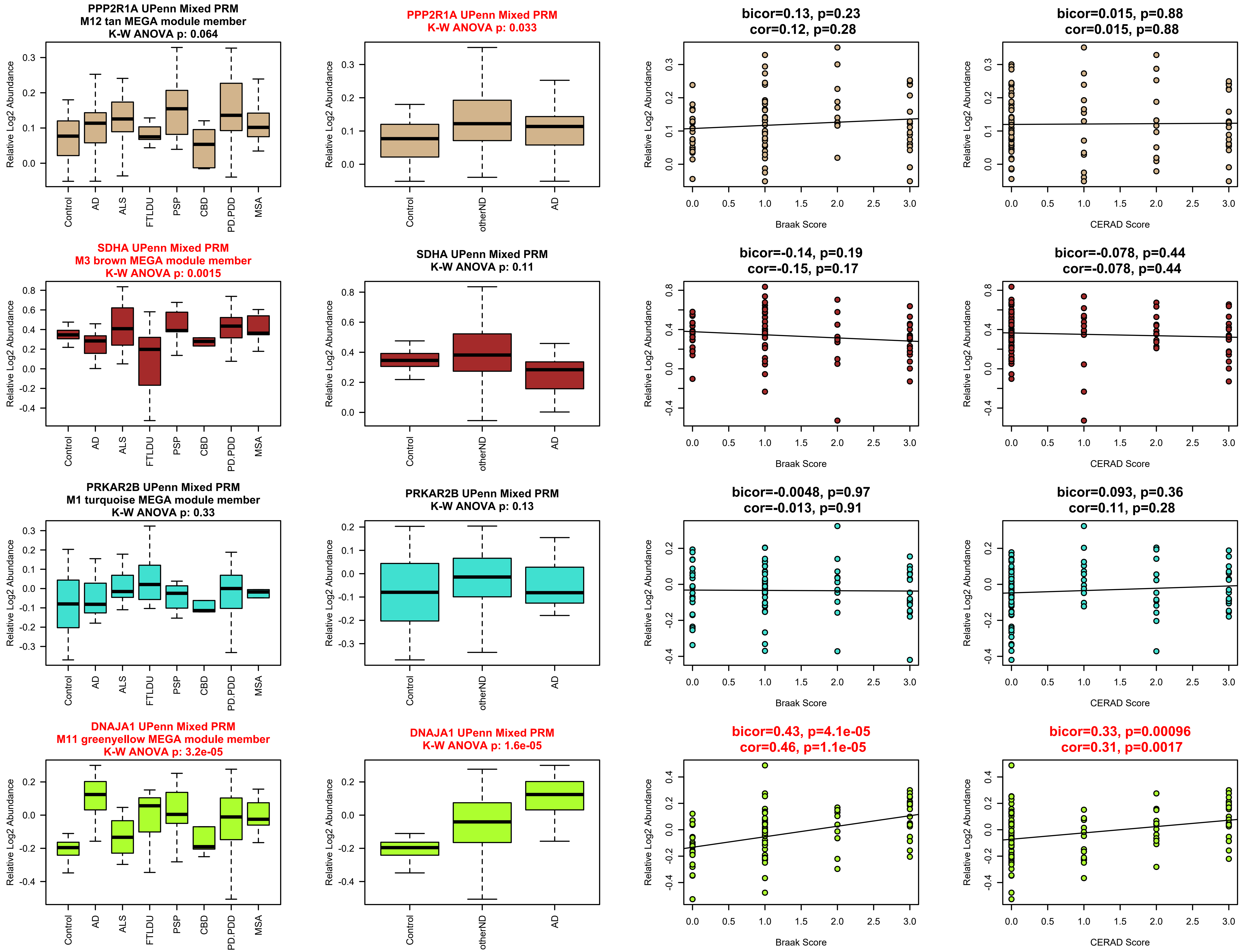
PPP2R1A UPenn Mixed PRM
M12 tan MEGA module member
K-W ANOVA p: 0.064
PPP2R1A UPenn Mixed PRM
K-W ANOVA p: 0.033
bicor=0.13, p=0.23
cor=0.12, p=0.28
bicor=0.015, p=0.88
cor=0.015, p=0.88
SDHA UPenn Mixed PRM
M3 brown MEGA module member
K-W ANOVA p: 0.0015
SDHA UPenn Mixed PRM
K-W ANOVA p: 0.11
bicor=-0.14, p=0.19
cor=-0.15, p=0.17
bicor=-0.078, p=0.44
cor=-0.078, p=0.44
PRKAR2B UPenn Mixed PRM
M1 turquoise MEGA module member
K-W ANOVA p: 0.33
PRKAR2B UPenn Mixed PRM
K-W ANOVA p: 0.13
bicor=-0.0048, p=0.97
cor=-0.013, p=0.91
bicor=0.093, p=0.36
cor=0.11, p=0.28
DNAJA1 UPenn Mixed PRM
M11 greenyellow MEGA module member
K-W ANOVA p: 3.2e-05
DNAJA1 UPenn Mixed PRM
K-W ANOVA p: 1.6e-05
bicor=0.43, p=4.1e-05
cor=0.46, p=1.1e-05
bicor=0.33, p=0.00096
cor=0.31, p=0.0017
Relative Log2 Abundance
Control
AD
ALS
FTLDU
PSP
CBD
PD.PDD
MSA
otherND
Braak Score
CERAD Score

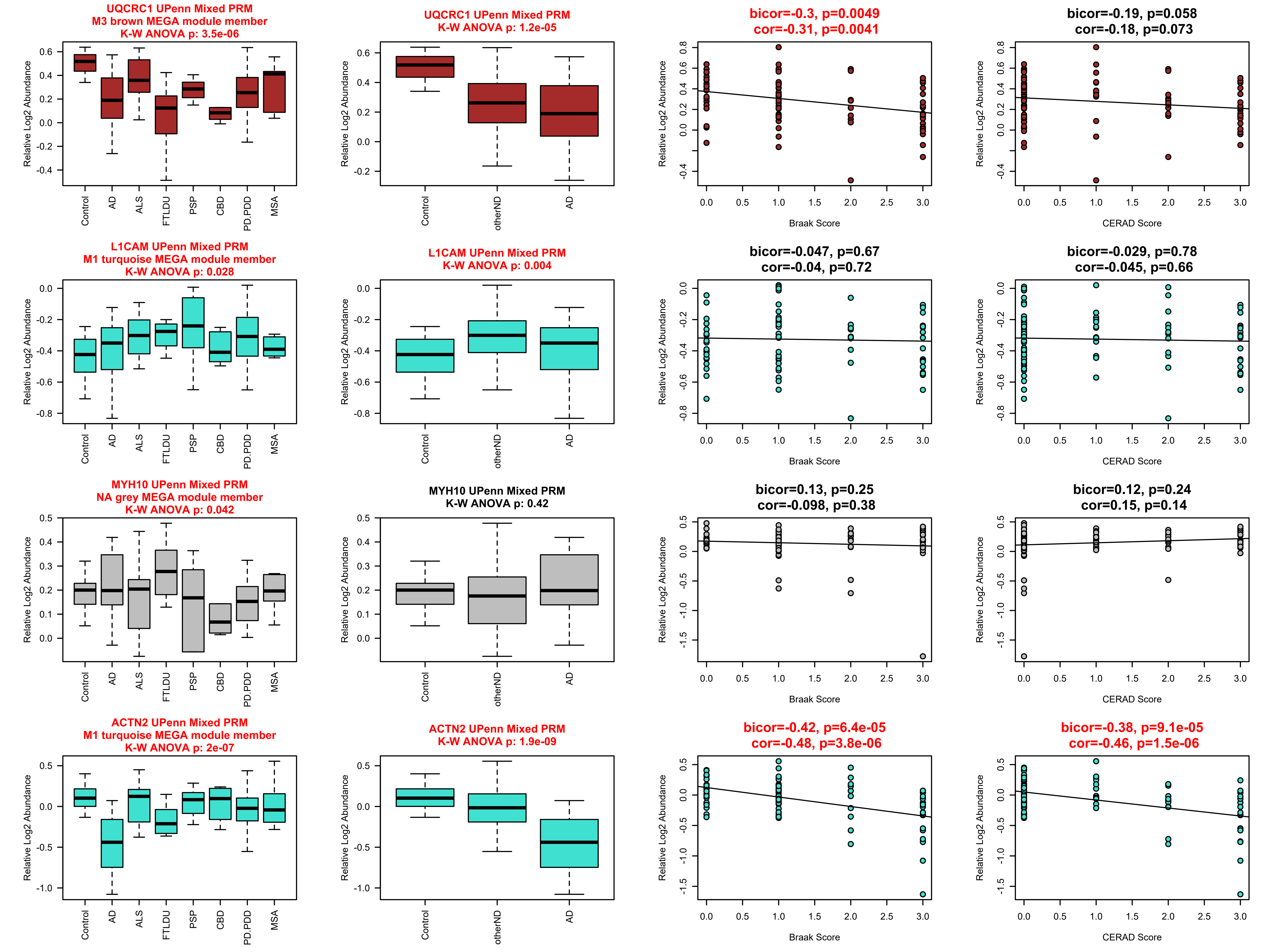

UQCRC1 UPenn Mixed PRM
M3 brown MEGA module member
K-W ANOVA p: 3.5e-06
UQCRC1 UPenn Mixed PRM
K-W ANOVA p: 1.2e-05
bicor=-0.3, p=0.0049
cor=-0.31, p=0.0041
bicor=-0.19, p=0.058
cor=-0.18, p=0.073
L1CAM UPenn Mixed PRM
M1 turquoise MEGA module member
K-W ANOVA p: 0.028
L1CAM UPenn Mixed PRM
K-W ANOVA p: 0.004
bicor=-0.047, p=0.67
cor=-0.04, p=0.72
bicor=-0.029, p=0.78
cor=-0.045, p=0.66
MYH10 UPenn Mixed PRM
NA grey MEGA module member
K-W ANOVA p: 0.042
MYH10 UPenn Mixed PRM
K-W ANOVA p: 0.42
bicor=0.13, p=0.25
cor=-0.098, p=0.38
bicor=0.12, p=0.24
cor=0.15, p=0.14
ACTN2 UPenn Mixed PRM
M1 turquoise MEGA module member
K-W ANOVA p: 2e-07
ACTN2 UPenn Mixed PRM
K-W ANOVA p: 1.9e-09
bicor=-0.42, p=6.4e-05
cor=-0.48, p=3.8e-06
bicor=-0.38, p=9.1e-05
cor=-0.46, p=1.5e-06
Relative Log2 Abundance
Control
AD
ALS
FTLDU
PSP
CBD
PD.PDD
MSA
otherND
Braak Score
CERAD Score

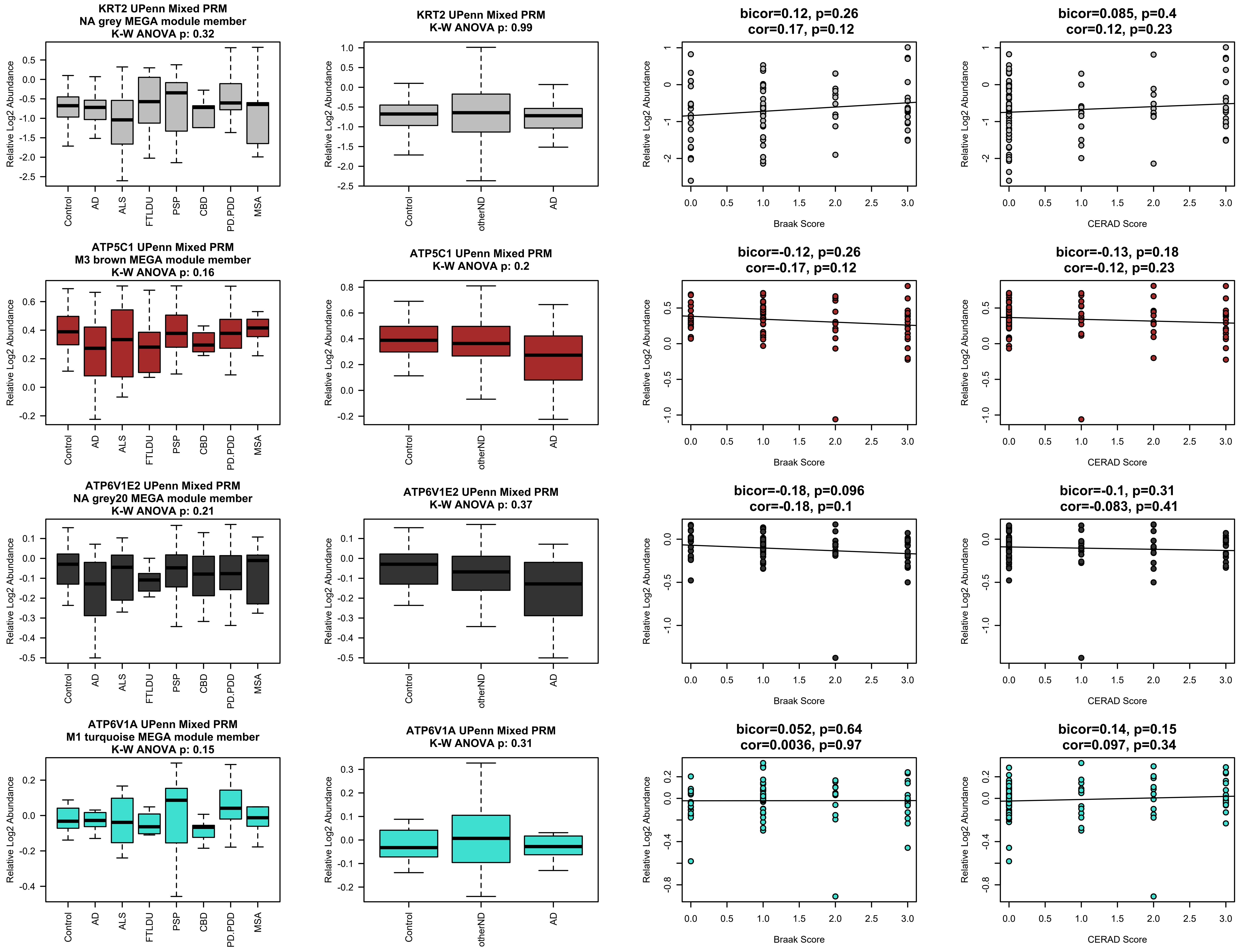
KRT2 UPenn Mixed PRM
NA grey MEGA module member
K-W ANOVA p: 0.32
KRT2 UPenn Mixed PRM
K-W ANOVA p: 0.99
bicor=0.12, p=0.26
cor=0.17, p=0.12
bicor=0.085, p=0.4
cor=0.12, p=0.23
ATP5C1 UPenn Mixed PRM
M3 brown MEGA module member
K-W ANOVA p: 0.16
ATP5C1 UPenn Mixed PRM
K-W ANOVA p: 0.2
bicor=-0.12, p=0.26
cor=-0.17, p=0.12
bicor=-0.13, p=0.18
cor=-0.12, p=0.23
ATP6V1E2 UPenn Mixed PRM
NA grey20 MEGA module member
K-W ANOVA p: 0.21
ATP6V1E2 UPenn Mixed PRM
K-W ANOVA p: 0.37
bicor=-0.18, p=0.096
cor=-0.18, p=0.1
bicor=-0.1, p=0.31
cor=-0.083, p=0.41
ATP6V1A UPenn Mixed PRM
M1 turquoise MEGA module member
K-W ANOVA p: 0.15
ATP6V1A UPenn Mixed PRM
K-W ANOVA p: 0.31
bicor=0.052, p=0.64
cor=0.0036, p=0.97
bicor=0.14, p=0.15
cor=0.097, p=0.34
Relative Log2 Abundance
Control
AD
ALS
FTLDU
PSP
CBD
PD.PDD
MSA
otherND
Braak Score
CERAD Score

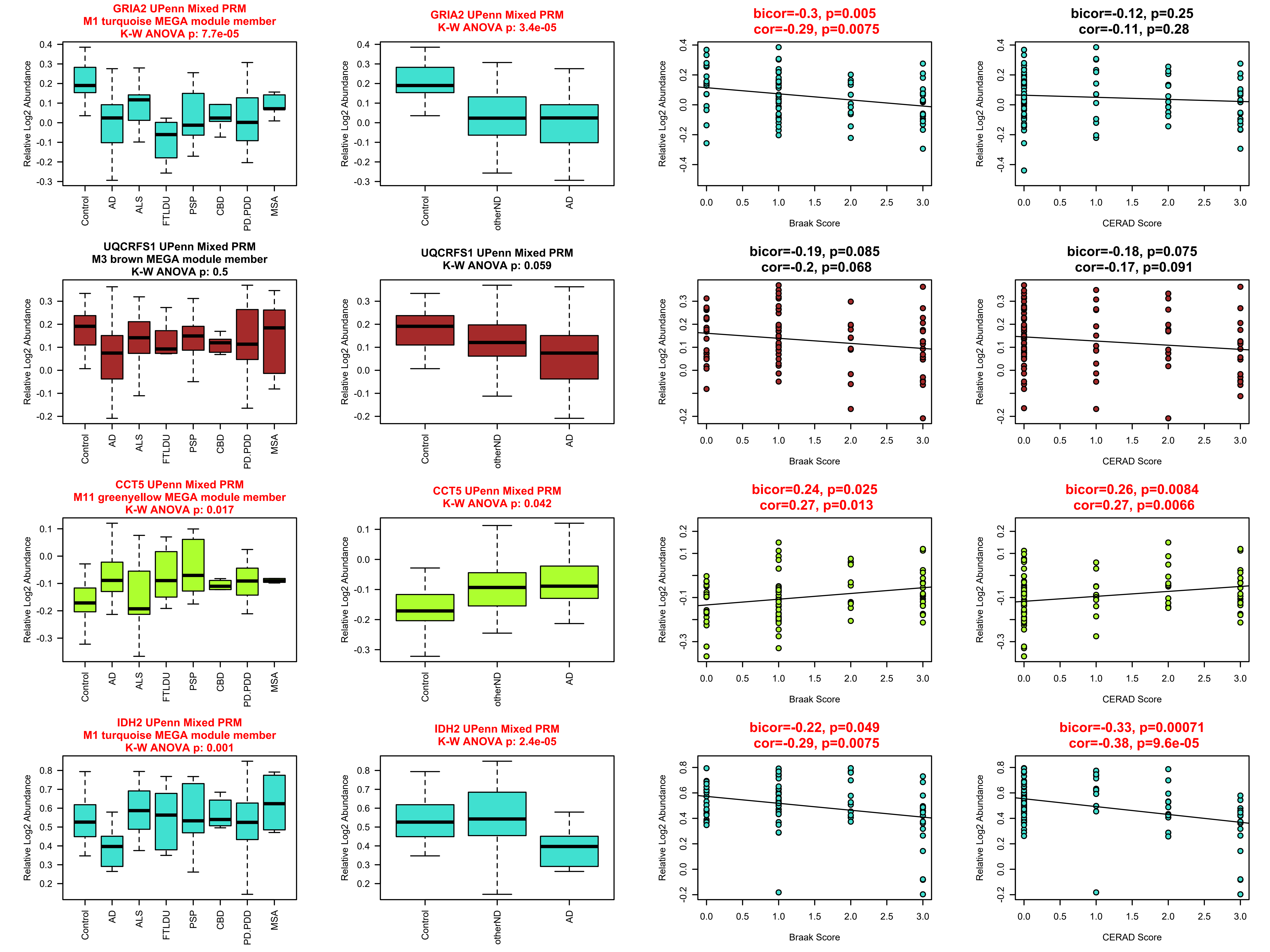
GRIA2 UPenn Mixed PRM
M1 turquoise MEGA module member
K-W ANOVA p: 7.7e-05
GRIA2 UPenn Mixed PRM
K-W ANOVA p: 3.4e-05
bicor=-0.3, p=0.005
cor=-0.29, p=0.0075
bicor=-0.12, p=0.25
cor=-0.11, p=0.28
UQCRFS1 UPenn Mixed PRM
M3 brown MEGA module member
K-W ANOVA p: 0.5
UQCRFS1 UPenn Mixed PRM
K-W ANOVA p: 0.059
bicor=-0.19, p=0.085
cor=-0.2, p=0.068
bicor=-0.18, p=0.075
cor=-0.17, p=0.091
CCT5 UPenn Mixed PRM
M11 greenyellow MEGA module member
K-W ANOVA p: 0.017
CCT5 UPenn Mixed PRM
K-W ANOVA p: 0.042
bicor=0.24, p=0.025
cor=0.27, p=0.013
bicor=0.26, p=0.0084
cor=0.27, p=0.0066
IDH2 UPenn Mixed PRM
M1 turquoise MEGA module member
K-W ANOVA p: 0.001
IDH2 UPenn Mixed PRM
K-W ANOVA p: 2.4e-05
bicor=-0.22, p=0.049
cor=-0.29, p=0.0075
bicor=-0.33, p=0.00071
cor=-0.38, p=9.6e-05
Relative Log2 Abundance
Control
AD
ALS
FTLDU
PSP
CBD
PD.PDD
MSA
otherND
Braak Score
CERAD Score

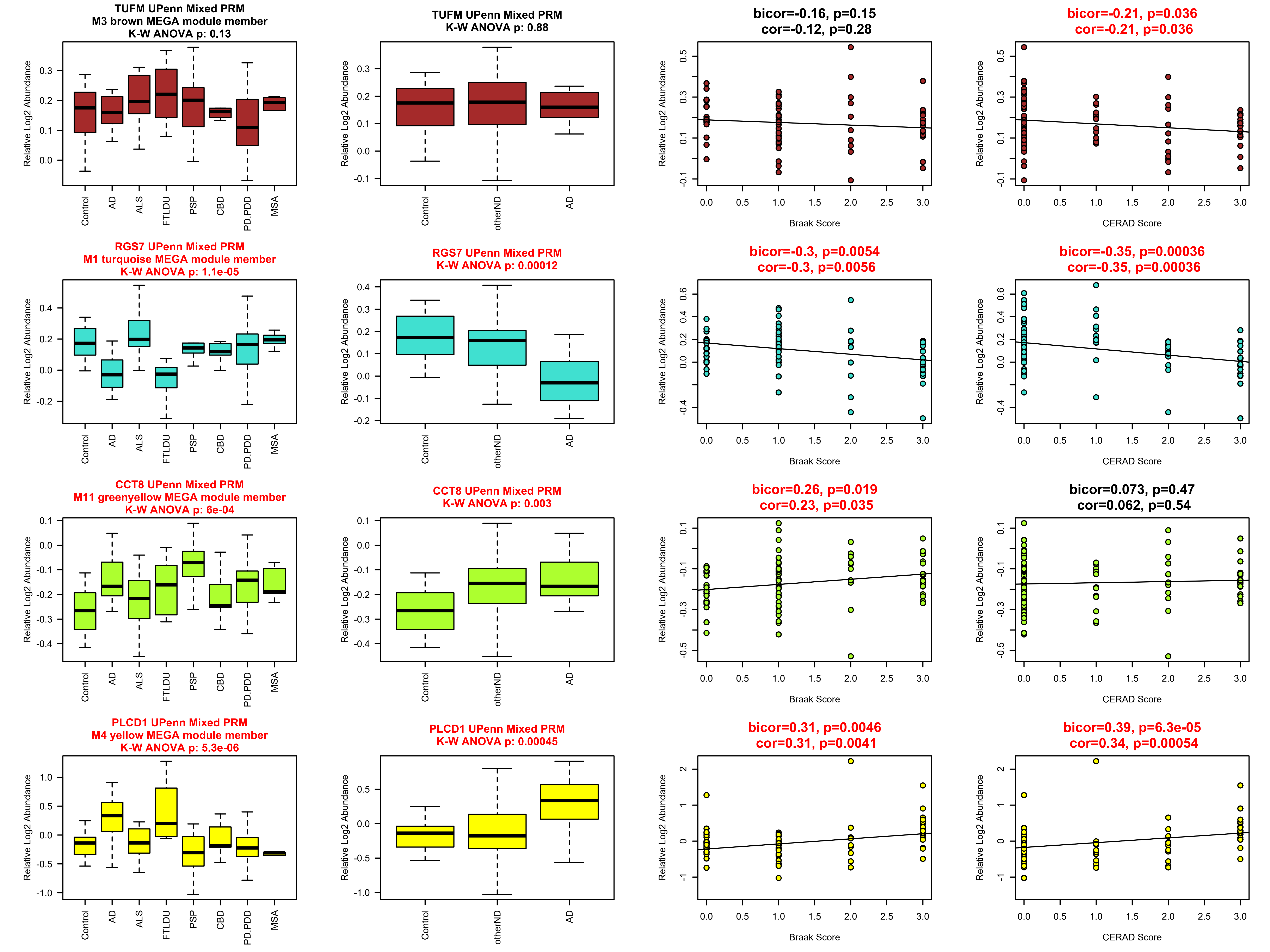

TUFM UPenn Mixed PRM
M3 brown MEGA module member
K-W ANOVA p: 0.13
TUFM UPenn Mixed PRM
K-W ANOVA p: 0.88
bicor=-0.16, p=0.15
cor=-0.12, p=0.28
bicor=-0.21, p=0.036
cor=-0.21, p=0.036
RGS7 UPenn Mixed PRM
M1 turquoise MEGA module member
K-W ANOVA p: 1.1e-05
RGS7 UPenn Mixed PRM
K-W ANOVA p: 0.00012
bicor=-0.3, p=0.0054
cor=-0.3, p=0.0056
bicor=-0.35, p=0.00036
cor=-0.35, p=0.00036
CCT8 UPenn Mixed PRM
M11 greenyellow MEGA module member
K-W ANOVA p: 6e-04
CCT8 UPenn Mixed PRM
K-W ANOVA p: 0.003
bicor=0.26, p=0.019
cor=0.23, p=0.035
bicor=0.073, p=0.47
cor=0.062, p=0.54
PLCD1 UPenn Mixed PRM
M4 yellow MEGA module member
K-W ANOVA p: 5.3e-06
PLCD1 UPenn Mixed PRM
K-W ANOVA p: 0.00045
bicor=0.31, p=0.0046
cor=0.31, p=0.0041
bicor=0.39, p=6.3e-05
cor=0.34, p=0.00054
Relative Log2 Abundance
Control
AD
ALS
FTLDU
PSP
CBD
PD.PDD
MSA
otherND
Braak Score
CERAD Score

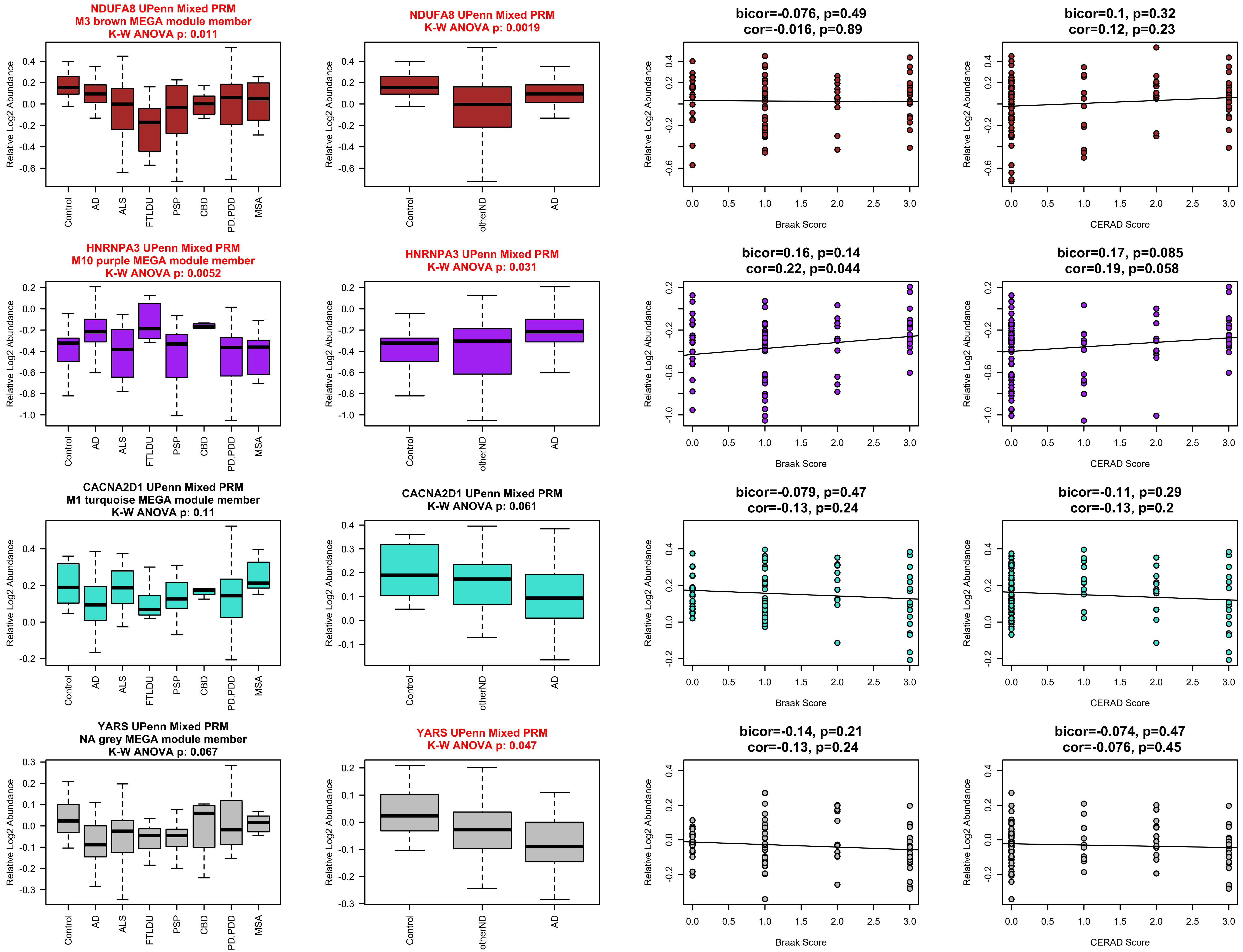

NDUFA8 UPenn Mixed PRM
M3 brown MEGA module member
K-W ANOVA p: 0.011
NDUFA8 UPenn Mixed PRM
K-W ANOVA p: 0.0019
bicor=-0.076, p=0.49
cor=-0.016, p=0.89
bicor=0.1, p=0.32
cor=0.12, p=0.23
HNRNPA3 UPenn Mixed PRM
M10 purple MEGA module member
K-W ANOVA p: 0.0052
HNRNPA3 UPenn Mixed PRM
K-W ANOVA p: 0.031
bicor=0.16, p=0.14
cor=0.22, p=0.044
bicor=0.17, p=0.085
cor=0.19, p=0.058
CACNA2D1 UPenn Mixed PRM
M1 turquoise MEGA module member
K-W ANOVA p: 0.11
CACNA2D1 UPenn Mixed PRM
K-W ANOVA p: 0.061
bicor=-0.079, p=0.47
cor=-0.13, p=0.24
bicor=-0.11, p=0.29
cor=-0.13, p=0.2
YARS UPenn Mixed PRM
NA grey MEGA module member
K-W ANOVA p: 0.067
YARS UPenn Mixed PRM
K-W ANOVA p: 0.047
bicor=-0.14, p=0.21
cor=-0.13, p=0.24
bicor=-0.074, p=0.47
cor=-0.076, p=0.45
Relative Log2 Abundance
Control
AD
ALS
FTLDU
PSP
CBD
PD.PDD
MSA
otherND
Braak Score
CERAD Score

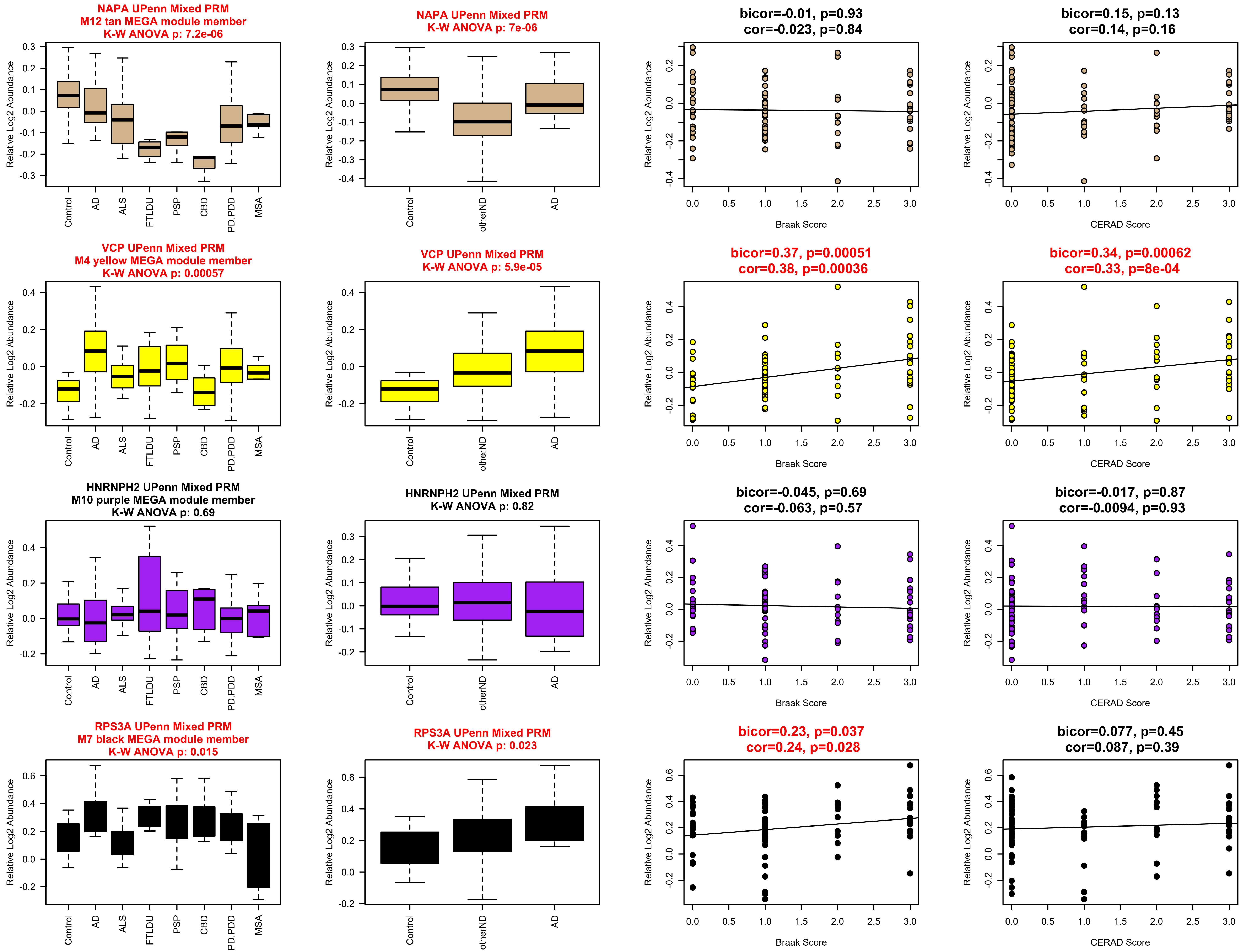
NAPA UPenn Mixed PRM
M12 tan MEGA module member
K-W ANOVA p: 7.2e-06
NAPA UPenn Mixed PRM
K-W ANOVA p: 7e-06
bicor=-0.01, p=0.93
cor=-0.023, p=0.84
bicor=0.15, p=0.13
cor=0.14, p=0.16
VCP UPenn Mixed PRM
M4 yellow MEGA module member
K-W ANOVA p: 0.00057
VCP UPenn Mixed PRM
K-W ANOVA p: 5.9e-05
bicor=0.37, p=0.00051
cor=0.38, p=0.00036
bicor=0.34, p=0.00062
cor=0.33, p=8e-04
HNRNPH2 UPenn Mixed PRM
M10 purple MEGA module member
K-W ANOVA p: 0.69
HNRNPH2 UPenn Mixed PRM
K-W ANOVA p: 0.82
bicor=-0.045, p=0.69
cor=-0.063, p=0.57
bicor=-0.017, p=0.87
cor=-0.0094, p=0.93
RPS3A UPenn Mixed PRM
M7 black MEGA module member
K-W ANOVA p: 0.015
RPS3A UPenn Mixed PRM
K-W ANOVA p: 0.023
bicor=0.23, p=0.037
cor=0.24, p=0.028
bicor=0.077, p=0.45
cor=0.087, p=0.39
Relative Log2 Abundance
Control
AD
ALS
FTLDU
PSP
CBD
PD.PDD
MSA
otherND
Braak Score
CERAD Score

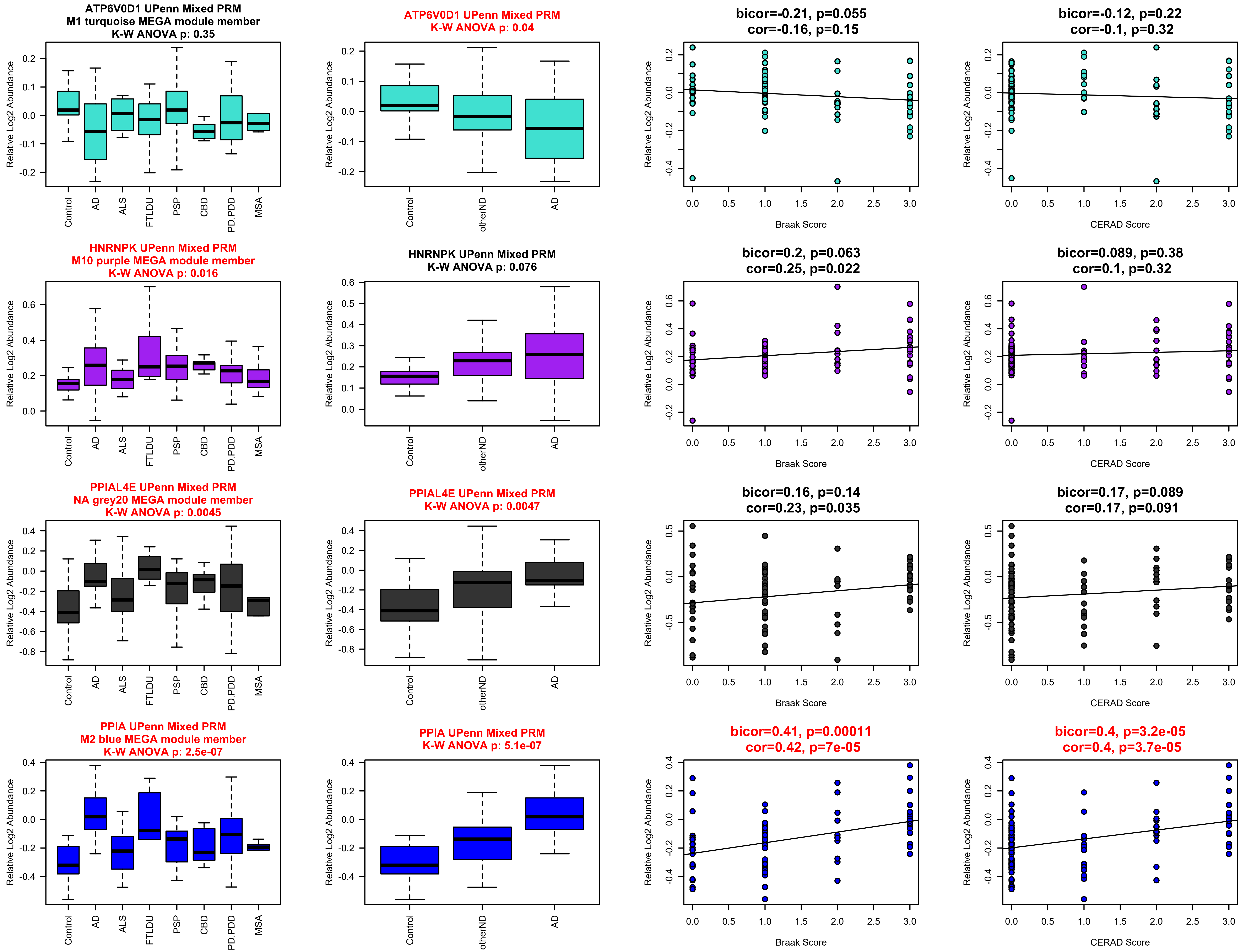
ATP6V0D1 UPenn Mixed PRM
M1 turquoise MEGA module member
K-W ANOVA p: 0.35
Relative Log2 Abundance
Control
AD
ALS
FTLDU
PSP
CBD
PD.PDD
MSA
ATP6V0D1 UPenn Mixed PRM
K-W ANOVA p: 0.04
Relative Log2 Abundance
Control
otherND
AD
bicor=-0.21, p=0.055
cor=-0.16, p=0.15
Relative Log2 Abundance
Braak Score
bicor=-0.12, p=0.22
cor=-0.1, p=0.32
Relative Log2 Abundance
CERAD Score
HNRNPK UPenn Mixed PRM
M10 purple MEGA module member
K-W ANOVA p: 0.016
Relative Log2 Abundance
Control
AD
ALS
FTLDU
PSP
CBD
PD.PDD
MSA
HNRNPK UPenn Mixed PRM
K-W ANOVA p: 0.076
Relative Log2 Abundance
Control
otherND
AD
bicor=0.2, p=0.063
cor=0.25, p=0.022
Relative Log2 Abundance
Braak Score
bicor=0.089, p=0.38
cor=0.1, p=0.32
Relative Log2 Abundance
CERAD Score
PPIAL4E UPenn Mixed PRM
NA grey20 MEGA module member
K-W ANOVA p: 0.0045
Relative Log2 Abundance
Control
AD
ALS
FTLDU
PSP
CBD
PD.PDD
MSA
PPIAL4E UPenn Mixed PRM
K-W ANOVA p: 0.0047
Relative Log2 Abundance
Control
otherND
AD
bicor=0.16, p=0.14
cor=0.23, p=0.035
Relative Log2 Abundance
Braak Score
bicor=0.17, p=0.089
cor=0.17, p=0.091
Relative Log2 Abundance
CERAD Score
PPIA UPenn Mixed PRM
M2 blue MEGA module member
K-W ANOVA p: 2.5e-07
Relative Log2 Abundance
Control
AD
ALS
FTLDU
PSP
CBD
PD.PDD
MSA
PPIA UPenn Mixed PRM
K-W ANOVA p: 5.1e-07
Relative Log2 Abundance
Control
otherND
AD
bicor=0.41, p=0.00011
cor=0.42, p=7e-05
Relative Log2 Abundance
Braak Score
bicor=0.4, p=3.2e-05
cor=0.4, p=3.7e-05
Relative Log2 Abundance
CERAD Score

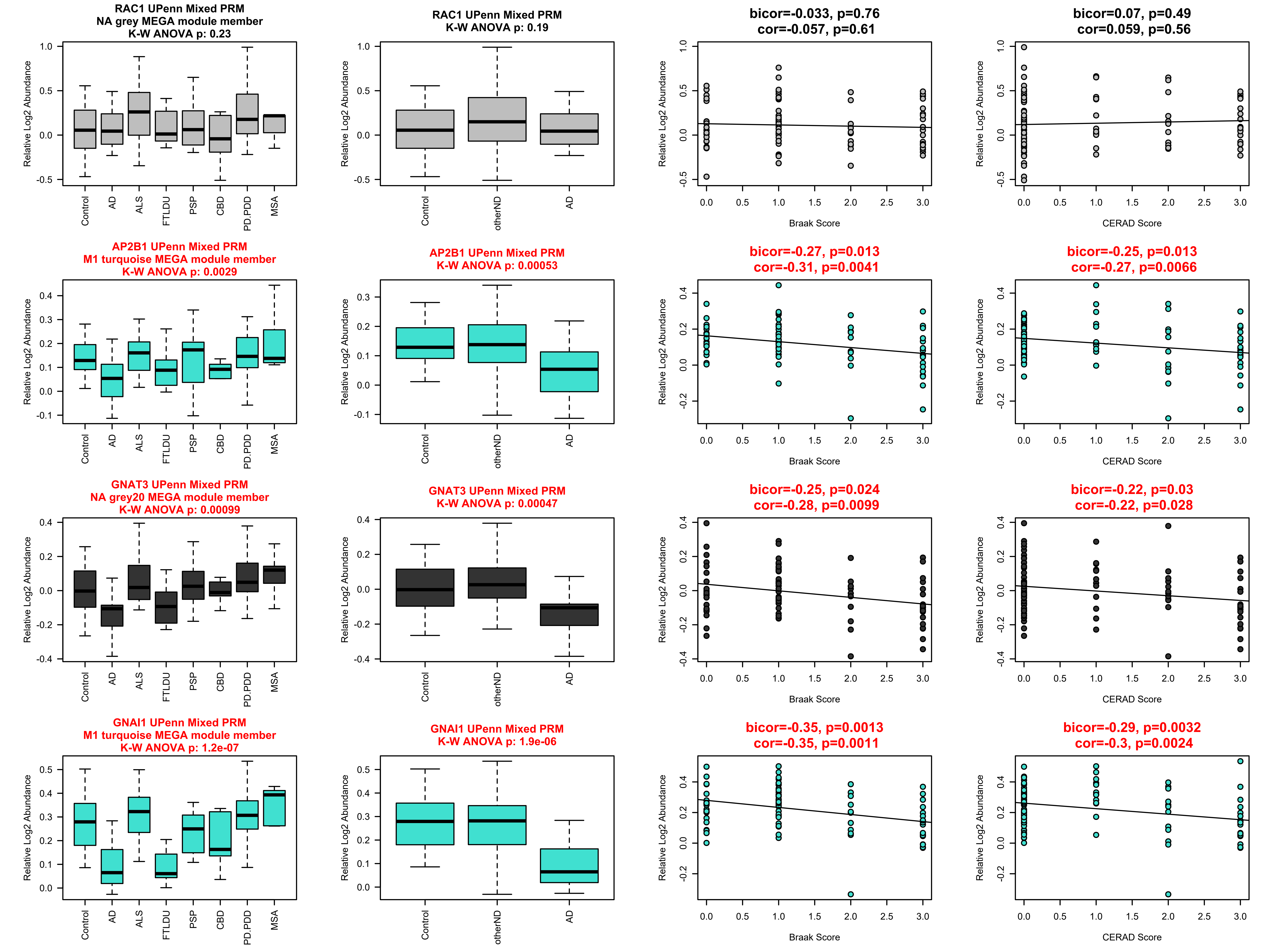
RAC1 UPenn Mixed PRM
NA grey MEGA module member
K-W ANOVA p: 0.23
RAC1 UPenn Mixed PRM
K-W ANOVA p: 0.19
bicor=-0.033, p=0.76
cor=-0.057, p=0.61
bicor=0.07, p=0.49
cor=0.059, p=0.56
AP2B1 UPenn Mixed PRM
M1 turquoise MEGA module member
K-W ANOVA p: 0.0029
AP2B1 UPenn Mixed PRM
K-W ANOVA p: 0.00053
bicor=-0.27, p=0.013
cor=-0.31, p=0.0041
bicor=-0.25, p=0.013
cor=-0.27, p=0.0066
GNAT3 UPenn Mixed PRM
NA grey20 MEGA module member
K-W ANOVA p: 0.00099
GNAT3 UPenn Mixed PRM
K-W ANOVA p: 0.00047
bicor=-0.25, p=0.024
cor=-0.28, p=0.0099
bicor=-0.22, p=0.03
cor=-0.22, p=0.028
GNAI1 UPenn Mixed PRM
M1 turquoise MEGA module member
K-W ANOVA p: 1.2e-07
GNAI1 UPenn Mixed PRM
K-W ANOVA p: 1.9e-06
bicor=-0.35, p=0.0013
cor=-0.35, p=0.0011
bicor=-0.29, p=0.0032
cor=-0.3, p=0.0024
Relative Log2 Abundance
Control
AD
ALS
FTLDU
PSP
CBD
PD.PDD
MSA
otherND
Braak Score
CERAD Score

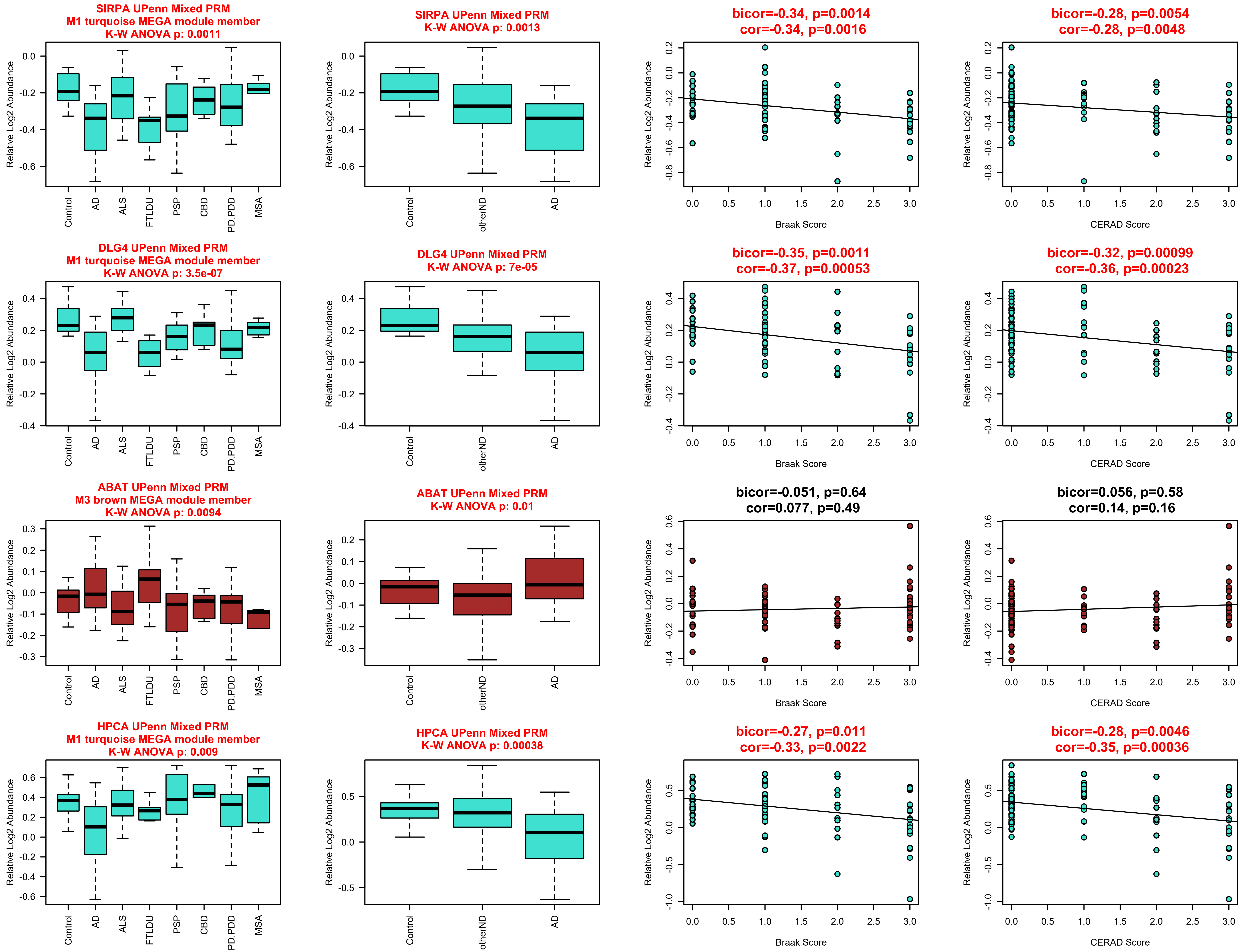
SIRPA UPenn Mixed PRM
M1 turquoise MEGA module member
K-W ANOVA p: 0.0011
SIRPA UPenn Mixed PRM
K-W ANOVA p: 0.0013
bicor=-0.34, p=0.0014
cor=-0.34, p=0.0016
bicor=-0.28, p=0.0054
cor=-0.28, p=0.0048
DLG4 UPenn Mixed PRM
M1 turquoise MEGA module member
K-W ANOVA p: 3.5e-07
DLG4 UPenn Mixed PRM
K-W ANOVA p: 7e-05
bicor=-0.35, p=0.0011
cor=-0.37, p=0.00053
bicor=-0.32, p=0.00099
cor=-0.36, p=0.00023
ABAT UPenn Mixed PRM
M3 brown MEGA module member
K-W ANOVA p: 0.0094
ABAT UPenn Mixed PRM
K-W ANOVA p: 0.01
bicor=-0.051, p=0.64
cor=0.077, p=0.49
bicor=0.056, p=0.58
cor=0.14, p=0.16
HPCA UPenn Mixed PRM
M1 turquoise MEGA module member
K-W ANOVA p: 0.009
HPCA UPenn Mixed PRM
K-W ANOVA p: 0.00038
bicor=-0.27, p=0.011
cor=-0.33, p=0.0022
bicor=-0.28, p=0.0046
cor=-0.35, p=0.00036
Relative Log2 Abundance
Control
AD
ALS
FTLDU
PSP
CBD
PD.PDD
MSA
otherND
Braak Score
CERAD Score

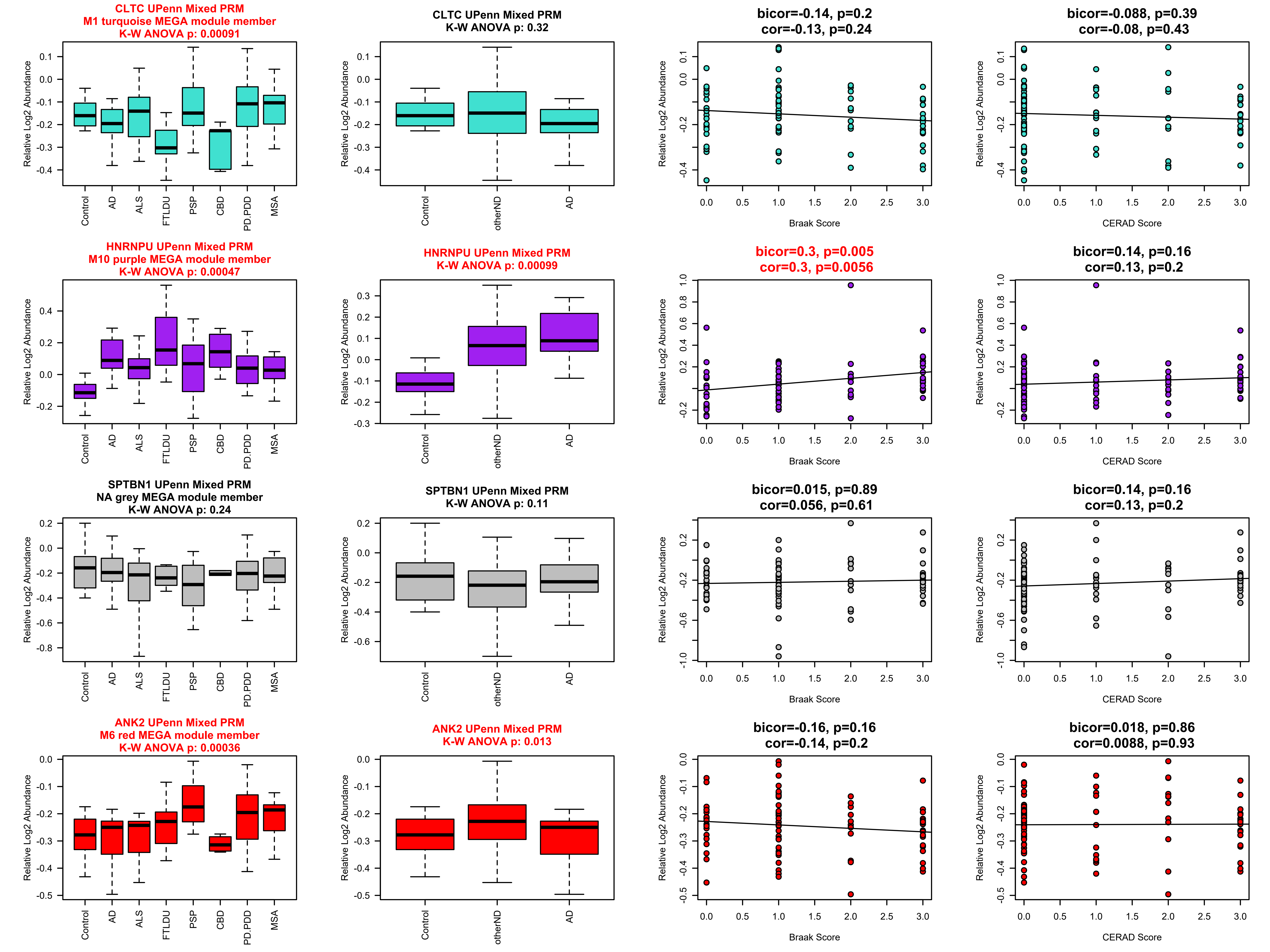

CLTC UPenn Mixed PRM
M1 turquoise MEGA module member
K-W ANOVA p: 0.00091
CLTC UPenn Mixed PRM
K-W ANOVA p: 0.32
bicor=-0.14, p=0.2
cor=-0.13, p=0.24
bicor=-0.088, p=0.39
cor=-0.08, p=0.43
HNRNPU UPenn Mixed PRM
M10 purple MEGA module member
K-W ANOVA p: 0.00047
HNRNPU UPenn Mixed PRM
K-W ANOVA p: 0.00099
bicor=0.3, p=0.005
cor=0.3, p=0.0056
bicor=0.14, p=0.16
cor=0.13, p=0.2
SPTBN1 UPenn Mixed PRM
NA grey MEGA module member
K-W ANOVA p: 0.24
SPTBN1 UPenn Mixed PRM
K-W ANOVA p: 0.11
bicor=0.015, p=0.89
cor=0.056, p=0.61
bicor=0.14, p=0.16
cor=0.13, p=0.2
ANK2 UPenn Mixed PRM
M6 red MEGA module member
K-W ANOVA p: 0.00036
ANK2 UPenn Mixed PRM
K-W ANOVA p: 0.013
bicor=-0.16, p=0.16
cor=-0.14, p=0.2
bicor=0.018, p=0.86
cor=0.0088, p=0.93
Relative Log2 Abundance
Control
AD
ALS
FTLDU
PSP
CBD
PD.PDD
MSA
otherND
Braak Score
CERAD Score

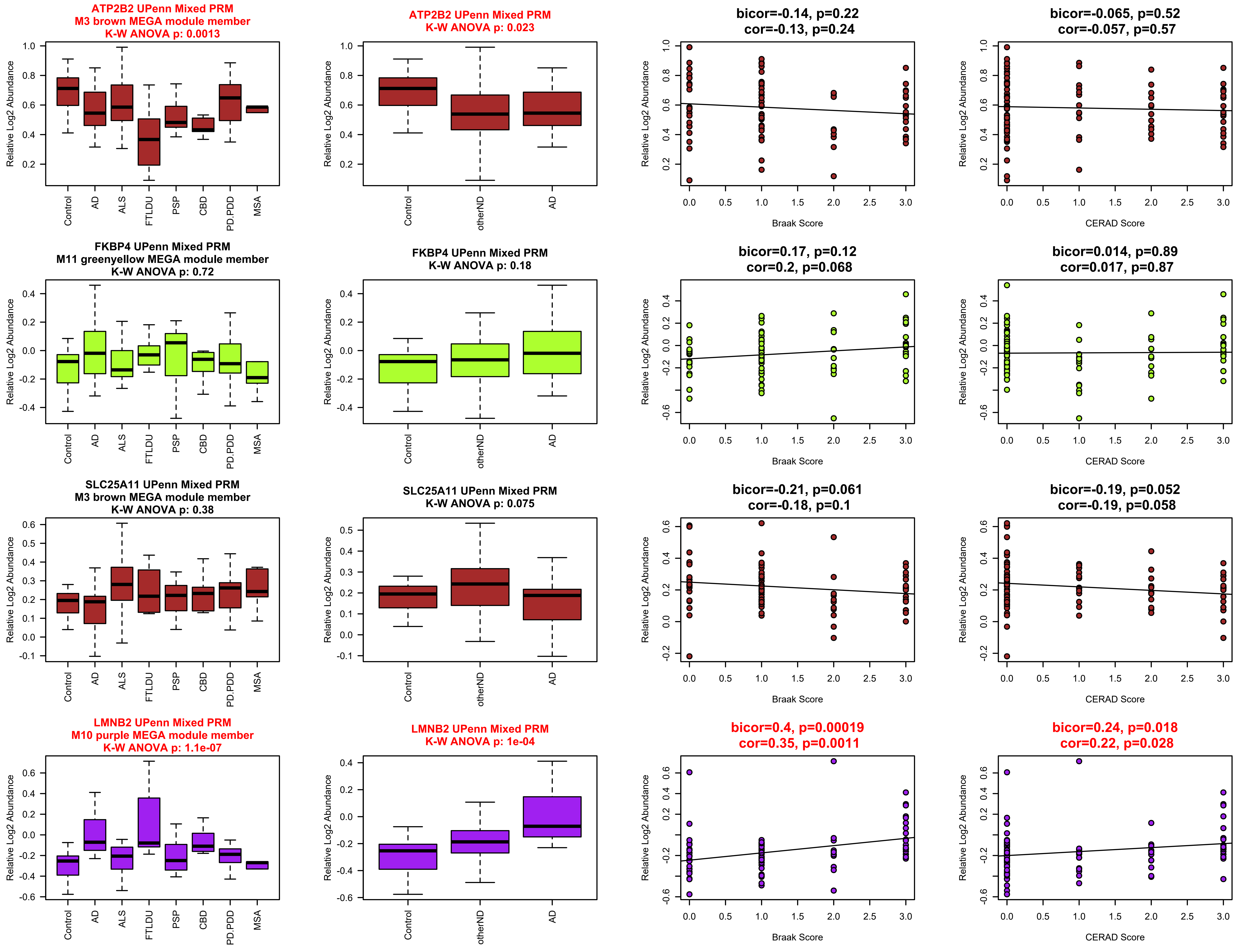
ATP2B2 UPenn Mixed PRM
M3 brown MEGA module member
K-W ANOVA p: 0.0013
ATP2B2 UPenn Mixed PRM
K-W ANOVA p: 0.023
bicor=-0.14, p=0.22
cor=-0.13, p=0.24
bicor=-0.065, p=0.52
cor=-0.057, p=0.57
FKBP4 UPenn Mixed PRM
M11 greenyellow MEGA module member
K-W ANOVA p: 0.72
FKBP4 UPenn Mixed PRM
K-W ANOVA p: 0.18
bicor=0.17, p=0.12
cor=0.2, p=0.068
bicor=0.014, p=0.89
cor=0.017, p=0.87
SLC25A11 UPenn Mixed PRM
M3 brown MEGA module member
K-W ANOVA p: 0.38
SLC25A11 UPenn Mixed PRM
K-W ANOVA p: 0.075
bicor=-0.21, p=0.061
cor=-0.18, p=0.1
bicor=-0.19, p=0.052
cor=-0.19, p=0.058
LMNB2 UPenn Mixed PRM
M10 purple MEGA module member
K-W ANOVA p: 1.1e-07
LMNB2 UPenn Mixed PRM
K-W ANOVA p: 1e-04
bicor=0.4, p=0.00019
cor=0.35, p=0.0011
bicor=0.24, p=0.018
cor=0.22, p=0.028
Relative Log2 Abundance
Control
AD
ALS
FTLDU
PSP
CBD
PD.PDD
MSA
otherND
Braak Score
CERAD Score

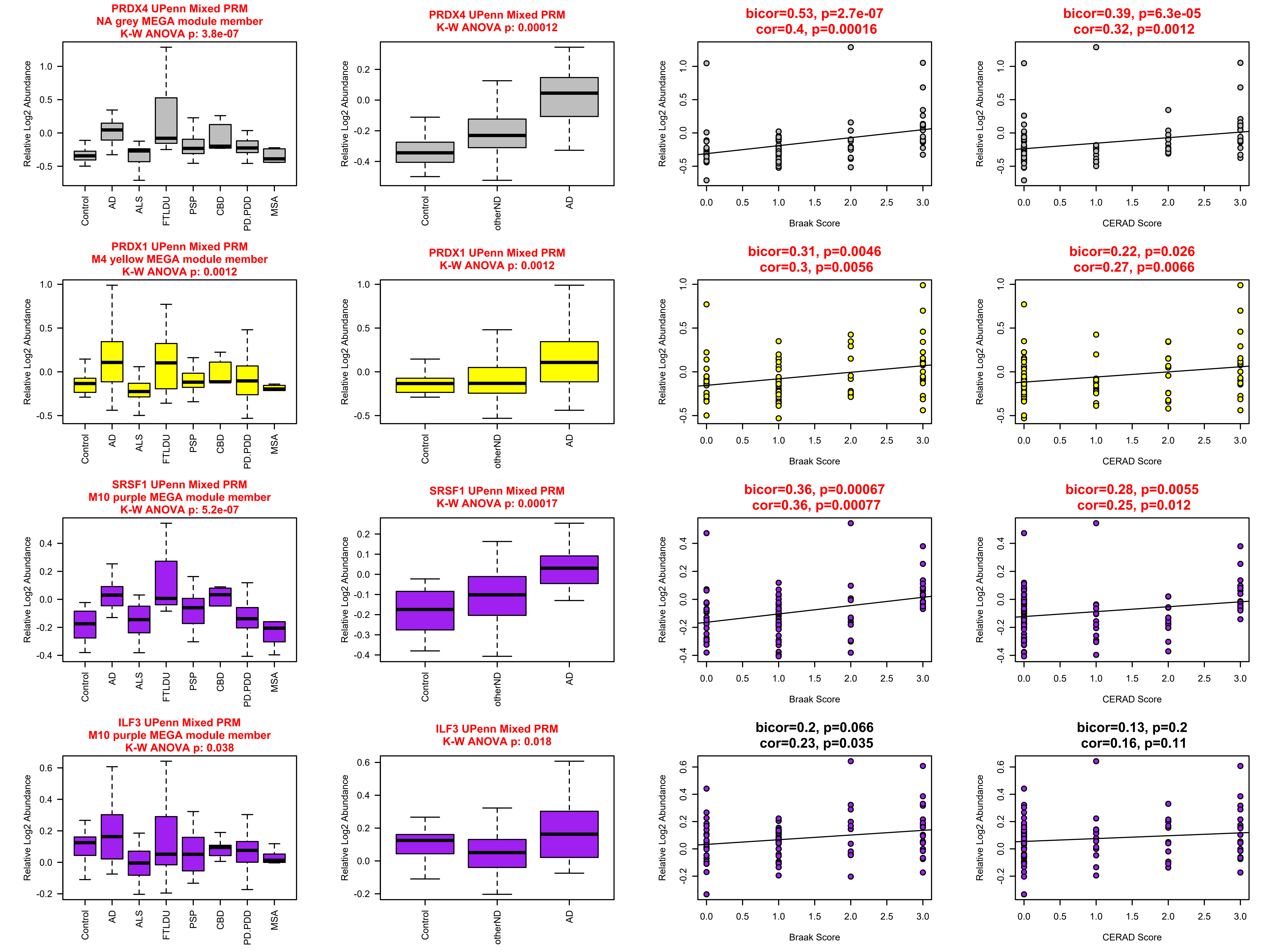
PRDX4 UPenn Mixed PRM
NA grey MEGA module member
K-W ANOVA p: 3.8e-07
PRDX4 UPenn Mixed PRM
K-W ANOVA p: 0.00012
bicor=0.53, p=2.7e-07
cor=0.4, p=0.00016
bicor=0.39, p=6.3e-05
cor=0.32, p=0.0012
PRDX1 UPenn Mixed PRM
M4 yellow MEGA module member
K-W ANOVA p: 0.0012
PRDX1 UPenn Mixed PRM
K-W ANOVA p: 0.0012
bicor=0.31, p=0.0046
cor=0.3, p=0.0056
bicor=0.22, p=0.026
cor=0.27, p=0.0066
SRSF1 UPenn Mixed PRM
M10 purple MEGA module member
K-W ANOVA p: 5.2e-07
SRSF1 UPenn Mixed PRM
K-W ANOVA p: 0.00017
bicor=0.36, p=0.00067
cor=0.36, p=0.00077
bicor=0.28, p=0.0055
cor=0.25, p=0.012
ILF3 UPenn Mixed PRM
M10 purple MEGA module member
K-W ANOVA p: 0.038
ILF3 UPenn Mixed PRM
K-W ANOVA p: 0.018
bicor=0.2, p=0.066
cor=0.23, p=0.035
bicor=0.13, p=0.2
cor=0.16, p=0.11
Relative Log2 Abundance
Control
AD
ALS
FTLDU
PSP
CBD
PD.PDD
MSA
otherND
Braak Score
CERAD Score

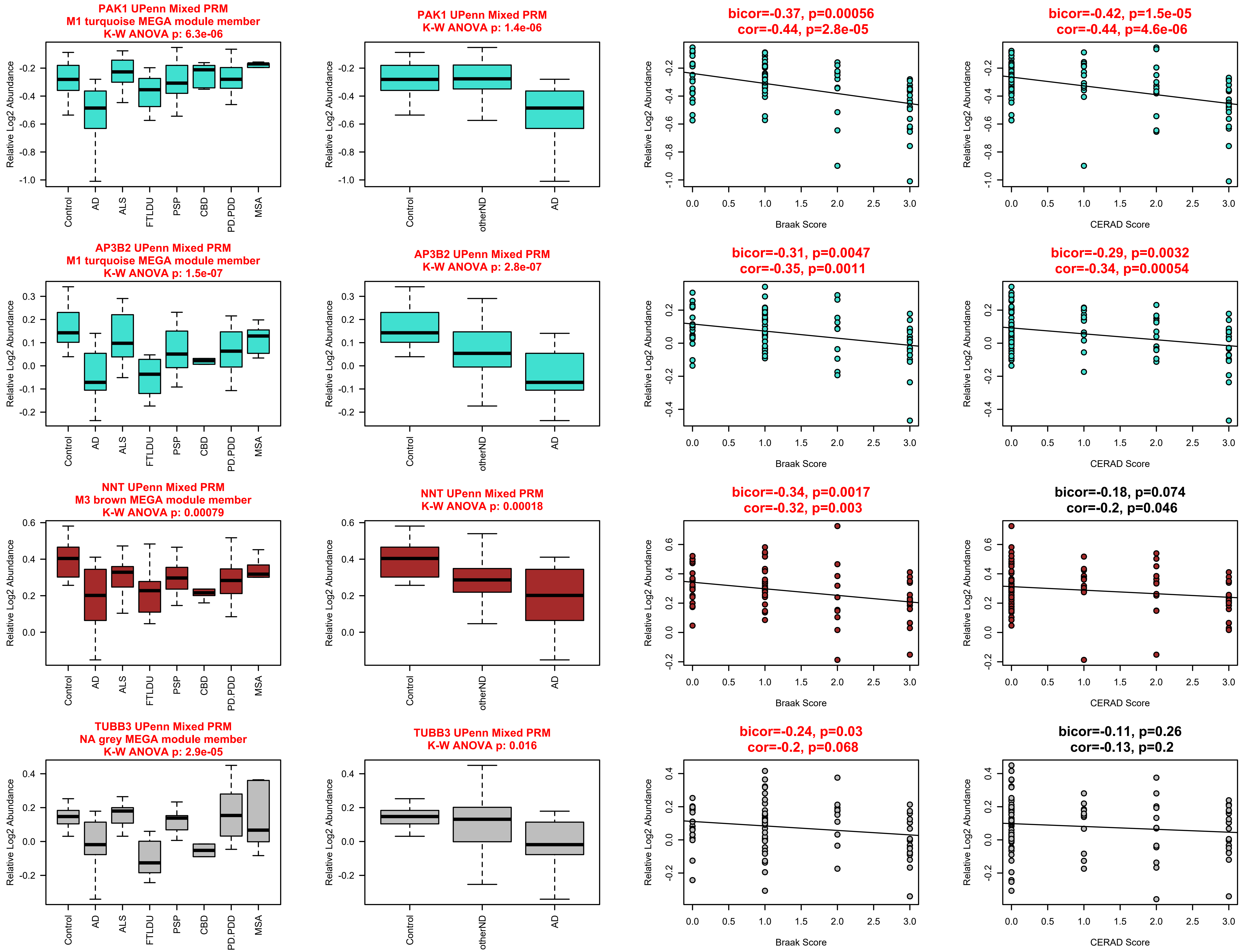
PAK1 UPenn Mixed PRM
M1 turquoise MEGA module member
K-W ANOVA p: 6.3e-06
PAK1 UPenn Mixed PRM
K-W ANOVA p: 1.4e-06
bicor=-0.37, p=0.00056
cor=-0.44, p=2.8e-05
bicor=-0.42, p=1.5e-05
cor=-0.44, p=4.6e-06
AP3B2 UPenn Mixed PRM
M1 turquoise MEGA module member
K-W ANOVA p: 1.5e-07
AP3B2 UPenn Mixed PRM
K-W ANOVA p: 2.8e-07
bicor=-0.31, p=0.0047
cor=-0.35, p=0.0011
bicor=-0.29, p=0.0032
cor=-0.34, p=0.00054
NNT UPenn Mixed PRM
M3 brown MEGA module member
K-W ANOVA p: 0.00079
NNT UPenn Mixed PRM
K-W ANOVA p: 0.00018
bicor=-0.34, p=0.0017
cor=-0.32, p=0.003
bicor=-0.18, p=0.074
cor=-0.2, p=0.046
TUBB3 UPenn Mixed PRM
NA grey MEGA module member
K-W ANOVA p: 2.9e-05
TUBB3 UPenn Mixed PRM
K-W ANOVA p: 0.016
bicor=-0.24, p=0.03
cor=-0.2, p=0.068
bicor=-0.11, p=0.26
cor=-0.13, p=0.2
Relative Log2 Abundance
Control
AD
ALS
FTLDU
PSP
CBD
PD.PDD
MSA
otherND
Braak Score
CERAD Score

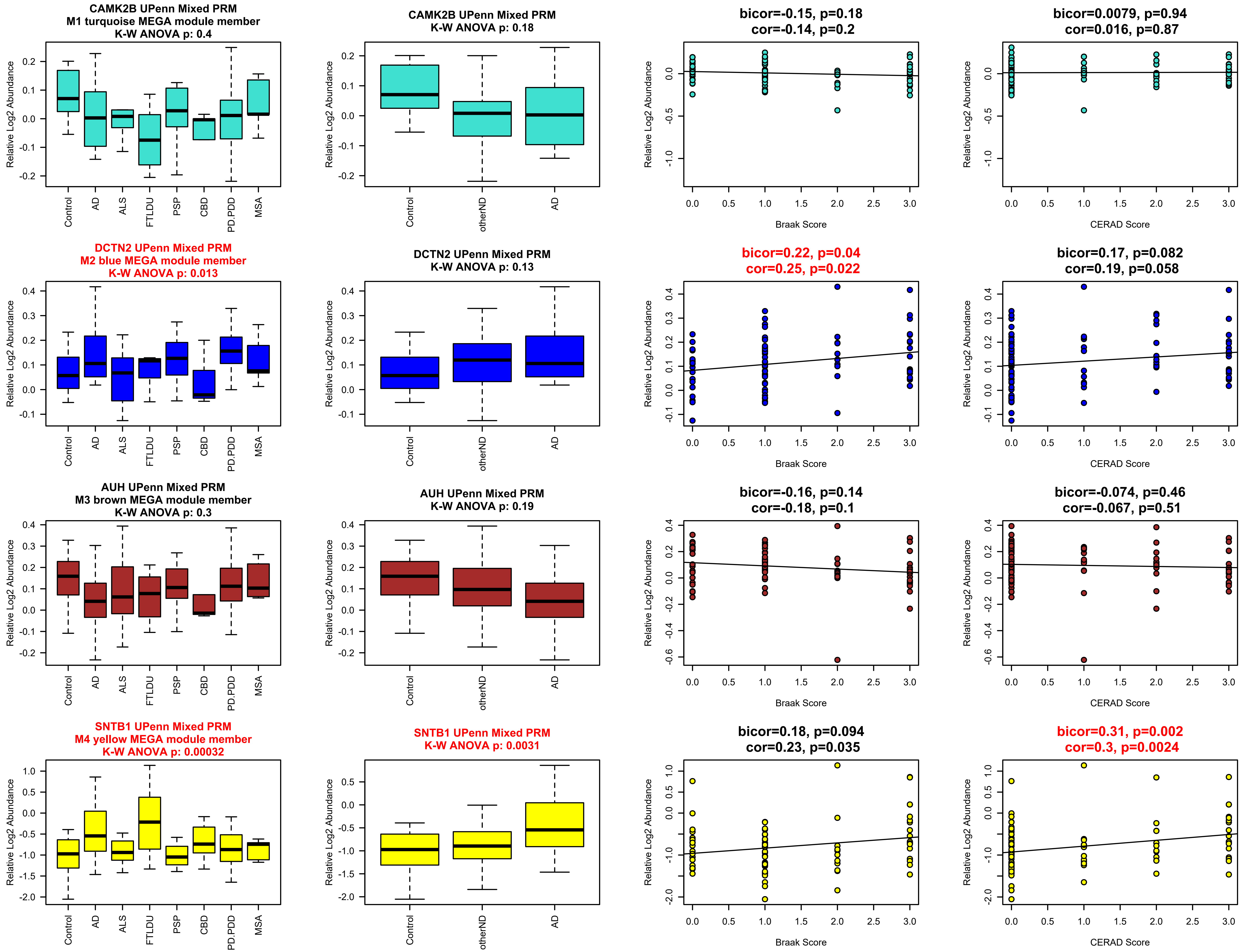
CAMK2B UPenn Mixed PRM
M1 turquoise MEGA module member
K-W ANOVA p: 0.4
CAMK2B UPenn Mixed PRM
K-W ANOVA p: 0.18
bicor=-0.15, p=0.18
cor=-0.14, p=0.2
bicor=0.0079, p=0.94
cor=0.016, p=0.87
DCTN2 UPenn Mixed PRM
M2 blue MEGA module member
K-W ANOVA p: 0.013
DCTN2 UPenn Mixed PRM
K-W ANOVA p: 0.13
bicor=0.22, p=0.04
cor=0.25, p=0.022
bicor=0.17, p=0.082
cor=0.19, p=0.058
AUH UPenn Mixed PRM
M3 brown MEGA module member
K-W ANOVA p: 0.3
AUH UPenn Mixed PRM
K-W ANOVA p: 0.19
bicor=-0.16, p=0.14
cor=-0.18, p=0.1
bicor=-0.074, p=0.46
cor=-0.067, p=0.51
SNTB1 UPenn Mixed PRM
M4 yellow MEGA module member
K-W ANOVA p: 0.00032
SNTB1 UPenn Mixed PRM
K-W ANOVA p: 0.0031
bicor=0.18, p=0.094
cor=0.23, p=0.035
bicor=0.31, p=0.002
cor=0.3, p=0.0024
Relative Log2 Abundance
Control
AD
ALS
FTLDU
PSP
CBD
PD.PDD
MSA
otherND
Braak Score
CERAD Score

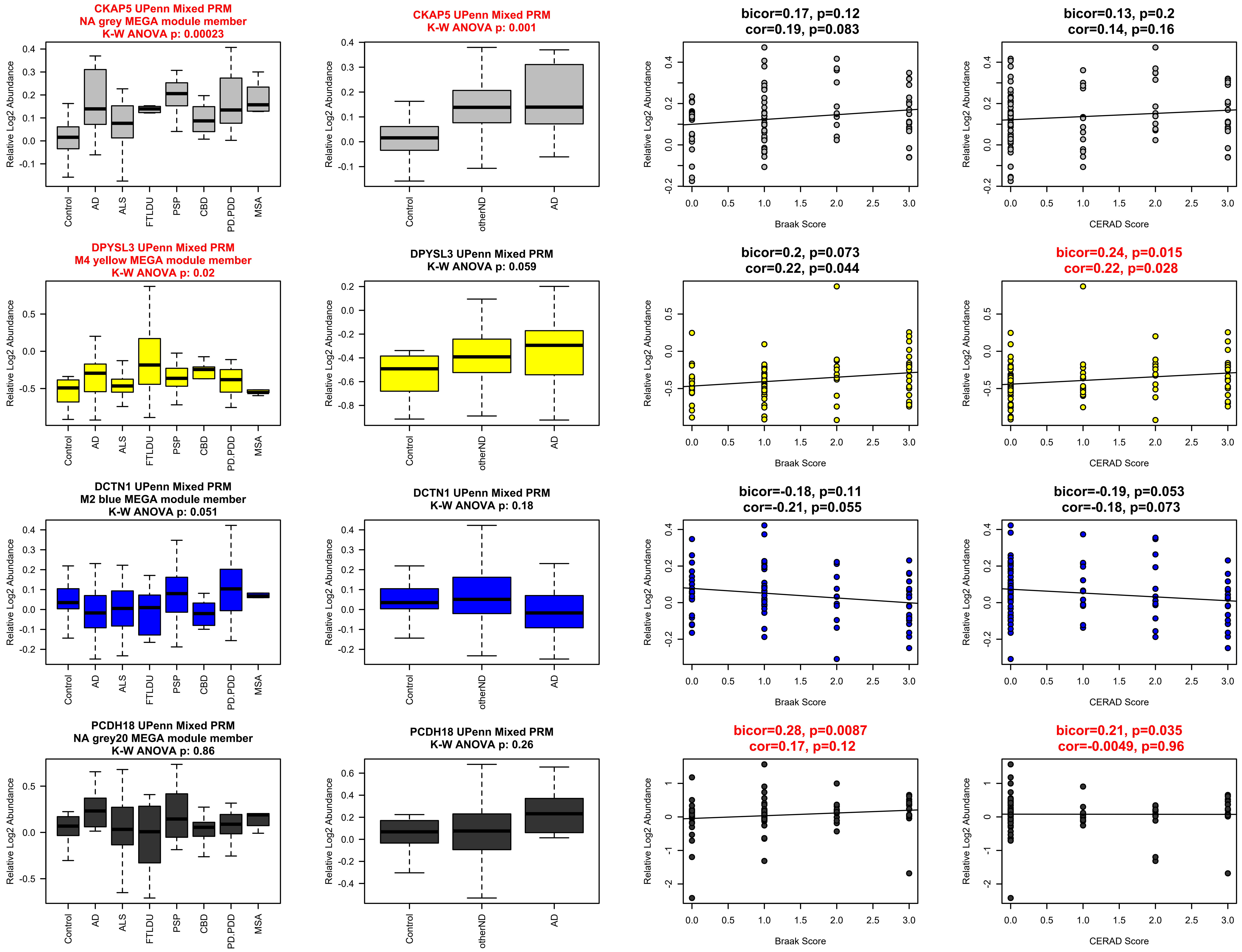
CKAP5 UPenn Mixed PRM
NA grey MEGA module member
K-W ANOVA p: 0.00023
CKAP5 UPenn Mixed PRM
K-W ANOVA p: 0.001
bicor=0.17, p=0.12
cor=0.19, p=0.083
bicor=0.13, p=0.2
cor=0.14, p=0.16
DPYSL3 UPenn Mixed PRM
M4 yellow MEGA module member
K-W ANOVA p: 0.02
DPYSL3 UPenn Mixed PRM
K-W ANOVA p: 0.059
bicor=0.2, p=0.073
cor=0.22, p=0.044
bicor=0.24, p=0.015
cor=0.22, p=0.028
DCTN1 UPenn Mixed PRM
M2 blue MEGA module member
K-W ANOVA p: 0.051
DCTN1 UPenn Mixed PRM
K-W ANOVA p: 0.18
bicor=-0.18, p=0.11
cor=-0.21, p=0.055
bicor=-0.19, p=0.053
cor=-0.18, p=0.073
PCDH18 UPenn Mixed PRM
NA grey20 MEGA module member
K-W ANOVA p: 0.86
PCDH18 UPenn Mixed PRM
K-W ANOVA p: 0.26
bicor=0.28, p=0.0087
cor=0.17, p=0.12
bicor=0.21, p=0.035
cor=-0.0049, p=0.96
Relative Log2 Abundance
Control
AD
ALS
FTLDU
PSP
CBD
PD.PDD
MSA
otherND
Braak Score
CERAD Score

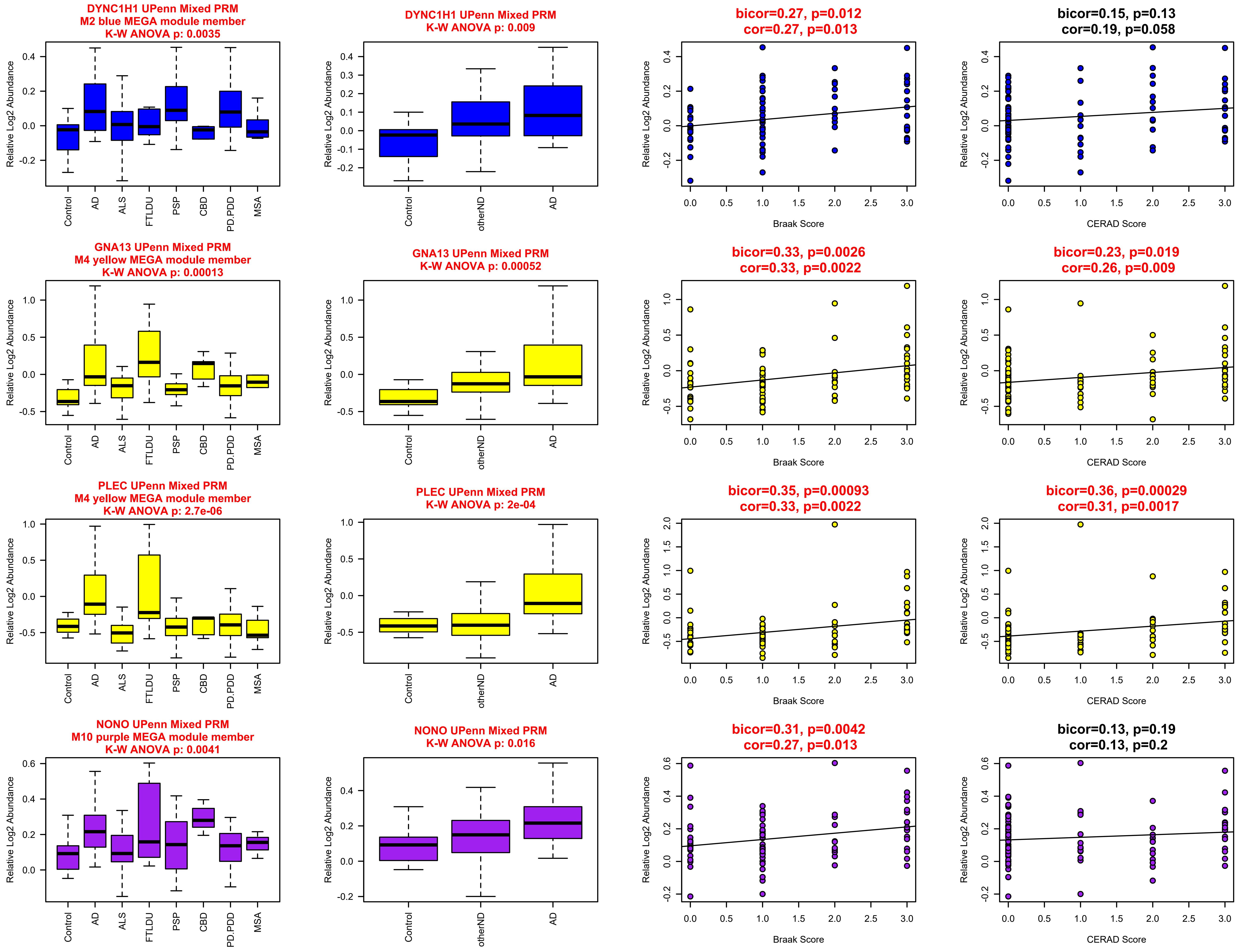

DYNC1H1 UPenn Mixed PRM
M2 blue MEGA module member
K-W ANOVA p: 0.0035
DYNC1H1 UPenn Mixed PRM
K-W ANOVA p: 0.009
bicor=0.27, p=0.012
cor=0.27, p=0.013
bicor=0.15, p=0.13
cor=0.19, p=0.058
GNA13 UPenn Mixed PRM
M4 yellow MEGA module member
K-W ANOVA p: 0.00013
GNA13 UPenn Mixed PRM
K-W ANOVA p: 0.00052
bicor=0.33, p=0.0026
cor=0.33, p=0.0022
bicor=0.23, p=0.019
cor=0.26, p=0.009
PLEC UPenn Mixed PRM
M4 yellow MEGA module member
K-W ANOVA p: 2.7e-06
PLEC UPenn Mixed PRM
K-W ANOVA p: 2e-04
bicor=0.35, p=0.00093
cor=0.33, p=0.0022
bicor=0.36, p=0.00029
cor=0.31, p=0.0017
NONO UPenn Mixed PRM
M10 purple MEGA module member
K-W ANOVA p: 0.0041
NONO UPenn Mixed PRM
K-W ANOVA p: 0.016
bicor=0.31, p=0.0042
cor=0.27, p=0.013
bicor=0.13, p=0.19
cor=0.13, p=0.2
Relative Log2 Abundance
Control
AD
ALS
FTLDU
PSP
CBD
PD.PDD
MSA
otherND
Braak Score
CERAD Score
0.0
0.5
1.0
1.5
2.0
2.5
3.0

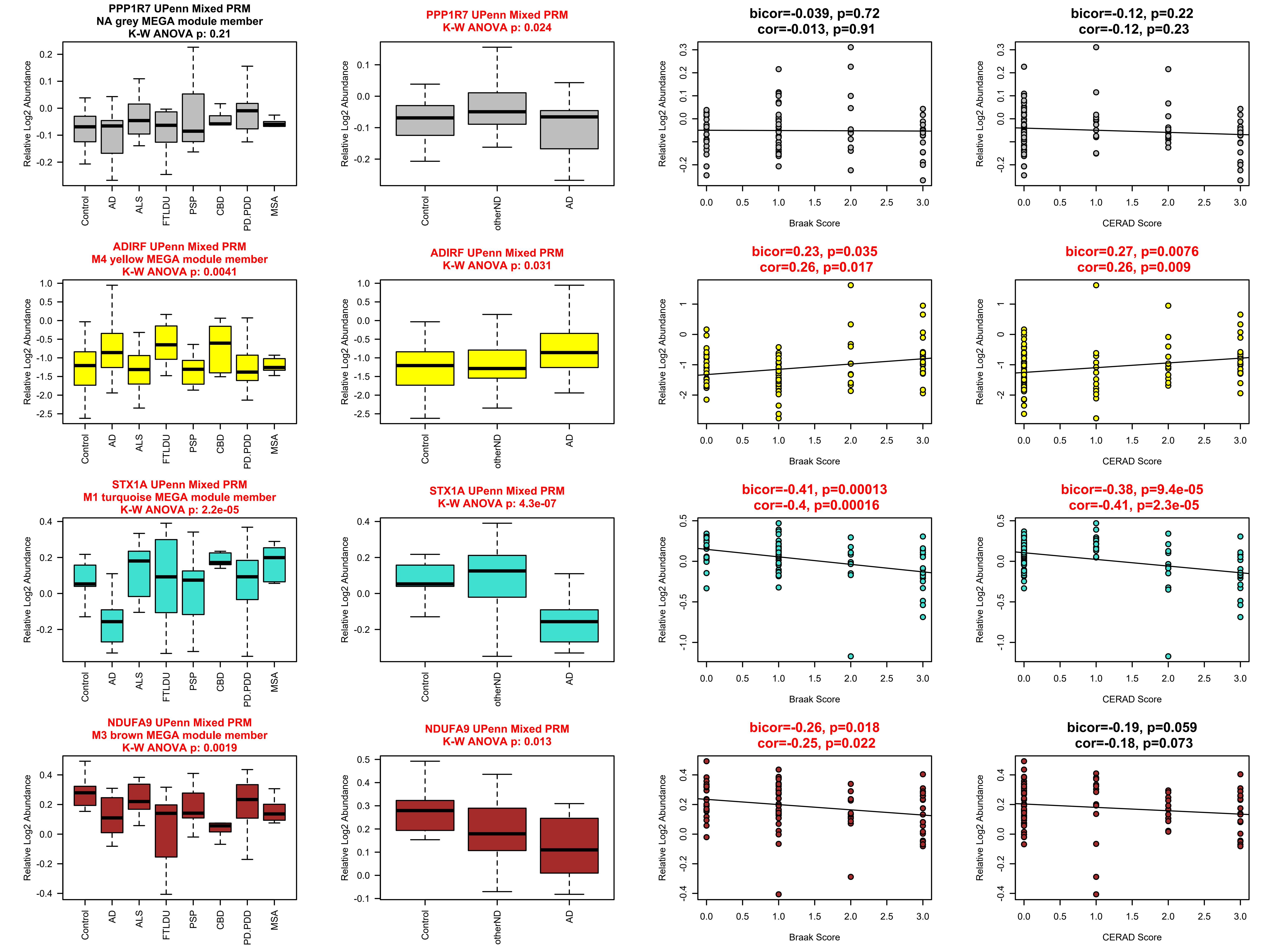
PPP1R7 UPenn Mixed PRM
NA grey MEGA module member
K-W ANOVA p: 0.21
PPP1R7 UPenn Mixed PRM
K-W ANOVA p: 0.024
bicor=-0.039, p=0.72
cor=-0.013, p=0.91
bicor=-0.12, p=0.22
cor=-0.12, p=0.23
ADIRF UPenn Mixed PRM
M4 yellow MEGA module member
K-W ANOVA p: 0.0041
ADIRF UPenn Mixed PRM
K-W ANOVA p: 0.031
bicor=0.23, p=0.035
cor=0.26, p=0.017
bicor=0.27, p=0.0076
cor=0.26, p=0.009
STX1A UPenn Mixed PRM
M1 turquoise MEGA module member
K-W ANOVA p: 2.2e-05
STX1A UPenn Mixed PRM
K-W ANOVA p: 4.3e-07
bicor=-0.41, p=0.00013
cor=-0.4, p=0.00016
bicor=-0.38, p=9.4e-05
cor=-0.41, p=2.3e-05
NDUFA9 UPenn Mixed PRM
M3 brown MEGA module member
K-W ANOVA p: 0.0019
NDUFA9 UPenn Mixed PRM
K-W ANOVA p: 0.013
bicor=-0.26, p=0.018
cor=-0.25, p=0.022
bicor=-0.19, p=0.059
cor=-0.18, p=0.073
Relative Log2 Abundance
Control
AD
ALS
FTLDU
PSP
CBD
PD.PDD
MSA
otherND
Braak Score
CERAD Score

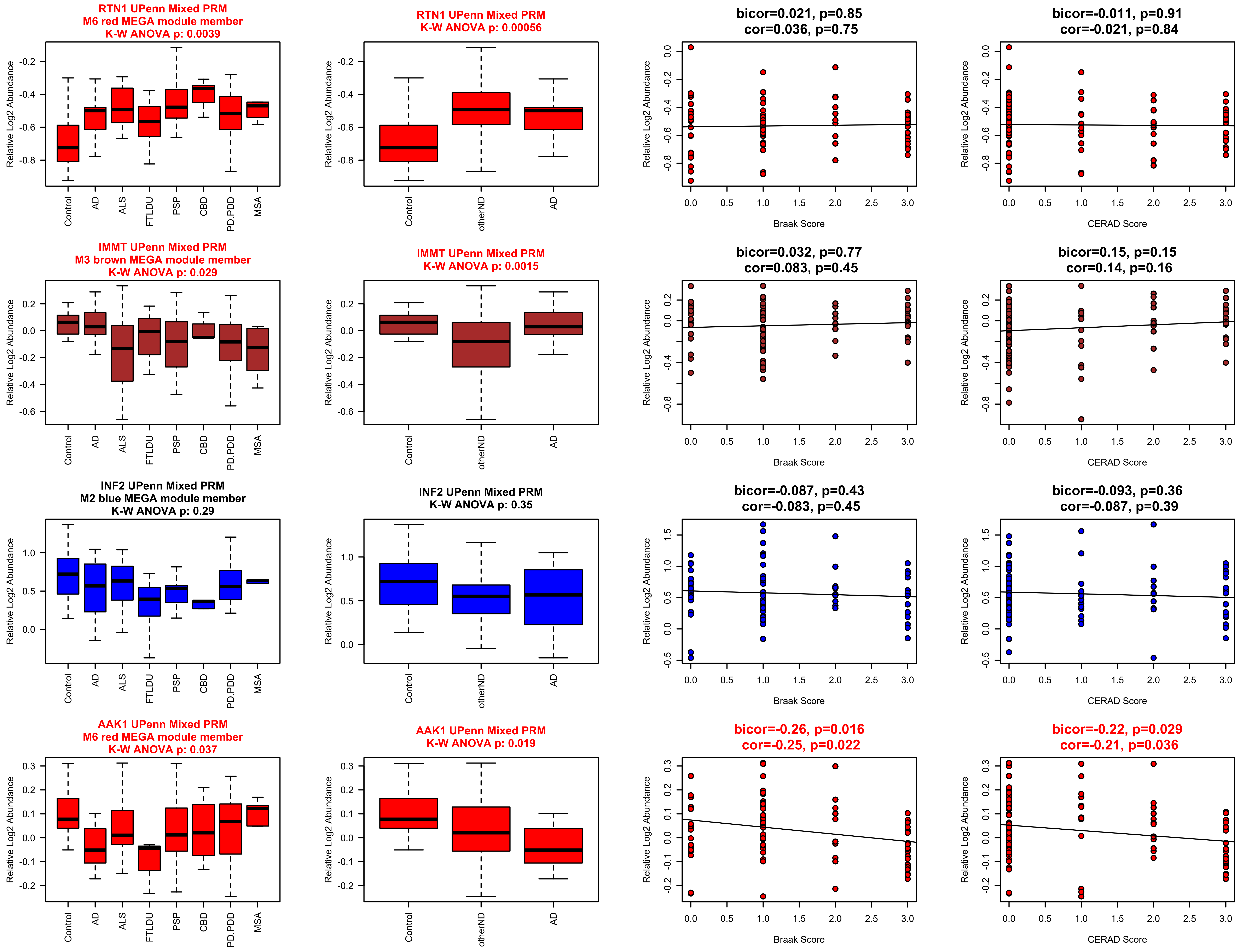

RTN1 UPenn Mixed PRM
M6 red MEGA module member
K-W ANOVA p: 0.0039
RTN1 UPenn Mixed PRM
K-W ANOVA p: 0.00056
bicor=0.021, p=0.85
cor=0.036, p=0.75
bicor=-0.011, p=0.91
cor=-0.021, p=0.84
IMMT UPenn Mixed PRM
M3 brown MEGA module member
K-W ANOVA p: 0.029
IMMT UPenn Mixed PRM
K-W ANOVA p: 0.0015
bicor=0.032, p=0.77
cor=0.083, p=0.45
bicor=0.15, p=0.15
cor=0.14, p=0.16
INF2 UPenn Mixed PRM
M2 blue MEGA module member
K-W ANOVA p: 0.29
INF2 UPenn Mixed PRM
K-W ANOVA p: 0.35
bicor=-0.087, p=0.43
cor=-0.083, p=0.45
bicor=-0.093, p=0.36
cor=-0.087, p=0.39
AAK1 UPenn Mixed PRM
M6 red MEGA module member
K-W ANOVA p: 0.037
AAK1 UPenn Mixed PRM
K-W ANOVA p: 0.019
bicor=-0.26, p=0.016
cor=-0.25, p=0.022
bicor=-0.22, p=0.029
cor=-0.21, p=0.036
Relative Log2 Abundance
Control
AD
ALS
FTLDU
PSP
CBD
PD.PDD
MSA
otherND
Braak Score
CERAD Score

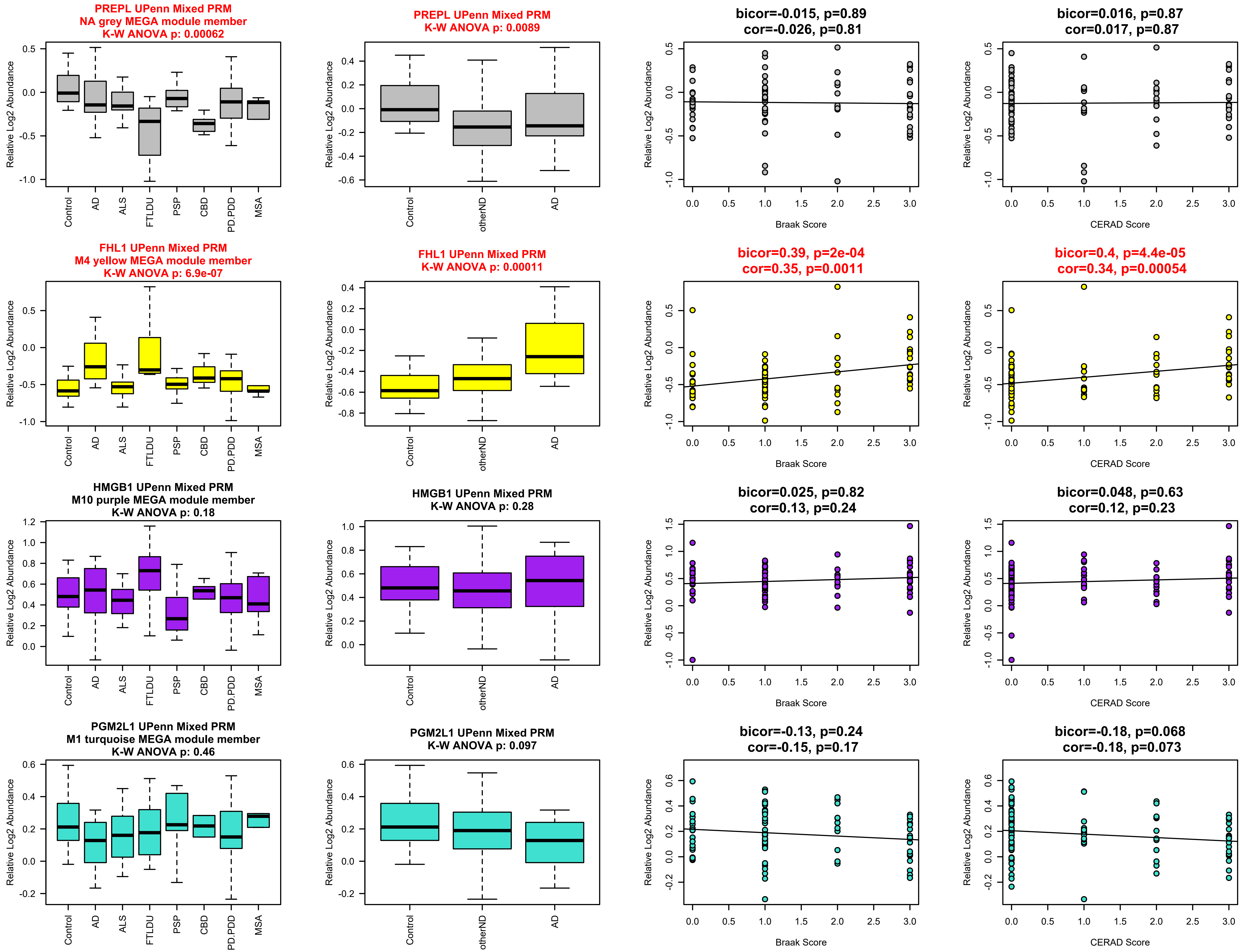
PREPL UPenn Mixed PRM
NA grey MEGA module member
K-W ANOVA p: 0.00062
PREPL UPenn Mixed PRM
K-W ANOVA p: 0.0089
bicor=-0.015, p=0.89
cor=-0.026, p=0.81
bicor=0.016, p=0.87
cor=0.017, p=0.87
FHL1 UPenn Mixed PRM
M4 yellow MEGA module member
K-W ANOVA p: 6.9e-07
FHL1 UPenn Mixed PRM
K-W ANOVA p: 0.00011
bicor=0.39, p=2e-04
cor=0.35, p=0.0011
bicor=0.4, p=4.4e-05
cor=0.34, p=0.00054
HMGB1 UPenn Mixed PRM
M10 purple MEGA module member
K-W ANOVA p: 0.18
HMGB1 UPenn Mixed PRM
K-W ANOVA p: 0.28
bicor=0.025, p=0.82
cor=0.13, p=0.24
bicor=0.048, p=0.63
cor=0.12, p=0.23
PGM2L1 UPenn Mixed PRM
M1 turquoise MEGA module member
K-W ANOVA p: 0.46
PGM2L1 UPenn Mixed PRM
K-W ANOVA p: 0.097
bicor=-0.13, p=0.24
cor=-0.15, p=0.17
bicor=-0.18, p=0.068
cor=-0.18, p=0.073
Relative Log2 Abundance
Control
AD
ALS
FTLDU
PSP
CBD
PD.PDD
MSA
otherND
Braak Score
CERAD Score

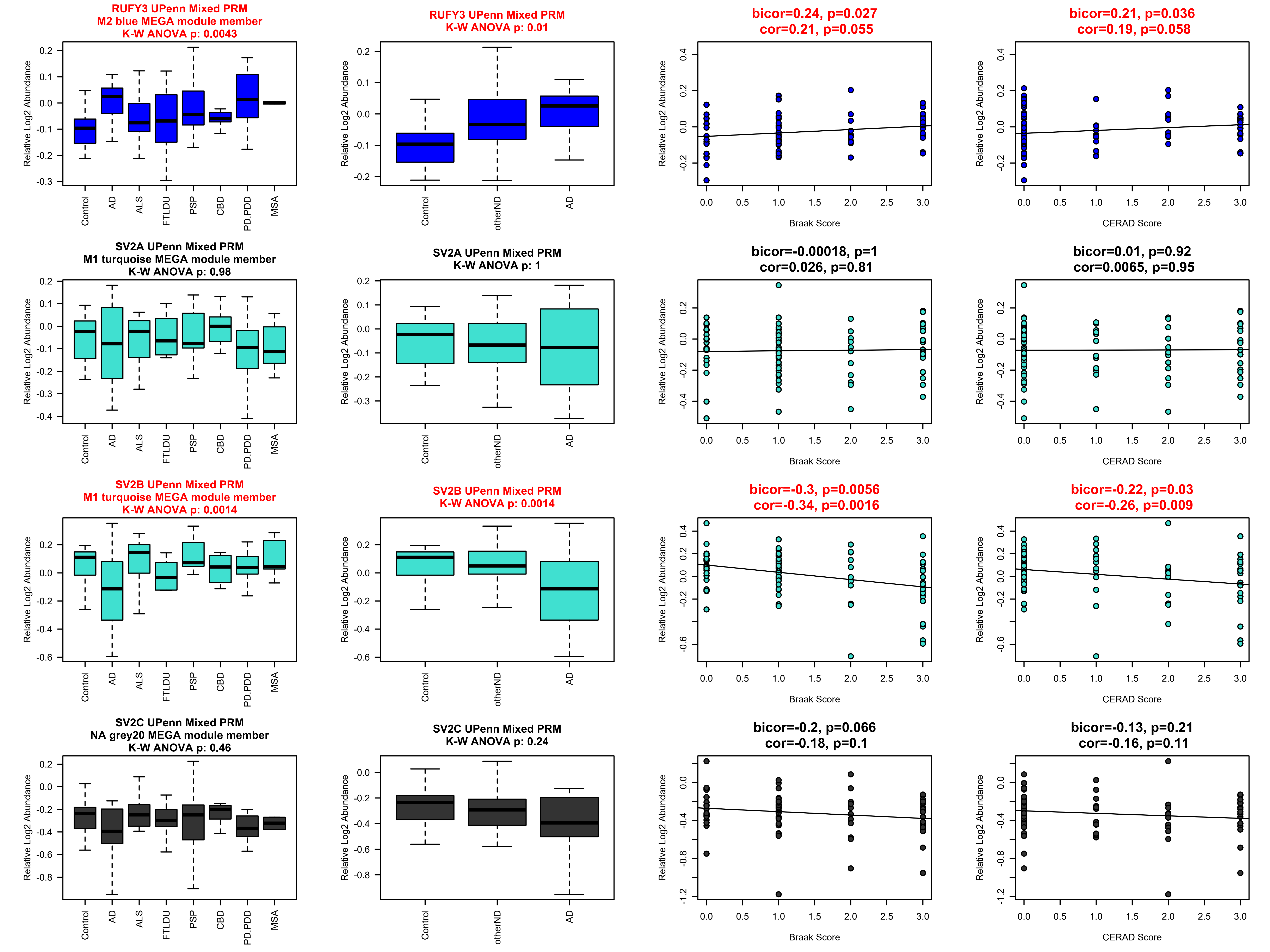

RUFY3 UPenn Mixed PRM
M2 blue MEGA module member
K-W ANOVA p: 0.0043
RUFY3 UPenn Mixed PRM
K-W ANOVA p: 0.01
bicor=0.24, p=0.027
cor=0.21, p=0.055
bicor=0.21, p=0.036
cor=0.19, p=0.058
SV2A UPenn Mixed PRM
M1 turquoise MEGA module member
K-W ANOVA p: 0.98
SV2A UPenn Mixed PRM
K-W ANOVA p: 1
bicor=-0.00018, p=1
cor=0.026, p=0.81
bicor=0.01, p=0.92
cor=0.0065, p=0.95
SV2B UPenn Mixed PRM
M1 turquoise MEGA module member
K-W ANOVA p: 0.0014
SV2B UPenn Mixed PRM
K-W ANOVA p: 0.0014
bicor=-0.3, p=0.0056
cor=-0.34, p=0.0016
bicor=-0.22, p=0.03
cor=-0.26, p=0.009
SV2C UPenn Mixed PRM
NA grey20 MEGA module member
K-W ANOVA p: 0.46
SV2C UPenn Mixed PRM
K-W ANOVA p: 0.24
bicor=-0.2, p=0.066
cor=-0.18, p=0.1
bicor=-0.13, p=0.21
cor=-0.16, p=0.11
Relative Log2 Abundance
Control
AD
ALS
FTLDU
PSP
CBD
PD.PDD
MSA
otherND
Braak Score
CERAD Score

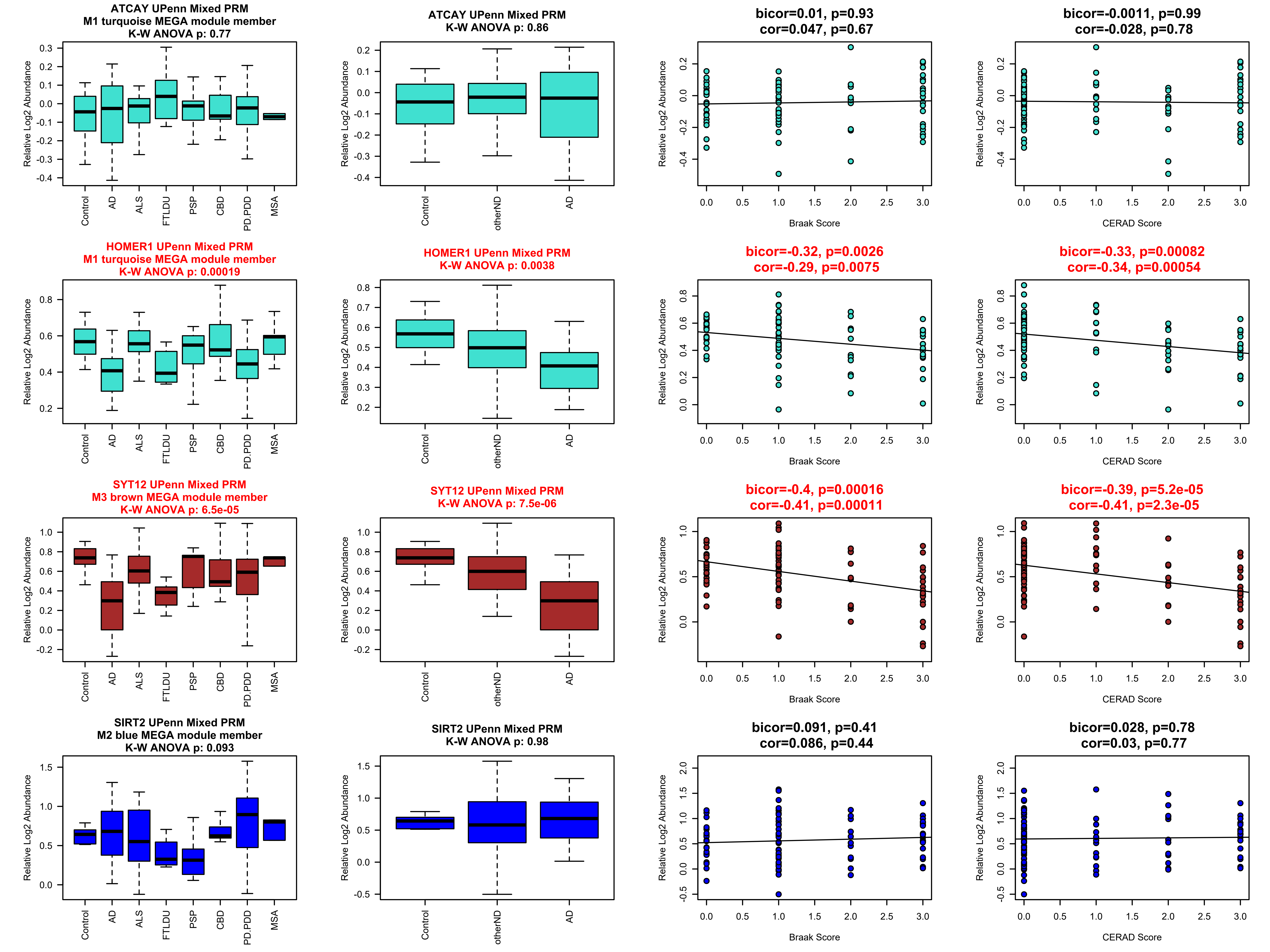
ATCAY UPenn Mixed PRM
M1 turquoise MEGA module member
K-W ANOVA p: 0.77
ATCAY UPenn Mixed PRM
K-W ANOVA p: 0.86
bicor=0.01, p=0.93
cor=0.047, p=0.67
bicor=-0.0011, p=0.99
cor=-0.028, p=0.78
HOMER1 UPenn Mixed PRM
M1 turquoise MEGA module member
K-W ANOVA p: 0.00019
HOMER1 UPenn Mixed PRM
K-W ANOVA p: 0.0038
bicor=-0.32, p=0.0026
cor=-0.29, p=0.0075
bicor=-0.33, p=0.00082
cor=-0.34, p=0.00054
SYT12 UPenn Mixed PRM
M3 brown MEGA module member
K-W ANOVA p: 6.5e-05
SYT12 UPenn Mixed PRM
K-W ANOVA p: 7.5e-06
bicor=-0.4, p=0.00016
cor=-0.41, p=0.00011
bicor=-0.39, p=5.2e-05
cor=-0.41, p=2.3e-05
SIRT2 UPenn Mixed PRM
M2 blue MEGA module member
K-W ANOVA p: 0.093
SIRT2 UPenn Mixed PRM
K-W ANOVA p: 0.98
bicor=0.091, p=0.41
cor=0.086, p=0.44
bicor=0.028, p=0.78
cor=0.03, p=0.77
Relative Log2 Abundance
Control
AD
ALS
FTLDU
PSP
CBD
PD.PDD
MSA
otherND
Braak Score
CERAD Score

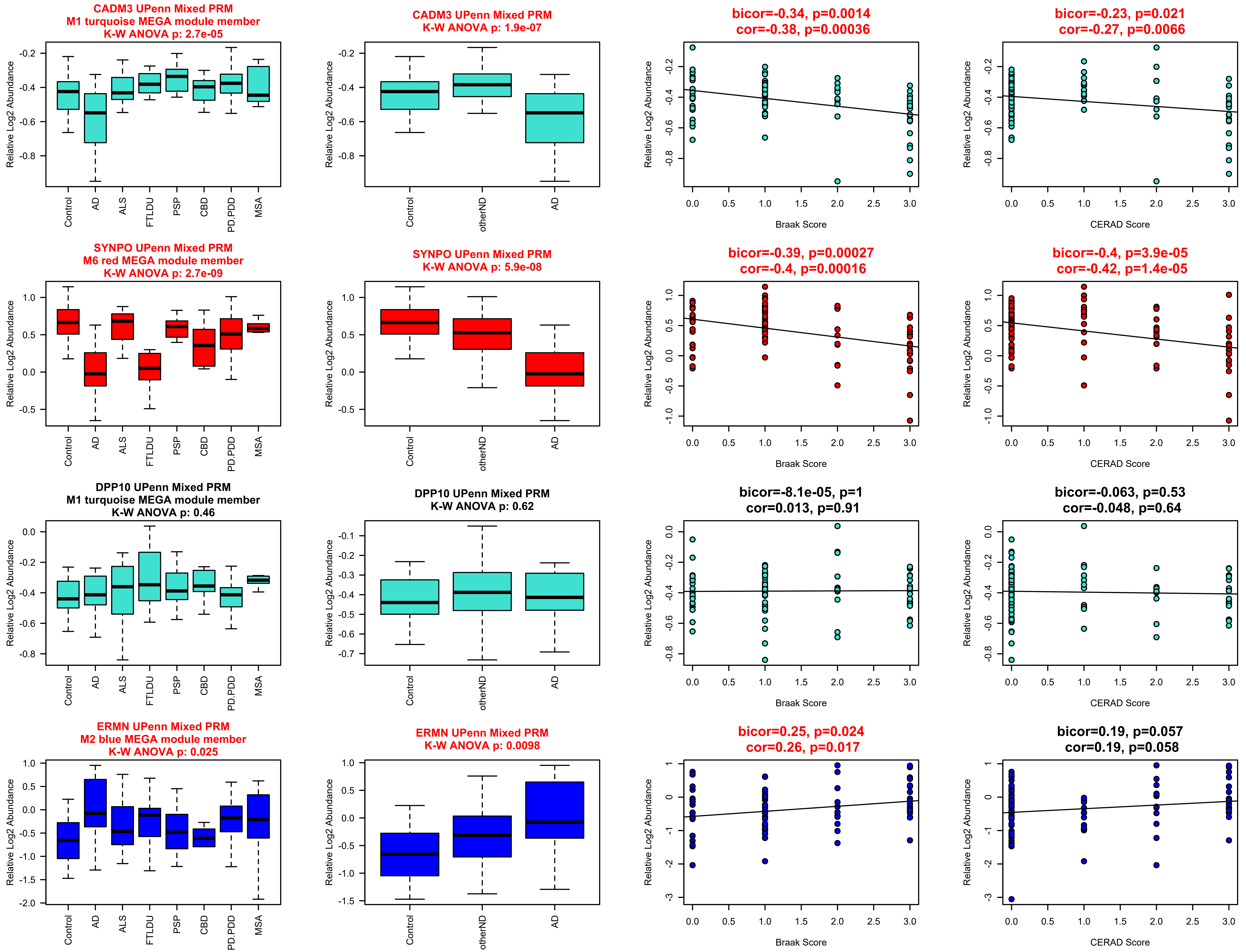

CADM3 UPenn Mixed PRM
M1 turquoise MEGA module member
K-W ANOVA p: 2.7e-05
CADM3 UPenn Mixed PRM
K-W ANOVA p: 1.9e-07
bicor=-0.34, p=0.0014
cor=-0.38, p=0.00036
bicor=-0.23, p=0.021
cor=-0.27, p=0.0066
SYNPO UPenn Mixed PRM
M6 red MEGA module member
K-W ANOVA p: 2.7e-09
SYNPO UPenn Mixed PRM
K-W ANOVA p: 5.9e-08
bicor=-0.39, p=0.00027
cor=-0.4, p=0.00016
bicor=-0.4, p=3.9e-05
cor=-0.42, p=1.4e-05
DPP10 UPenn Mixed PRM
M1 turquoise MEGA module member
K-W ANOVA p: 0.46
DPP10 UPenn Mixed PRM
K-W ANOVA p: 0.62
bicor=-8.1e-05, p=1
cor=0.013, p=0.91
bicor=-0.063, p=0.53
cor=-0.048, p=0.64
ERMN UPenn Mixed PRM
M2 blue MEGA module member
K-W ANOVA p: 0.025
ERMN UPenn Mixed PRM
K-W ANOVA p: 0.0098
bicor=0.25, p=0.024
cor=0.26, p=0.017
bicor=0.19, p=0.057
cor=0.19, p=0.058
Relative Log2 Abundance
Control
AD
ALS
FTLDU
PSP
CBD
PD.PDD
MSA
otherND
Braak Score
CERAD Score

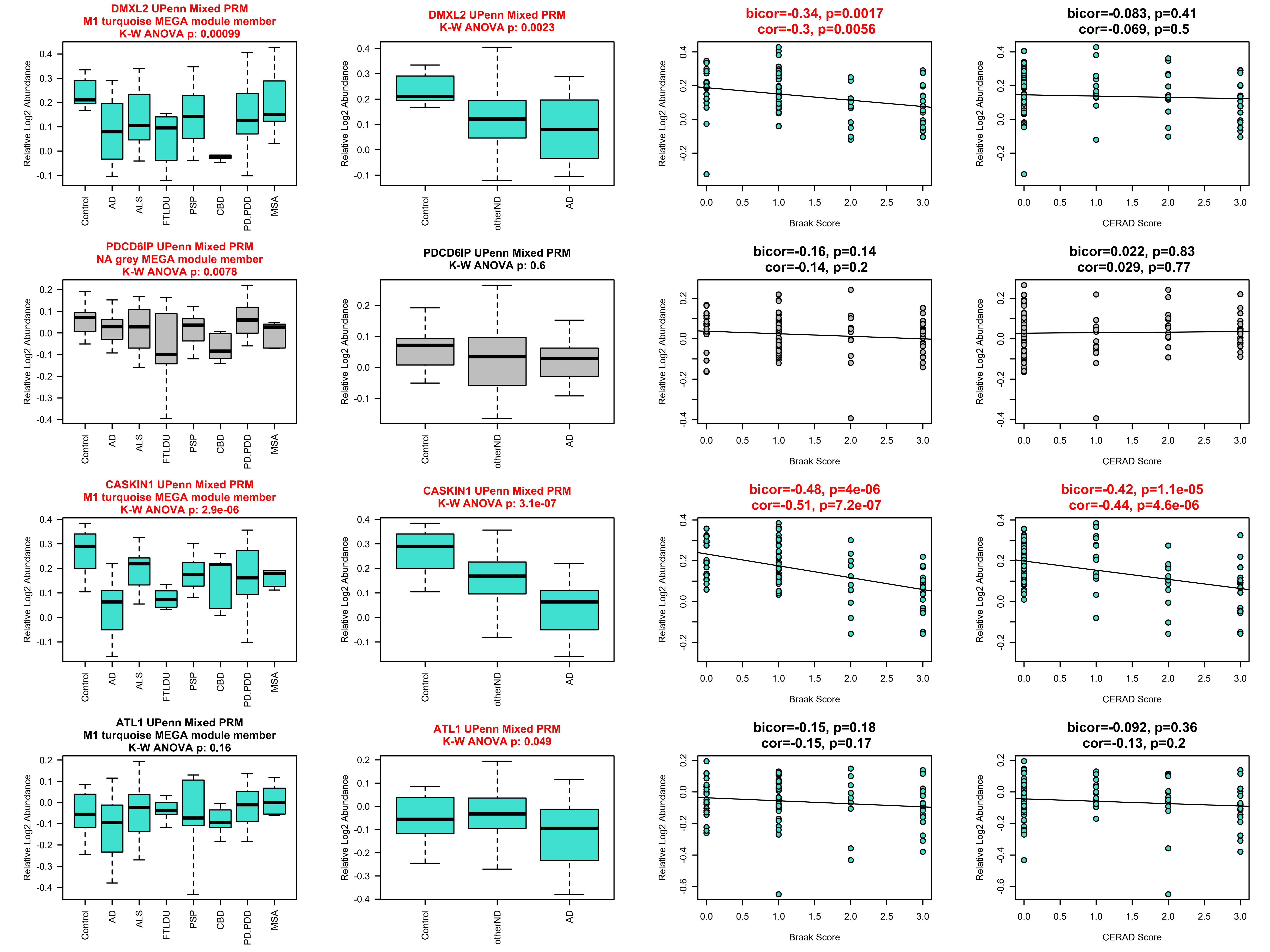
DMXL2 UPenn Mixed PRM
M1 turquoise MEGA module member
K-W ANOVA p: 0.00099
Relative Log2 Abundance
Control
AD
ALS
FTLDU
PSP
CBD
PD.PDD
MSA
DMXL2 UPenn Mixed PRM
K-W ANOVA p: 0.0023
Relative Log2 Abundance
Control
otherND
AD
bicor=-0.34, p=0.0017
cor=-0.3, p=0.0056
Relative Log2 Abundance
Braak Score
bicor=-0.083, p=0.41
cor=-0.069, p=0.5
Relative Log2 Abundance
CERAD Score
PDCD6IP UPenn Mixed PRM
NA grey MEGA module member
K-W ANOVA p: 0.0078
Relative Log2 Abundance
Control
AD
ALS
FTLDU
PSP
CBD
PD.PDD
MSA
PDCD6IP UPenn Mixed PRM
K-W ANOVA p: 0.6
Relative Log2 Abundance
Control
otherND
AD
bicor=-0.16, p=0.14
cor=-0.14, p=0.2
Relative Log2 Abundance
Braak Score
bicor=0.022, p=0.83
cor=0.029, p=0.77
Relative Log2 Abundance
CERAD Score
CASKIN1 UPenn Mixed PRM
M1 turquoise MEGA module member
K-W ANOVA p: 2.9e-06
Relative Log2 Abundance
Control
AD
ALS
FTLDU
PSP
CBD
PD.PDD
MSA
CASKIN1 UPenn Mixed PRM
K-W ANOVA p: 3.1e-07
Relative Log2 Abundance
Control
otherND
AD
bicor=-0.48, p=4e-06
cor=-0.51, p=7.2e-07
Relative Log2 Abundance
Braak Score
bicor=-0.42, p=1.1e-05
cor=-0.44, p=4.6e-06
Relative Log2 Abundance
CERAD Score
ATL1 UPenn Mixed PRM
M1 turquoise MEGA module member
K-W ANOVA p: 0.16
Relative Log2 Abundance
Control
AD
ALS
FTLDU
PSP
CBD
PD.PDD
MSA
ATL1 UPenn Mixed PRM
K-W ANOVA p: 0.049
Relative Log2 Abundance
Control
otherND
AD
bicor=-0.15, p=0.18
cor=-0.15, p=0.17
Relative Log2 Abundance
Braak Score
bicor=-0.092, p=0.36
cor=-0.13, p=0.2
Relative Log2 Abundance
CERAD Score

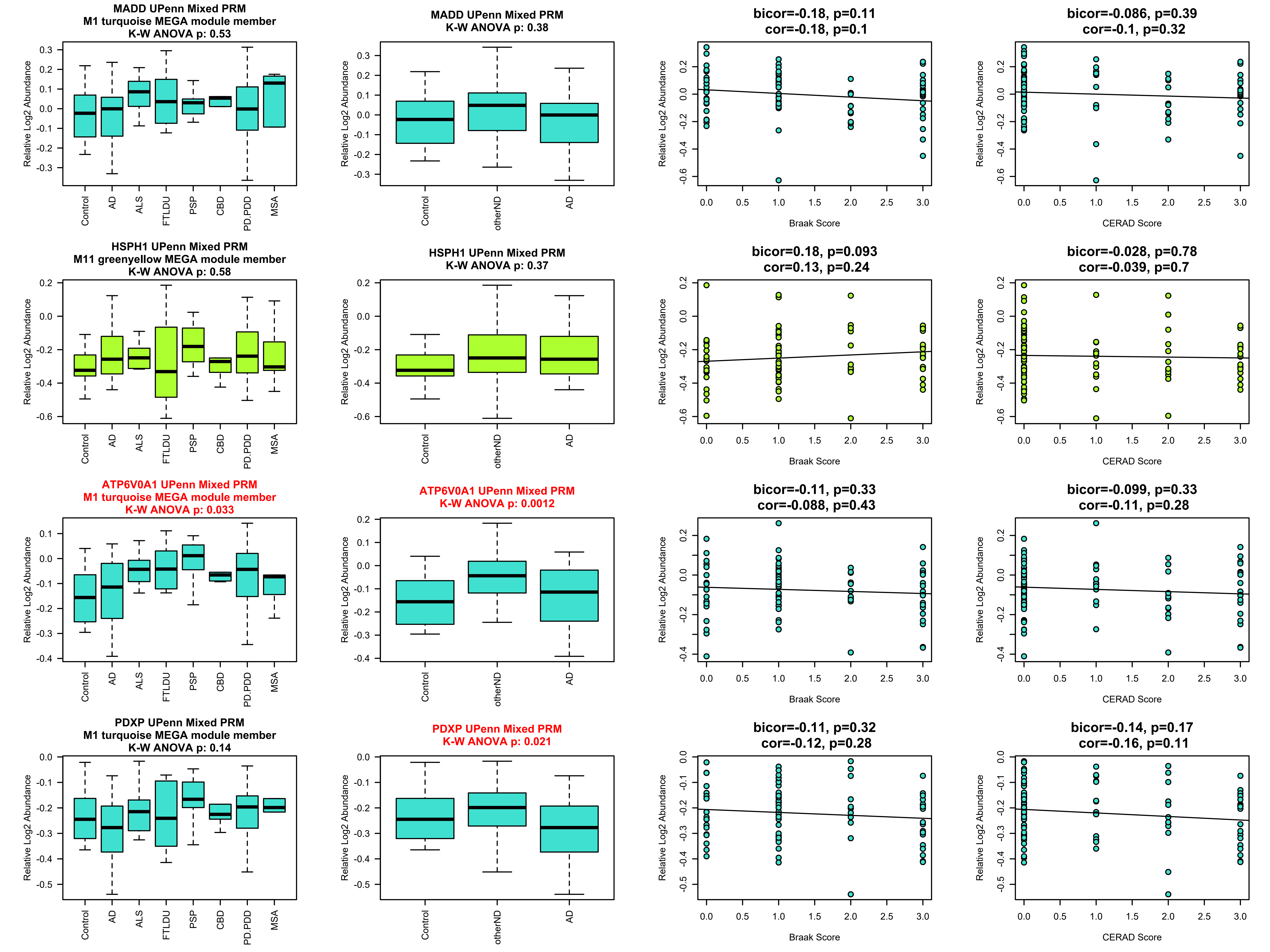

MADD UPenn Mixed PRM
M1 turquoise MEGA module member
K-W ANOVA p: 0.53
MADD UPenn Mixed PRM
K-W ANOVA p: 0.38
bicor=-0.18, p=0.11
cor=-0.18, p=0.1
bicor=-0.086, p=0.39
cor=-0.1, p=0.32
HSPH1 UPenn Mixed PRM
M11 greenyellow MEGA module member
K-W ANOVA p: 0.58
HSPH1 UPenn Mixed PRM
K-W ANOVA p: 0.37
bicor=0.18, p=0.093
cor=0.13, p=0.24
bicor=-0.028, p=0.78
cor=-0.039, p=0.7
ATP6V0A1 UPenn Mixed PRM
M1 turquoise MEGA module member
K-W ANOVA p: 0.033
ATP6V0A1 UPenn Mixed PRM
K-W ANOVA p: 0.0012
bicor=-0.11, p=0.33
cor=-0.088, p=0.43
bicor=-0.099, p=0.33
cor=-0.11, p=0.28
PDXP UPenn Mixed PRM
M1 turquoise MEGA module member
K-W ANOVA p: 0.14
PDXP UPenn Mixed PRM
K-W ANOVA p: 0.021
bicor=-0.11, p=0.32
cor=-0.12, p=0.28
bicor=-0.14, p=0.17
cor=-0.16, p=0.11
Relative Log2 Abundance
Control
AD
ALS
FTLDU
PSP
CBD
PD.PDD
MSA
otherND
Braak Score
CERAD Score

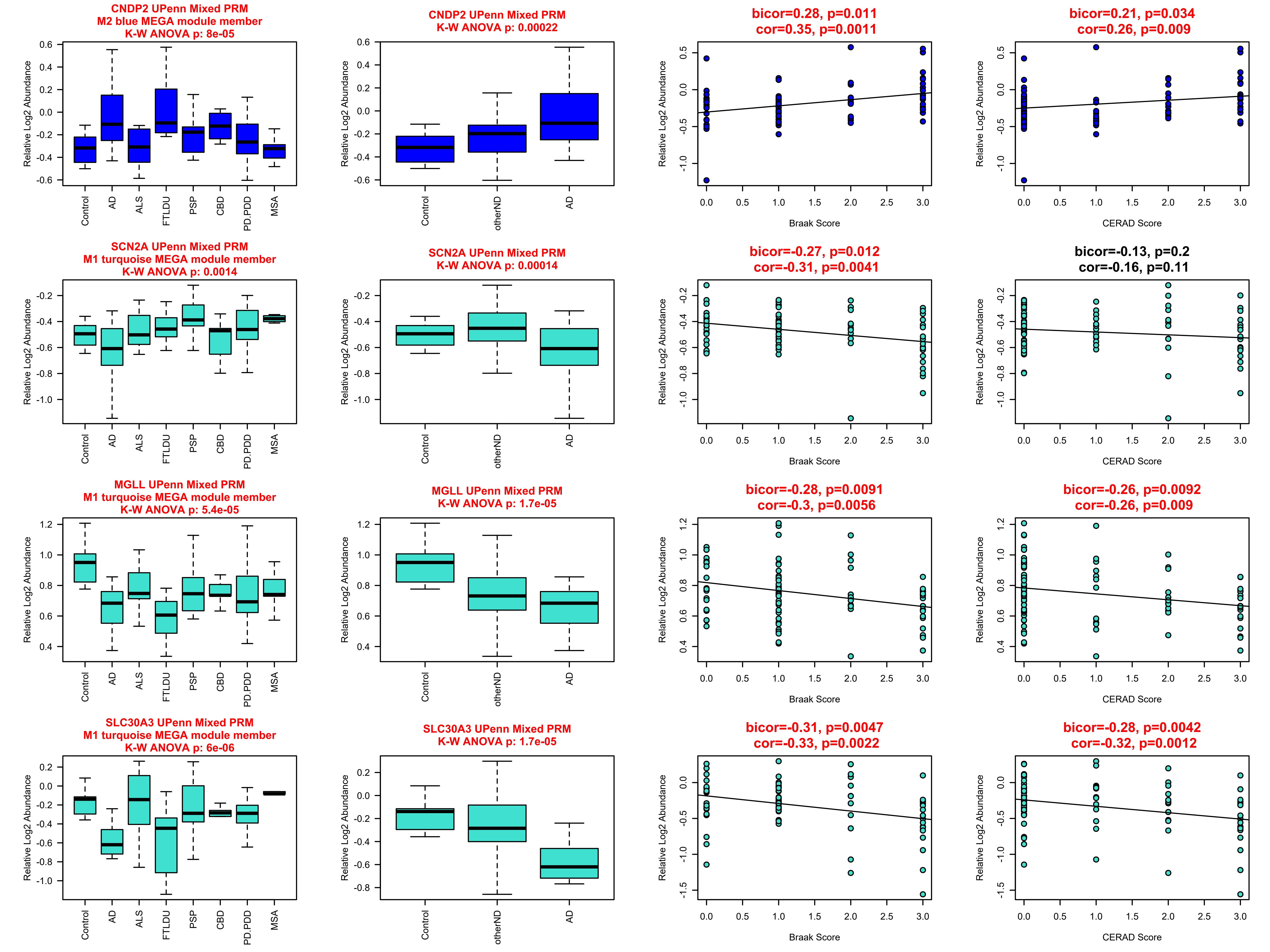
CNDP2 UPenn Mixed PRM
M2 blue MEGA module member
K-W ANOVA p: 8e-05
CNDP2 UPenn Mixed PRM
K-W ANOVA p: 0.00022
bicor=0.28, p=0.011
cor=0.35, p=0.0011
bicor=0.21, p=0.034
cor=0.26, p=0.009
SCN2A UPenn Mixed PRM
M1 turquoise MEGA module member
K-W ANOVA p: 0.0014
SCN2A UPenn Mixed PRM
K-W ANOVA p: 0.00014
bicor=-0.27, p=0.012
cor=-0.31, p=0.0041
bicor=-0.13, p=0.2
cor=-0.16, p=0.11
MGLL UPenn Mixed PRM
M1 turquoise MEGA module member
K-W ANOVA p: 5.4e-05
MGLL UPenn Mixed PRM
K-W ANOVA p: 1.7e-05
bicor=-0.28, p=0.0091
cor=-0.3, p=0.0056
bicor=-0.26, p=0.0092
cor=-0.26, p=0.009
SLC30A3 UPenn Mixed PRM
M1 turquoise MEGA module member
K-W ANOVA p: 6e-06
SLC30A3 UPenn Mixed PRM
K-W ANOVA p: 1.7e-05
bicor=-0.31, p=0.0047
cor=-0.33, p=0.0022
bicor=-0.28, p=0.0042
cor=-0.32, p=0.0012
Relative Log2 Abundance
Control
AD
ALS
FTLDU
PSP
CBD
PD.PDD
MSA
otherND
Braak Score
CERAD Score

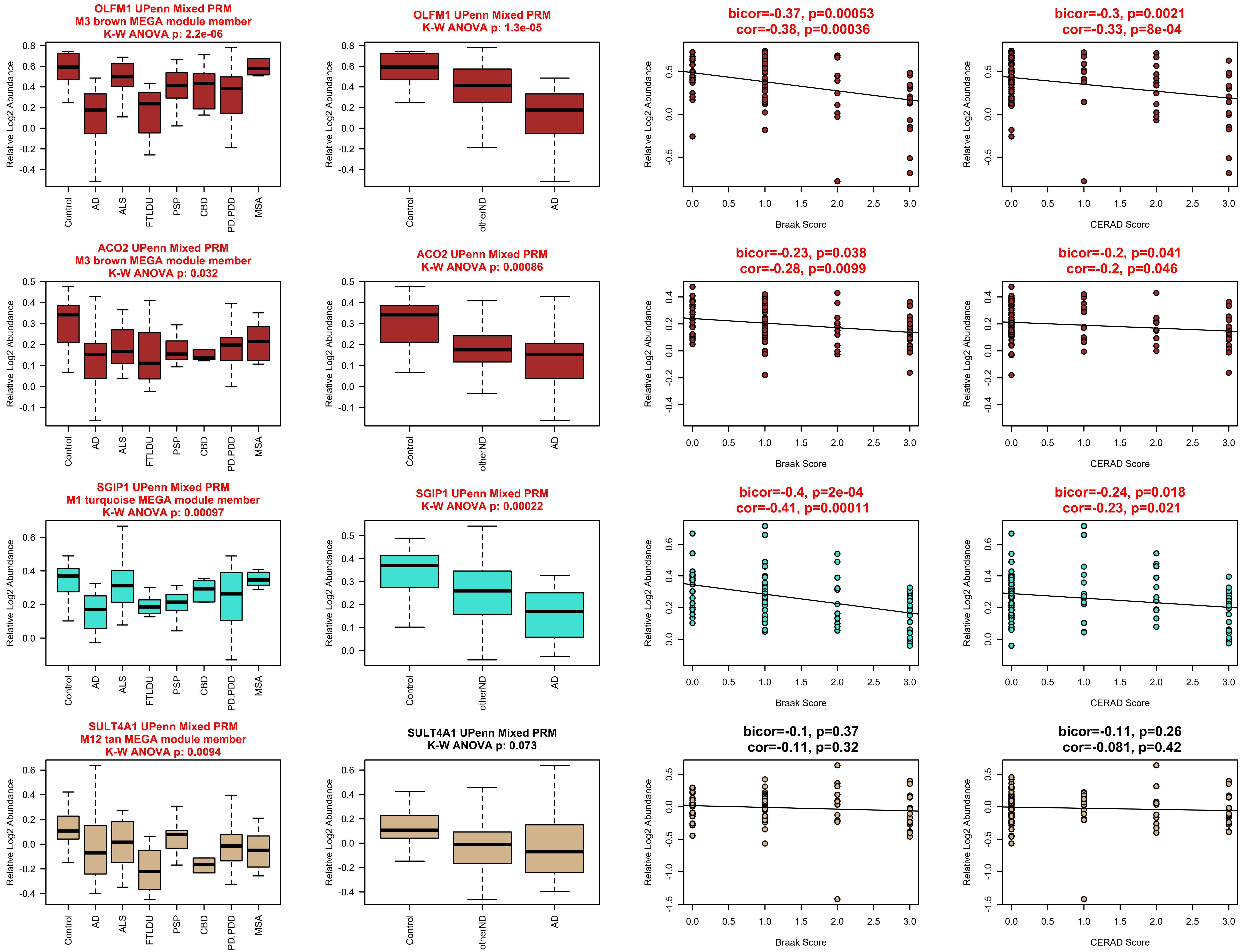
OLFM1 UPenn Mixed PRM
M3 brown MEGA module member
K-W ANOVA p: 2.2e-06
OLFM1 UPenn Mixed PRM
K-W ANOVA p: 1.3e-05
bicor=-0.37, p=0.00053
cor=-0.38, p=0.00036
bicor=-0.3, p=0.0021
cor=-0.33, p=8e-04
ACO2 UPenn Mixed PRM
M3 brown MEGA module member
K-W ANOVA p: 0.032
ACO2 UPenn Mixed PRM
K-W ANOVA p: 0.00086
bicor=-0.23, p=0.038
cor=-0.28, p=0.0099
bicor=-0.2, p=0.041
cor=-0.2, p=0.046
SGIP1 UPenn Mixed PRM
M1 turquoise MEGA module member
K-W ANOVA p: 0.00097
SGIP1 UPenn Mixed PRM
K-W ANOVA p: 0.00022
bicor=-0.4, p=2e-04
cor=-0.41, p=0.00011
bicor=-0.24, p=0.018
cor=-0.23, p=0.021
SULT4A1 UPenn Mixed PRM
M12 tan MEGA module member
K-W ANOVA p: 0.0094
SULT4A1 UPenn Mixed PRM
K-W ANOVA p: 0.073
bicor=-0.1, p=0.37
cor=-0.11, p=0.32
bicor=-0.11, p=0.26
cor=-0.081, p=0.42
Relative Log2 Abundance
Control
AD
ALS
FTLDU
PSP
CBD
PD.PDD
MSA
otherND
Braak Score
CERAD Score

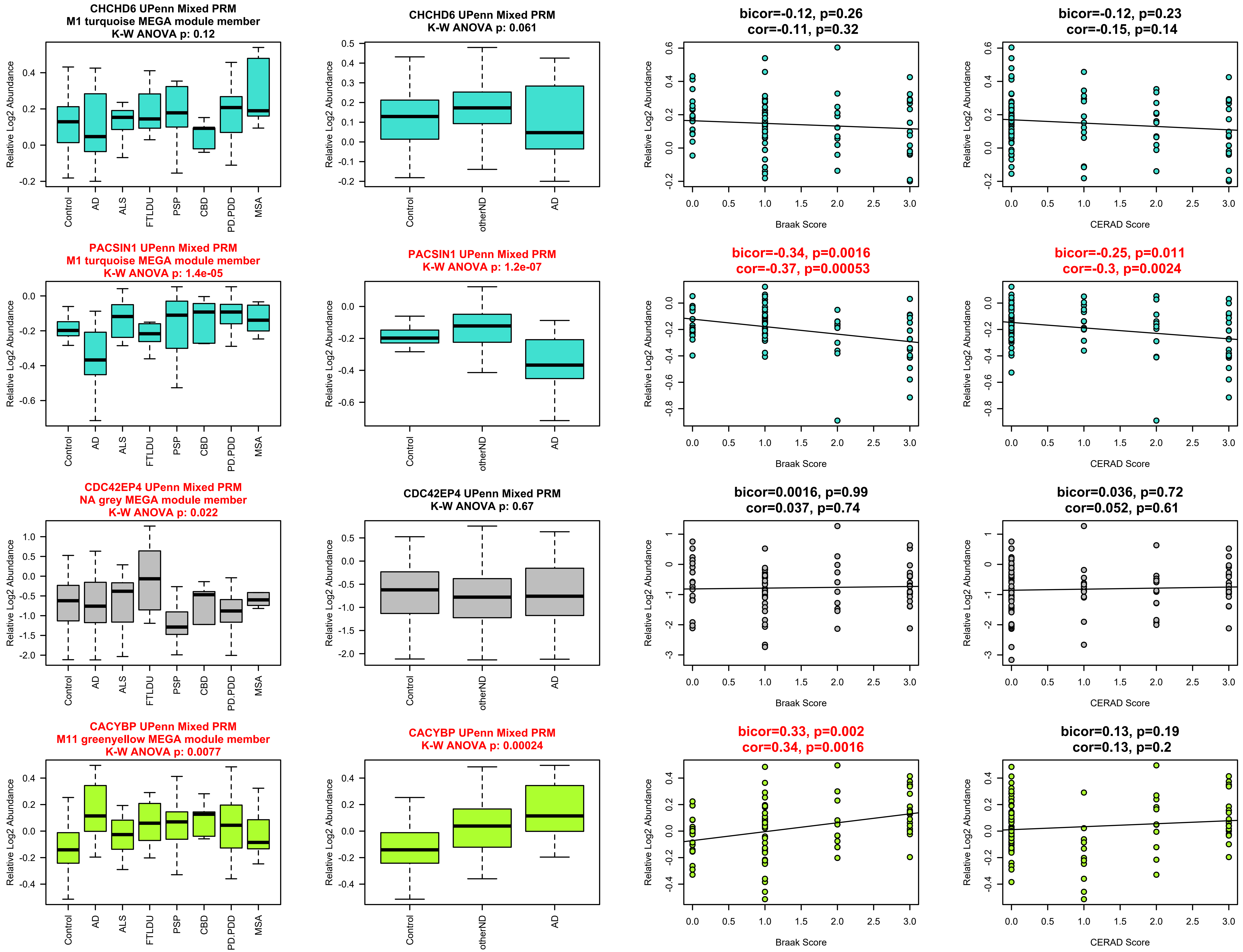
CHCHD6 UPenn Mixed PRM
M1 turquoise MEGA module member
K-W ANOVA p: 0.12
CHCHD6 UPenn Mixed PRM
K-W ANOVA p: 0.061
bicor=-0.12, p=0.26
cor=-0.11, p=0.32
bicor=-0.12, p=0.23
cor=-0.15, p=0.14
PACSIN1 UPenn Mixed PRM
M1 turquoise MEGA module member
K-W ANOVA p: 1.4e-05
PACSIN1 UPenn Mixed PRM
K-W ANOVA p: 1.2e-07
bicor=-0.34, p=0.0016
cor=-0.37, p=0.00053
bicor=-0.25, p=0.011
cor=-0.3, p=0.0024
CDC42EP4 UPenn Mixed PRM
NA grey MEGA module member
K-W ANOVA p: 0.022
CDC42EP4 UPenn Mixed PRM
K-W ANOVA p: 0.67
bicor=0.0016, p=0.99
cor=0.037, p=0.74
bicor=0.036, p=0.72
cor=0.052, p=0.61
CACYBP UPenn Mixed PRM
M11 greenyellow MEGA module member
K-W ANOVA p: 0.0077
CACYBP UPenn Mixed PRM
K-W ANOVA p: 0.00024
bicor=0.33, p=0.002
cor=0.34, p=0.0016
bicor=0.13, p=0.19
cor=0.13, p=0.2
Relative Log2 Abundance
Control
AD
ALS
FTLDU
PSP
CBD
PD.PDD
MSA
otherND
Braak Score
CERAD Score

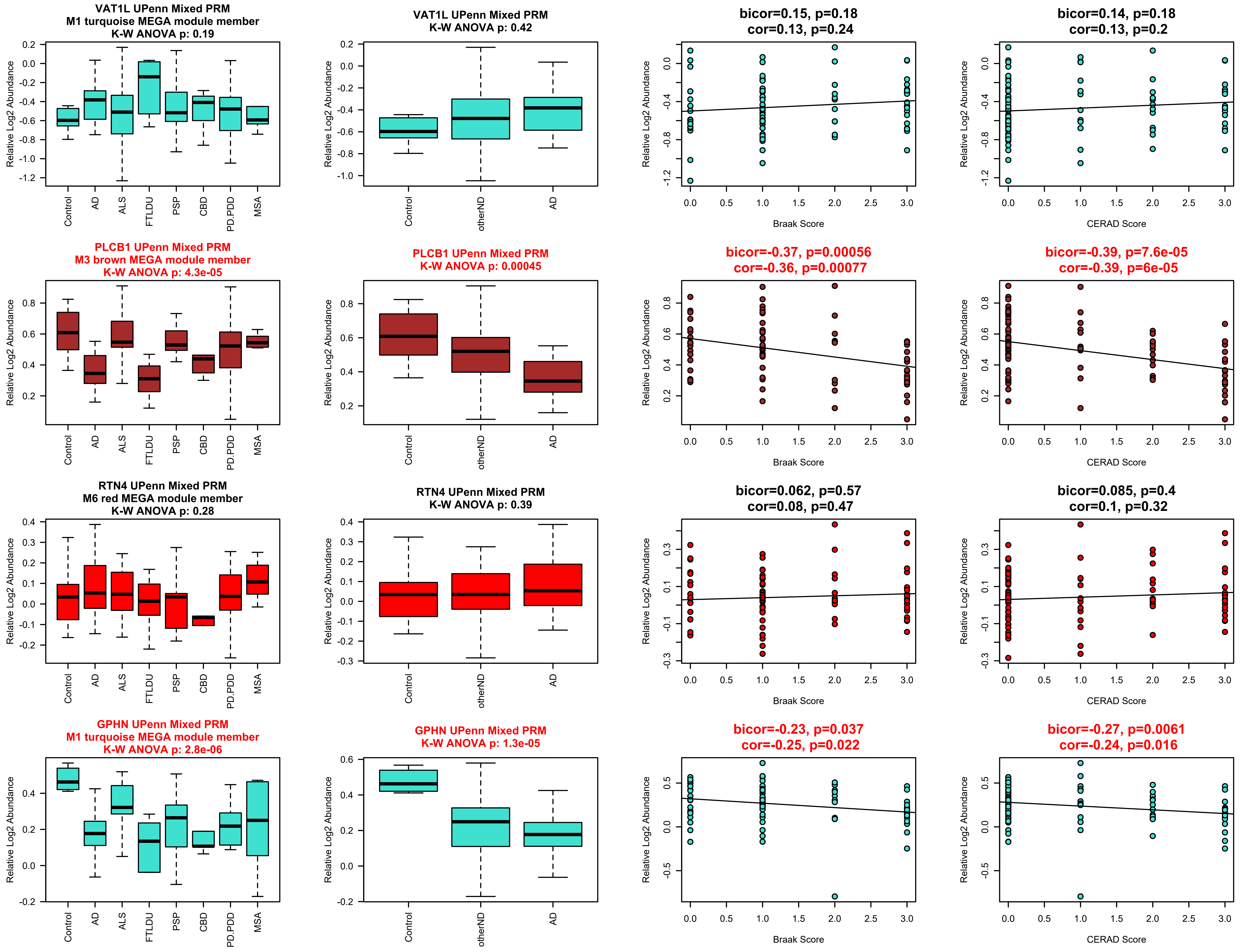
VAT1L UPenn Mixed PRM
M1 turquoise MEGA module member
K-W ANOVA p: 0.19
VAT1L UPenn Mixed PRM
K-W ANOVA p: 0.42
bicor=0.15, p=0.18
cor=0.13, p=0.24
bicor=0.14, p=0.18
cor=0.13, p=0.2
PLCB1 UPenn Mixed PRM
M3 brown MEGA module member
K-W ANOVA p: 4.3e-05
PLCB1 UPenn Mixed PRM
K-W ANOVA p: 0.00045
bicor=-0.37, p=0.00056
cor=-0.36, p=0.00077
bicor=-0.39, p=7.6e-05
cor=-0.39, p=6e-05
RTN4 UPenn Mixed PRM
M6 red MEGA module member
K-W ANOVA p: 0.28
RTN4 UPenn Mixed PRM
K-W ANOVA p: 0.39
bicor=0.062, p=0.57
cor=0.08, p=0.47
bicor=0.085, p=0.4
cor=0.1, p=0.32
GPHN UPenn Mixed PRM
M1 turquoise MEGA module member
K-W ANOVA p: 2.8e-06
GPHN UPenn Mixed PRM
K-W ANOVA p: 1.3e-05
bicor=-0.23, p=0.037
cor=-0.25, p=0.022
bicor=-0.27, p=0.0061
cor=-0.24, p=0.016
Relative Log2 Abundance
Control
AD
ALS
FTLDU
PSP
CBD
PD.PDD
MSA
otherND
Braak Score
CERAD Score

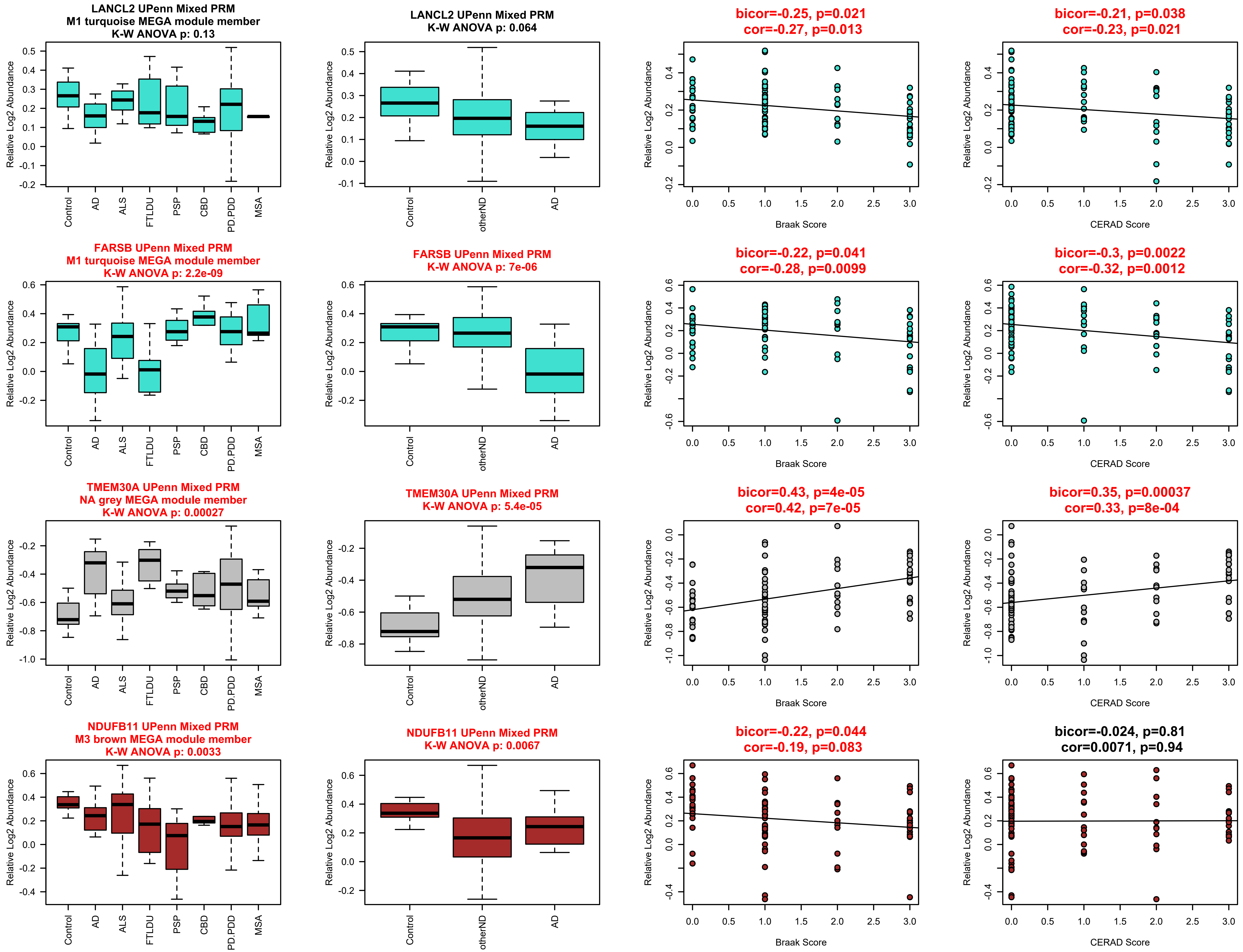
LANCL2 UPenn Mixed PRM
M1 turquoise MEGA module member
K-W ANOVA p: 0.13
LANCL2 UPenn Mixed PRM
K-W ANOVA p: 0.064
bicor=-0.25, p=0.021
cor=-0.27, p=0.013
bicor=-0.21, p=0.038
cor=-0.23, p=0.021
FARSB UPenn Mixed PRM
M1 turquoise MEGA module member
K-W ANOVA p: 2.2e-09
FARSB UPenn Mixed PRM
K-W ANOVA p: 7e-06
bicor=-0.22, p=0.041
cor=-0.28, p=0.0099
bicor=-0.3, p=0.0022
cor=-0.32, p=0.0012
TMEM30A UPenn Mixed PRM
NA grey MEGA module member
K-W ANOVA p: 0.00027
TMEM30A UPenn Mixed PRM
K-W ANOVA p: 5.4e-05
bicor=0.43, p=4e-05
cor=0.42, p=7e-05
bicor=0.35, p=0.00037
cor=0.33, p=8e-04
NDUFB11 UPenn Mixed PRM
M3 brown MEGA module member
K-W ANOVA p: 0.0033
NDUFB11 UPenn Mixed PRM
K-W ANOVA p: 0.0067
bicor=-0.22, p=0.044
cor=-0.19, p=0.083
bicor=-0.024, p=0.81
cor=0.0071, p=0.94
Relative Log2 Abundance
Control
AD
ALS
FTLDU
PSP
CBD
PD.PDD
MSA
otherND
Braak Score
CERAD Score

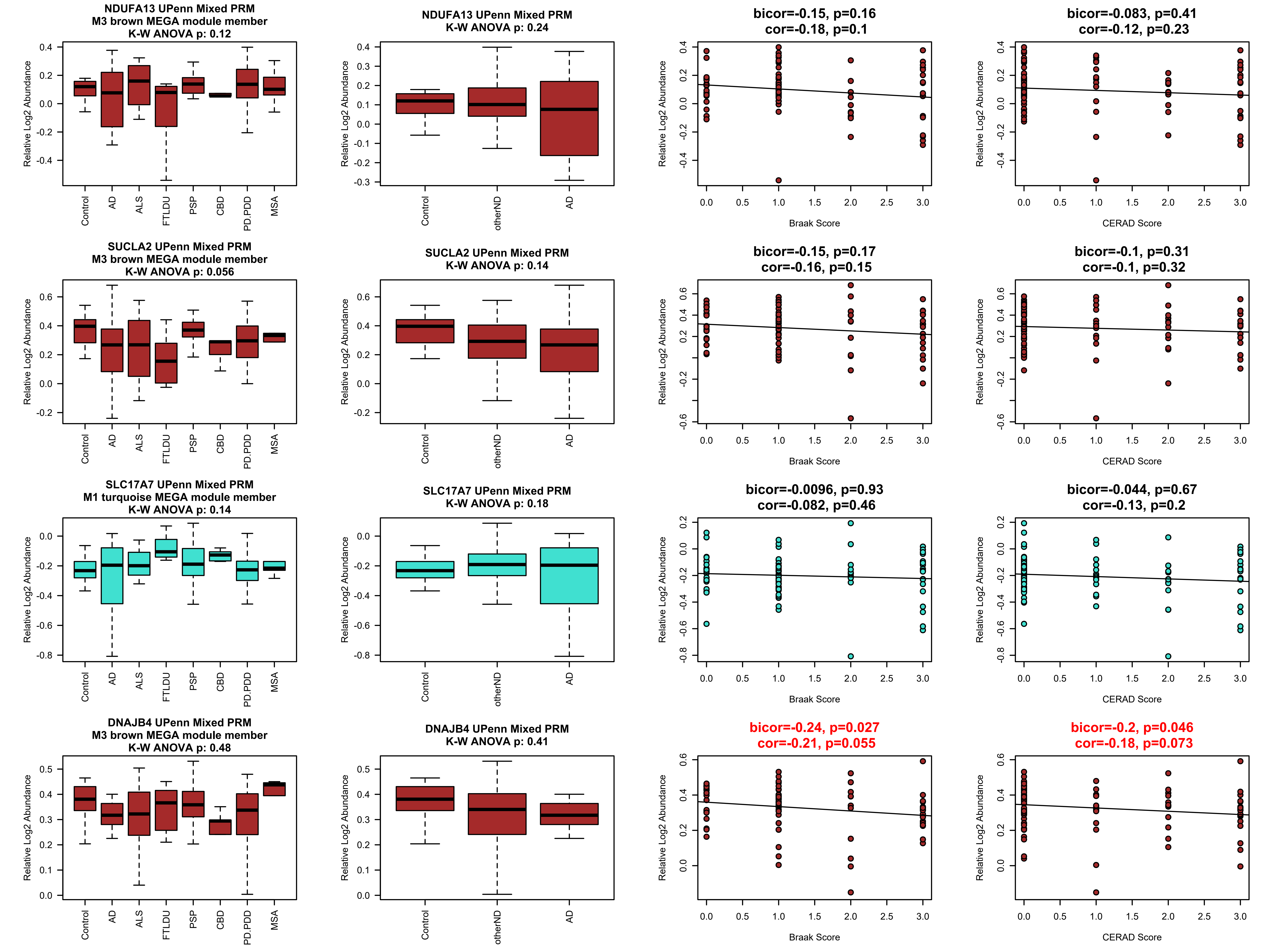
NDUFA13 UPenn Mixed PRM
M3 brown MEGA module member
K-W ANOVA p: 0.12
Relative Log2 Abundance
Control
AD
ALS
FTLDU
PSP
CBD
PD.PDD
MSA
NDUFA13 UPenn Mixed PRM
K-W ANOVA p: 0.24
Relative Log2 Abundance
Control
otherND
AD
bicor=-0.15, p=0.16
cor=-0.18, p=0.1
Relative Log2 Abundance
Braak Score
bicor=-0.083, p=0.41
cor=-0.12, p=0.23
Relative Log2 Abundance
CERAD Score
SUCLA2 UPenn Mixed PRM
M3 brown MEGA module member
K-W ANOVA p: 0.056
Relative Log2 Abundance
Control
AD
ALS
FTLDU
PSP
CBD
PD.PDD
MSA
SUCLA2 UPenn Mixed PRM
K-W ANOVA p: 0.14
Relative Log2 Abundance
Control
otherND
AD
bicor=-0.15, p=0.17
cor=-0.16, p=0.15
Relative Log2 Abundance
Braak Score
bicor=-0.1, p=0.31
cor=-0.1, p=0.32
Relative Log2 Abundance
CERAD Score
SLC17A7 UPenn Mixed PRM
M1 turquoise MEGA module member
K-W ANOVA p: 0.14
Relative Log2 Abundance
Control
AD
ALS
FTLDU
PSP
CBD
PD.PDD
MSA
SLC17A7 UPenn Mixed PRM
K-W ANOVA p: 0.18
Relative Log2 Abundance
Control
otherND
AD
bicor=-0.0096, p=0.93
cor=-0.082, p=0.46
Relative Log2 Abundance
Braak Score
bicor=-0.044, p=0.67
cor=-0.13, p=0.2
Relative Log2 Abundance
CERAD Score
DNAJB4 UPenn Mixed PRM
M3 brown MEGA module member
K-W ANOVA p: 0.48
Relative Log2 Abundance
Control
AD
ALS
FTLDU
PSP
CBD
PD.PDD
MSA
DNAJB4 UPenn Mixed PRM
K-W ANOVA p: 0.41
Relative Log2 Abundance
Control
otherND
AD
bicor=-0.24, p=0.027
cor=-0.21, p=0.055
Relative Log2 Abundance
Braak Score
bicor=-0.2, p=0.046
cor=-0.18, p=0.073
Relative Log2 Abundance
CERAD Score

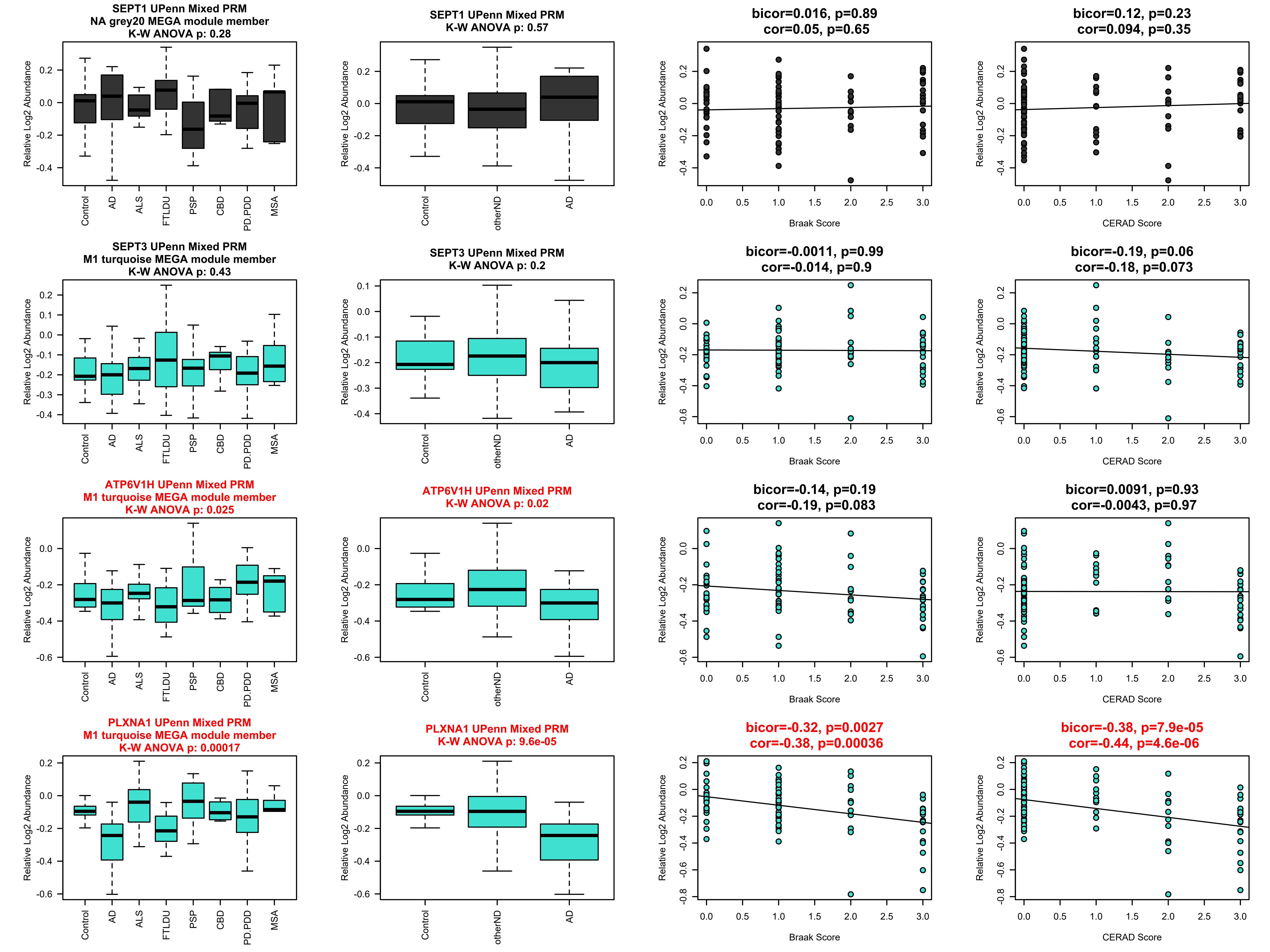

SEPT1 UPenn Mixed PRM
NA grey20 MEGA module member
K-W ANOVA p: 0.28
SEPT1 UPenn Mixed PRM
K-W ANOVA p: 0.57
bicor=0.016, p=0.89
cor=0.05, p=0.65
bicor=0.12, p=0.23
cor=0.094, p=0.35
SEPT3 UPenn Mixed PRM
M1 turquoise MEGA module member
K-W ANOVA p: 0.43
SEPT3 UPenn Mixed PRM
K-W ANOVA p: 0.2
bicor=-0.0011, p=0.99
cor=-0.014, p=0.9
bicor=-0.19, p=0.06
cor=-0.18, p=0.073
ATP6V1H UPenn Mixed PRM
M1 turquoise MEGA module member
K-W ANOVA p: 0.025
ATP6V1H UPenn Mixed PRM
K-W ANOVA p: 0.02
bicor=-0.14, p=0.19
cor=-0.19, p=0.083
bicor=0.0091, p=0.93
cor=-0.0043, p=0.97
PLXNA1 UPenn Mixed PRM
M1 turquoise MEGA module member
K-W ANOVA p: 0.00017
PLXNA1 UPenn Mixed PRM
K-W ANOVA p: 9.6e-05
bicor=-0.32, p=0.0027
cor=-0.38, p=0.00036
bicor=-0.38, p=7.9e-05
cor=-0.44, p=4.6e-06
Relative Log2 Abundance
Control
AD
ALS
FTLDU
PSP
CBD
PD.PDD
MSA
otherND
Braak Score
CERAD Score

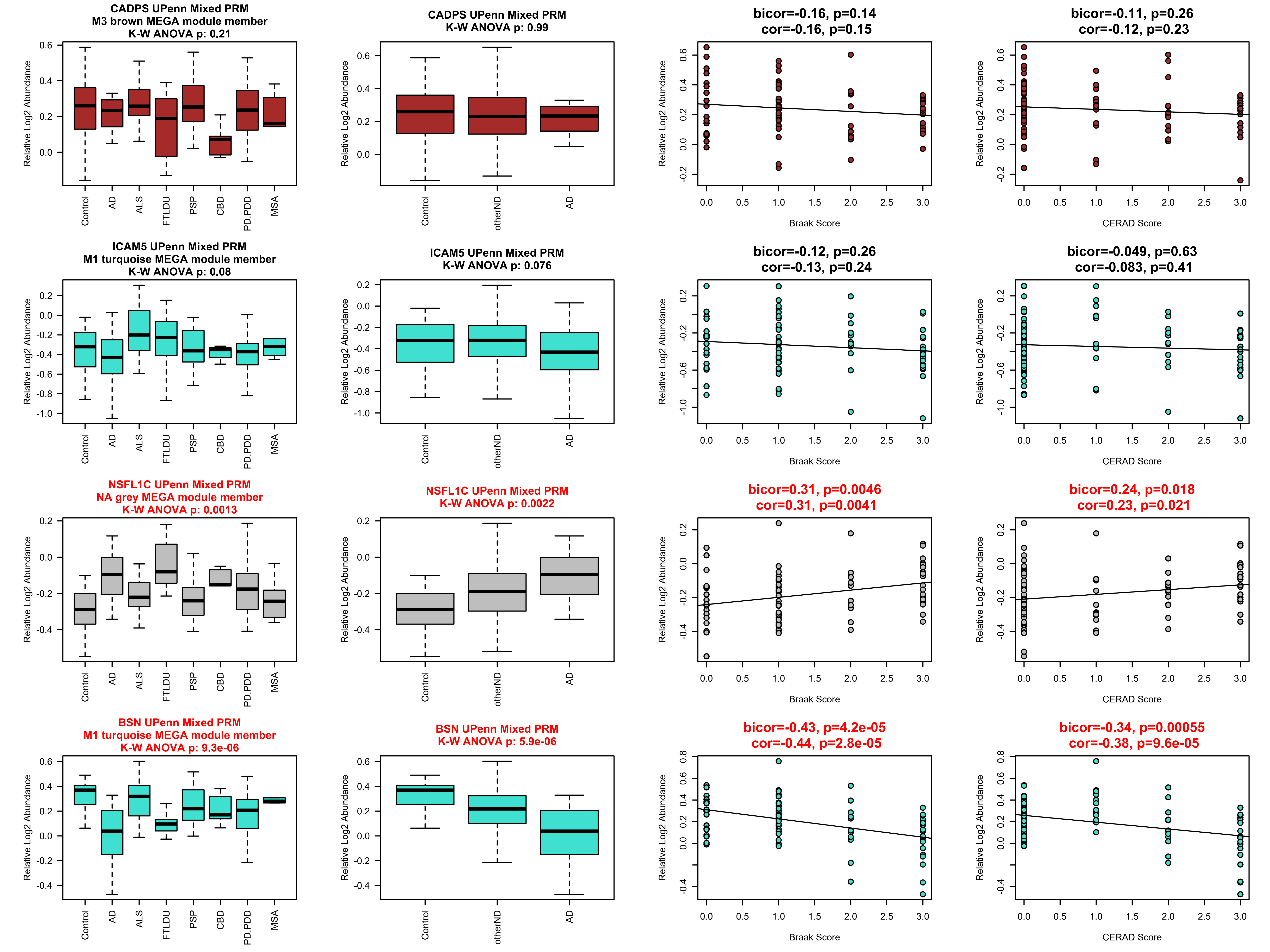
CADPS UPenn Mixed PRM
M3 brown MEGA module member
K-W ANOVA p: 0.21
CADPS UPenn Mixed PRM
K-W ANOVA p: 0.99
bicor=-0.16, p=0.14
cor=-0.16, p=0.15
bicor=-0.11, p=0.26
cor=-0.12, p=0.23
ICAM5 UPenn Mixed PRM
M1 turquoise MEGA module member
K-W ANOVA p: 0.08
ICAM5 UPenn Mixed PRM
K-W ANOVA p: 0.076
bicor=-0.12, p=0.26
cor=-0.13, p=0.24
bicor=-0.049, p=0.63
cor=-0.083, p=0.41
NSFL1C UPenn Mixed PRM
NA grey MEGA module member
K-W ANOVA p: 0.0013
NSFL1C UPenn Mixed PRM
K-W ANOVA p: 0.0022
bicor=0.31, p=0.0046
cor=0.31, p=0.0041
bicor=0.24, p=0.018
cor=0.23, p=0.021
BSN UPenn Mixed PRM
M1 turquoise MEGA module member
K-W ANOVA p: 9.3e-06
BSN UPenn Mixed PRM
K-W ANOVA p: 5.9e-06
bicor=-0.43, p=4.2e-05
cor=-0.44, p=2.8e-05
bicor=-0.34, p=0.00055
cor=-0.38, p=9.6e-05
Relative Log2 Abundance
Control
AD
ALS
FTLDU
PSP
CBD
PD.PDD
MSA
otherND
Braak Score
CERAD Score

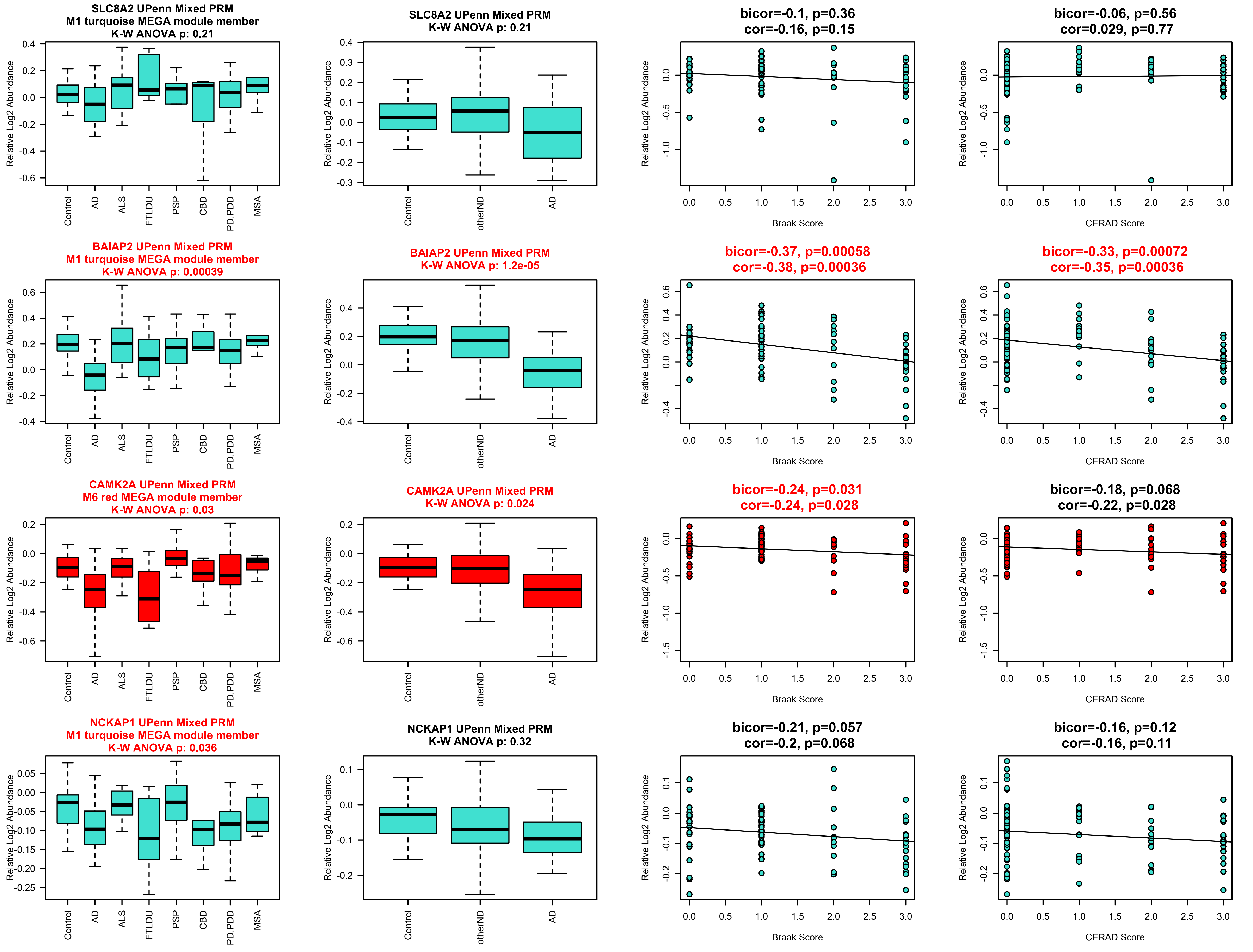

SLC8A2 UPenn Mixed PRM
M1 turquoise MEGA module member
K-W ANOVA p: 0.21
SLC8A2 UPenn Mixed PRM
K-W ANOVA p: 0.21
bicor=-0.1, p=0.36
cor=-0.16, p=0.15
bicor=-0.06, p=0.56
cor=0.029, p=0.77
BAIAP2 UPenn Mixed PRM
M1 turquoise MEGA module member
K-W ANOVA p: 0.00039
BAIAP2 UPenn Mixed PRM
K-W ANOVA p: 1.2e-05
bicor=-0.37, p=0.00058
cor=-0.38, p=0.00036
bicor=-0.33, p=0.00072
cor=-0.35, p=0.00036
CAMK2A UPenn Mixed PRM
M6 red MEGA module member
K-W ANOVA p: 0.03
CAMK2A UPenn Mixed PRM
K-W ANOVA p: 0.024
bicor=-0.24, p=0.031
cor=-0.24, p=0.028
bicor=-0.18, p=0.068
cor=-0.22, p=0.028
NCKAP1 UPenn Mixed PRM
M1 turquoise MEGA module member
K-W ANOVA p: 0.036
NCKAP1 UPenn Mixed PRM
K-W ANOVA p: 0.32
bicor=-0.21, p=0.057
cor=-0.2, p=0.068
bicor=-0.16, p=0.12
cor=-0.16, p=0.11
Relative Log2 Abundance
Control
AD
ALS
FTLDU
PSP
CBD
PD.PDD
MSA
otherND
Braak Score
CERAD Score

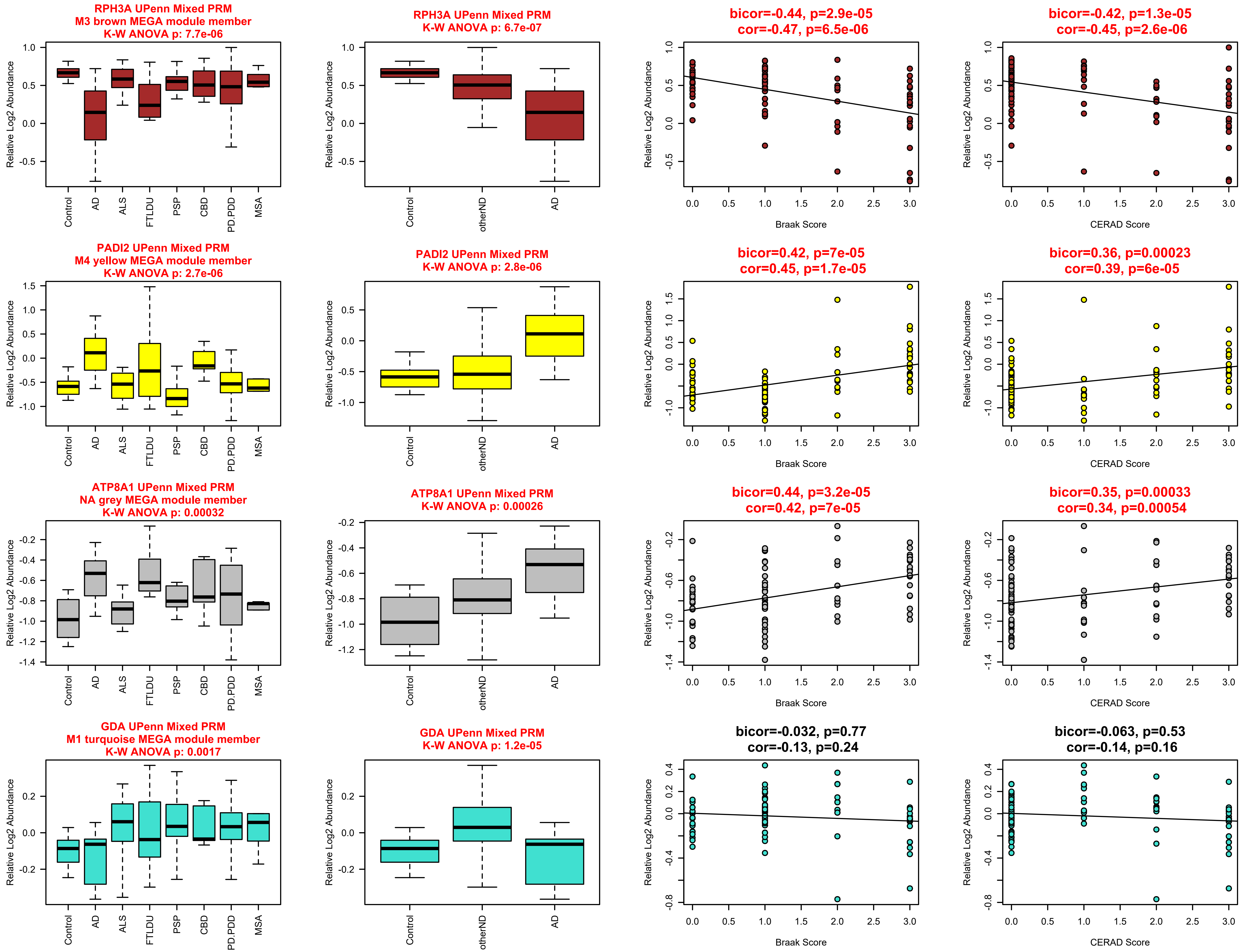

RPH3A UPenn Mixed PRM
M3 brown MEGA module member
K-W ANOVA p: 7.7e-06
RPH3A UPenn Mixed PRM
K-W ANOVA p: 6.7e-07
bicor=-0.44, p=2.9e-05
cor=-0.47, p=6.5e-06
bicor=-0.42, p=1.3e-05
cor=-0.45, p=2.6e-06
PADI2 UPenn Mixed PRM
M4 yellow MEGA module member
K-W ANOVA p: 2.7e-06
PADI2 UPenn Mixed PRM
K-W ANOVA p: 2.8e-06
bicor=0.42, p=7e-05
cor=0.45, p=1.7e-05
bicor=0.36, p=0.00023
cor=0.39, p=6e-05
ATP8A1 UPenn Mixed PRM
NA grey MEGA module member
K-W ANOVA p: 0.00032
ATP8A1 UPenn Mixed PRM
K-W ANOVA p: 0.00026
bicor=0.44, p=3.2e-05
cor=0.42, p=7e-05
bicor=0.35, p=0.00033
cor=0.34, p=0.00054
GDA UPenn Mixed PRM
M1 turquoise MEGA module member
K-W ANOVA p: 0.0017
GDA UPenn Mixed PRM
K-W ANOVA p: 1.2e-05
bicor=-0.032, p=0.77
cor=-0.13, p=0.24
bicor=-0.063, p=0.53
cor=-0.14, p=0.16
Relative Log2 Abundance
Control
AD
ALS
FTLDU
PSP
CBD
PD.PDD
MSA
otherND
Braak Score
CERAD Score

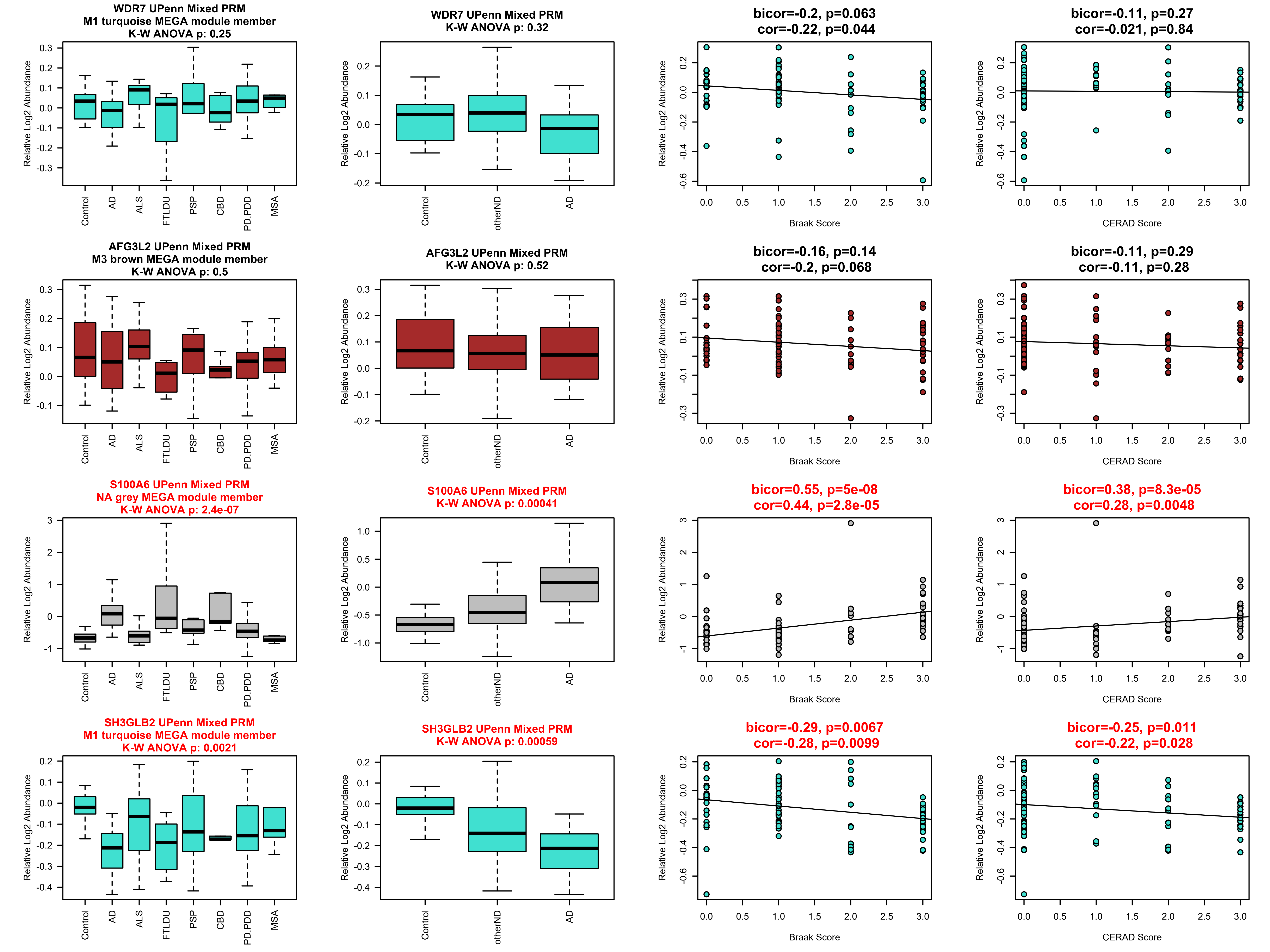
WDR7 UPenn Mixed PRM
M1 turquoise MEGA module member
K-W ANOVA p: 0.25
WDR7 UPenn Mixed PRM
K-W ANOVA p: 0.32
bicor=-0.2, p=0.063
cor=-0.22, p=0.044
bicor=-0.11, p=0.27
cor=-0.021, p=0.84
AFG3L2 UPenn Mixed PRM
M3 brown MEGA module member
K-W ANOVA p: 0.5
AFG3L2 UPenn Mixed PRM
K-W ANOVA p: 0.52
bicor=-0.16, p=0.14
cor=-0.2, p=0.068
bicor=-0.11, p=0.29
cor=-0.11, p=0.28
S100A6 UPenn Mixed PRM
NA grey MEGA module member
K-W ANOVA p: 2.4e-07
S100A6 UPenn Mixed PRM
K-W ANOVA p: 0.00041
bicor=0.55, p=5e-08
cor=0.44, p=2.8e-05
bicor=0.38, p=8.3e-05
cor=0.28, p=0.0048
SH3GLB2 UPenn Mixed PRM
M1 turquoise MEGA module member
K-W ANOVA p: 0.0021
SH3GLB2 UPenn Mixed PRM
K-W ANOVA p: 0.00059
bicor=-0.29, p=0.0067
cor=-0.28, p=0.0099
bicor=-0.25, p=0.011
cor=-0.22, p=0.028
Relative Log2 Abundance
Control
AD
ALS
FTLDU
PSP
CBD
PD.PDD
MSA
otherND
Braak Score
CERAD Score

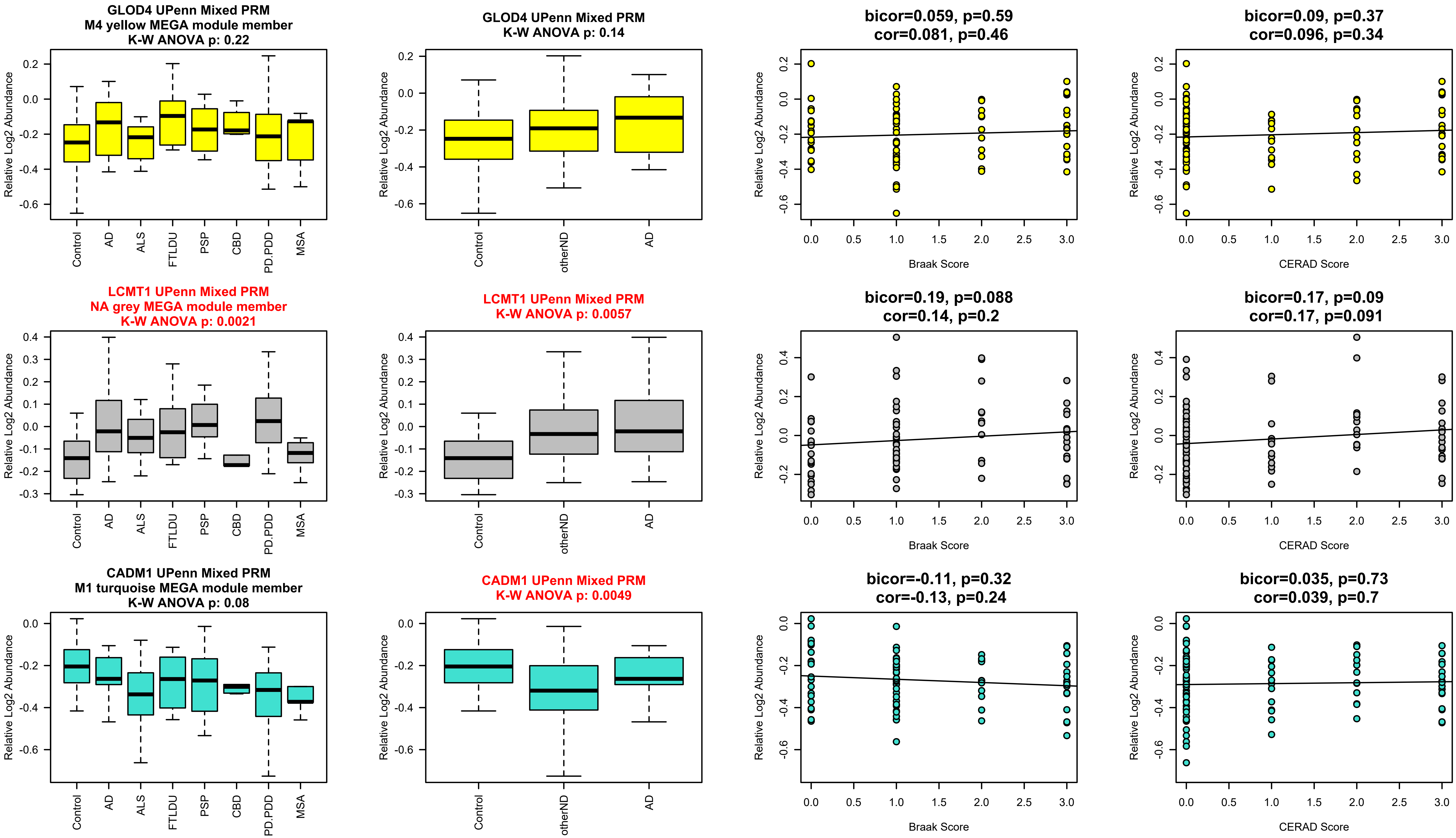
GLOD4 UPenn Mixed PRM
M4 yellow MEGA module member
K-W ANOVA p: 0.22
GLOD4 UPenn Mixed PRM
K-W ANOVA p: 0.14
bicor=0.059, p=0.59
cor=0.081, p=0.46
bicor=0.09, p=0.37
cor=0.096, p=0.34
LCMT1 UPenn Mixed PRM
NA grey MEGA module member
K-W ANOVA p: 0.0021
LCMT1 UPenn Mixed PRM
K-W ANOVA p: 0.0057
bicor=0.19, p=0.088
cor=0.14, p=0.2
bicor=0.17, p=0.09
cor=0.17, p=0.091
CADM1 UPenn Mixed PRM
M1 turquoise MEGA module member
K-W ANOVA p: 0.08
CADM1 UPenn Mixed PRM
K-W ANOVA p: 0.0049
bicor=-0.11, p=0.32
cor=-0.13, p=0.24
bicor=0.035, p=0.73
cor=0.039, p=0.7
Relative Log2 Abundance
Control
AD
ALS
FTLDU
PSP
CBD
PD.PDD
MSA
otherND
Braak Score
CERAD Score

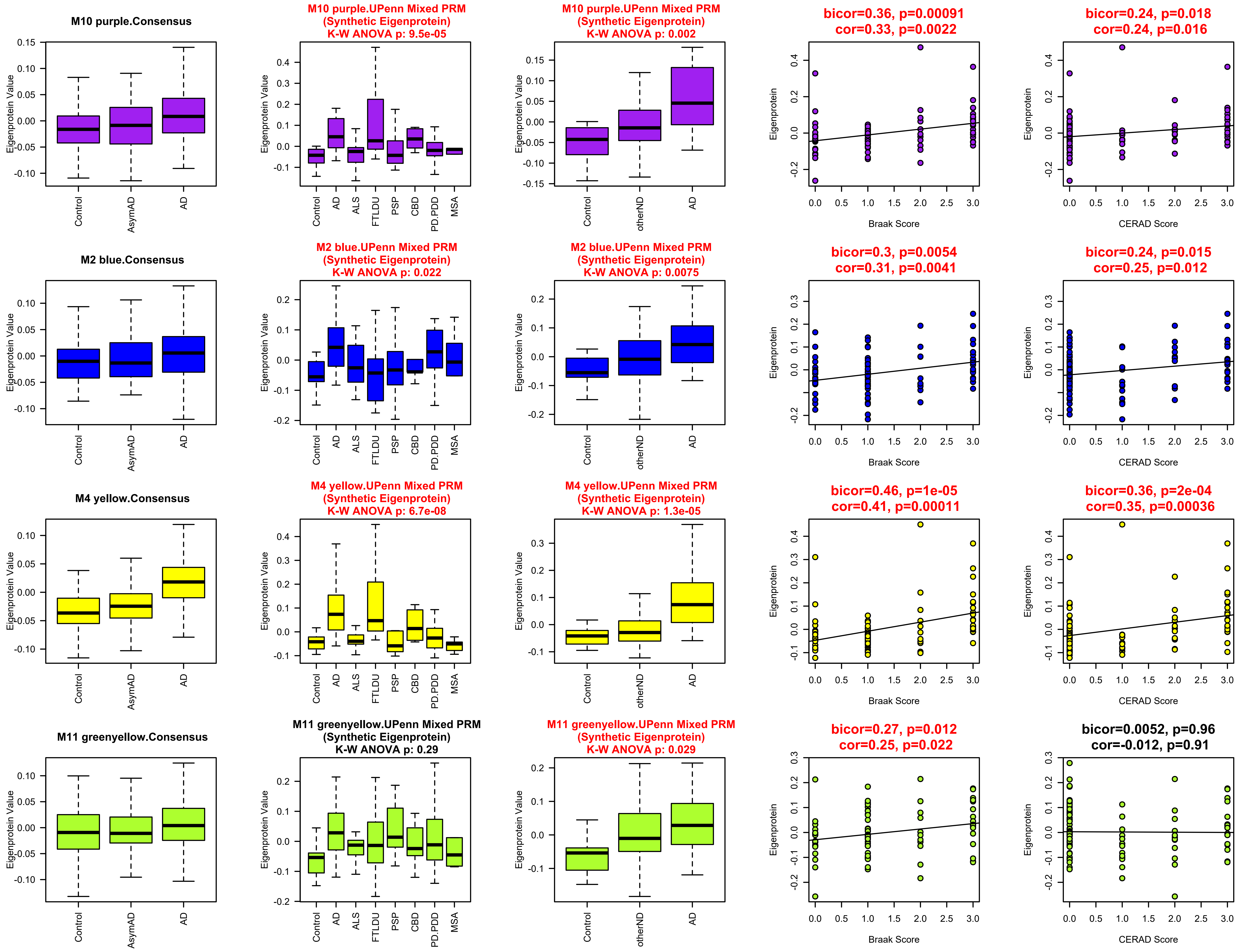
M10 purple.Consensus
M10 purple.UPenn Mixed PRM (Synthetic Eigenprotein) K-W ANOVA p: 9.5e-05
M10 purple.UPenn Mixed PRM (Synthetic Eigenprotein) K-W ANOVA p: 0.002
bicor=0.36, p=0.00091 cor=0.33, p=0.0022
bicor=0.24, p=0.018 cor=0.24, p=0.016
M2 blue.Consensus
M2 blue.UPenn Mixed PRM (Synthetic Eigenprotein) K-W ANOVA p: 0.022
M2 blue.UPenn Mixed PRM (Synthetic Eigenprotein) K-W ANOVA p: 0.0075
bicor=0.3, p=0.0054 cor=0.31, p=0.0041
bicor=0.24, p=0.015 cor=0.25, p=0.012
M4 yellow.Consensus
M4 yellow.UPenn Mixed PRM (Synthetic Eigenprotein) K-W ANOVA p: 6.7e-08
M4 yellow.UPenn Mixed PRM (Synthetic Eigenprotein) K-W ANOVA p: 1.3e-05
bicor=0.46, p=1e-05 cor=0.41, p=0.00011
bicor=0.36, p=2e-04 cor=0.35, p=0.00036
M11 greenyellow.Consensus
M11 greenyellow.UPenn Mixed PRM (Synthetic Eigenprotein) K-W ANOVA p: 0.29
M11 greenyellow.UPenn Mixed PRM (Synthetic Eigenprotein) K-W ANOVA p: 0.029
bicor=0.27, p=0.012 cor=0.25, p=0.022
bicor=0.0052, p=0.96 cor=-0.012, p=0.91
Eigenprotein Value
Eigenprotein
Control
AsymAD
AD
ALS
FTLDU
PSP
CBD
PD.PDD
MSA
otherND
Braak Score
CERAD Score

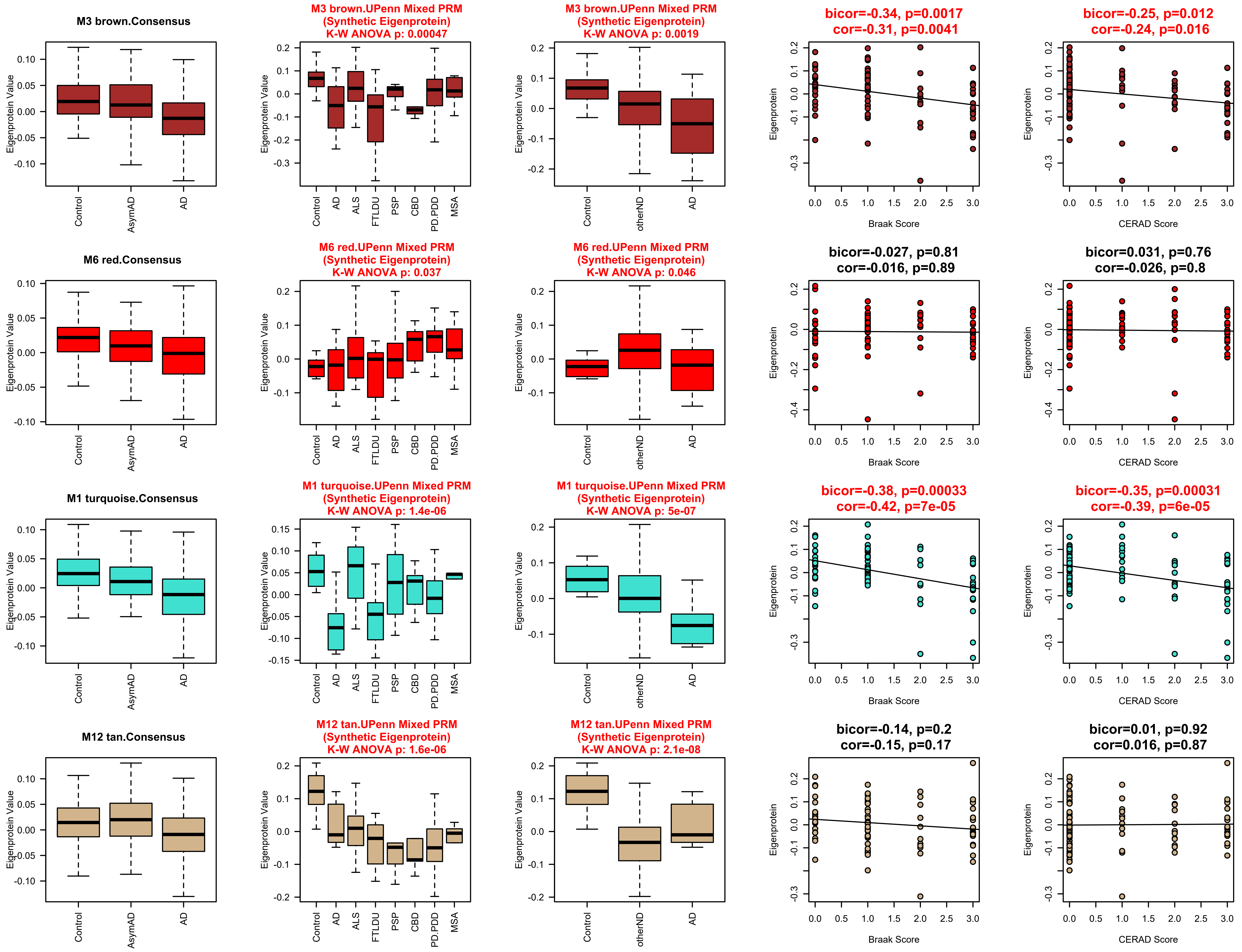
M3 brown.Consensus
M3 brown.UPenn Mixed PRM (Synthetic Eigenprotein) K-W ANOVA p: 0.00047
M3 brown.UPenn Mixed PRM (Synthetic Eigenprotein) K-W ANOVA p: 0.0019
bicor=-0.34, p=0.0017 cor=-0.31, p=0.0041
bicor=-0.25, p=0.012 cor=-0.24, p=0.016
M6 red.Consensus
M6 red.UPenn Mixed PRM (Synthetic Eigenprotein) K-W ANOVA p: 0.037
M6 red.UPenn Mixed PRM (Synthetic Eigenprotein) K-W ANOVA p: 0.046
bicor=-0.027, p=0.81 cor=-0.016, p=0.89
bicor=0.031, p=0.76 cor=-0.026, p=0.8
M1 turquoise.Consensus
M1 turquoise.UPenn Mixed PRM (Synthetic Eigenprotein) K-W ANOVA p: 1.4e-06
M1 turquoise.UPenn Mixed PRM (Synthetic Eigenprotein) K-W ANOVA p: 5e-07
bicor=-0.38, p=0.00033 cor=-0.42, p=7e-05
bicor=-0.35, p=0.00031 cor=-0.39, p=6e-05
M12 tan.Consensus
M12 tan.UPenn Mixed PRM (Synthetic Eigenprotein) K-W ANOVA p: 1.6e-06
M12 tan.UPenn Mixed PRM (Synthetic Eigenprotein) K-W ANOVA p: 2.1e-08
bicor=-0.14, p=0.2 cor=-0.15, p=0.17
bicor=0.01, p=0.92 cor=0.016, p=0.87
Eigenprotein Value
Eigenprotein
Control
AsymAD
AD
ALS
FTLDU
PSP
CBD
PD.PDD
MSA
otherND
Braak Score
CERAD Score

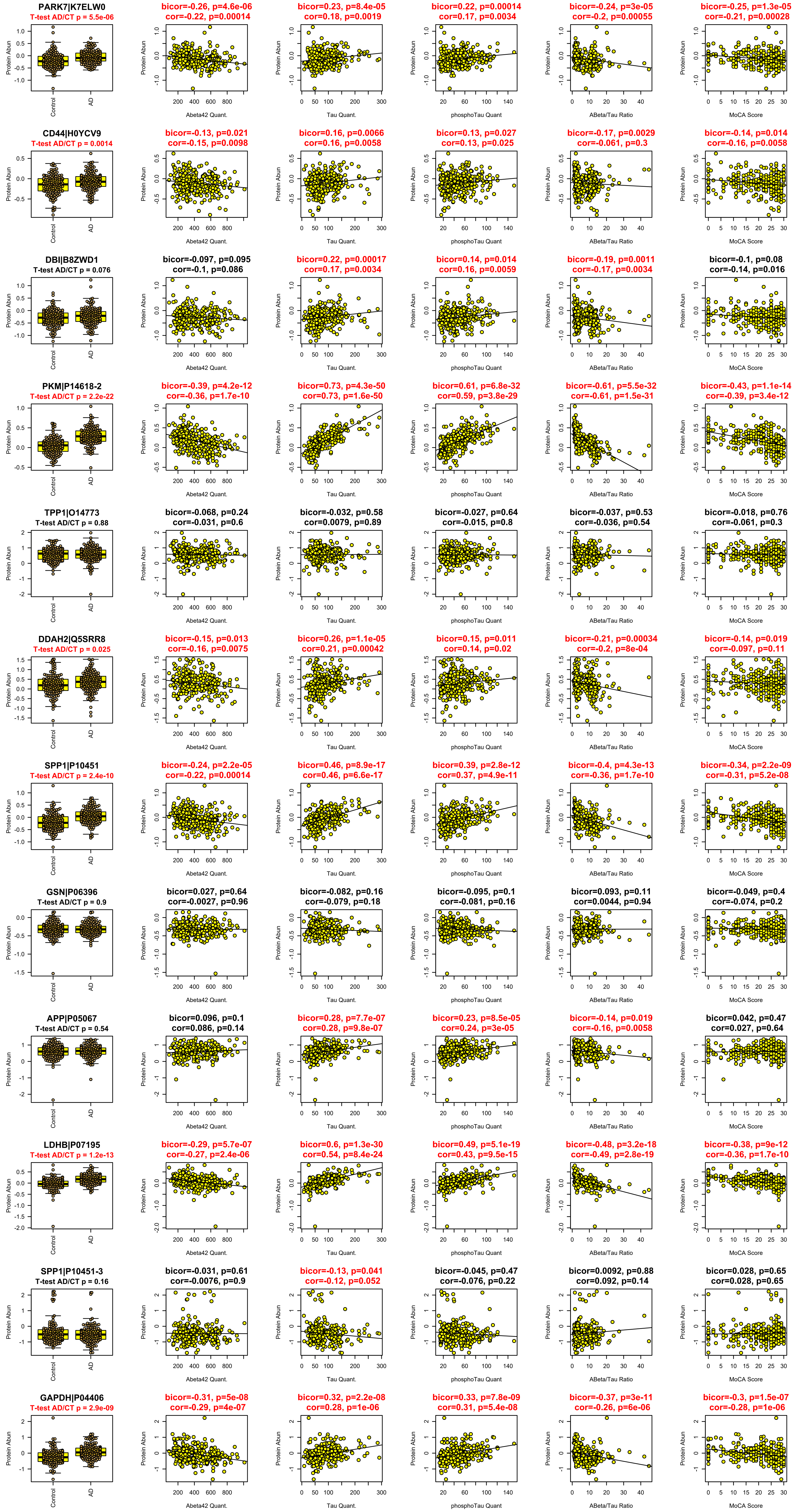

PARK7|K7ELW0
T-test AD/CT p = 5.5e-06
bicor=-0.26, p=4.6e-06
cor=-0.22, p=0.00014
bicor=0.23, p=8.4e-05
cor=0.18, p=0.0019
bicor=0.22, p=0.00014
cor=0.17, p=0.0034
bicor=-0.24, p=3e-05
cor=-0.2, p=0.00055
bicor=-0.25, p=1.3e-05
cor=-0.21, p=0.00028
CD44|H0YCV9
T-test AD/CT p = 0.0014
bicor=-0.13, p=0.021
cor=-0.15, p=0.0098
bicor=0.16, p=0.0066
cor=0.16, p=0.0058
bicor=0.13, p=0.027
cor=0.13, p=0.025
bicor=-0.17, p=0.0029
cor=-0.061, p=0.3
bicor=-0.14, p=0.014
cor=-0.16, p=0.0058
DBI|B8ZWD1
T-test AD/CT p = 0.076
bicor=-0.097, p=0.095
cor=-0.1, p=0.086
bicor=0.22, p=0.00017
cor=0.17, p=0.0034
bicor=0.14, p=0.014
cor=0.16, p=0.0059
bicor=-0.19, p=0.0011
cor=-0.17, p=0.0034
bicor=-0.1, p=0.08
cor=-0.14, p=0.016
PKM|P14618-2
T-test AD/CT p = 2.2e-22
bicor=-0.39, p=4.2e-12
cor=-0.36, p=1.7e-10
bicor=0.73, p=4.3e-50
cor=0.73, p=1.6e-50
bicor=0.61, p=6.8e-32
cor=0.59, p=3.8e-29
bicor=-0.61, p=5.5e-32
cor=-0.61, p=1.5e-31
bicor=-0.43, p=1.1e-14
cor=-0.39, p=3.4e-12
TPP1|O14773
T-test AD/CT p = 0.88
bicor=-0.068, p=0.24
cor=-0.031, p=0.6
bicor=-0.032, p=0.58
cor=0.0079, p=0.89
bicor=-0.027, p=0.64
cor=-0.015, p=0.8
bicor=-0.037, p=0.53
cor=-0.036, p=0.54
bicor=-0.018, p=0.76
cor=-0.061, p=0.3
DDAH2|Q5SRR8
T-test AD/CT p = 0.025
bicor=-0.15, p=0.013
cor=-0.16, p=0.0075
bicor=0.26, p=1.1e-05
cor=0.21, p=0.00042
bicor=0.15, p=0.011
cor=0.14, p=0.02
bicor=-0.21, p=0.00034
cor=-0.2, p=8e-04
bicor=-0.14, p=0.019
cor=-0.097, p=0.11
SPP1|P10451
T-test AD/CT p = 2.4e-10
bicor=-0.24, p=2.2e-05
cor=-0.22, p=0.00014
bicor=0.46, p=8.9e-17
cor=0.46, p=6.6e-17
bicor=0.39, p=2.8e-12
cor=0.37, p=4.9e-11
bicor=-0.4, p=4.3e-13
cor=-0.36, p=1.7e-10
bicor=-0.34, p=2.2e-09
cor=-0.31, p=5.2e-08
GSN|P06396
T-test AD/CT p = 0.9
bicor=0.027, p=0.64
cor=-0.0027, p=0.96
bicor=-0.082, p=0.16
cor=-0.079, p=0.18
bicor=-0.095, p=0.1
cor=-0.081, p=0.16
bicor=0.093, p=0.11
cor=0.0044, p=0.94
bicor=-0.049, p=0.4
cor=-0.074, p=0.2
APP|P05067
T-test AD/CT p = 0.54
bicor=0.096, p=0.1
cor=0.086, p=0.14
bicor=0.28, p=7.7e-07
cor=0.28, p=9.8e-07
bicor=0.23, p=8.5e-05
cor=0.24, p=3e-05
bicor=-0.14, p=0.019
cor=-0.16, p=0.0058
bicor=0.042, p=0.47
cor=0.027, p=0.64
LDHB|P07195
T-test AD/CT p = 1.2e-13
bicor=-0.29, p=5.7e-07
cor=-0.27, p=2.4e-06
bicor=0.6, p=1.3e-30
cor=0.54, p=8.4e-24
bicor=0.49, p=5.1e-19
cor=0.43, p=9.5e-15
bicor=-0.48, p=3.2e-18
cor=-0.49, p=2.8e-19
bicor=-0.38, p=9e-12
cor=-0.36, p=1.7e-10
SPP1|P10451-3
T-test AD/CT p = 0.16
bicor=-0.031, p=0.61
cor=-0.0076, p=0.9
bicor=-0.13, p=0.041
cor=-0.12, p=0.052
bicor=-0.045, p=0.47
cor=-0.076, p=0.22
bicor=0.0092, p=0.88
cor=0.092, p=0.14
bicor=0.028, p=0.65
cor=0.028, p=0.65
GAPDH|P04406
T-test AD/CT p = 2.9e-09
bicor=-0.31, p=5e-08
cor=-0.29, p=4e-07
bicor=0.32, p=2.2e-08
cor=0.28, p=1e-06
bicor=0.33, p=7.8e-09
cor=0.31, p=5.4e-08
bicor=-0.37, p=3e-11
cor=-0.26, p=6e-06
bicor=-0.3, p=1.5e-07
cor=-0.28, p=1e-06
Protein Abun
Control
AD
Abeta42 Quant.
Tau Quant.
phosphoTau Quant
ABeta/Tau Ratio
MoCA Score

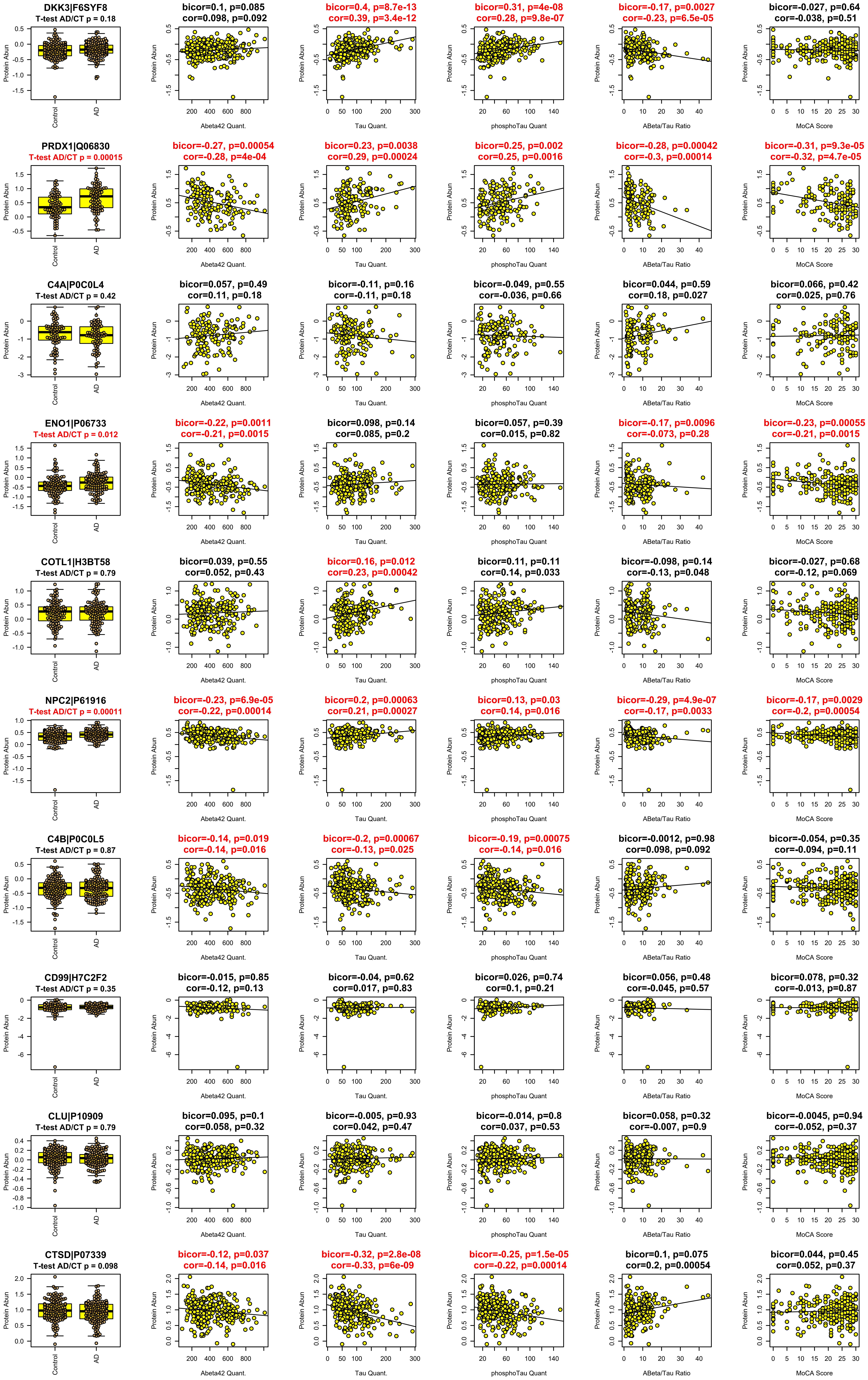
DKK3|F6SYF8
T-test AD/CT p = 0.18
bicor=0.1, p=0.085
cor=0.098, p=0.092
bicor=0.4, p=8.7e-13
cor=0.39, p=3.4e-12
bicor=0.31, p=4e-08
cor=0.28, p=9.8e-07
bicor=-0.17, p=0.0027
cor=-0.23, p=6.5e-05
bicor=-0.027, p=0.64
cor=-0.038, p=0.51
PRDX1|Q06830
T-test AD/CT p = 0.00015
bicor=-0.27, p=0.00054
cor=-0.28, p=4e-04
bicor=0.23, p=0.0038
cor=0.29, p=0.00024
bicor=0.25, p=0.002
cor=0.25, p=0.0016
bicor=-0.28, p=0.00042
cor=-0.3, p=0.00014
bicor=-0.31, p=9.3e-05
cor=-0.32, p=4.7e-05
C4A|P0C0L4
T-test AD/CT p = 0.42
bicor=0.057, p=0.49
cor=0.11, p=0.18
bicor=-0.11, p=0.16
cor=-0.11, p=0.18
bicor=-0.049, p=0.55
cor=-0.036, p=0.66
bicor=0.044, p=0.59
cor=0.18, p=0.027
bicor=0.066, p=0.42
cor=0.025, p=0.76
ENO1|P06733
T-test AD/CT p = 0.012
bicor=-0.22, p=0.0011
cor=-0.21, p=0.0015
bicor=0.098, p=0.14
cor=0.085, p=0.2
bicor=0.057, p=0.39
cor=0.015, p=0.82
bicor=-0.17, p=0.0096
cor=-0.073, p=0.28
bicor=-0.23, p=0.00055
cor=-0.21, p=0.0015
COTL1|H3BT58
T-test AD/CT p = 0.79
bicor=0.039, p=0.55
cor=0.052, p=0.43
bicor=0.16, p=0.012
cor=0.23, p=0.00042
bicor=0.11, p=0.11
cor=0.14, p=0.033
bicor=-0.098, p=0.14
cor=-0.13, p=0.048
bicor=-0.027, p=0.68
cor=-0.12, p=0.069
NPC2|P61916
T-test AD/CT p = 0.00011
bicor=-0.23, p=6.9e-05
cor=-0.22, p=0.00014
bicor=0.2, p=0.00063
cor=0.21, p=0.00027
bicor=0.13, p=0.03
cor=0.14, p=0.016
bicor=-0.29, p=4.9e-07
cor=-0.17, p=0.0033
bicor=-0.17, p=0.0029
cor=-0.2, p=0.00054
C4B|P0C0L5
T-test AD/CT p = 0.87
bicor=-0.14, p=0.019
cor=-0.14, p=0.016
bicor=-0.2, p=0.00067
cor=-0.13, p=0.025
bicor=-0.19, p=0.00075
cor=-0.14, p=0.016
bicor=-0.0012, p=0.98
cor=0.098, p=0.092
bicor=-0.054, p=0.35
cor=-0.094, p=0.11
CD99|H7C2F2
T-test AD/CT p = 0.35
bicor=-0.015, p=0.85
cor=-0.12, p=0.13
bicor=-0.04, p=0.62
cor=0.017, p=0.83
bicor=0.026, p=0.74
cor=0.1, p=0.21
bicor=0.056, p=0.48
cor=-0.045, p=0.57
bicor=0.078, p=0.32
cor=-0.013, p=0.87
CLU|P10909
T-test AD/CT p = 0.79
bicor=0.095, p=0.1
cor=0.058, p=0.32
bicor=-0.005, p=0.93
cor=0.042, p=0.47
bicor=-0.014, p=0.8
cor=0.037, p=0.53
bicor=0.058, p=0.32
cor=-0.007, p=0.9
bicor=-0.0045, p=0.94
cor=-0.052, p=0.37
CTSD|P07339
T-test AD/CT p = 0.098
bicor=-0.12, p=0.037
cor=-0.14, p=0.016
bicor=-0.32, p=2.8e-08
cor=-0.33, p=6e-09
bicor=-0.25, p=1.5e-05
cor=-0.22, p=0.00014
bicor=0.1, p=0.075
cor=0.2, p=0.00054
bicor=0.044, p=0.45
cor=0.052, p=0.37
Protein Abun
Control
AD
Abeta42 Quant.
Tau Quant.
phosphoTau Quant
ABeta/Tau Ratio
MoCA Score

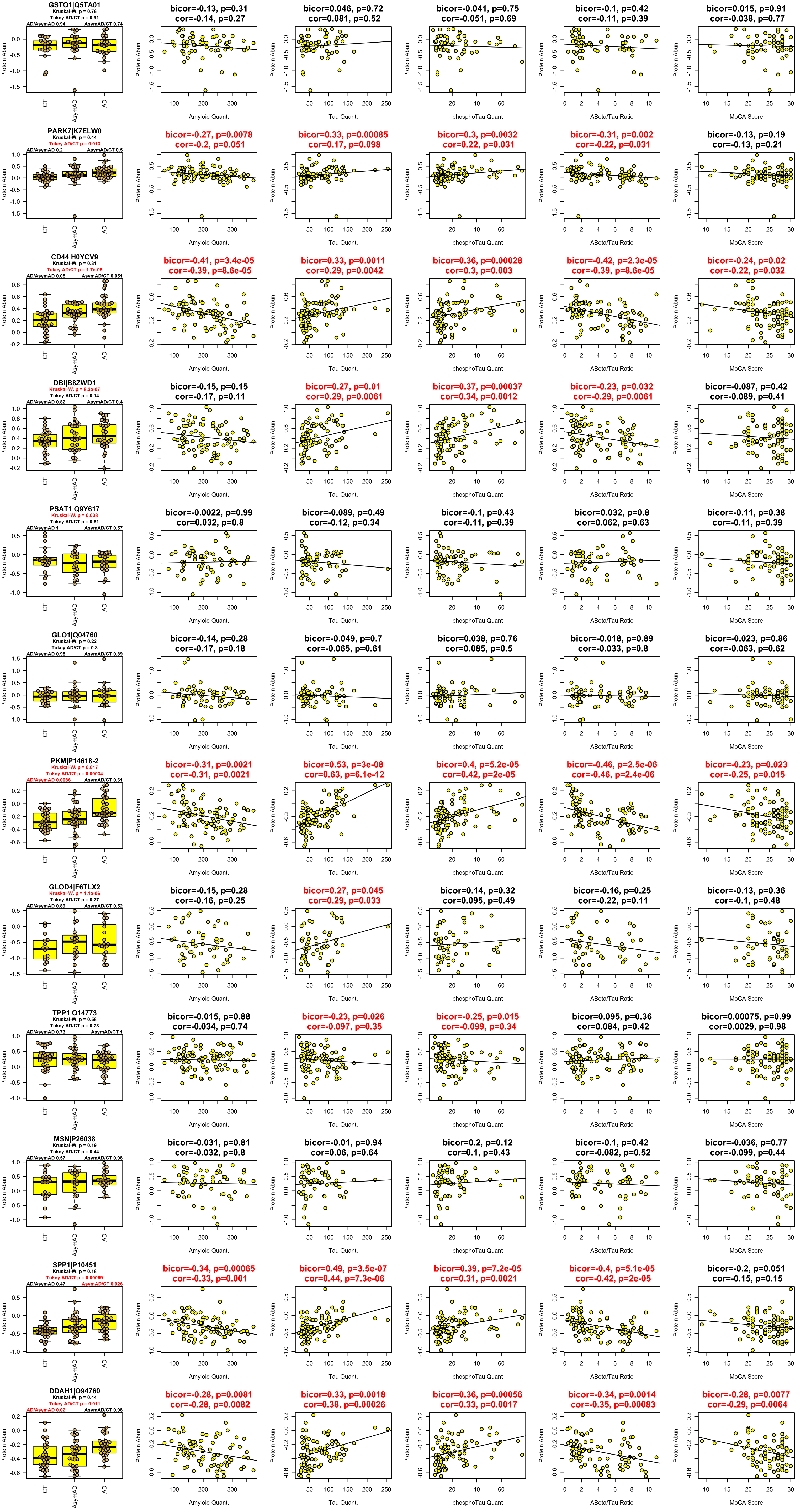

GSTO1|Q5TA01
Kruskal-W. p = 0.76
Tukey AD/CT p = 0.91
AD/AsymAD 0.94
AsymAD/CT 0.74
bicor=-0.13, p=0.31
cor=-0.14, p=0.27
bicor=0.046, p=0.72
cor=0.081, p=0.52
bicor=-0.041, p=0.75
cor=-0.051, p=0.69
bicor=-0.1, p=0.42
cor=-0.11, p=0.39
bicor=0.015, p=0.91
cor=-0.038, p=0.77
PARK7|K7ELW0
Kruskal-W. p = 0.44
Tukey AD/CT p = 0.013
AD/AsymAD 0.2
AsymAD/CT 0.5
bicor=-0.27, p=0.0078
cor=-0.2, p=0.051
bicor=0.33, p=0.00085
cor=0.17, p=0.098
bicor=0.3, p=0.0032
cor=0.22, p=0.031
bicor=-0.31, p=0.002
cor=-0.22, p=0.031
bicor=-0.13, p=0.19
cor=-0.13, p=0.21
CD44|H0YCV9
Kruskal-W. p = 0.31
Tukey AD/CT p = 1.7e-05
AD/AsymAD 0.05
AsymAD/CT 0.051
bicor=-0.41, p=3.4e-05
cor=-0.39, p=8.6e-05
bicor=0.33, p=0.0011
cor=0.29, p=0.0042
bicor=0.36, p=0.00028
cor=0.3, p=0.003
bicor=-0.42, p=2.3e-05
cor=-0.39, p=8.6e-05
bicor=-0.24, p=0.02
cor=-0.22, p=0.032
DBI|B8ZWD1
Kruskal-W. p = 8.2e-07
Tukey AD/CT p = 0.14
AD/AsymAD 0.82
AsymAD/CT 0.4
bicor=-0.15, p=0.15
cor=-0.17, p=0.11
bicor=0.27, p=0.01
cor=0.29, p=0.0061
bicor=0.37, p=0.00037
cor=0.34, p=0.0012
bicor=-0.23, p=0.032
cor=-0.29, p=0.0061
bicor=-0.087, p=0.42
cor=-0.089, p=0.41
PSAT1|Q9Y617
Kruskal-W. p = 0.038
Tukey AD/CT p = 0.61
AD/AsymAD 1
AsymAD/CT 0.57
bicor=-0.0022, p=0.99
cor=0.032, p=0.8
bicor=-0.089, p=0.49
cor=-0.12, p=0.34
bicor=-0.1, p=0.43
cor=-0.11, p=0.39
bicor=0.032, p=0.8
cor=0.062, p=0.63
bicor=-0.11, p=0.38
cor=-0.11, p=0.39
GLO1|Q04760
Kruskal-W. p = 0.22
Tukey AD/CT p = 0.8
AD/AsymAD 0.98
AsymAD/CT 0.89
bicor=-0.14, p=0.28
cor=-0.17, p=0.18
bicor=-0.049, p=0.7
cor=-0.065, p=0.61
bicor=0.038, p=0.76
cor=0.085, p=0.5
bicor=-0.018, p=0.89
cor=-0.033, p=0.8
bicor=-0.023, p=0.86
cor=-0.063, p=0.62
PKM|P14618-2
Kruskal-W. p = 0.017
Tukey AD/CT p = 0.00034
AD/AsymAD 0.0086
AsymAD/CT 0.61
bicor=-0.31, p=0.0021
cor=-0.31, p=0.0021
bicor=0.53, p=3e-08
cor=0.63, p=6.1e-12
bicor=0.4, p=5.2e-05
cor=0.42, p=2e-05
bicor=-0.46, p=2.5e-06
cor=-0.46, p=2.4e-06
bicor=-0.23, p=0.023
cor=-0.25, p=0.015
GLOD4|F6TLX2
Kruskal-W. p = 1.1e-06
Tukey AD/CT p = 0.27
AD/AsymAD 0.89
AsymAD/CT 0.52
bicor=-0.15, p=0.28
cor=-0.16, p=0.25
bicor=0.27, p=0.045
cor=0.29, p=0.033
bicor=0.14, p=0.32
cor=0.095, p=0.49
bicor=-0.16, p=0.25
cor=-0.22, p=0.11
bicor=-0.13, p=0.36
cor=-0.1, p=0.48
TPP1|O14773
Kruskal-W. p = 0.58
Tukey AD/CT p = 0.73
AD/AsymAD 0.73
AsymAD/CT 1
bicor=-0.015, p=0.88
cor=-0.034, p=0.74
bicor=-0.23, p=0.026
cor=-0.097, p=0.35
bicor=-0.25, p=0.015
cor=-0.099, p=0.34
bicor=0.095, p=0.36
cor=0.084, p=0.42
bicor=0.00075, p=0.99
cor=0.0029, p=0.98
MSN|P26038
Kruskal-W. p = 0.19
Tukey AD/CT p = 0.44
AD/AsymAD 0.57
AsymAD/CT 0.98
bicor=-0.031, p=0.81
cor=-0.032, p=0.8
bicor=-0.01, p=0.94
cor=0.06, p=0.64
bicor=0.2, p=0.12
cor=0.1, p=0.43
bicor=-0.1, p=0.42
cor=-0.082, p=0.52
bicor=-0.036, p=0.77
cor=-0.099, p=0.44
SPP1|P10451
Kruskal-W. p = 0.18
Tukey AD/CT p = 0.00059
AD/AsymAD 0.47
AsymAD/CT 0.026
bicor=-0.34, p=0.00065
cor=-0.33, p=0.001
bicor=0.49, p=3.5e-07
cor=0.44, p=7.3e-06
bicor=0.39, p=7.2e-05
cor=0.31, p=0.0021
bicor=-0.4, p=5.1e-05
cor=-0.42, p=2e-05
bicor=-0.2, p=0.051
cor=-0.15, p=0.15
DDAH1|O94760
Kruskal-W. p = 0.44
Tukey AD/CT p = 0.011
AD/AsymAD 0.02
AsymAD/CT 0.98
bicor=-0.28, p=0.0081
cor=-0.28, p=0.0082
bicor=0.33, p=0.0018
cor=0.38, p=0.00026
bicor=0.36, p=0.00056
cor=0.33, p=0.0017
bicor=-0.34, p=0.0014
cor=-0.35, p=0.00083
bicor=-0.28, p=0.0077
cor=-0.29, p=0.0064
Protein Abun
CT
AsymAD
AD
Amyloid Quant.
Tau Quant.
phosphoTau Quant
ABeta/Tau Ratio
MoCA Score

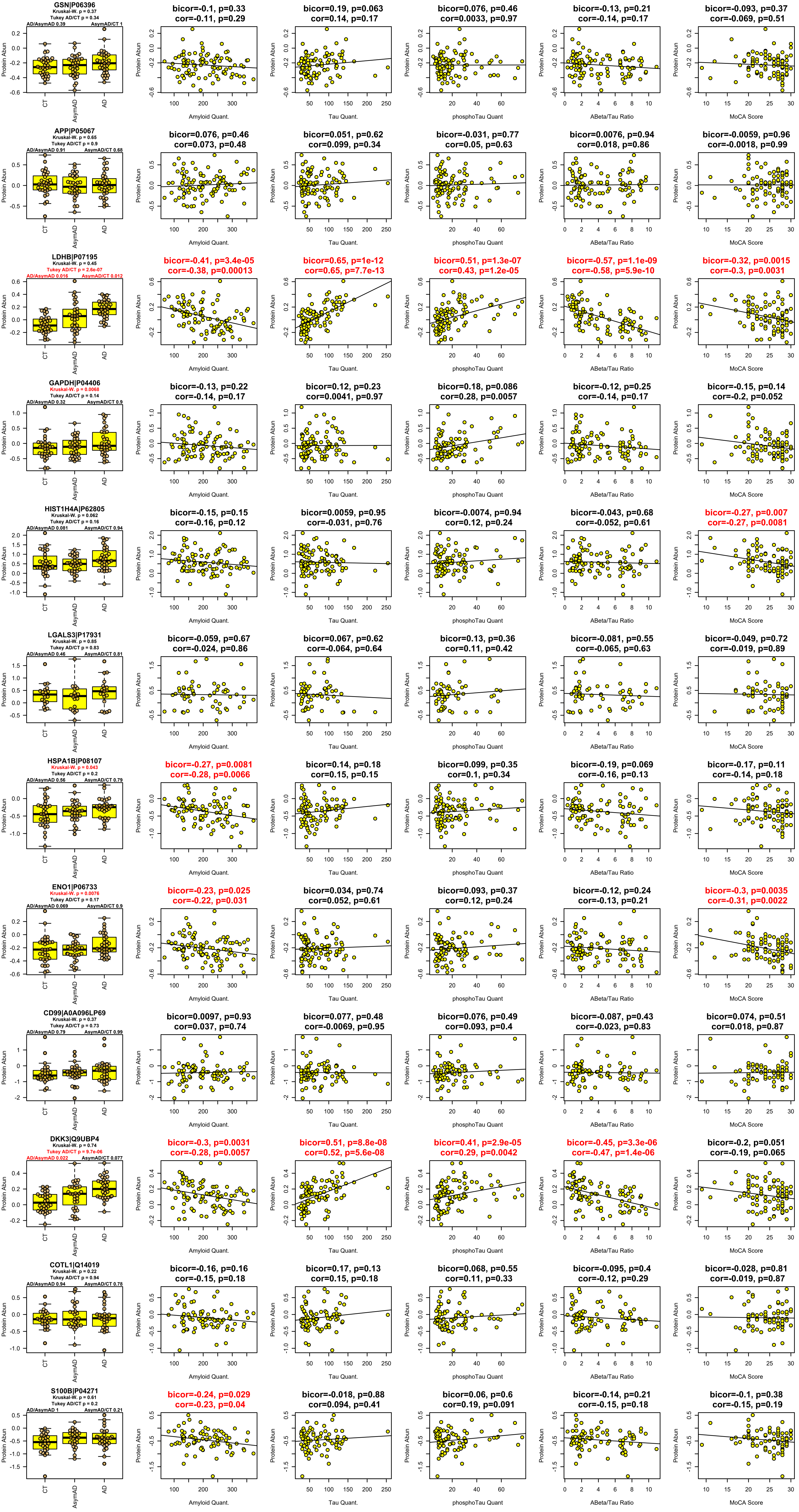

GSN|P06396
Kruskal-W. p = 0.37
Tukey AD/CT p = 0.34
AD/AsymAD 0.39
AsymAD/CT 1
bicor=-0.1, p=0.33
cor=-0.11, p=0.29
bicor=0.19, p=0.063
cor=0.14, p=0.17
bicor=0.076, p=0.46
cor=0.0033, p=0.97
bicor=-0.13, p=0.21
cor=-0.14, p=0.17
bicor=-0.093, p=0.37
cor=-0.069, p=0.51
APP|P05067
Kruskal-W. p = 0.65
Tukey AD/CT p = 0.9
AD/AsymAD 0.91
AsymAD/CT 0.68
bicor=0.076, p=0.46
cor=0.073, p=0.48
bicor=0.051, p=0.62
cor=0.099, p=0.34
bicor=-0.031, p=0.77
cor=0.05, p=0.63
bicor=0.0076, p=0.94
cor=0.018, p=0.86
bicor=-0.0059, p=0.96
cor=-0.0018, p=0.99
LDHB|P07195
Kruskal-W. p = 0.45
Tukey AD/CT p = 2.6e-07
AD/AsymAD 0.016
AsymAD/CT 0.012
bicor=-0.41, p=3.4e-05
cor=-0.38, p=0.00013
bicor=0.65, p=1e-12
cor=0.65, p=7.7e-13
bicor=0.51, p=1.3e-07
cor=0.43, p=1.2e-05
bicor=-0.57, p=1.1e-09
cor=-0.58, p=5.9e-10
bicor=-0.32, p=0.0015
cor=-0.3, p=0.0031
GAPDH|P04406
Kruskal-W. p = 0.0068
Tukey AD/CT p = 0.14
AD/AsymAD 0.32
AsymAD/CT 0.9
bicor=-0.13, p=0.22
cor=-0.14, p=0.17
bicor=0.12, p=0.23
cor=0.0041, p=0.97
bicor=0.18, p=0.086
cor=0.28, p=0.0057
bicor=-0.12, p=0.25
cor=-0.14, p=0.17
bicor=-0.15, p=0.14
cor=-0.2, p=0.052
HIST1H4A|P62805
Kruskal-W. p = 0.062
Tukey AD/CT p = 0.16
AD/AsymAD 0.081
AsymAD/CT 0.94
bicor=-0.15, p=0.15
cor=-0.16, p=0.12
bicor=0.0059, p=0.95
cor=-0.031, p=0.76
bicor=-0.0074, p=0.94
cor=0.12, p=0.24
bicor=-0.043, p=0.68
cor=-0.052, p=0.61
bicor=-0.27, p=0.007
cor=-0.27, p=0.0081
LGALS3|P17931
Kruskal-W. p = 0.85
Tukey AD/CT p = 0.83
AD/AsymAD 0.46
AsymAD/CT 0.81
bicor=-0.059, p=0.67
cor=-0.024, p=0.86
bicor=0.067, p=0.62
cor=-0.064, p=0.64
bicor=0.13, p=0.36
cor=0.11, p=0.42
bicor=-0.081, p=0.55
cor=-0.065, p=0.63
bicor=-0.049, p=0.72
cor=-0.019, p=0.89
HSPA1B|P08107
Kruskal-W. p = 0.043
Tukey AD/CT p = 0.2
AD/AsymAD 0.56
AsymAD/CT 0.79
bicor=-0.27, p=0.0081
cor=-0.28, p=0.0066
bicor=0.14, p=0.18
cor=0.15, p=0.15
bicor=0.099, p=0.35
cor=0.1, p=0.34
bicor=-0.19, p=0.069
cor=-0.16, p=0.13
bicor=-0.17, p=0.11
cor=-0.14, p=0.18
ENO1|P06733
Kruskal-W. p = 0.0076
Tukey AD/CT p = 0.17
AD/AsymAD 0.069
AsymAD/CT 0.9
bicor=-0.23, p=0.025
cor=-0.22, p=0.031
bicor=0.034, p=0.74
cor=0.052, p=0.61
bicor=0.093, p=0.37
cor=0.12, p=0.24
bicor=-0.12, p=0.24
cor=-0.13, p=0.21
bicor=-0.3, p=0.0035
cor=-0.31, p=0.0022
CD99|A0A096LP69
Kruskal-W. p = 0.37
Tukey AD/CT p = 0.73
AD/AsymAD 0.79
AsymAD/CT 0.99
bicor=0.0097, p=0.93
cor=0.037, p=0.74
bicor=0.077, p=0.48
cor=-0.0069, p=0.95
bicor=0.076, p=0.49
cor=0.093, p=0.4
bicor=-0.087, p=0.43
cor=-0.023, p=0.83
bicor=0.074, p=0.51
cor=0.018, p=0.87
DKK3|Q9UBP4
Kruskal-W. p = 0.74
Tukey AD/CT p = 9.7e-06
AD/AsymAD 0.022
AsymAD/CT 0.077
bicor=-0.3, p=0.0031
cor=-0.28, p=0.0057
bicor=0.51, p=8.8e-08
cor=0.52, p=5.6e-08
bicor=0.41, p=2.9e-05
cor=0.29, p=0.0042
bicor=-0.45, p=3.3e-06
cor=-0.47, p=1.4e-06
bicor=-0.2, p=0.051
cor=-0.19, p=0.065
COTL1|Q14019
Kruskal-W. p = 0.22
Tukey AD/CT p = 0.94
AD/AsymAD 0.94
AsymAD/CT 0.78
bicor=-0.16, p=0.16
cor=-0.15, p=0.18
bicor=0.17, p=0.13
cor=0.15, p=0.18
bicor=0.068, p=0.55
cor=0.11, p=0.33
bicor=-0.095, p=0.4
cor=-0.12, p=0.29
bicor=-0.028, p=0.81
cor=-0.019, p=0.87
S100B|P04271
Kruskal-W. p = 0.61
Tukey AD/CT p = 0.2
AD/AsymAD 1
AsymAD/CT 0.21
bicor=-0.24, p=0.029
cor=-0.23, p=0.04
bicor=-0.018, p=0.88
cor=0.094, p=0.41
bicor=0.06, p=0.6
cor=0.19, p=0.091
bicor=-0.14, p=0.21
cor=-0.15, p=0.18
bicor=-0.1, p=0.38
cor=-0.15, p=0.19
Protein Abun
CT
AsymAD
AD
Amyloid Quant.
Tau Quant.
phosphoTau Quant
ABeta/Tau Ratio
MoCA Score

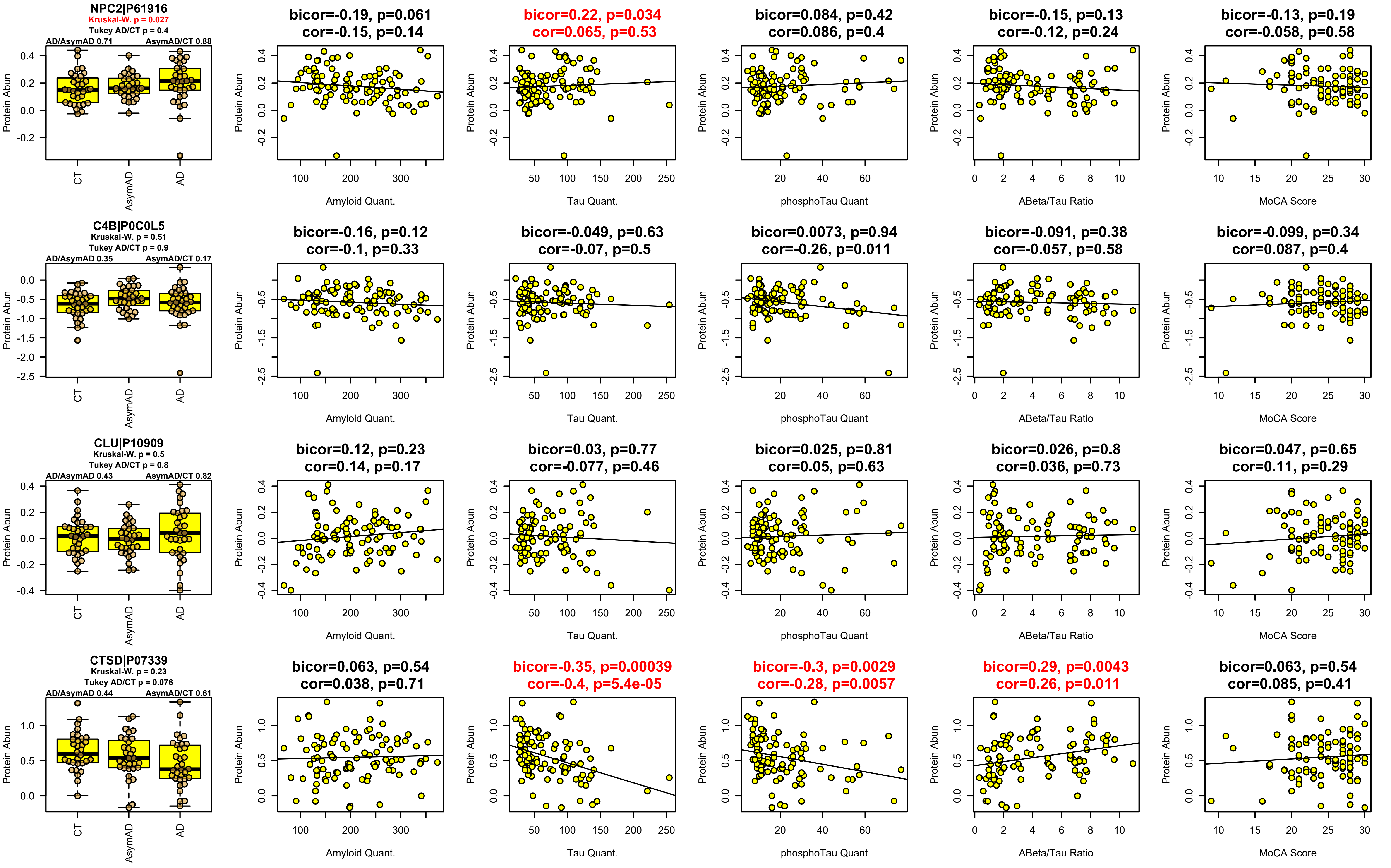
NPC2|P61916
Kruskal-W. p = 0.027
Tukey AD/CT p = 0.4
AD/AsymAD 0.71
AsymAD/CT 0.88
bicor=-0.19, p=0.061
cor=-0.15, p=0.14
bicor=0.22, p=0.034
cor=0.065, p=0.53
bicor=0.084, p=0.42
cor=0.086, p=0.4
bicor=-0.15, p=0.13
cor=-0.12, p=0.24
bicor=-0.13, p=0.19
cor=-0.058, p=0.58
C4B|P0C0L5
Kruskal-W. p = 0.51
Tukey AD/CT p = 0.9
AD/AsymAD 0.35
AsymAD/CT 0.17
bicor=-0.16, p=0.12
cor=-0.1, p=0.33
bicor=-0.049, p=0.63
cor=-0.07, p=0.5
bicor=0.0073, p=0.94
cor=-0.26, p=0.011
bicor=-0.091, p=0.38
cor=-0.057, p=0.58
bicor=-0.099, p=0.34
cor=0.087, p=0.4
CLU|P10909
Kruskal-W. p = 0.5
Tukey AD/CT p = 0.8
AD/AsymAD 0.43
AsymAD/CT 0.82
bicor=0.12, p=0.23
cor=0.14, p=0.17
bicor=0.03, p=0.77
cor=-0.077, p=0.46
bicor=0.025, p=0.81
cor=0.05, p=0.63
bicor=0.026, p=0.8
cor=0.036, p=0.73
bicor=0.047, p=0.65
cor=0.11, p=0.29
CTSD|P07339
Kruskal-W. p = 0.23
Tukey AD/CT p = 0.076
AD/AsymAD 0.44
AsymAD/CT 0.61
bicor=0.063, p=0.54
cor=0.038, p=0.71
bicor=-0.35, p=0.00039
cor=-0.4, p=5.4e-05
bicor=-0.3, p=0.0029
cor=-0.28, p=0.0057
bicor=0.29, p=0.0043
cor=0.26, p=0.011
bicor=0.063, p=0.54
cor=0.085, p=0.41
Protein Abun
CT
AsymAD
AD
Amyloid Quant.
Tau Quant.
phosphoTau Quant
ABeta/Tau Ratio
MoCA Score
